# Identification of four linear B-cell epitopes on the SARS-CoV-2 spike protein able to elicit neutralizing antibodies

**DOI:** 10.1101/2020.12.13.422550

**Authors:** Lin Li, Zhongpeng Zhao, Xiaolan Yang, Wendong Li, Shaolong Chen, Ting Sun, Lu Wang, Yufei He, Guang Liu, Xiaohan Han, Hao Wen, Yong Liu, Yifan Chen, Haoyu Wang, Jing Li, Zhongyi Su, Chen Du, Yiting Wang, Xinyang Li, Zeqian Yang, Jie Wang, Min Li, Tiecheng Wang, Ying Wang, Yubo Fan, Hui Wang, Jing Zhang

**Author notes:** Lin Li, Zhongpeng Zhao and Xiaolan Yang contributed equally to this work. Correspondence to: Prof. Jing Zhang, Beijing Advanced Innovation Centre for Biomedical Engineering, Key Laboratory for Biomechanics and Mechanobiology of Ministry of Education, School of Biological Science and Medical Engineering, Beihang University, Beijing 100083, P. R. China, No. 37 Xueyuan Road, Haidian District, Beijing, 100082, China., Prof. Hui Wang, State Key Laboratory of Pathogen and Biosecurity, Beijing Institute of Microbiology and Epidemiology, Academy of Military Medical Sciences, Beijing. No. 20Dongdajie Street, Fengtai District, Beijing 100071, China., Prof. Yubo Fan, Beijing Advanced Innovation Centre for Biomedical Engineering, Key Laboratory for Biomechanics and Mechanobiology of Ministry of Education, School of Biological Science and Medical Engineering, Beihang University, Beijing 100083, P. R. China, No. 37 Xueyuan Road, Haidian District, Beijing, 100082, China., Assoc. Prof. Ying Wang, Beijing Advanced Innovation Centre for Biomedical Engineering, Key Laboratory for Biomechanics and Mechanobiology of Ministry of Education, School of Biological Science and Medical Engineering, Beihang University, Beijing 100083, P. R. China, No. 37 Xueyuan Road, Haidian District, Beijing, 100082, China.

## Abstract

SARS-CoV-2 unprecedentedly threatens the public health at worldwide level. There is an urgent need to develop an effective vaccine within a highly accelerated time. Here, we present the most comprehensive S-protein-based linear B-cell epitope candidate list by combining epitopes predicted by eight widely-used immune-informatics methods with the epitopes curated from literature published between Feb 6, 2020 and July 10, 2020. We find four top prioritized linear B-cell epitopes in the hotspot regions of S protein can specifically bind with serum antibodies from horse, mouse, and monkey inoculated with different SARS-CoV-2 vaccine candidates or a patient recovering from COVID-19. The four linear B-cell epitopes can induce neutralizing antibodies against both pseudo and live SARS-CoV-2 virus in immunized wild-type BALB/c mice. This study suggests that the four linear B-cell epitopes are potentially important candidates for serological assay or vaccine development.

## Introduction

The new coronavirus SARS-CoV-2 with documented person-to-person transmission has caused millions of confirmed cases and poses unprecedented threat to human lives^1–3^. The clinical features associated with SARS-CoV-2 infection include non-symptomatic infection, mild flu-like symptoms to pneumonia, severe acute respiratory distress syndrome or even deaths^3,4^. It has been declared as a pandemic by World Health Organization (WHO) on 11^th^ March, 2020. To date, no effective treatment is available yet, even though great efforts are being put into the discovery of antiviral drugs. There is an urgent need to understand and develop vaccines against SARS-CoV-2.

At present, more than one hundred of vaccine development projects being carried out globally ranged from viral vector-based vaccines, mRNA and DNA vaccines, subunit vaccines, nano-particle-based vaccines, to inactivated-whole virus vaccines^5–8^. Some vaccine candidates such as inactivated vaccine can inhibit virus replication and protect against upper respiratory tract disease, and other vaccine candidates such as Ad5-nCoV encoding the full spike of SARS-CoV-2 demonstrate good characteristics in both safety and immunogenicity^5–9^. However, some mild and self-limiting adverse reactions are still observed in some clinical trials^10^. It is necessary to carefully evaluate the effects and side-effects of these vaccine candidates, and meanwhile to have ongoing new vaccine development.

SARS-CoV-2 fuses and enters into the host cells through the spike (S) protein binding to the angiotensin-converting enzyme 2 (ACE2) receptor^11^. It would be efficient to prevent virus entry by blocking the binding of S protein to ACE2^12–14^. The S protein consists of S1-subunit and the membrane fusion S2-subunit. The significant role of the S1 subunit receptor-binding domain (RBD) in the entry of virus to the host cell makes it wildly-investigated target for developing therapeutic antibodies and vaccines^15–17^. Other area of S protein may also elicit neutralizing antibodies^18,19^.

Various immune-informatics methods have been used to predict large quantities of linear B-cell epitopes with high immunogenicity^20–43^, but the selection of the in-silicon methods is mostly based on the researchers’ experiences and preferences. Due to lack of validations from biological experiments, it is difficult to focus on a limited number of candidates for diagnostics and linear B-cell epitope-based vaccine development.

In this study, we presented a comprehensive S-protein-based linear B-cell epitope candidate list by combining epitopes predicted by eight widely-used immune-informatics methods with the epitopes curated from literature published between Feb 6, 2020 and July 10, 2020. The antigenicity, toxicity, stability and physiochemical properties were explored for all the identified linear B-cell epitopes. Based on integrative information from epitopes properties and 3D structure of S protein, the top prioritized linear B-cell epitopes were investigated for their binding affinity with serum antibodies from horse, mouse, and monkey inoculated with different SARS-CoV-2 vaccine candidates (S1-based vaccine for horse, RBD-based vaccines for mouse and monkey) and a patient recovering from COVID-19. The induction of neutralizing antibodies against both pseudo and live SARS-CoV-2 virus has been examined following immunizations in wild-type BALB/c mice. Four linear B-cell epitopes found able to elicit neutralizing antibodies are potentially important candidates for serological assay or vaccine development.

## Results

### Identification of linear B-cell epitopes on the SARS-CoV-2 Spike protein

Spike protein is an important target for vaccine development due to its indispensable function in helping SARS-CoV-2 gain entry into host cells. B-cells can be guided through linear B-cell epitopes to recognize and activate defense responses against viral infection. To construct a comprehensive linear B-cell epitope candidate list, we first performed in-silicon prediction of B-cell epitopes from S protein through eight methods, obtaining a total of 4044 linear B-cell epitopes (256 for Bepipred and Bepipred2.0 with default parameter settings, Kolaskar and Tongaonkar antigenicity, Parker hydrophilicity, Chou and Fasman beta-turn, and Karplus and Schulz flexibility provided by IEDB (Immune-Epitope-Database And Analysis-Resource)^44^; 128 for BcePred ^45^ using accessibility, antigenic propensity, exposed surface, flexibility, hydrophilicity, polarity, and turns; 3007 for the ANNpred-based server ABCpred^46^; 44 for Ellipro^47^; 176 for BCPREDS^48^; 191 for AAP^49^; 215 for FBCPRED^50^; and 27 for COVIDep^51^). We additionally extracted 279 linear B-cell epitopes from 24 articles or preprints^19–43^ published between Feb 6, 2020 and July 10, 2020. We finally established a full list of 3836 unique linear B-cell epitope candidates by combining predictions by the eight methods and those curated from literature (Supplementary Table 1).

To obtain linear B-cell epitopes with high potential to initiate a defensive immune reaction, we adopted a series of high stringent criteria to filter out epitopes with low antigenicity. 614 linear B-cell epitopes were estimated by VaxiJen 2.0^52^ to have high antigenicity scores (larger than 0.9 viewed adequate to initiate a defensive immune reaction) (Fig. 1A) with length varying from 6 amino acids to 30 amino acids (Fig. 1B) (Supplementary Table 1). The positions of the 614 linear B-cell epitopes on the S protein amino acid sequence were examined to exclude the epitopes locating on the non-outer surface based on the transmembrane topology of SARS-CoV-2 S protein predicted by TMHMM v2.0 (outside:1-1213; transmembrane:1214-1236; inside: 1237-1273). A total of 14 regions locating on the outer surface of SARS-CoV-2 (4-32, 167-189, 369-402, 404-431, 482-523, 526-563, 568-605, 628-678, 697-741, 890-916, 1027-1051, 1057-1081, 1155-1181, 1196-1213) were identified as hotspot regions containing highly antigenic linear B-cell epitopes identified by multiple methods (Fig. 1C). Based on the conservation status of each residue of the S protein predicted by ConSurf using seven known coronaviruses incorporating SARS-CoV-2 (YP_009724390.1), SARS-CoV (NP_828851.1), MERS-CoV (YP_009047204.1), alpha coronavirus 229E (NP_073551.1), alpha coronavirus NL63 (AFV53148.1), beta coronavirus OC43 (YP_009555241.1) and beta coronavirus HKU1 (AAT98580.1), we discovered that RBD region (319-514) of the S protein was not conserved among the seven coronaviruses, consistent with the observation that more mutations were found in the RBD region by performing sequence alignment of S protein against 118,694 (20200927) sequences of SARS-CoV-2 in the NGDC database (Fig. 1C). The linear B-cell epitopes from the 14 hotspot regions were mapped to the 3D structure of the SARS-CoV-2 S protein (PDB ID:6VSB), suggesting that hotspot regions locating on the exposed area of spike stem or spike head would harbor good B-cell epitope candidates (Fig. 1D and E).

**Fig. 1.**
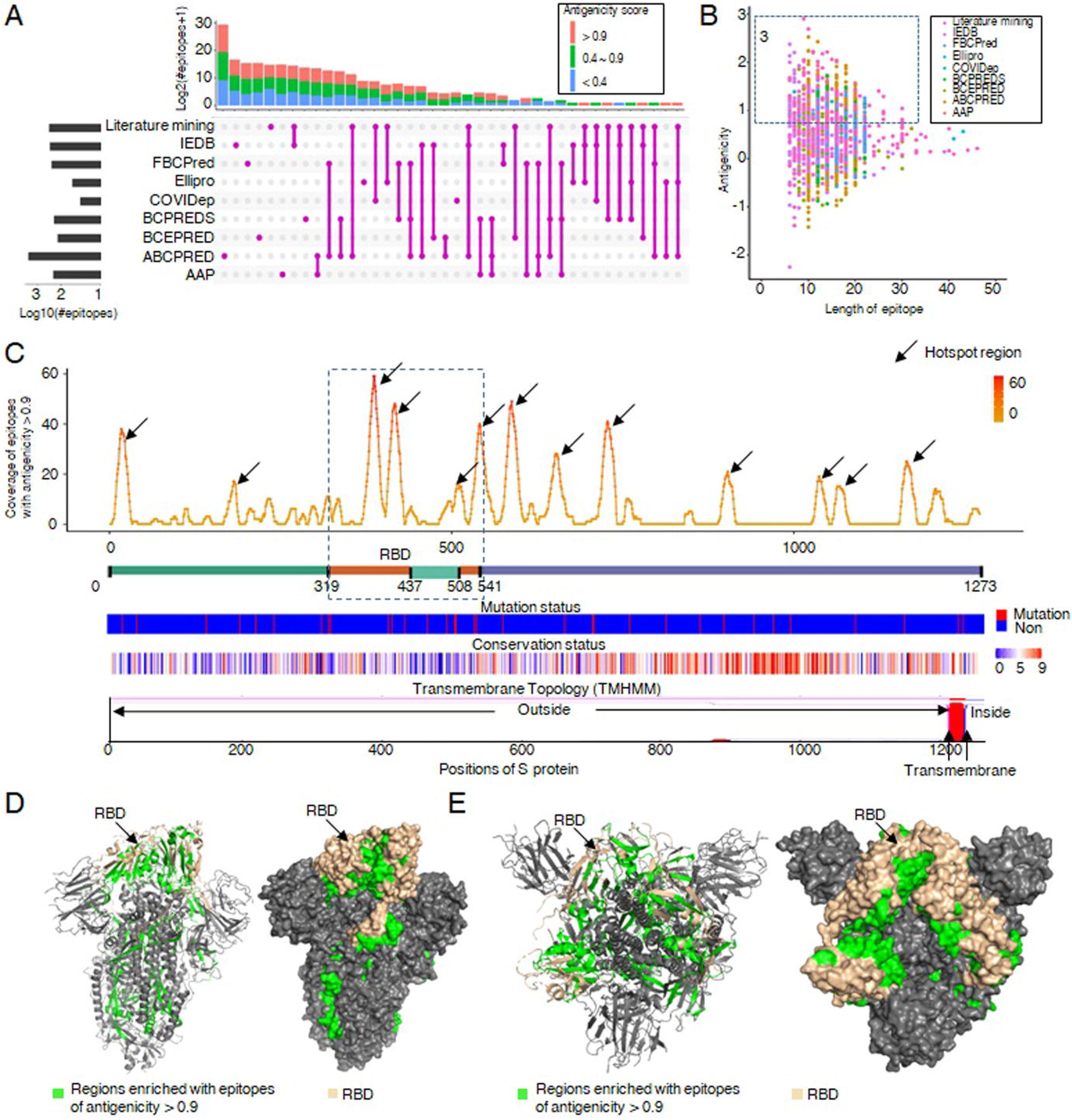
The predicted linear B-cell epitopes in the Spike protein of SARS-CoV-2. **a**, The number of linear B-cell epitopes shared among the distinct methods and literature mining. The pink, green and light blue represent epitopes with antigenicity scores >0.9, 0.4 and 0.9, and <0.4, respectively. **b**, The antigenicity score and peptide length distribution of the predicted linear B-cell epitopes. **c**, The positions of the linear B-cell epitopes with high antigenicity scores (>0.9) on the S protein amino acid sequence. The y-axis is the coverage of predicted linear B-cell epitopes. The bars below are the protein domain, mutation sites, conservative scores across seven known coronaviruses viruses, and the transmembrane domain predicted by TMHMM, respectively. **d-e**, The localizations of regions enriched with linear B-cell epitopes (antigenicity score>0.9) on SARS-CoV-2 S (PDB: 6VSB) protein. The grey is protein, the green is the regions enriched with linear B-cell epitopes (antigenicity score>0.9), and the wheat color is the RBD regions.

### Characterization of the 18 linear B-cell epitopes selected with predicted high immunigenicity

From the hotspot regions of the exposed area of S protein, we selected 18 linear B-cell epitopes with few mutations reported in less than ten out of a total of 118,694 SARS-CoV-2 virus strains as the potentially optimal linear B-cell epitope candidates (Fig. 2A). As IgG and IgA are the most abundant isotypes in blood^53^, we examined the recently available target profiles (20-mer peptides tiling every 5 amino acids across the SARS-CoV-2 proteome) of IgG and IgA antibodies from 232 COVID-19 patient sera (101 for hospitalized, 131 for non-hospitalized) and 190 pre-COVID-19 era controls ^53^. We discovered that the 20-mer peptides in the target profiles with more than 75% overlapping with the 18 linear B-cell epitopes exhibited non-negligible enrichment score in some hospitalized or non-hospitalized COVID-19 patient IgG, or IgA samples, suggesting that the regions harboring the 18 linear B-cell epitopes have immunogenic potentials. Particularly, compared with pre-COVID-19 era controls, ‘GDEVRQIAPGQTGKIADYNYKLP’, ‘DEVRQIAPGQTGKIADYNYKLPDDFT’, ‘VRQIAPGQTGKIAD’, ‘APGQTGK’, ‘APGQTGKIADYNYKL’, ‘CVNFNFNGL’, ‘LGQSKR’, ‘GQSKRVDF’, and ‘GQSKRVDFC’ showed a statistically significantly higher enrichment score in hospitalized COVID-19 patient IgG samples, while ‘APGQTGKIADYNYKLPDDFT’, ‘KIADYNYKLPDDFT’, and ‘VVFLHVTYV’ having significantly higher enrichment score in non-hospitalized COVID-19 patient IgG samples (Fig. 2A). Similarly, ‘KLNDLCFTNVYAD’ and ‘CVNFNFNGL’ had significantly higher enrichment score in hospitalized COVID-19 IgA samples than negative controls, while ‘KLNDLCFTNVYAD’, ‘GDEVRQIAPGQTGKIADYNYKLP’, ‘CVNFNFNGL’, and ‘YQPYRVVVLSFELLH’ showed significantly higher enrichment score in non-hospitalized COVID-19 IgA samples than negative controls (Fig. 2A).

**Fig. 2.**
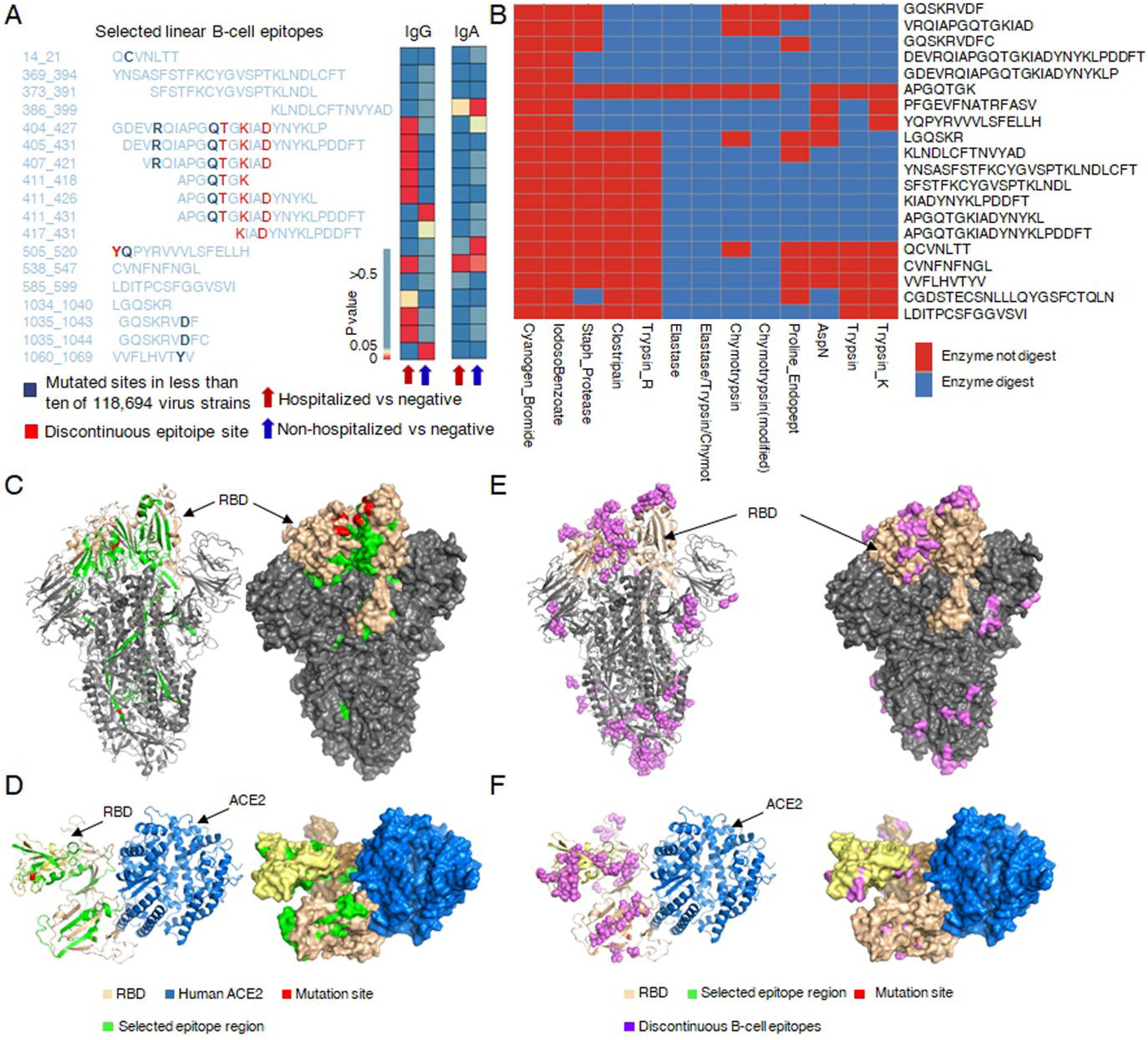
The characteristics of the 18 selected linear B cell epitopes. **a**, The sequences of 18 selected linear B-cell epitopes. The bold is the mutated site in less than ten of 118,694 virus strains; The red is the predicted discontinuous residues. The bars on the right side are the Wilcoxon test *p* value for the comparisons of IgG or IgA antibody enrichment scores associated with each linear B-cell epitope between COVID-19 patients and negative controls. **b**, The digesting enzymes profile of the epitope sequence. Red indicated not digest, blue indicated digest. **c-d**, The localization of the 18 selected epitopes mapped on SARS-CoV-2 S (PDB: 6VSB) protein (c) and ACE-RBD complex (d). **e-f**, The localizations of B cell discontinuous epitopes on SARS-CoV-2 S (PDB: 6VSB) protein (e) and ACE-RBD complex (f). The spike protein is grey, the RBD region is wheat color, the selected epitopes are green, the mutation sites are red, the human ACE domain is blue, and the discontinuous B-cell epitopes are purple.

To evaluate the stability of the 18 linear B-cell epitopes, we estimated the number of peptide-digesting enzymes through the protein digest server with13 enzymes available (See Method). A linear B-cell epitope would have potentially higher stability if more enzymes were predicted to be unable to digest it. All the 18 linear B-cell epitopes were found having at least two non-digesting enzymes varying from 2 to 12 enzymes (Fig. 2B). The 18 linear B-cell epitopes were mapped to the 3D structure of the SARS-CoV-2 S protein (PDB ID: 6VSB), showing that ‘YNSASFSTFKCYGVSPTKLNDLCFT’, ‘SFSTFKCYGVSPTKLNDL’, ‘KLNDLCFTNVYAD ‘, ‘GDEVRQIAPGQTGKIADYNYKLP’, ‘DEVRQIAPGQTGKIADYNYKLPDDFT’, ‘VRQIAPGQTGKIAD’, ‘APGQTGK’, ‘APGQTGKIADYNYKL’, ‘APGQTGKIADYNYKLPDDFT’, ‘KIADYNYKLPDDFT’, ‘YQPYRVVVLSFELLH’, and ‘CVNFNFNGL’ located in the most exposed RBD region of the spike head (Fig. 2C), and ‘QCVNLTT’, ‘LDITPCSFGGVSVI’, ‘LGQSKR’, ‘GQSKRVDF’, ‘GQSKRVDFC’, and ‘VVFLHVTYV’ were in the spike stem region (Fig. 2C). The 12 linear B-cell epitopes in the spike head were found to substantially overlap with the predicted interacting surface where ACE-2 binds to the RBD of SARS-CoV-2 S protein (Fig. 2D)^54–57^, suggesting that an antibody binding to this surface may block viral entry into cells. Discontinuous B-cell epitopes were also predicted by Discotope 2.0 using A, B, and C chain of the 3D structure of S protein (PDB ID:6VSB). The positions of discontinuous B-cell epitopes mapped on the 3D structure of S protein, revealing that RBD region is enriched with discontinuous B-cell epitopes (Fig. 3E). Similarly, the discontinuous B-cell epitopes in the RBD regions overlapped with the interacting surface between ACE2 and the RBD of S protein (Fig. 3F). The linear B-cell epitopes locating in the RBD regions (‘GDEVRQIAPGQTGKIADYNYKLP’, ‘DEVRQIAPGQTGKIADYNYKLPDDFT’, ‘VRQIAPGQTGKIAD’, ‘APGQTGK’, ‘APGQTGKIADYNYKL’, ‘APGQTGKIADYNYKLPDDFT’, ‘KIADYNYKLPDDFT’, and ‘YQPYRVVVLSFELLH’) incorporated multiple discontinuous B-cell epitopes (Fig. 2A, E). By comparing the mutations documented in 53,969 SARS-CoV-2 virus strains (NGDC), we discovered that eight linear B-cell epitopes in the RBD region incorporated one or two mutation sites in few virus strains (C mutated in 11 virus strain for ‘QCVNLTT’; R mutated in one virus strain and Q mutated in 10 virus strain for ‘GDEVRQIAPGQTGKIADYNYKLP’, ‘DEVRQIAPGQTGKIADYNYKLPDDFT’, and ‘VRQIAPGQTGKIAD’; Q mutated in 10 virus strain for ‘APGQTGK’, ‘APGQTGKIADYNYKL’, ‘APGQTGKIADYNYKLPDDFT’; Y mutated in 4 virus strain and Q mutated in 2 virus strain for ‘YQPYRVVVLSFELLH’; D mutated in one virus strain for ‘GQSKRVDF’ and ‘GQSKRVDFC’; Y mutated in 11 virus strain for ‘VVFLHVTYV’) (Fig. 2A). None of the 18 linear B-cell epitopes were found toxic.

**Fig. 3.**
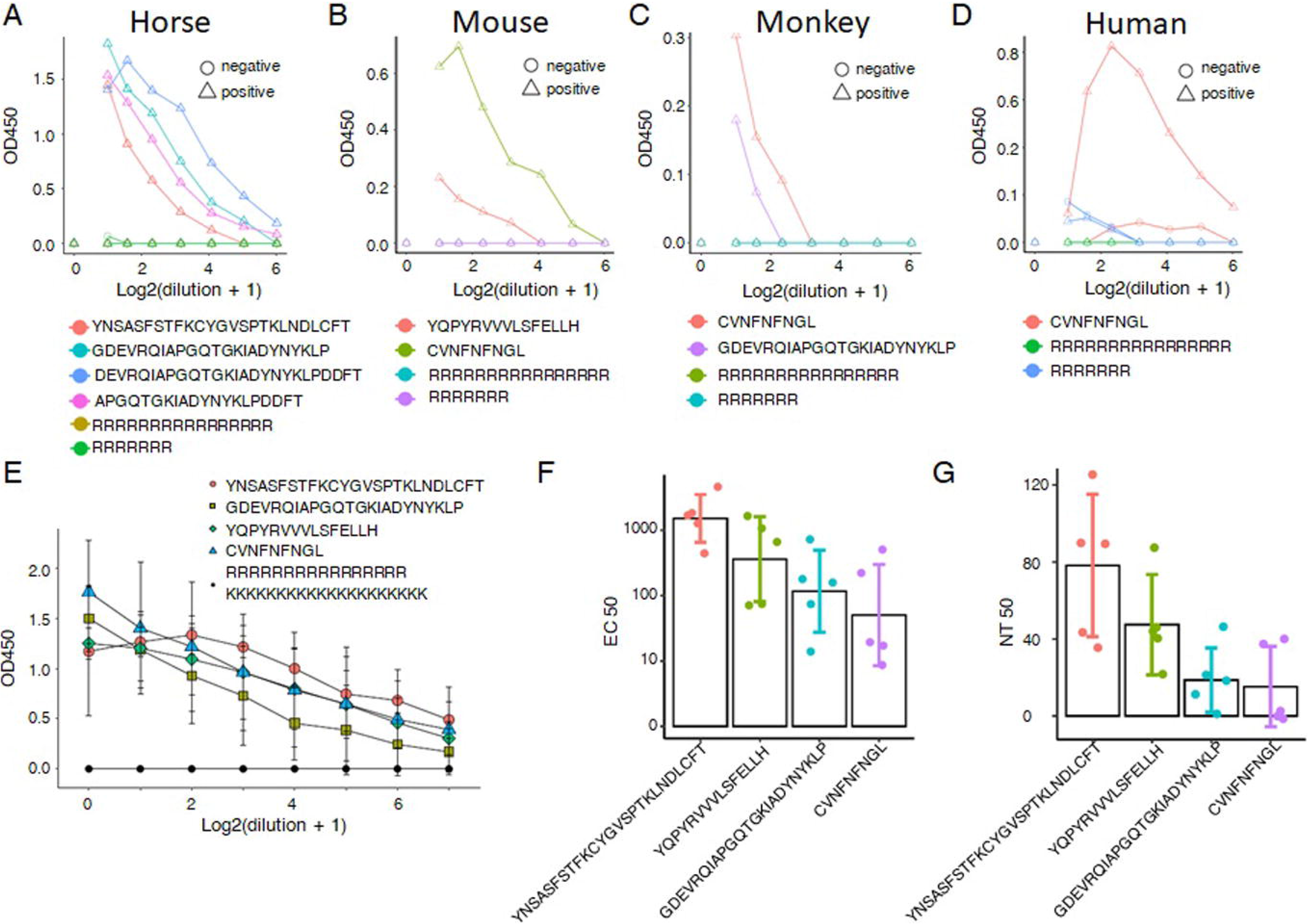
Measurements of the selected Linear B cell epitope binding to antibody and neutralization efficiency of selected epitopes against SARS-CoV-2. a-d, The binding affinity assessed by ELISA between linear B-cell epitopes and serum antibodies from immunized horse with S1-based vaccines (a), immunized mouse with RBD-based vaccines (b), immunized monkey with RBD-based vaccines (c), and a patient recovering from COVID-19 (d). e, The binding affinity assessed by ELISA between the linear B-cell epitopes and serum antibodies from immunized mice with corresponding epitopes of ‘YNSASFSTFKCYGVSPTKLNDLCFT’, ‘GDEVRQIAPGQTGKIADYNYKLP’, ‘YQPYRVVVLSFELLH’, and ‘CVNFNFNGL’. f, Neutralization assay against SARS-CoV-2 pseudovirus in ‘YNSASFSTFKCYGVSPTKLNDLCFT’, ‘GDEVRQIAPGQTGKIADYNYKLP’, ‘YQPYRVVVLSFELLH’, and ‘CVNFNFNGL’. y-axis is the value of EC_50_. g, Neutralization assay against SARS-CoV-2 live virus in ‘YNSASFSTFKCYGVSPTKLNDLCFT’, ‘GDEVRQIAPGQTGKIADYNYKLP’, ‘YQPYRVVVLSFELLH’, and ‘CVNFNFNGL’. y-axis is the value of NT_50_.

### Four linear B-cell epitopes specifically binding with serum antibodies from the animals inoculated with different SARS-CoV-2 vaccine candidates or a patient recovering from COVID-19

Animal models are necessary to demonstrate efficacy and safety in the development of vaccines against SARS-CoV-2 infection^58,59^. BALB/c mice is a good animal model for investigating SARS-CoV-2 infection in both upper and lower respiratory tracts^5^. Monkey, phylogenetically close to humans, has been used to test whether seroconversion provides protective immunity against SARS-CoV-2^58^. To assess the binding of the 18 linear B-cell epitopes with serum IgG antibodies against SARS-CoV-2, we immunized three model animals including horse, mouse, and monkey. Indirect enzyme-linked immunosorbent assay (ELISA) was performed between serum antibodies (including animals inoculated with different SARS-CoV-2 vaccine candidates and normal control animal models), and the 18 linear B-cell epitopes and arbitrary control peptides (‘RRRRRRRRRRRRRRRR’ and ‘RRRRRRR’). We discovered that four linear B-cell epitopes reacted specifically and dose dependently with serum antibodies from the vaccinated horse (‘YNSASFSTFKCYGVSPTKLNDLCFT’, and three highly overlapped linear B-cell epitopes including ‘GDEVRQIAPGQTGKIADYNYKLP’, ‘DEVRQIAPGQTGKIADYNYKLPDDFT’ and ‘APGQTGKIADYNYKLPDDFT’ for horse (Fig. 3A); ‘YQPYRVVVLSFELLH’ and ‘CVNFNFNGL’ for mouse (Fig. 3B); ‘CVNFNFNGL’ and ‘GDEVRQIAPGQTGKIADYNYKLP’ for monkey (Fig. 3C)), whereas the arbitrary control peptides had no effects. The normal sera from horse, mouse, monkey and human were not reactive. In addition, sera from a patient recovering from COVID-19 and the healthy persons were provided by Dr. Bin Su at Beijing Youan Hospital. Indirect ELISA results demonstrated that ‘CVNFNFNGL’ specifically bond to serum IgG antibodies in the patient recovering from COVID-19, but not in healthy human (Fig. 3D). These results suggest that the four linear B-cell epitopes (‘YNSASFSTFKCYGVSPTKLNDLCFT’, ‘GDEVRQIAPGQTGKIADYNYKLP’, ‘YQPYRVVVLSFELLH’, and ‘CVNFNFNGL’) may be able to induce antibodies against SARS-CoV-2.

### Antibodies against the four linear B-cell epitopes neutralize SARS-CoV-2

To confirm that the four linear B-cell epitopes generated antibodies against SARS-CoV-2, we immunized six-to eight-week-old female BALB/c mice through five consequent subcutaneous injections of five microgram dose of linear B-cell epitope-based synthetic peptides with 7 days interval between two consecutive injections (five mice per linear B-cell epitope). Serum samples collected from each synthetic peptide immunized mice were assessed for binding to corresponding linear B-cell epitope by ELISA, as well as for neutralization against SARS-CoV-2 pseudovirus and live virus. It was evident that a substantial percentage of antibodies generated in vaccinated mice against the linear B-cell epitope antigens as demonstrated in the ELISA results (Fig. 3E), suggesting that antibodies directed at the four linear B-cell epitopes could bind SARS-CoV-2.

In SARS-CoV and MERS-CoV^60^, pseudovirus neutralization assay is a sensitive and quantitative method. We therefore tested the concentration of neutralizing antibodies in immune sera from wild type mice 7 days after the fifth vaccination against SARS-CoV-2 pseudovirus using a pseudotyped virus-based neutralization assay developed recently for SARS-CoV-2^61^. The neutralizing antibodies were detected in all mice immunized with each of the four linear B-cell epitopes (mean EC50 is 1963 for ‘YNSASFSTFKCYGVSPTKLNDLCFT’; mean EC50 is 704 for ‘YQPYRVVVLSFELLH’; mean EC50 is 230 for ‘GDEVRQIAPGQTGKIADYNYKLP’; mean EC50 is 154 for ‘CVNFNFNGL’) (Fig. 3F), whereas no neutralizing antibodies were found in the two control group.

We also performed neutralization assay against SARS-CoV-2 live virus (BetaCoV/Beijing/IMEBJ01/2020) using immune sera from vaccinated mice 7 days after the fifth vaccination. We found the strongest neutralization activity with mean NT50 of 78.2 for ‘YNSASFSTFKCYGVSPTKLNDLCFT’; ‘YQPYRVVVLSFELLH’ also showed strong neutralization activity (mean NT50 = 47.4); we observed the neutralization activities for ‘GDEVRQIAPGQTGKIADYNYKLP’ (mean NT50 = 18.6) and ‘CVNFNFNGL’ (mean NT50 = 15.2) (Fig. 3G). Overall, these results demonstrated that the four linear B-cell epitopes can induce neutralizing antibodies against SARS-CoV-2.

## Conclusion

SARS-CoV-2 has caused a serious pandemic all over the world. Safe and effective vaccines are urgently needed to be developed and deployed in a rapid but highly reliable manner. More than one hundred projects are being executed in the WHO draft landscape of COVID-19 candidate vaccines including a variety of vaccine types such as viral vector-based vaccines, mRNA and DNA vaccines, subunit vaccines, nanoparticle-based vaccines, to inactivated-whole virus vaccines^5–9^. However, even for vaccines in clinical trial phase III, some mild side effects are still observed and sporadically reported. Ongoing efforts are necessary for developing new vaccines. Rigorous selection of epitopes most likely to elicit neutralizing antibodies against virus can accelerate this process.

S protein is an important target in vaccine development for its critical functions for SARS-CoV-2 to fuse and enter into the host cells^15–17^. A large number of predicted epitopes are generated within a highly accelerated time frame^20–43^, but the selection of in-silicon prediction methods is based on the experiences and preferences of each individual researcher. There is no comprehensive benchmark about the accuracy and specificity of these epitopes prediction methods yet. Lack of limited number of target epitopes with supports from biological validation experiments makes the epitope-based vaccine development in the middle of nowhere.

In this study, we predicted linear B-cell epitopes from S protein by eight widely-used immune-informatics methods including Bepipred and Bepipred2.0 with default parameter settings, Kolaskar and Tongaonkar antigenicity, Parker hydrophilicity, Chou and Fasman beta-turn, and Karplus and Schulz flexibility provided by IEDB (Immune-Epitope-Database And Analysis-Resource)^44^, BcePred^45^ using accessibility, antigenic propensity, exposed surface, flexibility, hydrophilicity, polarity, and turns, ANNpred-based server ABCpred^46^, Ellipro^47^, BCPREDS^48^, AAP^49^, FBCPRED^50^ and COVIDep^51^. To generate a comprehensive linear B-cell epitope candidate list, we incorporated 279 linear B-cell epitopes curated from 24 articles or preprints^19–43^ published between Feb 6, 2020 and July 10, 2020. Interestingly, the linear B-cell epitopes predicted by different methods converged to some hotspot regions in the S protein, suggesting the pivotal role of these regions in diagnostics assay and vaccine development. Integrating antigenicity, toxicity, stability and physiochemical properties, 3D structure of S protein, the 3D conformation structure between RBD of S protein and ACE2, we selected 18 top prioritized linear B-cell epitopes for further investigation. Four out of 18 linear B-cell epitope-based synthetic peptides were found to specifically bind with serum antibodies from horse, mouse, and monkey inoculated with different SARS-CoV-2 vaccine candidates or a patient recovering from COVID-19. The serum antibodies we used here were generated by different regions of SARS-CoV-2 spike protein (S1-based vaccine for horse, RBD-based vaccines for mouse and monkey, unknown regions for a patient recovering from COVID-19), leading to the observation that the four peptides didn’t consistently show specific binding with all antibodies produced across horse, mouse, monkey and a patient recovering from COVID-19. However, the four peptides were able to elicit neutralizing antibodies in immunized wild-type BALB/c mice against both pseudo and live SARS-CoV-2 virus.

To the best of our knowledge, the linear B-cell epitope candidate list we presented is one of the most comprehensive and valuable source for developing vaccines. Importantly, the four linear B-cell epitopes we identified able to elicit neutralizing antibodies against SARS-CoV-2 are promising and valuable candidates which have immediate usefulness for developing vaccines against SARS-CoV-2. Besides, more linear B-cell epitopes in the candidate list may also deserve being examined. The analysis workflow we adopted can be broadly applied to identify B-cell epitopes for a large repertoire of virus, not limited to coronavirus. In short, the results we presented here will be useful to guide the identification and prioritization of linear B-cell epitope-based diagnostics and vaccine designs during this unprecedented pandemic.

## Methods

### Data retrieval and structural analysis

SARS-CoV-2 protein sequence (Accession number MN908947.3)^62^ was extracted from the NCBI database. Experimentally solved 3D structure of SARS-CoV-2 S protein (PDB ID: 6VSB)^11^ was retrieved from Protein Data Bank. The predicted 3D structure of protein interactions between SARS-CoV-2 S protein RBD domain and human ACE-2 was obtained from recent reports^54–57,63,64^. The transmembrane topology of S protein was examined by TMHMM.v2.0 (http://www.cbs.dtu.dk/services/TMHMM/). VaxiJen v2.0^52^ was used to estimate the antigenicity of full-length SARS-CoV-2 S protein.

### Linear B-cell epitope prediction for SARS-CoV-2 S protein

A total of eight algorithms were used to predict linear B-cell epitopes for SARS-CoV-2 S protein. Bepipred and Bepipred2.0 with default parameter settings, Kolaskar and Tongaonkar antigenicity, Parker hydrophilicity, Chou and Fasman beta-turn, and Karplus and Schulz flexibility provided by IEDB (Immune-Epitope-Database And Analysis-Resource)^44^ were applied upon SARS-CoV-2 S protein sequence to predict linear B-cell epitopes. Linear B-cell epitopes were also predicted by BcePred^45^ using accessibility, antigenic propensity, exposed surface, flexibility, hydrophilicity, polarity, and turns. The ANNpred-based server ABCpred^46^, Ellipro^47^, BCPREDS^48^, AAP^49^, FBCPRED^50^, and COVIDep^51^ were also employed to predict linear B-cell epitopes. 25 published articles or preprints^19–43^ about epitope predictions from Feb 6, 2020 to July 10, 2020 were curated to obtain a total of 214 B-cell epitopes. All the linear B-cell epitopes including those predicted by the eight methods and those curated from literature were combined to the linear B-cell epitope candidate list. According to the transmembrane topology of SARS-CoV-2 S protein predicted by TMHMM v2.0, the linear B-cell epitopes on the outer surface were retained for downstream analysis with intracellular epitopes eliminated. The linear B-cell epitopes consisting of less than six amino acids or more than 50 amino acids were further removed. The antigenicity of the remained linear B-cell epitopes was evaluated by VaxiJen 2.0^52^. A stringent criterion was employed to have linear B-cell epitopes with an antigenicity score larger than 0.9 viewed adequate to initiate a defensive immune reaction.

### Characterization of the 18 selected linear B-cell epitopes

The target profiles of IgG or IgA antibodies from 232 COVID-19 patient sera (101 for hospitalized, 131 for non-hospitalized) and 190 pre-COVID-19 era controls were generated in duplicates by a recent research^53^, using coronavirus 20-mer libraries which included 20-mer peptides tiling every 5 amino acids across the SARS-CoV-2 proteome. The target profiles for IgG/IgA IPs using coronavirus 20-mer libraries were in the form of Z-score, directly obtained from the literature^53^. The Z-scores of the 20-mer peptides in the target profiles were first averaged for the two duplicates of each sample. For each of the 18 linear B-cell epitopes, the Z-scores of 20-mer peptides with more than 75% overlapping with the linear B-cell epitope were compared between hospitalized, non-hospitalized COVID-19 patient groups and negative controls using Mann Whitney U test. The significance was determined if the p value was less than or equal to 0.05.

The locations of the selected linear B-cell epitopes on the 3D structure of SARS-CoV-2 S protein or the interacting conformation of S protein RBD and human ACE2^54–56^ were examined through open-source Pymol. The chemicals and physical properties (Grand average of hydropathicity, half-life, molecular weight, stability index and amino acid atomic composition) of the selected linear B-cell epitopes were analyzed through Protparam^65^. ToxinPred (https://webs.iiitd.edu.in/raghava/toxinpred/index.html) was employed to evaluate the toxicity of linear B-cell epitopes along with hydrophobicity, hydropathicity, hydrophilicity, and charge. As the stable peptidesdigested by fewer enzymes are more favorable candidate vaccines^66^, the digestion of all the selected B-cell epitopes by 13 enzymes (Trypsin, Chymotrpsin, Clostripain, Cyanogen Bromide, IodosoBenzoate, Proline Endopept, Staph Protease, Trypsin K, Trypsin R, AspN, Chymotrypsin (modified), Elastase, and Elastase/Trypsin/Chymotryp) were examined by protein digest server (http://db.systemsbiology.net:8080/proteomicsToolkit/proteinDigest.html).

### Conservation analysis of selected B-cell epitopes

The conservation status for each residue of SARS-CoV-2 were investigated by ConSurf^67^ using the amino acid sequences of S protein from seven known coronaviruses including SARS-CoV-2 (YP_009724390.1), SARS-CoV (NP_828851.1), MERS-CoV (YP_009047204.1), alpha coronavirus 229E (NP_073551.1), alpha coronavirus NL63 (AFV53148.1), beta coronavirus OC43 (YP_009555241.1) and beta coronavirus HKU1 (AAT98580.1). The S protein sequences of different SARS-CoV-2 virus strains were extracted from an open-access database NGDC (https://bigd.big.ac.cn/ncov/), where 118,694 SARS-CoV-2 virus strains’ sequences were documented with 53,969 mutations reported in the virus genome until September27, 2020.

### Peptide Synthesis

The selected 18 linear B-cell epitopes were synthesized by Scilight-Peptide Inc., Beijing, China via a practical approach of Fmoc solid-phase peptide synthesis. The unsophisticated peptides were purified using a Varian ProStar 218 high-performance liquid chromatography (HPLC) instrument with an Agilent Venusil MP C18 reversed phase column. Peptides were eluted with a linear gradient of water, H2O, and acetonitrile, CAN, (both having 0.05% TFA) at a flow rate of 1 mL/min. The separation was monitored at 220 nm using UV detection. Then peptides were subjected to Voyager-DE STR mass spectrometric (MS) analysis. The solvents for gradient elution HPLC are: solvent A, CAN 2%, TFA 0.05% and solvent B, CAN 90%, TFA 0.05%. Peptides were dissolved in deionized H2O at a final concentration of 20mg/ml and stored at −20°C until further use.

### Cell lines and viruses

Vero cells (ATCC,CCL-81) and 293T cells (ATCC,CRL-11268) were maintained in Dulbecco’s modified Eagle’s medium (Gibco, USA) supplemented with 10% fetal bovine serum (Hyclone, USA), penicillin (Hyclone, USA, 100 units/mL) and streptomycin (Hyclone, USA, 100 μg/mL) (complete medium) in 5% CO2 environment at 37 °C and passaged every 2-3 days. BetaCoV/Beijing/IMEBJ01/2020 strain (Genome Warehouse Accession No. GWHACAX01000000) was isolated from a COVID-19 patient and propagated in Vero cells. Virus titres were determined on Vero cells, and virus stocks were stored in aliquots at −80 °C until use(100 CCID50/0.05ml). SARS-CoV-2 pseudoviruses were provided by Dr. Weijin Huang at Institute for Biological Product Control, National Institutes for Food and Drug Control (NIFDC) of China^68^.

The experiments with infectious SARS-CoV-2 were performed at the biosafety level 3 facilities in Beijing Institute of Microbiology and Epidemiology, Academy of Military Sciences, China.

### Animal experiments

All animal experiments were approved by and carried out in accordance with the guidelines of the Institutional Experimental Animal Welfare and Ethics Committee. Specific pathogen-free female BALB/c mice aged 6–8 weeks were obtained from Beijing Vital River Laboratory Animal Technologies Co., Ltd (Beijing, China) and were housed and bred in the temperature-, humidity and light cycle-controlled animal facility (20 ± 2 °C; 50 ± 10%; light, 7:00–19:00; dark,19:00–7:00). The rhesus monkeys (2.0±0.5 kg, 1.5–2.5 years) were provided by Laboratory Animal Center, Academy of Military Medical Science. The animals were housed under standard laboratory conditions. The 4-6 year old, healthy brown horses (300-350kg in weight) that had no detectable antibodies against SARS-CoV-2, were provided by Chifeng Boen Pharmacy Co., LTD (InnerMongolia, China)

The Balb/C mice and monkeys were intramuscularly (i.m.) immunized with the RBD-IgG1 Fc subunit vaccine candidate (10μg per mouse, 40μg per monkey), or Al(OH)3 adjuvant as a control. After the primary injection, all animals received two booster injections with 14 days intervals. The horses were inoculated subcutaneously with SARS-CoV-2 Spike Protein (S1 Subunit, His tag) (Sino Biological, China, cat no:40591-V08H) antigens containing 1.0mg, 1.5mg, 2.0mg and 3.0mg on day 0, 14, 28 and 42, respectively, or Freund′s adjuvant as a control. For each inoculation, 5ml of antigen suspension was mixed with an equal volume of Freund′s adjuvant according to the directions and injected into the several sites near the submandibular and inguinal lumph nodes. Blood samples were collected and used for ELISA and neutralization assays.

### SARS-CoV-2 S1-specific IgG assay in horse, mouse, monkey and human (Indirect ELISA)

96-well polystyrene microplates (Oriental Ocean Global Health, China) were coated with 2 μg/mL (50μL/well) SARS-CoV-2 Spike Protein (S1 Subunit, His tag) (Sino Biological, China, cat no:40591-V08H) in carbonate bicarbonate buffer pH9.6 and the plates were incubated at 4°C overnight. The plates were then blocked at 37 °C for one hour with PBS (Solarbio, China, cat no:A8020) pH 7.4 in 5% skim milk (blocking buffer) and washed with PBST(0.1% Tween-80) three times. Serial dilutions of sera in dilution buffer (Solarbio, China, cat no: P1010) were added to the plates and incubatedat 37°C for 30 minutes. HRP-conjugated goat anti-mouse IgG (Southern Biotech, cat no:1030-05, 1:5000 dilution), HRP-conjuated goat anti-horse IgG (Bellancom life H15034,1:2000 dilution) or HRP-conjugated goat anti-monkey IgG (Solarbio, China, cat no: SE241, 1:3000 dilution for monkey, 1:2000 dilution for human) was added to the plates, and the plates were incubated in incubator (ZHCHENG, China, ZXDP-B2050) at 37°C for 30 minutes and washed with PBST three times. The assay was developed for 10min at 37°C with 50 μL of TMB substrate solution (Solarbio,China, cat no: PR1200), stopped by the addition of 50 μL of stop solution (Solarbio, China, cat no:C1058), and then the optical density (OD) was measured at 450 nm (TECAN, INFINITE F50). The endpoint titre was defined as the highest reciprocal serum dilution that yielded anabsorbance ≥2.1-fold over negative control serum values (OD value is set as 0.05, if it is less than 0.05).The indirect ELISA were performed at the biosafety level 2 facilities in Beijing Institute of Microbiology and Epidemiology, Academy of Military Medical Sciences, China.

### Vaccination of mice

Linear B-cell epitopes (‘YNSASFSTFKCYGVSPTKLNDLCFT’, ‘GDEVRQIAPGQTGKIADYNYKLP’, ‘YQPYRVVVLSFELLH’, and ‘CVNFNFNGL’) were respectively used to immunize BALB/c mice (n = 5 per linear B-cell epitope) through five consequent subcutaneous injections of five microgram dose with 7 days interval between two consecutive injections. Sera were collected for SARS-CoV-2 live virus neutralization assay titration (NT_50_) and SARS-CoV-2 pseudovirus neutralization assay titration (EC50) at different time point.

### SARS-CoV-2 pseudovirus neutralization assay

A pseudotyped virus-based neutralization assay against SARS-CoV-2 in biosafety level 2 facilities was performed as previously described^61^. Briefly, serial dilutions of the samples to be tested were mixed with 325-1,300 TCID_50_/ml of pseudotyped virus. The target cells were incubated for 24 hours, and then the neutralizing antibody content of the sample was obtained by calculating the amount of pseudotyped virus entering the target cells which were detected by the expression of luciferase. The half maximal effective concentration (EC_50_) was calculated for the tested samples using the Reed-Muench method in GraphPad Prism 8 (GraphPad software, Inc., San Diego, CA, USA). The assays were performed at the biosafety level 2 facilities in Beijing Institute of Microbiology and Epidemiology, Academy of Military Medical Sciences, China.

### SARS-CoV-2 neutralization assay

Microneutralization (MN) assay was performed to assess the neutralizing activity of sera from the mice. 50μl (100 CCID_50_/0.05ml) of SARS-CoV-2 IME-BJ01 strain was incubated with serial dilution of heat-inactivated sera in 5% CO_2_ environment at 37°C for one hour. The complexes of antibody-virus (100TCID50/50μl) were added to pre-plated Vero cell monolayers in 96-well plates and incubated for 72 hours. The Reed-Muench method was applied to estimate the dilution of sera required for NT_50_. The initial dilution of sera (1:16) was set as the confidence limit of the assay. Seropositivity was defined as a titre ≥ 16.

## Acknowledgements

We thank Dr. Weijin Huang at Institute for Biological Product Control, National Institutes for Food and Drug Control (NIFDC) of China for providing the SARS-CoV-2 pseudovirus and Dr. Bin Su at Beijing Youan Hospital for providing human Sera. We also thank Dr. Xiliang Wang at State Key Laboratory of Pathogen and Biosecurity, Beijing Institute of Microbiology and Epidemiology, Academy of Military Medical Sciences, China for giving valuable suggestions about the immunization of horse. We are especially grateful for the constructive advice about the immunization of mouse and monkey given by the late professor, Yusen Zhou at State Key Laboratory of Pathogen and Biosecurity, Beijing Institute of Microbiology and Epidemiology, Academy of Military Medical Sciences, China. This work was supported by grants from the State Key Program of National Natural Science Foundation of China (11932017 to YW), the National Natural Science Foundation of China (NSFC No.11421202,11827803 to YBF, and 8204100482 to YSZ), the National Key Research and Development Program of China (2018YFC1200100, and 2020YFC0860100 to ZPZ), National Science and Technology Major Project of the 13^th^ Five-Year Plan (2017ZX10304402003 to ZPZ),the Youth Thousand Scholar Program of China (J.Z.), Program for High-Level Overseas Talents, Beihang University (J.Z.) and Outstanding and innovative program in medicine and engineering, Beihang University (J.Z).

## Author contributions

L.L., W.D.L., T.S., L.W., Y.F.H., G.L., X.H.H., H.W., Y.L., Y.F.C., H.Y.W., J.L., Z.Y.S., C.D., Y.T.W., X.Y.L., Z.Q.Y., and J.W. performed the immune-informatics analysis, collected and mined literature, and filtered peptides. Z.P.Z., X.L.Y., S.L.C., M.L., and T.C.W. performed the biological experiments. Y.W., Y.B.F., Z.P.Z., H.W. and J.Z. conceived the project, supervised the work, and interpreted the data. L.L., Z.P.Z., X.L.Y., W.D.L., S.L.C., T.S., L.W., Y.F.H., G.L., X.H.H., H.W., Y.L., Y.F.C., H.Y.W., J.L., Z.Y.S., C.D., Y.T.W., X.Y.L., Z.Q.Y., J.W., M.L., T.C.W., Y.W., Y.B.F., H.W., and J.Z. drafted the manuscript, edited the draft and prepared the final manuscript, which was approved by all co-authors.

## Additional information

L.L., Z.P.Z, and X.L.Y. contribute equally to this work.

Correspondence should be addressed to Y.W., Y.B.F., H.W., or J.Z.

