## Supplementary Table 1 for "Identification of four linear B-cell epitopes on the SARS-CoV-2 spike protein able to elicit neutralizing antibodies"

### Description of supplementary materials

#### Supplementary Data

Supplementary  Table 1. The full list of linear B-cell epitopes from spike protein of SARS-CoV-2.

**Supplementary Table 1. The full list of linear B-cell epitopes from spike protein of SARS-CoV-2.**

| start | end | peptide | Antigenicity (score) | Peptide Length | methods | methods_number | Allergenicity | Toxicity | Hydrophobicity | Hydropathicity | Hydrophilicity | Charge | pI | Mol wt | topology |
|---|---|---|---|---|---|---|---|---|---|---|---|---|---|---|---|
| 538 | 547 | CVNFNFNGL | 1.7985 | 9 | BCEPRED | 1 | ALLERGEN | Non-Toxin | 0.06 | 0.58 | -0.97 | 0 | 5.85 | 1027.29 | outside |
| 369 | 394 | YNSASFSTFKCYGVSPTKLNDLCFT | 1.4031 | 25 | IEDB,literature | 2 | NON-ALLERGEN | Non-Toxin | -0.08 | -0.05 | -0.41 | 1 | 8.33 | 2794.49 | outside |
| 505 | 520 | YQPYRVVVLSFELLH | 0.9711 | 15 | BCEPRED,literature | 2 | NON-ALLERGEN | Non-Toxin | 0 | 0.47 | -0.73 | 0.5 | 7.1 | 1863.42 | outside |
| 404 | 427 | GDEVRQIAPGQTGKIADYNYKLP | 1.1017 | 23 | IEDB,literature | 2 | NON-ALLERGEN | Non-Toxin | -0.21 | -0.89 | 0.25 | 0 | 6.46 | 2534.17 | outside |
| 411 | 418 | APGQTGK | 1.7614 | 7 | IEDB | 1 | ALLERGEN | Non-Toxin | -0.21 | -1.24 | 0.33 | 1 | 9.11 | 657.82 | outside |
| 14 | 21 | QCVNLTT | 1.5916 | 7 | IEDB | 1 | ALLERGEN | Non-Toxin | -0.08 | 0.3 | -0.67 | 0 | 5.85 | 777.99 | outside |
| 1034 | 1040 | LGQSKR | 2.0011 | 6 | IEDB | 1 | ALLERGEN | Non-Toxin | -0.52 | -1.55 | 0.78 | 2 | 11.01 | 687.88 | outside |
| 1035 | 1043 | GQSKRVDF | 1.9298 | 8 | BCEPRED | 1 | ALLERGEN | Non-Toxin | -0.4 | -1.2 | 0.69 | 1 | 9.1 | 936.14 | outside |
| 407 | 421 | VRQIAPGQTGKIAD | 1.2606 | 14 | IEDB,literature | 2 | NON-ALLERGEN | Non-Toxin | -0.17 | -0.37 | 0.21 | 1 | 9.1 | 1453.87 | outside |
| 411 | 426 | APGQTGKIADYNYKL | 1.4441 | 15 | IEDB,literature | 2 | NON-ALLERGEN | Non-Toxin | -0.16 | -0.81 | -0.01 | 1 | 8.76 | 1639.06 | outside |
| 411 | 431 | APGQTGKIADYNYKLPDDFT | 1.0425 | 20 | ABCPRED,IEDB,literature | 3 | NON-ALLERGEN | Non-Toxin | -0.17 | -0.93 | 0.15 | -1 | 4.43 | 2214.7 | outside |
| 417 | 431 | KIADYNYKLPDDFT | 0.9567 | 14 | BCEPRED,literature | 2 | NON-ALLERGEN | Non-Toxin | -0.22 | -0.99 | 0.26 | -1 | 4.43 | 1703.07 | outside |
| 1035 | 1044 | GQSKRVDFC | 1.779 | 9 | IEDB,literature | 2 | ALLERGEN | Non-Toxin | -0.35 | -0.79 | 0.5 | 1 | 8.57 | 1039.29 | outside |
| 405 | 431 | DEVRQIAPGQTGKIADYNYKLPDDFT | 0.9773 | 26 | literature | 1 | NON-ALLERGEN | Non-Toxin | -0.23 | -0.96 | 0.34 | -2 | 4.37 | 2955.61 | outside |
| 1060 | 1069 | VVFLHVTYV | 1.5122 | 9 | literature | 1 | ALLERGEN | Non-Toxin | 0.3 | 2.02 | -1.5 | 0.5 | 7.09 | 1076.44 | outside |
| 585 | 599 | LDITPCSFGGVSVI | 1.6124 | 14 | ABCPRED,literature | 2 | NON-ALLERGEN | Non-Toxin | 0.18 | 1.31 | -0.62 | -1 | 3.8 | 1407.85 | outside |
| 386 | 399 | KLNDLCFTNVYAD | 1.6735 | 13 | literature | 1 | NON-ALLERGEN | Non-Toxin | -0.11 | -0.08 | -0.18 | -1 | 4.21 | 1515.88 | outside |
| 373 | 391 | SFSTFKCYGVSPTKLNDL | 1.0676 | 18 | literature | 1 | NON-ALLERGEN | Non-Toxin | -0.1 | -0.11 | -0.23 | 1 | 8.51 | 2007.54 | outside |
| 775 | 795 | DKNTQEVFAQVKQIYKTPPI | 0.3752 | 20 | ABCPRED | 1 | NON-ALLERGEN | Non-Toxin | -0.22 | -0.83 | 0.16 | 1 | 8.76 | 2348 | outside |
| 470 | 484 | TEIYQAGSTPCNGV | -0.1224 | 14 | ABCPRED | 1 | NON-ALLERGEN | Non-Toxin | -0.05 | -0.24 | -0.3 | -1 | 4 | 1439.76 | outside |
| 590 | 606 | CSFGGVSVITPGTNTS | 0.7052 | 16 | AAP | 1 | NON-ALLERGEN | Non-Toxin | 0.06 | 0.46 | -0.53 | 0 | 5.85 | 1526.91 | outside |
| 1146 | 1166 | DSFKEELDKYFKNHTSPDVD | -0.4964 | 20 | ABCPRED | 1 | ALLERGEN | Non-Toxin | -0.35 | -1.55 | 0.81 | -2.5 | 4.52 | 2414.84 | outside |
| 156 | 170 | EFRVYSSANNCTFE | 0.0642 | 14 | ABCPRED | 1 | NON-ALLERGEN | Non-Toxin | -0.21 | -0.57 | -0.05 | -1 | 4.54 | 1666.96 | outside |
| 453 | 467 | YRLFRKSNLKPFER | 0.113 | 14 | ABCPRED | 1 | ALLERGEN | Non-Toxin | -0.48 | -1.34 | 0.54 | 4 | 11.01 | 1854.38 | outside |
| 607 | 621 | QVAVLYQDVNCTEV | 0.9497 | 14 | ABCPRED | 1 | NON-ALLERGEN | Non-Toxin | -0.04 | 0.39 | -0.39 | -2 | 3.67 | 1580.98 | outside |
| 1089 | 1103 | FPREGVFVSNGTHW | 0.516 | 14 | ABCPRED | 1 | NON-ALLERGEN | Non-Toxin | -0.07 | -0.39 | -0.41 | 0.5 | 7.1 | 1633 | outside |
| 1143 | 1163 | PELDSFKEELDKYFKNHTSP | -0.5547 | 20 | ABCPRED | 1 | NON-ALLERGEN | Non-Toxin | -0.31 | -1.47 | 0.65 | -1.5 | 4.9 | 2424.93 | outside |
| 492 | 508 | LQSYGFQPTNGVGYQP | 0.5258 | 16 | ABCPRED,literature | 2 | NON-ALLERGEN | Non-Toxin | -0.07 | -0.73 | -0.61 | 0 | 5.87 | 1756.15 | outside |
| 1165 | 1181 | DLGDISGINASVVNIQ | 1.0543 | 16 | FBCPred | 1 | NON-ALLERGEN | Non-Toxin | 0.03 | 0.48 | -0.22 | -2 | 3.57 | 1615.02 | outside |
| 962 | 976 | LVKQLSSNFGAISS | 0.3662 | 14 | ABCPRED | 1 | NON-ALLERGEN | Non-Toxin | -0.01 | 0.46 | -0.38 | 1 | 9.11 | 1450.86 | outside |
| 1248 | 1268 | CSCGSCCKFDEDDSEPVLKG | 0.3036 | 20 | ABCPRED,literature | 2 | NON-ALLERGEN | Non-Toxin | -0.21 | -0.47 | 0.6 | -3 | 4.11 | 2122.6 | inside |
| 343 | 359 | NATRFASVYAWNRKRI | 0.2999 | 16 | ABCPRED,BCPREDS,FBCPred | 3 | NON-ALLERGEN | Non-Toxin | -0.32 | -0.7 | -0.04 | 4 | 11.73 | 1953.45 | outside |
| 306 | 326 | FTVEKGIYQTSNFRVQPTES | 0.6248 | 20 | ABCPRED | 1 | NON-ALLERGEN | Non-Toxin | -0.2 | -0.72 | 0 | 0 | 6.49 | 2331.86 | outside |
| 916 | 936 | LYENQKLIANQFNSAIGKIQ | 0.0495 | 20 | ABCPRED | 1 | NON-ALLERGEN | Non-Toxin | -0.13 | -0.37 | -0.21 | 1 | 8.83 | 2292.94 | outside |
| 340 | 354 | EVFNATRFASVYAW | -0.0464 | 14 | ABCPRED | 1 | NON-ALLERGEN | Non-Toxin | 0 | 0.3 | -0.65 | 0 | 6.35 | 1661.03 | outside |
| 33 | 49 | TRGVYYPDKVFRSSVL | 0.2413 | 16 | ABCPRED,AAP | 2 | NON-ALLERGEN | Non-Toxin | -0.2 | -0.26 | -0.07 | 2 | 9.72 | 1887.39 | outside |
| 437 | 457 | NSNNLDSKVGGNYNYLYRLF | 0.3061 | 20 | ABCPRED | 1 | ALLERGEN | Non-Toxin | -0.21 | -0.86 | -0.29 | 1 | 8.76 | 2351.86 | outside |
| 383 | 403 | SPTKLNDLCFTNVYADSFVI | 0.9482 | 20 | ABCPRED | 1 | NON-ALLERGEN | Non-Toxin | -0.02 | 0.33 | -0.4 | -1 | 4.21 | 2247.83 | outside |
| 337 | 351 | PFGEVFNATRFASV | 0.1918 | 14 | ABCPRED | 1 | NON-ALLERGEN | Non-Toxin | 0 | 0.39 | -0.39 | 0 | 6.36 | 1541.91 | outside |
| 743 | 765 | CGDSTECSNLLLQYGSFCTQLN | 0.5822 | 22 | AAP | 1 | NON-ALLERGEN | Non-Toxin | -0.09 | -0.06 | -0.37 | -2 | 3.67 | 2396.97 | outside |
| 150 | 170 | KSWMESEFRVYSSANNCTFE | 0.0349 | 20 | ABCPRED | 1 | NON-ALLERGEN | Non-Toxin | -0.23 | -0.8 | 0.06 | -1 | 4.79 | 2415.9 | outside |
| 1081 | 1101 | ICHDGKAHFPREGVFVSNGT | 0.0643 | 20 | ABCPRED | 1 | NON-ALLERGEN | Non-Toxin | -0.12 | -0.34 | -0.01 | 1 | 7.22 | 2171.71 | outside |
| 739 | 755 | TMYICGDSTECSNLLL | -0.2091 | 16 | ABCPRED | 1 | NON-ALLERGEN | Non-Toxin | 0 | 0.47 | -0.42 | -2 | 3.67 | 1763.26 | outside |
| 35 | 49 | GVYYPDKVFRSSVL | -0.1109 | 14 | ABCPRED | 1 | NON-ALLERGEN | Non-Toxin | -0.09 | 0.08 | -0.27 | 1 | 8.83 | 1630.07 | outside |
| 597 | 613 | VITPGTNTSNQVAVLY | 0.4189 | 16 | FBCPred | 1 | NON-ALLERGEN | Non-Toxin | 0.03 | 0.37 | -0.7 | 0 | 5.88 | 1677.13 | outside |
| 1199 | 1215 | DLQELGKYEQYIKWPW | 0.8261 | 16 | FBCPred | 1 | NON-ALLERGEN | Non-Toxin | -0.18 | -1.22 | -0.09 | -1 | 4.68 | 2096.62 | outside_transmembrane-region |
| 447 | 461 | GNYNYLYRLFRKSN | 0.0603 | 14 | ABCPRED | 1 | NON-ALLERGEN | Non-Toxin | -0.35 | -1.29 | -0.22 | 3 | 10 | 1808.23 | outside |
| 1133 | 1155 | VNNTVYDPLQPELDSFKEELDK | -0.2876 | 22 | BCPREDS | 1 | ALLERGEN | Non-Toxin | -0.25 | -1.03 | 0.51 | -4 | 4.02 | 2594.15 | outside |
| 628 | 642 | QLTPTWRVYSTGSN | 0.9924 | 14 | ABCPRED | 1 | NON-ALLERGEN | Non-Toxin | -0.19 | -0.81 | -0.44 | 1 | 9.1 | 1609.96 | outside |
| 680 | 700 | SPRRARSVASQSIIAYTMSL | 0.3956 | 20 | ABCPRED | 1 | ALLERGEN | Non-Toxin | -0.2 | -0.02 | -0.08 | 3 | 11.72 | 2194.81 | outside |
| 417 | 447 | KIADYNYKLPDDFTGCVIAWNSNNLDSKVG | 0.7009 | 30 | literature | 1 | NON-ALLERGEN | Non-Toxin | -0.13 | -0.47 | -0.02 | -1 | 4.58 | 3362.17 | outside |
| 653 | 673 | AEHVNNSYECDIPIGAGICA | 0.765 | 20 | ABCPRED | 1 | NON-ALLERGEN | Non-Toxin | 0 | 0.14 | -0.17 | -2.5 | 4.14 | 2076.57 | outside |
| 443 | 463 | SKVGGNYNYLYRLFRKSNLK | 0.1598 | 20 | ABCPRED | 1 | NON-ALLERGEN | Non-Toxin | -0.31 | -0.95 | 0 | 5 | 10.3 | 2421.1 | outside |
| 674 | 690 | YQTQTNSPRRARSVAS | 0.1927 | 16 | ABCPRED,BCPREDS | 2 | NON-ALLERGEN | Non-Toxin | -0.47 | -1.43 | 0.31 | 3 | 11.72 | 1822.18 | outside |
| 8 | 24 | LPLVSSQCVNLTTRTQ | 1.3416 | 16 | ABCPRED,FBCPred | 2 | NON-ALLERGEN | Non-Toxin | -0.14 | 0.12 | -0.4 | 1 | 8.6 | 1760.28 | outside |
| 424 | 444 | KLPDDFTGCVIAWNSNNLDS | 0.0285 | 20 | ABCPRED | 1 | NON-ALLERGEN | Non-Toxin | -0.11 | -0.34 | -0.07 | -2 | 3.94 | 2209.7 | outside |
| 841 | 855 | LGDIAARDLICAQK | 1.1817 | 14 | ABCPRED | 1 | ALLERGEN | Non-Toxin | -0.11 | 0.37 | 0.18 | 0 | 6.28 | 1486.95 | outside |
| 888 | 910 | FGAGAALQIPFAMQMAYRFNGI | 0.76 | 22 | literature | 1 | NON-ALLERGEN | Non-Toxin | 0.1 | 0.68 | -0.76 | 1 | 9.1 | 2375.13 | outside |
| 450 | 464 | NYLYRLFRKSNLKP | 0.2318 | 14 | ABCPRED | 1 | NON-ALLERGEN | Non-Toxin | -0.36 | -1.04 | 0.01 | 4 | 10.45 | 1812.35 | outside |
| 1089 | 1109 | FPREGVFVSNGTHWFVTQRN | 0.441 | 20 | ABCPRED | 1 | NON-ALLERGEN | Non-Toxin | -0.15 | -0.54 | -0.34 | 1.5 | 9.95 | 2378.93 | outside |
| 814 | 828 | KRSFIEDLLFNKVT | -0.041 | 14 | ABCPRED | 1 | ALLERGEN | Non-Toxin | -0.2 | -0.17 | 0.23 | 1 | 8.93 | 1710.21 | outside |
| 480 | 500 | CNGVEGFNCYFPLQSYGFQP | 0.5841 | 20 | ABCPRED | 1 | NON-ALLERGEN | Non-Toxin | -0.01 | -0.2 | -0.67 | -1 | 4 | 2270.81 | outside |
| 881 | 895 | TITSGWTFGAGAAL | 0.288 | 14 | ABCPRED | 1 | NON-ALLERGEN | Non-Toxin | 0.19 | 0.82 | -0.85 | 0 | 5.88 | 1352.71 | outside |
| 960 | 980 | NTLVKQLSSNFGAISSVLND | 0.0693 | 20 | ABCPRED | 1 | NON-ALLERGEN | Non-Toxin | -0.06 | 0.16 | -0.28 | 0 | 6.19 | 2107.65 | outside |
| 161 | 181 | SSANNCTFEYVSQPFLMDLE | 0.3881 | 20 | ABCPRED | 1 | ALLERGEN | Non-Toxin | -0.08 | -0.17 | -0.26 | -3 | 3.58 | 2295.8 | outside |
| 56 | 195 | WIFGTTLDSKTQSLLIVNNATNVVIKVCEFQFCN | 0.1404 | 103 | ellipro | 1 | NON-ALLERGEN | too long | | | | | | | outside |
| 87 | 101 | NDGVYFASTEKSNI | 0.4766 | 14 | ABCPRED | 1 | NON-ALLERGEN | Non-Toxin | -0.15 | -0.61 | 0.07 | -1 | 4.38 | 1544.83 | outside |
| 743 | 759 | CGDSTECSNLLLQYGS | 0.2817 | 16 | AAP | 1 | NON-ALLERGEN | Non-Toxin | -0.1 | -0.17 | -0.17 | -2 | 3.67 | 1690.06 | outside |
| 740 | 754 | MYICGDSTECSNLL | -0.2196 | 14 | ABCPRED | 1 | NON-ALLERGEN | Toxin | -0.03 | 0.32 | -0.33 | -2 | 3.67 | 1548.96 | outside |
| 410 | 432 | IAPGQTGKIADYNYKLPDDFTG | 1.1317 | 22 | BCPREDS | 1 | ALLERGEN | Toxin | -0.12 | -0.66 | 0.05 | -1 | 4.43 | 2384.95 | outside |
| 1234 | 1250 | LCCMTSCCSCLKGCCS | -0.2867 | 16 | ABCPRED | 1 | ALLERGEN | Non-Toxin | -0.02 | 1.22 | -0.53 | 1 | 7.99 | 1645.27 | transmembrane-region_inside |
| 1010 | 1030 | QQLIRAAEIRASANLAATKM | 0.301 | 20 | ABCPRED | 1 | ALLERGEN | Non-Toxin | -0.17 | 0.05 | 0.05 | 2 | 10.84 | 2156.82 | outside |
| 618 | 638 | TEVPVAIHADQLTPTWRVYS | 0.3834 | 20 | ABCPRED | 1 | NON-ALLERGEN | Non-Toxin | -0.07 | -0.1 | -0.35 | -0.5 | 5.33 | 2283.86 | outside |
| 578 | 592 | DPQTLEILDITPCS | 1.252 | 14 | ABCPRED | 1 | NON-ALLERGEN | Non-Toxin | -0.07 | -0.02 | 0.04 | -3 | 3.5 | 1544.94 | outside |
| 292 | 306 | ALDPLSETKCTLKS | 1.017 | 14 | ABCPRED | 1 | NON-ALLERGEN | Non-Toxin | -0.19 | -0.26 | 0.35 | 0 | 6.38 | 1505.94 | outside |

| | | | | | | | | | | | | | | | |
|---|---|---|---|---|---|---|---|---|---|---|---|---|---|---|---|
| 31 | 51 | SFTRGVYYPDKVFRSSVLHS | 0.0861 | 20 | ABCPRED | 1 | NON-ALLERGEN | Non-Toxin | -0.17 | -0.31 | -0.18 | 2.5 | 9.72 | 2345.92 | outside |
| 412 | 432 | PGQTGKIADYNYKLPDDFTG | 1.0123 | 20 | ABCPRED | 1 | NON-ALLERGEN | Non-Toxin | -0.18 | -1.04 | 0.17 | -1 | 4.43 | 2200.68 | outside |
| 1108 | 1124 | NFYEPQIITTDNTFVS | 0.4447 | 16 | literature | 1 | ALLERGEN | Non-Toxin | -0.06 | -0.28 | -0.42 | -2 | 3.67 | 1889.28 | outside |
| 538 | 558 | CVNFNFNGLTGTGVLTESNK | 1.1352 | 20 | ABCPRED | 1 | NON-ALLERGEN | Non-Toxin | -0.06 | -0.07 | -0.34 | 0 | 6.32 | 2115.65 | outside |
| 482 | 502 | GVEGFNCYFPLQSYGFQPTN | 0.9121 | 20 | ABCPRED | 1 | ALLERGEN | Non-Toxin | -0.02 | -0.35 | -0.64 | -1 | 4 | 2268.78 | outside |
| 330 | 344 | PNITNLCPFGEVFN | 0.7053 | 14 | ABCPRED | 1 | NON-ALLERGEN | Non-Toxin | 0.03 | 0.16 | -0.56 | -1 | 4 | 1564.98 | outside |
| 300 | 314 | KCTLKSFTVEKGIY | 0.1527 | 14 | ABCPRED | 1 | NON-ALLERGEN | Non-Toxin | -0.14 | -0.09 | 0.04 | 2 | 9.17 | 1617.14 | outside |
| 93 | 115 | ASTEKSNIIRGWIFGTTLDSKT | -0.0057 | 22 | AAP,BCPREDS | 2 | NON-ALLERGEN | Non-Toxin | -0.14 | -0.35 | 0.04 | 1 | 8.93 | 2426.05 | outside |
| 50 | 64 | STQDLFLPFFSNVT | 0.3443 | 14 | ABCPRED | 1 | NON-ALLERGEN | Non-Toxin | 0.03 | 0.36 | -0.67 | -1 | 3.8 | 1616 | outside |
| 61 | 81 | NVTWFHAIHVSGTNGTKRFD | 0.7547 | 20 | ABCPRED | 1 | ALLERGEN | Non-Toxin | -0.12 | -0.48 | -0.31 | 2 | 9.11 | 2287.82 | outside |
| 651 | 667 | IGAEHVNNSYECDIPI | 0.9115 | 16 | FBCPred | 1 | NON-ALLERGEN | Non-Toxin | -0.05 | -0.17 | -0.09 | -2.5 | 4.14 | 1774.17 | outside |
| 639 | 653 | GSNVFQTRAGCLIG | 0.8547 | 14 | ABCPRED | 1 | NON-ALLERGEN | Non-Toxin | -0.02 | 0.39 | -0.41 | 1 | 8.6 | 1422.83 | outside |
| 979 | 993 | DILSRLDKVEAEVQ | 0.3337 | 14 | ABCPRED | 1 | NON-ALLERGEN | Non-Toxin | -0.24 | -0.31 | 0.69 | -2 | 4.32 | 1615.02 | outside |
| 76 | 90 | TKRFDNPVLPFNDG | 0.2892 | 14 | ABCPRED | 1 | NON-ALLERGEN | Non-Toxin | -0.25 | -0.94 | 0.26 | 0 | 6.31 | 1619.99 | outside |
| 239 | 259 | QTLLALHRSYLTPGDSSSGW | 0.5163 | 20 | ABCPRED | 1 | ALLERGEN | Non-Toxin | -0.1 | -0.35 | -0.37 | 0.5 | 7.09 | 2189.71 | outside |
| 682 | 696 | RRARSVASQSIIAY | 0.3552 | 14 | ABCPRED | 1 | NON-ALLERGEN | Non-Toxin | -0.28 | -0.15 | 0.09 | 3 | 11.72 | 1577.99 | outside |
| 461 | 475 | LKPFERDISTEIYQ | -0.1738 | 14 | ABCPRED | 1 | NON-ALLERGEN | Non-Toxin | -0.24 | -0.8 | 0.35 | -1 | 4.68 | 1739.16 | outside |
| 1095 | 1123 | FVSNGTHWFVTQRNFYEPQIITTDNTFVSK | 0.3761 | 30 | literature | 1 | ALLERGEN | Non-Toxin | -0.11 | -0.42 | -0.42 | 0.5 | 7.1 | 3578.41 | outside |
| 568 | 590 | DIADTTDAVRDPQTLEILDITP | 1.191 | 22 | FBCPred | 1 | NON-ALLERGEN | Non-Toxin | -0.15 | -0.28 | 0.37 | -5 | 3.62 | 2412.95 | outside |
| 1224 | 1244 | LIAIVMVTIMLCCMTSCCSC | 0.1643 | 20 | ABCPRED | 1 | NON-ALLERGEN | Non-Toxin | 0.23 | 2.33 | -1.08 | 0 | 5.75 | 2139.09 | transmembrane-region_inside |
| 499 | 515 | PTNGVGYQPYRVVVLS | 0.5326 | 16 | AAP | 1 | NON-ALLERGEN | Non-Toxin | -0.04 | 0.06 | -0.57 | 1 | 8.93 | 1749.24 | outside |
| 1068 | 1088 | VPAQEKNFTTAPAICHDGKA | 0.3708 | 20 | ABCPRED | 1 | NON-ALLERGEN | Non-Toxin | -0.13 | -0.44 | 0.12 | 0.5 | 7.06 | 2098.64 | outside |
| 897 | 913 | PFAMQMAYRFNGIGVT | 1.3871 | 16 | ABCPRED | 1 | NON-ALLERGEN | Non-Toxin | 0.03 | 0.36 | -0.7 | 1 | 9.1 | 1803.36 | outside |
| 213 | 229 | VRDLPQGFSALEPLVD | 0.5953 | 16 | FBCPred | 1 | NON-ALLERGEN | Non-Toxin | -0.08 | 0.09 | 0.07 | -2 | 4.03 | 1756.22 | outside |
| 1029 | 1043 | MSECVLGQSKRVDF | 1.0345 | 14 | ABCPRED | 1 | NON-ALLERGEN | Non-Toxin | -0.2 | -0.11 | 0.23 | 0 | 6.39 | 1599.05 | outside |
| 720 | 734 | ISVTTEILPVSMTK | 1.286 | 14 | ABCPRED | 1 | NON-ALLERGEN | Non-Toxin | 0.03 | 0.74 | -0.31 | 0 | 6.35 | 1519.04 | outside |
| 226 | 246 | LVDLPIGINITRFQTLLALH | 0.867 | 20 | ABCPRED | 1 | NON-ALLERGEN | Non-Toxin | 0.09 | 0.99 | -0.69 | 0.5 | 7.1 | 2248.04 | outside |
| 298 | 312 | ETKCTLKSFTVEKG | 0.6533 | 14 | ABCPRED | 1 | NON-ALLERGEN | Non-Toxin | -0.25 | -0.62 | 0.52 | 1 | 8.51 | 1571.02 | outside |
| 640 | 660 | SNVFQTRAGCLIGAEHVNNS | 0.8696 | 20 | ABCPRED | 1 | NON-ALLERGEN | Non-Toxin | -0.11 | -0.14 | -0.23 | 0.5 | 7.07 | 2117.62 | outside |
| 890 | 904 | AGAALQIPFAMQMA | 0.7743 | 14 | ABCPRED | 1 | NON-ALLERGEN | Non-Toxin | 0.17 | 1.06 | -0.77 | 0 | 5.88 | 1419.92 | outside |
| 948 | 968 | LQDVVNQNAQALNTLVKQLS | 0.077 | 20 | ABCPRED | 1 | NON-ALLERGEN | Non-Toxin | -0.14 | -0.1 | -0.27 | 0 | 6.19 | 2196.81 | outside |
| 1129 | 1149 | VIGIVNNTVYDPLQPELDSF | 0.4442 | 20 | ABCPRED | 1 | ALLERGEN | Non-Toxin | 0.02 | 0.23 | -0.35 | -3 | 3.5 | 2233.81 | outside |
| 945 | 959 | LGKLQDVVNQNAQA | 0.456 | 14 | ABCPRED | 1 | NON-ALLERGEN | Non-Toxin | -0.17 | -0.41 | -0.04 | 0 | 6.19 | 1497.89 | outside |
| 536 | 550 | NKCVNFNFNGLTGT | 1.3691 | 14 | ABCPRED | 1 | NON-ALLERGEN | Non-Toxin | -0.1 | -0.29 | -0.45 | 1 | 8.57 | 1528.91 | outside |
| 528 | 548 | KKSTNLVKNKCVNFNFNGLT | 1.0818 | 20 | ABCPRED | 1 | ALLERGEN | Non-Toxin | -0.23 | -0.58 | 0 | 4 | 10.05 | 2269.95 | outside |
| 315 | 335 | TSNFRVQPTESIVRFPNITN | 0.5629 | 20 | ABCPRED | 1 | NON-ALLERGEN | Non-Toxin | -0.21 | -0.52 | -0.12 | 1 | 9.95 | 2320.88 | outside |
| 135 | 149 | FCNDPFLGVYYHKN | 0.5621 | 14 | ABCPRED | 1 | NON-ALLERGEN | Toxin | -0.07 | -0.44 | -0.57 | 0.5 | 7.05 | 1717.13 | outside |
| 429 | 443 | FTGCVIAWNSNNLD | 0.2471 | 14 | ABCPRED | 1 | NON-ALLERGEN | Non-Toxin | 0.01 | 0.2 | -0.64 | -1 | 3.8 | 1553.91 | outside |
| 753 | 773 | LLQYGSFCTQLNRALTGIAV | 0.7445 | 20 | ABCPRED | 1 | NON-ALLERGEN | Non-Toxin | 0.02 | 0.68 | -0.71 | 1 | 8.57 | 2168.85 | outside |
| 1160 | 1176 | TSPDVDLGDISGINAS | 1.0756 | 16 | AAP | 1 | NON-ALLERGEN | Non-Toxin | -0.05 | -0.04 | 0.14 | -3 | 3.43 | 1560.86 | outside |
| 796 | 816 | DFGGFNFSQILPDPSKPSKR | 0.6234 | 20 | ABCPRED | 1 | ALLERGEN | Non-Toxin | -0.22 | -0.88 | 0.26 | 1 | 8.93 | 2237.78 | outside |
| 766 | 780 | ALTGIAVEQDKNTQ | 0.795 | 14 | ABCPRED | 1 | NON-ALLERGEN | Non-Toxin | -0.17 | -0.51 | 0.19 | -1 | 4.38 | 1487.84 | outside |
| 280 | 296 | NENGTITDAVDCALDP | 0.5804 | 16 | ABCPRED | 1 | NON-ALLERGEN | Non-Toxin | -0.12 | -0.36 | 0.28 | -4 | 3.38 | 1647.95 | outside |
| 909 | 923 | IGVTQNVLYENQKL | 0.7174 | 14 | ABCPRED | 1 | NON-ALLERGEN | Non-Toxin | -0.11 | -0.24 | -0.31 | 0 | 6.35 | 1619.08 | outside |
| 1233 | 1249 | MLCCMTSCCSCLKGCC | -0.2449 | 16 | ABCPRED,AAP | 2 | NON-ALLERGEN | Non-Toxin | 0.01 | 1.39 | -0.62 | 1 | 7.99 | 1689.39 | transmembrane-region_inside |
| 1090 | 1106 | PREGVFVSNGTHWFVT | 0.3563 | 16 | BCPREDS | 1 | NON-ALLERGEN | Non-Toxin | -0.04 | -0.12 | -0.48 | 0.5 | 7.1 | 1833.27 | outside |
| 1067 | 1089 | YVPAQEKNFTTAPAICHDGKAH | 0.327 | 22 | AAP | 1 | NON-ALLERGEN | Non-Toxin | -0.14 | -0.6 | -0.02 | 1 | 7.22 | 2398.99 | outside |
| 544 | 564 | NGLTGTGVLTESNKKFLPFQ | 0.8196 | 20 | ABCPRED | 1 | NON-ALLERGEN | Non-Toxin | -0.09 | -0.32 | -0.16 | 1 | 8.94 | 2151.76 | outside |
| 354 | 374 | NRKRISNCVADYSVLYNSAS | 0.0274 | 20 | ABCPRED | 1 | NON-ALLERGEN | Non-Toxin | -0.27 | -0.5 | 0.03 | 2 | 9.25 | 2260.78 | outside |
| 1088 | 1102 | HFPREGVFVSNGTH | 0.4946 | 14 | ABCPRED | 1 | NON-ALLERGEN | Non-Toxin | -0.12 | -0.56 | -0.21 | 1 | 7.26 | 1583.93 | outside |
| 1052 | 1066 | FPQSAPHGVVFLHV | 0.1725 | 14 | ABCPRED | 1 | NON-ALLERGEN | Non-Toxin | 0.14 | 0.68 | -0.88 | 1 | 7.26 | 1534.99 | outside |
| 498 | 514 | QPTNGVGYQPYRVVVL | 0.4774 | 16 | ABCPRED,FBCPred | 2 | NON-ALLERGEN | Non-Toxin | -0.07 | -0.11 | -0.57 | 1 | 8.93 | 1790.3 | outside |
| 1070 | 1147 | REGVFVSNGTHWFVTQRNFYEPQIITTDNTFVS[illegible] | 0.3685 | 77 | ellipro | 1 | ALLERGEN | too long | | | | | | | outside |
| 649 | 669 | CLIGAEHVNNSYECDIPIGA | 1.0187 | 20 | ABCPRED | 1 | ALLERGEN | Non-Toxin | 0.01 | 0.25 | -0.24 | -2.5 | 4.14 | 2118.66 | outside |
| 559 | 573 | FLPFQQFGRDIADT | 0.233 | 14 | ABCPRED | 1 | NON-ALLERGEN | Non-Toxin | -0.09 | -0.19 | -0.19 | -1 | 4.21 | 1655.04 | outside |
| 577 | 591 | RDPQTLEILDITPC | 1.2642 | 14 | ABCPRED | 1 | NON-ALLERGEN | Non-Toxin | -0.17 | -0.29 | 0.23 | -2 | 4.03 | 1614.05 | outside |
| 103 | 117 | GWIFGTTLDSKTQS | 0.4831 | 14 | ABCPRED | 1 | NON-ALLERGEN | Non-Toxin | -0.07 | -0.37 | -0.28 | 0 | 6.19 | 1540.9 | outside |
| 598 | 620 | ITPGTNTSNQVAVLYQDVNCTE | 0.8412 | 22 | FBCPred | 1 | ALLERGEN | Toxin | -0.11 | -0.28 | -0.28 | -2 | 3.67 | 2367.9 | outside |
| 861 | 875 | LPPLLTDEMIAQYT | 0.3726 | 14 | ABCPRED | 1 | NON-ALLERGEN | Non-Toxin | 0.02 | 0.23 | -0.42 | -2 | 3.67 | 1605.09 | outside |
| 972 | 994 | AISSVLNDILSRLDKVEAEVQI | 0.1315 | 22 | FBCPred | 1 | NON-ALLERGEN | Non-Toxin | -0.08 | 0.42 | 0.14 | -2 | 4.32 | 2413.1 | outside |
| 763 | 777 | LNRALTGIAVEQDK | 0.3722 | 14 | ABCPRED | 1 | NON-ALLERGEN | Non-Toxin | -0.19 | -0.26 | 0.29 | 0 | 6.42 | 1527.95 | outside |
| 990 | 1010 | EVQIDRLITGRLQSLQTYVT | -0.2899 | 20 | ABCPRED | 1 | NON-ALLERGEN | Non-Toxin | -0.17 | -0.12 | -0.13 | 0 | 6.42 | 2333.99 | outside |
| 1129 | 1145 | VIGIVNNTVYDPLQPE | 0.7133 | 16 | ABCPRED,literature | 2 | NON-ALLERGEN | Non-Toxin | 0.01 | 0.14 | -0.38 | -2 | 3.67 | 1771.25 | outside |
| 824 | 838 | NKVTLADAGFIKQY | 0.5511 | 14 | ABCPRED | 1 | NON-ALLERGEN | Non-Toxin | -0.1 | -0.13 | -0.14 | 1 | 8.83 | 1568.01 | outside |
| 797 | 811 | FGGFNFSQILPDPS | 0.5927 | 14 | ABCPRED | 1 | NON-ALLERGEN | Non-Toxin | 0.05 | 0.04 | -0.51 | -1 | 3.8 | 1525.88 | outside |
| 568 | 584 | DIADTTDAVRDPQTLE | 0.6532 | 16 | ABCPRED,FBCPred | 2 | NON-ALLERGEN | Toxin | -0.27 | -0.82 | 0.68 | -4 | 3.72 | 1760.06 | outside |
| 693 | 713 | IAYTMSLGAENSVAYSNNSI | 0.7231 | 20 | ABCPRED | 1 | ALLERGEN | Non-Toxin | 0 | 0.17 | -0.5 | -1 | 4 | 2105.59 | outside |
| 806 | 821 | LPDPSKPSKRSFIED | 0.0476 | 15 | BCEPRED,literature | 2 | NON-ALLERGEN | Non-Toxin | -0.34 | -1.26 | 0.85 | 0 | 6.47 | 1716.12 | outside |
| 246 | 262 | RSYLTPGDSSSGWTAG | 0.4016 | 16 | BCPREDS,FBCPred | 2 | NON-ALLERGEN | Non-Toxin | -0.14 | -0.75 | -0.1 | 0 | 6.19 | 1641.93 | outside |
| 35 | 55 | GVYYPDKVFRSSVLHSTQDL | 0.119 | 20 | ABCPRED | 1 | ALLERGEN | Non-Toxin | -0.15 | -0.34 | -0.15 | 0.5 | 7.09 | 2311.87 | outside |
| 203 | 217 | IYSKHTPINLVRDL | 0.606 | 14 | ABCPRED | 1 | NON-ALLERGEN | Non-Toxin | -0.15 | -0.16 | -0.17 | 1.5 | 8.94 | 1669.17 | outside |
| 760 | 774 | CTQLNRALTGIAVE | 0.7783 | 14 | ABCPRED | 1 | NON-ALLERGEN | Non-Toxin | -0.07 | 0.4 | -0.24 | 0 | 6.32 | 1488.93 | outside |
| 752 | 772 | LLLQYGSFCTQLNRALTGIA | 0.7592 | 20 | ABCPRED | 1 | NON-ALLERGEN | Non-Toxin | 0.02 | 0.66 | -0.73 | 1 | 8.57 | 2182.88 | outside |
| 604 | 620 | TSNQVAVLYQDVNCTE | 0.7593 | 16 | ABCPRED | 1 | ALLERGEN | Non-Toxin | -0.13 | -0.24 | -0.24 | -2 | 3.67 | 1784.16 | outside |
| 338 | 358 | FGEVFNATRFASVYAWNRKR | 0.2936 | 20 | ABCPRED | 1 | ALLERGEN | Non-Toxin | -0.23 | -0.49 | -0.12 | 3 | 10.91 | 2420 | outside |
| 961 | 975 | TLVKQLSSNFGAIS | 0.4688 | 14 | ABCPRED | 1 | NON-ALLERGEN | Non-Toxin | 0 | 0.46 | -0.43 | 1 | 9.11 | 1464.89 | outside |
| 58 | 78 | FFSNVTWFHAIHVSGTNGTK | 0.669 | 20 | ABCPRED | 1 | ALLERGEN | Non-Toxin | 0.02 | 0.02 | -0.72 | 2 | 9.11 | 2250.8 | outside |

| | | | | | | | | | | | | | | | |
|---|---|---|---|---|---|---|---|---|---|---|---|---|---|---|---|
| 860 | 880 | VLPPLLTDEMIAQYTSALLA | 0.2806 | 20 | ABCPRED | 1 | NON-ALLERGEN | Non-Toxin | 0.11 | 0.89 | -0.59 | -2 | 3.67 | 2159.87 | outside |
| 878 | 898 | LAGTITSGWTFGAGAALQIP | 0.2138 | 20 | ABCPRED | 1 | ALLERGEN | Non-Toxin | 0.18 | 0.81 | -0.79 | 0 | 5.88 | 1932.51 | outside |
| 537 | 557 | KCVNFNFNGLTGTGVLTESN | 1.2355 | 20 | ABCPRED | 1 | ALLERGEN | Non-Toxin | -0.06 | -0.07 | -0.34 | 0 | 6.32 | 2115.65 | outside |
| 761 | 781 | TQLNRALTGIAVEQDKNTQE | 0.4131 | 20 | ABCPRED | 1 | NON-ALLERGEN | Non-Toxin | -0.29 | -0.95 | 0.35 | -1 | 4.68 | 2229.74 | outside |
| 1195 | 1211 | ESLIDLQELGKYEQYI | 0.6105 | 16 | ABCPRED | 1 | ALLERGEN | Non-Toxin | -0.13 | -0.52 | 0.13 | -3 | 4.01 | 1941.42 | outside |
| 993 | 1007 | IDRLITGRLQSLQT | -0.4976 | 14 | ABCPRED | 1 | NON-ALLERGEN | Non-Toxin | -0.22 | -0.12 | -0.01 | 1 | 9.95 | 1614.1 | outside |
| 1069 | 1089 | PAQEKNFTTAPAICHDGKAH | 0.2868 | 20 | ABCPRED | 1 | NON-ALLERGEN | Non-Toxin | -0.18 | -0.81 | 0.17 | 1 | 7.22 | 2136.65 | outside |
| 470 | 486 | TEIYQAGSTPCNGVEG | -0.0105 | 16 | ABCPRED | 1 | NON-ALLERGEN | Non-Toxin | -0.07 | -0.46 | -0.07 | -2 | 3.8 | 1625.96 | outside |
| 692 | 712 | IIAYTMSLGAENSVAYSNNS | 0.5973 | 20 | ABCPRED | 1 | ALLERGEN | Non-Toxin | 0 | 0.17 | -0.49 | -1 | 4 | 2105.59 | outside |
| 587 | 607 | ITPCSFGGVSVITPGTNTSN | 0.8062 | 20 | ABCPRED | 1 | NON-ALLERGEN | Non-Toxin | 0.04 | 0.3 | -0.52 | 0 | 5.85 | 1952.46 | outside |
| 809 | 829 | PSKPSKRSFIEDLLFNKVTL | 0.3112 | 20 | ABCPRED | 1 | NON-ALLERGEN | Non-Toxin | -0.2 | -0.36 | 0.25 | 2 | 9.72 | 2320.02 | outside |
| 834 | 854 | IKQYGDCLGDIAARDLICAQ | 0.4955 | 20 | ABCPRED | 1 | NON-ALLERGEN | Non-Toxin | -0.1 | 0.17 | 0.03 | -1 | 4.43 | 2166.79 | outside |
| 633 | 653 | WRVYSTGSNVFQTRAGCLIG | 0.6893 | 20 | ABCPRED | 1 | NON-ALLERGEN | Non-Toxin | -0.08 | 0.07 | -0.51 | 2 | 9.55 | 2215.81 | outside |
| 732 | 748 | TKTSVDCTMYICGDST | 0.0937 | 16 | ABCPRED | 1 | NON-ALLERGEN | Non-Toxin | -0.12 | -0.09 | -0.06 | -1 | 4.21 | 1725.15 | outside |
| 1154 | 1174 | KYFKNHTSPDVDLGDISGIN | 0.6427 | 20 | ABCPRED | 1 | ALLERGEN | Non-Toxin | -0.17 | -0.74 | 0.17 | -0.5 | 5.31 | 2220.72 | outside |
| 69 | 85 | HVSGTNGTKRFDNPVL | 0.4599 | 16 | ABCPRED,FBCPred | 2 | ALLERGEN | Non-Toxin | -0.21 | -0.73 | 0.07 | 1.5 | 9.1 | 1742.16 | outside |
| 299 | 319 | TKCTLKSFTVEKGIYQTSNF | 0.3214 | 20 | ABCPRED | 1 | NON-ALLERGEN | Non-Toxin | -0.16 | -0.39 | -0.1 | 2 | 9.17 | 2295.93 | outside |
| 8 | 22 | LPLVSSQCVNLTTR | 1.3565 | 14 | ABCPRED | 1 | NON-ALLERGEN | Non-Toxin | -0.09 | 0.44 | -0.44 | 1 | 8.6 | 1531.01 | outside |
| 1148 | 1168 | FKEELDKYFKNHTSPDVDLG | 0.0852 | 20 | ABCPRED | 1 | NON-ALLERGEN | Non-Toxin | -0.26 | -1.17 | 0.56 | -1.5 | 4.83 | 2382.9 | outside |
| 113 | 133 | KTQSLLIVNNATNVVIKVCE | 0.2896 | 20 | ABCPRED | 1 | NON-ALLERGEN | Non-Toxin | -0.05 | 0.51 | -0.27 | 1 | 8.54 | 2186.91 | outside |
| 257 | 271 | GWTAGAAAYYVGYL | 0.5673 | 14 | ABCPRED | 1 | NON-ALLERGEN | Non-Toxin | 0.2 | 0.61 | -1.14 | 0 | 5.87 | 1462.82 | outside |
| 606 | 620 | NQVAVLYQDVNCTE | 0.8274 | 14 | ABCPRED | 1 | NON-ALLERGEN | Non-Toxin | -0.12 | -0.16 | -0.26 | -2 | 3.67 | 1595.95 | outside |
| 882 | 902 | ITSGWTFGAGAALQIPFAMQ | 0.6031 | 20 | ABCPRED | 1 | NON-ALLERGEN | Non-Toxin | 0.16 | 0.73 | -0.86 | 0 | 5.88 | 2067.69 | outside |
| 832 | 846 | GFIKQYGDCLGDIA | -0.1475 | 14 | ABCPRED | 1 | NON-ALLERGEN | Non-Toxin | 0.01 | 0.21 | -0.18 | -1 | 4.21 | 1499.91 | outside |
| 1097 | 1117 | SNGTHWFVTQRNFYEPQIIT | 0.2369 | 20 | ABCPRED | 1 | NON-ALLERGEN | Toxin | -0.13 | -0.68 | -0.52 | 0.5 | 7.1 | 2438.99 | outside |
| 464 | 480 | FERDISTEIYQAGSTP | -0.2904 | 16 | ABCPRED | 1 | NON-ALLERGEN | Non-Toxin | -0.18 | -0.7 | 0.19 | -2 | 4.14 | 1814.16 | outside |
| 701 | 721 | AENSVAYSNNSIAIPTNFTI | 0.6168 | 20 | ABCPRED | 1 | ALLERGEN | Non-Toxin | -0.01 | 0.08 | -0.47 | -1 | 4 | 2126.59 | outside |
| 523 | 537 | TVCGPKKSTNLVKN | 0.0085 | 14 | ABCPRED | 1 | NON-ALLERGEN | Non-Toxin | -0.25 | -0.59 | 0.22 | 3 | 9.81 | 1488.97 | outside |
| 259 | 273 | TAGAAAYYVGYLQP | 0.7174 | 14 | ABCPRED | 1 | NON-ALLERGEN | Non-Toxin | 0.11 | 0.34 | -0.89 | 0 | 5.87 | 1444.8 | outside |
| 415 | 431 | TGKIADYNYKLPDDFT | 0.9642 | 16 | ABCPRED | 1 | NON-ALLERGEN | Non-Toxin | -0.19 | -0.93 | 0.2 | -1 | 4.43 | 1861.26 | outside |
| 729 | 749 | VSMTKTSVDCTMYICGDSTE | 0.3832 | 20 | ABCPRED | 1 | NON-ALLERGEN | Non-Toxin | -0.1 | 0.02 | -0.02 | -2 | 4.03 | 2171.73 | outside |
| 1135 | 1155 | NTVYDPLQPELDSFKEELDKYKKKKK | -0.1996 | 26 | literature | 1 | NON-ALLERGEN | Non-Toxin | -0.42 | -1.7 | 0.97 | 1 | 8.56 | 3185.02 | outside |
| 1068 | 1097 | VPAQEKNFTTAPAICHDGKAHFPREGVFV | 0.2211 | 29 | literature | 1 | ALLERGEN | Non-Toxin | -0.11 | -0.28 | -0.01 | 1 | 7.22 | 3168.01 | outside |
| 357 | 377 | RISNCVADYSVLYNSASFST | -0.0478 | 20 | ABCPRED | 1 | NON-ALLERGEN | Non-Toxin | -0.09 | 0.16 | -0.41 | 0 | 6.15 | 2197.67 | outside |
| 240 | 254 | TLLALHRSYLTPGD | 0.8079 | 14 | ABCPRED | 1 | NON-ALLERGEN | Non-Toxin | -0.07 | 0.02 | -0.36 | 0.5 | 7.09 | 1556.99 | outside |
| 916 | 930 | LYENQKLIANQFNS | -0.1448 | 14 | ABCPRED | 1 | ALLERGEN | Non-Toxin | -0.19 | -0.74 | -0.24 | 0 | 6.35 | 1682.08 | outside |
| 386 | 406 | KLNDLCFTNVYADSFVIRGD | 0.932 | 20 | ABCPRED | 1 | NON-ALLERGEN | Non-Toxin | -0.11 | 0.06 | -0.09 | -1 | 4.43 | 2290.86 | outside |
| 122 | 142 | NATNVVIKVCEFQFCNDPFL | -0.1646 | 20 | ABCPRED | 1 | NON-ALLERGEN | Non-Toxin | -0.01 | 0.45 | -0.43 | -1 | 4.38 | 2301.95 | outside |
| 1101 | 1121 | HWFVTQRNFYEPQIITTDNT | 0.5486 | 20 | ABCPRED | 1 | NON-ALLERGEN | Non-Toxin | -0.17 | -0.82 | -0.41 | -0.5 | 5.33 | 2511.05 | outside |
| 77 | 91 | KRFDNPVLPFNDGV | 0.239 | 14 | ABCPRED | 1 | NON-ALLERGEN | Non-Toxin | -0.2 | -0.59 | 0.19 | 0 | 6.31 | 1618.02 | outside |
| 290 | 304 | DCALDPLSETKCTL | 0.7415 | 14 | ABCPRED | 1 | NON-ALLERGEN | Non-Toxin | -0.14 | 0 | 0.26 | -2 | 4.03 | 1508.91 | outside |
| 287 | 301 | DAVDCALDPLSETK | 0.8368 | 14 | ABCPRED | 1 | NON-ALLERGEN | Non-Toxin | -0.16 | -0.22 | 0.56 | -3 | 3.84 | 1476.8 | outside |
| 965 | 985 | QLSSNFGAISSVLNDILSRL | 0.1593 | 20 | ABCPRED | 1 | NON-ALLERGEN | Non-Toxin | -0.03 | 0.51 | -0.36 | 0 | 6.19 | 2134.72 | outside |
| 57 | 73 | PFFSNVTWFHAIHVSG | 0.4499 | 16 | AAP | 1 | NON-ALLERGEN | Non-Toxin | 0.14 | 0.5 | -1.05 | 1 | 7.26 | 1846.31 | outside |
| 863 | 877 | PLLTDEMIAQYTSA | 0.1459 | 14 | ABCPRED | 1 | NON-ALLERGEN | Non-Toxin | -0.01 | 0.14 | -0.31 | -2 | 3.67 | 1552.96 | outside |
| 55 | 69 | FLPFFSNVTWFHAI | 0.4357 | 14 | ABCPRED | 1 | NON-ALLERGEN | Non-Toxin | 0.24 | 1.06 | -1.39 | 0.5 | 7.1 | 1726.21 | outside |
| 512 | 532 | VLSFELLHAPATVCGPKKST | 0.3253 | 20 | ABCPRED | 1 | NON-ALLERGEN | Non-Toxin | -0.01 | 0.38 | -0.23 | 1.5 | 8.54 | 2098.78 | outside |
| 795 | 809 | KDFGGFNFSQILPD | 0.4321 | 14 | ABCPRED | 1 | NON-ALLERGEN | Non-Toxin | -0.06 | -0.31 | -0.1 | -1 | 4.21 | 1584.95 | outside |
| 757 | 777 | GSFCTQLNRALTGIAVEQDK | 0.6153 | 20 | ABCPRED | 1 | NON-ALLERGEN | Toxin | -0.15 | -0.19 | 0.04 | 0 | 6.39 | 2151.72 | outside |
| 144 | 158 | YYHKNNKSWMESEF | 0.4604 | 14 | ABCPRED | 1 | NON-ALLERGEN | Non-Toxin | -0.31 | -1.81 | 0.05 | 0.5 | 7.09 | 1863.23 | outside |
| 618 | 632 | TEVPVAIHADQLTP | 0.2341 | 14 | ABCPRED | 1 | NON-ALLERGEN | Non-Toxin | -0.01 | 0.14 | -0.19 | -1.5 | 4.36 | 1490.88 | outside |
| 1137 | 1157 | VYDPLQPELDSFKEELDKYF | -0.5197 | 20 | ABCPRED | 1 | NON-ALLERGEN | Non-Toxin | -0.2 | -0.89 | 0.4 | -4 | 4.02 | 2476.02 | outside |
| 301 | 315 | CTLKSFTVEKGIYQ | -0.0869 | 14 | ABCPRED | 1 | NON-ALLERGEN | Non-Toxin | -0.11 | -0.06 | -0.16 | 1 | 8.51 | 1617.1 | outside |
| 779 | 801 | QEVFAQVKQIYKTPPIKDFGGF | -0.0911 | 22 | AAP | 1 | NON-ALLERGEN | Non-Toxin | -0.09 | -0.35 | -0.08 | 1 | 8.76 | 2541.28 | outside |
| 1021 | 1035 | SANLAATKMSECVL | 0.4064 | 14 | ABCPRED | 1 | NON-ALLERGEN | Non-Toxin | -0.03 | 0.6 | -0.18 | 0 | 6.32 | 1437.88 | outside |
| 594 | 614 | GVSVITPGTNTSNQVAVLYQ | 0.4621 | 20 | ABCPRED | 1 | NON-ALLERGEN | Non-Toxin | 0.01 | 0.28 | -0.61 | 0 | 5.88 | 2048.59 | outside |
| 232 | 252 | GINITRFQTLLALHRSYLTP | 0.6602 | 20 | ABCPRED | 1 | NON-ALLERGEN | Non-Toxin | -0.07 | 0.17 | -0.56 | 2.5 | 10.84 | 2315.03 | outside |
| 274 | 294 | TFLLKYNENGTITDAVDCAL | 0.5434 | 20 | ABCPRED | 1 | NON-ALLERGEN | Toxin | -0.04 | 0.19 | -0.21 | -2 | 4.03 | 2201.77 | outside |
| 1025 | 1039 | AATKMSECVLGQSK | 0.3171 | 14 | ABCPRED | 1 | NON-ALLERGEN | Non-Toxin | -0.16 | -0.11 | 0.2 | 1 | 8.54 | 1452.9 | outside |
| 54 | 68 | LFLPFFSNVTWFHA | 0.2973 | 14 | ABCPRED | 1 | NON-ALLERGEN | Non-Toxin | 0.22 | 1.01 | -1.39 | 0.5 | 7.1 | 1726.21 | outside |
| 827 | 847 | TLADAGFIKQYGDCLGDIAA | 0.2227 | 20 | ABCPRED | 1 | ALLERGEN | Non-Toxin | 0.03 | 0.4 | -0.16 | -2 | 3.94 | 2042.58 | outside |
| 90 | 104 | VYFASTEKSNIIRG | 0.2015 | 14 | ABCPRED | 1 | NON-ALLERGEN | Non-Toxin | -0.13 | -0.11 | -0.07 | 1 | 8.93 | 1584.99 | outside |
| 831 | 847 | AGFIKQYGDCLGDIAA | -0.0348 | 16 | ABCPRED,FBCPred | 2 | NON-ALLERGEN | Non-Toxin | 0.04 | 0.41 | -0.22 | -1 | 4.21 | 1642.09 | outside |
| 121 | 141 | NNATNVVIKVCEFQFCNDPF | -0.2748 | 20 | ABCPRED | 1 | NON-ALLERGEN | Non-Toxin | -0.07 | 0.08 | -0.33 | -1 | 4.38 | 2302.89 | outside |
| 438 | 458 | SNNLDSKVGGNYNYLYRLFR | 0.0982 | 20 | ABCPRED | 1 | NON-ALLERGEN | Non-Toxin | -0.27 | -0.92 | -0.14 | 2 | 9.55 | 2393.94 | outside |
| 319 | 339 | RVQPTESIVRFPNITNLCPF | 0.7177 | 20 | ABCPRED | 1 | ALLERGEN | Non-Toxin | -0.13 | -0.04 | -0.27 | 1 | 8.6 | 2332.01 | outside |
| 241 | 255 | LLALHRSYLTPGDS | 0.562 | 14 | ABCPRED | 1 | NON-ALLERGEN | Non-Toxin | -0.08 | 0.01 | -0.31 | 0.5 | 7.09 | 1542.96 | outside |
| 439 | 459 | NNLDSKVGGNYNYLYRLFRK | 0.1093 | 20 | ABCPRED | 1 | NON-ALLERGEN | Non-Toxin | -0.31 | -1.07 | -0.01 | 3 | 9.83 | 2435.04 | outside |
| 303 | 323 | LKSFTVEKGIYQTSNFRVQP | 0.4756 | 20 | ABCPRED | 1 | ALLERGEN | Non-Toxin | -0.19 | -0.51 | -0.07 | 2 | 9.72 | 2342.98 | outside |
| 829 | 843 | ADAGFIKQYGDCLG | 0.2099 | 14 | ABCPRED | 1 | NON-ALLERGEN | Non-Toxin | -0.02 | 0.02 | -0.09 | -1 | 4.21 | 1457.82 | outside |
| 585 | 605 | LDITPCSFGGVSVITPGTNT | 1.0966 | 20 | ABCPRED | 1 | NON-ALLERGEN | Non-Toxin | 0.07 | 0.53 | -0.49 | -1 | 3.8 | 1979.53 | outside |
| 1007 | 1027 | YVTQQLIRAAEIRASANLAA | 0.2351 | 20 | ABCPRED | 1 | NON-ALLERGEN | Non-Toxin | -0.1 | 0.29 | -0.22 | 1 | 9.1 | 2159.76 | outside |
| 220 | 240 | FSALEPLVDLPIGINITRFQ | 0.7946 | 20 | ABCPRED | 1 | NON-ALLERGEN | Non-Toxin | 0.05 | 0.65 | -0.42 | -1 | 4.38 | 2243.94 | outside |
| 313 | 333 | YQTSNFRVQPTESIVRFPNI | 0.5074 | 20 | ABCPRED | 1 | ALLERGEN | Non-Toxin | -0.2 | -0.55 | -0.21 | 1 | 9.1 | 2396.98 | outside |
| 17 | 31 | NLTTRTQLPPAYTN | 1.1058 | 14 | ABCPRED | 1 | NON-ALLERGEN | Non-Toxin | -0.23 | -0.92 | -0.31 | 1 | 9.1 | 1589.97 | outside |
| 806 | 820 | LPDPSKPSKRSFIE | 0.1516 | 14 | ABCPRED | 1 | NON-ALLERGEN | Non-Toxin | -0.32 | -1.1 | 0.7 | 1 | 8.93 | 1601.02 | outside |
| 229 | 243 | LPIGINITRFQTLL | 0.8157 | 14 | ABCPRED | 1 | NON-ALLERGEN | Non-Toxin | 0.07 | 0.91 | -0.76 | 1 | 10.11 | 1599.18 | outside |

| | | | | | | | | | | | | | | | |
|---|---|---|---|---|---|---|---|---|---|---|---|---|---|---|---|
| 722 | 738 | VTTEILPVSMTKTSVD | 1.1143 | 16 | ABCPRED,AAP | 2 | ALLERGEN | Non-Toxin | -0.04 | 0.37 | -0.09 | -1 | 4.38 | 1721.23 | outside |
| 225 | 245 | PLVDLPIGINITRFQTLLAL | 0.7693 | 20 | ABCPRED | 1 | NON-ALLERGEN | Non-Toxin | 0.1 | 1.07 | -0.67 | 0 | 6.19 | 2208.01 | outside |
| 725 | 739 | EILPVSMTKTSVDC | 1.617 | 14 | ABCPRED | 1 | NON-ALLERGEN | Non-Toxin | -0.05 | 0.4 | -0.01 | -1 | 4.38 | 1522.99 | outside |
| 607 | 623 | QVAVLYQDVNCTEVPV | 0.6886 | 16 | AAP | 1 | NON-ALLERGEN | Non-Toxin | 0 | 0.5 | -0.43 | -2 | 3.67 | 1777.26 | outside |
| 200 | 216 | YFKIYSKHTPINLVRD | 0.657 | 16 | ABCPRED | 1 | NON-ALLERGEN | Toxin | -0.19 | -0.53 | -0.15 | 2.5 | 9.55 | 1994.56 | outside |
| 937 | 951 | SLSSTASALGKLQD | 0.8857 | 14 | ABCPRED | 1 | NON-ALLERGEN | Non-Toxin | -0.11 | -0.01 | 0.04 | 0 | 6.19 | 1377.71 | outside |
| 930 | 950 | AIGKIQDSLSSTASALGKLQ | 0.631 | 20 | ABCPRED | 1 | NON-ALLERGEN | Non-Toxin | -0.07 | 0.14 | -0.02 | 1 | 8.94 | 1988.57 | outside |
| 910 | 924 | GVTQNVLYENQKLI | 0.2141 | 14 | ABCPRED | 1 | NON-ALLERGEN | Non-Toxin | -0.11 | -0.24 | -0.31 | 0 | 6.35 | 1619.08 | outside |
| 1229 | 1249 | MVTIMLCCMTSCCSCLKGCC | -0.2013 | 20 | ABCPRED | 1 | NON-ALLERGEN | Non-Toxin | 0.08 | 1.61 | -0.75 | 1 | 7.99 | 2134.05 | transmembrane-region_inside |
| 151 | 165 | SWMESEFRVYSSAN | 0.3605 | 14 | ABCPRED | 1 | NON-ALLERGEN | Non-Toxin | -0.19 | -0.69 | -0.08 | -1 | 4.54 | 1693 | outside |
| 1092 | 1106 | EGVFVSNGTHWFVT | 0.2346 | 14 | ABCPRED | 1 | NON-ALLERGEN | Non-Toxin | 0.09 | 0.29 | -0.76 | -0.5 | 5.25 | 1579.94 | outside |
| 772 | 786 | VEQDKNTQEVFAQV | 0.0998 | 14 | ABCPRED | 1 | NON-ALLERGEN | Non-Toxin | -0.25 | -0.85 | 0.35 | -2 | 4.14 | 1634.97 | outside |
| 19 | 35 | TTRTQLPPAYTNSFTR | 0.3477 | 16 | ABCPRED,BCPREDS,IEDB | 3 | NON-ALLERGEN | Non-Toxin | -0.3 | -1.03 | -0.15 | 2 | 10.84 | 1854.27 | outside |
| 304 | 318 | KSFTVEKGIYQTSN | 0.1422 | 14 | ABCPRED | 1 | NON-ALLERGEN | Non-Toxin | -0.21 | -0.82 | 0.08 | 1 | 8.83 | 1601.98 | outside |
| 189 | 203 | LREFVFKNIDGYFK | -0.3638 | 14 | ABCPRED | 1 | NON-ALLERGEN | Non-Toxin | -0.15 | -0.26 | 0.02 | 1 | 8.83 | 1776.27 | outside |
| 213 | 227 | VRDLPQGFSALEPL | 0.9407 | 14 | ABCPRED | 1 | NON-ALLERGEN | Non-Toxin | -0.07 | 0.06 | -0.03 | -1 | 4.38 | 1541.97 | outside |
| 554 | 568 | ESNKKFLPFQQFGR | 0.4184 | 14 | ABCPRED | 1 | NON-ALLERGEN | Non-Toxin | -0.32 | -1.21 | 0.26 | 2 | 10.01 | 1726.17 | outside |
| 847 | 863 | RDLICAQKFNGLTVLP | 0.4118 | 16 | ABCPRED | 1 | NON-ALLERGEN | Non-Toxin | -0.08 | 0.35 | -0.23 | 1 | 8.57 | 1788.38 | outside |
| 172 | 186 | SQPFLMDLEGKQGN | 0.9603 | 14 | ABCPRED | 1 | ALLERGEN | Toxin | -0.18 | -0.88 | 0.18 | -1 | 4.38 | 1563.96 | outside |
| 163 | 185 | ANNCTFEYVSQPFLMDLEGKQG | 0.5793 | 22 | AAP | 1 | NON-ALLERGEN | Toxin | -0.11 | -0.45 | -0.11 | -2 | 4.14 | 2492.1 | outside |
| 634 | 648 | RVYSTGSNVFQTRA | 0.3248 | 14 | ABCPRED | 1 | NON-ALLERGEN | Non-Toxin | -0.26 | -0.55 | -0.15 | 2 | 10.84 | 1585.93 | outside |
| 221 | 235 | SALEPLVDLPIGIN | 0.6695 | 14 | ABCPRED | 1 | NON-ALLERGEN | Non-Toxin | 0.12 | 0.82 | -0.32 | -2 | 3.67 | 1450.91 | outside |
| 30 | 50 | NSFTRGVYYPDKVFRSSVLH | -0.0735 | 20 | ABCPRED | 1 | NON-ALLERGEN | Non-Toxin | -0.19 | -0.44 | -0.18 | 2.5 | 9.72 | 2372.95 | outside |
| 1059 | 1073 | GVVFLHVTYVPAQE | 1.0085 | 14 | ABCPRED | 1 | NON-ALLERGEN | Non-Toxin | 0.13 | 0.79 | -0.77 | -0.5 | 5.25 | 1559.01 | outside |
| 312 | 332 | IYQTSNFRVQPTESIVRFPN | 0.2983 | 20 | ABCPRED | 1 | NON-ALLERGEN | Non-Toxin | -0.2 | -0.55 | -0.21 | 1 | 9.1 | 2396.98 | outside |
| 25 | 41 | PPAYTNSFTRGVYYPD | 0.1245 | 16 | ABCPRED,FBCPred | 2 | NON-ALLERGEN | Non-Toxin | -0.15 | -0.88 | -0.36 | 0 | 6.18 | 1848.21 | outside |
| 56 | 196 | WIFGTTLDSKTQSLLIVNNATNVVIKVCEFQFCN | 0.112 | 99 | ellipro | 1 | NON-ALLERGEN | too long | | | | | | | outside |
| 165 | 179 | NCTFEYVSQPFLMD | 0.423 | 14 | ABCPRED | 1 | ALLERGEN | Non-Toxin | -0.04 | -0.03 | -0.47 | -2 | 3.67 | 1694.1 | outside |
| 1161 | 1181 | SPDVDLGDISGINASVVNIQ | 1.0657 | 20 | ABCPRED | 1 | NON-ALLERGEN | Non-Toxin | 0 | 0.29 | -0.08 | -3 | 3.43 | 2013.49 | outside |
| 579 | 599 | PQTLEILDITPCSFGGVSVI | 1.3978 | 20 | ABCPRED | 1 | ALLERGEN | Non-Toxin | 0.11 | 0.86 | -0.47 | -2 | 3.67 | 2089.74 | outside |
| 804 | 824 | QILPDPSKPSKRSFIEDLLF | 0.1014 | 20 | ABCPRED | 1 | NON-ALLERGEN | Non-Toxin | -0.17 | -0.37 | 0.25 | 0 | 6.47 | 2331 | outside |
| 315 | 331 | TSNFRVQPTESIVRFP | 0.2732 | 16 | ABCPRED,AAP | 2 | NON-ALLERGEN | Non-Toxin | -0.22 | -0.45 | -0.04 | 1 | 9.95 | 1878.34 | outside |
| 774 | 788 | QDKNTQEVFAQVKQ | 0.4896 | 14 | ABCPRED | 1 | NON-ALLERGEN | Non-Toxin | -0.37 | -1.43 | 0.47 | 0 | 6.42 | 1663.03 | outside |
| 750 | 770 | SNLLLQYGSFCTQLNRALTG | 0.5992 | 20 | ABCPRED | 1 | ALLERGEN | Non-Toxin | -0.07 | 0.12 | -0.59 | 1 | 8.57 | 2199.82 | outside |
| 187 | 207 | KNLREFVFKNIDGYFKIYSK | 0.0312 | 20 | ABCPRED | 1 | NON-ALLERGEN | Non-Toxin | -0.22 | -0.63 | 0.14 | 3 | 9.71 | 2510.23 | outside |
| 245 | 265 | HRSYLTPGDSSSGWTAGAAA | 0.5037 | 20 | ABCPRED | 1 | NON-ALLERGEN | Non-Toxin | -0.1 | -0.49 | -0.18 | 0.5 | 7.09 | 1992.36 | outside |
| 255 | 275 | SSGWTAGAAAYYVGYLQPRT | 0.6473 | 20 | ABCPRED | 1 | NON-ALLERGEN | Non-Toxin | -0.02 | -0.17 | -0.63 | 1 | 8.83 | 2119.6 | outside |
| 643 | 657 | FQTRAGCLIGAEHV | 1.27 | 14 | ABCPRED | 1 | NON-ALLERGEN | Non-Toxin | -0.03 | 0.37 | -0.31 | 0.5 | 7.07 | 1501.93 | outside |
| 304 | 326 | KSFTVEKGIYQTSNFRVQPTES | 0.5575 | 22 | AAP | 1 | NON-ALLERGEN | Non-Toxin | -0.25 | -0.87 | 0.15 | 1 | 8.83 | 2547.14 | outside |
| 530 | 550 | STNLVKNKCVNFNFNGLTGT | 1.0756 | 20 | ABCPRED | 1 | ALLERGEN | Non-Toxin | -0.12 | -0.25 | -0.33 | 2 | 9.36 | 2171.76 | outside |
| 833 | 853 | FIKQYGDCLGDIAARDLICA | 0.2604 | 20 | ABCPRED | 1 | NON-ALLERGEN | Non-Toxin | -0.03 | 0.49 | -0.1 | -1 | 4.43 | 2185.83 | outside |
| 1195 | 1215 | ESLIDLQELGKYEQYIKWPW | 0.8237 | 20 | ABCPRED | 1 | NON-ALLERGEN | Non-Toxin | -0.12 | -0.78 | -0.08 | -2 | 4.41 | 2539.2 | outside_transmembrane-region |
| 600 | 616 | PGTNTSNQVAVLYQDV | 0.3366 | 16 | ABCPRED | 1 | ALLERGEN | Non-Toxin | -0.09 | -0.3 | -0.36 | -1 | 3.8 | 1706.08 | outside |
| 204 | 218 | YSKHTPINLVRDLP | 0.4022 | 14 | ABCPRED | 1 | NON-ALLERGEN | Non-Toxin | -0.2 | -0.59 | -0.04 | 1.5 | 8.94 | 1653.12 | outside |
| 508 | 522 | YRVVVLSFELLHAP | 0.8065 | 14 | ABCPRED | 1 | NON-ALLERGEN | Non-Toxin | 0.07 | 0.98 | -0.67 | 0.5 | 7.1 | 1643.17 | outside |
| 65 | 85 | FHAIHVSGTNGTKRFDNPVL | 0.5642 | 20 | ABCPRED | 1 | NON-ALLERGEN | Non-Toxin | -0.11 | -0.29 | -0.21 | 2 | 9.11 | 2210.78 | outside |
| 476 | 492 | GSTPCNGVEGFNCYFP | 0.2489 | 16 | ABCPRED,FBCPred | 2 | NON-ALLERGEN | Non-Toxin | -0.01 | -0.18 | -0.47 | -1 | 4 | 1692.07 | outside |
| 917 | 931 | YENQKLIANQFNSA | -0.0151 | 14 | ABCPRED | 1 | ALLERGEN | Non-Toxin | -0.21 | -0.88 | -0.15 | 0 | 6.35 | 1639.99 | outside |
| 1247 | 1263 | CCSCGSCCKFDEDDSE | 0.1273 | 16 | ABCPRED | 1 | NON-ALLERGEN | Non-Toxin | -0.27 | -0.56 | 0.71 | -4 | 3.77 | 1731.03 | inside |
| 1105 | 1125 | TQRNFYEPQIITTDNTFVSG | 0.415 | 20 | ABCPRED | 1 | ALLERGEN | Non-Toxin | -0.17 | -0.68 | -0.2 | -1 | 4.38 | 2331.82 | outside |
| 572 | 588 | TTDAVRDPQTLEILDI | 0.8254 | 16 | ABCPRED,AAP | 2 | NON-ALLERGEN | Non-Toxin | -0.16 | -0.19 | 0.3 | -3 | 3.84 | 1800.23 | outside |
| 611 | 625 | LYQDVNCTEVPVAI | 0.6816 | 14 | ABCPRED | 1 | NON-ALLERGEN | Non-Toxin | 0.02 | 0.54 | -0.42 | -2 | 3.67 | 1563.99 | outside |
| 988 | 1002 | EAEVQIDRLITGRL | -0.5103 | 14 | ABCPRED | 1 | NON-ALLERGEN | Non-Toxin | -0.21 | -0.11 | 0.4 | -1 | 4.68 | 1613.06 | outside |
| 576 | 590 | VRDPQTLEILDITP | 1.4571 | 14 | ABCPRED | 1 | ALLERGEN | Non-Toxin | -0.14 | -0.16 | 0.19 | -2 | 4.03 | 1610.05 | outside |
| 97 | 113 | KSNIIRGWIFGTTLDS | -0.5366 | 16 | ABCPRED | 1 | NON-ALLERGEN | Non-Toxin | -0.07 | 0 | -0.26 | 1 | 9.1 | 1808.31 | outside |
| 734 | 748 | TSVDCTMYICGDST | -0.2033 | 14 | ABCPRED | 1 | NON-ALLERGEN | Non-Toxin | -0.05 | 0.23 | -0.25 | -2 | 3.57 | 1495.84 | outside |
| 1019 | 1033 | RASANLAATKMSEC | 0.3999 | 14 | ABCPRED | 1 | NON-ALLERGEN | Non-Toxin | -0.21 | -0.16 | 0.24 | 1 | 8.57 | 1452.84 | outside |
| 896 | 916 | IPFAMQMAYRFNGIGVTQNV | 1.2009 | 20 | ABCPRED | 1 | NON-ALLERGEN | Non-Toxin | 0.02 | 0.38 | -0.71 | 1 | 9.1 | 2257.96 | outside |
| 379 | 395 | CYGVSPTKLNDLCFTN | 2.0134 | 16 | ABCPRED,literature | 2 | NON-ALLERGEN | Non-Toxin | -0.08 | -0.02 | -0.38 | 0 | 6.12 | 1775.25 | outside |
| 793 | 813 | PIKDFGGFNFSQILPDPSKP | -0.0321 | 20 | ABCPRED | 1 | ALLERGEN | Non-Toxin | -0.08 | -0.47 | 0.01 | 0 | 6.31 | 2204.8 | outside |
| 402 | 418 | IRGDEVRQIAPGQTGK | 0.7338 | 16 | AAP | 1 | ALLERGEN | Non-Toxin | -0.3 | -0.96 | 0.59 | 1 | 9.1 | 1725.18 | outside |
| 157 | 171 | FRVYSSANNCTFEY | -0.1237 | 14 | ABCPRED | 1 | NON-ALLERGEN | Non-Toxin | -0.16 | -0.41 | -0.43 | 0 | 6.32 | 1701.02 | outside |
| 1064 | 1086 | HVTYVPAQEKNFTTAPAICHDG | 0.4614 | 22 | BCPREDS,FBCPred | 2 | NON-ALLERGEN | Non-Toxin | -0.08 | -0.35 | -0.22 | 0 | 6.01 | 2399.98 | outside |
| 612 | 626 | YQDVNCTEVPVAIH | 1.0383 | 14 | ABCPRED | 1 | NON-ALLERGEN | Non-Toxin | -0.05 | 0.04 | -0.33 | -1.5 | 4.36 | 1587.97 | outside |
| 397 | 419 | ADSFVIRGDEVRQIAPGQTGKI | 0.5327 | 22 | AAP | 1 | ALLERGEN | Non-Toxin | -0.17 | -0.29 | 0.29 | 0 | 6.47 | 2357.98 | outside |
| 963 | 983 | VKQLSSNFGAISSVLNDILS | 0.2036 | 20 | ABCPRED | 1 | NON-ALLERGEN | Non-Toxin | 0 | 0.56 | -0.34 | 0 | 6.19 | 2092.68 | outside |
| 893 | 913 | ALQIPFAMQMAYRFNGIGVT | 1.2262 | 20 | ABCPRED | 1 | NON-ALLERGEN | Non-Toxin | 0.07 | 0.62 | -0.76 | 1 | 9.1 | 2228.96 | outside |
| 154 | 170 | ESEFRVYSSANNCTFE | 0.0621 | 16 | ABCPRED,BCPREDS,FBCPred | 3 | ALLERGEN | Non-Toxin | -0.24 | -0.77 | 0.16 | -2 | 4.26 | 1883.18 | outside |
| 900 | 920 | MQMAYRFNGIGVTQNVLYEN | 0.809 | 20 | ABCPRED | 1 | NON-ALLERGEN | Non-Toxin | -0.09 | -0.23 | -0.51 | 0 | 6.35 | 2348.99 | outside |
| 1102 | 1116 | WFVTQRNFYEPQII | 0.5412 | 14 | ABCPRED | 1 | NON-ALLERGEN | Non-Toxin | -0.07 | -0.3 | -0.69 | 0 | 6.35 | 1841.31 | outside |
| 941 | 955 | TASALGKLQDVVNQ | 0.5172 | 14 | ABCPRED | 1 | NON-ALLERGEN | Non-Toxin | -0.11 | -0.01 | -0.08 | 0 | 6.19 | 1443.83 | outside |
| 305 | 325 | SFTVEKGIYQTSNFRVQPTE | 0.7705 | 20 | ABCPRED | 1 | NON-ALLERGEN | Non-Toxin | -0.2 | -0.72 | 0 | 0 | 6.49 | 2331.86 | outside |
| 841 | 861 | LGDIAARDLICAQKFNGLTV | 0.7345 | 20 | ABCPRED | 1 | ALLERGEN | Non-Toxin | -0.02 | 0.57 | -0.17 | 0 | 6.28 | 2118.78 | outside |
| 735 | 749 | SVDCTMYICGDSTE | -0.1613 | 14 | ABCPRED | 1 | NON-ALLERGEN | Non-Toxin | -0.08 | 0.03 | -0.01 | -3 | 3.5 | 1523.85 | outside |
| 896 | 910 | IPFAMQMAYRFNGI | 1.0623 | 14 | ABCPRED | 1 | NON-ALLERGEN | Non-Toxin | 0.05 | 0.51 | -0.79 | 1 | 9.1 | 1659.2 | outside |
| 350 | 370 | VYAWNRKRISNCVADYSVLY | 0.1216 | 20 | ABCPRED | 1 | NON-ALLERGEN | Non-Toxin | -0.16 | -0.14 | -0.37 | 2 | 9.16 | 2421.05 | outside |
| 886 | 900 | WTFGAGAALQIPFA | 0.6005 | 14 | ABCPRED | 1 | NON-ALLERGEN | Non-Toxin | 0.23 | 0.97 | -1.01 | 0 | 5.88 | 1449.87 | outside |

| | | | | | | | | | | | | | | | |
|---|---|---|---|---|---|---|---|---|---|---|---|---|---|---|---|
| 667 | 681 | GAGICASYQTQTNS | 0.6038 | 14 | ABCPRED | 1 | NON-ALLERGEN | Non-Toxin | -0.09 | -0.36 | -0.41 | 0 | 5.84 | 1400.68 | outside |
| 450 | 470 | NYLYRLFRKSNLKPFERDIS | 0.1611 | 20 | ABCPRED | 1 | ALLERGEN | Non-Toxin | -0.36 | -0.98 | 0.26 | 3 | 10 | 2560.24 | outside |
| 1134 | 1156 | NNTVYDPLQPELDSFKEELDKY | -0.3853 | 22 | AAP,FBCPred | 2 | NON-ALLERGEN | Non-Toxin | -0.27 | -1.28 | 0.48 | -4 | 4.02 | 2658.19 | outside |
| 139 | 161 | PFLGVYYHKNNKSWMESEFRVY | 0.5148 | 22 | BCPREDS | 1 | NON-ALLERGEN | Non-Toxin | -0.17 | -0.83 | -0.27 | 1.5 | 8.71 | 2795.49 | outside |
| 497 | 511 | FQPTNGVGYQPYRV | 0.6799 | 14 | ABCPRED | 1 | NON-ALLERGEN | Non-Toxin | -0.15 | -0.79 | -0.49 | 1 | 8.93 | 1626.01 | outside |
| 849 | 869 | LICAQKFNGLTVLPPLLTDE | 0.3207 | 20 | ABCPRED | 1 | ALLERGEN | Non-Toxin | 0.04 | 0.6 | -0.39 | -1 | 4.38 | 2185.92 | outside |
| 1153 | 1181 | DKYFKNHTSPDVDLGDISGINASVVNIQKK | 0.7715 | 30 | literature | 1 | NON-ALLERGEN | Non-Toxin | -0.19 | -0.64 | 0.26 | 0.5 | 7.09 | 3304.13 | outside |
| 331 | 345 | NITNLCPFGEVFNA | 0.6106 | 14 | ABCPRED | 1 | NON-ALLERGEN | Non-Toxin | 0.05 | 0.41 | -0.6 | -1 | 4 | 1538.94 | outside |
| 1153 | 1169 | DKYFKNHTSPDVDLGD | 0.614 | 16 | ABCPRED,literature | 2 | ALLERGEN | Non-Toxin | -0.3 | -1.41 | 0.59 | -1.5 | 4.56 | 1851.18 | outside |
| 861 | 881 | LPPLLTDEMIAQYTSALLAG | 0.3373 | 20 | ABCPRED | 1 | NON-ALLERGEN | Non-Toxin | 0.09 | 0.66 | -0.51 | -2 | 3.67 | 2117.79 | outside |
| 57 | 77 | PFFSNVTWFHAIHVSGTNGT | 0.6969 | 20 | ABCPRED | 1 | NON-ALLERGEN | Non-Toxin | 0.07 | 0.13 | -0.87 | 1 | 7.26 | 2219.74 | outside |
| 555 | 575 | SNKKFLPFQQFGRDIADTTD | 0.4281 | 20 | ABCPRED | 1 | NON-ALLERGEN | Non-Toxin | -0.27 | -0.95 | 0.33 | 0 | 6.39 | 2328.85 | outside |
| 223 | 237 | LEPLVDLPIGINIT | 1.1627 | 14 | ABCPRED | 1 | NON-ALLERGEN | Non-Toxin | 0.16 | 1.02 | -0.46 | -2 | 3.67 | 1507.03 | outside |
| 70 | 92 | VSGTNGTKRFDNPVLPFNDGVY | 0.4291 | 22 | AAP | 1 | ALLERGEN | Non-Toxin | -0.14 | -0.54 | -0.07 | 0 | 6.31 | 2397.95 | outside |
| 119 | 135 | IVNNATNVVIKVCEFQ | -0.0057 | 16 | FBCPred | 1 | ALLERGEN | Non-Toxin | 0 | 0.68 | -0.45 | 0 | 6.32 | 1791.34 | outside |
| 326 | 340 | IVRFPNITNLCPFG | 0.8566 | 14 | ABCPRED | 1 | NON-ALLERGEN | Non-Toxin | 0.04 | 0.66 | -0.71 | 1 | 8.6 | 1591.11 | outside |
| 115 | 135 | QSLLIVNNATNVVIKVCEFQ | 0.1923 | 20 | ABCPRED | 1 | NON-ALLERGEN | Non-Toxin | 0.01 | 0.71 | -0.52 | 0 | 6.32 | 2232.94 | outside |
| 1093 | 1113 | GVFVSNGTHWFVTQRNFYEP | 0.2915 | 20 | ABCPRED | 1 | NON-ALLERGEN | Non-Toxin | -0.06 | -0.38 | -0.61 | 0.5 | 7.1 | 2385.92 | outside |
| 913 | 927 | QNVLYENQKLIANQ | 0.0761 | 14 | ABCPRED | 1 | NON-ALLERGEN | Non-Toxin | -0.22 | -0.83 | -0.18 | 0 | 6.35 | 1675.1 | outside |
| 1048 | 1068 | HLMSFPQSAPHGVVFLHVTY | 0.4454 | 20 | ABCPRED | 1 | NON-ALLERGEN | Non-Toxin | 0.09 | 0.46 | -0.92 | 1.5 | 7.35 | 2267.94 | outside |
| 745 | 765 | DSTECSNLLLQYGSFCTQLN | 0.7636 | 20 | ABCPRED | 1 | NON-ALLERGEN | Toxin | -0.11 | -0.18 | -0.35 | -2 | 3.67 | 2236.75 | outside |
| 881 | 897 | TITSGWTFGAGAALQI | 0.3908 | 16 | ABCPRED | 1 | NON-ALLERGEN | Non-Toxin | 0.17 | 0.78 | -0.84 | 0 | 5.88 | 1594.04 | outside |
| 1026 | 1040 | ATKMSECVLGQSKR | 0.5357 | 14 | ABCPRED | 1 | NON-ALLERGEN | Non-Toxin | -0.3 | -0.56 | 0.45 | 2 | 9.36 | 1538.01 | outside |
| 720 | 742 | ISVTTEILPVSMTKTSVDCTMY | 1.2581 | 22 | FBCPred | 1 | NON-ALLERGEN | Non-Toxin | 0 | 0.55 | -0.36 | -1 | 4.38 | 2420.17 | outside |
| 1199 | 1221 | DLQELGKYEQYIKWPWYIWLGF | 0.9678 | 22 | FBCPred | 1 | ALLERGEN | Non-Toxin | -0.02 | -0.5 | -0.6 | -1 | 4.68 | 2876.66 | outside_transmembrane-region |
| 678 | 692 | TNSPRRARSVASQS | 0.1779 | 14 | ABCPRED | 1 | NON-ALLERGEN | Non-Toxin | -0.49 | -1.3 | 0.55 | 3 | 12.31 | 1516.81 | outside |
| 1159 | 1181 | HTSPDVDLGDISGINASVVNIQ | 1.0856 | 22 | FBCPred | 1 | NON-ALLERGEN | Non-Toxin | -0.03 | 0.09 | -0.12 | -2.5 | 3.93 | 2251.77 | outside |
| 101 | 117 | IRGWIFGTTLDSKTQS | 0.1657 | 16 | ABCPRED | 1 | NON-ALLERGEN | Non-Toxin | -0.13 | -0.33 | -0.17 | 1 | 9.1 | 1810.28 | outside |
| 170 | 184 | YVSQPFLMDLEGKQ | 0.5381 | 14 | ABCPRED | 1 | NON-ALLERGEN | Non-Toxin | -0.11 | -0.39 | -0.11 | -1 | 4.38 | 1655.11 | outside |
| 80 | 100 | DNPVLPFNDGVYFASTEKSN | 0.4672 | 20 | ABCPRED | 1 | NON-ALLERGEN | Non-Toxin | -0.13 | -0.63 | 0.01 | -2 | 4.03 | 2214.65 | outside |
| 179 | 193 | LEGKQGNFKNLREF | 0.6975 | 14 | ABCPRED | 1 | NON-ALLERGEN | Non-Toxin | -0.33 | -1.24 | 0.5 | 1 | 8.93 | 1680.11 | outside |
| 105 | 125 | IFGTTLDSKTQSLLIVNNAT | 0.5121 | 20 | ABCPRED | 1 | ALLERGEN | Non-Toxin | -0.02 | 0.32 | -0.4 | 0 | 6.19 | 2136.74 | outside |
| 507 | 521 | PYRVVVLSFELLHA | 1.0185 | 14 | ABCPRED | 1 | NON-ALLERGEN | Non-Toxin | 0.07 | 0.98 | -0.67 | 0.5 | 7.1 | 1643.17 | outside |
| 872 | 892 | QYTSALLAGTITSGWTFGAG | 0.3912 | 20 | ABCPRED | 1 | NON-ALLERGEN | Non-Toxin | 0.11 | 0.43 | -0.8 | 0 | 5.88 | 2002.51 | outside |
| 532 | 552 | NLVKNKCVNFNFNGLTGTGV | 1.084 | 20 | ABCPRED | 1 | NON-ALLERGEN | Non-Toxin | -0.07 | 0.02 | -0.4 | 2 | 9.36 | 2139.77 | outside |
| 540 | 554 | NFNFNGLTGTGVLT | 1.1216 | 14 | ABCPRED | 1 | ALLERGEN | Non-Toxin | 0.06 | 0.26 | -0.76 | 0 | 5.88 | 1454.82 | outside |
| 756 | 770 | YGSFCTQLNRALTG | 0.5511 | 14 | ABCPRED | 1 | NON-ALLERGEN | Non-Toxin | -0.1 | -0.08 | -0.5 | 1 | 8.57 | 1530.92 | outside |
| 245 | 261 | HRSYLTPGDSSSGWTA | 0.6017 | 16 | ABCPRED | 1 | NON-ALLERGEN | Non-Toxin | -0.18 | -0.93 | -0.13 | 0.5 | 7.09 | 1722.02 | outside |
| 769 | 783 | GIAVEQDKNTQEVF | 0.5109 | 14 | ABCPRED | 1 | NON-ALLERGEN | Toxin | -0.17 | -0.61 | 0.31 | -2 | 4.14 | 1577.92 | outside |
| 242 | 262 | LALHRSYLTPGDSSSGWTAG | 0.4411 | 20 | ABCPRED | 1 | NON-ALLERGEN | Non-Toxin | -0.07 | -0.29 | -0.31 | 0.5 | 7.09 | 2076.54 | outside |
| 984 | 998 | LDKVEAEVQIDRLI | -0.2016 | 14 | ABCPRED | 1 | NON-ALLERGEN | Non-Toxin | -0.17 | 0.06 | 0.54 | -2 | 4.32 | 1641.11 | outside |
| 244 | 258 | LHRSYLTPGDSSSG | 0.4766 | 14 | ABCPRED | 1 | NON-ALLERGEN | Toxin | -0.2 | -0.8 | 0.03 | 0.5 | 7.09 | 1476.76 | outside |
| 502 | 516 | GVGYQPYRVVVLSF | 1.2391 | 14 | ABCPRED | 1 | NON-ALLERGEN | Non-Toxin | 0.06 | 0.69 | -0.81 | 1 | 8.93 | 1584.06 | outside |
| 1082 | 1096 | CHDGKAHFPREGVF | -0.2683 | 14 | ABCPRED | 1 | NON-ALLERGEN | Non-Toxin | -0.19 | -0.72 | 0.21 | 1 | 7.22 | 1599.98 | outside |
| 853 | 867 | QKFNGLTVLPPLLT | 0.3372 | 14 | ABCPRED | 1 | NON-ALLERGEN | Non-Toxin | 0.04 | 0.45 | -0.61 | 1 | 9.11 | 1541.09 | outside |
| 341 | 363 | VFNATRFASVYAWNRKRISNCV | 0.2643 | 22 | FBCPred | 1 | ALLERGEN | Non-Toxin | -0.19 | -0.08 | -0.3 | 4 | 10.92 | 2603.3 | outside |
| 761 | 775 | TQLNRALTGIAVEQ | 0.4849 | 14 | ABCPRED | 1 | NON-ALLERGEN | Non-Toxin | -0.13 | -0.03 | -0.15 | 0 | 6.36 | 1513.93 | outside |
| 805 | 833 | ILPDPSKPSKRSFIEDLLFNKVTLADAGFK | 0.4162 | 30 | literature | 1 | NON-ALLERGEN | Non-Toxin | -0.14 | -0.18 | 0.23 | 1 | 8.76 | 3348.34 | outside |
| 180 | 200 | EGKQGNFKNLREFVFKNIDG | 0.2498 | 20 | ABCPRED | 1 | NON-ALLERGEN | Non-Toxin | -0.28 | -1.05 | 0.46 | 1 | 8.83 | 2340.93 | outside |
| 56 | 196 | /IFGTTLDSKTQSLLIVNNATNVVIKVCEFQFCND | 0.1835 | 103 | ellipro,literature | 2 | NON-ALLERGEN | too long | | | | | | | outside |
| 47 | 61 | VLHSTQDLFLPFFS | 0.2442 | 14 | ABCPRED | 1 | NON-ALLERGEN | Non-Toxin | 0.1 | 0.71 | -0.82 | -0.5 | 5.09 | 1651.1 | outside |
| 63 | 77 | TWFHAIHVSGTNGT | 0.533 | 14 | ABCPRED | 1 | NON-ALLERGEN | Non-Toxin | 0.04 | -0.09 | -0.81 | 1 | 7.26 | 1527.87 | outside |
| 991 | 1005 | VQIDRLITGRLQSL | -0.2271 | 14 | ABCPRED | 1 | NON-ALLERGEN | Non-Toxin | -0.16 | 0.23 | -0.09 | 1 | 9.95 | 1612.13 | outside |
| 520 | 540 | APATVCGPKKSTNLVKNKCV | 0.2034 | 20 | ABCPRED | 1 | NON-ALLERGEN | Non-Toxin | -0.18 | -0.17 | 0.13 | 4 | 9.72 | 2058.77 | outside |
| 954 | 968 | QNAQALNTLVKQLS | 0.0161 | 14 | ABCPRED | 1 | NON-ALLERGEN | Non-Toxin | -0.16 | -0.26 | -0.29 | 1 | 9.11 | 1527.96 | outside |
| 938 | 952 | LSSTASALGKLQDV | 0.8753 | 14 | ABCPRED | 1 | NON-ALLERGEN | Non-Toxin | -0.05 | 0.34 | -0.09 | 0 | 6.19 | 1389.77 | outside |
| 837 | 851 | YGDCLGDIAARDLI | 0.2458 | 14 | ABCPRED | 1 | NON-ALLERGEN | Non-Toxin | -0.04 | 0.4 | 0.04 | -2 | 3.94 | 1494.88 | outside |
| 429 | 451 | FTGCVIAWNSNNLDSKVGGNYN | 0.4339 | 22 | AAP | 1 | NON-ALLERGEN | Non-Toxin | -0.07 | -0.31 | -0.41 | 0 | 6.15 | 2373.91 | outside |
| 509 | 529 | RVVVLSFELLHAPATVCGPK | 0.3924 | 20 | ABCPRED | 1 | NON-ALLERGEN | Non-Toxin | 0.03 | 0.85 | -0.38 | 1.5 | 8.57 | 2136.88 | outside |
| 635 | 655 | VYSTGSNVFQTRAGCLIGAE | 0.7107 | 20 | ABCPRED | 1 | ALLERGEN | Non-Toxin | -0.03 | 0.26 | -0.36 | 0 | 6.32 | 2073.6 | outside |
| 491 | 505 | PLQSYGFQPTNGVG | 0.3687 | 14 | ABCPRED | 1 | ALLERGEN | Non-Toxin | -0.03 | -0.49 | -0.54 | 0 | 5.88 | 1464.81 | outside |
| 1065 | 1079 | VTYVPAQEKNFTTA | 0.7333 | 14 | ABCPRED | 1 | NON-ALLERGEN | Non-Toxin | -0.1 | -0.33 | -0.26 | 0 | 6.35 | 1568.94 | outside |
| 728 | 742 | PVSMTKTSVDCTMY | 1.3667 | 14 | ABCPRED | 1 | NON-ALLERGEN | Non-Toxin | -0.09 | 0.05 | -0.25 | 0 | 6.15 | 1563.02 | outside |
| 356 | 370 | KRISNCVADYSVLY | -0.0485 | 14 | ABCPRED | 1 | NON-ALLERGEN | Non-Toxin | -0.15 | 0.1 | -0.21 | 1 | 8.51 | 1631.07 | outside |
| 473 | 489 | YQAGSTPCNGVEGFNC | 0.3341 | 16 | ABCPRED,AAP | 2 | ALLERGEN | Non-Toxin | -0.07 | -0.36 | -0.33 | -1 | 4 | 1646.99 | outside |
| 878 | 892 | LAGTITSGWTFGAG | 0.2906 | 14 | ABCPRED | 1 | NON-ALLERGEN | Non-Toxin | 0.18 | 0.66 | -0.81 | 0 | 5.88 | 1338.69 | outside |
| 506 | 520 | QPYRVVVLSFELLH | 0.9675 | 14 | ABCPRED | 1 | NON-ALLERGEN | Non-Toxin | 0 | 0.6 | -0.62 | 0.5 | 7.1 | 1700.23 | outside |
| 584 | 598 | ILDITPCSFGGVSV | 1.6035 | 14 | ABCPRED | 1 | NON-ALLERGEN | Non-Toxin | 0.18 | 1.31 | -0.62 | -1 | 3.8 | 1407.85 | outside |
| 488 | 508 | CYFPLQSYGFQPTNGVGYQP | 0.681 | 20 | ABCPRED | 1 | ALLERGEN | Non-Toxin | -0.02 | -0.47 | -0.78 | 0 | 5.84 | 2266.81 | outside |
| 271 | 287 | QPRTFLLKYNENGTIT | 0.2688 | 16 | FBCPred | 1 | NON-ALLERGEN | Non-Toxin | -0.22 | -0.81 | -0.11 | 1 | 8.93 | 1895.39 | outside |
| 1259 | 1274 | DDSEPVLKGVKLHYT | 1.1849 | 15 | COVIDep | 1 | NON-ALLERGEN | Non-Toxin | -0.19 | -0.69 | 0.37 | -0.5 | 5.39 | 1701.13 | inside |
| 266 | 282 | YVGYLQPRTFLLKYNE | 0.5108 | 16 | ABCPRED | 1 | NON-ALLERGEN | Non-Toxin | -0.13 | -0.44 | -0.46 | 1 | 8.76 | 2004.56 | outside |
| 12 | 32 | SSQCVNLTTRTQLPPAYTNS | 1.1345 | 20 | ABCPRED | 1 | NON-ALLERGEN | Non-Toxin | -0.21 | -0.61 | -0.29 | 1 | 8.57 | 2181.69 | outside |
| 48 | 62 | LHSTQDLFLPFFSN | 0.1556 | 14 | ABCPRED | 1 | NON-ALLERGEN | Non-Toxin | 0.01 | 0.16 | -0.7 | -0.5 | 5.09 | 1666.07 | outside |
| 786 | 830 | KTPPIKDFGGFNFSQILPDPSKPSKRSFIEDLLFNK | 0.1733 | 44 | literature | 1 | NON-ALLERGEN | Non-Toxin | -0.14 | -0.4 | 0.07 | 3 | 9.62 | 5027.52 | outside |
| 468 | 482 | ISTEIYQAGSTPCN | 0.1359 | 14 | ABCPRED | 1 | NON-ALLERGEN | Non-Toxin | -0.07 | -0.25 | -0.3 | -1 | 4 | 1483.81 | outside |
| 927 | 947 | FNSAIGKIQDSLSSTASALG | 0.4804 | 20 | ABCPRED | 1 | NON-ALLERGEN | Non-Toxin | -0.02 | 0.24 | -0.18 | 0 | 6.19 | 1967.45 | outside |

| | | | | | | | | | | | | | | | |
|---|---|---|---|---|---|---|---|---|---|---|---|---|---|---|---|
| 562 | 582 | FQQFGRDIADTTDAVRDPQT | 0.071 | 20 | ABCPRED | 1 | NON-ALLERGEN | Non-Toxin | -0.3 | -0.98 | 0.4 | -2 | 4.14 | 2281.7 | outside |
| 1248 | 1264 | CSCGSCCKFDEDDSEP | 0.1542 | 16 | ABCPRED,FBCPred | 2 | NON-ALLERGEN | Non-Toxin | -0.28 | -0.81 | 0.78 | -4 | 3.77 | 1725.01 | inside |
| 24 | 38 | LPPAYTNSFTRGVY | 0.1299 | 14 | ABCPRED | 1 | NON-ALLERGEN | Non-Toxin | -0.07 | -0.27 | -0.59 | 1 | 8.93 | 1585.97 | outside |
| 602 | 616 | TNTSNQVAVLYQDV | 0.3582 | 14 | ABCPRED | 1 | NON-ALLERGEN | Non-Toxin | -0.11 | -0.2 | -0.41 | -1 | 3.8 | 1551.88 | outside |
| 1069 | 1083 | PAQEKNFTTAPAIC | 0.5959 | 14 | ABCPRED | 1 | ALLERGEN | Non-Toxin | -0.1 | -0.27 | -0.09 | 0 | 6.32 | 1490.88 | outside |
| 1017 | 1037 | EIRASANLAATKMSECVLGQ | 0.5216 | 20 | ABCPRED | 1 | ALLERGEN | Non-Toxin | -0.12 | 0.14 | 0.07 | 0 | 6.46 | 2092.7 | outside |
| 492 | 514 | LQSYGFQPTNGVGYQPYRVVVL | 0.6433 | 22 | FBCPred | 1 | NON-ALLERGEN | Non-Toxin | -0.03 | -0.05 | -0.7 | 1 | 8.83 | 2486.17 | outside |
| 922 | 936 | LIANQFNSAIGKIQ | 0.1674 | 14 | ABCPRED | 1 | NON-ALLERGEN | Non-Toxin | 0 | 0.33 | -0.47 | 1 | 9.11 | 1516.98 | outside |
| 75 | 99 | GTKRFDNPVLPFNDGVYFASTEKSNK | 0.4286 | 26 | literature | 1 | ALLERGEN | Non-Toxin | -0.23 | -0.89 | 0.24 | 1 | 8.76 | 2932.61 | outside |
| 694 | 708 | AYTMSLGAENSVAY | 0.5218 | 14 | ABCPRED | 1 | ALLERGEN | Non-Toxin | 0.02 | 0.21 | -0.52 | -1 | 4 | 1476.81 | outside |
| 690 | 704 | QSIIAYTMSLGAEN | 0.629 | 14 | ABCPRED | 1 | NON-ALLERGEN | Non-Toxin | 0.02 | 0.27 | -0.46 | -1 | 4 | 1497.89 | outside |
| 586 | 606 | DITPCSFGGVSVITPGTNTS | 0.9823 | 20 | ABCPRED | 1 | NON-ALLERGEN | Non-Toxin | 0.03 | 0.31 | -0.38 | -1 | 3.8 | 1953.44 | outside |
| 791 | 811 | TPPIKDFGGFNFSQILPDPS | 0.2481 | 20 | ABCPRED | 1 | NON-ALLERGEN | Non-Toxin | -0.04 | -0.31 | -0.16 | -1 | 4.21 | 2177.73 | outside |
| 431 | 445 | GCVIAWNSNNLDSK | 0.6118 | 14 | ABCPRED | 1 | NON-ALLERGEN | Non-Toxin | -0.12 | -0.29 | -0.2 | 0 | 6.16 | 1520.88 | outside |
| 955 | 975 | NAQALNTLVKQLSSNFGAIS | 0.3505 | 20 | ABCPRED | 1 | ALLERGEN | Non-Toxin | -0.05 | 0.17 | -0.41 | 1 | 9.11 | 2076.64 | outside |
| 1097 | 1113 | SNGTHWFVTQRNFYEP | 0.1457 | 16 | ABCPRED,FBCPred | 2 | NON-ALLERGEN | Non-Toxin | -0.2 | -1.14 | -0.41 | 0.5 | 7.1 | 1983.36 | outside |
| 581 | 595 | TLEILDITPCSFGG | 1.5811 | 14 | ABCPRED | 1 | NON-ALLERGEN | Non-Toxin | 0.1 | 0.74 | -0.37 | -2 | 3.67 | 1465.89 | outside |
| 430 | 450 | TGCVIAWNSNNLDSKVGGNY | 0.5742 | 20 | ABCPRED | 1 | NON-ALLERGEN | Non-Toxin | -0.08 | -0.3 | -0.34 | 0 | 6.15 | 2112.6 | outside |
| 818 | 840 | IEDLLFNKVTLADAGFIKQYGD | 0.3688 | 22 | AAP | 1 | NON-ALLERGEN | Non-Toxin | -0.04 | 0.1 | -0.04 | -2 | 4.23 | 2471.14 | outside |
| 770 | 790 | IAVEQDKNTQEVFAQVKQIY | 0.3611 | 20 | ABCPRED | 1 | NON-ALLERGEN | Non-Toxin | -0.18 | -0.49 | 0.09 | -1 | 4.68 | 2351.95 | outside |
| 784 | 798 | QVKQIYKTPPIKDF | 0.1794 | 14 | ABCPRED | 1 | NON-ALLERGEN | Non-Toxin | -0.22 | -0.81 | 0.15 | 2 | 9.55 | 1705.24 | outside |
| 1008 | 1024 | VTQQLIRAAEIRASAN | 0.0867 | 16 | AAP | 1 | ALLERGEN | Non-Toxin | -0.19 | -0.02 | 0.04 | 1 | 9.95 | 1741.21 | outside |
| 359 | 379 | SNCVADYSVLYNSASFSTFK | -0.0103 | 20 | ABCPRED | 1 | ALLERGEN | Non-Toxin | -0.06 | 0.11 | -0.45 | 0 | 6.15 | 2203.67 | outside |
| 322 | 376 | TNLCPFGEVFNATRFASVYAWNRKRISNCVADY | 0.2542 | 54 | ellipro,literature | 2 | ALLERGEN | too long | | | | | | | outside |
| 915 | 929 | VLYENQKLIANQFN | 0.0805 | 14 | ABCPRED | 1 | NON-ALLERGEN | Non-Toxin | -0.13 | -0.38 | -0.37 | 0 | 6.35 | 1694.14 | outside |
| 601 | 641 | 'SNQVAVLYQDVNCTEVPVAIHADQLTPTWRVY | 0.6197 | 40 | literature | 1 | NON-ALLERGEN | Non-Toxin | -0.09 | -0.2 | -0.34 | -1.5 | 4.54 | 4337.36 | outside |
| 241 | 261 | LLALHRSYLTPGDSSSGWTA | 0.5843 | 20 | ABCPRED | 1 | NON-ALLERGEN | Non-Toxin | -0.05 | -0.08 | -0.4 | 0.5 | 7.09 | 2132.65 | outside |
| 579 | 593 | PQTLEILDITPCSF | 1.3489 | 14 | ABCPRED | 1 | NON-ALLERGEN | Non-Toxin | 0.03 | 0.43 | -0.36 | -2 | 3.67 | 1577.03 | outside |
| 10 | 24 | LVSSQCVNLTTRTQ | 1.2893 | 14 | ABCPRED | 1 | NON-ALLERGEN | Non-Toxin | -0.19 | -0.01 | -0.33 | 1 | 8.6 | 1549.97 | outside |
| 892 | 906 | AALQIPFAMQMAYR | 1.0216 | 14 | ABCPRED | 1 | NON-ALLERGEN | Non-Toxin | 0.01 | 0.55 | -0.69 | 1 | 9.1 | 1611.15 | outside |
| 961 | 981 | TLVKQLSSNFGAISSVLNDI | 0.2086 | 20 | ABCPRED | 1 | NON-ALLERGEN | Non-Toxin | 0.01 | 0.56 | -0.38 | 0 | 6.19 | 2106.71 | outside |
| 327 | 368 | NITNLCPFGEVFNATRFASVYAWNRKRISNCVA | 0.4113 | 41 | ellipro | 1 | NON-ALLERGEN | Non-Toxin | -0.13 | 0 | -0.31 | 3 | 9.41 | 4727.98 | outside |
| 664 | 686 | IPIGAGICASYQTQTNSPRRAR | 0.6077 | 22 | AAP,BCPREDS | 2 | ALLERGEN | Non-Toxin | -0.23 | -0.5 | -0.04 | 3 | 10.77 | 2360.99 | outside |
| 453 | 473 | YRLFRKSNLKPFERDISTEI | 0.0376 | 20 | ABCPRED | 1 | ALLERGEN | Non-Toxin | -0.35 | -0.92 | 0.49 | 2 | 9.71 | 2513.18 | outside |
| 706 | 726 | AYSNNSIAIPTNFTISVTTE | 0.7773 | 20 | ABCPRED | 1 | ALLERGEN | Non-Toxin | -0.01 | 0.1 | -0.49 | -1 | 4 | 2143.62 | outside |
| 456 | 472 | FRKSNLKPFERDISTE | 0.3795 | 16 | ABCPRED,AAP | 2 | ALLERGEN | Non-Toxin | -0.41 | -1.3 | 0.8 | 1 | 8.93 | 1967.43 | outside |
| 214 | 234 | RDLPQGFSALEPLVDLPIGI | 0.8015 | 20 | ABCPRED | 1 | NON-ALLERGEN | Non-Toxin | 0.02 | 0.4 | -0.14 | -2 | 4.03 | 2150.81 | outside |
| 647 | 661 | AGCLIGAEHVNNSY | 0.829 | 14 | ABCPRED | 1 | NON-ALLERGEN | Non-Toxin | 0.01 | 0.14 | -0.44 | -0.5 | 5.25 | 1447.79 | outside |
| 14 | 34 | QCVNLTTRTQLPPAYTNSFT | 0.9669 | 20 | ABCPRED | 1 | ALLERGEN | Non-Toxin | -0.16 | -0.42 | -0.47 | 1 | 8.57 | 2255.82 | outside |
| 51 | 65 | TQDLFLPFFSNVTW | 0.5668 | 14 | ABCPRED | 1 | NON-ALLERGEN | Non-Toxin | 0.08 | 0.36 | -0.94 | -1 | 3.8 | 1715.14 | outside |
| 335 | 355 | LCPFGEVFNATRFASVYAWN | 0.1673 | 20 | ABCPRED | 1 | NON-ALLERGEN | Non-Toxin | 0.03 | 0.39 | -0.71 | 0 | 6.32 | 2292.87 | outside |
| 414 | 434 | QTGKIADYNYKLPDDFTGCV | 0.6255 | 20 | ABCPRED | 1 | NON-ALLERGEN | Non-Toxin | -0.15 | -0.6 | 0.05 | -1 | 4.43 | 2248.78 | outside |
| 21 | 35 | RTQLPPAYTNSFTR | 0.2033 | 14 | ABCPRED | 1 | NON-ALLERGEN | Non-Toxin | -0.31 | -1.07 | -0.11 | 2 | 10.84 | 1652.03 | outside |
| 445 | 478 | VGGNYNYLYRLFRKSNLKPFERDISTEIYQAGS | 0.1172 | 33 | literature | 1 | NON-ALLERGEN | Non-Toxin | -0.23 | -0.75 | 0 | 2 | 9.34 | 3900.86 | outside |
| 637 | 651 | STGSNVFQTRAGCL | 0.9742 | 14 | ABCPRED | 1 | NON-ALLERGEN | Non-Toxin | -0.12 | -0.01 | -0.29 | 1 | 8.6 | 1440.79 | outside |
| 152 | 166 | WMESEFRVYSSANN | 0.2181 | 14 | ABCPRED | 1 | NON-ALLERGEN | Non-Toxin | -0.22 | -0.89 | -0.09 | -1 | 4.54 | 1720.03 | outside |
| 946 | 960 | GKLQDVVNQNAQAL | 0.4958 | 14 | ABCPRED | 1 | ALLERGEN | Non-Toxin | -0.17 | -0.41 | -0.04 | 0 | 6.19 | 1497.89 | outside |
| 1080 | 1094 | AICHDGKAHFPREG | -0.3671 | 14 | ABCPRED | 1 | NON-ALLERGEN | Non-Toxin | -0.21 | -0.77 | 0.34 | 1 | 7.22 | 1537.91 | outside |
| 1031 | 1045 | ECVLGQSKRVDFCG | 0.9433 | 14 | ABCPRED | 1 | NON-ALLERGEN | Non-Toxin | -0.18 | -0.04 | 0.23 | 0 | 6.36 | 1540.97 | outside |
| 1214 | 1234 | WYIWLGFIAGLIAIVMVTIM | 0.6547 | 20 | ABCPRED | 1 | NON-ALLERGEN | Non-Toxin | 0.42 | 2.2 | -1.56 | 0 | 5.88 | 2311.26 | transmembrane-region |
| 648 | 668 | GCLIGAEHVNNSYECDIPIG | 0.9157 | 20 | ABCPRED | 1 | NON-ALLERGEN | Non-Toxin | 0.01 | 0.14 | -0.21 | -2.5 | 4.14 | 2104.64 | outside |
| 787 | 807 | QIYKTPPIKDFGGFNFSQIL | 0.0666 | 20 | ABCPRED | 1 | NON-ALLERGEN | Non-Toxin | -0.03 | -0.14 | -0.37 | 1 | 8.83 | 2313.99 | outside |
| 1116 | 1136 | TTDNTFVSGNCDVVIGIVNN | 0.4741 | 20 | ABCPRED | 1 | NON-ALLERGEN | Non-Toxin | -0.01 | 0.32 | -0.36 | -2 | 3.57 | 2082.57 | outside |
| 1200 | 1216 | LQELGKYEQYIKWPWY | 0.7729 | 16 | ABCPRED,AAP | 2 | NON-ALLERGEN | Non-Toxin | -0.13 | -1.09 | -0.42 | 0 | 6.48 | 2144.71 | outside_transmembrane-region |
| 950 | 970 | DVVNQNAQALNTLVKQLSSN | -0.0214 | 20 | ABCPRED | 1 | NON-ALLERGEN | Non-Toxin | -0.17 | -0.33 | -0.16 | 0 | 6.19 | 2156.69 | outside |
| 676 | 692 | TQTNSPRRARSVASQS | 0.1585 | 16 | ABCPRED,FBCPred | 2 | NON-ALLERGEN | Non-Toxin | -0.48 | -1.4 | 0.47 | 3 | 12.31 | 1746.08 | outside |
| 546 | 560 | LTGTGVLTESNKKF | 0.7508 | 14 | ABCPRED | 1 | NON-ALLERGEN | Non-Toxin | -0.12 | -0.28 | 0.05 | 1 | 8.94 | 1494.92 | outside |
| 775 | 791 | DKNTQEVFAQVKQIYK | 0.119 | 16 | FBCPred | 1 | NON-ALLERGEN | Non-Toxin | -0.3 | -1.07 | 0.33 | 1 | 8.76 | 1939.44 | outside |
| 402 | 416 | IRGDEVRQIAPGQT | 0.2932 | 14 | ABCPRED | 1 | NON-ALLERGEN | Non-Toxin | -0.28 | -0.79 | 0.46 | 0 | 6.42 | 1539.92 | outside |
| 665 | 679 | PIGAGICASYQTQT | 0.8996 | 14 | ABCPRED | 1 | NON-ALLERGEN | Non-Toxin | 0.02 | 0.16 | -0.57 | 0 | 5.84 | 1409.78 | outside |
| 180 | 196 | EGKQGNFKNLREFVFK | 0.3563 | 16 | AAP | 1 | NON-ALLERGEN | Non-Toxin | -0.32 | -1.13 | 0.49 | 2 | 9.72 | 1941.46 | outside |
| 275 | 291 | FLLKYNENGTITDAVD | 0.5419 | 16 | AAP | 1 | ALLERGEN | Non-Toxin | -0.09 | -0.23 | -0.04 | -2 | 4.03 | 1813.23 | outside |
| 1103 | 1119 | FVTQRNFYEPQIITTD | 0.5271 | 16 | ABCPRED,AAP | 2 | NON-ALLERGEN | Non-Toxin | -0.16 | -0.51 | -0.25 | -1 | 4.38 | 1972.42 | outside |
| 1103 | 1117 | FVTQRNFYEPQIIT | 0.4614 | 14 | ABCPRED | 1 | NON-ALLERGEN | Non-Toxin | -0.11 | -0.29 | -0.47 | 0 | 6.35 | 1756.2 | outside |
| 828 | 848 | LADAGFIKQYGDCLGDIAAR | 0.3902 | 20 | ABCPRED | 1 | NON-ALLERGEN | Toxin | -0.05 | 0.21 | 0.01 | -1 | 4.43 | 2097.66 | outside |
| 471 | 491 | EIYQAGSTPCNGVEGFNCYF | 0.1777 | 20 | ABCPRED | 1 | ALLERGEN | Non-Toxin | -0.02 | -0.17 | -0.44 | -2 | 3.8 | 2199.68 | outside |
| 612 | 632 | YQDVNCTEVPVAIHADQLTP | 0.7026 | 20 | ABCPRED | 1 | NON-ALLERGEN | Non-Toxin | -0.08 | -0.16 | -0.2 | -2.5 | 4.03 | 2213.74 | outside |
| 205 | 225 | SKHTPINLVRDLPQGFSALE | 0.6781 | 20 | ABCPRED | 1 | NON-ALLERGEN | Non-Toxin | -0.14 | -0.34 | 0.02 | 0.5 | 7.1 | 2222.83 | outside |
| 859 | 875 | TVLPPLLTDEMIAQYT | 0.2683 | 16 | FBCPred | 1 | NON-ALLERGEN | Non-Toxin | 0.04 | 0.42 | -0.49 | -2 | 3.67 | 1805.36 | outside |
| 1213 | 1233 | PWYIWLGFIAGLIAIVMVTI | 0.7642 | 20 | ABCPRED | 1 | NON-ALLERGEN | Non-Toxin | 0.4 | 2.03 | -1.5 | 0 | 5.88 | 2277.18 | transmembrane-region |
| 730 | 744 | SMTKTSVDCTMYIC | 0.5397 | 14 | ABCPRED | 1 | ALLERGEN | Non-Toxin | -0.07 | 0.36 | -0.34 | 0 | 6.12 | 1583.07 | outside |
| 446 | 466 | GGNYNYLYRLFRKSNLKPFE | 0.2192 | 20 | ABCPRED | 1 | NON-ALLERGEN | Non-Toxin | -0.27 | -1.04 | -0.07 | 3 | 9.83 | 2480.12 | outside |
| 237 | 251 | RFQTLLALHRSYLT | 0.3481 | 14 | ABCPRED | 1 | NON-ALLERGEN | Non-Toxin | -0.16 | 0.04 | -0.52 | 2.5 | 10.84 | 1719.23 | outside |
| 894 | 914 | LQIPFAMQMAYRFNGIGVTQ | 1.2425 | 20 | ABCPRED | 1 | NON-ALLERGEN | Non-Toxin | 0.02 | 0.36 | -0.72 | 1 | 9.1 | 2286.02 | outside |
| 185 | 205 | NFKNLREFVFKNIDGYFKIY | 0.0338 | 20 | ABCPRED | 1 | NON-ALLERGEN | Non-Toxin | -0.16 | -0.43 | -0.15 | 2 | 9.43 | 2556.26 | outside |
| 813 | 833 | SKRSFIEDLLFNKVTLADAG | 0.3463 | 20 | ABCPRED | 1 | ALLERGEN | Non-Toxin | -0.13 | 0.02 | 0.19 | 0 | 6.47 | 2224.83 | outside |
| 312 | 326 | IYQTSNFRVQPTES | 0.8629 | 14 | ABCPRED,literature | 2 | NON-ALLERGEN | Non-Toxin | -0.25 | -0.92 | -0.12 | 0 | 6.35 | 1670.01 | outside |

| | | | | | | | | | | | | | | | |
|---|---|---|---|---|---|---|---|---|---|---|---|---|---|---|---|
| 689 | 703 | SQSIIAYTMSLGAE | 0.5614 | 14 | ABCPRED | 1 | NON-ALLERGEN | Non-Toxin | 0.05 | 0.46 | -0.45 | -1 | 4 | 1470.86 | outside |
| 388 | 410 | NDLCFTNVYADSFVIRGDEVRQ | 0.4304 | 22 | FBCPred | 1 | ALLERGEN | Non-Toxin | -0.19 | -0.27 | 0.07 | -2 | 4.23 | 2562.12 | outside |
| 987 | 1001 | VEAEVQIDRLITGR | -0.3089 | 14 | ABCPRED | 1 | NON-ALLERGEN | Non-Toxin | -0.2 | -0.08 | 0.42 | -1 | 4.68 | 1599.03 | outside |
| 977 | 991 | LNDILSRLDKVEAE | 0.0675 | 14 | ABCPRED | 1 | NON-ALLERGEN | Non-Toxin | -0.24 | -0.34 | 0.66 | -2 | 4.32 | 1615.02 | outside |
| 881 | 901 | TITSGWTFGAGAALQIPFAM | 0.4779 | 20 | ABCPRED | 1 | NON-ALLERGEN | Non-Toxin | 0.19 | 0.87 | -0.89 | 0 | 5.88 | 2040.66 | outside |
| 324 | 344 | ESIVRFPNITNLCPFGEVFN | 0.2993 | 20 | ABCPRED | 1 | NON-ALLERGEN | Non-Toxin | -0.02 | 0.25 | -0.37 | -1 | 4.54 | 2296.92 | outside |
| 1088 | 1108 | HFPREGVFVSNGTHWFVTQR | 0.3144 | 20 | ABCPRED | 1 | NON-ALLERGEN | Non-Toxin | -0.14 | -0.52 | -0.38 | 2 | 9.96 | 2401.97 | outside |
| 889 | 903 | GAGAALQIPFAMQM | 0.6971 | 14 | ABCPRED | 1 | NON-ALLERGEN | Non-Toxin | 0.16 | 0.91 | -0.74 | 0 | 5.88 | 1405.9 | outside |
| 562 | 576 | FQQFGRDIADTTDA | 0.1813 | 14 | ABCPRED | 1 | ALLERGEN | Non-Toxin | -0.22 | -0.72 | 0.27 | -2 | 3.94 | 1584.85 | outside |
| 1050 | 1066 | MSFPQSAPHGVVFLHV | 0.251 | 16 | ABCPRED | 1 | NON-ALLERGEN | Non-Toxin | 0.12 | 0.66 | -0.83 | 1 | 7.26 | 1753.29 | outside |
| 553 | 567 | TESNKKFLPFQQFG | 0.6269 | 14 | ABCPRED | 1 | NON-ALLERGEN | Non-Toxin | -0.2 | -0.94 | 0.01 | 1 | 8.94 | 1671.09 | outside |
| 833 | 847 | FIKQYGDCLGDIAA | -0.2548 | 14 | ABCPRED | 1 | NON-ALLERGEN | Non-Toxin | 0.02 | 0.37 | -0.21 | -1 | 4.21 | 1513.93 | outside |
| 307 | 323 | TVEKGIYQTSNFRVQP | 0.6733 | 16 | ABCPRED | 1 | ALLERGEN | Toxin | -0.22 | -0.76 | -0.03 | 1 | 8.93 | 1867.33 | outside |
| 1252 | 1274 | SCCKFDEDDSEPVLKGVKLHYT | 0.8094 | 22 | AAP | 1 | ALLERGEN | Non-Toxin | -0.22 | -0.65 | 0.47 | -1.5 | 4.83 | 2514.13 | inside |
| 465 | 509 | ERDISTENCYFPLQSYGFQVGYQPYR | 0.8125 | 26 | ellipro,literature | 2 | ALLERGEN | Non-Toxin | -0.21 | -0.94 | -0.17 | -1 | 4.68 | 3161.83 | outside |
| 1087 | 1101 | AHFPREGVFVSNGT | 0.5016 | 14 | ABCPRED | 1 | NON-ALLERGEN | Non-Toxin | -0.08 | -0.2 | -0.21 | 0.5 | 7.1 | 1517.86 | outside |
| 1140 | 1156 | PLQPELDSFKEELDKY | -0.5822 | 16 | FBCPred | 1 | NON-ALLERGEN | Toxin | -0.27 | -1.24 | 0.71 | -3 | 4.18 | 1951.39 | outside |
| 4 | 19 | FLVLLPLVSSQCVNL | 0.8302 | 15 | IEDB,literature | 2 | NON-ALLERGEN | Non-Toxin | 0.2 | 1.78 | -1.07 | 0 | 5.85 | 1645.27 | outside |
| 193 | 213 | VFKNIDGYFKIYSKHTPINL | 0.3566 | 20 | ABCPRED | 1 | NON-ALLERGEN | Non-Toxin | -0.08 | -0.22 | -0.33 | 2.5 | 9.43 | 2398.11 | outside |
| 404 | 424 | GDEVRQIAPGQTGKIADYNY | 1.219 | 20 | ABCPRED | 1 | ALLERGEN | Non-Toxin | -0.21 | -0.93 | 0.22 | -1 | 4.56 | 2195.67 | outside |
| 564 | 578 | QFGRDIADTTDAVR | 0.1003 | 14 | ABCPRED | 1 | NON-ALLERGEN | Non-Toxin | -0.3 | -0.69 | 0.54 | -1 | 4.43 | 1564.86 | outside |
| 1054 | 1074 | QSAPHGVVFLHVTYVPAQEK | 0.794 | 20 | ABCPRED | 1 | NON-ALLERGEN | Non-Toxin | -0.02 | -0.01 | -0.42 | 1 | 7.25 | 2207.82 | outside |
| 202 | 222 | KIYSKHTPINLVRDLPQGFS | 0.5542 | 20 | ABCPRED | 1 | NON-ALLERGEN | Non-Toxin | -0.17 | -0.48 | -0.07 | 2.5 | 9.72 | 2313.99 | outside |
| 1009 | 1023 | TQQLIRAAEIRASA | 0.0932 | 14 | ABCPRED | 1 | NON-ALLERGEN | Non-Toxin | -0.21 | -0.07 | 0.14 | 1 | 9.95 | 1527.94 | outside |
| 556 | 570 | NKKFLPFQQFGRDI | 0.37 | 14 | ABCPRED | 1 | NON-ALLERGEN | Non-Toxin | -0.25 | -0.83 | 0.11 | 2 | 10.01 | 1738.23 | outside |
| 125 | 139 | NVVIKVCEFQFCND | -0.6418 | 14 | ABCPRED | 1 | NON-ALLERGEN | Non-Toxin | -0.05 | 0.45 | -0.26 | -1 | 4.38 | 1658.12 | outside |
| 903 | 917 | AYRFNGIGVTQNVL | 0.7506 | 14 | ABCPRED | 1 | NON-ALLERGEN | Non-Toxin | -0.03 | 0.25 | -0.62 | 1 | 9.1 | 1551.98 | outside |
| 287 | 318 | DAVDCALDPLSETKCTLKSFTVEKGIYQTSN | 0.5013 | 31 | literature | 1 | NON-ALLERGEN | Non-Toxin | -0.16 | -0.29 | 0.18 | -2 | 4.44 | 3378.23 | outside |
| 1086 | 1102 | KAHFPREGVFVSNGTH | 0.2948 | 16 | ABCPRED,AAP | 2 | ALLERGEN | Non-Toxin | -0.16 | -0.62 | -0.03 | 2 | 9.11 | 1783.21 | outside |
| 197 | 211 | IDGYFKIYSKHTPI | 0.5093 | 14 | ABCPRED | 1 | NON-ALLERGEN | Non-Toxin | -0.06 | -0.31 | -0.29 | 1.5 | 8.76 | 1682.16 | outside |
| 1063 | 1077 | LHVTYVPAQEKNFT | 1.2382 | 14 | ABCPRED | 1 | NON-ALLERGEN | Non-Toxin | -0.1 | -0.36 | -0.36 | 0.5 | 7.1 | 1647.07 | outside |
| 257 | 277 | GWTAGAAAYYVGYLQPRTFL | 0.533 | 20 | ABCPRED | 1 | NON-ALLERGEN | Non-Toxin | 0.06 | 0.24 | -0.88 | 1 | 8.83 | 2205.79 | outside |
| 501 | 517 | NGVGYQPYRVVVLSFE | 0.9736 | 16 | ABCPRED,BCPREDS | 2 | NON-ALLERGEN | Non-Toxin | -0.02 | 0.16 | -0.51 | 0 | 6.35 | 1827.31 | outside |
| 391 | 407 | CFTNVYADSFVIRGDE | 0.0861 | 16 | ABCPRED | 1 | NON-ALLERGEN | Non-Toxin | -0.09 | 0.07 | -0.09 | -2 | 4.03 | 1836.22 | outside |
| 370 | 384 | NSASFSTFKCYGVS | 0.6982 | 14 | ABCPRED | 1 | ALLERGEN | Non-Toxin | -0.05 | 0.08 | -0.45 | 1 | 8.54 | 1497.82 | outside |
| 380 | 395 | YGVSPTKLNDLCFTN | 2.0602 | 15 | literature | 1 | NON-ALLERGEN | Non-Toxin | -0.09 | -0.19 | -0.33 | 0 | 6.15 | 1672.1 | outside |
| 1069 | 1147 | REGVFVSNGTHWFVTQRNFYEPQIITTDNTFVS | 0.3566 | 78 | ellipro,literature | 2 | ALLERGEN | too long | | | | | | | outside |
| 713 | 727 | AIPTNFTISVTTEI | 0.8574 | 14 | ABCPRED | 1 | NON-ALLERGEN | Non-Toxin | 0.09 | 0.72 | -0.57 | -1 | 4 | 1506.92 | outside |
| 593 | 613 | GGVSVITPGTNTSNQVAVLY | 0.4308 | 20 | ABCPRED | 1 | ALLERGEN | Non-Toxin | 0.05 | 0.43 | -0.62 | 0 | 5.88 | 1977.51 | outside |
| 572 | 586 | TTDAVRDPQTLEIL | 0.384 | 14 | ABCPRED | 1 | ALLERGEN | Non-Toxin | -0.18 | -0.29 | 0.26 | -2 | 4.03 | 1571.95 | outside |
| 390 | 404 | LCFTNVYADSFVIR | 0.5206 | 14 | ABCPRED | 1 | NON-ALLERGEN | Non-Toxin | 0.02 | 0.88 | -0.66 | 0 | 6.15 | 1648.1 | outside |
| 906 | 920 | FNGIGVTQNVLYEN | 0.498 | 14 | ABCPRED | 1 | NON-ALLERGEN | Non-Toxin | -0.01 | -0.06 | -0.57 | -1 | 4 | 1567.94 | outside |
| 1084 | 1104 | DGKAHFPREGVFVSNGTHWF | 0.1005 | 20 | ABCPRED | 1 | NON-ALLERGEN | Non-Toxin | -0.11 | -0.59 | -0.16 | 1 | 7.26 | 2288.8 | outside |
| 187 | 201 | KNLREFVFKNIDGY | -0.4161 | 14 | ABCPRED | 1 | NON-ALLERGEN | Non-Toxin | -0.24 | -0.71 | 0.21 | 1 | 8.83 | 1743.2 | outside |
| 462 | 495 | KPFERDISTEIYQAGSTPCNGVEGFNCYFPLQS | 0.1632 | 33 | literature | 1 | ALLERGEN | Non-Toxin | -0.13 | -0.5 | -0.08 | -2 | 4.41 | 3699.56 | outside |
| 14 | 44 | QCVNLTTRTQLPPAYTNSFTRGVYYPDKVF | 0.6331 | 30 | literature | 1 | NON-ALLERGEN | Non-Toxin | -0.17 | -0.46 | -0.35 | 2 | 9.16 | 3481.38 | outside |
| 582 | 602 | LEILDITPCSFGGVSVITPG | 1.2631 | 20 | ABCPRED | 1 | NON-ALLERGEN | Non-Toxin | 0.15 | 1.02 | -0.49 | -2 | 3.67 | 2018.66 | outside |
| 330 | 350 | PNITNLCPFGEVFNATRFAS | 0.5575 | 20 | ABCPRED | 1 | ALLERGEN | Non-Toxin | -0.04 | 0.13 | -0.42 | 0 | 6.32 | 2198.76 | outside |
| 870 | 884 | IAQYTSALLAGTIT | 0.4542 | 14 | ABCPRED | 1 | NON-ALLERGEN | Non-Toxin | 0.14 | 0.99 | -0.84 | 0 | 5.88 | 1422.85 | outside |
| 444 | 458 | KVGGNYNYLYRLFR | -0.2536 | 14 | ABCPRED | 1 | NON-ALLERGEN | Non-Toxin | -0.24 | -0.71 | -0.36 | 3 | 10 | 1763.24 | outside |
| 640 | 654 | SNVFQTRAGCLIGA | 0.7968 | 14 | ABCPRED | 1 | NON-ALLERGEN | Non-Toxin | -0.02 | 0.54 | -0.45 | 1 | 8.6 | 1436.85 | outside |
| 16 | 36 | VNLTTRTQLPPAYTNSFTRG | 0.6822 | 20 | ABCPRED | 1 | NON-ALLERGEN | Non-Toxin | -0.21 | -0.62 | -0.28 | 2 | 10.84 | 2237.79 | outside |
| 1182 | 1210 | EIDRLNEVAKNLNESLIDLQELGKYEQY | 0.4662 | 28 | COVIDep,literature | 2 | NON-ALLERGEN | Non-Toxin | -0.25 | -0.86 | 0.43 | -4 | 4.26 | 3338.13 | outside |
| 846 | 860 | ARDLICAQKFNGLT | 0.199 | 14 | ABCPRED | 1 | NON-ALLERGEN | Non-Toxin | -0.14 | 0.07 | -0.06 | 1 | 8.57 | 1550.01 | outside |
| 279 | 299 | YNENGTITDAVDCALDPLSE | 0.4678 | 20 | ABCPRED | 1 | NON-ALLERGEN | Non-Toxin | -0.12 | -0.38 | 0.18 | -5 | 3.34 | 2140.54 | outside |
| 268 | 282 | GYLQPRTFLLKYNE | 0.3921 | 14 | ABCPRED | 1 | NON-ALLERGEN | Non-Toxin | -0.19 | -0.71 | -0.25 | 1 | 8.83 | 1742.22 | outside |
| 342 | 356 | FNATRFASVYAWNR | 0.1267 | 14 | ABCPRED | 1 | ALLERGEN | Non-Toxin | -0.17 | -0.32 | -0.53 | 2 | 10.84 | 1703.07 | outside |
| 539 | 559 | VNFNFNGLTGTGVLTESNKK | 1.1457 | 20 | ABCPRED | 1 | NON-ALLERGEN | Non-Toxin | -0.12 | -0.39 | -0.14 | 1 | 8.94 | 2140.69 | outside |
| 28 | 48 | YTNSFTRGVYYPDKVFRSSV | -0.1994 | 20 | ABCPRED | 1 | NON-ALLERGEN | Toxin | -0.21 | -0.57 | -0.2 | 2 | 9.55 | 2386.92 | outside |
| 318 | 338 | FRVQPTESIVRFPNITNLCP | 0.8067 | 20 | ABCPRED | 1 | ALLERGEN | Non-Toxin | -0.13 | -0.03 | -0.27 | 1 | 8.6 | 2332.01 | outside |
| 153 | 169 | MESEFRVYSSANNCTF | 0.1397 | 16 | ABCPRED,AAP | 2 | ALLERGEN | Non-Toxin | -0.18 | -0.43 | -0.11 | -1 | 4.54 | 1885.26 | outside |
| 76 | 92 | TKRFDNPVLPFNDGVY | 0.2219 | 16 | ABCPRED,AAP | 2 | NON-ALLERGEN | Non-Toxin | -0.18 | -0.64 | -0.01 | 0 | 6.31 | 1882.33 | outside |
| 609 | 629 | AVLYQDVNCTEVPVAIHADQ | 0.4849 | 20 | ABCPRED | 1 | ALLERGEN | Non-Toxin | -0.02 | 0.26 | -0.28 | -2.5 | 4.03 | 2185.73 | outside |
| 862 | 882 | PPLLTDEMIAQYTSALLAGT | 0.3552 | 20 | ABCPRED | 1 | NON-ALLERGEN | Non-Toxin | 0.05 | 0.43 | -0.44 | -2 | 3.67 | 2105.73 | outside |
| 882 | 898 | ITSGWTFGAGAALQIP | 0.2071 | 16 | ABCPRED | 1 | ALLERGEN | Non-Toxin | 0.18 | 0.72 | -0.82 | 0 | 5.88 | 1590.05 | outside |
| 714 | 734 | IPTNFTISVTTEILPVSMTK | 1.0688 | 20 | ABCPRED | 1 | NON-ALLERGEN | Non-Toxin | 0.04 | 0.56 | -0.46 | 0 | 6.35 | 2192.9 | outside |
| 942 | 956 | ASALGKLQDVVNQN | 0.5261 | 14 | ABCPRED | 1 | NON-ALLERGEN | Non-Toxin | -0.14 | -0.21 | -0.04 | 0 | 6.19 | 1456.83 | outside |
| 564 | 580 | QFGRDIADTTDAVRDP | 0.0689 | 16 | ABCPRED | 1 | NON-ALLERGEN | Non-Toxin | -0.31 | -0.92 | 0.66 | -2 | 4.14 | 1777.09 | outside |
| 169 | 189 | EYVSQPFLMDLEGKQGNFKN | 0.8014 | 20 | ABCPRED | 1 | ALLERGEN | Non-Toxin | -0.19 | -0.88 | 0.12 | -1 | 4.68 | 2344.93 | outside |
| 953 | 967 | NQNAQALNTLVKQL | -0.0658 | 14 | ABCPRED | 1 | NON-ALLERGEN | Non-Toxin | -0.19 | -0.46 | -0.29 | 1 | 9.11 | 1554.99 | outside |
| 867 | 887 | DEMIAQYTSALLAGTITSGW | 0.2418 | 20 | ABCPRED | 1 | NON-ALLERGEN | Non-Toxin | 0.06 | 0.33 | -0.51 | -2 | 3.67 | 2128.68 | outside |
| 289 | 309 | VDCALDPLSETKCTLKSFTV | 0.7027 | 20 | ABCPRED | 1 | NON-ALLERGEN | Non-Toxin | -0.09 | 0.29 | 0.05 | -1 | 4.56 | 2170.8 | outside |
| 242 | 256 | LALHRSYLTPGDSS | 0.6223 | 14 | ABCPRED | 1 | NON-ALLERGEN | Non-Toxin | -0.13 | -0.31 | -0.16 | 0.5 | 7.09 | 1516.87 | outside |
| 316 | 338 | SNFRVQPTESIVRFPNITNLCP | 0.8386 | 22 | FBCPred | 1 | NON-ALLERGEN | Non-Toxin | -0.16 | -0.23 | -0.22 | 1 | 8.6 | 2533.22 | outside |
| 605 | 619 | SNQVAVLYQDVNCT | 0.7849 | 14 | ABCPRED | 1 | NON-ALLERGEN | Non-Toxin | -0.1 | 0.03 | -0.46 | -1 | 3.8 | 1553.91 | outside |
| 501 | 521 | NGVGYQPYRVVVLSFELLHA | 0.9212 | 20 | ABCPRED | 1 | NON-ALLERGEN | Non-Toxin | 0.03 | 0.44 | -0.64 | 0.5 | 7.1 | 2261.92 | outside |
| 152 | 168 | WMESEFRVYSSANNCT | 0.022 | 16 | ABCPRED | 1 | NON-ALLERGEN | Non-Toxin | -0.2 | -0.66 | -0.16 | -1 | 4.54 | 1924.3 | outside |

| | | | | | | | | | | | | | | | |
|---|---|---|---|---|---|---|---|---|---|---|---|---|---|---|---|
| 1001 | 1015 | LQSLQTYVTQQLIR | -0.3524 | 14 | ABCPRED | 1 | NON-ALLERGEN | Non-Toxin | -0.16 | -0.14 | -0.55 | 1 | 9.1 | 1691.19 | outside |
| 199 | 213 | GYFKIYSKHTPINL | 0.9308 | 14 | ABCPRED | 1 | ALLERGEN | Non-Toxin | -0.07 | -0.36 | -0.49 | 2.5 | 9.55 | 1681.18 | outside |
| 872 | 894 | QYTSALLAGTITSGWTFGAGAA | 0.4277 | 22 | AAP | 1 | NON-ALLERGEN | Non-Toxin | 0.12 | 0.55 | -0.77 | 0 | 5.88 | 2144.69 | outside |
| 236 | 252 | TRFQTLLALHRSYLTP | 0.5115 | 16 | ABCPRED | 1 | ALLERGEN | Non-Toxin | -0.16 | -0.11 | -0.48 | 2.5 | 10.84 | 1917.48 | outside |
| 325 | 339 | SIVRFPNITNLCPF | 0.7956 | 14 | ABCPRED | 1 | NON-ALLERGEN | Non-Toxin | 0.01 | 0.64 | -0.69 | 1 | 8.6 | 1621.13 | outside |
| 282 | 304 | NGTITDAVDCALDPLSETKCTL | 0.6769 | 22 | FBCPred | 1 | NON-ALLERGEN | Non-Toxin | -0.09 | 0.08 | 0.1 | -3 | 3.84 | 2280.86 | outside |
| 295 | 309 | PLSETKCTLKSFTV | 0.7254 | 14 | ABCPRED | 1 | NON-ALLERGEN | Non-Toxin | -0.12 | 0.04 | -0.01 | 1 | 8.54 | 1554.03 | outside |
| 871 | 885 | AQYTSALLAGTITS | 0.6271 | 14 | ABCPRED | 1 | NON-ALLERGEN | Non-Toxin | 0.07 | 0.61 | -0.69 | 0 | 5.88 | 1396.76 | outside |
| 575 | 595 | AVRDPQTLEILDITPCSFGG | 1.1985 | 20 | ABCPRED | 1 | NON-ALLERGEN | Non-Toxin | -0.05 | 0.16 | -0.05 | -2 | 4.03 | 2132.71 | outside |
| 127 | 147 | VIKVCEFQFCNDPFLGVYYH | -0.1316 | 20 | ABCPRED | 1 | NON-ALLERGEN | Toxin | 0.04 | 0.43 | -0.67 | -0.5 | 5.33 | 2422.11 | outside |
| 772 | 794 | VEQDKNTQEVFAQVKQIYKTPP | 0.2192 | 22 | BCPREDS | 1 | NON-ALLERGEN | Non-Toxin | -0.27 | -1.09 | 0.3 | 0 | 6.52 | 2591.25 | outside |
| 70 | 86 | VSGTNGTKRFDNPVLP | 0.5162 | 16 | ABCPRED | 1 | NON-ALLERGEN | Non-Toxin | -0.19 | -0.63 | 0.1 | 1 | 9.1 | 1702.13 | outside |
| 685 | 699 | RSVASQSIIAYTMS | 0.1589 | 14 | ABCPRED | 1 | NON-ALLERGEN | Toxin | -0.06 | 0.39 | -0.41 | 1 | 9.1 | 1513.92 | outside |
| 822 | 842 | LFNKVTLADAGFIKQYGDCL | 0.3335 | 20 | ABCPRED | 1 | NON-ALLERGEN | Non-Toxin | -0.01 | 0.36 | -0.3 | 0 | 6.27 | 2216.88 | outside |
| 673 | 687 | SYQTQTNSPRRARS | 0.1125 | 14 | ABCPRED | 1 | NON-ALLERGEN | Non-Toxin | -0.59 | -2.06 | 0.49 | 3 | 11.72 | 1651.94 | outside |
| 782 | 796 | FAQVKQIYKTPPIK | 0.3354 | 14 | ABCPRED | 1 | NON-ALLERGEN | Non-Toxin | -0.15 | -0.44 | -0.1 | 3 | 10.01 | 1661.23 | outside |
| 109 | 129 | TLDSKTQSLLIVNNATNVVI | 0.678 | 20 | ABCPRED | 1 | NON-ALLERGEN | Non-Toxin | -0.03 | 0.48 | -0.39 | 0 | 6.19 | 2143.78 | outside |
| 877 | 893 | LLAGTITSGWTFGAGA | 0.3702 | 16 | AAP,FBCPred | 2 | NON-ALLERGEN | Non-Toxin | 0.21 | 0.93 | -0.86 | 0 | 5.88 | 1522.96 | outside |
| 626 | 648 | ADQLTPTWRVYSTGSNVFQTRA | 0.5336 | 22 | FBCPred | 1 | NON-ALLERGEN | Non-Toxin | -0.2 | -0.59 | -0.25 | 1 | 9.1 | 2499.05 | outside |
| 601 | 607 | GTNTSN | 1.2831 | 6 | IEDB,literature | 2 | ALLERGEN | Non-Toxin | -0.29 | -1.6 | -0.02 | 0 | 5.88 | 592.64 | outside |
| 797 | 817 | FGGFNFSQILPDPSKPSKRS | 0.6909 | 20 | ABCPRED | 1 | ALLERGEN | Non-Toxin | -0.19 | -0.74 | 0.12 | 2 | 10.01 | 2209.77 | outside |
| 683 | 697 | RARSVASQSIIAYT | 0.4747 | 14 | ABCPRED | 1 | NON-ALLERGEN | Non-Toxin | -0.17 | 0.12 | -0.16 | 2 | 10.84 | 1522.91 | outside |
| 345 | 365 | TRFASVYAWNRKRISNCVAD | 0.2264 | 20 | ABCPRED | 1 | NON-ALLERGEN | Non-Toxin | -0.27 | -0.44 | 0.01 | 3 | 10.06 | 2357.94 | outside |
| 1093 | 1107 | GVFVSNGTHWFVTQ | 0.184 | 14 | ABCPRED | 1 | NON-ALLERGEN | Non-Toxin | 0.08 | 0.29 | -0.96 | 0.5 | 7.1 | 1578.96 | outside |
| 530 | 544 | STNLVKNKCVNFNF | 1.289 | 14 | ABCPRED | 1 | NON-ALLERGEN | Non-Toxin | -0.17 | -0.21 | -0.29 | 2 | 9.36 | 1628.08 | outside |
| 584 | 604 | ILDITPCSFGGVSVITPGTN | 0.9838 | 20 | ABCPRED | 1 | NON-ALLERGEN | Non-Toxin | 0.12 | 0.79 | -0.56 | -1 | 3.8 | 1991.59 | outside |
| 715 | 735 | PTNFTISVTTEILPVSMTKT | 1.2666 | 20 | ABCPRED | 1 | NON-ALLERGEN | Non-Toxin | -0.01 | 0.3 | -0.39 | 0 | 6.35 | 2180.84 | outside |
| 56 | 76 | LPFFSNVTWFHAIHVSGTNG | 0.6458 | 20 | ABCPRED | 1 | NON-ALLERGEN | Non-Toxin | 0.1 | 0.36 | -0.94 | 1 | 7.26 | 2231.8 | outside |
| 890 | 910 | AGAALQIPFAMQMAYRFNGI | 0.7475 | 20 | ABCPRED | 1 | NON-ALLERGEN | Non-Toxin | 0.07 | 0.63 | -0.71 | 1 | 9.1 | 2170.87 | outside |
| 100 | 120 | IIRGWIFGTTLDSKTQSLLI | 0.1427 | 20 | ABCPRED | 1 | NON-ALLERGEN | Non-Toxin | 0.02 | 0.57 | -0.5 | 1 | 9.1 | 2263 | outside |
| 641 | 661 | NVFQTRAGCLIGAEHVNNSY | 0.9383 | 20 | ABCPRED | 1 | NON-ALLERGEN | Non-Toxin | -0.1 | -0.16 | -0.36 | 0.5 | 7.06 | 2193.72 | outside |
| 580 | 600 | QTLEILDITPCSFGGVSVIT | 1.3356 | 20 | ABCPRED | 1 | NON-ALLERGEN | Non-Toxin | 0.11 | 0.91 | -0.49 | -2 | 3.67 | 2093.73 | outside |
| 737 | 757 | DCTMYICGDSTECSNLLLQY | -0.2104 | 20 | ABCPRED | 1 | NON-ALLERGEN | Non-Toxin | -0.07 | 0.09 | -0.34 | -3 | 3.5 | 2272.85 | outside |
| 1068 | 1092 | VPAQEKNFTTAPAICHDGKAHFPR | 0.0607 | 24 | literature | 1 | ALLERGEN | Non-Toxin | -0.18 | -0.64 | 0.1 | 2 | 8.54 | 2636.32 | outside |
| 1127 | 1147 | DVVIGIVNNTVYDPLQPELD | 0.7802 | 20 | ABCPRED | 1 | ALLERGEN | Non-Toxin | -0.01 | 0.16 | -0.16 | -4 | 3.38 | 2213.78 | outside |
| 266 | 286 | YVGYLQPRTFLLKYNENGTI | 0.5395 | 20 | ABCPRED | 1 | ALLERGEN | Non-Toxin | -0.1 | -0.36 | -0.47 | 1 | 8.76 | 2390.05 | outside |
| 639 | 659 | GSNVFQTRAGCLIGAEHVNN | 0.8762 | 20 | ABCPRED | 1 | NON-ALLERGEN | Non-Toxin | -0.09 | -0.12 | -0.24 | 0.5 | 7.07 | 2087.6 | outside |
| 721 | 735 | SVTTEILPVSMTKT | 1.1868 | 14 | ABCPRED | 1 | NON-ALLERGEN | Non-Toxin | -0.03 | 0.37 | -0.21 | 0 | 6.35 | 1506.98 | outside |
| 537 | 553 | KCVNFNFNGLTGTGVL | 1.4136 | 16 | AAP | 1 | NON-ALLERGEN | Non-Toxin | 0.03 | 0.44 | -0.61 | 1 | 8.57 | 1684.19 | outside |
| 973 | 987 | ISSVLNDILSRLDK | -0.3367 | 14 | ABCPRED | 1 | NON-ALLERGEN | Non-Toxin | -0.15 | 0.24 | 0.19 | 0 | 6.31 | 1573.03 | outside |
| 163 | 183 | ANNCTFEYVSQPFLMDLEGK | 0.4346 | 20 | ABCPRED | 1 | NON-ALLERGEN | Non-Toxin | -0.1 | -0.3 | -0.14 | -2 | 4.14 | 2306.88 | outside |
| 670 | 684 | ICASYQTQTNSPRR | 0.2317 | 14 | ABCPRED | 1 | NON-ALLERGEN | Non-Toxin | -0.39 | -1.19 | 0.06 | 2 | 9.55 | 1624.98 | outside |
| 569 | 583 | IADTTDAVRDPQTL | 0.3617 | 14 | ABCPRED | 1 | NON-ALLERGEN | Non-Toxin | -0.21 | -0.44 | 0.35 | -2 | 3.94 | 1515.83 | outside |
| 473 | 495 | YQAGSTPCNGVEGFNCYFPLQS | 0.475 | 22 | AAP,BCPREDS | 2 | NON-ALLERGEN | Non-Toxin | -0.04 | -0.29 | -0.52 | -1 | 4 | 2382.92 | outside |
| 1104 | 1126 | VTQRNFYEPQIITTDNTFVSGN | 0.5011 | 22 | AAP | 1 | ALLERGEN | Non-Toxin | -0.16 | -0.59 | -0.24 | -1 | 4.38 | 2545.09 | outside |
| 253 | 267 | DSSSGWTAGAAAYY | 0.4142 | 14 | ABCPRED | 1 | NON-ALLERGEN | Non-Toxin | 0 | -0.26 | -0.46 | -1 | 3.8 | 1406.6 | outside |
| 1201 | 1217 | QELGKYEQYIKWPWYI | 0.841 | 16 | ABCPRED,BCPREDS | 2 | NON-ALLERGEN | Non-Toxin | -0.12 | -1.04 | -0.42 | 0 | 6.48 | 2144.71 | outside_transmembrane-region |
| 1050 | 1064 | MSFPQSAPHGVVFL | 0.1283 | 14 | ABCPRED | 1 | ALLERGEN | Non-Toxin | 0.13 | 0.69 | -0.81 | 0.5 | 7.1 | 1516.98 | outside |
| 1152 | 1172 | LDKYFKNHTSPDVDLGDISG | 0.4537 | 20 | ABCPRED | 1 | ALLERGEN | Non-Toxin | -0.18 | -0.77 | 0.31 | -1.5 | 4.56 | 2221.7 | outside |
| 116 | 136 | SLLIVNNATNVVIKVCEFQF | 0.2947 | 20 | ABCPRED | 1 | NON-ALLERGEN | Non-Toxin | 0.07 | 1.02 | -0.65 | 0 | 6.32 | 2251.98 | outside |
| 1050 | 1070 | MSFPQSAPHGVVFLHVTYVP | 0.6215 | 20 | ABCPRED | 1 | ALLERGEN | Non-Toxin | 0.11 | 0.56 | -0.88 | 1 | 7.25 | 2213.88 | outside |
| 578 | 598 | DPQTLEILDITPCSFGGVSV | 1.5468 | 20 | ABCPRED | 1 | NON-ALLERGEN | Non-Toxin | 0.04 | 0.47 | -0.24 | -3 | 3.5 | 2091.66 | outside |
| 83 | 97 | VLPFNDGVYFASTE | 0.5346 | 14 | ABCPRED | 1 | NON-ALLERGEN | Non-Toxin | 0.06 | 0.31 | -0.46 | -2 | 3.67 | 1558.9 | outside |
| 642 | 656 | VFQTRAGCLIGAEH | 1.2294 | 14 | ABCPRED | 1 | NON-ALLERGEN | Non-Toxin | -0.03 | 0.37 | -0.31 | 0.5 | 7.07 | 1501.93 | outside |
| 658 | 678 | NSYECDIPIGAGICASYQTQ | 0.7339 | 20 | ABCPRED | 1 | NON-ALLERGEN | Non-Toxin | -0.05 | -0.13 | -0.31 | -2 | 3.67 | 2133.62 | outside |
| 1051 | 1065 | SFPQSAPHGVVFLH | 0.1221 | 14 | ABCPRED | 1 | ALLERGEN | Non-Toxin | 0.08 | 0.32 | -0.75 | 1 | 7.26 | 1522.93 | outside |
| 252 | 266 | GDSSSGWTAGAAAY | 0.2846 | 14 | ABCPRED | 1 | NON-ALLERGEN | Non-Toxin | 0.01 | -0.2 | -0.3 | -1 | 3.8 | 1300.48 | outside |
| 101 | 115 | IRGWIFGTTLDSKT | 0.009 | 14 | ABCPRED | 1 | NON-ALLERGEN | Non-Toxin | -0.08 | -0.06 | -0.23 | 1 | 9.1 | 1595.04 | outside |
| 269 | 291 | YLQPRTFLLKYNENGTITDAVD | 0.3694 | 22 | AAP | 1 | NON-ALLERGEN | Non-Toxin | -0.16 | -0.52 | -0.09 | -1 | 4.56 | 2572.2 | outside |
| 487 | 507 | NCYFPLQSYGFQPTNGVGYQ | 0.6877 | 20 | ABCPRED | 1 | ALLERGEN | Non-Toxin | -0.05 | -0.56 | -0.77 | 0 | 5.84 | 2283.8 | outside |
| 590 | 604 | CSFGGVSVITPGTN | 0.6824 | 14 | ABCPRED | 1 | NON-ALLERGEN | Non-Toxin | 0.1 | 0.64 | -0.59 | 0 | 5.85 | 1338.7 | outside |
| 505 | 535 | YQPYRVVVLSFELLHAPATVCGPKKSTNLV | 0.5147 | 30 | literature | 1 | NON-ALLERGEN | Non-Toxin | -0.04 | 0.28 | -0.4 | 2.5 | 9.17 | 3331.39 | outside |
| 418 | 432 | IADYNYKLPDDFTG | 0.7024 | 14 | ABCPRED | 1 | ALLERGEN | Non-Toxin | -0.13 | -0.74 | 0.04 | -2 | 3.94 | 1631.95 | outside |
| 773 | 793 | EQDKNTQEVFAQVKQIYKTP | 0.1742 | 20 | ABCPRED | 1 | NON-ALLERGEN | Non-Toxin | -0.32 | -1.32 | 0.4 | 0 | 6.52 | 2394.97 | outside |
| 1106 | 1126 | QRNFYEPQIITTDNTFVSGN | 0.3656 | 20 | ABCPRED | 1 | ALLERGEN | Non-Toxin | -0.19 | -0.82 | -0.16 | -1 | 4.38 | 2344.82 | outside |
| 327 | 388 | ATRFASVYAWNRKRISNCVADYSVLYNSASFSTI | 0.4565 | 56 | ellipro | 1 | ALLERGEN | too long | | | | | | | outside |
| 886 | 906 | WTFGAGAALQIPFAMQMAYR | 0.8479 | 20 | ABCPRED | 1 | ALLERGEN | Non-Toxin | 0.08 | 0.5 | -0.82 | 1 | 9.1 | 2230.92 | outside |
| 741 | 757 | YICGDSTECSNLLLQY | 0.0499 | 16 | ABCPRED | 1 | NON-ALLERGEN | Non-Toxin | -0.05 | 0.1 | -0.45 | -2 | 3.67 | 1822.27 | outside |
| 364 | 386 | DYSVLYNSASFSTFKCYGVSPT | 0.6349 | 22 | FBCPred | 1 | NON-ALLERGEN | Non-Toxin | -0.04 | 0 | -0.51 | 0 | 6.15 | 2436.97 | outside |
| 813 | 827 | SKRSFIEDLLFNKV | -0.0521 | 14 | ABCPRED | 1 | NON-ALLERGEN | Non-Toxin | -0.21 | -0.18 | 0.28 | 1 | 8.93 | 1696.18 | outside |
| 560 | 574 | LPFQQFGRDIADTT | 0.2497 | 14 | ABCPRED | 1 | NON-ALLERGEN | Non-Toxin | -0.15 | -0.44 | -0.04 | -1 | 4.21 | 1608.97 | outside |
| 1253 | 1273 | CCKFDEDDSEPVLKGVKLHY | 0.8282 | 20 | ABCPRED | 1 | NON-ALLERGEN | Toxin | -0.22 | -0.64 | 0.52 | -1.5 | 4.83 | 2325.92 | inside |
| 604 | 618 | TSNQVAVLYQDVNC | 0.6476 | 14 | ABCPRED | 1 | NON-ALLERGEN | Non-Toxin | -0.1 | 0.03 | -0.46 | -1 | 3.8 | 1553.91 | outside |
| 353 | 367 | WNRKRISNCVADYS | 0.2138 | 14 | ABCPRED | 1 | NON-ALLERGEN | Non-Toxin | -0.37 | -1.01 | 0.18 | 2 | 9.36 | 1712.1 | outside |
| 951 | 965 | VVNQNAQALNTLVK | 0.0937 | 14 | ABCPRED | 1 | NON-ALLERGEN | Non-Toxin | -0.1 | 0.12 | -0.39 | 1 | 9.11 | 1511.96 | outside |
| 82 | 96 | PVLPFNDGVYFAST | 0.3918 | 14 | ABCPRED | 1 | NON-ALLERGEN | Non-Toxin | 0.09 | 0.44 | -0.68 | -1 | 3.8 | 1526.9 | outside |
| 714 | 728 | IPTNFTISVTTEIL | 0.6489 | 14 | ABCPRED | 1 | NON-ALLERGEN | Non-Toxin | 0.11 | 0.86 | -0.66 | -1 | 4 | 1549.01 | outside |

| | | | | | | | | | | | | | | | |
|---|---|---|---|---|---|---|---|---|---|---|---|---|---|---|---|
| 141 | 161 | LGVYYHKNNKSWMESEFRVY | 0.5308 | 20 | ABCPRED | 1 | NON-ALLERGEN | Non-Toxin | -0.22 | -0.97 | -0.17 | 1.5 | 8.71 | 2551.17 | outside |
| 1114 | 1130 | IITTDNTFVSGNCDVV | 0.1669 | 16 | ABCPRED,FBCPred | 2 | NON-ALLERGEN | Non-Toxin | 0.02 | 0.6 | -0.38 | -2 | 3.57 | 1698.11 | outside |
| 879 | 895 | AGTITSGWTFGAGAAL | 0.2868 | 16 | ABCPRED,literature | 2 | NON-ALLERGEN | Non-Toxin | 0.19 | 0.81 | -0.78 | 0 | 5.88 | 1480.87 | outside |
| 423 | 437 | YKLPDDFTGCVIAW | -0.321 | 14 | ABCPRED | 1 | NON-ALLERGEN | Non-Toxin | 0.03 | 0.27 | -0.44 | -1 | 4.21 | 1628.07 | outside |
| 934 | 954 | IQDSLSSTASALGKLQDVVN | 0.5355 | 20 | ABCPRED | 1 | NON-ALLERGEN | Non-Toxin | -0.09 | 0.11 | -0.04 | -1 | 4.21 | 2046.56 | outside |
| 598 | 614 | ITPGTNTSNQVAVLYQ | 0.7026 | 16 | ABCPRED,BCPREDS | 2 | NON-ALLERGEN | Non-Toxin | -0.05 | -0.11 | -0.59 | 0 | 5.88 | 1706.13 | outside |
| 400 | 414 | FVIRGDEVRQIAPG | 0.5476 | 14 | ABCPRED | 1 | NON-ALLERGEN | Non-Toxin | -0.14 | 0.01 | 0.19 | 0 | 6.42 | 1556.99 | outside |
| 1235 | 1255 | CCMTSCCSCLKGCCSCGSCC | -0.2485 | 20 | ABCPRED | 1 | ALLERGEN | Non-Toxin | -0.04 | 1.1 | -0.47 | 1 | 7.87 | 1985.7 | transmembrane-region_inside |
| 359 | 373 | SNCVADYSVLYNSA | -0.1801 | 14 | ABCPRED | 1 | NON-ALLERGEN | Non-Toxin | -0.04 | 0.2 | -0.51 | -1 | 3.8 | 1505.8 | outside |
| 818 | 832 | IEDLLFNKVTLADA | 0.5976 | 14 | ABCPRED | 1 | ALLERGEN | Non-Toxin | 0 | 0.56 | -0.03 | -2 | 4.03 | 1562 | outside |
| 725 | 745 | EILPVSMTKTSVDCTMYICG | 0.7398 | 20 | ABCPRED | 1 | NON-ALLERGEN | Toxin | 0.01 | 0.6 | -0.34 | -1 | 4.38 | 2191.91 | outside |
| 522 | 538 | ATVCGPKKSTNLVKNK | 0.1893 | 16 | ABCPRED,ellipro | 2 | NON-ALLERGEN | Non-Toxin | -0.27 | -0.64 | 0.35 | 4 | 10.05 | 1688.25 | outside |
| 610 | 624 | VLYQDVNCTEVPVA | 0.6762 | 14 | ABCPRED | 1 | NON-ALLERGEN | Non-Toxin | 0.01 | 0.52 | -0.4 | -2 | 3.67 | 1549.96 | outside |
| 595 | 609 | VSVITPGTNTSNQV | 0.3548 | 14 | ABCPRED | 1 | NON-ALLERGEN | Non-Toxin | -0.04 | 0.06 | -0.45 | 0 | 5.88 | 1416.76 | outside |
| 1218 | 1238 | LGFIAGLIAIVMVTIMLCCM | 0.6641 | 20 | ABCPRED | 1 | ALLERGEN | Non-Toxin | 0.39 | 2.67 | -1.27 | 0 | 5.82 | 2113.12 | transmembrane-region |
| 380 | 400 | YGVSPTKLNDLCFTNVYADS | 1.361 | 20 | ABCPRED | 1 | NON-ALLERGEN | Non-Toxin | -0.08 | -0.12 | -0.3 | -1 | 4.21 | 2207.72 | outside |
| 659 | 679 | SYECDIPIGAGICASYQTQT | 0.7894 | 20 | ABCPRED | 1 | NON-ALLERGEN | Non-Toxin | -0.03 | 0 | -0.34 | -2 | 3.67 | 2120.62 | outside |
| 59 | 73 | FSNVTWFHAIHVSG | 0.5564 | 14 | ABCPRED | 1 | NON-ALLERGEN | Non-Toxin | 0.12 | 0.49 | -1.02 | 1 | 7.26 | 1601.99 | outside |
| 626 | 642 | ADQLTPTWRVYSTGSN | 0.6074 | 16 | FBCPred | 1 | NON-ALLERGEN | Non-Toxin | -0.19 | -0.82 | -0.23 | 0 | 6.19 | 1796.15 | outside |
| 672 | 686 | ASYQTQTNSPRRAR | 0.2419 | 14 | ABCPRED | 1 | NON-ALLERGEN | Non-Toxin | -0.55 | -1.88 | 0.44 | 3 | 11.72 | 1635.94 | outside |
| 1155 | 1175 | YFKNHTSPDVDLGDISGINA | 0.8499 | 20 | ABCPRED | 1 | ALLERGEN | Non-Toxin | -0.1 | -0.45 | -0.01 | -1.5 | 4.42 | 2163.62 | outside |
| 776 | 796 | KNTQEVFAQVKQIYKTPPIK | 0.3427 | 20 | ABCPRED | 1 | NON-ALLERGEN | Non-Toxin | -0.24 | -0.85 | 0.16 | 3 | 9.84 | 2361.09 | outside |
| 1018 | 1038 | IRASANLAATKMSECVLGQS | 0.379 | 20 | ABCPRED | 1 | NON-ALLERGEN | Non-Toxin | -0.1 | 0.27 | -0.07 | 1 | 8.57 | 2050.66 | outside |
| 292 | 308 | ALDPLSETKCTLKSFT | 0.7272 | 16 | FBCPred | 1 | ALLERGEN | Non-Toxin | -0.14 | -0.1 | 0.13 | 0 | 6.38 | 1754.25 | outside |
| 762 | 776 | QLNRALTGIAVEQD | 0.3177 | 14 | ABCPRED | 1 | NON-ALLERGEN | Non-Toxin | -0.16 | -0.23 | 0.09 | -1 | 4.38 | 1527.91 | outside |
| 409 | 423 | QIAPGQTGKIADYN | 1.5448 | 14 | ABCPRED | 1 | NON-ALLERGEN | Non-Toxin | -0.13 | -0.69 | -0.05 | 0 | 6.18 | 1475.83 | outside |
| 786 | 827 | IYKTPPIKDFGGFNFSQILPDPSKPSKRSFIEDLLFN | 0.0915 | 41 | literature | 1 | NON-ALLERGEN | Non-Toxin | -0.17 | -0.54 | 0.14 | 3 | 9.62 | 4742.13 | outside |
| 981 | 1001 | LSRLDKVEAEVQIDRLITGR | -0.1351 | 20 | ABCPRED | 1 | NON-ALLERGEN | Non-Toxin | -0.28 | -0.31 | 0.58 | 0 | 6.56 | 2311.97 | outside |
| 741 | 755 | YICGDSTECSNLLL | -0.0653 | 14 | ABCPRED | 1 | NON-ALLERGEN | Non-Toxin | -0.01 | 0.46 | -0.36 | -2 | 3.67 | 1530.93 | outside |
| 800 | 820 | FNFSQILPDPSKPSKRSFIE | 0.41 | 20 | ABCPRED | 1 | NON-ALLERGEN | Non-Toxin | -0.2 | -0.66 | 0.18 | 1 | 8.93 | 2337.94 | outside |
| 1077 | 1091 | TAPAICHDGKAHFP | 0.1183 | 14 | ABCPRED | 1 | NON-ALLERGEN | Non-Toxin | -0.05 | -0.21 | -0.16 | 1 | 7.22 | 1464.85 | outside |
| 48 | 68 | LHSTQDLFLPFFSNVTWFHA | 0.397 | 20 | ABCPRED | 1 | NON-ALLERGEN | Non-Toxin | 0.07 | 0.31 | -0.93 | 0 | 5.99 | 2408.01 | outside |
| 94 | 114 | STEKSNIIRGWIFGTTLDSK | 0.0044 | 20 | ABCPRED | 1 | NON-ALLERGEN | Non-Toxin | -0.16 | -0.45 | 0.09 | 1 | 8.93 | 2253.84 | outside |
| 44 | 58 | RSSVLHSTQDLFLP | 0.3358 | 14 | ABCPRED | 1 | NON-ALLERGEN | Non-Toxin | -0.13 | -0.07 | -0.23 | 0.5 | 7.1 | 1600.01 | outside |
| 1018 | 1032 | IRASANLAATKMSE | 0.6488 | 14 | ABCPRED | 1 | ALLERGEN | Non-Toxin | -0.16 | -0.02 | 0.18 | 1 | 9.1 | 1462.87 | outside |
| 303 | 317 | LKSFTVEKGIYQTS | -0.0219 | 14 | ABCPRED | 1 | NON-ALLERGEN | Non-Toxin | -0.13 | -0.3 | -0.06 | 1 | 8.83 | 1601.04 | outside |
| 931 | 947 | IGKIQDSLSSTASALG | 0.654 | 16 | ABCPRED | 1 | NON-ALLERGEN | Non-Toxin | -0.02 | 0.29 | -0.08 | 0 | 6.19 | 1547.96 | outside |
| 828 | 842 | LADAGFIKQYGDCL | 0.2417 | 14 | ABCPRED | 1 | NON-ALLERGEN | Non-Toxin | 0 | 0.32 | -0.21 | -1 | 4.21 | 1513.93 | outside |
| 1133 | 1153 | VNNTVYDPLQPELDSFKEEL | 0.0255 | 20 | ABCPRED | 1 | NON-ALLERGEN | Non-Toxin | -0.18 | -0.77 | 0.27 | -4 | 3.84 | 2350.86 | outside |
| 1109 | 1129 | FYEPQIITTDNTFVSGNCDV | 0.1857 | 20 | ABCPRED | 1 | ALLERGEN | Non-Toxin | -0.05 | -0.09 | -0.31 | -3 | 3.5 | 2263.75 | outside |
| 137 | 157 | NDPFLGVYYHKNNKSWMESE | 0.606 | 20 | ABCPRED | 1 | ALLERGEN | Toxin | -0.23 | -1.32 | 0.03 | -0.5 | 5.46 | 2458.98 | outside |
| 423 | 445 | YKLPDDFTGCVIAWNSNNLDSK | 0.2192 | 22 | FBCPred | 1 | ALLERGEN | Non-Toxin | -0.15 | -0.54 | -0.04 | -1 | 4.43 | 2501.08 | outside |
| 1203 | 1223 | LGKYEQYIKWPWYIWLGFIA | 0.9901 | 20 | ABCPRED | 1 | NON-ALLERGEN | Non-Toxin | 0.1 | 0.09 | -1 | 1 | 8.71 | 2575.37 | outside_transmembrane-region |
| 38 | 52 | YPDKVFRSSVLHST | -0.0472 | 14 | ABCPRED | 1 | NON-ALLERGEN | Non-Toxin | -0.2 | -0.44 | -0.04 | 1.5 | 8.94 | 1636.03 | outside |
| 393 | 515 | GKIADYNYKLPDDFTGCVIAWNSNNLDSYNYLYI | 0.4477 | 80 | ellipro,literature | 2 | ALLERGEN | too long | | | | | | | outside |
| 70 | 90 | VSGTNGTKRFDNPVLPFNDG | 0.4955 | 20 | ABCPRED | 1 | ALLERGEN | Non-Toxin | -0.18 | -0.74 | 0.11 | 0 | 6.31 | 2135.61 | outside |
| 825 | 845 | KVTLADAGFIKQYGDCLGDI | 0.3935 | 20 | ABCPRED | 1 | NON-ALLERGEN | Toxin | -0.03 | 0.23 | -0.03 | -1 | 4.43 | 2127.74 | outside |
| 594 | 608 | GVSVITPGTNTSNQ | 0.4772 | 14 | ABCPRED | 1 | NON-ALLERGEN | Non-Toxin | -0.07 | -0.26 | -0.34 | 0 | 5.88 | 1374.68 | outside |
| 959 | 973 | LNTLVKQLSSNFGA | -0.1222 | 14 | ABCPRED | 1 | NON-ALLERGEN | Non-Toxin | -0.04 | 0.22 | -0.44 | 1 | 9.11 | 1491.92 | outside |
| 522 | 536 | ATVCGPKKSTNLVK | 0.0422 | 14 | ABCPRED,IEDB | 2 | NON-ALLERGEN | Non-Toxin | -0.18 | -0.21 | 0.17 | 3 | 9.81 | 1445.94 | outside |
| 1046 | 1066 | GYHLMSFPQSAPHGVVFLHV | 0.2445 | 20 | ABCPRED | 1 | NON-ALLERGEN | Non-Toxin | 0.11 | 0.47 | -0.89 | 1.5 | 7.35 | 2223.89 | outside |
| 420 | 434 | DYNYKLPDDFTGCV | 0.2941 | 14 | ABCPRED | 1 | ALLERGEN | Non-Toxin | -0.16 | -0.71 | 0.03 | -2 | 3.94 | 1649.98 | outside |
| 286 | 300 | TDAVDCALDPLSET | 0.615 | 14 | ABCPRED | 1 | NON-ALLERGEN | Non-Toxin | -0.09 | 0.01 | 0.31 | -4 | 3.38 | 1449.73 | outside |
| 1106 | 1128 | QRNFYEPQIITTDNTFVSGNCD | 0.2875 | 22 | BCPREDS | 1 | NON-ALLERGEN | Non-Toxin | -0.21 | -0.79 | -0.06 | -2 | 4.03 | 2563.07 | outside |
| 599 | 619 | TPGTNTSNQVAVLYQDVNCT | 0.6814 | 20 | ABCPRED | 1 | ALLERGEN | Non-Toxin | -0.12 | -0.36 | -0.37 | -1 | 3.8 | 2125.59 | outside |
| 627 | 643 | DQLTPTWRVYSTGSNV | 0.6483 | 16 | ABCPRED | 1 | NON-ALLERGEN | Non-Toxin | -0.17 | -0.67 | -0.29 | 0 | 6.19 | 1824.21 | outside |
| 1258 | 1274 | EDDSEPVLKGVKLHYT | 1.125 | 16 | ABCPRED,AAP | 2 | NON-ALLERGEN | Non-Toxin | -0.22 | -0.86 | 0.53 | -1.5 | 4.76 | 1830.26 | inside |
| 268 | 288 | GYLQPRTFLLKYNENGTITD | 0.5657 | 20 | ABCPRED | 1 | NON-ALLERGEN | Non-Toxin | -0.17 | -0.71 | -0.14 | 0 | 6.41 | 2343.93 | outside |
| 177 | 197 | MDLEGKQGNFKNLREFVFKN | 0.6174 | 20 | ABCPRED | 1 | ALLERGEN | Non-Toxin | -0.28 | -0.97 | 0.4 | 1 | 8.83 | 2415.07 | outside |
| 414 | 428 | QTGKIADYNYKLPD | 1.2934 | 14 | ABCPRED | 1 | NON-ALLERGEN | Non-Toxin | -0.25 | -1.21 | 0.24 | 0 | 6.3 | 1626 | outside |
| 308 | 322 | VEKGIYQTSNFRVQ | 0.5508 | 14 | ABCPRED | 1 | NON-ALLERGEN | Non-Toxin | -0.24 | -0.71 | -0.01 | 1 | 8.93 | 1669.08 | outside |
| 952 | 974 | VNQNAQALNTLVKQLSSNFGAI | 0.1884 | 22 | AAP | 1 | NON-ALLERGEN | Non-Toxin | -0.07 | 0.06 | -0.44 | 1 | 9.11 | 2330.97 | outside |
| 815 | 835 | RSFIEDLLFNKVTLADAGFI | 0.3128 | 20 | ABCPRED | 1 | NON-ALLERGEN | Non-Toxin | 0 | 0.61 | -0.2 | -1 | 4.56 | 2269.92 | outside |
| 121 | 135 | NNATNVVIKVCEFQ | 0.0357 | 14 | ABCPRED | 1 | NON-ALLERGEN | Non-Toxin | -0.09 | 0.15 | -0.28 | 0 | 6.32 | 1579.01 | outside |
| 464 | 478 | FERDISTEIYQAGS | -0.4408 | 14 | ABCPRED | 1 | NON-ALLERGEN | Non-Toxin | -0.19 | -0.64 | 0.25 | -2 | 4.14 | 1615.91 | outside |
| 691 | 705 | SIIAYTMSLGAENS | 0.717 | 14 | ABCPRED | 1 | NON-ALLERGEN | Non-Toxin | 0.05 | 0.46 | -0.45 | -1 | 4 | 1456.83 | outside |
| 967 | 981 | SSNFGAISSVLNDI | 0.5245 | 14 | ABCPRED | 1 | NON-ALLERGEN | Non-Toxin | 0.04 | 0.54 | -0.38 | -1 | 3.8 | 1423.74 | outside |
| 1201 | 1221 | QELGKYEQYIKWPWYIWLGF | 0.9359 | 20 | ABCPRED | 1 | ALLERGEN | Non-Toxin | -0.01 | -0.57 | -0.72 | 0 | 6.48 | 2648.38 | outside_transmembrane-region |
| 65 | 79 | FHAIHVSGTNGTKR | 0.8667 | 14 | ABCPRED | 1 | ALLERGEN | Non-Toxin | -0.18 | -0.57 | -0.11 | 3 | 11.01 | 1524.91 | outside |
| 846 | 866 | ARDLICAQKFNGLTVLPPLL | 0.374 | 20 | ABCPRED | 1 | NON-ALLERGEN | Non-Toxin | 0 | 0.67 | -0.39 | 1 | 8.57 | 2182.96 | outside |
| 92 | 106 | FASTEKSNIIRGWI | -0.3847 | 14 | ABCPRED | 1 | NON-ALLERGEN | Non-Toxin | -0.09 | -0.06 | -0.17 | 1 | 9.1 | 1622.06 | outside |
| 338 | 354 | FGEVFNATRFASVYAW | -0.0702 | 16 | ABCPRED,AAP | 2 | NON-ALLERGEN | Non-Toxin | 0.05 | 0.41 | -0.72 | 0 | 6.35 | 1865.29 | outside |
| 798 | 812 | GGFNFSQILPDPSK | 0.3944 | 14 | ABCPRED | 1 | NON-ALLERGEN | Non-Toxin | -0.07 | -0.44 | -0.11 | 0 | 6.19 | 1506.88 | outside |
| 66 | 80 | HAIHVSGTNGTKRF | 0.4466 | 14 | ABCPRED | 1 | NON-ALLERGEN | Non-Toxin | -0.18 | -0.57 | -0.11 | 3 | 11.01 | 1524.91 | outside |
| 315 | 329 | TSNFRVQPTESIVR | 0.4901 | 14 | ABCPRED | 1 | NON-ALLERGEN | Non-Toxin | -0.29 | -0.6 | 0.14 | 1 | 9.95 | 1634.02 | outside |
| 477 | 491 | STPCNGVEGFNCYF | 0.2571 | 14 | ABCPRED | 1 | NON-ALLERGEN | Non-Toxin | -0.02 | -0.06 | -0.54 | -1 | 4 | 1537.87 | outside |
| 605 | 625 | SNQVAVLYQDVNCTEVPVAI | 0.7095 | 20 | ABCPRED | 1 | NON-ALLERGEN | Non-Toxin | 0 | 0.5 | -0.43 | -2 | 3.67 | 2162.74 | outside |

| | | | | | | | | | | | | | | | |
|---|---|---|---|---|---|---|---|---|---|---|---|---|---|---|---|
| 462 | 482 | KPFERDISTEIYQAGSTPCN | -0.0922 | 20 | ABCPRED | 1 | NON-ALLERGEN | Non-Toxin | -0.23 | -0.89 | 0.26 | -1 | 4.68 | 2256.75 | outside |
| 537 | 551 | KCVNFNFNGLTGTG | 1.5894 | 14 | ABCPRED | 1 | NON-ALLERGEN | Non-Toxin | -0.04 | -0.06 | -0.46 | 1 | 8.57 | 1471.86 | outside |
| 496 | 510 | GFQPTNGVGYQPYR | 0.7217 | 14 | ABCPRED | 1 | NON-ALLERGEN | Non-Toxin | -0.17 | -1.12 | -0.39 | 1 | 8.93 | 1583.93 | outside |
| 252 | 285 | GDSSSGWTAGAAAYYVGYLQPRTFLLKYNENGT | 0.3826 | 33 | literature | 1 | ALLERGEN | Non-Toxin | -0.08 | -0.44 | -0.35 | 0 | 6.41 | 3559.35 | outside |
| 1028 | 1048 | KMSECVLGQSKRVDFCGKGY | 0.6829 | 20 | ABCPRED | 1 | NON-ALLERGEN | Non-Toxin | -0.23 | -0.45 | 0.29 | 2 | 8.9 | 2235.91 | outside |
| 697 | 711 | MSLGAENSVAYSNN | 0.8278 | 14 | ABCPRED | 1 | NON-ALLERGEN | Non-Toxin | -0.09 | -0.33 | -0.24 | -1 | 4 | 1456.74 | outside |
| 731 | 751 | MTKTSVDCTMYICGDSTECS | 0.1817 | 20 | ABCPRED | 1 | NON-ALLERGEN | Non-Toxin | -0.13 | -0.07 | 0.01 | -2 | 4.03 | 2175.73 | outside |
| 1084 | 1100 | DGKAHFPREGVFVSNG | 0.1337 | 16 | ABCPRED | 1 | ALLERGEN | Non-Toxin | -0.16 | -0.62 | 0.22 | 0.5 | 7.1 | 1717.1 | outside |
| 59 | 79 | FSNVTWFHAIHVSGTNGTKR | 0.7724 | 20 | ABCPRED | 1 | ALLERGEN | Non-Toxin | -0.1 | -0.35 | -0.45 | 3 | 11.01 | 2259.81 | outside |
| 186 | 200 | FKNLREFVFKNIDG | -0.2695 | 14 | ABCPRED | 1 | NON-ALLERGEN | Non-Toxin | -0.2 | -0.41 | 0.2 | 1 | 8.93 | 1727.2 | outside |
| 706 | 720 | AYSNNSIAIPTNFT | 0.575 | 14 | ABCPRED | 1 | ALLERGEN | Non-Toxin | -0.02 | -0.07 | -0.64 | 0 | 5.88 | 1512.83 | outside |
| 686 | 706 | SVASQSIIAYTMSLGAENSV | 0.4849 | 20 | ABCPRED | 1 | NON-ALLERGEN | Non-Toxin | 0.04 | 0.58 | -0.45 | -1 | 4 | 2028.55 | outside |
| 393 | 409 | TNVYADSFVIRGDEVR | -0.2427 | 16 | FBCPred | 1 | ALLERGEN | Non-Toxin | -0.2 | -0.28 | 0.22 | -1 | 4.56 | 1841.23 | outside |
| 136 | 152 | CNDPFLGVYYHKNNKS | 0.6111 | 16 | ABCPRED,AAP | 2 | NON-ALLERGEN | Non-Toxin | -0.23 | -1.07 | -0.12 | 1.5 | 8.48 | 1899.34 | outside |
| 1242 | 1264 | SCLKGCCSCGSCCKFDEDDSEP | 0.1368 | 22 | FBCPred | 1 | ALLERGEN | Non-Toxin | -0.23 | -0.42 | 0.54 | -3 | 4.11 | 2316.84 | inside |
| 691 | 713 | SIIAYTMSLGAENSVAYSNNSI | 0.6111 | 22 | BCPREDS | 1 | ALLERGEN | Non-Toxin | 0.02 | 0.32 | -0.52 | -1 | 4 | 2305.86 | outside |
| 367 | 381 | VLYNSASFSTFKCY | 0.1769 | 14 | ABCPRED | 1 | NON-ALLERGEN | Non-Toxin | -0.01 | 0.34 | -0.76 | 1 | 8.51 | 1630.03 | outside |
| 100 | 116 | IIRGWIFGTTLDSKTQ | 0.0658 | 16 | ABCPRED,AAP | 2 | NON-ALLERGEN | Non-Toxin | -0.07 | 0.01 | -0.3 | 1 | 9.1 | 1836.37 | outside |
| 678 | 698 | TNSPRRARSVASQSIIAYTM | 0.195 | 20 | ABCPRED | 1 | ALLERGEN | Non-Toxin | -0.25 | -0.37 | -0.02 | 3 | 11.72 | 2209.78 | outside |
| 786 | 806 | KQIYKTPPIKDFGGFNFSQI | 0.1735 | 20 | ABCPRED | 1 | NON-ALLERGEN | Non-Toxin | -0.11 | -0.53 | -0.13 | 2 | 9.55 | 2329 | outside |
| 707 | 721 | YSNNSIAIPTNFTI | 0.6764 | 14 | ABCPRED | 1 | NON-ALLERGEN | Non-Toxin | 0.01 | 0.12 | -0.74 | 0 | 5.88 | 1554.92 | outside |
| 708 | 730 | SNNSIAIPTNFTISVTTEILPV | 0.7747 | 22 | AAP | 1 | NON-ALLERGEN | Non-Toxin | 0.06 | 0.56 | -0.55 | -1 | 4 | 2331.98 | outside |
| 871 | 891 | AQYTSALLAGTITSGWTFGA | 0.46 | 20 | ABCPRED | 1 | ALLERGEN | Non-Toxin | 0.12 | 0.54 | -0.82 | 0 | 5.88 | 2016.53 | outside |
| 169 | 183 | EYVSQPFLMDLEGK | 0.5925 | 14 | ABCPRED | 1 | NON-ALLERGEN | Non-Toxin | -0.1 | -0.39 | 0.09 | -2 | 4.14 | 1656.09 | outside |
| 679 | 693 | NSPRRARSVASQSI | 0.0895 | 14 | ABCPRED | 1 | NON-ALLERGEN | Non-Toxin | -0.42 | -0.93 | 0.45 | 3 | 12.31 | 1528.87 | outside |
| 786 | 800 | KQIYKTPPIKDFGG | -0.4785 | 14 | ABCPRED | 1 | NON-ALLERGEN | Non-Toxin | -0.19 | -0.92 | 0.24 | 2 | 9.55 | 1592.08 | outside |
| 708 | 728 | SNNSIAIPTNFTISVTTEIL | 0.5764 | 20 | ABCPRED | 1 | NON-ALLERGEN | Non-Toxin | 0.04 | 0.49 | -0.53 | -1 | 4 | 2135.7 | outside |
| 791 | 807 | TPPIKDFGGFNFSQIL | 0.2437 | 16 | ABCPRED | 1 | NON-ALLERGEN | Toxin | 0.03 | 0.08 | -0.41 | 0 | 6.19 | 1781.28 | outside |
| 130 | 144 | VCEFQFCNDPFLGV | -0.0637 | 14 | ABCPRED | 1 | ALLERGEN | Non-Toxin | 0.07 | 0.69 | -0.56 | -2 | 3.67 | 1618.05 | outside |
| 365 | 379 | YSVLYNSASFSTFK | 0.0673 | 14 | ABCPRED | 1 | NON-ALLERGEN | Non-Toxin | -0.03 | 0.11 | -0.67 | 1 | 8.83 | 1613.97 | outside |
| 1084 | 1098 | DGKAHFPREGVFVS | 0.0556 | 14 | ABCPRED | 1 | NON-ALLERGEN | Non-Toxin | -0.15 | -0.43 | 0.24 | 0.5 | 7.1 | 1545.91 | outside |
| 351 | 365 | YAWNRKRISNCVAD | 0.2786 | 14 | ABCPRED | 1 | NON-ALLERGEN | Non-Toxin | -0.33 | -0.83 | 0.12 | 2 | 9.36 | 1696.1 | outside |
| 1078 | 1092 | APAICHDGKAHFPR | -0.5688 | 14 | ABCPRED | 1 | NON-ALLERGEN | Non-Toxin | -0.16 | -0.48 | 0.09 | 2 | 8.57 | 1519.93 | outside |
| 436 | 450 | WNSNNLDSKVGGNY | 0.7148 | 14 | ABCPRED | 1 | NON-ALLERGEN | Non-Toxin | -0.22 | -1.29 | -0.11 | 0 | 6.18 | 1567.84 | outside |
| 927 | 941 | FNSAIGKIQDSLSS | 0.1795 | 14 | ABCPRED | 1 | NON-ALLERGEN | Non-Toxin | -0.08 | -0.04 | -0.06 | 0 | 6.19 | 1466.81 | outside |
| 857 | 877 | GLTVLPPLLTDEMIAQYTSA | 0.3868 | 20 | ABCPRED | 1 | ALLERGEN | Non-Toxin | 0.07 | 0.56 | -0.49 | -2 | 3.67 | 2133.79 | outside |
| 469 | 483 | STEIYQAGSTPCNG | -0.1578 | 14 | ABCPRED | 1 | NON-ALLERGEN | Non-Toxin | -0.11 | -0.6 | -0.17 | -1 | 4 | 1427.7 | outside |
| 72 | 86 | GTNGTKRFDNPVLP | 0.448 | 14 | ABCPRED | 1 | NON-ALLERGEN | Non-Toxin | -0.24 | -0.96 | 0.2 | 1 | 9.1 | 1515.89 | outside |
| 273 | 287 | RTFLLKYNENGTIT | 0.1914 | 14 | ABCPRED | 1 | NON-ALLERGEN | Non-Toxin | -0.19 | -0.56 | -0.14 | 1 | 8.93 | 1670.11 | outside |
| 733 | 753 | KTSVDCTMYICGDSTECSNL | 0.1987 | 20 | ABCPRED | 1 | NON-ALLERGEN | Non-Toxin | -0.14 | -0.11 | 0.01 | -2 | 4.03 | 2170.7 | outside |
| 1175 | 1195 | SVVNIQKEIDRLNEVAKNLN | 0.3536 | 20 | ABCPRED | 1 | NON-ALLERGEN | Non-Toxin | -0.25 | -0.51 | 0.35 | 0 | 6.53 | 2296.92 | outside |
| 338 | 352 | FGEVFNATRFASVY | 0.0181 | 14 | ABCPRED | 1 | NON-ALLERGEN | Non-Toxin | 0.01 | 0.41 | -0.55 | 0 | 6.35 | 1607.97 | outside |
| 963 | 977 | VKQLSSNFGAISSV | 0.5947 | 14 | ABCPRED | 1 | NON-ALLERGEN | Non-Toxin | -0.01 | 0.49 | -0.36 | 1 | 9.11 | 1436.83 | outside |
| 734 | 754 | TSVDCTMYICGDSTECSNLL | -0.0168 | 20 | ABCPRED | 1 | NON-ALLERGEN | Non-Toxin | -0.06 | 0.28 | -0.23 | -3 | 3.5 | 2155.69 | outside |
| 1198 | 1220 | IDLQELGKYEQYIKWPWYIWLG | 0.9637 | 22 | BCPREDS | 1 | ALLERGEN | Non-Toxin | -0.02 | -0.43 | -0.57 | -1 | 4.68 | 2842.65 | outside_transmembrane-region |
| 1100 | 1114 | THWFVTQRNFYEPQ | 0.3664 | 14 | ABCPRED | 1 | NON-ALLERGEN | Non-Toxin | -0.22 | -1.22 | -0.49 | 0.5 | 7.1 | 1853.23 | outside |
| 565 | 585 | FGRDIADTTDAVRDPQTLEI | 0.359 | 20 | ABCPRED | 1 | NON-ALLERGEN | Non-Toxin | -0.23 | -0.53 | 0.48 | -3 | 4.04 | 2233.7 | outside |
| 186 | 210 | FKNLREFVFKNIDGYFKIYSKHTP | 0.1429 | 24 | literature | 1 | NON-ALLERGEN | Non-Toxin | -0.19 | -0.63 | -0.03 | 3.5 | 9.71 | 2992.83 | outside |
| 696 | 712 | TMSLGAENSVAYSNNS | 0.6687 | 16 | ABCPRED,FBCPred | 2 | ALLERGEN | Non-Toxin | -0.11 | -0.38 | -0.22 | -1 | 4 | 1644.95 | outside |
| 458 | 478 | KSNLKPFERDISTEIYQAGS | 0.2212 | 20 | ABCPRED | 1 | NON-ALLERGEN | Non-Toxin | -0.26 | -0.94 | 0.41 | 0 | 6.52 | 2283.81 | outside |
| 522 | 542 | ATVCGPKKSTNLVKNKCVNF | 0.5546 | 20 | ABCPRED | 1 | NON-ALLERGEN | Non-Toxin | -0.19 | -0.21 | 0.04 | 4 | 9.72 | 2151.86 | outside |
| 895 | 909 | QIPFAMQMAYRFNG | 0.7789 | 14 | ABCPRED | 1 | ALLERGEN | Non-Toxin | -0.05 | -0.06 | -0.65 | 1 | 9.1 | 1674.17 | outside |
| 1228 | 1248 | VMVTIMLCCMTSCCSCLKGC | -0.1297 | 20 | ABCPRED | 1 | NON-ALLERGEN | Non-Toxin | 0.1 | 1.69 | -0.78 | 1 | 8.04 | 2130.05 | transmembrane-region_inside |
| 1047 | 1067 | YHLMSFPQSAPHGVVFLHVT | 0.4275 | 20 | ABCPRED | 1 | ALLERGEN | Non-Toxin | 0.09 | 0.46 | -0.91 | 1.5 | 7.35 | 2267.94 | outside |
| 426 | 442 | PDDFTGCVIAWNSNNL | 0.1077 | 16 | FBCPred | 1 | NON-ALLERGEN | Non-Toxin | -0.04 | -0.14 | -0.37 | -2 | 3.57 | 1766.14 | outside |
| 1240 | 1256 | CCSCLKGCCSCGSCCK | -0.0187 | 16 | ABCPRED | 1 | NON-ALLERGEN | Non-Toxin | -0.11 | 0.8 | -0.18 | 2 | 8.33 | 1588.17 | inside |
| 71 | 91 | SGTNGTKRFDNPVLPFNDGV | 0.3666 | 20 | ABCPRED | 1 | NON-ALLERGEN | Non-Toxin | -0.18 | -0.74 | 0.12 | 0 | 6.31 | 2135.61 | outside |
| 563 | 577 | QQFGRDIADTTDAV | 0.1325 | 14 | ABCPRED | 1 | ALLERGEN | Non-Toxin | -0.22 | -0.62 | 0.34 | -2 | 3.94 | 1536.81 | outside |
| 1065 | 1081 | VTYVPAQEKNFTTAPA | 0.6987 | 16 | AAP,BCPREDS,FBCPred | 3 | NON-ALLERGEN | Non-Toxin | -0.08 | -0.27 | -0.26 | 0 | 6.35 | 1737.16 | outside |
| 281 | 295 | ENGTITDAVDCALD | 0.492 | 14 | ABCPRED | 1 | NON-ALLERGEN | Non-Toxin | -0.09 | -0.05 | 0.31 | -4 | 3.38 | 1436.7 | outside |
| 210 | 230 | INLVRDLPQGFSALEPLVDL | 0.4501 | 20 | ABCPRED | 1 | NON-ALLERGEN | Non-Toxin | 0 | 0.51 | -0.21 | -2 | 4.03 | 2209.88 | outside |
| 632 | 646 | TWRVYSTGSNVFQT | 0.1548 | 14 | ABCPRED | 1 | NON-ALLERGEN | Non-Toxin | -0.14 | -0.47 | -0.6 | 1 | 9.1 | 1645.99 | outside |
| 978 | 992 | NDILSRLDKVEAEV | -0.0165 | 14 | ABCPRED | 1 | NON-ALLERGEN | Non-Toxin | -0.24 | -0.31 | 0.69 | -2 | 4.32 | 1600.99 | outside |
| 513 | 527 | LSFELLHAPATVCG | 0.6842 | 14 | ABCPRED | 1 | ALLERGEN | Non-Toxin | 0.14 | 1.02 | -0.64 | -0.5 | 5.25 | 1457.91 | outside |
| 498 | 518 | QPTNGVGYQPYRVVVLSFEL | 0.7659 | 20 | ABCPRED | 1 | ALLERGEN | Non-Toxin | -0.04 | 0.03 | -0.51 | 0 | 6.35 | 2266.89 | outside |
| 898 | 918 | FAMQMAYRFNGIGVTQNVLY | 0.9281 | 20 | ABCPRED | 1 | NON-ALLERGEN | Non-Toxin | 0.02 | 0.35 | -0.82 | 1 | 8.93 | 2324.02 | outside |
| 1131 | 1151 | GIVNNTVYDPLQPELDSFKE | 0.1033 | 20 | ABCPRED | 1 | NON-ALLERGEN | Non-Toxin | -0.13 | -0.57 | 0.12 | -3 | 3.92 | 2278.8 | outside |
| 547 | 569 | TGTGVLTESNKKFLPFQQFGRD | 0.5666 | 22 | AAP | 1 | ALLERGEN | Non-Toxin | -0.19 | -0.7 | 0.1 | 1 | 8.93 | 2471.1 | outside |
| 164 | 184 | NNCTFEYVSQPFLMDLEGKQ | 0.4715 | 20 | ABCPRED | 1 | NON-ALLERGEN | Toxin | -0.15 | -0.57 | -0.1 | -2 | 4.14 | 2363.94 | outside |
| 322 | 338 | PTESIVRFPNITNLCP | 0.4947 | 16 | ABCPRED,BCPREDS,FBCPred | 3 | ALLERGEN | Non-Toxin | -0.08 | 0.02 | -0.28 | 0 | 6.32 | 1801.32 | outside |
| 572 | 592 | TTDAVRDPQTLEILDITPCS | 0.804 | 20 | ABCPRED | 1 | NON-ALLERGEN | Non-Toxin | -0.15 | -0.18 | 0.18 | -3 | 3.84 | 2188.72 | outside |
| 372 | 392 | ASFSTFKCYGVSPTKLNDLC | 1.227 | 20 | ABCPRED | 1 | ALLERGEN | Non-Toxin | -0.07 | 0.11 | -0.28 | 1 | 8.34 | 2181.78 | outside |
| 687 | 701 | VASQSIIAYTMSLG | 0.5256 | 14 | ABCPRED | 1 | NON-ALLERGEN | Non-Toxin | 0.13 | 1.01 | -0.77 | 0 | 5.88 | 1440.88 | outside |
| 207 | 229 | HTPINLVRDLPQGFSALEPLVD | 0.3666 | 22 | FBCPred | 1 | ALLERGEN | Non-Toxin | -0.06 | 0.04 | -0.15 | -1.5 | 4.54 | 2432.11 | outside |
| 40 | 60 | DKVFRSSVLHSTQDLFLPFF | -0.0649 | 20 | ABCPRED | 1 | ALLERGEN | Non-Toxin | -0.07 | 0.21 | -0.31 | 0.5 | 7.1 | 2384.02 | outside |
| 772 | 788 | VEQDKNTQEVFAQVKQ | 0.2983 | 16 | AAP | 1 | NON-ALLERGEN | Non-Toxin | -0.33 | -1.21 | 0.51 | -1 | 4.68 | 1891.31 | outside |
| 1236 | 1265 | KKKKCCSCLKGCCSCGSCCKFDEDDSEPVL | 0.4611 | 30 | literature | 1 | ALLERGEN | Non-Toxin | -0.28 | -0.4 | 0.62 | 1 | 7.92 | 3248.23 | transmembrane-region_inside |

| | | | | | | | | | | | | | | | |
|---|---|---|---|---|---|---|---|---|---|---|---|---|---|---|---|
| 222 | 242 | ALEPLVDLPIGINITRFQTL | 0.6921 | 20 | ABCPRED | 1 | NON-ALLERGEN | Non-Toxin | 0.05 | 0.7 | -0.42 | -1 | 4.38 | 2223.96 | outside |
| 32 | 52 | FTRGVYYPDKVFRSSVLHST | 0.1739 | 20 | ABCPRED | 1 | NON-ALLERGEN | Non-Toxin | -0.17 | -0.3 | -0.22 | 2.5 | 9.72 | 2359.95 | outside |
| 53 | 67 | DLFLPFFSNVTWFH | 0.3301 | 14 | ABCPRED | 1 | NON-ALLERGEN | Non-Toxin | 0.15 | 0.63 | -1.14 | -0.5 | 5.09 | 1770.22 | outside |
| 947 | 961 | KLQDVVNQNAQALN | 0.6585 | 14 | ABCPRED | 1 | NON-ALLERGEN | Non-Toxin | -0.23 | -0.63 | -0.03 | 0 | 6.19 | 1554.94 | outside |
| 653 | 669 | AEHVNNSYECDIPIGA | 0.829 | 16 | ABCPRED,BCPREDS | 2 | NON-ALLERGEN | Non-Toxin | -0.08 | -0.34 | -0.01 | -2.5 | 4.14 | 1732.08 | outside |
| 255 | 269 | SSGWTAGAAAYYVG | 0.3269 | 14 | ABCPRED | 1 | NON-ALLERGEN | Non-Toxin | 0.12 | 0.31 | -0.81 | 0 | 5.87 | 1360.63 | outside |
| 680 | 694 | SPRRARSVASQSII | 0.2593 | 14 | ABCPRED | 1 | NON-ALLERGEN | Non-Toxin | -0.33 | -0.36 | 0.31 | 3 | 12.31 | 1527.93 | outside |
| 184 | 204 | GNFKNLREFVFKNIDGYFKI | 0.0264 | 20 | ABCPRED | 1 | NON-ALLERGEN | Non-Toxin | -0.15 | -0.38 | -0.03 | 2 | 9.55 | 2450.14 | outside |
| 19 | 47 | TTRTQLPPAYTNSFTRGVYYPDKVFRSS | 0.2845 | 28 | literature | 1 | NON-ALLERGEN | Non-Toxin | -0.25 | -0.83 | -0.1 | 3 | 10 | 3254.01 | outside |
| 619 | 633 | EVPVAIHADQLTPT | 0.3123 | 14 | ABCPRED | 1 | NON-ALLERGEN | Non-Toxin | -0.01 | 0.14 | -0.19 | -1.5 | 4.36 | 1490.88 | outside |
| 4 | 24 | FLVLLPLVSSQCVNLTTRTQ | 1.0702 | 20 | ABCPRED | 1 | NON-ALLERGEN | Non-Toxin | 0 | 0.83 | -0.7 | 1 | 8.6 | 2232.98 | outside |
| 715 | 731 | PTNFTISVTTEILPVS | 1.1529 | 16 | FBCPred | 1 | NON-ALLERGEN | Non-Toxin | 0.07 | 0.59 | -0.54 | -1 | 4 | 1719.2 | outside |
| 864 | 878 | LLTDEMIAQYTSAL | -0.0502 | 14 | ABCPRED | 1 | NON-ALLERGEN | Non-Toxin | 0.03 | 0.53 | -0.44 | -2 | 3.67 | 1569.01 | outside |
| 389 | 403 | DLCFTNVYADSFVI | 0.6352 | 14 | ABCPRED | 1 | NON-ALLERGEN | Non-Toxin | 0.1 | 0.95 | -0.66 | -2 | 3.57 | 1607 | outside |
| 805 | 819 | ILPDPSKPSKRSFI | 0.2767 | 14 | ABCPRED | 1 | NON-ALLERGEN | Non-Toxin | -0.22 | -0.53 | 0.36 | 2 | 10.01 | 1585.07 | outside |
| 797 | 813 | FGGFNFSQILPDPSKP | 0.3112 | 16 | AAP | 1 | ALLERGEN | Toxin | -0.03 | -0.31 | -0.26 | 0 | 6.19 | 1751.2 | outside |
| 242 | 264 | LALHRSYLTPGDSSSGWTAGAA | 0.4485 | 22 | BCPREDS | 1 | NON-ALLERGEN | Non-Toxin | -0.04 | -0.1 | -0.33 | 0.5 | 7.09 | 2218.72 | outside |
| 481 | 501 | NGVEGFNCYFPLQSYGFQPT | 0.7422 | 20 | ABCPRED | 1 | ALLERGEN | Non-Toxin | -0.02 | -0.36 | -0.64 | -1 | 4 | 2268.78 | outside |
| 20 | 34 | TRTQLPPAYTNSFT | 0.5883 | 14 | ABCPRED | 1 | ALLERGEN | Non-Toxin | -0.2 | -0.8 | -0.36 | 1 | 9.1 | 1596.95 | outside |
| 727 | 747 | LPVSMTKTSVDCTMYICGDS | 0.3164 | 20 | ABCPRED | 1 | ALLERGEN | Non-Toxin | -0.04 | 0.34 | -0.24 | -1 | 4.21 | 2151.79 | outside |
| 506 | 526 | QPYRVVVLSFELLHAPATVC | 0.719 | 20 | ABCPRED | 1 | NON-ALLERGEN | Non-Toxin | 0.04 | 0.82 | -0.63 | 0.5 | 7.06 | 2242.96 | outside |
| 16 | 32 | VNLTTRTQLPPAYTNS | 1.1248 | 16 | ABCPRED,AAP | 2 | NON-ALLERGEN | Non-Toxin | -0.19 | -0.59 | -0.35 | 1 | 9.1 | 1776.21 | outside |
| 246 | 268 | RSYLTPGDSSSGWTAGAAAYYV | 0.4413 | 22 | AAP | 1 | ALLERGEN | Non-Toxin | -0.04 | -0.23 | -0.42 | 0 | 6.18 | 2280.73 | outside |
| 894 | 910 | LQIPFAMQMAYRFNGI | 0.8899 | 16 | AAP | 1 | NON-ALLERGEN | Non-Toxin | 0.03 | 0.47 | -0.79 | 1 | 9.1 | 1900.53 | outside |
| 384 | 404 | PTKLNDLCFTNVYADSFVIR | 0.9772 | 20 | ABCPRED | 1 | NON-ALLERGEN | Non-Toxin | -0.09 | 0.15 | -0.27 | 0 | 6.28 | 2316.94 | outside |
| 151 | 167 | SWMESEFRVYSSANNC | 0.1724 | 16 | ABCPRED | 1 | NON-ALLERGEN | Toxin | -0.2 | -0.67 | -0.12 | -1 | 4.54 | 1910.27 | outside |
| 1081 | 1095 | ICHDGKAHFPREGV | -0.4185 | 14 | ABCPRED | 1 | NON-ALLERGEN | Non-Toxin | -0.18 | -0.6 | 0.26 | 1 | 7.22 | 1565.97 | outside |
| 1231 | 1247 | TIMLCCMTSCCSCLKG | -0.1119 | 16 | FBCPred | 1 | NON-ALLERGEN | Non-Toxin | 0.04 | 1.32 | -0.64 | 1 | 8.1 | 1697.39 | transmembrane-region_inside |
| 534 | 548 | VKNKCVNFNFNGLT | 1.5749 | 14 | ABCPRED | 1 | NON-ALLERGEN | Non-Toxin | -0.14 | -0.19 | -0.31 | 2 | 9.36 | 1598.06 | outside |
| 677 | 691 | QTNSPRRARSVASQ | 0.0709 | 14 | ABCPRED | 1 | NON-ALLERGEN | Non-Toxin | -0.52 | -1.49 | 0.54 | 3 | 12.31 | 1557.87 | outside |
| 174 | 188 | PFLMDLEGKQGNFK | 1.1174 | 14 | ABCPRED | 1 | NON-ALLERGEN | Non-Toxin | -0.15 | -0.65 | 0.18 | 0 | 6.42 | 1624.1 | outside |
| 64 | 86 | WFHAIHVSGTNGTKRFDNPVLP | 0.4415 | 22 | BCPREDS | 1 | NON-ALLERGEN | Non-Toxin | -0.09 | -0.38 | -0.35 | 2 | 9.11 | 2494.14 | outside |
| 167 | 181 | TFEYVSQPFLMDLE | 0.7984 | 14 | ABCPRED | 1 | NON-ALLERGEN | Non-Toxin | 0 | 0.06 | -0.33 | -3 | 3.58 | 1719.14 | outside |
| 494 | 524 | SYGFQPTNGVGYQPYRVVVLSFELLHAPAT | 0.8107 | 30 | literature | 1 | NON-ALLERGEN | Non-Toxin | 0 | 0.09 | -0.61 | 0.5 | 7.09 | 3312.2 | outside |
| 130 | 150 | VCEFQFCNDPFLGVYYHKNN | 0.2084 | 20 | ABCPRED | 1 | ALLERGEN | Toxin | -0.09 | -0.36 | -0.48 | -0.5 | 5.33 | 2438.02 | outside |
| 660 | 674 | YECDIPIGAGICAS | 0.5804 | 14 | ABCPRED | 1 | ALLERGEN | Non-Toxin | 0.1 | 0.76 | -0.31 | -2 | 3.67 | 1411.8 | outside |
| 705 | 719 | VAYSNNSIAIPTNF | 0.5729 | 14 | ABCPRED | 1 | NON-ALLERGEN | Non-Toxin | 0.03 | 0.28 | -0.72 | 0 | 5.88 | 1510.86 | outside |
| 608 | 628 | VAVLYQDVNCTEVPVAIHAD | 0.6258 | 20 | ABCPRED | 1 | NON-ALLERGEN | Non-Toxin | 0.04 | 0.65 | -0.37 | -2.5 | 4.03 | 2156.73 | outside |
| 1035 | 1055 | GQSKRVDFCGKGYHLMSFPQ | 0.7776 | 20 | ABCPRED | 1 | NON-ALLERGEN | Non-Toxin | -0.2 | -0.69 | -0.02 | 2.5 | 9.25 | 2285.92 | outside |
| 397 | 417 | ADSFVIRGDEVRQIAPGQTG | 0.5457 | 20 | ABCPRED | 1 | NON-ALLERGEN | Non-Toxin | -0.17 | -0.35 | 0.26 | -1 | 4.56 | 2116.61 | outside |
| 967 | 987 | SSNFGAISSVLNDILSRLDK | 0.1255 | 20 | ABCPRED | 1 | ALLERGEN | Non-Toxin | -0.11 | 0.12 | 0.02 | 0 | 6.31 | 2136.68 | outside |
| 435 | 451 | AWNSNNLDSKVGGNYN | 0.7971 | 16 | AAP | 1 | NON-ALLERGEN | Non-Toxin | -0.22 | -1.23 | -0.12 | 0 | 6.18 | 1753.05 | outside |
| 600 | 622 | PGTNTSNQVAVLYQDVNCTEVP | 0.6405 | 22 | AAP | 1 | NON-ALLERGEN | Non-Toxin | -0.11 | -0.34 | -0.25 | -2 | 3.67 | 2349.88 | outside |
| 528 | 542 | KKSTNLVKNKCVNF | 1.0894 | 14 | ABCPRED | 1 | NON-ALLERGEN | Non-Toxin | -0.32 | -0.72 | 0.3 | 4 | 10.05 | 1623.15 | outside |
| 588 | 602 | TPCSFGGVSVITPG | 0.9486 | 14 | ABCPRED | 1 | NON-ALLERGEN | Non-Toxin | 0.14 | 0.77 | -0.61 | 0 | 5.85 | 1321.71 | outside |
| 754 | 768 | LQYGSFCTQLNRAL | 1.0734 | 14 | ABCPRED | 1 | NON-ALLERGEN | Non-Toxin | -0.11 | 0.02 | -0.59 | 1 | 8.57 | 1614.06 | outside |
| 214 | 228 | RDLPQGFSALEPLV | 0.7136 | 14 | ABCPRED | 1 | NON-ALLERGEN | Non-Toxin | -0.07 | 0.06 | -0.03 | -1 | 4.38 | 1541.97 | outside |
| 844 | 858 | IAARDLICAQKFNG | 0.3468 | 14 | ABCPRED | 1 | NON-ALLERGEN | Non-Toxin | -0.1 | 0.3 | -0.07 | 1 | 8.57 | 1519.98 | outside |
| 524 | 544 | VCGPKKSTNLVKNKCVNFNF | 0.8346 | 20 | ABCPRED | 1 | NON-ALLERGEN | Non-Toxin | -0.19 | -0.31 | -0.03 | 4 | 9.72 | 2240.96 | outside |
| 275 | 289 | FLLKYNENGTITDA | 0.6641 | 14 | ABCPRED | 1 | NON-ALLERGEN | Non-Toxin | -0.09 | -0.31 | -0.15 | -1 | 4.38 | 1598.98 | outside |
| 283 | 303 | GTITDAVDCALDPLSETKCT | 0.7528 | 20 | ABCPRED | 1 | NON-ALLERGEN | Non-Toxin | -0.09 | 0.07 | 0.19 | -3 | 3.84 | 2053.56 | outside |
| 1237 | 1257 | MTSCCSCLKGCCSCGSCCKF | -0.0422 | 20 | ABCPRED | 1 | ALLERGEN | Toxin | -0.07 | 0.8 | -0.34 | 2 | 8.33 | 2054.78 | inside |
| 228 | 248 | DLPIGINITRFQTLLALHRS | 0.864 | 20 | ABCPRED | 1 | NON-ALLERGEN | Non-Toxin | -0.07 | 0.32 | -0.36 | 1.5 | 9.95 | 2279 | outside |
| 570 | 584 | ADTTDAVRDPQTLE | 0.4052 | 14 | ABCPRED | 1 | ALLERGEN | Non-Toxin | -0.3 | -1.01 | 0.69 | -3 | 3.84 | 1531.78 | outside |
| 483 | 503 | VEGFNCYFPLQSYGFQPTNG | 0.5561 | 20 | ABCPRED | 1 | ALLERGEN | Non-Toxin | -0.02 | -0.36 | -0.64 | -1 | 4 | 2268.78 | outside |
| 436 | 460 | WNSNNLDSYNYLYRN | 0.2089 | 15 | ellipro | 1 | NON-ALLERGEN | Non-Toxin | -0.31 | -1.62 | -0.42 | 0 | 6.18 | 1936.24 | outside |
| 19 | 41 | TTRTQLPPAYTNSFTRGVYYPD | 0.4709 | 22 | BCPREDS,FBCPred | 2 | NON-ALLERGEN | Non-Toxin | -0.22 | -0.92 | -0.25 | 1 | 8.83 | 2549.1 | outside |
| 291 | 305 | CALDPLSETKCTLK | 0.7406 | 14 | ABCPRED | 1 | NON-ALLERGEN | Non-Toxin | -0.17 | -0.03 | 0.26 | 0 | 6.36 | 1522 | outside |
| 1115 | 1135 | ITTDNTFVSGNCDVVIGIVN | 0.508 | 20 | ABCPRED | 1 | ALLERGEN | Non-Toxin | 0.06 | 0.72 | -0.46 | -2 | 3.57 | 2081.63 | outside |
| 972 | 986 | AISSVLNDILSRLD | -0.1005 | 14 | ABCPRED | 1 | NON-ALLERGEN | Non-Toxin | -0.06 | 0.64 | -0.06 | -1 | 4.21 | 1515.93 | outside |
| 695 | 709 | YTMSLGAENSVAYS | 0.633 | 14 | ABCPRED | 1 | NON-ALLERGEN | Non-Toxin | -0.01 | 0.03 | -0.46 | -1 | 4 | 1492.81 | outside |
| 194 | 214 | FKNIDGYFKIYSKHTPINLV | 0.6171 | 20 | ABCPRED | 1 | ALLERGEN | Non-Toxin | -0.08 | -0.22 | -0.33 | 2.5 | 9.43 | 2398.11 | outside |
| 983 | 997 | RLDKVEAEVQIDRL | -0.0567 | 14 | ABCPRED | 1 | NON-ALLERGEN | Non-Toxin | -0.35 | -0.58 | 0.88 | -1 | 4.79 | 1684.13 | outside |
| 702 | 718 | ENSVAYSNNSIAIPTN | 0.3501 | 16 | BCPREDS | 1 | ALLERGEN | Non-Toxin | -0.11 | -0.42 | -0.26 | -1 | 4 | 1694.01 | outside |
| 539 | 553 | VNFNFNGLTGTGVL | 1.2915 | 14 | ABCPRED | 1 | NON-ALLERGEN | Non-Toxin | 0.11 | 0.61 | -0.84 | 0 | 5.88 | 1452.85 | outside |
| 281 | 301 | ENGTITDAVDCALDPLSETK | 0.7029 | 20 | ABCPRED | 1 | ALLERGEN | Non-Toxin | -0.15 | -0.37 | 0.42 | -4 | 3.77 | 2092.54 | outside |
| 21 | 43 | RTQLPPAYTNSFTRGVYYPDKV | 0.4077 | 22 | AAP | 1 | NON-ALLERGEN | Non-Toxin | -0.23 | -0.85 | -0.15 | 2 | 9.55 | 2574.2 | outside |
| 645 | 665 | TRAGCLIGAEHVNNSYECDI | 1.0613 | 20 | ABCPRED | 1 | ALLERGEN | Non-Toxin | -0.12 | -0.16 | -0.02 | -1.5 | 4.66 | 2165.67 | outside |
| 352 | 366 | AWNRKRISNCVADY | 0.2831 | 14 | ABCPRED | 1 | NON-ALLERGEN | Non-Toxin | -0.33 | -0.83 | 0.12 | 2 | 9.36 | 1696.1 | outside |
| 514 | 534 | SFELLHAPATVCGPKKSTNL | 0.404 | 20 | ABCPRED | 1 | NON-ALLERGEN | Non-Toxin | -0.07 | -0.01 | -0.15 | 1.5 | 8.54 | 2113.75 | outside |
| 527 | 547 | PKKSTNLVKNKCVNFNFNGL | 1.0582 | 20 | ABCPRED | 1 | ALLERGEN | Non-Toxin | -0.23 | -0.62 | 0.02 | 4 | 10.05 | 2265.96 | outside |
| 631 | 645 | PTWRVYSTGSNVFQ | 0.1826 | 14 | ABCPRED | 1 | ALLERGEN | Non-Toxin | -0.13 | -0.54 | -0.57 | 1 | 9.1 | 1642 | outside |
| 201 | 270 | SKHTPPQGFSALEPLVDLPIGINITRFQTLLAAYY | 0.5624 | 43 | ellipro | 1 | NON-ALLERGEN | Non-Toxin | 0.03 | 0.31 | -0.57 | 1.5 | 8.66 | 4857.36 | outside |
| 408 | 422 | RQIAPGQTGKIADY | 1.504 | 14 | ABCPRED | 1 | NON-ALLERGEN | Non-Toxin | -0.21 | -0.76 | 0.15 | 1 | 8.93 | 1517.91 | outside |
| 364 | 378 | DYSVLYNSASFSTF | 0.225 | 14 | ABCPRED | 1 | ALLERGEN | Non-Toxin | 0 | 0.14 | -0.67 | -1 | 3.8 | 1600.88 | outside |
| 1211 | 1227 | KWPWYIWLGFIAGLIA | 0.7111 | 16 | ABCPRED | 1 | NON-ALLERGEN | Non-Toxin | 0.29 | 1.07 | -1.38 | 1 | 8.94 | 1934.61 | outside_transmembrane-region |
| 277 | 291 | LKYNENGTITDAVD | 0.6707 | 14 | ABCPRED | 1 | NON-ALLERGEN | Non-Toxin | -0.18 | -0.73 | 0.26 | -2 | 4.03 | 1552.86 | outside |

| | | | | | | | | | | | | | | | |
|---|---|---|---|---|---|---|---|---|---|---|---|---|---|---|---|
| 198 | 270 | PINLVRDLPQGFSALEPLVDLPIGINITRFQTLLALI | 0.5344 | 58 | ellipro | 1 | NON-ALLERGEN | too long | | | | | | | outside |
| 851 | 871 | CAQKFNGLTVLPPLLTDEMI | 0.046 | 20 | ABCPRED | 1 | NON-ALLERGEN | Non-Toxin | 0.02 | 0.5 | -0.36 | -1 | 4.38 | 2203.95 | outside |
| 943 | 957 | SALGKLQDVVNQNA | 0.3999 | 14 | ABCPRED | 1 | NON-ALLERGEN | Non-Toxin | -0.14 | -0.21 | -0.04 | 0 | 6.19 | 1456.83 | outside |
| 372 | 386 | ASFSTFKCYGVSPT | 0.8761 | 14 | ABCPRED | 1 | NON-ALLERGEN | Non-Toxin | -0.01 | 0.22 | -0.51 | 1 | 8.54 | 1494.86 | outside |
| 736 | 750 | VDCTMYICGDSTEC | -0.2858 | 14 | ABCPRED | 1 | NON-ALLERGEN | Non-Toxin | -0.06 | 0.26 | -0.1 | -3 | 3.5 | 1539.91 | outside |
| 310 | 326 | KGIYQTSNFRVQPTES | 0.7664 | 16 | ABCPRED | 1 | NON-ALLERGEN | Non-Toxin | -0.27 | -1.07 | 0.08 | 1 | 8.93 | 1855.27 | outside |
| 1108 | 1130 | NFYEPQIITTDNTFVSGNCDVV | 0.2307 | 22 | FBCPred | 1 | ALLERGEN | Non-Toxin | -0.05 | -0.05 | -0.34 | -3 | 3.5 | 2477.02 | outside |
| 373 | 387 | SFSTFKCYGVSPTK | 0.9678 | 14 | ABCPRED | 1 | NON-ALLERGEN | Non-Toxin | -0.1 | -0.19 | -0.26 | 2 | 9.25 | 1551.96 | outside |
| 594 | 610 | GVSVITPGTNTSNQVA | 0.4651 | 16 | ABCPRED | 1 | NON-ALLERGEN | Non-Toxin | -0.01 | 0.14 | -0.43 | 0 | 5.88 | 1544.92 | outside |
| 332 | 346 | ITNLCPFGEVFNAT | 0.4524 | 14 | ABCPRED | 1 | NON-ALLERGEN | Non-Toxin | 0.08 | 0.61 | -0.64 | -1 | 4 | 1525.94 | outside |
| 444 | 460 | KVGGNYNYLYRLFRKS | -0.0651 | 16 | ABCPRED,FBCPred | 2 | NON-ALLERGEN | Non-Toxin | -0.29 | -0.92 | -0.11 | 4 | 10.17 | 1978.52 | outside |
| 1091 | 1105 | REGVFVSNGTHWFV | 0.3977 | 14 | ABCPRED | 1 | NON-ALLERGEN | Non-Toxin | -0.02 | 0.02 | -0.52 | 0.5 | 7.1 | 1635.02 | outside |
| 163 | 177 | ANNCTFEYVSQPFL | 0.0903 | 14 | ABCPRED | 1 | ALLERGEN | Non-Toxin | -0.04 | -0.04 | -0.61 | -1 | 4 | 1633 | outside |
| 327 | 341 | VRFPNITNLCPFGE | 1.0597 | 14 | ABCPRED | 1 | NON-ALLERGEN | Non-Toxin | -0.05 | 0.09 | -0.36 | 0 | 6.32 | 1607.06 | outside |
| 405 | 433 | DEVRQIAPGQTGKIADYNYKLPDDFTGKKK | 1.0411 | 30 | literature | 1 | NON-ALLERGEN | Non-Toxin | -0.3 | -1.23 | 0.59 | 1 | 8.62 | 3397.25 | outside |
| 525 | 541 | CGPKKSTNLVKNKCVN | 0.2006 | 16 | ABCPRED | 1 | NON-ALLERGEN | Non-Toxin | -0.31 | -0.77 | 0.36 | 4 | 9.72 | 1733.31 | outside |
| 968 | 982 | SNFGAISSVLNDIL | 0.3271 | 14 | ABCPRED | 1 | NON-ALLERGEN | Non-Toxin | 0.09 | 0.86 | -0.53 | -1 | 3.8 | 1449.83 | outside |
| 89 | 103 | GVYFASTEKSNIIR | 0.0505 | 14 | ABCPRED | 1 | NON-ALLERGEN | Non-Toxin | -0.13 | -0.11 | -0.07 | 1 | 8.93 | 1584.99 | outside |
| 227 | 241 | VDLPIGINITRFQT | 0.9138 | 14 | ABCPRED | 1 | NON-ALLERGEN | Non-Toxin | -0.02 | 0.42 | -0.4 | 0 | 6.19 | 1587.07 | outside |
| 807 | 827 | PDPSKPSKRSFIEDLLFNKV | 0.12 | 20 | ABCPRED | 1 | NON-ALLERGEN | Non-Toxin | -0.26 | -0.77 | 0.51 | 1 | 8.83 | 2317.95 | outside |
| 379 | 399 | CYGVSPTKLNDLCFTNVYAD | 1.5222 | 20 | ABCPRED | 1 | NON-ALLERGEN | Non-Toxin | -0.06 | 0.04 | -0.37 | -1 | 4.21 | 2223.78 | outside |
| 15 | 35 | CVNLTTRTQLPPAYTNSFTR | 0.7376 | 20 | ABCPRED | 1 | NON-ALLERGEN | Non-Toxin | -0.21 | -0.47 | -0.33 | 2 | 9.55 | 2283.87 | outside |
| 583 | 597 | EILDITPCSFGGVS | 1.6193 | 14 | ABCPRED | 1 | NON-ALLERGEN | Non-Toxin | 0.1 | 0.76 | -0.3 | -2 | 3.67 | 1437.83 | outside |
| 117 | 131 | LLIVNNATNVVIKV | 0.4113 | 14 | ABCPRED | 1 | NON-ALLERGEN | Non-Toxin | 0.12 | 1.44 | -0.75 | 1 | 9.11 | 1510.08 | outside |
| 111 | 125 | DSKTQSLLIVNNAT | 0.5334 | 14 | ABCPRED | 1 | NON-ALLERGEN | Non-Toxin | -0.15 | -0.2 | -0.07 | 0 | 6.19 | 1503.88 | outside |
| 288 | 302 | AVDCALDPLSETKC | 0.8534 | 14 | ABCPRED | 1 | NON-ALLERGEN | Non-Toxin | -0.11 | 0.21 | 0.27 | -2 | 4.03 | 1464.85 | outside |
| 216 | 230 | LPQGFSALEPLVDL | 0.437 | 14 | ABCPRED | 1 | NON-ALLERGEN | Non-Toxin | 0.09 | 0.65 | -0.37 | -2 | 3.67 | 1498.95 | outside |
| 834 | 850 | IKQYGDCLGDIAARDL | 0.4298 | 16 | AAP | 1 | NON-ALLERGEN | Non-Toxin | -0.14 | -0.11 | 0.23 | -1 | 4.43 | 1751.22 | outside |
| 371 | 385 | SASFSTFKCYGVSP | 0.7798 | 14 | ABCPRED | 1 | NON-ALLERGEN | Non-Toxin | -0.01 | 0.21 | -0.46 | 1 | 8.54 | 1480.83 | outside |
| 112 | 126 | SKTQSLLIVNNATN | 0.558 | 14 | ABCPRED | 1 | ALLERGEN | Non-Toxin | -0.14 | -0.2 | -0.27 | 1 | 9.11 | 1502.9 | outside |
| 1190 | 1217 | AKNLNESLIDLQELGKYEQYIKWPWYIWKK | 0.6619 | 30 | literature | 1 | ALLERGEN | Non-Toxin | -0.17 | -0.88 | -0.07 | 1 | 8.59 | 3798.88 | outside_transmembrane-region |
| 646 | 660 | RAGCLIGAEHVNNS | 1.1927 | 14 | ABCPRED | 1 | NON-ALLERGEN | Non-Toxin | -0.12 | -0.09 | -0.06 | 0.5 | 7.07 | 1440.8 | outside |
| 85 | 99 | PFNDGVYFASTEKS | 0.5515 | 14 | ABCPRED | 1 | NON-ALLERGEN | Non-Toxin | -0.12 | -0.6 | 0.01 | -1 | 4.38 | 1561.85 | outside |
| 380 | 402 | YGVSPTKLNDLCFTNVYADSFV | 1.1404 | 22 | BCPREDS | 1 | ALLERGEN | Non-Toxin | -0.02 | 0.21 | -0.45 | -1 | 4.21 | 2454.06 | outside |
| 771 | 799 | AVEQDKNTQEVFAQVKQIYKTPPIKDFGGK | 0.0782 | 30 | literature | 1 | NON-ALLERGEN | Non-Toxin | -0.23 | -0.9 | 0.36 | 1 | 8.66 | 3408.33 | outside |
| 630 | 646 | TPTWRVYSTGSNVFQT | 0.1876 | 16 | ABCPRED,BCPREDS | 2 | NON-ALLERGEN | Non-Toxin | -0.14 | -0.56 | -0.55 | 1 | 9.1 | 1844.24 | outside |
| 742 | 756 | ICGDSTECSNLLLQ | 0.0093 | 14 | ABCPRED | 1 | NON-ALLERGEN | Toxin | -0.06 | 0.3 | -0.19 | -2 | 3.67 | 1495.89 | outside |
| 20 | 40 | TRTQLPPAYTNSFTRGVYYP | 0.4303 | 20 | ABCPRED | 1 | NON-ALLERGEN | Non-Toxin | -0.19 | -0.81 | -0.4 | 2 | 9.72 | 2332.88 | outside |
| 780 | 794 | EVFAQVKQIYKTPP | 0.1563 | 14 | ABCPRED | 1 | NON-ALLERGEN | Non-Toxin | -0.13 | -0.43 | -0.08 | 1 | 8.83 | 1648.14 | outside |
| 514 | 528 | SFELLHAPATVCGP | 0.261 | 14 | ABCPRED | 1 | NON-ALLERGEN | Non-Toxin | 0.09 | 0.64 | -0.51 | -0.5 | 5.25 | 1441.86 | outside |
| 661 | 681 | ECDIPIGAGICASYQTQTNS | 0.6705 | 20 | ABCPRED | 1 | NON-ALLERGEN | Toxin | -0.06 | -0.11 | -0.22 | -2 | 3.67 | 2071.55 | outside |
| 10 | 30 | LVSSQCVNLTTRTQLPPAYT | 1.1791 | 20 | ABCPRED | 1 | NON-ALLERGEN | Non-Toxin | -0.11 | 0.01 | -0.48 | 1 | 8.57 | 2192.81 | outside |
| 523 | 543 | TVCGPKKSTNLVKNKCVNFN | 0.6333 | 20 | ABCPRED | 1 | NON-ALLERGEN | Non-Toxin | -0.23 | -0.48 | 0.08 | 4 | 9.72 | 2194.89 | outside |
| 464 | 511 | FERDISTEIYNCYFPLQSYGFQPTVGYQPYRVV | 0.3876 | 33 | ellipro,literature | 2 | NON-ALLERGEN | Non-Toxin | -0.1 | -0.37 | -0.43 | -1 | 4.68 | 3981.94 | outside |
| 139 | 159 | PFLGVYYHKNNKSWMESEFR | 0.6801 | 20 | ABCPRED | 1 | NON-ALLERGEN | Non-Toxin | -0.22 | -1.05 | -0.1 | 1.5 | 8.76 | 2533.15 | outside |
| 1008 | 1022 | VTQQLIRAAEIRAS | 0.1575 | 14 | ABCPRED | 1 | NON-ALLERGEN | Non-Toxin | -0.19 | 0.1 | 0.06 | 1 | 9.95 | 1556 | outside |
| 228 | 242 | DLPIGINITRFQTL | 1.1058 | 14 | ABCPRED | 1 | NON-ALLERGEN | Non-Toxin | -0.02 | 0.39 | -0.42 | 0 | 6.19 | 1601.1 | outside |
| 1103 | 1123 | FVTQRNFYEPQIITTDNTFV | 0.504 | 20 | ABCPRED | 1 | ALLERGEN | Non-Toxin | -0.11 | -0.27 | -0.41 | -1 | 4.38 | 2434 | outside |
| 755 | 769 | QYGSFCTQLNRALT | 0.8913 | 14 | ABCPRED | 1 | NON-ALLERGEN | Non-Toxin | -0.16 | -0.3 | -0.49 | 1 | 8.57 | 1602 | outside |
| 652 | 666 | GAEHVNNSYECDIP | 0.6407 | 14 | ABCPRED | 1 | NON-ALLERGEN | Non-Toxin | -0.16 | -0.84 | 0.15 | -2.5 | 4.14 | 1547.81 | outside |
| 360 | 376 | NCVADYSVLYNSASFS | 0.1775 | 16 | ABCPRED,FBCPred | 2 | NON-ALLERGEN | Non-Toxin | -0.01 | 0.3 | -0.58 | -1 | 3.8 | 1740.08 | outside |
| 1083 | 1097 | HDGKAHFPREGVFV | -0.0372 | 14 | ABCPRED | 1 | NON-ALLERGEN | Non-Toxin | -0.16 | -0.6 | 0.18 | 1 | 7.26 | 1595.98 | outside |
| 299 | 313 | TKCTLKSFTVEKGI | 0.2686 | 14 | ABCPRED | 1 | NON-ALLERGEN | Non-Toxin | -0.15 | -0.05 | 0.18 | 2 | 9.25 | 1555.07 | outside |
| 676 | 696 | TQTNSPRRARSVASQSIIAY | 0.1189 | 20 | ABCPRED | 1 | ALLERGEN | Non-Toxin | -0.3 | -0.65 | 0.06 | 3 | 11.72 | 2206.72 | outside |
| 1246 | 1262 | GCCSCGSCCKFDEDDS | 0.0009 | 16 | ABCPRED,BCPREDS | 2 | NON-ALLERGEN | Non-Toxin | -0.22 | -0.36 | 0.53 | -3 | 3.84 | 1658.97 | inside |
| 19 | 44 | TTRTQLPPAYTNSFTRGVYYPDKVF | 0.3642 | 25 | literature | 1 | NON-ALLERGEN | Non-Toxin | -0.19 | -0.69 | -0.26 | 2 | 9.55 | 2923.63 | outside |
| 310 | 324 | KGIYQTSNFRVQPT | 0.962 | 14 | ABCPRED | 1 | NON-ALLERGEN | Non-Toxin | -0.25 | -0.92 | -0.14 | 2 | 10.01 | 1639.05 | outside |
| 1227 | 1247 | IVMVTIMLCCMTSCCSCLKG | 0.13 | 20 | ABCPRED | 1 | NON-ALLERGEN | Non-Toxin | 0.14 | 1.8 | -0.82 | 1 | 8.1 | 2140.08 | transmembrane-region_inside |
| 1230 | 1250 | VTIMLCCMTSCCSCLKGCCS | -0.2622 | 20 | ABCPRED | 1 | NON-ALLERGEN | Non-Toxin | 0.05 | 1.48 | -0.67 | 1 | 7.99 | 2089.93 | transmembrane-region_inside |
| 284 | 298 | TITDAVDCALDPLS | 0.5939 | 14 | ABCPRED | 1 | NON-ALLERGEN | Non-Toxin | 0 | 0.58 | -0.03 | -3 | 3.43 | 1433.78 | outside |
| 475 | 489 | AGSTPCNGVEGFNC | 0.2288 | 14 | ABCPRED | 1 | NON-ALLERGEN | Non-Toxin | -0.03 | -0.07 | -0.23 | -1 | 4 | 1355.65 | outside |
| 616 | 636 | NCTEVPVAIHADQLTPTWRV | 0.4175 | 20 | ABCPRED | 1 | NON-ALLERGEN | Non-Toxin | -0.09 | -0.05 | -0.29 | -0.5 | 5.33 | 2250.85 | outside |
| 500 | 520 | TNGVGYQPYRVVVLSFELLH | 0.8678 | 20 | ABCPRED | 1 | NON-ALLERGEN | Non-Toxin | 0.01 | 0.32 | -0.64 | 0.5 | 7.1 | 2291.95 | outside |
| 894 | 908 | LQIPFAMQMAYRFN | 1.0747 | 14 | ABCPRED | 1 | NON-ALLERGEN | Non-Toxin | -0.02 | 0.24 | -0.78 | 1 | 9.1 | 1730.28 | outside |
| 208 | 228 | TPINLVRDLPQGFSALEPLV | 0.5719 | 20 | ABCPRED | 1 | NON-ALLERGEN | Non-Toxin | -0.01 | 0.38 | -0.29 | -1 | 4.38 | 2179.85 | outside |
| 758 | 778 | SFCTQLNRALTGIAVEQDKN | 0.6095 | 20 | ABCPRED | 1 | ALLERGEN | Non-Toxin | -0.19 | -0.34 | 0.05 | 0 | 6.39 | 2208.77 | outside |
| 971 | 991 | GAISSVLNDILSRLDKVEAE | 0.0342 | 20 | ABCPRED | 1 | NON-ALLERGEN | Non-Toxin | -0.11 | 0.19 | 0.31 | -2 | 4.32 | 2129.69 | outside |
| 1139 | 1155 | DPLQPELDSFKEELDK | -0.5669 | 16 | ABCPRED,BCPREDS | 2 | NON-ALLERGEN | Non-Toxin | -0.32 | -1.38 | 1.04 | -4 | 4.02 | 1903.3 | outside |
| 804 | 818 | QILPDPSKPSKRSF | 0.4021 | 14 | ABCPRED | 1 | NON-ALLERGEN | Non-Toxin | -0.32 | -1.1 | 0.5 | 2 | 10.01 | 1600.04 | outside |
| 1194 | 1216 | NESLIDLQELGKYEQYIKWPWY | 0.88 | 22 | AAP | 1 | NON-ALLERGEN | Non-Toxin | -0.14 | -0.93 | -0.17 | -2 | 4.41 | 2816.51 | outside_transmembrane-region |
| 64 | 84 | WFHAIHVSGTNGTKRFDNPV | 0.41 | 20 | ABCPRED | 1 | ALLERGEN | Non-Toxin | -0.12 | -0.53 | -0.29 | 2 | 9.11 | 2283.83 | outside |
| 905 | 925 | RFNGIGVTQNVLYENQKLIA | 0.6196 | 20 | ABCPRED | 1 | NON-ALLERGEN | Non-Toxin | -0.11 | -0.13 | -0.29 | 1 | 8.93 | 2277.93 | outside |
| 402 | 428 | IRGDEVRQIAPGQTGKIADYNYKLPD | 0.9322 | 26 | literature | 1 | ALLERGEN | Non-Toxin | -0.25 | -0.92 | 0.38 | 0 | 6.5 | 2918.65 | outside |
| 1003 | 1017 | SLQTYVTQQLIRAA | -0.1937 | 14 | ABCPRED | 1 | NON-ALLERGEN | Non-Toxin | -0.11 | 0.1 | -0.51 | 1 | 9.1 | 1592.04 | outside |
| 407 | 427 | VRQIAPGQTGKIADYNYKLP | 1.2386 | 20 | ABCPRED | 1 | NON-ALLERGEN | Non-Toxin | -0.18 | -0.65 | -0.01 | 2 | 9.55 | 2232.87 | outside |
| 1247 | 1267 | CCSCGSCCKFDEDDSEPVLK | 0.2158 | 20 | ABCPRED | 1 | NON-ALLERGEN | Toxin | -0.22 | -0.32 | 0.55 | -3 | 4.11 | 2168.68 | inside |
| 1156 | 1176 | FKNHTSPDVDLGDISGINAS | 0.9142 | 20 | ABCPRED | 1 | ALLERGEN | Non-Toxin | -0.11 | -0.42 | 0.12 | -1.5 | 4.42 | 2087.52 | outside |

| | | | | | | | | | | | | | | | |
|---|---|---|---|---|---|---|---|---|---|---|---|---|---|---|---|
| 160 | 174 | YSSANNCTFEYVSQ | 0.1528 | 14 | ABCPRED | 1 | NON-ALLERGEN | Non-Toxin | -0.15 | -0.6 | -0.43 | -1 | 4 | 1612.87 | outside |
| 696 | 718 | TMSLGAENSVAYSNNSIAIPTN | 0.5154 | 22 | FBCPred | 1 | NON-ALLERGEN | Non-Toxin | -0.04 | -0.05 | -0.35 | -1 | 4 | 2254.77 | outside |
| 251 | 267 | PGDSSSGWTAGAAAYY | 0.2562 | 16 | ABCPRED,AAP | 2 | ALLERGEN | Non-Toxin | 0.01 | -0.36 | -0.41 | -1 | 3.8 | 1560.8 | outside |
| 669 | 683 | GICASYQTQTNSPR | 0.4272 | 14 | ABCPRED | 1 | NON-ALLERGEN | Non-Toxin | -0.25 | -0.89 | -0.16 | 1 | 8.57 | 1525.85 | outside |
| 181 | 201 | GKQGNFKNLREFVFKNIDGY | 0.1265 | 20 | ABCPRED | 1 | ALLERGEN | Non-Toxin | -0.24 | -0.94 | 0.2 | 2 | 9.55 | 2374.99 | outside |
| 794 | 814 | IKDFGGFNFSQILPDPSKPS | 0.1781 | 20 | ABCPRED | 1 | NON-ALLERGEN | Non-Toxin | -0.09 | -0.43 | 0.02 | 0 | 6.31 | 2194.76 | outside |
| 789 | 815 | YKTPPIKDFGGFNFSQILPDPSKR | 0.2569 | 24 | ellipro,literature | 2 | NON-ALLERGEN | Non-Toxin | -0.19 | -0.83 | 0.14 | 2 | 9.55 | 2753.5 | outside |
| 276 | 290 | LLKYNENGTITDAV | 0.3992 | 14 | ABCPRED | 1 | NON-ALLERGEN | Non-Toxin | -0.09 | -0.21 | -0.08 | -1 | 4.38 | 1550.94 | outside |
| 900 | 916 | MQMAYRFNGIGVTQNV | 1.1008 | 16 | ABCPRED | 1 | NON-ALLERGEN | Non-Toxin | -0.07 | 0 | -0.58 | 1 | 9.1 | 1829.37 | outside |
| 490 | 510 | FPLQSYGFQPTNGVGYQPYR | 0.6275 | 20 | ABCPRED | 1 | NON-ALLERGEN | Non-Toxin | -0.11 | -0.82 | -0.58 | 1 | 8.83 | 2319.86 | outside |
| 814 | 834 | KRSFIEDLLFNKVTLADAGF | 0.445 | 20 | ABCPRED | 1 | ALLERGEN | Non-Toxin | -0.09 | 0.19 | 0.05 | 0 | 6.47 | 2284.93 | outside |
| 469 | 489 | STEIYQAGSTPCNGVEGFNC | 0.1782 | 20 | ABCPRED | 1 | NON-ALLERGEN | Non-Toxin | -0.07 | -0.32 | -0.21 | -2 | 3.8 | 2077.51 | outside |
| 219 | 233 | GFSALEPLVDLPIG | 0.7572 | 14 | ABCPRED | 1 | NON-ALLERGEN | Non-Toxin | 0.16 | 0.92 | -0.39 | -2 | 3.67 | 1427.87 | outside |
| 254 | 268 | SSSGWTAGAAAYYV | 0.3218 | 14 | ABCPRED | 1 | NON-ALLERGEN | Non-Toxin | 0.09 | 0.29 | -0.79 | 0 | 5.87 | 1390.65 | outside |
| 857 | 871 | GLTVLPPLLTDEMI | 0.4082 | 14 | ABCPRED | 1 | NON-ALLERGEN | Non-Toxin | 0.14 | 0.99 | -0.47 | -2 | 3.67 | 1512.06 | outside |
| 868 | 888 | EMIAQYTSALLAGTITSGWT | 0.2956 | 20 | ABCPRED | 1 | NON-ALLERGEN | Non-Toxin | 0.08 | 0.47 | -0.68 | -1 | 4 | 2114.7 | outside |
| 87 | 107 | NDGVYFASTEKSNIIRGWIF | -0.2393 | 20 | ABCPRED | 1 | NON-ALLERGEN | Non-Toxin | -0.06 | -0.13 | -0.28 | 0 | 6.42 | 2317.88 | outside |
| 316 | 336 | SNFRVQPTESIVRFPNITNL | 0.6511 | 20 | ABCPRED | 1 | NON-ALLERGEN | Non-Toxin | -0.17 | -0.3 | -0.19 | 1 | 9.95 | 2332.94 | outside |
| 863 | 883 | PLLTDEMIAQYTSALLAGTI | 0.2489 | 20 | ABCPRED | 1 | NON-ALLERGEN | Non-Toxin | 0.09 | 0.74 | -0.53 | -2 | 3.67 | 2121.78 | outside |
| 524 | 599 | NGLTGTGVLTESNKKFLPFQQFGRDIADTTDAV | 0.7634 | 75 | literature | 1 | NON-ALLERGEN | too long | | | | | | | outside |
| 1001 | 1021 | LQSLQTYVTQQLIRAAEIRA | 0.0026 | 20 | ABCPRED | 1 | NON-ALLERGEN | Non-Toxin | -0.16 | 0 | -0.25 | 1 | 9.1 | 2302.97 | outside |
| 136 | 150 | CNDPFLGVYYHKNN | 0.68 | 14 | ABCPRED | 1 | NON-ALLERGEN | Non-Toxin | -0.16 | -0.89 | -0.38 | 0.5 | 7.05 | 1684.06 | outside |
| 413 | 427 | GQTGKIADYNYKLP | 1.2756 | 14 | ABCPRED | 1 | NON-ALLERGEN | Non-Toxin | -0.19 | -0.99 | 0.02 | 1 | 8.76 | 1567.97 | outside |
| 406 | 420 | EVRQIAPGQTGKIA | 1.3182 | 14 | ABCPRED | 1 | NON-ALLERGEN | Non-Toxin | -0.16 | -0.37 | 0.21 | 1 | 9.1 | 1467.9 | outside |
| 9 | 29 | PLVSSQCVNLTTRTQLPPAY | 1.2186 | 20 | ABCPRED | 1 | NON-ALLERGEN | Non-Toxin | -0.1 | -0.04 | -0.46 | 1 | 8.57 | 2188.82 | outside |
| 134 | 148 | QFCNDPFLGVYYHK | 0.4646 | 14 | ABCPRED | 1 | NON-ALLERGEN | Non-Toxin | -0.08 | -0.44 | -0.57 | 0.5 | 7.05 | 1731.16 | outside |
| 57 | 71 | PFFSNVTWFHAIHV | 0.6603 | 14 | ABCPRED | 1 | NON-ALLERGEN | Non-Toxin | 0.17 | 0.66 | -1.22 | 1 | 7.26 | 1702.15 | outside |
| 329 | 345 | FPNITNLCPFGEVFNA | 0.6058 | 16 | ABCPRED | 1 | ALLERGEN | Non-Toxin | 0.08 | 0.43 | -0.68 | -1 | 4 | 1783.26 | outside |
| 819 | 833 | EDLLFNKVTLADAG | 0.7009 | 14 | ABCPRED | 1 | NON-ALLERGEN | Non-Toxin | -0.04 | 0.21 | 0.1 | -2 | 4.03 | 1505.89 | outside |
| 824 | 844 | NKVTLADAGFIKQYGDCLGD | 0.2805 | 20 | ABCPRED | 1 | NON-ALLERGEN | Non-Toxin | -0.09 | -0.17 | 0.07 | -1 | 4.43 | 2128.68 | outside |
| 764 | 784 | NRALTGIAVEQDKNTQEVFA | 0.4147 | 20 | ABCPRED | 1 | ALLERGEN | Non-Toxin | -0.2 | -0.49 | 0.22 | -1 | 4.68 | 2204.72 | outside |
| 81 | 95 | NPVLPFNDGVYFAS | 0.3095 | 14 | ABCPRED | 1 | NON-ALLERGEN | Non-Toxin | 0.06 | 0.24 | -0.64 | -1 | 3.8 | 1539.9 | outside |
| 1240 | 1262 | CCSCLKGCCSCGSCCKFDEDDS | 0.021 | 22 | BCPREDS | 1 | NON-ALLERGEN | Non-Toxin | -0.19 | 0.04 | 0.31 | -2 | 4.23 | 2296.88 | inside |
| 689 | 705 | SQSIIAYTMSLGAENS | 0.5628 | 16 | ABCPRED | 1 | NON-ALLERGEN | Non-Toxin | -0.01 | 0.14 | -0.36 | -1 | 4 | 1672.07 | outside |
| 173 | 193 | QPFLMDLEGKQGNFKNLREF | 0.6632 | 20 | ABCPRED | 1 | NON-ALLERGEN | Toxin | -0.23 | -0.88 | 0.23 | 0 | 6.53 | 2412.07 | outside |
| 86 | 106 | FNDGVYFASTEKSNIIRGWI | -0.2699 | 20 | ABCPRED | 1 | NON-ALLERGEN | Non-Toxin | -0.06 | -0.13 | -0.28 | 0 | 6.42 | 2317.88 | outside |
| 1006 | 1026 | TYVTQQLIRAAEIRASANLA | 0.2689 | 20 | ABCPRED | 1 | ALLERGEN | Non-Toxin | -0.12 | 0.16 | -0.22 | 1 | 9.1 | 2189.79 | outside |
| 71 | 93 | SGTNGTKRFDNPVLPFNDGVYF | 0.3722 | 22 | FBCPred | 1 | ALLERGEN | Non-Toxin | -0.14 | -0.6 | -0.11 | 0 | 6.31 | 2445.99 | outside |
| 224 | 238 | EPLVDLPIGINITR | 0.9556 | 14 | ABCPRED | 1 | NON-ALLERGEN | Non-Toxin | -0.01 | 0.43 | -0.12 | -1 | 4.38 | 1550.05 | outside |
| 1060 | 1080 | VVFLHVTYVPAQEKNFTTAP | 0.9571 | 20 | ABCPRED | 1 | NON-ALLERGEN | Non-Toxin | 0.01 | 0.28 | -0.57 | 0.5 | 7.1 | 2261.9 | outside |
| 49 | 69 | HSTQDLFLPFFSNVTWFHAI | 0.3126 | 20 | ABCPRED | 1 | NON-ALLERGEN | Non-Toxin | 0.08 | 0.34 | -0.93 | 0 | 5.99 | 2408.01 | outside |
| 581 | 601 | TLEILDITPCSFGGVSVITP | 1.2987 | 20 | ABCPRED | 1 | NON-ALLERGEN | Non-Toxin | 0.14 | 1 | -0.51 | -2 | 3.67 | 2062.71 | outside |
| 853 | 873 | QKFNGLTVLPPLLTDEMIAQ | 0.199 | 20 | ABCPRED | 1 | NON-ALLERGEN | Non-Toxin | -0.01 | 0.2 | -0.3 | -1 | 4.38 | 2228.95 | outside |
| 99 | 119 | NIIRGWIFGTTLDSKTQSLL | 0.0538 | 20 | ABCPRED | 1 | NON-ALLERGEN | Non-Toxin | -0.04 | 0.17 | -0.4 | 1 | 9.1 | 2263.94 | outside |
| 402 | 422 | IRGDEVRQIAPGQTGKIADY | 0.8186 | 20 | ABCPRED | 1 | NON-ALLERGEN | Non-Toxin | -0.23 | -0.7 | 0.39 | 0 | 6.47 | 2187.74 | outside |
| 511 | 531 | VVLSFELLHAPATVCGPKKS | 0.4194 | 20 | ABCPRED | 1 | NON-ALLERGEN | Non-Toxin | 0.02 | 0.63 | -0.29 | 1.5 | 8.54 | 2096.81 | outside |
| 310 | 330 | KGIYQTSNFRVQPTESIVRF | 0.383 | 20 | ABCPRED | 1 | NON-ALLERGEN | Non-Toxin | -0.21 | -0.51 | -0.07 | 2 | 10 | 2370.99 | outside |
| 346 | 360 | RFASVYAWNRKRIS | 0.5368 | 14 | ABCPRED | 1 | NON-ALLERGEN | Non-Toxin | -0.34 | -0.69 | 0.02 | 4 | 11.73 | 1754.21 | outside |
| 257 | 273 | GWTAGAAAYYVGYLQP | 0.621 | 16 | ABCPRED | 1 | NON-ALLERGEN | Non-Toxin | 0.13 | 0.21 | -0.99 | 0 | 5.87 | 1688.1 | outside |
| 935 | 949 | QDSLSSTASALGKL | 0.4957 | 14 | ABCPRED | 1 | NON-ALLERGEN | Non-Toxin | -0.11 | -0.01 | 0.04 | 0 | 6.19 | 1377.71 | outside |
| 3 | 4 | V | short | 1 | literature | 1 | NON-ALLERGEN | Non-Toxin | 0.54 | 4.2 | -1.5 | 0 | 5.88 | 117.15 | outside |
| 385 | 405 | TKLNDLCFTNVYADSFVIRG | 1.049 | 20 | ABCPRED | 1 | NON-ALLERGEN | Non-Toxin | -0.08 | 0.2 | -0.27 | 0 | 6.28 | 2276.88 | outside |
| 60 | 74 | SNVTWFHAIHVSGT | 0.6819 | 14 | ABCPRED | 1 | NON-ALLERGEN | Non-Toxin | 0.06 | 0.24 | -0.87 | 1 | 7.26 | 1555.92 | outside |
| 1094 | 1108 | VFVSNGTHWFVTQR | 0.2064 | 14 | ABCPRED | 1 | NON-ALLERGEN | Non-Toxin | -0.05 | 0 | -0.75 | 1.5 | 10.11 | 1678.09 | outside |
| 270 | 292 | LQPRTFLLKYNENGTITDAVDC | 0.4892 | 22 | BCPREDS | 1 | ALLERGEN | Non-Toxin | -0.16 | -0.35 | -0.03 | -1 | 4.56 | 2512.16 | outside |
| 726 | 746 | ILPVSMTKTSVDCTMYICGD | 0.5044 | 20 | ABCPRED | 1 | NON-ALLERGEN | Non-Toxin | 0.01 | 0.6 | -0.35 | -1 | 4.21 | 2177.88 | outside |
| 840 | 860 | CLGDIAARDLICAQKFNGLT | 0.5933 | 20 | ABCPRED | 1 | NON-ALLERGEN | Non-Toxin | -0.05 | 0.48 | -0.15 | 0 | 6.25 | 2122.78 | outside |
| 1139 | 1159 | DPLQPELDSFKEELDKYFKN | -0.5291 | 20 | ABCPRED | 1 | NON-ALLERGEN | Non-Toxin | -0.31 | -1.4 | 0.75 | -3 | 4.3 | 2455.99 | outside |
| 1047 | 1061 | YHLMSFPQSAPHGV | 0.1729 | 14 | ABCPRED | 1 | NON-ALLERGEN | Non-Toxin | 0.02 | -0.14 | -0.72 | 1 | 7.25 | 1570.99 | outside |
| 454 | 474 | RLFRKSNLKPFERDISTEIY | -0.0299 | 20 | ABCPRED | 1 | NON-ALLERGEN | Non-Toxin | -0.35 | -0.92 | 0.49 | 2 | 9.71 | 2513.18 | outside |
| 201 | 215 | FKIYSKHTPINLVR | 0.9156 | 14 | ABCPRED | 1 | ALLERGEN | Non-Toxin | -0.17 | -0.26 | -0.22 | 3.5 | 10.3 | 1716.27 | outside |
| 460 | 474 | NLKPFERDISTEIY | 0.2921 | 14 | ABCPRED | 1 | NON-ALLERGEN | Non-Toxin | -0.24 | -0.8 | 0.35 | -1 | 4.68 | 1725.13 | outside |
| 162 | 182 | SANNCTFEYVSQPFLMDLEG | 0.1921 | 20 | ABCPRED | 1 | NON-ALLERGEN | Toxin | -0.06 | -0.15 | -0.27 | -3 | 3.58 | 2265.78 | outside |
| 980 | 1000 | ILSRLDKVEAEVQIDRLITG | -0.1035 | 20 | ABCPRED | 1 | NON-ALLERGEN | Non-Toxin | -0.16 | 0.14 | 0.34 | -1 | 4.79 | 2268.95 | outside |
| 1095 | 1115 | FVSNGTHWFVTQRNFYEPQI | 0.2942 | 20 | ABCPRED | 1 | ALLERGEN | Non-Toxin | -0.1 | -0.51 | -0.61 | 0.5 | 7.1 | 2471.03 | outside |
| 644 | 658 | QTRAGCLIGAEHVN | 1.3917 | 14 | ABCPRED | 1 | NON-ALLERGEN | Non-Toxin | -0.12 | -0.08 | -0.11 | 0.5 | 7.07 | 1468.86 | outside |
| 1035 | 1049 | GQSKRVDFCGKGYH | 1.1237 | 14 | ABCPRED | 1 | NON-ALLERGEN | Non-Toxin | -0.31 | -1.16 | 0.34 | 2.5 | 9.25 | 1581.97 | outside |
| 860 | 874 | VLPPLLTDEMIAQY | 0.2075 | 14 | ABCPRED | 1 | NON-ALLERGEN | Non-Toxin | 0.07 | 0.58 | -0.5 | -2 | 3.67 | 1603.12 | outside |
| 313 | 335 | YQTSNFRVQPTESIVRFPNITN | 0.5541 | 22 | BCPREDS | 1 | NON-ALLERGEN | Non-Toxin | -0.22 | -0.69 | -0.2 | 1 | 9.1 | 2612.22 | outside |
| 552 | 566 | LTESNKKFLPFQQF | 0.6257 | 14 | ABCPRED | 1 | ALLERGEN | Non-Toxin | -0.18 | -0.64 | -0.11 | 1 | 8.94 | 1727.2 | outside |
| 789 | 803 | YKTPPIKDFGGFNF | 0.2117 | 14 | ABCPRED,IEDB | 2 | NON-ALLERGEN | Non-Toxin | -0.07 | -0.56 | -0.2 | 1 | 8.83 | 1631.06 | outside |
| 965 | 979 | QLSSNFGAISSVLN | 0.6404 | 14 | ABCPRED | 1 | ALLERGEN | Non-Toxin | 0.02 | 0.49 | -0.58 | 0 | 5.88 | 1436.79 | outside |
| 321 | 341 | QPTESIVRFPNITNLCPFGE | 0.4358 | 20 | ABCPRED | 1 | NON-ALLERGEN | Non-Toxin | -0.09 | -0.22 | -0.19 | -1 | 4.54 | 2262.86 | outside |
| 97 | 117 | KSNIIRGWIFGTTLDSKTQS | -0.0695 | 20 | ABCPRED | 1 | NON-ALLERGEN | Non-Toxin | -0.17 | -0.45 | -0.05 | 2 | 10.01 | 2252.86 | outside |
| 551 | 571 | VLTESNKKFLPFQQFGRDIA | 0.3356 | 20 | ABCPRED | 1 | NON-ALLERGEN | Non-Toxin | -0.16 | -0.34 | 0.03 | 1 | 8.93 | 2338.99 | outside |
| 88 | 104 | DGVYFASTEKSNIIRG | 0.0323 | 16 | ABCPRED,BCPREDS | 2 | NON-ALLERGEN | Non-Toxin | -0.15 | -0.34 | 0.12 | 0 | 6.42 | 1757.16 | outside |
| 912 | 926 | TQNVLYENQKLIAN | 0.1556 | 14 | ABCPRED | 1 | NON-ALLERGEN | Non-Toxin | -0.19 | -0.63 | -0.22 | 0 | 6.35 | 1648.07 | outside |

| | | | | | | | | | | | | | | | |
|---|---|---|---|---|---|---|---|---|---|---|---|---|---|---|---|
| 128 | 142 | IKVCEFQFCNDPFL | -0.5546 | 14 | ABCPRED | 1 | NON-ALLERGEN | Non-Toxin | -0.01 | 0.46 | -0.37 | -1 | 4.38 | 1703.2 | outside |
| 1168 | 1188 | DISGINASVVNIQKEIDRLN | 0.4345 | 20 | ABCPRED | 1 | NON-ALLERGEN | Non-Toxin | -0.16 | -0.15 | 0.2 | -1 | 4.56 | 2198.77 | outside |
| 495 | 515 | YGFQPTNGVGYQPYRVVVLS | 0.8397 | 20 | ABCPRED | 1 | NON-ALLERGEN | Non-Toxin | -0.03 | -0.07 | -0.68 | 1 | 8.83 | 2244.84 | outside |
| 201 | 221 | FKIYSKHTPINLVRDLPQGF | 0.6125 | 20 | ABCPRED | 1 | NON-ALLERGEN | Toxin | -0.13 | -0.3 | -0.21 | 2.5 | 9.72 | 2374.09 | outside |
| 547 | 567 | TGTGVLTESNKKFLPFQQFG | 0.7524 | 20 | ABCPRED | 1 | NON-ALLERGEN | Non-Toxin | -0.09 | -0.36 | -0.2 | 1 | 8.94 | 2199.8 | outside |
| 102 | 122 | RGWIFGTTLDSKTQSLLIVN | 0.5414 | 20 | ABCPRED | 1 | NON-ALLERGEN | Non-Toxin | -0.05 | 0.15 | -0.38 | 1 | 9.1 | 2249.91 | outside |
| 86 | 100 | FNDGVYFASTEKSN | 0.5168 | 14 | ABCPRED | 1 | NON-ALLERGEN | Non-Toxin | -0.16 | -0.74 | 0.02 | -1 | 4.38 | 1578.84 | outside |
| 772 | 792 | VEQDKNTQEVFAQVKQIYKT | 0.1814 | 20 | ABCPRED | 1 | NON-ALLERGEN | Non-Toxin | -0.29 | -1.03 | 0.33 | 0 | 6.52 | 2396.99 | outside |
| 1221 | 1257 | AGLIAIVMVTIMLCCMTSCCSCLKGCCSCGSCCK | 0.143 | 36 | IEDB,literature | 2 | NON-ALLERGEN | Non-Toxin | 0.13 | 1.7 | -0.74 | 2 | 8.23 | 3701.25 | transmembrane-region_inside |
| 260 | 274 | AGAAAYYVGYLQPR | 1.0663 | 14 | ABCPRED | 1 | NON-ALLERGEN | Non-Toxin | 0 | 0.06 | -0.64 | 1 | 8.83 | 1499.88 | outside |
| 58 | 72 | FFSNVTWFHAIHVS | 0.6633 | 14 | ABCPRED | 1 | ALLERGEN | Non-Toxin | 0.15 | 0.71 | -1.2 | 1 | 7.26 | 1692.11 | outside |
| 603 | 609 | NTSNQV | 0.3619 | 6 | ellipro,literature | 2 | ALLERGEN | Non-Toxin | -0.31 | -1.3 | -0.17 | 0 | 5.88 | 661.75 | outside |
| 960 | 974 | NTLVKQLSSNFGAI | 0.0188 | 14 | ABCPRED | 1 | NON-ALLERGEN | Non-Toxin | -0.03 | 0.27 | -0.44 | 1 | 9.11 | 1491.92 | outside |
| 601 | 621 | GTNTSNQVAVLYQDVNCTEV | 0.8407 | 20 | ABCPRED | 1 | NON-ALLERGEN | Non-Toxin | -0.11 | -0.21 | -0.28 | -2 | 3.67 | 2155.62 | outside |
| 905 | 921 | RFNGIGVTQNVLYENQ | 0.7287 | 16 | FBCPred | 1 | NON-ALLERGEN | Non-Toxin | -0.16 | -0.55 | -0.3 | 0 | 6.35 | 1852.29 | outside |
| 348 | 362 | ASVYAWNRKRISNC | 0.3411 | 14 | ABCPRED | 1 | NON-ALLERGEN | Non-Toxin | -0.3 | -0.64 | -0.07 | 3 | 10.07 | 1668.09 | outside |
| 415 | 435 | TGKIADYNYKLPDDFTGCVI | 0.5848 | 20 | ABCPRED | 1 | NON-ALLERGEN | Non-Toxin | -0.08 | -0.2 | -0.06 | -1 | 4.43 | 2233.81 | outside |
| 358 | 372 | ISNCVADYSVLYNS | -0.4076 | 14 | ABCPRED | 1 | NON-ALLERGEN | Non-Toxin | -0.01 | 0.39 | -0.6 | -1 | 3.8 | 1547.89 | outside |
| 339 | 353 | GEVFNATRFASVYA | -0.0008 | 14 | ABCPRED | 1 | NON-ALLERGEN | Non-Toxin | -0.02 | 0.34 | -0.41 | 0 | 6.35 | 1531.87 | outside |
| 199 | 219 | GYFKIYSKHTPINLVRDLPQ | 0.3555 | 20 | ABCPRED | 1 | NON-ALLERGEN | Non-Toxin | -0.16 | -0.51 | -0.2 | 2.5 | 9.55 | 2390.09 | outside |
| 1197 | 1217 | LIDLQELGKYEQYIKWPWYI | 0.9949 | 20 | ABCPRED | 1 | ALLERGEN | Non-Toxin | -0.04 | -0.41 | -0.45 | -1 | 4.68 | 2599.35 | outside_transmembrane-region |
| 1131 | 1147 | GIVNNTVYDPLQPELD | 0.3695 | 16 | AAP | 1 | ALLERGEN | Non-Toxin | -0.08 | -0.38 | -0.09 | -3 | 3.5 | 1787.2 | outside |
| 1052 | 1072 | FPQSAPHGVVFLHVTYVPAQ | 0.564 | 20 | ABCPRED | 1 | NON-ALLERGEN | Non-Toxin | 0.09 | 0.42 | -0.84 | 1 | 7.25 | 2194.82 | outside |
| 471 | 485 | EIYQAGSTPCNGVE | -0.0233 | 14 | ABCPRED | 1 | NON-ALLERGEN | Non-Toxin | -0.08 | -0.44 | -0.06 | -2 | 3.8 | 1467.77 | outside |
| 156 | 176 | EFRVYSSANNCTFEYVSQPF | 0.1654 | 20 | ABCPRED | 1 | ALLERGEN | Toxin | -0.14 | -0.41 | -0.33 | -1 | 4.54 | 2388.86 | outside |
| 509 | 523 | RVVVLSFELLHAPA | 0.7038 | 14 | ABCPRED | 1 | NON-ALLERGEN | Non-Toxin | 0.09 | 1.2 | -0.54 | 0.5 | 7.1 | 1551.07 | outside |
| 267 | 287 | VGYLQPRTFLLKYNENGTIT | 0.6671 | 20 | ABCPRED | 1 | NON-ALLERGEN | Non-Toxin | -0.11 | -0.33 | -0.37 | 1 | 8.83 | 2327.98 | outside |
| 654 | 674 | EHVNNSYECDIPIGAGICAS | 0.7186 | 20 | ABCPRED | 1 | NON-ALLERGEN | Non-Toxin | -0.03 | 0.01 | -0.13 | -2.5 | 4.14 | 2092.57 | outside |
| 22 | 36 | TQLPPAYTNSFTRG | 0.1422 | 14 | ABCPRED | 1 | NON-ALLERGEN | Non-Toxin | -0.18 | -0.78 | -0.33 | 1 | 9.1 | 1552.9 | outside |
| 979 | 999 | DILSRLDKVEAEVQIDRLIT | -0.0973 | 20 | ABCPRED | 1 | NON-ALLERGEN | Non-Toxin | -0.2 | -0.01 | 0.49 | -2 | 4.44 | 2326.98 | outside |
| 129 | 143 | KVCEFQFCNDPFLG | -0.243 | 14 | ABCPRED | 1 | NON-ALLERGEN | Non-Toxin | -0.05 | 0.11 | -0.24 | -1 | 4.38 | 1647.09 | outside |
| 96 | 110 | EKSNIIRGWIFGTT | -0.3262 | 14 | ABCPRED | 1 | NON-ALLERGEN | Non-Toxin | -0.09 | -0.21 | -0.19 | 1 | 9.1 | 1622.07 | outside |
| 422 | 436 | NYKLPDDFTGCVIA | 0.0699 | 14 | ABCPRED | 1 | NON-ALLERGEN | Non-Toxin | -0.04 | 0.09 | -0.19 | -1 | 4.21 | 1555.96 | outside |
| 657 | 677 | NNSYECDIPIGAGICASYQT | 0.738 | 20 | ABCPRED | 1 | NON-ALLERGEN | Non-Toxin | -0.05 | -0.14 | -0.31 | -2 | 3.67 | 2119.59 | outside |
| 504 | 524 | GYQPYRVVVLSFELLHAPAT | 0.8415 | 20 | ABCPRED | 1 | NON-ALLERGEN | Non-Toxin | 0.02 | 0.4 | -0.62 | 0.5 | 7.1 | 2260.92 | outside |
| 722 | 742 | VTTEILPVSMTKTSVDCTMY | 1.2085 | 20 | ABCPRED | 1 | NON-ALLERGEN | Non-Toxin | -0.02 | 0.41 | -0.32 | -1 | 4.38 | 2219.9 | outside |
| 309 | 323 | EKGIYQTSNFRVQP | 0.8229 | 14 | ABCPRED | 1 | NON-ALLERGEN | Non-Toxin | -0.28 | -1.12 | 0.1 | 1 | 8.93 | 1667.06 | outside |
| 885 | 899 | GWTFGAGAALQIPF | 0.4208 | 14 | ABCPRED | 1 | ALLERGEN | Non-Toxin | 0.22 | 0.81 | -0.98 | 0 | 5.88 | 1435.85 | outside |
| 458 | 472 | KSNLKPFERDISTE | 0.4643 | 14 | ABCPRED | 1 | ALLERGEN | Non-Toxin | -0.39 | -1.36 | 0.88 | 0 | 6.53 | 1664.04 | outside |
| 1149 | 1169 | KEELDKYFKNHTSPDVDLGD | 0.2468 | 20 | ABCPRED | 1 | NON-ALLERGEN | Non-Toxin | -0.33 | -1.48 | 0.83 | -2.5 | 4.52 | 2350.81 | outside |
| 978 | 998 | NDILSRLDKVEAEVQIDRLI | -0.1996 | 20 | ABCPRED | 1 | ALLERGEN | Non-Toxin | -0.23 | -0.15 | 0.52 | -2 | 4.44 | 2339.98 | outside |
| 1052 | 1074 | FPQSAPHGVVFLHVTYVPAQEK | 0.6101 | 22 | literature | 1 | NON-ALLERGEN | Non-Toxin | 0 | 0.05 | -0.49 | 1 | 7.25 | 2452.14 | outside |
| 390 | 410 | LCFTNVYADSFVIRGDEVRQ | 0.2976 | 20 | ABCPRED | 1 | NON-ALLERGEN | Non-Toxin | -0.14 | 0.06 | -0.08 | -1 | 4.56 | 2332.9 | outside |
| 671 | 700 | CASYQTQTNSPRRARSVASQSIIAYTMSL | 0.3009 | 29 | literature | 1 | ALLERGEN | Non-Toxin | -0.22 | -0.34 | -0.19 | 3 | 10.06 | 3191.99 | outside |
| 479 | 493 | PCNGVEGFNCYFPL | 0.4399 | 14 | ABCPRED | 1 | NON-ALLERGEN | Non-Toxin | 0.05 | 0.2 | -0.66 | -1 | 4 | 1559.97 | outside |
| 974 | 988 | SSVLNDILSRLDKV | -0.3181 | 14 | ABCPRED | 1 | NON-ALLERGEN | Non-Toxin | -0.17 | 0.21 | 0.21 | 0 | 6.31 | 1559 | outside |
| 1183 | 1203 | IDRLNEVAKNLNESLIDLQE | 0.4322 | 20 | ABCPRED | 1 | NON-ALLERGEN | Non-Toxin | -0.23 | -0.53 | 0.47 | -3 | 4.18 | 2326.9 | outside |
| 220 | 234 | FSALEPLVDLPIGI | 0.7452 | 14 | ABCPRED | 1 | NON-ALLERGEN | Non-Toxin | 0.21 | 1.27 | -0.51 | -2 | 3.67 | 1483.98 | outside |
| 746 | 766 | STECSNLLLQYGSFCTQLNR | 0.7968 | 20 | ABCPRED | 1 | NON-ALLERGEN | Non-Toxin | -0.16 | -0.23 | -0.36 | 0 | 6.29 | 2277.85 | outside |
| 274 | 288 | TFLLKYNENGTITD | 0.668 | 14 | ABCPRED | 1 | NON-ALLERGEN | Non-Toxin | -0.12 | -0.49 | -0.14 | -1 | 4.38 | 1629.01 | outside |
| 773 | 787 | EQDKNTQEVFAQVK | 0.361 | 14 | ABCPRED | 1 | NON-ALLERGEN | Non-Toxin | -0.37 | -1.43 | 0.67 | -1 | 4.68 | 1664.01 | outside |
| 505 | 528 | YQPYRVVVLSFELLHAPATVCGP | 0.4697 | 23 | IEDB,literature | 2 | NON-ALLERGEN | Non-Toxin | 0.04 | 0.57 | -0.65 | 0.5 | 7.06 | 2560.35 | outside |
| 650 | 664 | LIGAEHVNNSYECD | 0.9242 | 14 | ABCPRED | 1 | ALLERGEN | Non-Toxin | -0.12 | -0.46 | 0.02 | -2.5 | 4.14 | 1563.86 | outside |
| 440 | 486 | GGNYNYLYRLFRKSNLKPFERDISTEIYQAGSTP | 0.21 | 46 | literature | 1 | NON-ALLERGEN | Non-Toxin | -0.22 | -0.79 | 0.11 | 1 | 8.42 | 5216.48 | outside |
| 1064 | 1078 | HVTYVPAQEKNFTT | 0.8277 | 14 | ABCPRED | 1 | NON-ALLERGEN | Non-Toxin | -0.15 | -0.69 | -0.26 | 0.5 | 7.1 | 1635.01 | outside |
| 474 | 488 | QAGSTPCNGVEGFN | 0.271 | 14 | ABCPRED | 1 | NON-ALLERGEN | Non-Toxin | -0.08 | -0.5 | -0.14 | -1 | 4 | 1380.65 | outside |
| 648 | 662 | GCLIGAEHVNNSYE | 0.8685 | 14 | ABCPRED | 1 | ALLERGEN | Non-Toxin | -0.05 | -0.24 | -0.19 | -1.5 | 4.51 | 1505.83 | outside |
| 870 | 890 | IAQYTSALLAGTITSGWTFG | 0.3239 | 20 | ABCPRED | 1 | NON-ALLERGEN | Toxin | 0.14 | 0.67 | -0.89 | 0 | 5.88 | 2058.62 | outside |
| 440 | 460 | NLDSKVGGNYNYLYRLFRKS | 0.2507 | 20 | ABCPRED | 1 | NON-ALLERGEN | Non-Toxin | -0.29 | -0.93 | 0 | 3 | 9.83 | 2408.01 | outside |
| 15 | 29 | CVNLTTRTQLPPAY | 1.483 | 14 | ABCPRED | 1 | NON-ALLERGEN | Non-Toxin | -0.13 | -0.14 | -0.48 | 1 | 8.57 | 1577.03 | outside |
| 223 | 243 | LEPLVDLPIGINITRFQTLL | 0.6048 | 20 | ABCPRED | 1 | NON-ALLERGEN | Non-Toxin | 0.06 | 0.8 | -0.49 | -1 | 4.38 | 2266.05 | outside |
| 201 | 270 | LVRDLPQGFSALEPLVDLPIGINITRFQTLLALHR | 0.6317 | 56 | ellipro | 1 | NON-ALLERGEN | too long | | | | | | | outside |
| 805 | 825 | ILPDPSKPSKRSFIEDLLFN | 0.0467 | 20 | ABCPRED | 1 | NON-ALLERGEN | Non-Toxin | -0.17 | -0.37 | 0.26 | 0 | 6.47 | 2316.97 | outside |
| 501 | 515 | NGVGYQPYRVVVLS | 0.804 | 14 | ABCPRED | 1 | NON-ALLERGEN | Non-Toxin | -0.03 | 0.24 | -0.62 | 1 | 8.93 | 1550.99 | outside |
| 1193 | 1213 | LNESLIDLQELGKYEQYIKW | 0.6851 | 20 | ABCPRED | 1 | NON-ALLERGEN | Non-Toxin | -0.15 | -0.64 | 0.01 | -2 | 4.41 | 2483.14 | outside |
| 902 | 922 | MAYRFNGIGVTQNVLYENQK | 0.8298 | 20 | ABCPRED | 1 | ALLERGEN | Non-Toxin | -0.16 | -0.51 | -0.3 | 1 | 8.83 | 2345.97 | outside |
| 596 | 618 | SVITPGTNTSNQVAVLYQDVNC | 0.5248 | 22 | BCPREDS | 1 | ALLERGEN | Non-Toxin | -0.06 | 0.06 | -0.45 | -1 | 3.8 | 2323.89 | outside |
| 547 | 561 | TGTGVLTESNKKFL | 0.498 | 14 | ABCPRED | 1 | NON-ALLERGEN | Non-Toxin | -0.12 | -0.28 | 0.05 | 1 | 8.94 | 1494.92 | outside |
| 970 | 990 | FGAISSVLNDILSRLDKVEA | 0.1201 | 20 | ABCPRED | 1 | NON-ALLERGEN | Non-Toxin | -0.05 | 0.5 | 0.03 | -1 | 4.56 | 2147.75 | outside |
| 155 | 169 | SEFRVYSSANNCTF | -0.0428 | 14 | ABCPRED | 1 | NON-ALLERGEN | Non-Toxin | -0.18 | -0.38 | -0.24 | 0 | 6.32 | 1624.92 | outside |
| 192 | 212 | FVFKNIDGYFKIYSKHTPIN | 0.4815 | 20 | ABCPRED | 1 | ALLERGEN | Non-Toxin | -0.07 | -0.27 | -0.36 | 2.5 | 9.43 | 2432.12 | outside |
| 1238 | 1258 | TSCCSCLKGCCSCGSCCKFD | 0.1024 | 20 | ABCPRED | 1 | NON-ALLERGEN | Non-Toxin | -0.12 | 0.53 | -0.12 | 1 | 7.94 | 2038.67 | inside |
| 621 | 635 | PVAIHADQLTPTWR | 0.6046 | 14 | ABCPRED | 1 | NON-ALLERGEN | Non-Toxin | -0.1 | -0.29 | -0.33 | 0.5 | 7.1 | 1605.03 | outside |
| 1174 | 1196 | ASVVNIQKEIDRLNEVAKNLNE | 0.2423 | 22 | BCPREDS | 1 | NON-ALLERGEN | Non-Toxin | -0.25 | -0.54 | 0.44 | -1 | 4.88 | 2497.14 | outside |
| 1112 | 1128 | PQIITTDNTFVSGNCD | 0.2404 | 16 | ABCPRED,BCPREDS | 2 | NON-ALLERGEN | Non-Toxin | -0.09 | -0.24 | -0.18 | -2 | 3.57 | 1725.09 | outside |
| 574 | 596 | DAVRDPQTLEILDITPCSFGGV | 1.0597 | 22 | AAP | 1 | NON-ALLERGEN | Non-Toxin | -0.05 | 0.18 | 0.02 | -3 | 3.84 | 2346.96 | outside |
| 520 | 536 | APATVCGPKKSTNLVK | 0.0812 | 16 | ABCPRED,AAP | 2 | NON-ALLERGEN | Non-Toxin | -0.15 | -0.17 | 0.12 | 3 | 9.81 | 1614.16 | outside |

| | | | | | | | | | | | | | | | |
|---|---|---|---|---|---|---|---|---|---|---|---|---|---|---|---|
| 403 | 417 | RGDEVRQIAPGQTG | 0.8295 | 14 | ABCPRED | 1 | NON-ALLERGEN | Non-Toxin | -0.32 | -1.14 | 0.59 | 0 | 6.42 | 1483.81 | outside |
| 856 | 870 | NGLTVLPPLLTDEM | 0.2278 | 14 | ABCPRED | 1 | NON-ALLERGEN | Non-Toxin | 0.04 | 0.41 | -0.33 | -2 | 3.67 | 1513 | outside |
| 551 | 565 | VLTESNKKFLPFQQ | 0.758 | 14 | ABCPRED | 1 | NON-ALLERGEN | Toxin | -0.18 | -0.54 | -0.04 | 1 | 8.94 | 1679.16 | outside |
| 194 | 208 | FKNIDGYFKIYSKH | 0.3365 | 14 | ABCPRED | 1 | NON-ALLERGEN | Non-Toxin | -0.17 | -0.79 | -0.09 | 2.5 | 9.43 | 1760.23 | outside |
| 954 | 974 | QNAQALNTLVKQLSSNFGAI | 0.2079 | 20 | ABCPRED | 1 | NON-ALLERGEN | Non-Toxin | -0.07 | 0.04 | -0.42 | 1 | 9.11 | 2117.7 | outside |
| 625 | 647 | HADQLTPTWRVYSTGSNVFQTR | 0.5124 | 22 | BCPREDS | 1 | NON-ALLERGEN | Non-Toxin | -0.23 | -0.82 | -0.25 | 1.5 | 9.1 | 2565.12 | outside |
| 429 | 449 | FTGCVIAWNSNNLDSKVGGN | 0.3614 | 20 | ABCPRED | 1 | NON-ALLERGEN | Non-Toxin | -0.05 | -0.1 | -0.35 | 0 | 6.16 | 2096.6 | outside |
| 67 | 81 | AIHVSGTNGTKRFD | 0.6856 | 14 | ABCPRED | 1 | NON-ALLERGEN | Non-Toxin | -0.2 | -0.59 | 0.14 | 1.5 | 9.1 | 1502.85 | outside |
| 943 | 963 | SALGKLQDVVNQNAQALNTL | 0.401 | 20 | ABCPRED | 1 | NON-ALLERGEN | Non-Toxin | -0.11 | -0.07 | -0.23 | 0 | 6.19 | 2097.67 | outside |
| 842 | 856 | GDIAARDLICAQKF | 0.7076 | 14 | ABCPRED | 1 | NON-ALLERGEN | Non-Toxin | -0.1 | 0.3 | 0.13 | 0 | 6.28 | 1520.96 | outside |
| 245 | 259 | HRSYLTPGDSSSGW | 0.5947 | 14 | ABCPRED | 1 | NON-ALLERGEN | Non-Toxin | -0.21 | -1.14 | -0.09 | 0.5 | 7.09 | 1549.81 | outside |
| 994 | 1008 | DRLITGRLQSLQTY | -0.4344 | 14 | ABCPRED | 1 | NON-ALLERGEN | Non-Toxin | -0.27 | -0.54 | -0.04 | 1 | 9.1 | 1664.11 | outside |
| 1085 | 1099 | GKAHFPREGVFVSN | 0.1213 | 14 | ABCPRED | 1 | NON-ALLERGEN | Non-Toxin | -0.14 | -0.43 | 0.04 | 1.5 | 9.1 | 1544.93 | outside |
| 535 | 549 | KNKCVNFNFNGLTG | 1.6052 | 14 | ABCPRED | 1 | NON-ALLERGEN | Non-Toxin | -0.16 | -0.51 | -0.21 | 2 | 9.36 | 1555.98 | outside |
| 108 | 122 | TTLDSKTQSLLIVN | 0.8976 | 14 | ABCPRED | 1 | NON-ALLERGEN | Non-Toxin | -0.1 | 0.14 | -0.21 | 0 | 6.19 | 1532.97 | outside |
| 976 | 996 | VLNDILSRLDKVEAEVQIDR | -0.1568 | 20 | ABCPRED | 1 | NON-ALLERGEN | Non-Toxin | -0.23 | -0.17 | 0.53 | -2 | 4.44 | 2325.95 | outside |
| 131 | 145 | CEFQFCNDPFLGVY | 0.007 | 14 | ABCPRED | 1 | NON-ALLERGEN | Non-Toxin | 0.03 | 0.29 | -0.62 | -2 | 3.67 | 1682.09 | outside |
| 952 | 972 | VNQNAQALNTLVKQLSSNFG | 0.1127 | 20 | ABCPRED | 1 | NON-ALLERGEN | Non-Toxin | -0.12 | -0.25 | -0.36 | 1 | 9.11 | 2146.7 | outside |
| 516 | 536 | ELLHAPATVCGPKKSTNLVK | 0.0205 | 20 | ABCPRED | 1 | NON-ALLERGEN | Non-Toxin | -0.12 | -0.09 | 0.04 | 2.5 | 9.25 | 2106.81 | outside |
| 792 | 812 | PPIKDFGGFNFSQILPDPSK | 0.1539 | 20 | ABCPRED | 1 | NON-ALLERGEN | Non-Toxin | -0.08 | -0.47 | 0.01 | 0 | 6.31 | 2204.8 | outside |
| 674 | 694 | YQTQTNSPRRARSVASQSII | 0.0916 | 20 | ABCPRED | 1 | NON-ALLERGEN | Non-Toxin | -0.35 | -0.91 | 0.09 | 3 | 11.72 | 2263.78 | outside |
| 433 | 453 | VIAWNSNNLDSKVGGNYNYL | 0.7955 | 20 | ABCPRED | 1 | NON-ALLERGEN | Non-Toxin | -0.08 | -0.42 | -0.46 | 0 | 6.18 | 2241.75 | outside |
| 587 | 601 | ITPCSFGGVSVITP | 0.9725 | 14 | ABCPRED | 1 | NON-ALLERGEN | Non-Toxin | 0.18 | 1.12 | -0.74 | 0 | 5.85 | 1377.82 | outside |
| 204 | 224 | YSKHTPINLVRDLPQGFSAL | 0.5839 | 20 | ABCPRED | 1 | NON-ALLERGEN | Non-Toxin | -0.11 | -0.23 | -0.25 | 1.5 | 8.94 | 2256.89 | outside |
| 170 | 190 | YVSQPFLMDLEGKQGNFKNL | 0.7661 | 20 | ABCPRED | 1 | NON-ALLERGEN | Non-Toxin | -0.13 | -0.51 | -0.12 | 0 | 6.41 | 2328.98 | outside |
| 732 | 752 | TKTSVDCTMYICGDSTECSN | 0.1878 | 20 | ABCPRED | 1 | NON-ALLERGEN | Non-Toxin | -0.17 | -0.34 | 0.08 | -2 | 4.03 | 2158.64 | outside |
| 377 | 391 | FKCYGVSPTKLNDL | 1.4748 | 14 | ABCPRED | 1 | NON-ALLERGEN | Non-Toxin | -0.12 | -0.18 | -0.13 | 1 | 8.51 | 1585.05 | outside |
| 610 | 630 | VLYQDVNCTEVPVAIHADQL | 0.4447 | 20 | ABCPRED | 1 | NON-ALLERGEN | Non-Toxin | -0.01 | 0.36 | -0.35 | -2.5 | 4.03 | 2227.82 | outside |
| 148 | 170 | NNKSWMESEFRVYSSANNCTFE | 0.222 | 22 | FBCPred | 1 | NON-ALLERGEN | Non-Toxin | -0.26 | -1.05 | 0.07 | -1 | 4.79 | 2644.14 | outside |
| 507 | 527 | PYRVVVLSFELLHAPATVCG | 0.5979 | 20 | ABCPRED | 1 | NON-ALLERGEN | Non-Toxin | 0.09 | 0.98 | -0.64 | 0.5 | 7.06 | 2171.88 | outside |
| 503 | 523 | VGYQPYRVVVLSFELLHAPA | 0.989 | 20 | ABCPRED | 1 | NON-ALLERGEN | Non-Toxin | 0.06 | 0.65 | -0.68 | 0.5 | 7.1 | 2258.95 | outside |
| 848 | 868 | DLICAQKFNGLTVLPPLLTD | 0.4771 | 20 | ABCPRED | 1 | ALLERGEN | Non-Toxin | 0.03 | 0.6 | -0.39 | -1 | 4.21 | 2171.89 | outside |
| 1114 | 1134 | IITTDNTFVSGNCDVVIGIV | 0.5096 | 20 | ABCPRED | 1 | ALLERGEN | Non-Toxin | 0.13 | 1.12 | -0.56 | -2 | 3.57 | 2080.69 | outside |
| 198 | 218 | DGYFKIYSKHTPINLVRDLP | 0.3949 | 20 | ABCPRED | 1 | NON-ALLERGEN | Toxin | -0.16 | -0.51 | -0.06 | 1.5 | 8.76 | 2377.04 | outside |
| 625 | 639 | HADQLTPTWRVYST | 0.5736 | 14 | ABCPRED | 1 | NON-ALLERGEN | Non-Toxin | -0.2 | -0.83 | -0.34 | 0.5 | 7.09 | 1675.03 | outside |
| 216 | 236 | LPQGFSALEPLVDLPIGINI | 0.7166 | 20 | ABCPRED | 1 | NON-ALLERGEN | Non-Toxin | 0.14 | 0.86 | -0.52 | -2 | 3.67 | 2106.81 | outside |
| 975 | 989 | SVLNDILSRLDKVE | -0.2257 | 14 | ABCPRED | 1 | ALLERGEN | Non-Toxin | -0.19 | 0.02 | 0.4 | -1 | 4.56 | 1601.04 | outside |
| 786 | 802 | KQIYKTPPIKDFGGFN | -0.2241 | 16 | ABCPRED | 1 | ALLERGEN | Non-Toxin | -0.17 | -0.85 | 0.07 | 2 | 9.55 | 1853.39 | outside |
| 63 | 83 | TWFHAIHVSGTNGTKRFDNP | 0.3963 | 20 | ABCPRED | 1 | ALLERGEN | Non-Toxin | -0.16 | -0.77 | -0.24 | 2 | 9.11 | 2285.8 | outside |
| 383 | 397 | SPTKLNDLCFTNVY | 1.6982 | 14 | ABCPRED | 1 | NON-ALLERGEN | Non-Toxin | -0.11 | -0.17 | -0.36 | 0 | 6.15 | 1615.03 | outside |
| 432 | 446 | CVIAWNSNNLDSKV | 0.595 | 14 | ABCPRED | 1 | NON-ALLERGEN | Non-Toxin | -0.09 | 0.04 | -0.31 | 0 | 6.16 | 1562.96 | outside |
| 1180 | 1196 | QKEIDRLNEVAKNLNE | 0.0684 | 16 | ABCPRED | 1 | NON-ALLERGEN | Non-Toxin | -0.41 | -1.39 | 0.9 | -1 | 4.88 | 1913.36 | outside |
| 633 | 647 | WRVYSTGSNVFQTR | 0.4314 | 14 | ABCPRED | 1 | NON-ALLERGEN | Non-Toxin | -0.25 | -0.74 | -0.36 | 2 | 10.84 | 1701.07 | outside |
| 416 | 436 | GKIADYNYKLPDDFTGCVIA | 0.5836 | 20 | ABCPRED | 1 | ALLERGEN | Non-Toxin | -0.06 | -0.08 | -0.06 | -1 | 4.43 | 2203.78 | outside |
| 666 | 680 | IGAGICASYQTQTN | 0.7798 | 14 | ABCPRED | 1 | NON-ALLERGEN | Non-Toxin | -0.02 | 0.02 | -0.56 | 0 | 5.84 | 1426.77 | outside |
| 28 | 42 | YTNSFTRGVYYPDK | 0.0251 | 14 | ABCPRED | 1 | NON-ALLERGEN | Non-Toxin | -0.25 | -1.18 | -0.16 | 1 | 8.76 | 1711.05 | outside |
| 369 | 391 | YNSASFSTFKCYGVSPTKLNDL | 0.9508 | 22 | AAP | 1 | NON-ALLERGEN | Non-Toxin | -0.11 | -0.26 | -0.29 | 1 | 8.48 | 2443.03 | outside |
| 899 | 921 | AMQMAYRFNGIGVTQNVLYENQ | 0.8238 | 22 | BCPREDS,FBCPred | 2 | NON-ALLERGEN | Non-Toxin | -0.1 | -0.28 | -0.48 | 0 | 6.35 | 2548.23 | outside |
| 788 | 808 | IYKTPPIKDFGGFNFSQILP | 0.1762 | 20 | ABCPRED | 1 | NON-ALLERGEN | Non-Toxin | 0 | -0.05 | -0.38 | 1 | 8.83 | 2282.97 | outside |
| 73 | 87 | TNGTKRFDNPVLPF | 0.1985 | 14 | ABCPRED | 1 | NON-ALLERGEN | Non-Toxin | -0.21 | -0.74 | 0.02 | 1 | 9.1 | 1606.01 | outside |
| 573 | 587 | TDAVRDPQTLEILD | 0.5785 | 14 | ABCPRED | 1 | NON-ALLERGEN | Non-Toxin | -0.22 | -0.49 | 0.5 | -3 | 3.84 | 1585.93 | outside |
| 347 | 361 | FASVYAWNRKRISN | 0.4597 | 14 | ABCPRED | 1 | NON-ALLERGEN | Non-Toxin | -0.26 | -0.61 | -0.18 | 3 | 11.01 | 1712.13 | outside |
| 981 | 997 | LSRLDKVEAEVQIDRL | 0.0645 | 16 | FBCPred | 1 | NON-ALLERGEN | Non-Toxin | -0.29 | -0.32 | 0.67 | -1 | 4.79 | 1884.4 | outside |
| 560 | 582 | LPFQQFGRDIADTTDAVRDPQT | 0.1838 | 22 | BCPREDS | 1 | NON-ALLERGEN | Toxin | -0.25 | -0.8 | 0.29 | -2 | 4.14 | 2492.01 | outside |
| 711 | 725 | SIAIPTNFTISVTT | 0.8408 | 14 | ABCPRED | 1 | NON-ALLERGEN | Non-Toxin | 0.12 | 0.91 | -0.76 | 0 | 5.88 | 1464.88 | outside |
| 989 | 1009 | AEVQIDRLITGRLQSLQTYV | -0.3501 | 20 | ABCPRED | 1 | NON-ALLERGEN | Non-Toxin | -0.15 | 0.01 | -0.13 | 0 | 6.42 | 2303.96 | outside |
| 1039 | 1059 | RVDFCGKGYHLMSFPQSAPH | 0.2998 | 20 | ABCPRED | 1 | NON-ALLERGEN | Non-Toxin | -0.13 | -0.45 | -0.23 | 2 | 8.54 | 2277.89 | outside |
| 530 | 552 | STNLVKNKCVNFNFNGLTGTGV | 1.0662 | 22 | BCPREDS | 1 | ALLERGEN | Non-Toxin | -0.08 | -0.05 | -0.36 | 2 | 9.36 | 2327.98 | outside |
| 667 | 689 | GAGICASYQTQTNSPRRARSVA | 0.4539 | 22 | FBCPred | 1 | NON-ALLERGEN | Non-Toxin | -0.27 | -0.6 | 0.05 | 3 | 10.77 | 2294.83 | outside |
| 823 | 837 | FNKVTLADAGFIKQ | 0.6472 | 14 | ABCPRED | 1 | NON-ALLERGEN | Non-Toxin | -0.05 | 0.16 | -0.15 | 1 | 8.94 | 1552.01 | outside |
| 1053 | 1059 | PQSAPH | 0.141 | 6 | IEDB | 1 | ALLERGEN | Non-Toxin | -0.21 | -1.48 | -0.08 | 0.5 | 7.1 | 635.75 | outside |
| 599 | 608 | TPGTNTSNQ | 0.5029 | 9 | IEDB | 1 | NON-ALLERGEN | Non-Toxin | -0.3 | -1.71 | -0.03 | 0 | 5.88 | 919.04 | outside |
| 498 | 504 | QPTNGV | 0.0716 | 6 | IEDB | 1 | ALLERGEN | Non-Toxin | -0.15 | -0.92 | -0.25 | 0 | 5.88 | 614.74 | outside |
| 49 | 55 | HSTQDL | 0.4956 | 6 | IEDB | 1 | ALLERGEN | Non-Toxin | -0.29 | -1.32 | 0.13 | -0.5 | 5.09 | 699.8 | outside |
| 1052 | 1059 | FPQSAPH | -0.3069 | 7 | IEDB,literature | 2 | ALLERGEN | Non-Toxin | -0.09 | -0.87 | -0.43 | 0.5 | 7.1 | 782.94 | outside |
| 599 | 607 | TPGTNTSN | 0.5249 | 8 | IEDB | 1 | ALLERGEN | Non-Toxin | -0.25 | -1.49 | -0.06 | 0 | 5.88 | 790.89 | outside |
| 599 | 609 | TPGTNTSNQV | 0.4224 | 10 | ABCPRED,BCEPRED | 2 | NON-ALLERGEN | Non-Toxin | -0.21 | -1.12 | -0.18 | 0 | 5.88 | 1018.19 | outside |
| 370 | 375 | NSASF | short | 5 | ellipro | 1 | ALLERGEN | Non-Toxin | -0.06 | -0.1 | -0.44 | 0 | 5.88 | 524.58 | outside |
| 601 | 611 | GTNTSNQVAV | 0.663 | 10 | ABCPRED | 1 | NON-ALLERGEN | Non-Toxin | -0.11 | -0.29 | -0.34 | 0 | 5.88 | 990.18 | outside |
| 672 | 694 | AVASQSIIA | 0.2729 | 9 | ellipro | 1 | NON-ALLERGEN | Non-Toxin | 0.17 | 1.5 | -0.64 | 0 | 5.88 | 859.11 | outside |
| 120 | 130 | VNNATNVVIK | 0.3623 | 10 | ABCPRED | 1 | ALLERGEN | Non-Toxin | -0.06 | 0.38 | -0.36 | 1 | 9.11 | 1071.39 | outside |
| 807 | 815 | PDPSKPSK | 0.0621 | 8 | IEDB,literature | 2 | ALLERGEN | Non-Toxin | -0.46 | -2.21 | 1.2 | 1 | 8.94 | 855.05 | outside |
| 161 | 168 | SSANNCT | -0.4197 | 7 | IEDB | 1 | ALLERGEN | Non-Toxin | -0.24 | -0.71 | -0.13 | 0 | 5.85 | 695.78 | outside |
| 687 | 694 | VASQSII | -0.0188 | 7 | ellipro,IEDB,literature | 3 | ALLERGEN | Non-Toxin | 0.15 | 1.41 | -0.69 | 0 | 5.88 | 716.93 | outside |
| 250 | 256 | TPGDSS | 0.3268 | 6 | IEDB | 1 | ALLERGEN | Non-Toxin | -0.22 | -1.3 | 0.53 | -1 | 3.8 | 562.6 | outside |
| 614 | 620 | DVNCTE | 2.375 | 6 | IEDB | 1 | ALLERGEN | Non-Toxin | -0.26 | -0.75 | 0.55 | -2 | 3.67 | 679.77 | outside |
| 1174 | 1180 | ASVVNI | 0.8671 | 6 | IEDB,literature | 2 | ALLERGEN | Non-Toxin | 0.19 | 1.73 | -0.8 | 0 | 5.88 | 601.78 | outside |

| | | | | | | | | | | | | | | | |
|---|---|---|---|---|---|---|---|---|---|---|---|---|---|---|---|
| 743 | 752 | CGDSTECSN | 0.1786 | 9 | BCEPRED | 1 | NON-ALLERGEN | Non-Toxin | -0.27 | -0.91 | 0.49 | -2 | 3.67 | 915.02 | outside |
| 68 | 77 | IHVSGTNGT | 0.8621 | 9 | BCEPRED | 1 | ALLERGEN | Non-Toxin | -0.01 | -0.11 | -0.46 | 0.5 | 7.1 | 885.08 | outside |
| 743 | 753 | CGDSTECSNL | 0.3973 | 10 | ABCPRED | 1 | NON-ALLERGEN | Non-Toxin | -0.19 | -0.44 | 0.26 | -2 | 3.67 | 1028.2 | outside |
| 744 | 751 | GDSTECS | 0.111 | 7 | IEDB | 1 | ALLERGEN | Non-Toxin | -0.26 | -1.03 | 0.74 | -2 | 3.67 | 697.75 | outside |
| 1096 | 1102 | VSNGTH | 0.7888 | 6 | IEDB | 1 | ALLERGEN | Non-Toxin | -0.13 | -0.73 | -0.32 | 0.5 | 7.1 | 613.71 | outside |
| 806 | 815 | LPDPSKPSK | -0.1058 | 9 | IEDB | 1 | ALLERGEN | Non-Toxin | -0.35 | -1.54 | 0.87 | 1 | 8.94 | 968.23 | outside |
| 743 | 751 | CGDSTECS | 0.0977 | 8 | IEDB | 1 | ALLERGEN | Non-Toxin | -0.23 | -0.59 | 0.53 | -2 | 3.67 | 800.9 | outside |
| 880 | 886 | GTITSG | -0.0576 | 6 | IEDB | 1 | ALLERGEN | Non-Toxin | 0.07 | 0.25 | -0.38 | 0 | 5.88 | 534.65 | outside |
| 667 | 675 | GAGICASY | 0.521 | 8 | COVIDep,IEDB,literature | 3 | NON-ALLERGEN | Non-Toxin | 0.17 | 0.96 | -0.72 | 0 | 5.84 | 740.93 | outside |
| 250 | 258 | TPGDSSSG | 0.0924 | 8 | BCEPRED | 1 | ALLERGEN | Non-Toxin | -0.18 | -1.12 | 0.44 | -1 | 3.8 | 706.76 | outside |
| 430 | 436 | TGCVIA | 0.4716 | 6 | IEDB,literature | 2 | ALLERGEN | Non-Toxin | 0.26 | 1.98 | -0.87 | 0 | 5.85 | 562.76 | outside |
| 3 | 5 | VF | short | 2 | literature | 1 | NON-ALLERGEN | Non-Toxin | 0.57 | 3.5 | -2 | 0 | 5.88 | 264.34 | outside |
| 687 | 692 | VASQS | short | 5 | ellipro,literature | 2 | ALLERGEN | Non-Toxin | -0.08 | 0.18 | -0.24 | 0 | 5.88 | 490.57 | outside |
| 655 | 661 | HVNNSY | 1.0629 | 6 | IEDB | 1 | ALLERGEN | Non-Toxin | -0.23 | -1.35 | -0.6 | 0.5 | 7.09 | 732.83 | outside |
| 653 | 660 | AEHVNNS | 0.8168 | 7 | ellipro | 1 | NON-ALLERGEN | Non-Toxin | -0.25 | -1.21 | 0.17 | -0.5 | 5.25 | 769.86 | outside |
| 475 | 485 | AGSTPCNGVE | 0.0244 | 10 | IEDB | 1 | ALLERGEN | Non-Toxin | -0.06 | -0.24 | 0.01 | -1 | 4 | 934.12 | outside |
| 936 | 948 | DSLSSTASALGK | 0.5805 | 12 | ABCPRED | 1 | NON-ALLERGEN | Non-Toxin | -0.11 | -0.04 | 0.18 | 0 | 6.19 | 1136.38 | outside |
| 665 | 675 | PIGAGICASY | 0.8411 | 10 | ABCPRED | 1 | NON-ALLERGEN | Non-Toxin | 0.2 | 1.06 | -0.76 | 0 | 5.84 | 951.24 | outside |
| 616 | 643 | NCTEVTGSNVF | 0.1838 | 11 | ellipro,literature | 2 | NON-ALLERGEN | Non-Toxin | -0.06 | 0.05 | -0.33 | -1 | 4 | 1170.41 | outside |
| 594 | 606 | GVSVITPGTNTS | 0.3921 | 12 | ABCPRED | 1 | NON-ALLERGEN | Non-Toxin | 0.03 | 0.28 | -0.43 | 0 | 5.88 | 1132.41 | outside |
| 114 | 126 | TQSLLIVNNATN | 0.3674 | 12 | ABCPRED | 1 | ALLERGEN | Non-Toxin | -0.05 | 0.16 | -0.59 | 0 | 5.88 | 1287.62 | outside |
| 568 | 578 | DIADTTDAVR | 0.8147 | 10 | ABCPRED | 1 | ALLERGEN | Non-Toxin | -0.25 | -0.41 | 0.69 | -2 | 3.94 | 1076.25 | outside |
| 614 | 621 | DVNCTEV | 2.2015 | 7 | IEDB | 1 | NON-ALLERGEN | Non-Toxin | -0.15 | -0.04 | 0.26 | -2 | 3.67 | 778.92 | outside |
| 593 | 605 | GGVSVITPGTNT | 0.2822 | 12 | ABCPRED | 1 | NON-ALLERGEN | Non-Toxin | 0.07 | 0.31 | -0.46 | 0 | 5.88 | 1102.39 | outside |
| 647 | 654 | AGCLIGA | 0.1743 | 7 | IEDB,literature | 2 | ALLERGEN | Non-Toxin | 0.3 | 1.94 | -0.8 | 0 | 5.85 | 603.83 | outside |
| 280 | 287 | NENGTIT | 0.6938 | 7 | ellipro,literature | 2 | NON-ALLERGEN | Non-Toxin | -0.2 | -1.11 | 0.11 | -1 | 4 | 747.86 | outside |
| 116 | 128 | SLLIVNNATNVV | 0.4935 | 12 | ABCPRED | 1 | NON-ALLERGEN | Non-Toxin | 0.11 | 1.21 | -0.82 | 0 | 5.88 | 1256.65 | outside |
| 115 | 127 | QSLLIVNNATNV | 0.4865 | 12 | ABCPRED | 1 | NON-ALLERGEN | Non-Toxin | 0.01 | 0.57 | -0.68 | 0 | 5.88 | 1285.65 | outside |
| 1225 | 1234 | IAIVMVTIM | 1.1339 | 9 | literature | 1 | NON-ALLERGEN | Non-Toxin | 0.43 | 2.98 | -1.32 | 0 | 5.88 | 990.47 | transmembrane-region |
| 1122 | 1135 | VSGNCDVVIGIVN | 0.7251 | 13 | BCEPRED | 1 | NON-ALLERGEN | Non-Toxin | 0.13 | 1.25 | -0.53 | -1 | 3.8 | 1288.68 | outside |
| 117 | 127 | LLIVNNATNV | 0.4718 | 10 | ABCPRED | 1 | ALLERGEN | Non-Toxin | 0.1 | 1.11 | -0.87 | 0 | 5.88 | 1070.41 | outside |
| 601 | 613 | GTNTSNQVAVLY | 0.6018 | 12 | ABCPRED | 1 | NON-ALLERGEN | Non-Toxin | -0.05 | -0.03 | -0.62 | 0 | 5.88 | 1266.55 | outside |
| 949 | 959 | QDVVNQNAQA | 0.2037 | 10 | IEDB | 1 | ALLERGEN | Non-Toxin | -0.25 | -0.9 | 0 | -1 | 3.8 | 1086.27 | outside |
| 114 | 124 | TQSLLIVNNA | 0.4399 | 10 | ABCPRED | 1 | NON-ALLERGEN | Non-Toxin | 0.02 | 0.61 | -0.69 | 0 | 5.88 | 1072.38 | outside |
| 10 | 22 | LVSSQCVNLTTR | 1.2957 | 12 | ABCPRED | 1 | NON-ALLERGEN | Non-Toxin | -0.15 | 0.33 | -0.37 | 1 | 8.6 | 1320.7 | outside |
| 114 | 128 | TQSLLIVNNATNVV | 0.4297 | 14 | ABCPRED,BCPREDS | 2 | NON-ALLERGEN | Non-Toxin | 0.03 | 0.74 | -0.72 | 0 | 5.88 | 1485.92 | outside |
| 181 | 187 | GKQGNF | 2.1342 | 6 | IEDB | 1 | ALLERGEN | Non-Toxin | -0.25 | -1.48 | 0.15 | 1 | 9.11 | 649.79 | outside |
| 544 | 554 | NGLTGTGVLT | 0.6212 | 10 | ABCPRED | 1 | NON-ALLERGEN | Non-Toxin | 0.09 | 0.5 | -0.61 | 0 | 5.88 | 932.2 | outside |
| 595 | 605 | VSVITPGTNT | 0.1777 | 10 | ABCPRED | 1 | ALLERGEN | Non-Toxin | 0.05 | 0.45 | -0.55 | 0 | 5.88 | 988.25 | outside |
| 28 | 34 | YTNSFT | -0.7122 | 6 | IEDB | 1 | ALLERGEN | Non-Toxin | -0.11 | -0.7 | -0.85 | 0 | 5.88 | 731.83 | outside |
| 594 | 604 | GVSVITPGTN | 0.2538 | 10 | ABCPRED | 1 | NON-ALLERGEN | Non-Toxin | 0.08 | 0.48 | -0.51 | 0 | 5.88 | 944.2 | outside |
| 598 | 608 | ITPGTNTSNQ | 0.941 | 10 | ABCPRED | 1 | NON-ALLERGEN | Non-Toxin | -0.2 | -1.09 | -0.21 | 0 | 5.88 | 1032.22 | outside |
| 597 | 609 | VITPGTNTSNQV | 0.3818 | 12 | ABCPRED,BCPREDS | 2 | NON-ALLERGEN | Non-Toxin | -0.07 | -0.21 | -0.42 | 0 | 5.88 | 1230.52 | outside |
| 118 | 130 | LIVNNATNVVIK | 0.2216 | 12 | ABCPRED | 1 | NON-ALLERGEN | Non-Toxin | 0.06 | 1.01 | -0.6 | 1 | 9.11 | 1297.75 | outside |
| 593 | 609 | GGVSVITPGTNTSNQV | 0.4008 | 16 | ABCPRED | 1 | NON-ALLERGEN | Non-Toxin | -0.02 | 0.01 | -0.39 | 0 | 5.88 | 1530.9 | outside |
| 599 | 611 | TPGTNTSNQVAV | 0.4567 | 12 | ABCPRED,AAP | 2 | NON-ALLERGEN | Non-Toxin | -0.11 | -0.43 | -0.32 | 0 | 5.88 | 1188.43 | outside |
| 1033 | 1039 | VLGQSK | 1.2255 | 6 | IEDB | 1 | ALLERGEN | Non-Toxin | -0.14 | -0.1 | 0.03 | 1 | 9.11 | 630.83 | outside |
| 936 | 942 | DSLSST | 0.7417 | 6 | IEDB | 1 | ALLERGEN | Non-Toxin | -0.19 | -0.47 | 0.28 | -1 | 3.8 | 608.67 | outside |
| 617 | 642 | CTEVNV | 0.4775 | 6 | ellipro | 1 | ALLERGEN | Non-Toxin | -0.05 | 0.53 | -0.2 | -1 | 4 | 663.82 | outside |
| 597 | 611 | VITPGTNTSNQVAV | 0.4277 | 14 | ABCPRED,FBCPred,literature | 3 | NON-ALLERGEN | Non-Toxin | -0.01 | 0.25 | -0.51 | 0 | 5.88 | 1400.76 | outside |
| 115 | 122 | QSLLIVN | 0.8168 | 7 | IEDB,literature | 2 | ALLERGEN | Non-Toxin | 0.11 | 1.21 | -0.89 | 0 | 5.88 | 786.05 | outside |
| 114 | 122 | TQSLLIVN | 0.6517 | 8 | BCEPRED | 1 | ALLERGEN | Non-Toxin | 0.07 | 0.98 | -0.82 | 0 | 5.88 | 887.17 | outside |
| 11 | 21 | VSSQCVNLTT | 1.2691 | 10 | ABCPRED | 1 | NON-ALLERGEN | Non-Toxin | -0.06 | 0.47 | -0.56 | 0 | 5.85 | 1051.32 | outside |
| 97 | 99 | KS | short | 2 | literature | 1 | NON-ALLERGEN | Non-Toxin | -0.68 | -2.35 | 1.65 | 1 | 9.11 | 233.28 | outside |
| 596 | 610 | SVITPGTNTSNQVA | 0.3688 | 14 | BCPREDS | 1 | NON-ALLERGEN | Non-Toxin | -0.06 | -0.11 | -0.38 | 0 | 5.88 | 1388.7 | outside |
| 988 | 993 | EAEVQ | short | 5 | literature | 1 | ALLERGEN | Non-Toxin | -0.23 | -0.9 | 0.84 | -2 | 3.8 | 574.65 | outside |
| 598 | 610 | ITPGTNTSNQVA | 0.8126 | 12 | ABCPRED,FBCPred | 2 | NON-ALLERGEN | Non-Toxin | -0.1 | -0.41 | -0.34 | 0 | 5.88 | 1202.46 | outside |
| 596 | 606 | SVITPGTNTS | 0.2244 | 10 | ABCPRED | 1 | NON-ALLERGEN | Non-Toxin | -0.03 | -0.05 | -0.37 | 0 | 5.88 | 976.19 | outside |
| 93 | 98 | ASTEK | short | 5 | literature | 1 | ALLERGEN | Non-Toxin | -0.38 | -1.42 | 1.08 | 0 | 6.35 | 534.62 | outside |
| 283 | 293 | GTITDAVDCA | 0.557 | 10 | ABCPRED | 1 | NON-ALLERGEN | Non-Toxin | 0.02 | 0.6 | -0.01 | -2 | 3.57 | 965.17 | outside |
| 652 | 660 | GAEHVNNS | 0.8804 | 8 | ellipro | 1 | ALLERGEN | Non-Toxin | -0.2 | -1.11 | 0.15 | -0.5 | 5.25 | 826.93 | outside |
| 10 | 20 | LVSSQCVNLT | 1.3797 | 10 | ABCPRED | 1 | NON-ALLERGEN | Non-Toxin | 0.02 | 0.92 | -0.7 | 0 | 5.85 | 1063.38 | outside |
| 595 | 607 | VSVITPGTNTSN | 0.3719 | 12 | ABCPRED | 1 | NON-ALLERGEN | Non-Toxin | -0.04 | 0.02 | -0.42 | 0 | 5.88 | 1189.46 | outside |
| 47 | 53 | VLHSTQ | 0.8948 | 6 | IEDB | 1 | NON-ALLERGEN | Non-Toxin | -0.08 | -0.03 | -0.62 | 0.5 | 7.1 | 683.85 | outside |
| 593 | 603 | GGVSVITPGT | 0.2377 | 10 | ABCPRED | 1 | NON-ALLERGEN | Non-Toxin | 0.16 | 0.79 | -0.53 | 0 | 5.88 | 887.15 | outside |
| 147 | 153 | KNNKSW | 0.4346 | 6 | IEDB | 1 | ALLERGEN | Non-Toxin | -0.56 | -2.75 | 0.55 | 2 | 10.02 | 775.94 | outside |
| 75 | 78 | TTKR | short | 4 | literature | 1 | NON-ALLERGEN | Non-Toxin | -0.8 | -2.45 | 1.3 | 2 | 11.01 | 504.63 | outside |
| 281 | 287 | ENGTIT | 0.9257 | 6 | ellipro | 1 | ALLERGEN | Non-Toxin | -0.12 | -0.72 | 0.1 | -1 | 4 | 633.74 | outside |
| 116 | 126 | SLLIVNNATN | 0.4332 | 10 | ABCPRED | 1 | NON-ALLERGEN | Non-Toxin | 0.02 | 0.61 | -0.69 | 0 | 5.88 | 1058.35 | outside |
| 597 | 606 | VITPGTNTS | 0.2753 | 9 | BCEPRED | 1 | ALLERGEN | Non-Toxin | -0.01 | 0.03 | -0.44 | 0 | 5.88 | 889.1 | outside |
| 282 | 294 | NGTITDAVDCAL | 0.5264 | 12 | ABCPRED | 1 | NON-ALLERGEN | Non-Toxin | 0.01 | 0.53 | -0.14 | -2 | 3.57 | 1192.47 | outside |
| 206 | 210 | KHTP | short | 4 | literature | 1 | ALLERGEN | Non-Toxin | -0.44 | -2.35 | 0.53 | 1.5 | 9.11 | 481.6 | outside |
| 519 | 529 | HAPATVCGPK | -0.3263 | 10 | ABCPRED | 1 | NON-ALLERGEN | Non-Toxin | -0.06 | -0.11 | -0.14 | 1.5 | 8.57 | 980.28 | outside |
| 580 | 584 | QTLE | short | 4 | literature | 1 | NON-ALLERGEN | Non-Toxin | -0.24 | -0.98 | 0.25 | -1 | 4 | 489.58 | outside |
| 596 | 612 | SVITPGTNTSNQVAVL | 0.4077 | 16 | ABCPRED | 1 | ALLERGEN | Non-Toxin | 0.01 | 0.41 | -0.54 | 0 | 5.88 | 1601.03 | outside |
| 1123 | 1133 | SGNCDVVIGI | 0.7421 | 10 | IEDB,literature | 2 | NON-ALLERGEN | Non-Toxin | 0.13 | 1.13 | -0.41 | -1 | 3.8 | 976.26 | outside |
| 597 | 607 | VITPGTNTSN | 0.4217 | 10 | ABCPRED,IEDB | 2 | NON-ALLERGEN | Non-Toxin | -0.07 | -0.32 | -0.38 | 0 | 5.88 | 1003.22 | outside |

| | | | | | | | | | | | | | | | |
|---|---|---|---|---|---|---|---|---|---|---|---|---|---|---|---|
| 617 | 642 | CTEVSNV | 1.3891 | 7 | ellipro | 1 | NON-ALLERGEN | Non-Toxin | -0.08 | 0.34 | -0.13 | -1 | 4 | 750.91 | outside |
| 282 | 288 | NGTITD | 1.1184 | 6 | IEDB | 1 | ALLERGEN | Non-Toxin | -0.14 | -0.72 | 0.1 | -1 | 3.8 | 619.71 | outside |
| 280 | 290 | NENGTITDAV | 0.3182 | 10 | ABCPRED | 1 | ALLERGEN | Non-Toxin | -0.13 | -0.53 | 0.18 | -2 | 3.67 | 1033.2 | outside |
| 723 | 731 | TTEILPVS | 1.2071 | 8 | IEDB,literature | 2 | ALLERGEN | Non-Toxin | 0.06 | 0.65 | -0.33 | -1 | 4 | 859.1 | outside |
| 1223 | 1243 | GLIAIVMVTIMLCCMTSCCS | 0.2381 | 20 | ABCPRED | 1 | NON-ALLERGEN | Non-Toxin | 0.24 | 2.18 | -1.03 | 0 | 5.77 | 2093.01 | transmembrane-region_inside |
| 333 | 339 | TNLCPF | 1.2508 | 6 | literature | 1 | ALLERGEN | Non-Toxin | 0.05 | 0.55 | -0.92 | 0 | 5.85 | 693.89 | outside |
| 527 | 531 | PKKS | short | 4 | literature | 1 | ALLERGEN | Non-Toxin | -0.63 | -2.55 | 1.57 | 2 | 10.02 | 458.6 | outside |
| 803 | 815 | SQILPDPSKPSK | -0.0245 | 12 | ABCPRED | 1 | ALLERGEN | Non-Toxin | -0.28 | -1.14 | 0.54 | 1 | 8.94 | 1296.65 | outside |
| 975 | 983 | SVLNDILS | -0.6172 | 8 | BCEPRED | 1 | ALLERGEN | Non-Toxin | 0.06 | 0.96 | -0.39 | -1 | 3.8 | 860.09 | outside |
| 686 | 698 | SVASQSIIAYTM | 0.3779 | 12 | ABCPRED | 1 | NON-ALLERGEN | Non-Toxin | 0.09 | 0.9 | -0.75 | 0 | 5.88 | 1270.63 | outside |
| 160 | 169 | YSSANNCTF | -0.1036 | 9 | BCEPRED | 1 | ALLERGEN | Non-Toxin | -0.12 | -0.39 | -0.63 | 0 | 5.84 | 1006.16 | outside |
| 70 | 79 | VSGTNGTKR | 0.9855 | 9 | BCEPRED | 1 | NON-ALLERGEN | Non-Toxin | -0.36 | -1.19 | 0.47 | 2 | 11.01 | 919.13 | outside |
| 281 | 291 | ENGTITDAVD | 0.4053 | 10 | ABCPRED | 1 | NON-ALLERGEN | Non-Toxin | -0.14 | -0.53 | 0.46 | -3 | 3.5 | 1034.18 | outside |
| 1196 | 1201 | SLIDL | short | 5 | literature | 1 | ALLERGEN | Non-Toxin | 0.16 | 1.56 | -0.42 | -1 | 3.8 | 559.73 | outside |
| 636 | 644 | YSTGSNVF | -0.1992 | 8 | IEDB | 1 | ALLERGEN | Non-Toxin | 0 | -0.06 | -0.74 | 0 | 5.88 | 874.02 | outside |
| 858 | 865 | LTVLPPL | 0.6786 | 7 | IEDB,literature | 2 | NON-ALLERGEN | Non-Toxin | 0.26 | 1.67 | -1.04 | 0 | 5.88 | 752.07 | outside |
| 650 | 660 | LIGAEHVNNS | 0.9735 | 10 | ABCPRED | 1 | ALLERGEN | Non-Toxin | -0.04 | -0.06 | -0.24 | -0.5 | 5.25 | 1053.29 | outside |
| 949 | 963 | QDVVNQNAQALNTL | 0.1966 | 14 | ABCPRED,AAP,BCPREDS | 3 | NON-ALLERGEN | Non-Toxin | -0.16 | -0.4 | -0.27 | -1 | 3.8 | 1527.87 | outside |
| 546 | 558 | LTGTGVLTESNK | 0.8122 | 12 | ABCPRED | 1 | NON-ALLERGEN | Non-Toxin | -0.1 | -0.23 | 0.02 | 0 | 6.35 | 1219.54 | outside |
| 367 | 375 | VLYNSASF | 0.1765 | 8 | IEDB | 1 | ALLERGEN | Non-Toxin | 0.1 | 0.78 | -0.97 | 0 | 5.88 | 900.1 | outside |
| 934 | 946 | IQDSLSSTASAL | 0.439 | 12 | ABCPRED | 1 | NON-ALLERGEN | Non-Toxin | -0.03 | 0.4 | -0.2 | -1 | 3.8 | 1192.45 | outside |
| 477 | 487 | STPCNGVEGF | -0.2614 | 10 | ABCPRED | 1 | NON-ALLERGEN | Non-Toxin | -0.03 | -0.14 | -0.19 | -1 | 4 | 1010.22 | outside |
| 1002 | 1012 | QSLQTYVTQQ | -0.1009 | 10 | IEDB | 1 | NON-ALLERGEN | Non-Toxin | -0.23 | -0.95 | -0.53 | 0 | 5.88 | 1195.45 | outside |
| 627 | 633 | DQLTPT | 0.7329 | 6 | IEDB | 1 | ALLERGEN | Non-Toxin | -0.22 | -1.03 | 0.1 | -1 | 3.8 | 673.8 | outside |
| 1165 | 1182 | DLGDISGINASVVNIQK | 0.8314 | 17 | COVIDep,literature | 2 | NON-ALLERGEN | Non-Toxin | -0.04 | 0.22 | -0.03 | -1 | 4.21 | 1743.21 | outside |
| 704 | 716 | SVAYSNNSIAIP | 0.7544 | 12 | ABCPRED | 1 | NON-ALLERGEN | Non-Toxin | 0.03 | 0.38 | -0.59 | 0 | 5.88 | 1235.52 | outside |
| 590 | 600 | CSFGGVSVIT | 1.0615 | 10 | ABCPRED | 1 | ALLERGEN | Non-Toxin | 0.21 | 1.51 | -0.81 | 0 | 5.85 | 969.26 | outside |
| 670 | 680 | ICASYQTQTN | 0.8252 | 10 | ABCPRED | 1 | ALLERGEN | Non-Toxin | -0.16 | -0.52 | -0.55 | 0 | 5.84 | 1128.36 | outside |
| 754 | 764 | LQYGSFCTQL | 1.4443 | 10 | ABCPRED | 1 | NON-ALLERGEN | Non-Toxin | 0.01 | 0.27 | -0.91 | 0 | 5.84 | 1159.47 | outside |
| 615 | 625 | VNCTEVPVAI | 0.9137 | 10 | ABCPRED | 1 | ALLERGEN | Non-Toxin | 0.11 | 1.21 | -0.5 | -1 | 4 | 1044.37 | outside |
| 280 | 293 | NENGTITDAVDCA | 0.5014 | 13 | IEDB | 1 | NON-ALLERGEN | Non-Toxin | -0.13 | -0.35 | 0.25 | -3 | 3.5 | 1322.54 | outside |
| 1050 | 1062 | MSFPQSAPHGVV | 0.1016 | 12 | FBCPred | 1 | ALLERGEN | Non-Toxin | 0.05 | 0.25 | -0.58 | 0.5 | 7.1 | 1256.61 | outside |
| 65 | 71 | FHAIHV | 1.6766 | 6 | IEDB,literature | 2 | NON-ALLERGEN | Non-Toxin | 0.22 | 1.15 | -1.22 | 1 | 7.26 | 722.93 | outside |
| 474 | 486 | QAGSTPCNGVEG | 0.2104 | 12 | FBCPred | 1 | NON-ALLERGEN | Non-Toxin | -0.1 | -0.53 | 0.02 | -1 | 4 | 1119.34 | outside |
| 595 | 613 | VSVITPGTNTSNQVAVLY | 0.3996 | 18 | ABCPRED | 1 | ALLERGEN | Non-Toxin | 0.04 | 0.52 | -0.69 | 0 | 5.88 | 1863.37 | outside |
| 120 | 132 | VNNATNVVIKVC | -0.0437 | 12 | BCPREDS | 1 | NON-ALLERGEN | Non-Toxin | 0 | 0.88 | -0.51 | 1 | 8.57 | 1273.69 | outside |
| 1003 | 1012 | SLQTYVTQQ | -0.0089 | 9 | literature | 1 | NON-ALLERGEN | Non-Toxin | -0.18 | -0.67 | -0.61 | 0 | 5.88 | 1067.3 | outside |
| 263 | 271 | AAYYVGYL | 0.5218 | 8 | IEDB,literature | 2 | NON-ALLERGEN | Non-Toxin | 0.22 | 0.91 | -1.4 | 0 | 5.87 | 919.15 | outside |
| 906 | 918 | FNGIGVTQNVLY | 0.6165 | 12 | ABCPRED,AAP,BCPREDS | 3 | NON-ALLERGEN | Non-Toxin | 0.1 | 0.52 | -0.93 | 0 | 5.88 | 1324.69 | outside |
| 1227 | 1243 | IVMVTIMLCCMTSCCS | -0.002 | 16 | ABCPRED | 1 | ALLERGEN | Non-Toxin | 0.2 | 2.12 | -1.03 | 0 | 5.77 | 1738.49 | transmembrane-region_inside |
| 7 | 19 | LLPLVSSQCVNL | 1.0673 | 12 | ABCPRED | 1 | NON-ALLERGEN | Non-Toxin | 0.11 | 1.32 | -0.85 | 0 | 5.85 | 1285.75 | outside |
| 279 | 286 | YNENGTI | 0.508 | 7 | IEDB | 1 | ALLERGEN | Non-Toxin | -0.17 | -1.2 | -0.16 | -1 | 4 | 809.93 | outside |
| 949 | 961 | QDVVNQNAQALN | 0.3694 | 12 | BCPREDS | 1 | NON-ALLERGEN | Non-Toxin | -0.22 | -0.72 | -0.13 | -1 | 3.8 | 1313.57 | outside |
| 1166 | 1182 | LGDISGINASVVNIQK | 0.6234 | 16 | ABCPRED | 1 | NON-ALLERGEN | Non-Toxin | 0 | 0.45 | -0.22 | 0 | 6.19 | 1628.11 | outside |
| 1190 | 1194 | AKNL | short | 4 | literature | 1 | ALLERGEN | Non-Toxin | -0.24 | -0.45 | 0.23 | 1 | 9.11 | 444.58 | outside |
| 750 | 764 | SNLLLQYGSFCTQL | 0.7599 | 14 | IEDB,literature | 2 | NON-ALLERGEN | Non-Toxin | 0.02 | 0.43 | -0.87 | 0 | 5.84 | 1587.04 | outside |
| 751 | 761 | NLLLQYGSFC | 1.1023 | 10 | ABCPRED | 1 | NON-ALLERGEN | Non-Toxin | 0.08 | 0.72 | -1.05 | 0 | 5.84 | 1157.5 | outside |
| 704 | 714 | SVAYSNNSIA | 0.8557 | 10 | ABCPRED | 1 | ALLERGEN | Non-Toxin | -0.03 | 0.16 | -0.53 | 0 | 5.88 | 1025.21 | outside |
| 720 | 730 | ISVTTEILPV | 1.1242 | 10 | ABCPRED | 1 | ALLERGEN | Non-Toxin | 0.18 | 1.39 | -0.59 | -1 | 4 | 1071.43 | outside |
| 389 | 397 | DLCFTNVY | 1.8569 | 8 | BCEPRED | 1 | ALLERGEN | Non-Toxin | 0.03 | 0.54 | -0.79 | -1 | 3.8 | 974.2 | outside |
| 1228 | 1244 | VMVTIMLCCMTSCCSC | -0.1507 | 16 | ABCPRED | 1 | NON-ALLERGEN | Non-Toxin | 0.15 | 1.99 | -0.98 | 0 | 5.75 | 1728.46 | transmembrane-region_inside |
| 886 | 891 | WTFGA | short | 5 | literature | 1 | ALLERGEN | Non-Toxin | 0.24 | 0.52 | -1.36 | 0 | 5.88 | 580.7 | outside |
| 1163 | 1181 | DVDLGDISGINASVVNIQ | 1.3153 | 18 | FBCPred | 1 | ALLERGEN | Non-Toxin | 0.01 | 0.46 | -0.11 | -3 | 3.43 | 1829.27 | outside |
| 1222 | 1240 | AGLIAIVMVTIMLCCMTS | 0.658 | 18 | ABCPRED | 1 | NON-ALLERGEN | Non-Toxin | 0.29 | 2.29 | -1.08 | 0 | 5.82 | 1870.71 | transmembrane-region_inside |
| 754 | 766 | LQYGSFCTQLNR | 1.383 | 12 | BCPREDS | 1 | ALLERGEN | Non-Toxin | -0.19 | -0.44 | -0.49 | 1 | 8.57 | 1429.79 | outside |
| 1046 | 1051 | GYHLM | short | 5 | literature | 1 | ALLERGEN | Non-Toxin | 0.11 | 0.16 | -1.18 | 0.5 | 7.09 | 619.81 | outside |
| 1223 | 1239 | GLIAIVMVTIMLCCMT | 0.6528 | 16 | ABCPRED | 1 | ALLERGEN | Non-Toxin | 0.33 | 2.51 | -1.2 | 0 | 5.82 | 1712.53 | transmembrane-region_inside |
| 721 | 731 | SVTTEILPVS | 0.9195 | 10 | ABCPRED | 1 | ALLERGEN | Non-Toxin | 0.08 | 0.86 | -0.38 | -1 | 4 | 1045.34 | outside |
| 668 | 682 | AGICASYQTQTNSP | 0.6263 | 14 | BCPREDS | 1 | NON-ALLERGEN | Non-Toxin | -0.11 | -0.44 | -0.41 | 0 | 5.84 | 1440.74 | outside |
| 666 | 682 | IGAGICASYQTQTNSP | 0.6076 | 16 | ABCPRED | 1 | NON-ALLERGEN | Non-Toxin | -0.04 | -0.13 | -0.47 | 0 | 5.84 | 1610.99 | outside |
| 803 | 809 | SQILPD | -0.1542 | 6 | IEDB,literature | 2 | ALLERGEN | Non-Toxin | -0.08 | -0.18 | -0.02 | -1 | 3.8 | 671.83 | outside |
| 364 | 372 | DYSVLYNS | 0.1757 | 8 | BCEPRED | 1 | ALLERGEN | Non-Toxin | -0.1 | -0.4 | -0.51 | -1 | 3.8 | 960.11 | outside |
| 1030 | 1038 | SECVLGQS | -0.011 | 8 | IEDB,literature | 2 | ALLERGEN | Non-Toxin | -0.07 | 0.19 | -0.06 | -1 | 4 | 822.01 | outside |
| 1165 | 1179 | DLGDISGINASVVN | 0.7884 | 14 | BCPREDS | 1 | NON-ALLERGEN | Non-Toxin | 0.03 | 0.47 | -0.14 | -2 | 3.57 | 1373.69 | outside |
| 494 | 504 | SYGFQPTNGV | 1.0549 | 10 | ABCPRED | 1 | NON-ALLERGEN | Non-Toxin | -0.03 | -0.52 | -0.6 | 0 | 5.88 | 1069.28 | outside |
| 288 | 296 | AVDCALDP | 0.773 | 8 | IEDB,literature | 2 | NON-ALLERGEN | Non-Toxin | 0.01 | 0.69 | 0.09 | -2 | 3.57 | 802.99 | outside |
| 1228 | 1246 | VMVTIMLCCMTSCCSCLK | 0.1367 | 18 | ABCPRED | 1 | NON-ALLERGEN | Non-Toxin | 0.1 | 1.77 | -0.81 | 1 | 8.1 | 1969.83 | transmembrane-region_inside |
| 1079 | 1086 | PAICHDG | -1.01 | 7 | IEDB,literature | 2 | ALLERGEN | Non-Toxin | 0 | 0.01 | -0.11 | -0.5 | 5.09 | 711.88 | outside |
| 9 | 19 | PLVSSQCVNL | 1.0958 | 10 | ABCPRED | 1 | ALLERGEN | Non-Toxin | 0.03 | 0.83 | -0.66 | 0 | 5.85 | 1059.39 | outside |
| 1164 | 1180 | VDLGDISGINASVVNI | 1.0784 | 16 | ABCPRED | 1 | NON-ALLERGEN | Non-Toxin | 0.1 | 0.96 | -0.32 | -2 | 3.57 | 1586.02 | outside |
| 525 | 533 | CGPKKSTN | -0.2855 | 8 | BCEPRED | 1 | NON-ALLERGEN | Non-Toxin | -0.39 | -1.54 | 0.64 | 2 | 9.36 | 834.06 | outside |
| 122 | 132 | NATNVVIKVC | 0.0919 | 10 | ABCPRED | 1 | NON-ALLERGEN | Non-Toxin | 0.01 | 0.98 | -0.48 | 1 | 8.57 | 1060.42 | outside |
| 1229 | 1245 | MVTIMLCCMTSCCSCL | -0.1171 | 16 | ABCPRED | 1 | ALLERGEN | Non-Toxin | 0.15 | 1.97 | -1 | 0 | 5.75 | 1742.49 | transmembrane-region_inside |
| 445 | 455 | VGGNYNYLYR | 0.3204 | 10 | ABCPRED | 1 | ALLERGEN | Non-Toxin | -0.16 | -0.82 | -0.68 | 1 | 8.83 | 1218.48 | outside |
| 493 | 503 | QSYGFQPTNG | 0.9097 | 10 | ABCPRED | 1 | ALLERGEN | Non-Toxin | -0.16 | -1.29 | -0.43 | 0 | 5.88 | 1098.28 | outside |
| 8 | 20 | LPLVSSQCVNLT | 1.4488 | 12 | ABCPRED | 1 | ALLERGEN | Non-Toxin | 0.05 | 0.95 | -0.73 | 0 | 5.85 | 1273.69 | outside |
| 669 | 681 | GICASYQTQTNS | 0.7595 | 12 | ABCPRED | 1 | NON-ALLERGEN | Non-Toxin | -0.14 | -0.53 | -0.43 | 0 | 5.84 | 1272.52 | outside |

| | | | | | | | | | | | | | | | |
|---|---|---|---|---|---|---|---|---|---|---|---|---|---|---|---|
| 617 | 627 | CTEVPVAIHA | 0.1753 | 10 | ABCPRED | 1 | ALLERGEN | Non-Toxin | 0.11 | 1 | -0.47 | -0.5 | 5.25 | 1039.35 | outside |
| 1227 | 1245 | IVMVTIMLCCMTSCCSCL | 0.0872 | 18 | ABCPRED | 1 | NON-ALLERGEN | Non-Toxin | 0.21 | 2.23 | -1.07 | 0 | 5.75 | 1954.82 | transmembrane-region_inside |
| 372 | 379 | ASFSTFK | 0.0865 | 7 | literature | 1 | ALLERGEN | Non-Toxin | -0.05 | 0.17 | -0.33 | 1 | 9.11 | 786.96 | outside |
| 160 | 170 | YSSANNCTFE | -0.0896 | 10 | ABCPRED | 1 | ALLERGEN | Non-Toxin | -0.17 | -0.7 | -0.27 | -1 | 4 | 1135.29 | outside |
| 966 | 973 | LSSNFGA | 0.6114 | 7 | IEDB | 1 | ALLERGEN | Non-Toxin | 0.06 | 0.41 | -0.57 | 0 | 5.88 | 694.83 | outside |
| 673 | 683 | SYQTQTNSPR | -0.0008 | 10 | ABCPRED | 1 | NON-ALLERGEN | Non-Toxin | -0.47 | -2.09 | 0.11 | 1 | 9.1 | 1181.36 | outside |
| 6 | 18 | VLLPLVSSQCVN | 0.6205 | 12 | ABCPRED | 1 | NON-ALLERGEN | Non-Toxin | 0.11 | 1.36 | -0.82 | 0 | 5.85 | 1271.72 | outside |
| 668 | 680 | AGICASYQTQTN | 0.8504 | 12 | ABCPRED | 1 | NON-ALLERGEN | Non-Toxin | -0.1 | -0.32 | -0.5 | 0 | 5.84 | 1256.52 | outside |
| 1002 | 1014 | QSLQTYVTQQLI | -0.2739 | 12 | BCPREDS | 1 | ALLERGEN | Non-Toxin | -0.09 | -0.1 | -0.74 | 0 | 5.88 | 1421.81 | outside |
| 592 | 600 | FGGVSVIT | 0.7715 | 8 | IEDB,literature | 2 | NON-ALLERGEN | Non-Toxin | 0.29 | 1.68 | -0.93 | 0 | 5.88 | 779.02 | outside |
| 495 | 504 | YGFQPTNGV | 1.0509 | 9 | literature | 1 | ALLERGEN | Non-Toxin | -0.01 | -0.49 | -0.7 | 0 | 5.88 | 982.19 | outside |
| 735 | 742 | SVDCTMY | 1.0932 | 7 | IEDB,literature | 2 | ALLERGEN | Non-Toxin | -0.04 | 0.33 | -0.46 | -1 | 3.8 | 818.01 | outside |
| 873 | 881 | YTSALLAG | 0.3798 | 8 | IEDB,literature | 2 | ALLERGEN | Non-Toxin | 0.16 | 1 | -0.88 | 0 | 5.88 | 795.01 | outside |
| 666 | 678 | IGAGICASYQTQ | 0.8956 | 12 | ABCPRED | 1 | NON-ALLERGEN | Toxin | 0.04 | 0.37 | -0.63 | 0 | 5.84 | 1211.53 | outside |
| 205 | 211 | SKHTPI | 0.7317 | 6 | IEDB | 1 | ALLERGEN | Non-Toxin | -0.21 | -0.95 | 0.1 | 1.5 | 9.11 | 681.87 | outside |
| 649 | 659 | CLIGAEHVNN | 1.1765 | 10 | ABCPRED | 1 | ALLERGEN | Non-Toxin | -0.01 | 0.27 | -0.37 | -0.5 | 5.25 | 1069.35 | outside |
| 1001 | 1014 | LQSLQTYVTQQLI | -0.1517 | 13 | BCEPRED | 1 | NON-ALLERGEN | Non-Toxin | -0.04 | 0.2 | -0.82 | 0 | 5.88 | 1534.99 | outside |
| 753 | 765 | LLQYGSFCTQLN | 0.9501 | 12 | AAP | 1 | NON-ALLERGEN | Non-Toxin | 0 | 0.25 | -0.89 | 0 | 5.84 | 1386.77 | outside |
| 526 | 534 | GPKKSTNL | 0.7072 | 8 | BCEPRED | 1 | ALLERGEN | Non-Toxin | -0.33 | -1.38 | 0.54 | 2 | 10.02 | 844.09 | outside |
| 687 | 697 | VASQSIIAYT | 0.3132 | 10 | ABCPRED | 1 | ALLERGEN | Non-Toxin | 0.11 | 0.97 | -0.8 | 0 | 5.88 | 1052.33 | outside |
| 284 | 296 | TITDAVDCALDP | 0.6497 | 12 | ABCPRED | 1 | NON-ALLERGEN | Non-Toxin | -0.02 | 0.42 | 0.09 | -3 | 3.43 | 1233.51 | outside |
| 622 | 632 | VAIHADQLTP | 0.2877 | 10 | ABCPRED | 1 | ALLERGEN | Non-Toxin | 0.02 | 0.36 | -0.38 | -0.5 | 5.09 | 1064.35 | outside |
| 604 | 614 | TSNQVAVLYQ | 0.4642 | 10 | ABCPRED | 1 | NON-ALLERGEN | Non-Toxin | -0.06 | 0.07 | -0.71 | 0 | 5.88 | 1122.39 | outside |
| 755 | 765 | QYGSFCTQLN | 1.3057 | 10 | ABCPRED | 1 | NON-ALLERGEN | Non-Toxin | -0.11 | -0.46 | -0.71 | 0 | 5.84 | 1160.41 | outside |
| 539 | 555 | VNFNFNGLTGTGVLTE | 1.1691 | 16 | ABCPRED | 1 | NON-ALLERGEN | Non-Toxin | 0.05 | 0.27 | -0.58 | -1 | 4 | 1683.1 | outside |
| 94 | 101 | STEKSNI | 0.6662 | 7 | IEDB | 1 | ALLERGEN | Non-Toxin | -0.33 | -1.24 | 0.66 | 0 | 6.35 | 777.92 | outside |
| 260 | 272 | AGAAAYYVGYLQ | 0.654 | 12 | ABCPRED | 1 | NON-ALLERGEN | Non-Toxin | 0.15 | 0.58 | -1 | 0 | 5.87 | 1246.55 | outside |
| 750 | 766 | SNLLLQYGSFCTQLNR | 0.8057 | 16 | ABCPRED | 1 | ALLERGEN | Non-Toxin | -0.14 | -0.13 | -0.56 | 1 | 8.57 | 1857.36 | outside |
| 541 | 551 | FNFNGLTGTG | 1.0567 | 10 | ABCPRED | 1 | NON-ALLERGEN | Non-Toxin | 0.06 | -0.02 | -0.72 | 0 | 5.88 | 1027.25 | outside |
| 364 | 376 | DYSVLYNSASFS | 0.5166 | 12 | ABCPRED,FBCPred | 2 | NON-ALLERGEN | Non-Toxin | -0.04 | -0.02 | -0.54 | -1 | 3.8 | 1352.57 | outside |
| 437 | 443 | NSNNLD | 1.1859 | 6 | IEDB | 1 | ALLERGEN | Non-Toxin | -0.4 | -1.83 | 0.35 | -1 | 3.8 | 675.73 | outside |
| 742 | 750 | ICGDSTEC | -0.2546 | 8 | BCEPRED | 1 | ALLERGEN | Non-Toxin | -0.1 | 0.07 | 0.26 | -2 | 3.67 | 826.99 | outside |
| 719 | 731 | TISVTTEILPVS | 1.036 | 12 | ABCPRED,FBCPred | 2 | NON-ALLERGEN | Non-Toxin | 0.11 | 1.03 | -0.5 | -1 | 4 | 1259.64 | outside |
| 495 | 501 | YGFQPT | 1.6231 | 6 | IEDB | 1 | ALLERGEN | Non-Toxin | -0.02 | -0.78 | -0.83 | 0 | 5.88 | 711.85 | outside |
| 648 | 660 | GCLIGAEHVNNS | 0.8445 | 12 | ABCPRED | 1 | NON-ALLERGEN | Non-Toxin | -0.01 | 0.13 | -0.28 | -0.5 | 5.25 | 1213.51 | outside |
| 94 | 100 | STEKSN | 1.0006 | 6 | IEDB | 1 | ALLERGEN | Non-Toxin | -0.51 | -2.2 | 1.07 | 0 | 6.35 | 664.74 | outside |
| 756 | 766 | YGSFCTQLNR | 0.9565 | 10 | ABCPRED | 1 | NON-ALLERGEN | Non-Toxin | -0.22 | -0.56 | -0.43 | 1 | 8.57 | 1188.46 | outside |
| 119 | 131 | IVNNATNVVIKV | 0.2468 | 12 | FBCPred | 1 | ALLERGEN | Non-Toxin | 0.06 | 1.04 | -0.58 | 1 | 9.11 | 1283.72 | outside |
| 836 | 842 | QYGDCL | 0.1281 | 6 | IEDB | 1 | ALLERGEN | Non-Toxin | -0.11 | -0.4 | -0.32 | -1 | 3.8 | 697.84 | outside |
| 367 | 379 | VLYNSASFSTFK | 0.0249 | 12 | ABCPRED | 1 | NON-ALLERGEN | Non-Toxin | -0.01 | 0.3 | -0.62 | 1 | 8.94 | 1363.69 | outside |
| 283 | 295 | GTITDAVDCALD | 0.4676 | 12 | ABCPRED | 1 | NON-ALLERGEN | Non-Toxin | 0 | 0.53 | 0.09 | -3 | 3.43 | 1193.45 | outside |
| 540 | 552 | NFNFNGLTGTGV | 1.3547 | 12 | ABCPRED,BCPREDS | 2 | ALLERGEN | Non-Toxin | 0.04 | 0.04 | -0.71 | 0 | 5.88 | 1240.52 | outside |
| 365 | 377 | YSVLYNSASFST | 0.2038 | 12 | ABCPRED | 1 | NON-ALLERGEN | Non-Toxin | 0.01 | 0.22 | -0.83 | 0 | 5.87 | 1338.59 | outside |
| 722 | 732 | VTTEILPVSM | 1.0712 | 10 | ABCPRED | 1 | ALLERGEN | Non-Toxin | 0.13 | 1.13 | -0.54 | -1 | 4 | 1089.46 | outside |
| 262 | 272 | AAAYYVGYLQ | 0.4697 | 10 | ABCPRED | 1 | NON-ALLERGEN | Non-Toxin | 0.14 | 0.56 | -1.15 | 0 | 5.87 | 1118.39 | outside |
| 647 | 657 | AGCLIGAEHV | 0.6223 | 10 | ABCPRED | 1 | ALLERGEN | Non-Toxin | 0.16 | 1.11 | -0.46 | -0.5 | 5.25 | 969.27 | outside |
| 1163 | 1179 | DVDLGDISGINASVVN | 1.1317 | 16 | ABCPRED | 1 | NON-ALLERGEN | Non-Toxin | 0.01 | 0.46 | -0.02 | -3 | 3.43 | 1587.94 | outside |
| 871 | 883 | AQYTSALLAGTI | 0.5417 | 12 | ABCPRED | 1 | NON-ALLERGEN | Non-Toxin | 0.12 | 0.84 | -0.79 | 0 | 5.88 | 1208.55 | outside |
| 669 | 679 | GICASYQTQT | 0.9671 | 10 | ABCPRED | 1 | NON-ALLERGEN | Non-Toxin | -0.08 | -0.21 | -0.57 | 0 | 5.84 | 1071.31 | outside |
| 368 | 378 | LYNSASFSTF | 0.0707 | 10 | ABCPRED | 1 | NON-ALLERGEN | Non-Toxin | 0.04 | 0.33 | -0.89 | 0 | 5.88 | 1136.35 | outside |
| 46 | 56 | SVLHSTQDLF | 0.2151 | 10 | ABCPRED | 1 | ALLERGEN | Non-Toxin | -0.03 | 0.21 | -0.47 | -0.5 | 5.09 | 1146.41 | outside |
| 926 | 932 | QFNSAI | -0.1194 | 6 | IEDB | 1 | ALLERGEN | Non-Toxin | 0 | 0.22 | -0.68 | 0 | 5.88 | 678.82 | outside |
| 666 | 676 | IGAGICASYQ | 0.8225 | 10 | ABCPRED | 1 | ALLERGEN | Non-Toxin | 0.14 | 0.87 | -0.74 | 0 | 5.84 | 982.26 | outside |
| 636 | 646 | YSTGSNVFQT | 0.0167 | 10 | ABCPRED | 1 | NON-ALLERGEN | Non-Toxin | -0.09 | -0.47 | -0.61 | 0 | 5.88 | 1103.29 | outside |
| 173 | 176 | SRTL | short | 4 | literature | 1 | ALLERGEN | Non-Toxin | -0.42 | -0.55 | 0.27 | 1 | 10.11 | 475.59 | outside |
| 602 | 614 | TNTSNQVAVLYQ | 0.5046 | 12 | ABCPRED | 1 | ALLERGEN | Non-Toxin | -0.12 | -0.29 | -0.61 | 0 | 5.88 | 1337.63 | outside |
| 5 | 17 | LVLLPLVSSQCV | 0.7049 | 12 | ABCPRED | 1 | ALLERGEN | Non-Toxin | 0.21 | 1.97 | -0.99 | 0 | 5.85 | 1270.78 | outside |
| 706 | 714 | AYSNNSIA | 0.5473 | 8 | BCEPRED | 1 | ALLERGEN | Non-Toxin | -0.07 | -0.22 | -0.51 | 0 | 5.88 | 838.97 | outside |
| 935 | 947 | QDSLSSTASALG | 0.6803 | 12 | ABCPRED | 1 | NON-ALLERGEN | Non-Toxin | -0.08 | -0.01 | -0.05 | -1 | 3.8 | 1136.34 | outside |
| 635 | 645 | VYSTGSNVFQ | -0.2512 | 10 | ABCPRED | 1 | NON-ALLERGEN | Non-Toxin | -0.02 | 0.02 | -0.72 | 0 | 5.88 | 1101.32 | outside |
| 550 | 558 | GVLTESNK | 0.7779 | 8 | BCEPRED,IEDB | 2 | ALLERGEN | Non-Toxin | -0.2 | -0.6 | 0.35 | 0 | 6.35 | 847.05 | outside |
| 662 | 674 | CDIPIGAGICAS | 0.6389 | 12 | ABCPRED | 1 | NON-ALLERGEN | Non-Toxin | 0.17 | 1.28 | -0.43 | -1 | 3.8 | 1119.48 | outside |
| 973 | 980 | ISSVLND | 0.0414 | 7 | IEDB,literature | 2 | ALLERGEN | Non-Toxin | -0.01 | 0.56 | -0.19 | -1 | 3.8 | 746.91 | outside |
| 67 | 79 | AIHVSGTNGTKR | 0.736 | 12 | ABCPRED | 1 | NON-ALLERGEN | Non-Toxin | -0.22 | -0.63 | 0.12 | 2.5 | 11.01 | 1240.56 | outside |
| 263 | 273 | AAYYVGYLQP | 0.6073 | 10 | ABCPRED | 1 | ALLERGEN | Non-Toxin | 0.1 | 0.22 | -1.1 | 0 | 5.87 | 1144.43 | outside |
| 934 | 944 | IQDSLSSTAS | 0.4491 | 10 | IEDB | 1 | NON-ALLERGEN | Non-Toxin | -0.11 | -0.08 | -0.01 | -1 | 3.8 | 1008.18 | outside |
| 588 | 595 | TPCSFGG | 1.6632 | 7 | IEDB | 1 | ALLERGEN | Non-Toxin | 0.07 | 0.2 | -0.51 | 0 | 5.85 | 667.82 | outside |
| 706 | 716 | AYSNNSIAIP | 0.5035 | 10 | ABCPRED | 1 | ALLERGEN | Non-Toxin | 0.01 | 0.11 | -0.59 | 0 | 5.88 | 1049.28 | outside |
| 475 | 487 | AGSTPCNGVEGF | -0.1412 | 12 | ABCPRED,BCPREDS | 2 | ALLERGEN | Non-Toxin | 0.01 | 0 | -0.2 | -1 | 4 | 1138.38 | outside |
| 757 | 765 | GSFCTQLN | 0.8144 | 8 | COVIDep,literature | 2 | ALLERGEN | Non-Toxin | -0.05 | 0.02 | -0.62 | 0 | 5.85 | 869.07 | outside |
| 542 | 552 | NFNGLTGTGV | 0.7827 | 10 | ABCPRED | 1 | ALLERGEN | Non-Toxin | 0.05 | 0.12 | -0.62 | 0 | 5.88 | 979.21 | outside |
| 742 | 752 | ICGDSTECSN | -0.0707 | 10 | ABCPRED | 1 | ALLERGEN | Non-Toxin | -0.17 | -0.37 | 0.26 | -2 | 3.67 | 1028.2 | outside |
| 159 | 169 | VYSSANNCTF | -0.1475 | 10 | ABCPRED | 1 | ALLERGEN | Non-Toxin | -0.05 | 0.07 | -0.72 | 0 | 5.84 | 1105.31 | outside |
| 705 | 715 | VAYSNNSIAI | 1.0545 | 10 | IEDB | 1 | NON-ALLERGEN | Non-Toxin | 0.07 | 0.69 | -0.74 | 0 | 5.88 | 1051.3 | outside |
| 1223 | 1241 | GLIAIVMVTIMLCCMTSC | 0.4583 | 18 | ABCPRED | 1 | NON-ALLERGEN | Non-Toxin | 0.28 | 2.33 | -1.11 | 0 | 5.79 | 1902.77 | transmembrane-region_inside |
| 614 | 624 | DVNCTEVPVA | 1.237 | 10 | ABCPRED | 1 | NON-ALLERGEN | Non-Toxin | -0.03 | 0.41 | -0.02 | -2 | 3.67 | 1046.29 | outside |
| 490 | 502 | FPLQSYGFQPTN | 0.9942 | 12 | ABCPRED | 1 | ALLERGEN | Non-Toxin | -0.06 | -0.62 | -0.72 | 0 | 5.88 | 1398.71 | outside |

| | | | | | | | | | | | | | | | |
|---|---|---|---|---|---|---|---|---|---|---|---|---|---|---|---|
| 528 | 538 | KKSTNLVKNK | 0.7187 | 10 | ABCPRED | 1 | NON-ALLERGEN | Non-Toxin | -0.51 | -1.61 | 0.9 | 4 | 10.49 | 1159.54 | outside |
| 93 | 103 | ASTEKSNIIR | -0.463 | 10 | ABCPRED | 1 | NON-ALLERGEN | Non-Toxin | -0.31 | -0.69 | 0.53 | 1 | 9.1 | 1118.39 | outside |
| 58 | 70 | FFSNVTWFHAIH | 0.3872 | 12 | ABCPRED | 1 | NON-ALLERGEN | Non-Toxin | 0.15 | 0.55 | -1.3 | 1 | 7.26 | 1505.87 | outside |
| 490 | 506 | FPLQSYGFQPTNGVGY | 0.5285 | 16 | ABCPRED | 1 | ALLERGEN | Non-Toxin | 0.01 | -0.34 | -0.78 | 0 | 5.87 | 1775.19 | outside |
| 1119 | 1137 | NTFVSGNCDVVIGIVNNT | 0.3298 | 18 | ABCPRED | 1 | NON-ALLERGEN | Non-Toxin | 0.04 | 0.59 | -0.54 | -1 | 3.8 | 1866.35 | outside |
| 1096 | 1108 | VSNGTHWFVTQR | 0.2743 | 12 | AAP | 1 | NON-ALLERGEN | Non-Toxin | -0.16 | -0.58 | -0.54 | 1.5 | 10.11 | 1431.75 | outside |
| 1159 | 1173 | HTSPDVDLGDISGI | 1.0779 | 14 | AAP | 1 | NON-ALLERGEN | Non-Toxin | -0.03 | -0.1 | 0.13 | -2.5 | 3.93 | 1425.72 | outside |
| 112 | 130 | SKTQSLLIVNNATNVVIK | 0.5618 | 18 | BCPREDS | 1 | NON-ALLERGEN | Non-Toxin | -0.07 | 0.34 | -0.31 | 2 | 10.02 | 1942.57 | outside |
| 687 | 699 | VASQSIIAYTMS | 0.3055 | 12 | ABCPRED | 1 | NON-ALLERGEN | Non-Toxin | 0.09 | 0.9 | -0.75 | 0 | 5.88 | 1270.63 | outside |
| 472 | 482 | IYQAGSTPCN | 0.0551 | 10 | ABCPRED | 1 | ALLERGEN | Non-Toxin | -0.06 | -0.3 | -0.53 | 0 | 5.84 | 1053.29 | outside |
| 61 | 71 | NVTWFHAIHV | 0.9051 | 10 | ABCPRED | 1 | ALLERGEN | Non-Toxin | 0.14 | 0.6 | -1.24 | 1 | 7.26 | 1223.55 | outside |
| 81 | 88 | NPVLPFN | 0.5863 | 7 | IEDB,literature | 2 | ALLERGEN | Non-Toxin | 0.04 | 0.09 | -0.77 | 0 | 5.88 | 800.02 | outside |
| 363 | 375 | ADYSVLYNSASF | 0.3625 | 12 | ABCPRED | 1 | NON-ALLERGEN | Non-Toxin | 0.01 | 0.2 | -0.61 | -1 | 3.8 | 1336.57 | outside |
| 329 | 341 | FPNITNLCPFGE | 1.0991 | 12 | ABCPRED | 1 | ALLERGEN | Non-Toxin | 0.04 | 0.13 | -0.55 | -1 | 4 | 1351.71 | outside |
| 711 | 723 | SIAIPTNFTISV | 0.7561 | 12 | ABCPRED | 1 | NON-ALLERGEN | Non-Toxin | 0.17 | 1.18 | -0.83 | 0 | 5.88 | 1262.64 | outside |
| 682 | 692 | RRARSVASQS | 0.5522 | 10 | ABCPRED | 1 | NON-ALLERGEN | Non-Toxin | -0.57 | -1.16 | 0.76 | 3 | 12.31 | 1117.35 | outside |
| 63 | 75 | TWFHAIHVSGTN | 0.4142 | 12 | ABCPRED | 1 | NON-ALLERGEN | Non-Toxin | 0.05 | -0.01 | -0.92 | 1 | 7.26 | 1369.68 | outside |
| 592 | 608 | FGGVSVITPGTNTSNQ | 0.5777 | 16 | ABCPRED | 1 | ALLERGEN | Non-Toxin | -0.01 | -0.08 | -0.46 | 0 | 5.88 | 1578.94 | outside |
| 886 | 898 | WTFGAGAALQIP | 0.2563 | 12 | ABCPRED | 1 | ALLERGEN | Non-Toxin | 0.2 | 0.75 | -0.93 | 0 | 5.88 | 1231.59 | outside |
| 253 | 265 | DSSSGWTAGAAA | 0.3658 | 12 | ABCPRED | 1 | NON-ALLERGEN | Non-Toxin | 0 | -0.09 | -0.16 | -1 | 3.8 | 1080.22 | outside |
| 700 | 716 | GAENSVAYSNNSIAIP | 0.4192 | 16 | ABCPRED | 1 | NON-ALLERGEN | Non-Toxin | -0.03 | -0.07 | -0.28 | -1 | 4 | 1606.93 | outside |
| 361 | 379 | CVADYSVLYNSASFSTFK | 0.0657 | 18 | ABCPRED | 1 | NON-ALLERGEN | Non-Toxin | -0.02 | 0.36 | -0.52 | 0 | 6.15 | 2002.46 | outside |
| 723 | 733 | TTEILPVSMT | 1.484 | 10 | ABCPRED | 1 | NON-ALLERGEN | Non-Toxin | 0.06 | 0.64 | -0.43 | -1 | 4 | 1091.43 | outside |
| 523 | 533 | TVCGPKKSTN | -0.0257 | 10 | ABCPRED,IEDB | 2 | ALLERGEN | Non-Toxin | -0.28 | -0.88 | 0.32 | 2 | 9.36 | 1034.33 | outside |
| 512 | 521 | VLSFELLHA | 1.0776 | 9 | literature | 1 | NON-ALLERGEN | Non-Toxin | 0.19 | 1.41 | -0.79 | -0.5 | 5.25 | 1028.35 | outside |
| 255 | 265 | SSGWTAGAAA | 0.2219 | 10 | ABCPRED | 1 | ALLERGEN | Non-Toxin | 0.1 | 0.32 | -0.52 | 0 | 5.88 | 878.03 | outside |
| 159 | 173 | VYSSANNCTFEYVS | 0.1398 | 14 | FBCPred | 1 | NON-ALLERGEN | Non-Toxin | -0.06 | -0.05 | -0.55 | -1 | 4 | 1583.87 | outside |
| 333 | 340 | TNLCPFG | 1.1812 | 7 | IEDB,literature | 2 | NON-ALLERGEN | Non-Toxin | 0.06 | 0.41 | -0.79 | 0 | 5.85 | 750.96 | outside |
| 873 | 889 | YTSALLAGTITSGWTF | 0.4178 | 16 | ABCPRED | 1 | NON-ALLERGEN | Non-Toxin | 0.15 | 0.69 | -0.98 | 0 | 5.88 | 1689.13 | outside |
| 161 | 171 | SSANNCTFEY | -0.0845 | 10 | ABCPRED | 1 | NON-ALLERGEN | Non-Toxin | -0.17 | -0.7 | -0.27 | -1 | 4 | 1135.29 | outside |
| 698 | 712 | SLGAENSVAYSNNS | 0.7089 | 14 | FBCPred | 1 | NON-ALLERGEN | Non-Toxin | -0.13 | -0.52 | -0.13 | -1 | 4 | 1412.62 | outside |
| 162 | 174 | SANNCTFEYVSQ | 0.1822 | 12 | ABCPRED | 1 | NON-ALLERGEN | Non-Toxin | -0.15 | -0.53 | -0.33 | -1 | 4 | 1362.59 | outside |
| 60 | 70 | SNVTWFHAIH | 0.569 | 10 | ABCPRED | 1 | ALLERGEN | Non-Toxin | 0.06 | 0.1 | -1.06 | 1 | 7.26 | 1211.49 | outside |
| 580 | 590 | QTLEILDITP | 2.027 | 10 | ABCPRED | 1 | NON-ALLERGEN | Non-Toxin | 0.01 | 0.31 | -0.18 | -2 | 3.67 | 1142.47 | outside |
| 688 | 700 | ASQSIIAYTMSL | 0.3219 | 12 | ABCPRED | 1 | NON-ALLERGEN | Non-Toxin | 0.09 | 0.87 | -0.78 | 0 | 5.88 | 1284.66 | outside |
| 950 | 968 | DVVNQNAQALNTLVKQLS | 0.1196 | 18 | AAP | 1 | NON-ALLERGEN | Non-Toxin | -0.14 | -0.13 | -0.21 | 0 | 6.19 | 1955.48 | outside |
| 393 | 403 | TNVYADSFVI | -0.4541 | 10 | ABCPRED | 1 | ALLERGEN | Non-Toxin | 0.09 | 0.77 | -0.7 | -1 | 3.8 | 1128.38 | outside |
| 1082 | 1089 | CHDGKAH | -0.2298 | 7 | IEDB | 1 | ALLERGEN | Non-Toxin | -0.31 | -1.41 | 0.5 | 1 | 7.22 | 766.92 | outside |
| 706 | 724 | AYSNNSIAIPTNFTISVT | 0.842 | 18 | ABCPRED | 1 | ALLERGEN | Non-Toxin | 0.03 | 0.34 | -0.69 | 0 | 5.88 | 1913.37 | outside |
| 163 | 173 | ANNCTFEYVS | 0.2319 | 10 | ABCPRED | 1 | ALLERGEN | Non-Toxin | -0.09 | -0.2 | -0.45 | -1 | 4 | 1147.35 | outside |
| 604 | 616 | TSNQVAVLYQDV | 0.2882 | 12 | ABCPRED | 1 | NON-ALLERGEN | Non-Toxin | -0.06 | 0.12 | -0.47 | -1 | 3.8 | 1336.64 | outside |
| 486 | 504 | FNCYFPLQSYGFQPTNGV | 0.8002 | 18 | ABCPRED | 1 | ALLERGEN | Non-Toxin | 0 | -0.18 | -0.87 | 0 | 5.84 | 2082.58 | outside |
| 624 | 634 | IHADQLTPTW | 0.2628 | 10 | ABCPRED | 1 | ALLERGEN | Non-Toxin | -0.04 | -0.4 | -0.56 | -0.5 | 5.09 | 1181.46 | outside |
| 710 | 722 | NSIAIPTNFTIS | 0.7228 | 12 | ABCPRED | 1 | NON-ALLERGEN | Non-Toxin | 0.07 | 0.54 | -0.68 | 0 | 5.88 | 1277.61 | outside |
| 1114 | 1126 | IITTDNTFVSGN | 0.3214 | 12 | FBCPred | 1 | NON-ALLERGEN | Toxin | 0 | 0.18 | -0.43 | -1 | 3.8 | 1281.56 | outside |
| 601 | 619 | GTNTSNQVAVLYQDVNCT | 0.8108 | 18 | ABCPRED | 1 | ALLERGEN | Non-Toxin | -0.12 | -0.27 | -0.39 | -1 | 3.8 | 1927.34 | outside |
| 589 | 605 | PCSFGGVSVITPGTNT | 0.76 | 16 | ABCPRED | 1 | NON-ALLERGEN | Non-Toxin | 0.07 | 0.41 | -0.54 | 0 | 5.85 | 1536.95 | outside |
| 481 | 491 | NGVEGFNCYF | 0.6268 | 10 | ABCPRED | 1 | ALLERGEN | Non-Toxin | 0.02 | -0.03 | -0.64 | -1 | 4 | 1149.38 | outside |
| 61 | 77 | NVTWFHAIHVSGTNGT | 0.8461 | 16 | ABCPRED,literature | 2 | NON-ALLERGEN | Non-Toxin | 0.03 | -0.03 | -0.79 | 1 | 7.26 | 1741.14 | outside |
| 1116 | 1126 | TTDNTFVSGN | 0.3035 | 10 | IEDB | 1 | ALLERGEN | Non-Toxin | -0.15 | -0.68 | -0.15 | -1 | 3.8 | 1055.2 | outside |
| 1114 | 1128 | IITTDNTFVSGNCD | 0.1793 | 14 | FBCPred | 1 | NON-ALLERGEN | Non-Toxin | -0.05 | 0.09 | -0.22 | -2 | 3.57 | 1499.81 | outside |
| 837 | 844 | YGDCLGD | -0.5555 | 7 | IEDB,literature | 2 | NON-ALLERGEN | Non-Toxin | -0.08 | -0.4 | 0.13 | -2 | 3.57 | 741.86 | outside |
| 1168 | 1184 | DISGINASVVNIQKEI | 0.6254 | 16 | ABCPRED | 1 | NON-ALLERGEN | Non-Toxin | -0.03 | 0.3 | -0.03 | -1 | 4.38 | 1700.17 | outside |
| 1054 | 1072 | QSAPHGVVFLHVTYVPAQ | 0.7664 | 18 | ABCPRED | 1 | NON-ALLERGEN | Non-Toxin | 0.07 | 0.4 | -0.79 | 1 | 7.25 | 1950.5 | outside |
| 1220 | 1236 | FIAGLIAIVMVTIMLC | 0.3648 | 16 | ABCPRED | 1 | NON-ALLERGEN | Non-Toxin | 0.42 | 2.85 | -1.33 | 0 | 5.85 | 1708.51 | transmembrane-region |
| 389 | 401 | DLCFTNVYADSF | 0.6691 | 12 | ABCPRED,FBCPred | 2 | NON-ALLERGEN | Non-Toxin | 0.01 | 0.38 | -0.5 | -2 | 3.57 | 1394.67 | outside |
| 883 | 897 | TSGWTFGAGAALQI | 0.2225 | 14 | ABCPRED,AAP | 2 | NON-ALLERGEN | Non-Toxin | 0.15 | 0.62 | -0.81 | 0 | 5.88 | 1379.74 | outside |
| 387 | 397 | LNDLCFTNVY | 1.6134 | 10 | ABCPRED | 1 | NON-ALLERGEN | Non-Toxin | 0.01 | 0.46 | -0.79 | -1 | 3.8 | 1201.5 | outside |
| 480 | 490 | CNGVEGFNCY | 0.4903 | 10 | ABCPRED | 1 | ALLERGEN | Non-Toxin | -0.03 | -0.06 | -0.49 | -1 | 4 | 1105.34 | outside |
| 712 | 724 | IAIPTNFTISVT | 1.0108 | 12 | ABCPRED | 1 | NON-ALLERGEN | Non-Toxin | 0.17 | 1.19 | -0.88 | 0 | 5.88 | 1276.67 | outside |
| 710 | 726 | NSIAIPTNFTISVTTE | 0.798 | 16 | ABCPRED | 1 | NON-ALLERGEN | Non-Toxin | 0.02 | 0.36 | -0.47 | -1 | 4 | 1708.13 | outside |
| 609 | 619 | AVLYQDVNCT | 0.8761 | 10 | ABCPRED | 1 | NON-ALLERGEN | Non-Toxin | -0.03 | 0.4 | -0.56 | -1 | 3.8 | 1125.4 | outside |
| 362 | 372 | VADYSVLYNS | 0.0734 | 10 | ABCPRED | 1 | ALLERGEN | Non-Toxin | 0 | 0.28 | -0.61 | -1 | 3.8 | 1130.35 | outside |
| 492 | 510 | LQSYGFQPTNGVGYQPYR | 0.594 | 18 | ABCPRED | 1 | NON-ALLERGEN | Non-Toxin | -0.16 | -0.97 | -0.5 | 1 | 8.83 | 2075.54 | outside |
| 607 | 616 | QVAVLYQDV | 0.265 | 9 | IEDB,literature | 2 | ALLERGEN | Non-Toxin | 0.04 | 0.71 | -0.63 | -1 | 3.8 | 1034.31 | outside |
| 530 | 540 | STNLVKNKCV | 0.205 | 10 | ABCPRED | 1 | ALLERGEN | Non-Toxin | -0.23 | -0.16 | 0.05 | 2 | 9.36 | 1105.46 | outside |
| 745 | 761 | DSTECSNLLLQYGSFC | 0.8382 | 16 | ABCPRED | 1 | NON-ALLERGEN | Non-Toxin | -0.07 | 0.02 | -0.33 | -2 | 3.67 | 1780.18 | outside |
| 606 | 618 | NQVAVLYQDVNC | 0.7078 | 12 | ABCPRED | 1 | ALLERGEN | Non-Toxin | -0.08 | 0.16 | -0.53 | -1 | 3.8 | 1365.7 | outside |
| 588 | 608 | TPCSFGGVSVITPGTNTSNQ | 0.8072 | 20 | ABCPRED | 1 | ALLERGEN | Non-Toxin | -0.03 | -0.09 | -0.42 | 0 | 5.85 | 1967.43 | outside |
| 493 | 509 | QSYGFQPTNGVGYQPY | 0.702 | 16 | ABCPRED | 1 | NON-ALLERGEN | Non-Toxin | -0.1 | -1.05 | -0.64 | 0 | 5.87 | 1806.16 | outside |
| 468 | 480 | ISTEIYQAGSTP | 0.0986 | 12 | ABCPRED | 1 | NON-ALLERGEN | Non-Toxin | -0.03 | -0.21 | -0.28 | -1 | 4 | 1266.54 | outside |
| 588 | 606 | TPCSFGGVSVITPGTNTS | 0.8124 | 18 | ABCPRED | 1 | NON-ALLERGEN | Non-Toxin | 0.04 | 0.28 | -0.49 | 0 | 5.85 | 1725.16 | outside |
| 61 | 73 | NVTWFHAIHVSG | 0.5465 | 12 | ABCPRED | 1 | NON-ALLERGEN | Non-Toxin | 0.11 | 0.4 | -1.01 | 1 | 7.26 | 1367.71 | outside |
| 1162 | 1182 | PDVDLGDISGINASVVNIQK | 0.9428 | 20 | FBCPred | 1 | NON-ALLERGEN | Non-Toxin | -0.05 | 0.14 | 0.05 | -2 | 3.94 | 2054.59 | outside |
| 603 | 619 | NTSNQVAVLYQDVNCT | 0.7566 | 16 | ABCPRED | 1 | NON-ALLERGEN | Non-Toxin | -0.14 | -0.24 | -0.41 | -1 | 3.8 | 1769.15 | outside |
| 491 | 509 | PLQSYGFQPTNGVGYQPY | 0.4777 | 18 | ABCPRED | 1 | ALLERGEN | Non-Toxin | -0.06 | -0.81 | -0.67 | 0 | 5.87 | 2016.47 | outside |
| 869 | 881 | MIAQYTSALLAG | 0.2649 | 12 | ABCPRED | 1 | NON-ALLERGEN | Non-Toxin | 0.15 | 1.06 | -0.87 | 0 | 5.88 | 1238.64 | outside |

| | | | | | | | | | | | | | | | |
|---|---|---|---|---|---|---|---|---|---|---|---|---|---|---|---|
| 1238 | 1250 | TSCCSCLKGCCS | -0.3486 | 12 | FBCPred | 1 | ALLERGEN | Non-Toxin | -0.1 | 0.74 | -0.27 | 1 | 8.1 | 1194.58 | inside |
| 707 | 717 | YSNNSIAIPT | 0.2831 | 10 | ABCPRED | 1 | ALLERGEN | Non-Toxin | -0.03 | -0.14 | -0.58 | 0 | 5.88 | 1079.31 | outside |
| 605 | 617 | SNQVAVLYQDVN | 0.3957 | 12 | ABCPRED | 1 | NON-ALLERGEN | Non-Toxin | -0.1 | -0.12 | -0.42 | -1 | 3.8 | 1349.64 | outside |
| 699 | 713 | LGAENSVAYSNNSI | 0.608 | 14 | IEDB | 1 | NON-ALLERGEN | Non-Toxin | -0.06 | -0.14 | -0.28 | -1 | 4 | 1438.71 | outside |
| 961 | 970 | TLVKQLSSN | -0.5178 | 9 | IEDB | 1 | NON-ALLERGEN | Non-Toxin | -0.17 | -0.16 | -0.17 | 1 | 9.11 | 989.27 | outside |
| 698 | 716 | SLGAENSVAYSNNSIAIP | 0.6736 | 18 | ABCPRED | 1 | NON-ALLERGEN | Non-Toxin | -0.01 | 0.11 | -0.33 | -1 | 4 | 1807.2 | outside |
| 699 | 711 | LGAENSVAYSNN | 0.6039 | 12 | ABCPRED | 1 | NON-ALLERGEN | Non-Toxin | -0.11 | -0.47 | -0.2 | -1 | 4 | 1238.44 | outside |
| 746 | 764 | STECSNLLLQYGSFCTQL | 0.7694 | 18 | AAP | 1 | NON-ALLERGEN | Non-Toxin | -0.04 | 0.19 | -0.57 | -1 | 4 | 2007.53 | outside |
| 529 | 539 | KSTNLVKNKC | 0.7023 | 10 | ABCPRED | 1 | NON-ALLERGEN | Non-Toxin | -0.39 | -0.97 | 0.5 | 3 | 9.81 | 1134.5 | outside |
| 262 | 274 | AAAYYVGYLQPR | 1.0262 | 12 | ABCPRED | 1 | NON-ALLERGEN | Non-Toxin | -0.04 | -0.04 | -0.71 | 1 | 8.83 | 1371.72 | outside |
| 485 | 501 | GFNCYFPLQSYGFQPT | 0.981 | 16 | ABCPRED | 1 | NON-ALLERGEN | Non-Toxin | 0.01 | -0.24 | -0.9 | 0 | 5.84 | 1869.31 | outside |
| 495 | 505 | YGFQPTNGVG | 0.7293 | 10 | ABCPRED | 1 | ALLERGEN | Non-Toxin | 0.01 | -0.48 | -0.63 | 0 | 5.88 | 1039.26 | outside |
| 596 | 614 | SVITPGTNTSNQVAVLYQ | 0.389 | 18 | BCPREDS | 1 | ALLERGEN | Non-Toxin | -0.03 | 0.09 | -0.59 | 0 | 5.88 | 1892.37 | outside |
| 360 | 378 | NCVADYSVLYNSASFSTF | 0.0071 | 18 | ABCPRED | 1 | NON-ALLERGEN | Non-Toxin | 0.01 | 0.38 | -0.68 | -1 | 3.8 | 1988.39 | outside |
| 388 | 402 | NDLCFTNVYADSFV | 0.6072 | 14 | FBCPred | 1 | NON-ALLERGEN | Non-Toxin | 0 | 0.38 | -0.52 | -2 | 3.57 | 1607.94 | outside |
| 409 | 419 | QIAPGQTGKI | 1.4416 | 10 | ABCPRED | 1 | ALLERGEN | Non-Toxin | -0.07 | -0.32 | -0.11 | 1 | 9.11 | 1012.33 | outside |
| 168 | 175 | FEYVSQP | 0.9073 | 7 | IEDB,literature | 2 | ALLERGEN | Non-Toxin | -0.07 | -0.53 | -0.4 | -1 | 4 | 869.03 | outside |
| 588 | 600 | TPCSFGGVSVIT | 1.1714 | 12 | ABCPRED | 1 | NON-ALLERGEN | Non-Toxin | 0.15 | 1.07 | -0.71 | 0 | 5.85 | 1167.51 | outside |
| 1159 | 1179 | HTSPDVDLGDISGINASVVN | 0.9177 | 20 | AAP | 1 | ALLERGEN | Non-Toxin | -0.03 | 0.05 | -0.05 | -2.5 | 3.93 | 2010.44 | outside |
| 373 | 383 | SFSTFKCYGV | 0.5939 | 10 | ABCPRED | 1 | NON-ALLERGEN | Non-Toxin | 0.02 | 0.44 | -0.66 | 1 | 8.54 | 1138.43 | outside |
| 888 | 900 | FGAGAALQIPFA | 0.6218 | 12 | ABCPRED | 1 | ALLERGEN | Non-Toxin | 0.25 | 1.27 | -0.87 | 0 | 5.88 | 1162.52 | outside |
| 113 | 131 | KTQSLLIVNNATNVVIKV | 0.5868 | 18 | ABCPRED | 1 | NON-ALLERGEN | Non-Toxin | -0.03 | 0.62 | -0.41 | 2 | 10.02 | 1954.63 | outside |
| 652 | 662 | GAEHVNNSYE | 0.8739 | 10 | ellipro,literature | 2 | ALLERGEN | Non-Toxin | -0.22 | -1.37 | 0.19 | -1.5 | 4.51 | 1119.25 | outside |
| 111 | 129 | DSKTQSLLIVNNATNVVI | 0.5418 | 18 | ABCPRED | 1 | NON-ALLERGEN | Non-Toxin | -0.05 | 0.37 | -0.31 | 0 | 6.19 | 1929.48 | outside |
| 587 | 603 | ITPCSFGGVSVITPGT | 0.854 | 16 | ABCPRED | 1 | NON-ALLERGEN | Non-Toxin | 0.15 | 0.91 | -0.67 | 0 | 5.85 | 1536.01 | outside |
| 1116 | 1130 | TTDNTFVSGNCDVV | 0.1255 | 14 | BCPREDS | 1 | NON-ALLERGEN | Non-Toxin | -0.08 | 0.04 | -0.18 | -2 | 3.57 | 1471.75 | outside |
| 621 | 633 | PVAIHADQLTPT | 0.5103 | 12 | ABCPRED | 1 | NON-ALLERGEN | Non-Toxin | 0 | 0.11 | -0.35 | -0.5 | 5.09 | 1262.6 | outside |
| 490 | 500 | FPLQSYGFQP | 0.6824 | 10 | ABCPRED | 1 | NON-ALLERGEN | Non-Toxin | 0.02 | -0.33 | -0.84 | 0 | 5.88 | 1183.47 | outside |
| 359 | 375 | SNCVADYSVLYNSASF | 0.0304 | 16 | ABCPRED | 1 | ALLERGEN | Non-Toxin | -0.01 | 0.3 | -0.58 | -1 | 3.8 | 1740.08 | outside |
| 951 | 969 | VVNQNAQALNTLVKQLSS | -0.0107 | 18 | ABCPRED | 1 | NON-ALLERGEN | Non-Toxin | -0.12 | 0.02 | -0.36 | 1 | 9.11 | 1927.47 | outside |
| 530 | 542 | STNLVKNKCVNF | 0.8907 | 12 | ABCPRED | 1 | ALLERGEN | Non-Toxin | -0.19 | -0.19 | -0.15 | 2 | 9.36 | 1366.77 | outside |
| 520 | 534 | APATVCGPKKSTNL | 0.2327 | 14 | ABCPRED,AAP | 2 | NON-ALLERGEN | Non-Toxin | -0.13 | -0.21 | 0.03 | 2 | 9.36 | 1386.82 | outside |
| 1220 | 1238 | FIAGLIAIVMVTIMLCCM | 0.4744 | 18 | ABCPRED | 1 | NON-ALLERGEN | Non-Toxin | 0.39 | 2.78 | -1.31 | 0 | 5.82 | 1942.87 | transmembrane-region |
| 623 | 633 | AIHADQLTPT | 0.4202 | 10 | ABCPRED | 1 | NON-ALLERGEN | Non-Toxin | -0.05 | -0.13 | -0.27 | -0.5 | 5.09 | 1066.32 | outside |
| 519 | 531 | HAPATVCGPKKS | -0.1664 | 12 | BCPREDS | 1 | NON-ALLERGEN | Non-Toxin | -0.16 | -0.48 | 0.16 | 2.5 | 9.36 | 1195.56 | outside |
| 663 | 679 | DIPIGAGICASYQTQT | 1.0235 | 16 | ABCPRED | 1 | NON-ALLERGEN | Non-Toxin | 0.02 | 0.2 | -0.43 | -1 | 3.8 | 1638.06 | outside |
| 589 | 607 | PCSFGGVSVITPGTNTSN | 0.7915 | 18 | ABCPRED | 1 | NON-ALLERGEN | Non-Toxin | 0.01 | 0.13 | -0.46 | 0 | 5.85 | 1738.16 | outside |
| 590 | 602 | CSFGGVSVITPG | 0.8136 | 12 | ABCPRED | 1 | NON-ALLERGEN | Non-Toxin | 0.18 | 1.09 | -0.67 | 0 | 5.85 | 1123.46 | outside |
| 1158 | 1172 | NHTSPDVDLGDISG | 1.0939 | 14 | FBCPred | 1 | NON-ALLERGEN | Non-Toxin | -0.13 | -0.67 | 0.27 | -2.5 | 3.93 | 1426.66 | outside |
| 490 | 508 | FPLQSYGFQPTNGVGYQP | 0.5761 | 18 | ABCPRED | 1 | NON-ALLERGEN | Non-Toxin | -0.03 | -0.58 | -0.68 | 0 | 5.87 | 2000.47 | outside |
| 286 | 298 | TDAVDCALDPLS | 0.6508 | 12 | ABCPRED | 1 | NON-ALLERGEN | Non-Toxin | -0.04 | 0.36 | 0.15 | -3 | 3.43 | 1219.48 | outside |
| 1051 | 1069 | SFPQSAPHGVVFLHVTYV | 0.5904 | 18 | ABCPRED | 1 | NON-ALLERGEN | Non-Toxin | 0.11 | 0.61 | -0.9 | 1 | 7.25 | 1985.54 | outside |
| 617 | 629 | CTEVPVAIHADQ | 0.0112 | 12 | ABCPRED | 1 | NON-ALLERGEN | Non-Toxin | -0.03 | 0.25 | -0.13 | -1.5 | 4.36 | 1282.6 | outside |
| 961 | 969 | TLVKQLSS | -0.4822 | 8 | BCEPRED | 1 | ALLERGEN | Non-Toxin | -0.11 | 0.26 | -0.21 | 1 | 9.11 | 875.15 | outside |
| 592 | 610 | FGGVSVITPGTNTSNQVA | 0.5541 | 18 | ABCPRED | 1 | NON-ALLERGEN | Non-Toxin | 0.03 | 0.26 | -0.52 | 0 | 5.88 | 1749.18 | outside |
| 254 | 270 | SSSGWTAGAAAYYVGY | 0.4248 | 16 | ABCPRED | 1 | NON-ALLERGEN | Non-Toxin | 0.09 | 0.14 | -0.83 | 0 | 5.87 | 1610.91 | outside |
| 1118 | 1136 | DNTFVSGNCDVVIGIVNN | 0.4207 | 18 | ABCPRED | 1 | NON-ALLERGEN | Non-Toxin | 0.01 | 0.43 | -0.36 | -2 | 3.57 | 1880.33 | outside |
| 529 | 541 | KSTNLVKNKCVN | 0.6943 | 12 | ABCPRED | 1 | NON-ALLERGEN | Non-Toxin | -0.33 | -0.75 | 0.31 | 3 | 9.81 | 1347.77 | outside |
| 1118 | 1138 | DNTFVSGNCDVVIGIVNNTV | 0.404 | 20 | ABCPRED | 1 | ALLERGEN | Non-Toxin | 0.03 | 0.57 | -0.42 | -2 | 3.57 | 2080.6 | outside |
| 489 | 509 | YFPLQSYGFQPTNGVGYQPY | 0.6186 | 20 | ABCPRED | 1 | ALLERGEN | Non-Toxin | -0.03 | -0.66 | -0.84 | 0 | 5.87 | 2326.85 | outside |
| 870 | 888 | IAQYTSALLAGTITSGWT | 0.3136 | 18 | ABCPRED | 1 | NON-ALLERGEN | Non-Toxin | 0.11 | 0.62 | -0.84 | 0 | 5.88 | 1854.36 | outside |
| 586 | 604 | DITPCSFGGVSVITPGTN | 1.0022 | 18 | ABCPRED | 1 | NON-ALLERGEN | Non-Toxin | 0.06 | 0.42 | -0.42 | -1 | 3.8 | 1765.23 | outside |
| 592 | 604 | FGGVSVITPGTN | 0.4578 | 12 | ABCPRED | 1 | NON-ALLERGEN | Non-Toxin | 0.13 | 0.6 | -0.63 | 0 | 5.88 | 1148.46 | outside |
| 430 | 442 | TGCVIAWNSNNL | 0.406 | 12 | FBCPred | 1 | NON-ALLERGEN | Non-Toxin | 0.02 | 0.29 | -0.79 | 0 | 5.85 | 1291.62 | outside |
| 390 | 402 | LCFTNVYADSFV | 0.3921 | 12 | ABCPRED | 1 | ALLERGEN | Non-Toxin | 0.11 | 1.03 | -0.88 | -1 | 3.8 | 1378.72 | outside |
| 873 | 893 | YTSALLAGTITSGWTFGAGA | 0.3854 | 20 | AAP | 1 | NON-ALLERGEN | Non-Toxin | 0.16 | 0.7 | -0.83 | 0 | 5.88 | 1945.45 | outside |
| 1238 | 1256 | TSCCSCLKGCCSCGSCCK | -0.0153 | 18 | ABCPRED | 1 | ALLERGEN | Toxin | -0.12 | 0.63 | -0.17 | 2 | 8.33 | 1776.38 | inside |
| 1121 | 1137 | FVSGNCDVVIGIVNNT | 0.5737 | 16 | ABCPRED | 1 | ALLERGEN | Non-Toxin | 0.09 | 0.92 | -0.6 | -1 | 3.8 | 1651.11 | outside |
| 112 | 122 | SKTQSLLIVN | 0.8487 | 10 | ABCPRED | 1 | ALLERGEN | Non-Toxin | -0.08 | 0.31 | -0.33 | 1 | 9.11 | 1102.45 | outside |
| 2 | 14 | FVFLVLLPLVSS | 0.6858 | 12 | ABCPRED | 1 | NON-ALLERGEN | Non-Toxin | 0.36 | 2.52 | -1.34 | 0 | 5.88 | 1333.86 | outside |
| 647 | 663 | AGCLIGAEHVNNSYEC | 0.914 | 16 | ABCPRED | 1 | NON-ALLERGEN | Non-Toxin | -0.03 | 0.06 | -0.26 | -1.5 | 4.51 | 1680.07 | outside |
| 53 | 63 | DLFLPFFSNV | -0.1949 | 10 | ABCPRED | 1 | ALLERGEN | Non-Toxin | 0.17 | 1.08 | -0.91 | -1 | 3.8 | 1198.52 | outside |
| 487 | 497 | NCYFPLQSYG | 0.5904 | 10 | ABCPRED | 1 | ALLERGEN | Non-Toxin | -0.03 | -0.33 | -0.92 | 0 | 5.84 | 1191.46 | outside |
| 64 | 74 | WFHAIHVSGT | 0.5138 | 10 | ABCPRED | 1 | NON-ALLERGEN | Non-Toxin | 0.14 | 0.41 | -1.08 | 1 | 7.26 | 1154.44 | outside |
| 161 | 175 | SSANNCTFEYVSQP | 0.0749 | 14 | AAP,BCPREDS | 2 | ALLERGEN | Non-Toxin | -0.15 | -0.62 | -0.26 | -1 | 4 | 1546.81 | outside |
| 619 | 629 | EVPVAIHADQ | -0.0056 | 10 | ABCPRED | 1 | NON-ALLERGEN | Non-Toxin | -0.02 | 0.12 | -0.01 | -1.5 | 4.36 | 1078.33 | outside |
| 617 | 628 | CTEVPVAIHAD | 0.0499 | 11 | IEDB,literature | 2 | ALLERGEN | Non-Toxin | 0.03 | 0.59 | -0.15 | -1.5 | 4.36 | 1154.45 | outside |
| 621 | 631 | PVAIHADQLT | 0.2095 | 10 | ABCPRED | 1 | NON-ALLERGEN | Non-Toxin | 0.02 | 0.36 | -0.38 | -0.5 | 5.09 | 1064.35 | outside |
| 469 | 479 | STEIYQAGST | 0.0944 | 10 | ABCPRED | 1 | NON-ALLERGEN | Non-Toxin | -0.1 | -0.54 | -0.16 | -1 | 4 | 1056.23 | outside |
| 880 | 892 | GTITSGWTFGAG | 0.2164 | 12 | FBCPred | 1 | ALLERGEN | Non-Toxin | 0.15 | 0.31 | -0.76 | 0 | 5.88 | 1154.42 | outside |
| 876 | 888 | ALLAGTITSGWT | 0.4151 | 12 | ABCPRED | 1 | ALLERGEN | Non-Toxin | 0.18 | 0.92 | -0.89 | 0 | 5.88 | 1190.54 | outside |
| 721 | 733 | SVTTEILPVSMT | 1.2175 | 12 | ABCPRED | 1 | NON-ALLERGEN | Non-Toxin | 0.07 | 0.82 | -0.46 | -1 | 4 | 1277.67 | outside |
| 160 | 172 | YSSANNCTFEYV | 0.0246 | 12 | ABCPRED | 1 | ALLERGEN | Non-Toxin | -0.09 | -0.34 | -0.54 | -1 | 4 | 1397.63 | outside |
| 959 | 967 | LNTLVKQL | -0.7591 | 8 | IEDB,literature | 2 | ALLERGEN | Non-Toxin | -0.06 | 0.5 | -0.49 | 1 | 9.11 | 928.27 | outside |
| 907 | 921 | NGIGVTQNVLYENQ | 0.5244 | 14 | ABCPRED,FBCPred | 2 | NON-ALLERGEN | Non-Toxin | -0.1 | -0.51 | -0.38 | -1 | 4 | 1548.9 | outside |
| 495 | 507 | YGFQPTNGVGYQ | 0.7136 | 12 | IEDB,literature | 2 | NON-ALLERGEN | Non-Toxin | -0.05 | -0.8 | -0.7 | 0 | 5.87 | 1330.6 | outside |
| 112 | 124 | SKTQSLLIVNNA | 0.6479 | 12 | ABCPRED | 1 | NON-ALLERGEN | Non-Toxin | -0.1 | 0.12 | -0.3 | 1 | 9.11 | 1287.66 | outside |

| | | | | | | | | | | | | | | | |
|---|---|---|---|---|---|---|---|---|---|---|---|---|---|---|---|
| 874 | 892 | TSALLAGTITSGWTFGAG | 0.3317 | 18 | ABCPRED | 1 | NON-ALLERGEN | Non-Toxin | 0.16 | 0.74 | -0.77 | 0 | 5.88 | 1711.17 | outside |
| 1114 | 1129 | IITTDNTFVSGNCDV | 0.1092 | 15 | IEDB | 1 | NON-ALLERGEN | Non-Toxin | -0.01 | 0.36 | -0.31 | -2 | 3.57 | 1598.96 | outside |
| 264 | 274 | AYYVGYLQPR | 1.3309 | 10 | ABCPRED | 1 | ALLERGEN | Non-Toxin | -0.1 | -0.41 | -0.75 | 1 | 8.83 | 1229.54 | outside |
| 953 | 971 | NQNAQALNTLVKQLSSNF | -0.018 | 18 | ABCPRED | 1 | NON-ALLERGEN | Non-Toxin | -0.18 | -0.48 | -0.32 | 1 | 9.11 | 1990.48 | outside |
| 486 | 502 | FNCYFPLQSYGFQPTN | 1.0229 | 16 | ABCPRED | 1 | ALLERGEN | Non-Toxin | -0.04 | -0.44 | -0.89 | 0 | 5.84 | 1926.36 | outside |
| 391 | 403 | CFTNVYADSFVI | -0.0283 | 12 | ABCPRED | 1 | ALLERGEN | Non-Toxin | 0.13 | 1.08 | -0.88 | -1 | 3.8 | 1378.72 | outside |
| 53 | 61 | DLFLPFFS | -0.3099 | 8 | IEDB,literature | 2 | ALLERGEN | Non-Toxin | 0.23 | 1.26 | -0.97 | -1 | 3.8 | 985.25 | outside |
| 520 | 530 | APATVCGPKK | -0.1417 | 10 | ABCPRED | 1 | NON-ALLERGEN | Non-Toxin | -0.13 | -0.18 | 0.21 | 2 | 9.36 | 971.31 | outside |
| 700 | 714 | GAENSVAYSNNSIA | 0.4195 | 14 | BCPREDS | 1 | NON-ALLERGEN | Non-Toxin | -0.08 | -0.29 | -0.19 | -1 | 4 | 1396.62 | outside |
| 869 | 887 | MIAQYTSALLAGTITSGW | 0.3352 | 18 | ABCPRED | 1 | ALLERGEN | Non-Toxin | 0.14 | 0.76 | -0.89 | 0 | 5.88 | 1884.45 | outside |
| 55 | 65 | FLPFFSNVTW | 0.8835 | 10 | ABCPRED | 1 | NON-ALLERGEN | Non-Toxin | 0.21 | 0.89 | -1.41 | 0 | 5.88 | 1257.59 | outside |
| 1055 | 1073 | SAPHGVVFLHVTYVPAQE | 0.6533 | 18 | ABCPRED | 1 | NON-ALLERGEN | Non-Toxin | 0.07 | 0.4 | -0.64 | 0 | 6.01 | 1951.48 | outside |
| 1160 | 1178 | TSPDVDLGDISGINASVV | 0.9371 | 18 | ABCPRED | 1 | ALLERGEN | Non-Toxin | 0.02 | 0.43 | -0.04 | -3 | 3.43 | 1759.16 | outside |
| 590 | 610 | CSFGGVSVITPGTNTSNQVA | 0.6784 | 20 | ABCPRED | 1 | NON-ALLERGEN | Non-Toxin | 0.02 | 0.32 | -0.5 | 0 | 5.85 | 1939.42 | outside |
| 881 | 893 | TITSGWTFGAGA | 0.3448 | 12 | ABCPRED | 1 | NON-ALLERGEN | Non-Toxin | 0.16 | 0.49 | -0.8 | 0 | 5.88 | 1168.44 | outside |
| 538 | 556 | CVNFNFNGLTGTGVLTES | 1.1292 | 18 | ABCPRED | 1 | NON-ALLERGEN | Non-Toxin | 0.03 | 0.33 | -0.55 | -1 | 4 | 1873.34 | outside |
| 338 | 346 | FGEVFNAT | -0.0731 | 8 | COVIDep | 1 | ALLERGEN | Non-Toxin | 0.09 | 0.44 | -0.53 | -1 | 4 | 884.06 | outside |
| 1113 | 1131 | QIITTDNTFVSGNCDVVI | 0.1387 | 18 | ABCPRED | 1 | ALLERGEN | Non-Toxin | 0.02 | 0.59 | -0.43 | -2 | 3.57 | 1939.44 | outside |
| 362 | 376 | VADYSVLYNSASFS | 0.3971 | 14 | ABCPRED,FBCPred | 2 | NON-ALLERGEN | Non-Toxin | 0.03 | 0.41 | -0.61 | -1 | 3.8 | 1522.81 | outside |
| 671 | 683 | CASYQTQTNSPR | 0.1696 | 12 | ABCPRED | 1 | NON-ALLERGEN | Non-Toxin | -0.37 | -1.38 | -0.03 | 1 | 8.57 | 1355.6 | outside |
| 283 | 297 | GTITDAVDCALDPL | 0.5939 | 14 | AAP | 1 | NON-ALLERGEN | Non-Toxin | 0.03 | 0.61 | -0.05 | -3 | 3.43 | 1403.76 | outside |
| 519 | 533 | HAPATVCGPKKSTN | -0.0107 | 14 | ABCPRED,literature | 2 | NON-ALLERGEN | Non-Toxin | -0.2 | -0.71 | 0.12 | 2.5 | 9.36 | 1410.8 | outside |
| 709 | 725 | NNSIAIPTNFTISVTT | 0.7532 | 16 | ABCPRED | 1 | NON-ALLERGEN | Non-Toxin | 0.02 | 0.36 | -0.64 | 0 | 5.88 | 1693.12 | outside |
| 777 | 787 | NTQEVFAQVK | 0.5676 | 10 | ABCPRED | 1 | ALLERGEN | Non-Toxin | -0.2 | -0.56 | 0.02 | 0 | 6.35 | 1163.44 | outside |
| 1120 | 1136 | TFVSGNCDVVIGIVNN | 0.423 | 16 | ABCPRED | 1 | ALLERGEN | Non-Toxin | 0.09 | 0.93 | -0.6 | -1 | 3.8 | 1651.11 | outside |
| 496 | 508 | GFQPTNGVGYQP | 0.6299 | 12 | ABCPRED,IEDB | 2 | ALLERGEN | Non-Toxin | -0.06 | -0.83 | -0.51 | 0 | 5.88 | 1264.54 | outside |
| 361 | 373 | CVADYSVLYNSA | -0.1052 | 12 | ABCPRED | 1 | ALLERGEN | Non-Toxin | 0.03 | 0.59 | -0.63 | -1 | 3.8 | 1304.59 | outside |
| 47 | 57 | VLHSTQDLFL | 0.3211 | 10 | ABCPRED | 1 | NON-ALLERGEN | Non-Toxin | 0.05 | 0.67 | -0.68 | -0.5 | 5.09 | 1172.5 | outside |
| 358 | 376 | ISNCVADYSVLYNSASFS | -0.0372 | 18 | ABCPRED | 1 | NON-ALLERGEN | Non-Toxin | 0.01 | 0.47 | -0.6 | -1 | 3.8 | 1940.35 | outside |
| 1117 | 1137 | TDNTFVSGNCDVVIGIVNNT | 0.5226 | 20 | ABCPRED,AAP | 2 | ALLERGEN | Non-Toxin | -0.01 | 0.32 | -0.36 | -2 | 3.57 | 2082.57 | outside |
| 489 | 499 | YFPLQSYGFQ | 0.5043 | 10 | ABCPRED | 1 | ALLERGEN | Non-Toxin | 0.02 | -0.3 | -1.07 | 0 | 5.87 | 1249.53 | outside |
| 159 | 175 | VYSSANNCTFEYVSQP | 0.0327 | 16 | ABCPRED | 1 | NON-ALLERGEN | Non-Toxin | -0.1 | -0.36 | -0.47 | -1 | 4 | 1809.15 | outside |
| 748 | 761 | ECSNLLLQYGSFC | 0.8661 | 13 | BCEPRED | 1 | NON-ALLERGEN | Non-Toxin | 0 | 0.42 | -0.63 | -1 | 4 | 1476.87 | outside |
| 60 | 72 | SNVTWFHAIHVS | 0.8768 | 12 | ABCPRED | 1 | NON-ALLERGEN | Non-Toxin | 0.08 | 0.37 | -0.98 | 1 | 7.26 | 1397.73 | outside |
| 1239 | 1253 | SCCSCLKGCCSCGS | -0.1566 | 14 | BCPREDS | 1 | NON-ALLERGEN | Non-Toxin | -0.07 | 0.78 | -0.26 | 1 | 8.04 | 1340.77 | inside |
| 165 | 175 | NCTFEYVSQP | 0.4213 | 10 | ABCPRED | 1 | ALLERGEN | Non-Toxin | -0.12 | -0.54 | -0.4 | -1 | 4 | 1187.42 | outside |
| 331 | 341 | NITNLCPFGE | 1.237 | 10 | ABCPRED | 1 | ALLERGEN | Non-Toxin | -0.01 | 0.04 | -0.41 | -1 | 4 | 1107.39 | outside |
| 1239 | 1255 | SCCSCLKGCCSCGSCC | -0.2439 | 16 | ABCPRED | 1 | ALLERGEN | Non-Toxin | -0.06 | 0.99 | -0.35 | 1 | 7.94 | 1547.07 | inside |
| 293 | 297 | LDPL | short | 4 | literature | 1 | NON-ALLERGEN | Non-Toxin | 0.07 | 0.62 | -0.15 | -1 | 3.8 | 456.59 | outside |
| 711 | 721 | SIAIPTNFTI | 0.7149 | 10 | ABCPRED | 1 | ALLERGEN | Non-Toxin | 0.17 | 1.08 | -0.87 | 0 | 5.88 | 1076.4 | outside |
| 882 | 894 | ITSGWTFGAGAA | 0.4273 | 12 | ABCPRED,BCPREDS | 2 | ALLERGEN | Non-Toxin | 0.19 | 0.7 | -0.81 | 0 | 5.88 | 1138.41 | outside |
| 885 | 897 | GWTFGAGAALQI | 0.4276 | 12 | ABCPRED | 1 | ALLERGEN | Non-Toxin | 0.22 | 0.85 | -0.93 | 0 | 5.88 | 1191.53 | outside |
| 1162 | 1174 | PDVDLGDISGIN | 1.2343 | 12 | FBCPred | 1 | NON-ALLERGEN | Non-Toxin | -0.02 | -0.02 | 0.22 | -3 | 3.43 | 1214.47 | outside |
| 1154 | 1163 | KYFKNHTSP | -0.2601 | 9 | literature | 1 | NON-ALLERGEN | Non-Toxin | -0.35 | -1.79 | 0.09 | 2.5 | 9.72 | 1121.38 | outside |
| 716 | 728 | TNFTISVTTEIL | 0.8447 | 12 | ABCPRED | 1 | NON-ALLERGEN | Non-Toxin | 0.07 | 0.77 | -0.63 | -1 | 4 | 1338.7 | outside |
| 807 | 815 | PDPSKR | 0.9887 | 6 | ellipro,literature | 2 | ALLERGEN | Non-Toxin | -0.66 | -2.65 | 1.55 | 1 | 9.1 | 698.84 | outside |
| 285 | 297 | ITDAVDCALDPL | 0.5927 | 12 | ABCPRED,AAP | 2 | ALLERGEN | Non-Toxin | 0.04 | 0.8 | -0.03 | -3 | 3.43 | 1245.57 | outside |
| 432 | 442 | CVIAWNSNNL | 0.5242 | 10 | ABCPRED | 1 | ALLERGEN | Non-Toxin | 0.03 | 0.46 | -0.91 | 0 | 5.85 | 1133.43 | outside |
| 53 | 65 | DLFLPFFSNVTW | 0.4627 | 12 | ABCPRED | 1 | NON-ALLERGEN | Non-Toxin | 0.16 | 0.77 | -1.07 | -1 | 3.8 | 1485.87 | outside |
| 254 | 264 | SSSGWTAGAA | 0.167 | 10 | ABCPRED | 1 | ALLERGEN | Non-Toxin | 0.05 | 0.06 | -0.44 | 0 | 5.88 | 894.03 | outside |
| 836 | 848 | QYGDCLGDIAAR | 0.5182 | 12 | ABCPRED | 1 | ALLERGEN | Non-Toxin | -0.15 | -0.22 | 0.11 | -1 | 4.21 | 1281.57 | outside |
| 225 | 237 | PLVDLPIGINIT | 1.3843 | 12 | ABCPRED | 1 | NON-ALLERGEN | Non-Toxin | 0.19 | 1.17 | -0.64 | -1 | 3.8 | 1264.72 | outside |
| 122 | 134 | NATNVVIKVCEF | 0.1918 | 12 | ABCPRED | 1 | NON-ALLERGEN | Non-Toxin | 0.01 | 0.76 | -0.36 | 0 | 6.32 | 1336.74 | outside |
| 682 | 694 | RRARSVASQSII | 0.3202 | 12 | ABCPRED | 1 | NON-ALLERGEN | Non-Toxin | -0.35 | -0.22 | 0.33 | 3 | 12.31 | 1343.71 | outside |
| 493 | 505 | QSYGFQPTNGVG | 0.6735 | 12 | ABCPRED | 1 | NON-ALLERGEN | Non-Toxin | -0.07 | -0.76 | -0.48 | 0 | 5.88 | 1254.5 | outside |
| 686 | 702 | SVASQSIIAYTMSLGA | 0.564 | 16 | ABCPRED | 1 | NON-ALLERGEN | Non-Toxin | 0.11 | 0.95 | -0.69 | 0 | 5.88 | 1599.06 | outside |
| 472 | 484 | IYQAGSTPCNGV | -0.208 | 12 | ABCPRED | 1 | ALLERGEN | Non-Toxin | 0.01 | 0.07 | -0.57 | 0 | 5.84 | 1209.51 | outside |
| 26 | 31 | PAYTN | short | 5 | literature | 1 | ALLERGEN | Non-Toxin | -0.12 | -1.06 | -0.6 | 0 | 5.88 | 564.65 | outside |
| 951 | 971 | VVNQNAQALNTLVKQLSSNF | 0.0245 | 20 | ABCPRED | 1 | NON-ALLERGEN | Non-Toxin | -0.11 | -0.02 | -0.44 | 1 | 9.11 | 2188.78 | outside |
| 590 | 608 | CSFGGVSVITPGTNTSNQ | 0.717 | 18 | ABCPRED | 1 | ALLERGEN | Non-Toxin | -0.02 | 0.02 | -0.44 | 0 | 5.85 | 1769.18 | outside |
| 388 | 404 | NDLCFTNVYADSFVIR | 0.6806 | 16 | ABCPRED | 1 | NON-ALLERGEN | Non-Toxin | -0.07 | 0.33 | -0.38 | -1 | 4.21 | 1877.32 | outside |
| 700 | 712 | GAENSVAYSNNS | 0.3565 | 12 | BCPREDS,FBCPred | 2 | NON-ALLERGEN | Non-Toxin | -0.17 | -0.86 | -0.02 | -1 | 4 | 1212.35 | outside |
| 111 | 121 | DSKTQSLLIV | 0.7617 | 10 | ABCPRED | 1 | NON-ALLERGEN | Non-Toxin | -0.09 | 0.31 | -0.05 | 0 | 6.19 | 1103.43 | outside |
| 486 | 496 | FNCYFPLQSY | 0.8219 | 10 | ABCPRED,BCEPRED | 2 | ALLERGEN | Non-Toxin | 0.02 | -0.01 | -1.17 | 0 | 5.84 | 1281.58 | outside |
| 429 | 439 | FTGCVIAWNS | -0.3731 | 10 | ABCPRED | 1 | ALLERGEN | Non-Toxin | 0.16 | 0.95 | -1.06 | 0 | 5.85 | 1097.39 | outside |
| 1122 | 1140 | VSGNCDVVIGIVNNTVYD | 0.5486 | 18 | ABCPRED | 1 | NON-ALLERGEN | Non-Toxin | 0.04 | 0.63 | -0.44 | -2 | 3.57 | 1881.36 | outside |
| 701 | 711 | AENSVAYSNN | 0.4697 | 10 | ABCPRED | 1 | ALLERGEN | Non-Toxin | -0.2 | -0.91 | -0.06 | -1 | 4 | 1068.19 | outside |
| 486 | 498 | FNCYFPLQSYGF | 0.7566 | 12 | ABCPRED | 1 | NON-ALLERGEN | Non-Toxin | 0.08 | 0.19 | -1.18 | 0 | 5.84 | 1485.84 | outside |
| 654 | 663 | EHVNNSYEC | 1.068 | 9 | IEDB | 1 | NON-ALLERGEN | Non-Toxin | -0.29 | -1.4 | 0.16 | -1.5 | 4.51 | 1094.24 | outside |
| 1052 | 1070 | FPQSAPHGVVFLHVTYVP | 0.6086 | 18 | ABCPRED | 1 | NON-ALLERGEN | Non-Toxin | 0.12 | 0.56 | -0.92 | 1 | 7.25 | 1995.58 | outside |
| 879 | 897 | AGTITSGWTFGAGAALQI | 0.3769 | 18 | ABCPRED | 1 | NON-ALLERGEN | Non-Toxin | 0.17 | 0.77 | -0.78 | 0 | 5.88 | 1722.2 | outside |
| 134 | 142 | QFCNDPFL | -0.243 | 8 | BCEPRED | 1 | ALLERGEN | Non-Toxin | -0.04 | -0.03 | -0.55 | -1 | 3.8 | 983.21 | outside |
| 489 | 505 | YFPLQSYGFQPTNGVG | 0.5744 | 16 | ABCPRED | 1 | ALLERGEN | Non-Toxin | 0.01 | -0.34 | -0.78 | 0 | 5.87 | 1775.19 | outside |
| 330 | 340 | PNITNLCPFG | 1.4887 | 10 | ABCPRED | 1 | NON-ALLERGEN | Non-Toxin | 0.05 | 0.23 | -0.71 | 0 | 5.85 | 1075.39 | outside |
| 387 | 396 | LNDLCFTNV | 2.0179 | 9 | literature | 1 | ALLERGEN | Non-Toxin | 0.01 | 0.66 | -0.62 | -1 | 3.8 | 1038.31 | outside |
| 953 | 969 | NQNAQALNTLVKQLSS | -0.0646 | 16 | ABCPRED | 1 | NON-ALLERGEN | Non-Toxin | -0.2 | -0.5 | -0.22 | 1 | 9.11 | 1729.17 | outside |
| 1051 | 1067 | SFPQSAPHGVVFLHVT | 0.4168 | 16 | ABCPRED | 1 | NON-ALLERGEN | Non-Toxin | 0.09 | 0.5 | -0.78 | 1 | 7.26 | 1723.2 | outside |

| | | | | | | | | | | | | | | | |
|---|---|---|---|---|---|---|---|---|---|---|---|---|---|---|---|
| 1053 | 1067 | PQSAPHGVVFLHVT | 0.6139 | 14 | ABCPRED,AAP | 2 | ALLERGEN | Non-Toxin | 0.08 | 0.43 | -0.73 | 1 | 7.26 | 1488.92 | outside |
| 111 | 127 | DSKTQSLLIVNNATNV | 0.5812 | 16 | ABCPRED | 1 | NON-ALLERGEN | Non-Toxin | -0.14 | -0.13 | -0.14 | 0 | 6.19 | 1717.15 | outside |
| 710 | 720 | NSIAIPTNFT | 0.5036 | 10 | ABCPRED | 1 | NON-ALLERGEN | Non-Toxin | 0.04 | 0.28 | -0.67 | 0 | 5.88 | 1077.34 | outside |
| 23 | 35 | QLPPAYTNSFTR | -0.0689 | 12 | ABCPRED,IEDB | 2 | NON-ALLERGEN | Non-Toxin | -0.2 | -0.82 | -0.35 | 1 | 9.1 | 1394.71 | outside |
| 773 | 781 | EQDKNTQE | 0.0364 | 8 | IEDB | 1 | ALLERGEN | Non-Toxin | -0.66 | -3.2 | 1.52 | -2 | 4.14 | 991.09 | outside |
| 1161 | 1179 | SPDVDLGDISGINASVVN | 0.872 | 18 | ABCPRED | 1 | NON-ALLERGEN | Non-Toxin | -0.01 | 0.27 | -0.01 | -3 | 3.43 | 1772.16 | outside |
| 747 | 765 | TECSNLLLQYGSFCTQLN | 0.7615 | 18 | ABCPRED | 1 | ALLERGEN | Toxin | -0.07 | 0.04 | -0.58 | -1 | 4 | 2034.56 | outside |
| 1159 | 1175 | HTSPDVDLGDISGINA | 1.0848 | 16 | ABCPRED | 1 | NON-ALLERGEN | Non-Toxin | -0.05 | -0.19 | 0.09 | -2.5 | 3.93 | 1610.93 | outside |
| 62 | 72 | VTWFHAIHVS | 0.7333 | 10 | ABCPRED | 1 | NON-ALLERGEN | Non-Toxin | 0.18 | 0.87 | -1.23 | 1 | 7.26 | 1196.52 | outside |
| 25 | 35 | PPAYTNSFTR | -0.4294 | 10 | ABCPRED | 1 | NON-ALLERGEN | Non-Toxin | -0.23 | -1.01 | -0.26 | 1 | 9.1 | 1153.38 | outside |
| 90 | 98 | VYFASTEK | 0.9206 | 8 | BCEPRED | 1 | NON-ALLERGEN | Non-Toxin | -0.09 | -0.17 | -0.11 | 0 | 6.35 | 944.15 | outside |
| 359 | 372 | SNCVADYSVLYNS | -0.1828 | 13 | IEDB,literature | 2 | ALLERGEN | Non-Toxin | -0.06 | 0.08 | -0.51 | -1 | 3.8 | 1434.71 | outside |
| 265 | 274 | YYVGYLQPR | 1.4692 | 9 | BCEPRED | 1 | NON-ALLERGEN | Non-Toxin | -0.14 | -0.66 | -0.78 | 1 | 8.83 | 1158.45 | outside |
| 857 | 869 | GLTVLPPLLTDE | 0.5505 | 12 | ABCPRED | 1 | ALLERGEN | Non-Toxin | 0.08 | 0.62 | -0.29 | -2 | 3.67 | 1267.67 | outside |
| 1119 | 1129 | NTFVSGNCDV | -0.5646 | 10 | IEDB | 1 | NON-ALLERGEN | Non-Toxin | -0.06 | 0.13 | -0.32 | -1 | 3.8 | 1055.26 | outside |
| 492 | 502 | LQSYGFQPTN | 1.043 | 10 | ABCPRED | 1 | ALLERGEN | Non-Toxin | -0.12 | -0.87 | -0.61 | 0 | 5.88 | 1154.39 | outside |
| 589 | 609 | PCSFGGVSVITPGTNTSNQV | 0.7278 | 20 | ABCPRED,BCPREDS | 2 | NON-ALLERGEN | Non-Toxin | 0 | 0.15 | -0.48 | 0 | 5.85 | 1965.46 | outside |
| 282 | 298 | NGTITDAVDCALDPLS | 0.5811 | 16 | ABCPRED | 1 | ALLERGEN | Non-Toxin | -0.03 | 0.26 | -0.01 | -3 | 3.43 | 1604.97 | outside |
| 837 | 843 | YGDCLG | -0.4941 | 6 | IEDB | 1 | ALLERGEN | Non-Toxin | 0.03 | 0.12 | -0.35 | -1 | 3.8 | 626.76 | outside |
| 427 | 437 | DDFTGCVIAW | -0.3876 | 10 | ABCPRED | 1 | ALLERGEN | Non-Toxin | 0.11 | 0.68 | -0.51 | -2 | 3.57 | 1126.38 | outside |
| 910 | 922 | GVTQNVLYENQK | 0.3633 | 12 | ABCPRED | 1 | NON-ALLERGEN | Non-Toxin | -0.23 | -0.97 | -0.06 | 0 | 6.35 | 1392.72 | outside |
| 876 | 894 | ALLAGTITSGWTFGAGAA | 0.3994 | 18 | ABCPRED | 1 | NON-ALLERGEN | Non-Toxin | 0.21 | 1.03 | -0.82 | 0 | 5.88 | 1665.14 | outside |
| 494 | 508 | SYGFQPTNGVGYQP | 0.7373 | 14 | literature | 1 | NON-ALLERGEN | Non-Toxin | -0.07 | -0.86 | -0.58 | 0 | 5.87 | 1514.82 | outside |
| 744 | 756 | GDSTECSNLLLQ | 0.2589 | 12 | ABCPRED | 1 | NON-ALLERGEN | Non-Toxin | -0.13 | -0.23 | 0.02 | -2 | 3.67 | 1279.56 | outside |
| 692 | 702 | IIAYTMSLGA | 0.681 | 10 | ABCPRED | 1 | ALLERGEN | Non-Toxin | 0.25 | 1.51 | -1.01 | 0 | 5.88 | 1039.4 | outside |
| 358 | 370 | ISNCVADYSVLY | -0.4142 | 12 | ABCPRED | 1 | NON-ALLERGEN | Non-Toxin | 0.07 | 0.82 | -0.74 | -1 | 3.8 | 1346.68 | outside |
| 747 | 763 | TECSNLLLQYGSFCTQ | 0.755 | 16 | ABCPRED | 1 | NON-ALLERGEN | Non-Toxin | -0.07 | 0.03 | -0.55 | -1 | 4 | 1807.26 | outside |
| 359 | 377 | SNCVADYSVLYNSASFST | 0.062 | 18 | ABCPRED | 1 | ALLERGEN | Non-Toxin | -0.04 | 0.18 | -0.52 | -1 | 3.8 | 1928.29 | outside |
| 64 | 76 | WFHAIHVSGTNG | 0.5744 | 12 | ABCPRED | 1 | ALLERGEN | Non-Toxin | 0.08 | 0.02 | -0.88 | 1 | 7.26 | 1325.63 | outside |
| 691 | 701 | SIIAYTMSLG | 0.746 | 10 | ABCPRED | 1 | ALLERGEN | Non-Toxin | 0.2 | 1.25 | -0.93 | 0 | 5.88 | 1055.4 | outside |
| 698 | 710 | SLGAENSVAYSN | 0.7274 | 12 | ABCPRED | 1 | NON-ALLERGEN | Non-Toxin | -0.08 | -0.25 | -0.19 | -1 | 4 | 1211.41 | outside |
| 118 | 132 | LIVNNATNVVIKVC | -0.0859 | 14 | FBCPred | 1 | NON-ALLERGEN | Non-Toxin | 0.09 | 1.34 | -0.69 | 1 | 8.57 | 1500.05 | outside |
| 23 | 33 | QLPPAYTNSF | 0.3474 | 10 | ABCPRED | 1 | NON-ALLERGEN | Non-Toxin | -0.05 | -0.46 | -0.68 | 0 | 5.88 | 1137.39 | outside |
| 1162 | 1180 | PDVDLGDISGINASVVNI | 1.0023 | 18 | ABCPRED | 1 | NON-ALLERGEN | Non-Toxin | 0.05 | 0.57 | -0.12 | -3 | 3.43 | 1798.25 | outside |
| 588 | 604 | TPCSFGGVSVITPGTN | 0.8122 | 16 | ABCPRED | 1 | NON-ALLERGEN | Non-Toxin | 0.07 | 0.41 | -0.54 | 0 | 5.85 | 1536.95 | outside |
| 937 | 949 | SLSSTASALGKL | 0.7104 | 12 | ABCPRED | 1 | ALLERGEN | Non-Toxin | -0.01 | 0.57 | -0.22 | 1 | 9.11 | 1134.46 | outside |
| 493 | 507 | QSYGFQPTNGVGYQ | 0.6697 | 14 | ABCPRED,literature | 2 | NON-ALLERGEN | Non-Toxin | -0.11 | -0.99 | -0.56 | 0 | 5.87 | 1545.84 | outside |
| 161 | 173 | SSANNCTFEYVS | 0.2066 | 12 | ABCPRED | 1 | NON-ALLERGEN | Non-Toxin | -0.12 | -0.3 | -0.32 | -1 | 4 | 1321.53 | outside |
| 872 | 890 | QYTSALLAGTITSGWTFG | 0.4406 | 18 | ABCPRED | 1 | NON-ALLERGEN | Non-Toxin | 0.1 | 0.4 | -0.86 | 0 | 5.88 | 1874.35 | outside |
| 1158 | 1174 | NHTSPDVDLGDISGIN | 1.1399 | 16 | ABCPRED | 1 | ALLERGEN | Non-Toxin | -0.11 | -0.53 | 0.14 | -2.5 | 3.93 | 1653.96 | outside |
| 873 | 891 | YTSALLAGTITSGWTFGA | 0.3925 | 18 | ABCPRED | 1 | NON-ALLERGEN | Non-Toxin | 0.15 | 0.69 | -0.89 | 0 | 5.88 | 1817.29 | outside |
| 699 | 715 | LGAENSVAYSNNSIAI | 0.7636 | 16 | ABCPRED | 1 | NON-ALLERGEN | Non-Toxin | 0.01 | 0.27 | -0.39 | -1 | 4 | 1622.98 | outside |
| 875 | 893 | SALLAGTITSGWTFGAGA | 0.3611 | 18 | ABCPRED | 1 | NON-ALLERGEN | Non-Toxin | 0.19 | 0.88 | -0.77 | 0 | 5.88 | 1681.14 | outside |
| 587 | 605 | ITPCSFGGVSVITPGTNT | 0.781 | 18 | ABCPRED | 1 | ALLERGEN | Non-Toxin | 0.09 | 0.58 | -0.61 | 0 | 5.85 | 1751.25 | outside |
| 857 | 868 | GLTVLPPLLTD | 0.5484 | 11 | BCEPRED | 1 | ALLERGEN | Non-Toxin | 0.15 | 0.99 | -0.59 | -1 | 3.8 | 1138.54 | outside |
| 914 | 921 | NVLYENQ | 0.4689 | 7 | IEDB | 1 | ALLERGEN | Non-Toxin | -0.21 | -1.04 | -0.29 | -1 | 4 | 879.04 | outside |
| 138 | 148 | DPFLGVYYHK | 0.9542 | 10 | ABCPRED | 1 | ALLERGEN | Non-Toxin | -0.04 | -0.44 | -0.49 | 0.5 | 7.09 | 1238.55 | outside |
| 510 | 522 | VVVLSFELLHAP | 0.8552 | 12 | ABCPRED | 1 | NON-ALLERGEN | Non-Toxin | 0.23 | 1.63 | -0.84 | -0.5 | 5.25 | 1323.78 | outside |
| 162 | 172 | SANNCTFEYV | 0.0177 | 10 | ABCPRED | 1 | ALLERGEN | Non-Toxin | -0.09 | -0.2 | -0.45 | -1 | 4 | 1147.35 | outside |
| 360 | 372 | NCVADYSVLYNS | -0.1824 | 12 | ABCPRED | 1 | ALLERGEN | Non-Toxin | -0.05 | 0.15 | -0.57 | -1 | 3.8 | 1347.62 | outside |
| 856 | 868 | NGLTVLPPLLTD | 0.2568 | 12 | ABCPRED | 1 | ALLERGEN | Non-Toxin | 0.08 | 0.62 | -0.53 | -1 | 3.8 | 1252.66 | outside |
| 1121 | 1139 | FVSGNCDVVIGIVNNTVY | 0.4356 | 18 | ABCPRED | 1 | NON-ALLERGEN | Non-Toxin | 0.11 | 0.98 | -0.74 | -1 | 3.8 | 1913.45 | outside |
| 495 | 509 | YGFQPTNGVGYQPY | 0.7464 | 14 | ABCPRED | 1 | ALLERGEN | Non-Toxin | -0.05 | -0.89 | -0.76 | 0 | 5.87 | 1590.92 | outside |
| 1192 | 1204 | NLNESLIDLQEL | 0.8502 | 12 | FBCPred | 1 | NON-ALLERGEN | Non-Toxin | -0.11 | -0.18 | 0.07 | -3 | 3.58 | 1400.74 | outside |
| 255 | 271 | SSGWTAGAAAYYVGYL | 0.417 | 16 | ABCPRED | 1 | NON-ALLERGEN | Non-Toxin | 0.14 | 0.43 | -0.96 | 0 | 5.87 | 1637 | outside |
| 87 | 97 | NDGVYFASTE | 0.2633 | 10 | ABCPRED | 1 | ALLERGEN | Non-Toxin | -0.08 | -0.49 | -0.07 | -2 | 3.67 | 1102.25 | outside |
| 653 | 662 | AEHVNNSYE | 0.8405 | 9 | BCEPRED | 1 | NON-ALLERGEN | Non-Toxin | -0.26 | -1.48 | 0.21 | -1.5 | 4.51 | 1062.18 | outside |
| 1057 | 1070 | PHGVVFLHVTYVP | 0.9305 | 13 | BCEPRED | 1 | ALLERGEN | Non-Toxin | 0.18 | 0.88 | -1.08 | 1 | 7.25 | 1464.93 | outside |
| 1115 | 1127 | ITTDNTFVSGNC | 0.1449 | 12 | BCPREDS | 1 | NON-ALLERGEN | Non-Toxin | -0.06 | 0.02 | -0.36 | -1 | 3.8 | 1271.53 | outside |
| 664 | 680 | IPIGAGICASYQTQTN | 0.9249 | 16 | ABCPRED | 1 | NON-ALLERGEN | Non-Toxin | 0.02 | 0.2 | -0.6 | 0 | 5.84 | 1637.08 | outside |
| 661 | 673 | ECDIPIGAGICA | 0.597 | 12 | BCPREDS | 1 | ALLERGEN | Non-Toxin | 0.14 | 1.06 | -0.2 | -2 | 3.67 | 1161.52 | outside |
| 359 | 369 | SNCVADYSVL | -0.1444 | 10 | ABCPRED | 1 | ALLERGEN | Non-Toxin | 0 | 0.66 | -0.48 | -1 | 3.8 | 1070.31 | outside |
| 908 | 920 | GIGVTQNVLYEN | 0.7229 | 12 | ABCPRED | 1 | NON-ALLERGEN | Non-Toxin | -0.01 | -0.01 | -0.47 | -1 | 4 | 1306.63 | outside |
| 1053 | 1069 | PQSAPHGVVFLHVTYV | 0.7839 | 16 | ABCPRED | 1 | NON-ALLERGEN | Non-Toxin | 0.1 | 0.56 | -0.88 | 1 | 7.25 | 1751.26 | outside |
| 1057 | 1071 | PHGVVFLHVTYVPA | 0.8058 | 14 | ABCPRED,IEDB,literature | 3 | ALLERGEN | Non-Toxin | 0.19 | 0.94 | -1.04 | 1 | 7.25 | 1536.02 | outside |
| 391 | 401 | CFTNVYADSF | -0.1717 | 10 | ABCPRED | 1 | ALLERGEN | Non-Toxin | 0.03 | 0.43 | -0.72 | -1 | 3.8 | 1166.39 | outside |
| 159 | 171 | VYSSANNCTFEY | -0.0967 | 12 | ABCPRED,BCPREDS,FBCPred | 3 | NON-ALLERGEN | Non-Toxin | -0.09 | -0.34 | -0.54 | -1 | 4 | 1397.63 | outside |
| 289 | 299 | VDCALDPLSE | 0.4584 | 10 | ABCPRED | 1 | ALLERGEN | Non-Toxin | -0.05 | 0.32 | 0.27 | -3 | 3.5 | 1061.3 | outside |
| 1159 | 1177 | HTSPDVDLGDISGINASV | 0.9036 | 18 | ABCPRED | 1 | NON-ALLERGEN | Non-Toxin | -0.03 | 0.02 | 0.02 | -2.5 | 3.93 | 1797.17 | outside |
| 591 | 607 | SFGGVSVITPGTNTSN | 0.6915 | 16 | ABCPRED | 1 | NON-ALLERGEN | Non-Toxin | 0.01 | 0.09 | -0.45 | 0 | 5.88 | 1537.88 | outside |
| 878 | 890 | LAGTITSGWTFG | 0.3445 | 12 | ABCPRED | 1 | NON-ALLERGEN | Non-Toxin | 0.18 | 0.66 | -0.91 | 0 | 5.88 | 1210.53 | outside |
| 773 | 780 | EQDKNTQ | 0.1017 | 7 | IEDB,literature | 2 | ALLERGEN | Non-Toxin | -0.66 | -3.16 | 1.31 | -1 | 4.38 | 861.96 | outside |
| 705 | 723 | VAYSNNSIAIPTNFTISV | 0.8012 | 18 | ABCPRED | 1 | NON-ALLERGEN | Non-Toxin | 0.07 | 0.62 | -0.75 | 0 | 5.88 | 1911.4 | outside |
| 950 | 967 | DVVNQNAQALNTLVKQL | 0.032 | 17 | COVIDep,literature | 2 | NON-ALLERGEN | Non-Toxin | -0.13 | -0.09 | -0.24 | 0 | 6.19 | 1868.39 | outside |
| 362 | 374 | VADYSVLYNSAS | 0.0838 | 12 | ABCPRED | 1 | NON-ALLERGEN | Non-Toxin | 0 | 0.32 | -0.53 | -1 | 3.8 | 1288.53 | outside |
| 487 | 505 | NCYFPLQSYGFQPTNGVG | 0.6928 | 18 | ABCPRED | 1 | ALLERGEN | Non-Toxin | -0.02 | -0.36 | -0.73 | 0 | 5.84 | 1992.46 | outside |
| 254 | 266 | SSSGWTAGAAAY | 0.3276 | 12 | ABCPRED | 1 | NON-ALLERGEN | Non-Toxin | 0.06 | 0.09 | -0.6 | 0 | 5.88 | 1128.31 | outside |

| | | | | | | | | | | | | | | | |
|---|---|---|---|---|---|---|---|---|---|---|---|---|---|---|---|
| 3 | 19 | VFLVLLPLVSSQCVNL | 0.7784 | 16 | ABCPRED | 1 | NON-ALLERGEN | Non-Toxin | 0.22 | 1.93 | -1.09 | 0 | 5.85 | 1744.42 | outside |
| 699 | 709 | LGAENSVAYS | 0.5507 | 10 | ABCPRED | 1 | ALLERGEN | Non-Toxin | 0 | 0.13 | -0.28 | -1 | 4 | 1010.2 | outside |
| 607 | 617 | QVAVLYQDVN | 0.4619 | 10 | ABCPRED | 1 | NON-ALLERGEN | Non-Toxin | -0.03 | 0.29 | -0.55 | -1 | 3.8 | 1148.43 | outside |
| 164 | 174 | NNCTFEYVSQ | 0.2109 | 10 | ABCPRED | 1 | NON-ALLERGEN | Non-Toxin | -0.18 | -0.73 | -0.38 | -1 | 4 | 1204.41 | outside |
| 258 | 268 | WTAGAAAYYV | 0.5371 | 10 | ABCPRED | 1 | ALLERGEN | Non-Toxin | 0.19 | 0.68 | -1.19 | 0 | 5.87 | 1072.31 | outside |
| 892 | 901 | AALQIPFAM | 0.7747 | 9 | ellipro | 1 | NON-ALLERGEN | Non-Toxin | 0.24 | 1.48 | -0.97 | 0 | 5.88 | 961.31 | outside |
| 292 | 299 | ALDPLSE | 0.5276 | 7 | IEDB | 1 | ALLERGEN | Non-Toxin | -0.05 | 0 | 0.31 | -2 | 3.67 | 743.9 | outside |
| 1161 | 1177 | SPDVDLGDISGINASV | 0.8446 | 16 | ABCPRED | 1 | NON-ALLERGEN | Non-Toxin | 0 | 0.26 | 0.08 | -3 | 3.43 | 1558.89 | outside |
| 145 | 153 | YHKNNKSW | 0.4099 | 8 | BCEPRED,IEDB | 2 | ALLERGEN | Non-Toxin | -0.47 | -2.62 | 0.06 | 2.5 | 9.72 | 1076.29 | outside |
| 591 | 609 | SFGGVSVITPGTNTSNQV | 0.6356 | 18 | ABCPRED,AAP | 2 | NON-ALLERGEN | Non-Toxin | 0 | 0.12 | -0.47 | 0 | 5.88 | 1765.18 | outside |
| 473 | 485 | YQAGSTPCNGVE | 0.224 | 12 | ABCPRED | 1 | NON-ALLERGEN | Non-Toxin | -0.11 | -0.6 | -0.17 | -1 | 4 | 1225.46 | outside |
| 390 | 400 | LCFTNVYADS | 0.7072 | 10 | ABCPRED | 1 | NON-ALLERGEN | Non-Toxin | 0.02 | 0.53 | -0.65 | -1 | 3.8 | 1132.38 | outside |
| 58 | 68 | FFSNVTWFHA | 0.4074 | 10 | ABCPRED | 1 | NON-ALLERGEN | Non-Toxin | 0.15 | 0.53 | -1.33 | 0.5 | 7.1 | 1255.53 | outside |
| 258 | 270 | WTAGAAAYYVGY | 0.6185 | 12 | ABCPRED | 1 | NON-ALLERGEN | Non-Toxin | 0.18 | 0.43 | -1.18 | 0 | 5.87 | 1292.57 | outside |
| 1239 | 1257 | SCCSCLKGCCSCGSCCKF | -0.0728 | 18 | ABCPRED | 1 | ALLERGEN | Non-Toxin | -0.08 | 0.82 | -0.28 | 2 | 8.33 | 1822.45 | inside |
| 112 | 128 | SKTQSLLIVNNATNVV | 0.5832 | 16 | ABCPRED | 1 | NON-ALLERGEN | Non-Toxin | -0.06 | 0.35 | -0.42 | 1 | 9.11 | 1701.2 | outside |
| 488 | 496 | CYFPLQSY | 0.9394 | 8 | IEDB,literature | 2 | ALLERGEN | Non-Toxin | 0.02 | 0.07 | -1.17 | 0 | 5.84 | 1020.27 | outside |
| 392 | 402 | FTNVYADSFV | -0.5014 | 10 | ABCPRED | 1 | ALLERGEN | Non-Toxin | 0.08 | 0.6 | -0.77 | -1 | 3.8 | 1162.39 | outside |
| 524 | 534 | VCGPKKSTNL | 0.0947 | 10 | ABCPRED | 1 | ALLERGEN | Non-Toxin | -0.21 | -0.43 | 0.18 | 2 | 9.36 | 1046.39 | outside |
| 46 | 58 | SVLHSTQDLFLP | 0.3035 | 12 | ABCPRED | 1 | NON-ALLERGEN | Non-Toxin | 0.01 | 0.36 | -0.54 | -0.5 | 5.09 | 1356.72 | outside |
| 618 | 628 | TEVPVAIHAD | 0.1126 | 10 | ABCPRED | 1 | ALLERGEN | Non-Toxin | 0.03 | 0.4 | -0.07 | -1.5 | 4.36 | 1051.3 | outside |
| 287 | 299 | DAVDCALDPLSE | 0.4675 | 12 | ABCPRED | 1 | ALLERGEN | Non-Toxin | -0.08 | 0.12 | 0.43 | -4 | 3.38 | 1247.49 | outside |
| 181 | 189 | GKQGNFKN | 1.0999 | 8 | IEDB | 1 | ALLERGEN | Non-Toxin | -0.41 | -2.04 | 0.51 | 2 | 10.02 | 892.1 | outside |
| 394 | 404 | NVYADSFVIR | -0.321 | 10 | ABCPRED | 1 | NON-ALLERGEN | Non-Toxin | -0.07 | 0.39 | -0.36 | 0 | 6.19 | 1183.46 | outside |
| 256 | 272 | SGWTAGAAAYYVGYLQ | 0.4265 | 16 | ABCPRED | 1 | ALLERGEN | Non-Toxin | 0.12 | 0.26 | -0.97 | 0 | 5.87 | 1678.06 | outside |
| 1256 | 1262 | FDEDDS | -0.0685 | 6 | IEDB | 1 | ALLERGEN | Non-Toxin | -0.4 | -2 | 1.63 | -4 | 3.38 | 726.71 | inside |
| 1203 | 1207 | LGKY | short | 4 | literature | 1 | ALLERGEN | Non-Toxin | -0.1 | -0.45 | -0.27 | 1 | 8.94 | 479.63 | outside |
| 473 | 484 | YQAGSTPCNGV | 0.0881 | 11 | IEDB,literature | 2 | NON-ALLERGEN | Non-Toxin | -0.06 | -0.34 | -0.45 | 0 | 5.84 | 1096.33 | outside |
| 1241 | 1257 | CSCLKGCCSCGSCCKF | -0.1298 | 16 | ABCPRED | 1 | NON-ALLERGEN | Toxin | -0.08 | 0.82 | -0.28 | 2 | 8.38 | 1632.21 | inside |
| 884 | 896 | SGWTFGAGAALQ | 0.267 | 12 | ABCPRED | 1 | NON-ALLERGEN | Non-Toxin | 0.13 | 0.41 | -0.76 | 0 | 5.88 | 1165.44 | outside |
| 256 | 266 | SGWTAGAAAY | 0.2756 | 10 | ABCPRED | 1 | NON-ALLERGEN | Non-Toxin | 0.13 | 0.27 | -0.78 | 0 | 5.88 | 954.13 | outside |
| 591 | 603 | SFGGVSVITPGT | 0.6877 | 12 | ABCPRED | 1 | NON-ALLERGEN | Non-Toxin | 0.16 | 0.83 | -0.63 | 0 | 5.88 | 1121.43 | outside |
| 522 | 532 | ATVCGPKKST | -0.1151 | 10 | BCEPRED | 1 | NON-ALLERGEN | Non-Toxin | -0.19 | -0.35 | 0.25 | 2 | 9.36 | 991.3 | outside |
| 703 | 721 | NSVAYSNNSIAIPTNFTI | 0.6865 | 18 | ABCPRED | 1 | NON-ALLERGEN | Non-Toxin | 0.01 | 0.19 | -0.66 | 0 | 5.88 | 1926.37 | outside |
| 879 | 893 | AGTITSGWTFGAGA | 0.3304 | 14 | ABCPRED,FBCPred | 2 | NON-ALLERGEN | Non-Toxin | 0.16 | 0.52 | -0.72 | 0 | 5.88 | 1296.6 | outside |
| 159 | 177 | VYSSANNCTFEYVSQPFL | 0.0464 | 18 | ABCPRED | 1 | ALLERGEN | Non-Toxin | -0.03 | 0.04 | -0.66 | -1 | 4 | 2069.52 | outside |
| 1227 | 1257 | IVMVTIMLCCMTSCCSCLKGCCSCGSCCKF | 0.0464 | 30 | BCEPRED | 1 | NON-ALLERGEN | Non-Toxin | 0.07 | 1.51 | -0.67 | 2 | 8.23 | 3162.46 | transmembrane-region_inside |
| 1136 | 1144 | TVYDPLQP | 0.3135 | 8 | BCEPRED | 1 | ALLERGEN | Non-Toxin | -0.08 | -0.53 | -0.35 | -1 | 3.8 | 932.15 | outside |
| 931 | 943 | IGKIQDSLSSTA | 0.5285 | 12 | ABCPRED | 1 | NON-ALLERGEN | Non-Toxin | -0.09 | 0.02 | 0.07 | 0 | 6.19 | 1219.53 | outside |
| 1214 | 1232 | WYIWLGFIAGLIAIVMVT | 0.7101 | 18 | ABCPRED | 1 | NON-ALLERGEN | Non-Toxin | 0.41 | 2.09 | -1.56 | 0 | 5.88 | 2066.87 | transmembrane-region |
| 566 | 578 | GRDIADTTDAVR | 0.4565 | 12 | ABCPRED | 1 | NON-ALLERGEN | Non-Toxin | -0.34 | -0.75 | 0.82 | -1 | 4.43 | 1289.52 | outside |
| 373 | 385 | SFSTFKCYGVSP | 0.8172 | 12 | ABCPRED | 1 | NON-ALLERGEN | Non-Toxin | -0.01 | 0.17 | -0.53 | 1 | 8.54 | 1322.65 | outside |
| 531 | 545 | TNLVKNKCVNFNFN | 1.2489 | 14 | ABCPRED,BCPREDS | 2 | NON-ALLERGEN | Non-Toxin | -0.19 | -0.41 | -0.3 | 2 | 9.36 | 1655.11 | outside |
| 517 | 535 | LLHAPATVCGPKKSTNLV | 0.0749 | 18 | ABCPRED | 1 | ALLERGEN | Non-Toxin | -0.04 | 0.31 | -0.29 | 2.5 | 9.36 | 1849.49 | outside |
| 1165 | 1183 | DLGDISGINASVVNIQKE | 0.9087 | 18 | ABCPRED | 1 | NON-ALLERGEN | Non-Toxin | -0.07 | 0.01 | 0.14 | -2 | 4.03 | 1872.34 | outside |
| 250 | 261 | TPGDSSSGWTA | 0.2473 | 11 | IEDB,literature | 2 | ALLERGEN | Non-Toxin | -0.09 | -0.8 | -0.07 | -1 | 3.8 | 1065.2 | outside |
| 545 | 559 | GLTGTGVLTESNKK | 1.0227 | 14 | IEDB | 1 | NON-ALLERGEN | Non-Toxin | -0.16 | -0.51 | 0.23 | 1 | 8.94 | 1404.8 | outside |
| 731 | 739 | MTKTSVDC | 1.5932 | 8 | IEDB | 1 | ALLERGEN | Non-Toxin | -0.2 | -0.12 | 0.21 | 0 | 6.16 | 884.13 | outside |
| 675 | 688 | QTQTNSPRRARSV | 0.1763 | 13 | IEDB,literature | 2 | NON-ALLERGEN | Non-Toxin | -0.57 | -1.74 | 0.57 | 3 | 12.31 | 1500.81 | outside |
| 468 | 484 | ISTEIYQAGSTPCNGV | -0.0518 | 16 | ABCPRED | 1 | NON-ALLERGEN | Non-Toxin | -0.02 | 0.02 | -0.36 | -1 | 4 | 1640.03 | outside |
| 744 | 762 | GDSTECSNLLLQYGSFCT | 0.5362 | 18 | ABCPRED | 1 | NON-ALLERGEN | Toxin | -0.07 | -0.04 | -0.32 | -2 | 3.67 | 1938.37 | outside |
| 716 | 730 | TNFTISVTTEILPV | 1.1495 | 14 | FBCPred | 1 | NON-ALLERGEN | Non-Toxin | 0.1 | 0.84 | -0.64 | -1 | 4 | 1534.98 | outside |
| 768 | 780 | TGIAVEQDKNTQ | 0.8219 | 12 | ABCPRED | 1 | NON-ALLERGEN | Non-Toxin | -0.26 | -1.06 | 0.42 | -1 | 4.38 | 1303.57 | outside |
| 103 | 113 | GWIFGTTLDS | -0.0845 | 10 | ABCPRED | 1 | ALLERGEN | Non-Toxin | 0.12 | 0.37 | -0.7 | -1 | 3.8 | 1096.35 | outside |
| 519 | 535 | HAPATVCGPKKSTNLV | 0.1852 | 16 | ABCPRED | 1 | NON-ALLERGEN | Non-Toxin | -0.11 | -0.12 | -0.1 | 2.5 | 9.36 | 1623.13 | outside |
| 693 | 711 | IAYTMSLGAENSVAYSNN | 0.74 | 18 | ABCPRED | 1 | NON-ALLERGEN | Non-Toxin | -0.03 | -0.02 | -0.47 | -1 | 4 | 1905.32 | outside |
| 959 | 977 | LNTLVKQLSSNFGAISSV | 0.2018 | 18 | ABCPRED | 1 | NON-ALLERGEN | Non-Toxin | 0.01 | 0.57 | -0.49 | 1 | 9.11 | 1878.43 | outside |
| 469 | 481 | STEIYQAGSTPC | 0.131 | 12 | ABCPRED | 1 | ALLERGEN | Non-Toxin | -0.09 | -0.38 | -0.22 | -1 | 4 | 1256.51 | outside |
| 251 | 261 | PGDSSSGWTA | 0.221 | 10 | ABCPRED | 1 | ALLERGEN | Non-Toxin | -0.08 | -0.81 | -0.04 | -1 | 3.8 | 964.08 | outside |
| 118 | 134 | LIVNNATNVVIKVCEF | -0.0046 | 16 | ABCPRED | 1 | NON-ALLERGEN | Non-Toxin | 0.08 | 1.13 | -0.57 | 0 | 6.32 | 1776.37 | outside |
| 676 | 688 | TQTNSPRRARSV | 0.1597 | 12 | FBCPred | 1 | NON-ALLERGEN | Non-Toxin | -0.56 | -1.59 | 0.6 | 3 | 12.31 | 1372.66 | outside |
| 734 | 746 | TSVDCTMYICGD | -0.4357 | 12 | ABCPRED | 1 | NON-ALLERGEN | Non-Toxin | -0.02 | 0.39 | -0.28 | -2 | 3.57 | 1307.63 | outside |
| 473 | 491 | YQAGSTPCNGVEGFNCYF | 0.3538 | 18 | ABCPRED | 1 | NON-ALLERGEN | Non-Toxin | -0.03 | -0.24 | -0.56 | -1 | 4 | 1957.37 | outside |
| 526 | 538 | GPKKSTNLVKNK | 0.4692 | 12 | ellipro,IEDB | 2 | NON-ALLERGEN | Non-Toxin | -0.41 | -1.51 | 0.75 | 4 | 10.49 | 1313.74 | outside |
| 533 | 547 | LVKNKCVNFNFNGL | 1.3128 | 14 | FBCPred | 1 | NON-ALLERGEN | Non-Toxin | -0.09 | 0.14 | -0.41 | 2 | 9.36 | 1610.12 | outside |
| 1157 | 1166 | KNHTSPDVD | 0.7809 | 9 | BCEPRED | 1 | NON-ALLERGEN | Non-Toxin | -0.39 | -1.83 | 0.79 | -0.5 | 5.22 | 1012.16 | outside |
| 657 | 667 | NNSYECDIPI | 0.4879 | 10 | ABCPRED | 1 | ALLERGEN | Non-Toxin | -0.14 | -0.62 | -0.02 | -2 | 3.67 | 1167.39 | outside |
| 929 | 941 | SAIGKIQDSLSS | 0.3792 | 12 | ABCPRED | 1 | NON-ALLERGEN | Non-Toxin | -0.1 | 0.01 | 0.12 | 0 | 6.19 | 1205.5 | outside |
| 544 | 560 | NGLTGTGVLTESNKKF | 0.6212 | 16 | ABCPRED | 1 | NON-ALLERGEN | Non-Toxin | -0.14 | -0.49 | 0.06 | 1 | 8.94 | 1666.11 | outside |
| 942 | 954 | ASALGKLQDVVN | 0.5413 | 12 | ABCPRED | 1 | ALLERGEN | Non-Toxin | -0.05 | 0.33 | -0.08 | 0 | 6.19 | 1214.56 | outside |
| 482 | 498 | GVEGFNCYFPLQSYGF | 0.7015 | 16 | ABCPRED | 1 | NON-ALLERGEN | Non-Toxin | 0.07 | 0.14 | -0.79 | -1 | 4 | 1828.26 | outside |
| 944 | 956 | ALGKLQDVVNQN | 0.5149 | 12 | ABCPRED | 1 | NON-ALLERGEN | Non-Toxin | -0.16 | -0.33 | -0.02 | 0 | 6.19 | 1298.65 | outside |
| 12 | 24 | SSQCVNLTTRTQ | 1.5168 | 12 | ABCPRED | 1 | NON-ALLERGEN | Non-Toxin | -0.31 | -0.68 | -0.11 | 1 | 8.6 | 1337.64 | outside |
| 533 | 551 | LVKNKCVNFNFNGLTGTG | 1.2501 | 18 | ABCPRED | 1 | NON-ALLERGEN | Non-Toxin | -0.07 | -0.02 | -0.37 | 2 | 9.36 | 1926.5 | outside |
| 1235 | 1251 | CCMTSCCSCLKGCCSC | -0.2666 | 16 | ABCPRED | 1 | ALLERGEN | Toxin | -0.05 | 1.14 | -0.48 | 1 | 7.94 | 1635.24 | transmembrane-region_inside |
| 943 | 961 | SALGKLQDVVNQNAQALN | 0.5362 | 18 | ABCPRED | 1 | NON-ALLERGEN | Non-Toxin | -0.14 | -0.24 | -0.13 | 0 | 6.19 | 1883.37 | outside |
| 535 | 553 | KNKCVNFNFNGLTGTGVL | 1.3849 | 18 | ABCPRED | 1 | NON-ALLERGEN | Non-Toxin | -0.07 | -0.02 | -0.37 | 2 | 9.36 | 1926.5 | outside |

| | | | | | | | | | | | | | | | |
|---|---|---|---|---|---|---|---|---|---|---|---|---|---|---|---|
| 770 | 779 | IAVEQDKNT | 0.4823 | 9 | BCEPRED | 1 | ALLERGEN | Non-Toxin | -0.27 | -0.9 | 0.58 | -1 | 4.38 | 1017.23 | outside |
| 942 | 960 | ASALGKLQDVVNQNAQAL | 0.4932 | 18 | ABCPRED | 1 | NON-ALLERGEN | Non-Toxin | -0.09 | 0.05 | -0.17 | 0 | 6.19 | 1840.34 | outside |
| 48 | 60 | LHSTQDLFLPFF | 0.3186 | 12 | ABCPRED | 1 | NON-ALLERGEN | Non-Toxin | 0.09 | 0.54 | -0.86 | -0.5 | 5.09 | 1464.86 | outside |
| 609 | 621 | AVLYQDVNCTEV | 0.9218 | 12 | ABCPRED | 1 | NON-ALLERGEN | Non-Toxin | -0.03 | 0.39 | -0.34 | -2 | 3.67 | 1353.68 | outside |
| 943 | 959 | SALGKLQDVVNQNAQA | 0.4707 | 16 | ABCPRED | 1 | ALLERGEN | Non-Toxin | -0.15 | -0.29 | -0.05 | 0 | 6.19 | 1656.07 | outside |
| 477 | 489 | STPCNGVEGFNC | 0.2147 | 12 | ABCPRED | 1 | NON-ALLERGEN | Non-Toxin | -0.07 | -0.2 | -0.23 | -1 | 4 | 1227.49 | outside |
| 108 | 120 | TTLDSKTQSLLI | 0.8325 | 12 | ABCPRED | 1 | NON-ALLERGEN | Non-Toxin | -0.1 | 0.11 | -0.13 | 0 | 6.19 | 1319.7 | outside |
| 133 | 145 | FQFCNDPFLGVY | 0.2495 | 12 | ABCPRED | 1 | NON-ALLERGEN | Non-Toxin | 0.08 | 0.43 | -0.89 | -1 | 3.8 | 1449.81 | outside |
| 369 | 385 | YNSASFSTFKCYGVSP | 0.7348 | 16 | ABCPRED | 1 | NON-ALLERGEN | Non-Toxin | -0.05 | -0.11 | -0.54 | 1 | 8.51 | 1758.14 | outside |
| 697 | 713 | MSLGAENSVAYSNNSI | 0.7951 | 16 | ABCPRED | 1 | NON-ALLERGEN | Non-Toxin | -0.05 | -0.06 | -0.31 | -1 | 4 | 1657.01 | outside |
| 474 | 508 | AGSTPCNGVEGFNCYFPLQSYGFQPTNGVGYC | 0.5024 | 34 | IEDB | 1 | ALLERGEN | Non-Toxin | -0.05 | -0.48 | -0.51 | -1 | 4 | 3629.46 | outside |
| 568 | 580 | DIADTTDAVRDP | 0.6079 | 12 | FBCPred | 1 | NON-ALLERGEN | Non-Toxin | -0.27 | -0.77 | 0.83 | -3 | 3.77 | 1288.48 | outside |
| 940 | 958 | STASALGKLQDVVNQNAQ | 0.557 | 18 | ABCPRED | 1 | ALLERGEN | Non-Toxin | -0.16 | -0.34 | -0.05 | 0 | 6.19 | 1844.28 | outside |
| 296 | 306 | LSETKCTLKS | 0.9434 | 10 | ABCPRED | 1 | NON-ALLERGEN | Non-Toxin | -0.26 | -0.42 | 0.42 | 1 | 8.54 | 1109.44 | outside |
| 1235 | 1249 | CCMTSCCSCLKGCC | -0.3983 | 14 | FBCPred | 1 | ALLERGEN | Non-Toxin | -0.04 | 1.19 | -0.49 | 1 | 7.99 | 1445 | transmembrane-region_inside |
| 375 | 385 | STFKCYGVSP | 1.104 | 10 | ABCPRED | 1 | ALLERGEN | Non-Toxin | -0.05 | 0 | -0.41 | 1 | 8.54 | 1088.37 | outside |
| 800 | 812 | FNFSQILPDPSK | 0.3682 | 12 | ABCPRED | 1 | NON-ALLERGEN | Non-Toxin | -0.11 | -0.44 | -0.13 | 0 | 6.19 | 1392.74 | outside |
| 1187 | 1195 | NEVAKNLN | -0.0646 | 8 | IEDB | 1 | ALLERGEN | Non-Toxin | -0.29 | -1.01 | 0.35 | 0 | 6.35 | 901.1 | outside |
| 108 | 118 | TTLDSKTQSL | 0.99 | 10 | BCEPRED | 1 | NON-ALLERGEN | Non-Toxin | -0.25 | -0.7 | 0.2 | 0 | 6.19 | 1093.34 | outside |
| 932 | 940 | GKIQDSLS | 0.2746 | 8 | BCEPRED | 1 | ALLERGEN | Non-Toxin | -0.2 | -0.57 | 0.4 | 0 | 6.19 | 847.05 | outside |
| 534 | 543 | VKNKCVNFN | 2.053 | 9 | literature | 1 | ALLERGEN | Non-Toxin | -0.27 | -0.51 | 0.01 | 2 | 9.36 | 1065.38 | outside |
| 524 | 542 | VCGPKKSTNLVKNKCVNF | 0.5286 | 18 | ABCPRED | 1 | NON-ALLERGEN | Non-Toxin | -0.21 | -0.3 | 0.09 | 4 | 9.72 | 1979.65 | outside |
| 476 | 494 | GSTPCNGVEGFNCYFPLQ | 0.4593 | 18 | FBCPred | 1 | NON-ALLERGEN | Non-Toxin | -0.02 | -0.14 | -0.51 | -1 | 4 | 1933.4 | outside |
| 1048 | 1064 | HLMSFPQSAPHGVVFL | 0.0469 | 16 | ABCPRED | 1 | NON-ALLERGEN | Non-Toxin | 0.12 | 0.64 | -0.85 | 1 | 7.26 | 1767.32 | outside |
| 741 | 751 | YICGDSTECS | -0.1828 | 10 | ABCPRED | 1 | ALLERGEN | Non-Toxin | -0.1 | -0.15 | 0.01 | -2 | 3.67 | 1077.27 | outside |
| 958 | 976 | ALNTLVKQLSSNFGAISS | 0.1815 | 18 | ABCPRED | 1 | NON-ALLERGEN | Non-Toxin | -0.01 | 0.43 | -0.43 | 1 | 9.11 | 1850.37 | outside |
| 179 | 186 | LEGKQGN | 1.8367 | 7 | IEDB | 1 | ALLERGEN | Non-Toxin | -0.31 | -1.63 | 0.66 | 0 | 6.35 | 744.91 | outside |
| 82 | 92 | PVLPFNDGVY | 0.3324 | 10 | ABCPRED | 1 | ALLERGEN | Toxin | 0.09 | 0.31 | -0.64 | -1 | 3.8 | 1120.41 | outside |
| 584 | 596 | ILDITPCSFGGV | 1.5192 | 12 | ABCPRED | 1 | ALLERGEN | Non-Toxin | 0.19 | 1.24 | -0.62 | -1 | 3.8 | 1221.61 | outside |
| 108 | 116 | TTLDSKTQ | 1.0912 | 8 | BCEPRED | 1 | NON-ALLERGEN | Non-Toxin | -0.35 | -1.25 | 0.44 | 0 | 6.19 | 893.07 | outside |
| 1081 | 1092 | ICHDGKAHFPR | -0.8542 | 11 | IEDB | 1 | NON-ALLERGEN | Non-Toxin | -0.24 | -0.79 | 0.2 | 2 | 8.57 | 1280.62 | outside |
| 691 | 709 | SIIAYTMSLGAENSVAYS | 0.6083 | 18 | ABCPRED | 1 | ALLERGEN | Non-Toxin | 0.07 | 0.58 | -0.57 | -1 | 4 | 1877.35 | outside |
| 867 | 879 | DEMIAQYTSALL | 0.0465 | 12 | ABCPRED | 1 | ALLERGEN | Non-Toxin | 0.01 | 0.36 | -0.32 | -2 | 3.67 | 1354.71 | outside |
| 370 | 382 | NSASFSTFKCYG | 0.2819 | 12 | ABCPRED | 1 | NON-ALLERGEN | Non-Toxin | -0.08 | -0.19 | -0.43 | 1 | 8.54 | 1311.58 | outside |
| 290 | 300 | DCALDPLSET | 0.5222 | 10 | ABCPRED | 1 | ALLERGEN | Non-Toxin | -0.12 | -0.17 | 0.38 | -3 | 3.5 | 1063.27 | outside |
| 604 | 622 | TSNQVAVLYQDVNCTEVP | 0.6941 | 18 | ABCPRED | 1 | NON-ALLERGEN | Non-Toxin | -0.09 | -0.07 | -0.29 | -2 | 3.67 | 1980.44 | outside |
| 600 | 620 | PGTNTSNQVAVLYQDVNCTE | 0.687 | 20 | ABCPRED | 1 | NON-ALLERGEN | Non-Toxin | -0.14 | -0.5 | -0.2 | -2 | 3.67 | 2153.6 | outside |
| 472 | 492 | IYQAGSTPCNGVEGFNCYFP | 0.1667 | 20 | BCPREDS | 1 | ALLERGEN | Non-Toxin | 0.01 | -0.07 | -0.59 | -1 | 4 | 2167.68 | outside |
| 816 | 821 | SFIED | short | 5 | literature | 1 | ALLERGEN | Non-Toxin | -0.05 | -0.1 | 0.4 | -2 | 3.67 | 609.69 | outside |
| 198 | 208 | DGYFKIYSKH | 0.4989 | 10 | ABCPRED | 1 | NON-ALLERGEN | Non-Toxin | -0.2 | -1.1 | -0.01 | 1.5 | 8.76 | 1257.55 | outside |
| 736 | 746 | VDCTMYICGD | -0.8689 | 10 | ABCPRED | 1 | NON-ALLERGEN | Non-Toxin | 0.02 | 0.62 | -0.33 | -2 | 3.57 | 1119.42 | outside |
| 252 | 268 | GDSSSGWTAGAAAYYV | 0.2821 | 16 | ABCPRED | 1 | NON-ALLERGEN | Non-Toxin | 0.05 | 0.01 | -0.5 | -1 | 3.8 | 1562.82 | outside |
| 19 | 29 | TTRTQLPPAY | 1.2001 | 10 | ABCPRED | 1 | ALLERGEN | Non-Toxin | -0.23 | -0.9 | -0.26 | 1 | 9.1 | 1147.43 | outside |
| 578 | 588 | DPQTLEILDI | 1.5757 | 10 | ABCPRED | 1 | ALLERGEN | Non-Toxin | -0.05 | 0.03 | 0.16 | -3 | 3.5 | 1156.45 | outside |
| 736 | 748 | VDCTMYICGDST | -0.4784 | 12 | ABCPRED,BCPREDS | 2 | ALLERGEN | Non-Toxin | -0.02 | 0.39 | -0.28 | -2 | 3.57 | 1307.63 | outside |
| 648 | 664 | GCLIGAEHVNNSYECD | 0.848 | 16 | ABCPRED,literature | 2 | NON-ALLERGEN | Non-Toxin | -0.09 | -0.27 | -0.04 | -2.5 | 4.14 | 1724.08 | outside |
| 714 | 732 | IPTNFTISVTTEILPVSM | 0.9956 | 18 | ABCPRED | 1 | NON-ALLERGEN | Non-Toxin | 0.11 | 0.88 | -0.66 | -1 | 4 | 1963.59 | outside |
| 114 | 134 | TQSLLIVNNATNVVIKVCEF | 0.204 | 20 | ABCPRED,BCPREDS | 2 | NON-ALLERGEN | Non-Toxin | 0.03 | 0.85 | -0.55 | 0 | 6.32 | 2205.91 | outside |
| 537 | 549 | KCVNFNFNGLTG | 1.6969 | 12 | IEDB,literature | 2 | NON-ALLERGEN | Non-Toxin | -0.05 | 0.02 | -0.51 | 1 | 8.57 | 1313.67 | outside |
| 647 | 665 | AGCLIGAEHVNNSYECDI | 0.7951 | 18 | ABCPRED | 1 | NON-ALLERGEN | Non-Toxin | -0.02 | 0.11 | -0.17 | -2.5 | 4.14 | 1908.35 | outside |
| 141 | 151 | LGVYYHKNNK | 0.9727 | 10 | ABCPRED | 1 | ALLERGEN | Non-Toxin | -0.26 | -1.3 | -0.2 | 2.5 | 9.55 | 1235.56 | outside |
| 478 | 488 | TPCNGVEGFN | 0.0446 | 10 | ABCPRED | 1 | ALLERGEN | Non-Toxin | -0.06 | -0.41 | -0.2 | -1 | 4 | 1037.25 | outside |
| 520 | 538 | APATVCGPKKSTNLVKNK | 0.2074 | 18 | ABCPRED,ellipro,literature | 3 | NON-ALLERGEN | Non-Toxin | -0.23 | -0.56 | 0.28 | 4 | 10.05 | 1856.47 | outside |
| 550 | 560 | GVLTESNKKF | 0.6528 | 10 | ABCPRED | 1 | NON-ALLERGEN | Non-Toxin | -0.21 | -0.59 | 0.33 | 1 | 8.94 | 1122.43 | outside |
| 761 | 771 | TQLNRALTGI | 0.2187 | 10 | ABCPRED | 1 | ALLERGEN | Non-Toxin | -0.12 | 0.06 | -0.33 | 1 | 10.11 | 1086.41 | outside |
| 299 | 309 | TKCTLKSFTV | 0.4314 | 10 | ABCPRED | 1 | NON-ALLERGEN | Non-Toxin | -0.13 | 0.26 | -0.17 | 2 | 9.36 | 1127.5 | outside |
| 164 | 176 | NNCTFEYVSQPF | 0.1629 | 12 | ABCPRED | 1 | NON-ALLERGEN | Non-Toxin | -0.11 | -0.51 | -0.53 | -1 | 4 | 1448.73 | outside |
| 730 | 742 | SMTKTSVDCTMY | 1.0622 | 12 | ABCPRED | 1 | NON-ALLERGEN | Non-Toxin | -0.15 | -0.16 | -0.17 | 0 | 6.15 | 1366.74 | outside |
| 335 | 347 | LCPFGEVFNATR | 0.1964 | 12 | ABCPRED | 1 | NON-ALLERGEN | Non-Toxin | -0.04 | 0.31 | -0.33 | 0 | 6.32 | 1353.72 | outside |
| 521 | 539 | PATVCGPKKSTNLVKNKC | 0.2012 | 18 | ABCPRED | 1 | NON-ALLERGEN | Non-Toxin | -0.24 | -0.52 | 0.26 | 4 | 9.72 | 1888.53 | outside |
| 694 | 704 | AYTMSLGAEN | 0.8326 | 10 | ABCPRED | 1 | ALLERGEN | Non-Toxin | -0.02 | -0.09 | -0.33 | -1 | 4 | 1056.29 | outside |
| 1216 | 1234 | IWLGFIAGLIAIVMVTIM | 0.6278 | 18 | ABCPRED | 1 | NON-ALLERGEN | Non-Toxin | 0.44 | 2.57 | -1.42 | 0 | 5.88 | 1961.84 | transmembrane-region |
| 1167 | 1185 | GDISGINASVVNIQKEID | 0.5521 | 18 | ABCPRED,BCPREDS | 2 | NON-ALLERGEN | Non-Toxin | -0.06 | 0.05 | 0.14 | -2 | 4.03 | 1872.34 | outside |
| 278 | 285 | KYNENGT | 0.9414 | 7 | IEDB,literature | 2 | ALLERGEN | Non-Toxin | -0.43 | -2.4 | 0.53 | 0 | 6.35 | 824.94 | outside |
| 614 | 634 | DVNCTEVPVAIHADQLTPTW | 0.7339 | 20 | ABCPRED | 1 | NON-ALLERGEN | Non-Toxin | -0.03 | 0.01 | -0.29 | -2.5 | 4.03 | 2209.75 | outside |
| 517 | 533 | LLHAPATVCGPKKSTN | -0.1057 | 16 | ABCPRED | 1 | ALLERGEN | Non-Toxin | -0.11 | -0.15 | -0.12 | 2.5 | 9.36 | 1637.16 | outside |
| 50 | 60 | STQDLFLPFF | 0.3011 | 10 | ABCPRED | 1 | NON-ALLERGEN | Non-Toxin | 0.1 | 0.59 | -0.8 | -1 | 3.8 | 1214.52 | outside |
| 781 | 789 | VFAQVKQI | 0.5854 | 8 | IEDB,literature | 2 | ALLERGEN | Non-Toxin | 0.02 | 0.82 | -0.55 | 1 | 9.11 | 932.25 | outside |
| 727 | 737 | LPVSMTKTSV | 1.2574 | 10 | ABCPRED | 1 | NON-ALLERGEN | Non-Toxin | -0.02 | 0.56 | -0.33 | 1 | 9.11 | 1062.43 | outside |
| 409 | 421 | QIAPGQTGKIAD | 1.4941 | 12 | literature | 1 | NON-ALLERGEN | Non-Toxin | -0.1 | -0.41 | 0.12 | 0 | 6.19 | 1198.52 | outside |
| 50 | 62 | STQDLFLPFFSN | 0.1041 | 12 | BCPREDS | 1 | NON-ALLERGEN | Non-Toxin | 0.01 | 0.13 | -0.62 | -1 | 3.8 | 1415.73 | outside |
| 279 | 293 | YNENGTITDAVDCA | 0.5541 | 14 | ABCPRED,FBCPred | 2 | NON-ALLERGEN | Non-Toxin | -0.12 | -0.41 | 0.07 | -3 | 3.5 | 1485.73 | outside |
| 49 | 59 | HSTQDLFLPF | 0.5162 | 10 | ABCPRED | 1 | NON-ALLERGEN | Non-Toxin | 0 | -0.01 | -0.6 | -0.5 | 5.09 | 1204.49 | outside |
| 700 | 720 | GAENSVAYSNNSIAIPTNFT | 0.4985 | 20 | BCPREDS,ellipro,literature | 3 | NON-ALLERGEN | Non-Toxin | -0.04 | -0.16 | -0.38 | -1 | 4 | 2070.48 | outside |
| 136 | 147 | CNDPFLGVYYH | 0.4109 | 11 | IEDB,literature | 2 | ALLERGEN | Non-Toxin | 0.01 | -0.14 | -0.79 | -0.5 | 5.09 | 1327.63 | outside |
| 681 | 691 | PRRARSVASQ | 0.33 | 10 | ABCPRED | 1 | NON-ALLERGEN | Non-Toxin | -0.55 | -1.24 | 0.73 | 3 | 12.31 | 1127.39 | outside |
| 411 | 421 | APGQTGKIAD | 1.2753 | 10 | ABCPRED | 1 | ALLERGEN | Non-Toxin | -0.12 | -0.59 | 0.3 | 0 | 6.19 | 957.19 | outside |

| | | | | | | | | | | | | | | | |
|---|---|---|---|---|---|---|---|---|---|---|---|---|---|---|---|
| 585 | 597 | LDITPCSFGGVS | 1.9283 | 12 | ABCPRED | 1 | NON-ALLERGEN | Non-Toxin | 0.11 | 0.8 | -0.45 | -1 | 3.8 | 1195.52 | outside |
| 924 | 936 | ANQFNSAIGKIQ | 0.1527 | 12 | ABCPRED | 1 | NON-ALLERGEN | Non-Toxin | -0.11 | -0.31 | -0.25 | 1 | 9.11 | 1290.62 | outside |
| 882 | 900 | ITSGWTFGAGAALQIPFA | 0.4686 | 18 | ABCPRED | 1 | NON-ALLERGEN | Non-Toxin | 0.2 | 0.9 | -0.89 | 0 | 5.88 | 1808.33 | outside |
| 731 | 741 | MTKTSVDCTM | 1.3958 | 10 | ABCPRED | 1 | NON-ALLERGEN | Non-Toxin | -0.15 | 0.02 | 0 | 0 | 6.16 | 1116.46 | outside |
| 109 | 125 | TLDSKTQSLLIVNNAT | 0.7116 | 16 | ABCPRED | 1 | ALLERGEN | Non-Toxin | -0.11 | 0.02 | -0.2 | 0 | 6.19 | 1718.18 | outside |
| 536 | 546 | NKCVNFNFNG | 1.3088 | 10 | ABCPRED | 1 | ALLERGEN | Non-Toxin | -0.17 | -0.6 | -0.37 | 1 | 8.57 | 1156.42 | outside |
| 961 | 977 | TLVKQLSSNFGAISSV | 0.388 | 16 | ABCPRED | 1 | NON-ALLERGEN | Non-Toxin | 0.02 | 0.62 | -0.45 | 1 | 9.11 | 1651.13 | outside |
| 335 | 345 | LCPFGEVFNA | -0.0095 | 10 | ABCPRED | 1 | NON-ALLERGEN | Non-Toxin | 0.14 | 0.89 | -0.66 | -1 | 4 | 1096.4 | outside |
| 729 | 739 | VSMTKTSVDC | 1.6231 | 10 | ABCPRED | 1 | NON-ALLERGEN | Non-Toxin | -0.13 | 0.24 | 0.05 | 0 | 6.16 | 1070.37 | outside |
| 426 | 436 | PDDFTGCVIA | -0.1092 | 10 | ABCPRED | 1 | NON-ALLERGEN | Non-Toxin | 0.06 | 0.61 | -0.17 | -2 | 3.57 | 1037.28 | outside |
| 250 | 260 | TPGDSSSGWT | 0.1741 | 10 | ABCPRED | 1 | ALLERGEN | Non-Toxin | -0.12 | -1.06 | -0.03 | -1 | 3.8 | 994.11 | outside |
| 433 | 443 | VIAWNSNNLD | 0.9177 | 10 | ABCPRED | 1 | NON-ALLERGEN | Non-Toxin | -0.05 | -0.14 | -0.51 | -1 | 3.8 | 1145.38 | outside |
| 676 | 690 | TQTNSPRRARSVAS | 0.1765 | 14 | FBCPred | 1 | NON-ALLERGEN | Non-Toxin | -0.48 | -1.29 | 0.5 | 3 | 12.31 | 1530.84 | outside |
| 935 | 953 | QDSLSSTASALGKLQDVV | 0.6096 | 18 | ABCPRED | 1 | NON-ALLERGEN | Non-Toxin | -0.1 | 0.07 | 0.04 | -1 | 4.21 | 1819.26 | outside |
| 615 | 633 | VNCTEVPVAIHADQLTPT | 0.5829 | 18 | AAP | 1 | ALLERGEN | Non-Toxin | -0.02 | 0.25 | -0.3 | -1.5 | 4.36 | 1908.42 | outside |
| 110 | 122 | LDSKTQSLLIVN | 0.9139 | 12 | ABCPRED | 1 | NON-ALLERGEN | Non-Toxin | -0.08 | 0.28 | -0.18 | 0 | 6.19 | 1330.73 | outside |
| 711 | 729 | SIAIPTNFTISVTTEILP | 0.7769 | 18 | ABCPRED | 1 | NON-ALLERGEN | Non-Toxin | 0.12 | 0.89 | -0.63 | -1 | 4 | 1917.5 | outside |
| 772 | 782 | VEQDKNTQEV | 0.2465 | 10 | ABCPRED,BCEPRED | 2 | NON-ALLERGEN | Non-Toxin | -0.42 | -1.72 | 0.92 | -2 | 4.14 | 1189.39 | outside |
| 530 | 548 | STNLVKNKCVNFNFNGLT | 0.9604 | 18 | ABCPRED | 1 | NON-ALLERGEN | Non-Toxin | -0.14 | -0.21 | -0.34 | 2 | 9.36 | 2013.57 | outside |
| 14 | 24 | QCVNLTTRTQ | 1.7824 | 10 | ABCPRED,literature | 2 | NON-ALLERGEN | Non-Toxin | -0.32 | -0.66 | -0.19 | 1 | 8.6 | 1163.46 | outside |
| 56 | 68 | LPFFSNVTWFHA | 0.5911 | 12 | ABCPRED | 1 | NON-ALLERGEN | Non-Toxin | 0.16 | 0.62 | -1.26 | 0.5 | 7.1 | 1465.84 | outside |
| 14 | 26 | QCVNLTTRTQLP | 1.2589 | 12 | ABCPRED | 1 | NON-ALLERGEN | Non-Toxin | -0.23 | -0.37 | -0.31 | 1 | 8.6 | 1373.77 | outside |
| 694 | 714 | AYTMSLGAENSVAYSNNSIA | 0.614 | 20 | ABCPRED,FBCPred | 2 | NON-ALLERGEN | Non-Toxin | -0.02 | 0.03 | -0.43 | -1 | 4 | 2063.5 | outside |
| 110 | 130 | LDSKTQSLLIVNNATNVVIK | 0.6342 | 20 | ABCPRED | 1 | NON-ALLERGEN | Non-Toxin | -0.07 | 0.32 | -0.22 | 1 | 8.94 | 2170.85 | outside |
| 686 | 704 | SVASQSIIAYTMSLGAEN | 0.6217 | 18 | ABCPRED | 1 | NON-ALLERGEN | Non-Toxin | 0.03 | 0.46 | -0.43 | -1 | 4 | 1842.31 | outside |
| 894 | 903 | LQIPFAMQM | 1.068 | 9 | literature | 1 | ALLERGEN | Non-Toxin | 0.13 | 0.9 | -0.98 | 0 | 5.88 | 1078.49 | outside |
| 613 | 633 | QDVNCTEVPVAIHADQLTPT | 0.7361 | 20 | ABCPRED | 1 | NON-ALLERGEN | Non-Toxin | -0.09 | -0.13 | -0.11 | -2.5 | 4.03 | 2151.67 | outside |
| 533 | 545 | LVKNKCVNFNFN | 1.6246 | 12 | FBCPred | 1 | NON-ALLERGEN | Non-Toxin | -0.16 | -0.13 | -0.33 | 2 | 9.36 | 1439.87 | outside |
| 313 | 321 | YQTSNFRV | 0.371 | 8 | IEDB | 1 | ALLERGEN | Non-Toxin | -0.3 | -0.91 | -0.38 | 1 | 9.1 | 1014.21 | outside |
| 649 | 665 | CLIGAEHVNNSYECDI | 1.0413 | 16 | ABCPRED | 1 | NON-ALLERGEN | Non-Toxin | -0.05 | 0.04 | -0.16 | -2.5 | 4.14 | 1780.19 | outside |
| 585 | 603 | LDITPCSFGGVSVITPGT | 1.2006 | 18 | ABCPRED | 1 | ALLERGEN | Non-Toxin | 0.13 | 0.83 | -0.53 | -1 | 3.8 | 1764.29 | outside |
| 936 | 952 | DSLSSTASALGKLQDV | 0.7391 | 16 | ABCPRED | 1 | ALLERGEN | Non-Toxin | -0.1 | 0.03 | 0.13 | -1 | 4.21 | 1591.96 | outside |
| 1 | 21 | MFVFLVLLPLVSSQCVNLTT | 0.8317 | 20 | BCEPRED | 1 | ALLERGEN | Non-Toxin | 0.2 | 1.71 | -1.1 | 0 | 5.85 | 2225.06 | outside |
| 940 | 952 | STASALGKLQDV | 0.8473 | 12 | ABCPRED | 1 | ALLERGEN | Non-Toxin | -0.08 | 0.15 | 0.02 | 0 | 6.19 | 1189.5 | outside |
| 946 | 958 | GKLQDVVNQNAQ | 0.524 | 12 | ABCPRED | 1 | NON-ALLERGEN | Non-Toxin | -0.26 | -0.94 | 0.14 | 0 | 6.19 | 1313.62 | outside |
| 214 | 221 | RDLPQGF | 1.1398 | 7 | IEDB | 1 | ALLERGEN | Non-Toxin | -0.28 | -0.99 | 0.27 | 0 | 6.19 | 832.02 | outside |
| 222 | 232 | ALEPLVDLPI | 0.0175 | 10 | ABCPRED | 1 | NON-ALLERGEN | Non-Toxin | 0.16 | 1.17 | -0.32 | -2 | 3.67 | 1079.45 | outside |
| 279 | 289 | YNENGTITDA | 0.602 | 10 | ABCPRED | 1 | ALLERGEN | Non-Toxin | -0.18 | -1.08 | 0.1 | -2 | 3.67 | 1097.24 | outside |
| 676 | 686 | TQTNSPRRAR | 0.0099 | 10 | ABCPRED | 1 | NON-ALLERGEN | Non-Toxin | -0.7 | -2.25 | 0.84 | 3 | 12.31 | 1186.42 | outside |
| 218 | 228 | QGFSALEPLV | 0.5402 | 10 | ABCPRED | 1 | ALLERGEN | Non-Toxin | 0.1 | 0.66 | -0.46 | -1 | 4 | 1060.36 | outside |
| 928 | 944 | NSAIGKIQDSLSSTAS | 0.4562 | 16 | ABCPRED | 1 | ALLERGEN | Non-Toxin | -0.12 | -0.19 | 0.07 | 0 | 6.19 | 1578.92 | outside |
| 858 | 870 | LTVLPPLLTDEM | 0.4043 | 12 | FBCPred | 1 | NON-ALLERGEN | Non-Toxin | 0.09 | 0.81 | -0.4 | -2 | 3.67 | 1341.81 | outside |
| 808 | 818 | DPSKPSKRSF | 0.8148 | 10 | BCEPRED,IEDB | 2 | ALLERGEN | Non-Toxin | -0.5 | -1.86 | 1.04 | 2 | 10.01 | 1148.4 | outside |
| 443 | 455 | SKVGGNYNYLYR | 0.2892 | 12 | ABCPRED,BCPREDS | 2 | NON-ALLERGEN | Non-Toxin | -0.25 | -1.07 | -0.29 | 2 | 9.55 | 1433.76 | outside |
| 523 | 536 | TVCGPKKSTNLVK | 0.0426 | 13 | IEDB | 1 | ALLERGEN | Non-Toxin | -0.22 | -0.36 | 0.22 | 3 | 9.81 | 1374.85 | outside |
| 521 | 537 | PATVCGPKKSTNLVKN | 0.0611 | 16 | ABCPRED | 1 | NON-ALLERGEN | Non-Toxin | -0.21 | -0.5 | 0.16 | 3 | 9.81 | 1657.19 | outside |
| 517 | 537 | LLHAPATVCGPKKSTNLVKN | -0.069 | 20 | ABCPRED | 1 | NON-ALLERGEN | Non-Toxin | -0.12 | -0.09 | -0.1 | 3.5 | 9.81 | 2091.8 | outside |
| 946 | 955 | GKLQDVVNQ | 0.3499 | 9 | BCEPRED | 1 | NON-ALLERGEN | Non-Toxin | -0.23 | -0.68 | 0.2 | 0 | 6.19 | 1000.26 | outside |
| 46 | 64 | SVLHSTQDLFLPFFSNVT | 0.3483 | 18 | ABCPRED | 1 | NON-ALLERGEN | Non-Toxin | 0.05 | 0.51 | -0.72 | -0.5 | 5.09 | 2052.58 | outside |
| 892 | 904 | AALQIPFAMQMA | 0.8846 | 12 | ellipro | 1 | NON-ALLERGEN | Non-Toxin | 0.16 | 1.13 | -0.86 | 0 | 5.88 | 1291.76 | outside |
| 888 | 904 | FGAGAALQIPFAMQMA | 0.7891 | 16 | ABCPRED | 1 | NON-ALLERGEN | Non-Toxin | 0.2 | 1.08 | -0.83 | 0 | 5.88 | 1624.18 | outside |
| 802 | 814 | FSQILPDPSKPS | -0.1744 | 12 | ABCPRED | 1 | NON-ALLERGEN | Non-Toxin | -0.14 | -0.58 | 0.08 | 0 | 6.19 | 1315.65 | outside |
| 715 | 727 | PTNFTISVTTEI | 1.0904 | 12 | BCPREDS | 1 | NON-ALLERGEN | Non-Toxin | 0.03 | 0.32 | -0.48 | -1 | 4 | 1322.65 | outside |
| 368 | 380 | LYNSASFSTFKC | 0.1109 | 12 | ABCPRED | 1 | ALLERGEN | Non-Toxin | -0.05 | 0.16 | -0.58 | 1 | 8.54 | 1367.69 | outside |
| 690 | 706 | QSIIAYTMSLGAENSV | 0.4478 | 16 | ABCPRED | 1 | NON-ALLERGEN | Non-Toxin | 0.04 | 0.45 | -0.48 | -1 | 4 | 1684.13 | outside |
| 145 | 154 | YHKNNKSWM | 0.3388 | 9 | IEDB | 1 | ALLERGEN | Non-Toxin | -0.39 | -2.12 | -0.09 | 2.5 | 9.72 | 1207.5 | outside |
| 946 | 962 | GKLQDVVNQNAQALNT | 0.4612 | 16 | ABCPRED | 1 | NON-ALLERGEN | Non-Toxin | -0.2 | -0.62 | -0.05 | 0 | 6.19 | 1713.13 | outside |
| 224 | 234 | EPLVDLPIGI | 0.7105 | 10 | ABCPRED | 1 | NON-ALLERGEN | Non-Toxin | 0.17 | 1.02 | -0.27 | -2 | 3.67 | 1065.43 | outside |
| 700 | 718 | GAENSVAYSNNSIAIPTN | 0.2798 | 18 | ABCPRED,AAP,BCPREDS,literatur | 4 | NON-ALLERGEN | Toxin | -0.07 | -0.29 | -0.26 | -1 | 4 | 1822.17 | outside |
| 56 | 74 | LPFFSNVTWFHAIHVSGT | 0.6306 | 18 | ABCPRED | 1 | NON-ALLERGEN | Non-Toxin | 0.14 | 0.62 | -1.06 | 1 | 7.26 | 2060.61 | outside |
| 535 | 555 | KNKCVNFNFNGLTGTGVLTE | 1.2763 | 20 | ABCPRED | 1 | NON-ALLERGEN | Non-Toxin | -0.1 | -0.23 | -0.2 | 1 | 8.54 | 2156.75 | outside |
| 297 | 307 | SETKCTLKSF | 0.6433 | 10 | ABCPRED | 1 | NON-ALLERGEN | Non-Toxin | -0.25 | -0.52 | 0.35 | 1 | 8.54 | 1143.45 | outside |
| 675 | 687 | QTQTNSPRRARS | -0.0122 | 12 | ABCPRED | 1 | NON-ALLERGEN | Non-Toxin | -0.67 | -2.23 | 0.74 | 3 | 12.31 | 1401.66 | outside |
| 687 | 705 | VASQSIIAYTMSLGAENS | 0.5471 | 18 | ABCPRED | 1 | ALLERGEN | Non-Toxin | 0.03 | 0.46 | -0.43 | -1 | 4 | 1842.31 | outside |
| 437 | 446 | NSNNLDSKV | 0.6019 | 9 | BCEPRED | 1 | ALLERGEN | Non-Toxin | -0.35 | -1.28 | 0.43 | 0 | 6.19 | 990.16 | outside |
| 1047 | 1063 | YHLMSFPQSAPHGVVF | 0.11 | 16 | ABCPRED | 1 | ALLERGEN | Non-Toxin | 0.09 | 0.32 | -0.88 | 1 | 7.25 | 1817.33 | outside |
| 851 | 863 | CAQKFNGLTVLP | -0.0043 | 12 | ABCPRED | 1 | ALLERGEN | Non-Toxin | 0 | 0.44 | -0.51 | 1 | 8.57 | 1290.72 | outside |
| 606 | 622 | NQVAVLYQDVNCTEVP | 0.7426 | 16 | ABCPRED | 1 | NON-ALLERGEN | Non-Toxin | -0.08 | 0.02 | -0.33 | -2 | 3.67 | 1792.23 | outside |
| 679 | 691 | NSPRRARSVASQ | 0.0552 | 12 | ABCPRED | 1 | ALLERGEN | Non-Toxin | -0.54 | -1.39 | 0.65 | 3 | 12.31 | 1328.6 | outside |
| 568 | 579 | DIADTTDAVRD | 0.8257 | 11 | BCEPRED | 1 | ALLERGEN | Non-Toxin | -0.29 | -0.69 | 0.9 | -3 | 3.77 | 1191.35 | outside |
| 131 | 141 | CEFQFCNDPF | -0.6637 | 10 | ABCPRED | 1 | ALLERGEN | Non-Toxin | -0.08 | -0.22 | -0.31 | -2 | 3.67 | 1249.5 | outside |
| 932 | 948 | GKIQDSLSSTASALGK | 0.59 | 16 | ABCPRED | 1 | NON-ALLERGEN | Non-Toxin | -0.14 | -0.24 | 0.22 | 1 | 8.94 | 1562.97 | outside |
| 519 | 537 | HAPATVCGPKKSTNLVKN | 0.009 | 18 | BCPREDS | 1 | NON-ALLERGEN | Non-Toxin | -0.19 | -0.52 | 0.09 | 3.5 | 9.81 | 1865.44 | outside |
| 945 | 961 | LGKLQDVVNQNAQALN | 0.5358 | 16 | ABCPRED | 1 | ALLERGEN | Non-Toxin | -0.15 | -0.34 | -0.14 | 0 | 6.19 | 1725.19 | outside |
| 475 | 493 | AGSTPCNGVEGFNCYFPL | 0.3341 | 18 | ABCPRED,AAP | 2 | NON-ALLERGEN | Non-Toxin | 0.04 | 0.15 | -0.54 | -1 | 4 | 1876.34 | outside |
| 479 | 491 | PCNGVEGFNCYF | 0.3733 | 12 | ABCPRED | 1 | NON-ALLERGEN | Non-Toxin | 0.02 | 0.05 | -0.62 | -1 | 4 | 1349.66 | outside |
| 107 | 127 | GTTLDSKTQSLLIVNNATNV | 0.6906 | 20 | ABCPRED,AAP | 2 | NON-ALLERGEN | Non-Toxin | -0.09 | -0.01 | -0.24 | 0 | 6.19 | 2089.64 | outside |

| | | | | | | | | | | | | | | | |
|---|---|---|---|---|---|---|---|---|---|---|---|---|---|---|---|
| 438 | 448 | SNNLDSKVGG | 0.7715 | 10 | ABCPRED | 1 | NON-ALLERGEN | Non-Toxin | -0.22 | -0.88 | 0.37 | 0 | 6.19 | 990.18 | outside |
| 427 | 445 | DDFTGCVIAWNSNNLDSK | 0.3503 | 18 | ABCPRED,FBCPred | 2 | NON-ALLERGEN | Non-Toxin | -0.15 | -0.49 | 0.02 | -2 | 3.94 | 1999.39 | outside |
| 470 | 482 | TEIYQAGSTPCN | 0.0966 | 12 | ABCPRED | 1 | ALLERGEN | Non-Toxin | -0.12 | -0.6 | -0.22 | -1 | 4 | 1283.54 | outside |
| 116 | 134 | SLLIVNNATNVVIKVCEF | 0.2114 | 18 | ABCPRED | 1 | NON-ALLERGEN | Non-Toxin | 0.08 | 1.17 | -0.59 | 0 | 6.32 | 1976.64 | outside |
| 928 | 946 | NSAIGKIQDSLSSTASAL | 0.4499 | 18 | ABCPRED | 1 | ALLERGEN | Non-Toxin | -0.07 | 0.14 | -0.07 | 0 | 6.19 | 1763.19 | outside |
| 223 | 231 | LEPLVDLP | -0.3271 | 8 | IEDB,literature | 2 | NON-ALLERGEN | Non-Toxin | 0.08 | 0.67 | -0.11 | -2 | 3.67 | 895.18 | outside |
| 519 | 539 | HAPATVCGPKKSTNLVKNKC | 0.1405 | 20 | ABCPRED,BCPREDS | 2 | NON-ALLERGEN | Non-Toxin | -0.22 | -0.54 | 0.18 | 4.5 | 9.72 | 2096.78 | outside |
| 529 | 547 | KSTNLVKNKCVNFNFNGL | 1.22 | 18 | ABCPRED | 1 | NON-ALLERGEN | Non-Toxin | -0.19 | -0.39 | -0.15 | 3 | 9.81 | 2040.64 | outside |
| 675 | 691 | QTQTNSPRRARSVASQ | 0.1434 | 16 | ABCPRED | 1 | NON-ALLERGEN | Non-Toxin | -0.51 | -1.57 | 0.46 | 3 | 12.31 | 1787.14 | outside |
| 597 | 615 | VITPGTNTSNQVAVLYQD | 0.3865 | 18 | ABCPRED,FBCPred | 2 | ALLERGEN | Toxin | -0.05 | -0.06 | -0.44 | -1 | 3.8 | 1920.38 | outside |
| 523 | 539 | TVCGPKKSTNLVKNKC | 0.1722 | 16 | ABCPRED | 1 | NON-ALLERGEN | Non-Toxin | -0.28 | -0.6 | 0.32 | 4 | 9.72 | 1720.31 | outside |
| 484 | 502 | EGFNCYFPLQSYGFQPTN | 0.8567 | 18 | ABCPRED | 1 | NON-ALLERGEN | Non-Toxin | -0.06 | -0.61 | -0.62 | -1 | 4 | 2112.56 | outside |
| 337 | 347 | PFGEVFNATR | -0.0048 | 10 | ABCPRED | 1 | NON-ALLERGEN | Non-Toxin | -0.11 | -0.26 | -0.12 | 0 | 6.36 | 1137.39 | outside |
| 868 | 886 | EMIAQYTSALLAGTITSG | 0.3295 | 18 | ABCPRED | 1 | NON-ALLERGEN | Non-Toxin | 0.08 | 0.62 | -0.54 | -1 | 4 | 1827.35 | outside |
| 279 | 288 | YNENGTITD | 0.765 | 9 | IEDB | 1 | ALLERGEN | Non-Toxin | -0.23 | -1.4 | 0.17 | -2 | 3.67 | 1026.15 | outside |
| 52 | 64 | QDLFLPFFSNVT | 0.2381 | 12 | ABCPRED | 1 | NON-ALLERGEN | Non-Toxin | 0.07 | 0.55 | -0.78 | -1 | 3.8 | 1427.79 | outside |
| 529 | 545 | KSTNLVKNKCVNFNFN | 1.4412 | 16 | ABCPRED | 1 | NON-ALLERGEN | Non-Toxin | -0.25 | -0.65 | -0.06 | 3 | 9.81 | 1870.39 | outside |
| 332 | 343 | ITNLCPFGEVF | 0.3094 | 11 | BCEPRED | 1 | NON-ALLERGEN | Non-Toxin | 0.16 | 0.99 | -0.75 | -1 | 4 | 1239.61 | outside |
| 474 | 492 | QAGSTPCNGVEGFNCYFP | 0.3595 | 18 | ABCPRED | 1 | NON-ALLERGEN | Non-Toxin | -0.03 | -0.26 | -0.43 | -1 | 4 | 1891.31 | outside |
| 946 | 964 | GKLQDVVNQNAQALNTLV | 0.2926 | 18 | ABCPRED | 1 | NON-ALLERGEN | Non-Toxin | -0.12 | -0.11 | -0.23 | 0 | 6.19 | 1925.46 | outside |
| 107 | 119 | GTTLDSKTQSLL | 0.7358 | 12 | BCPREDS | 1 | NON-ALLERGEN | Non-Toxin | -0.15 | -0.3 | 0.02 | 0 | 6.19 | 1263.59 | outside |
| 110 | 116 | LDSKTQ | 1.3071 | 6 | IEDB,literature | 2 | ALLERGEN | Non-Toxin | -0.4 | -1.43 | 0.72 | 0 | 6.19 | 690.83 | outside |
| 443 | 453 | SKVGGNYNYL | 0.6741 | 10 | ABCPRED | 1 | NON-ALLERGEN | Non-Toxin | -0.12 | -0.71 | -0.42 | 1 | 8.83 | 1114.37 | outside |
| 772 | 781 | VEQDKNTQE | 0.0684 | 9 | IEDB | 1 | ALLERGEN | Non-Toxin | -0.52 | -2.38 | 1.19 | -2 | 4.14 | 1090.24 | outside |
| 1169 | 1185 | ISGINASVVNIQKEID | 0.6591 | 16 | ABCPRED | 1 | NON-ALLERGEN | Non-Toxin | -0.03 | 0.3 | -0.03 | -1 | 4.38 | 1700.17 | outside |
| 567 | 577 | RDIADTTDAV | 0.6138 | 10 | ABCPRED | 1 | NON-ALLERGEN | Non-Toxin | -0.25 | -0.41 | 0.69 | -2 | 3.94 | 1076.25 | outside |
| 526 | 542 | GPKKSTNLVKNKCVNF | 0.8596 | 16 | ABCPRED | 1 | NON-ALLERGEN | Non-Toxin | -0.28 | -0.76 | 0.26 | 4 | 10.05 | 1777.35 | outside |
| 484 | 500 | EGFNCYFPLQSYGFQP | 0.687 | 16 | ABCPRED | 1 | NON-ALLERGEN | Non-Toxin | -0.02 | -0.42 | -0.69 | -1 | 4 | 1897.32 | outside |
| 763 | 773 | LNRALTGIAV | 0.2645 | 10 | ABCPRED | 1 | ALLERGEN | Non-Toxin | 0.04 | 1.08 | -0.51 | 1 | 10.11 | 1027.38 | outside |
| 482 | 500 | GVEGFNCYFPLQSYGFQP | 0.7751 | 18 | ABCPRED | 1 | NON-ALLERGEN | Non-Toxin | 0.02 | -0.16 | -0.69 | -1 | 4 | 2053.54 | outside |
| 354 | 364 | NRKRISNCVA | 0.2816 | 10 | ABCPRED | 1 | ALLERGEN | Non-Toxin | -0.46 | -0.77 | 0.49 | 3 | 10.87 | 1160.49 | outside |
| 936 | 954 | DSLSSTASALGKLQDVVN | 0.5678 | 18 | ABCPRED | 1 | NON-ALLERGEN | Non-Toxin | -0.1 | 0.07 | 0.04 | -1 | 4.21 | 1805.23 | outside |
| 695 | 711 | YTMSLGAENSVAYSNN | 0.6434 | 16 | ABCPRED,IEDB,literature | 3 | NON-ALLERGEN | Non-Toxin | -0.09 | -0.41 | -0.38 | -1 | 4 | 1721.05 | outside |
| 945 | 957 | LGKLQDVVNQNA | 0.3607 | 12 | ABCPRED | 1 | NON-ALLERGEN | Non-Toxin | -0.16 | -0.33 | -0.02 | 0 | 6.19 | 1298.65 | outside |
| 435 | 445 | AWNSNNLDSK | 1.0198 | 10 | BCEPRED | 1 | ALLERGEN | Non-Toxin | -0.31 | -1.48 | 0.15 | 0 | 6.19 | 1148.33 | outside |
| 1212 | 1230 | WPWYIWLGFIAGLIAIVM | 0.7583 | 18 | ABCPRED | 1 | NON-ALLERGEN | Toxin | 0.4 | 1.76 | -1.64 | 0 | 5.88 | 2149.96 | outside_transmembrane-region |
| 205 | 215 | SKHTPINLVR | 0.8329 | 10 | ABCPRED | 1 | NON-ALLERGEN | Non-Toxin | -0.26 | -0.57 | 0.05 | 2.5 | 11.01 | 1164.52 | outside |
| 428 | 444 | DFTGCVIAWNSNNLDS | 0.2716 | 16 | ABCPRED | 1 | NON-ALLERGEN | Non-Toxin | -0.05 | -0.09 | -0.36 | -2 | 3.57 | 1756.1 | outside |
| 890 | 902 | AGAALQIPFAMQ | 0.8023 | 12 | ABCPRED | 1 | NON-ALLERGEN | Non-Toxin | 0.15 | 0.93 | -0.75 | 0 | 5.88 | 1217.62 | outside |
| 890 | 906 | AGAALQIPFAMQMAYR | 0.9036 | 16 | ABCPRED | 1 | ALLERGEN | Non-Toxin | 0.04 | 0.57 | -0.63 | 1 | 9.1 | 1739.31 | outside |
| 887 | 903 | TFGAGAALQIPFAMQM | 0.678 | 16 | ABCPRED | 1 | NON-ALLERGEN | Non-Toxin | 0.17 | 0.92 | -0.83 | 0 | 5.88 | 1654.21 | outside |
| 692 | 704 | IIAYTMSLGAEN | 0.7766 | 12 | ABCPRED | 1 | NON-ALLERGEN | Non-Toxin | 0.1 | 0.67 | -0.57 | -1 | 4 | 1282.65 | outside |
| 741 | 759 | YICGDSTECSNLLLQYGS | 0.13 | 18 | ABCPRED | 1 | NON-ALLERGEN | Non-Toxin | -0.05 | 0.02 | -0.38 | -2 | 3.67 | 1966.43 | outside |
| 556 | 566 | NKKFLPFQQF | 0.6977 | 10 | ABCPRED | 1 | ALLERGEN | Non-Toxin | -0.19 | -0.77 | -0.27 | 2 | 10.02 | 1296.68 | outside |
| 107 | 123 | GTTLDSKTQSLLIVNN | 0.755 | 16 | ABCPRED | 1 | ALLERGEN | Non-Toxin | -0.11 | -0.12 | -0.17 | 0 | 6.19 | 1704.16 | outside |
| 525 | 537 | CGPKKSTNLVKN | -0.092 | 12 | ABCPRED | 1 | ALLERGEN | Non-Toxin | -0.32 | -0.97 | 0.42 | 3 | 9.81 | 1288.7 | outside |
| 1058 | 1074 | HGVVFLHVTYVPAQEK | 0.8847 | 16 | ABCPRED,literature | 2 | ALLERGEN | Non-Toxin | 0.02 | 0.24 | -0.52 | 1 | 7.25 | 1824.36 | outside |
| 439 | 447 | NNLDSKVG | 1.2952 | 8 | BCEPRED | 1 | ALLERGEN | Non-Toxin | -0.27 | -0.95 | 0.43 | 0 | 6.19 | 846.02 | outside |
| 662 | 678 | CDIPIGAGICASYQTQ | 0.8764 | 16 | ABCPRED | 1 | NON-ALLERGEN | Non-Toxin | 0.03 | 0.4 | -0.46 | -1 | 3.8 | 1640.09 | outside |
| 291 | 301 | CALDPLSETK | 0.9226 | 10 | ABCPRED | 1 | ALLERGEN | Non-Toxin | -0.16 | -0.21 | 0.38 | -1 | 4.38 | 1076.36 | outside |
| 518 | 534 | LHAPATVCGPKKSTNL | 0.0624 | 16 | ABCPRED | 1 | ALLERGEN | Non-Toxin | -0.11 | -0.15 | -0.12 | 2.5 | 9.36 | 1637.16 | outside |
| 771 | 781 | AVEQDKNTQE | 0.2792 | 10 | ABCPRED | 1 | ALLERGEN | Non-Toxin | -0.45 | -1.96 | 1.02 | -2 | 4.14 | 1161.33 | outside |
| 939 | 957 | SSTASALGKLQDVVNQNA | 0.4799 | 18 | ABCPRED | 1 | ALLERGEN | Non-Toxin | -0.13 | -0.19 | -0.04 | 0 | 6.19 | 1803.22 | outside |
| 611 | 621 | LYQDVNCTEV | 0.8859 | 10 | ABCPRED | 1 | ALLERGEN | Non-Toxin | -0.12 | -0.13 | -0.21 | -2 | 3.67 | 1183.44 | outside |
| 694 | 706 | AYTMSLGAENSV | 0.5192 | 12 | AAP | 1 | NON-ALLERGEN | Non-Toxin | 0 | 0.21 | -0.38 | -1 | 4 | 1242.53 | outside |
| 769 | 781 | GIAVEQDKNTQE | 0.6365 | 12 | ABCPRED | 1 | NON-ALLERGEN | Non-Toxin | -0.3 | -1.29 | 0.7 | -2 | 4.14 | 1331.58 | outside |
| 186 | 191 | FKNLR | short | 5 | literature | 1 | ALLERGEN | Non-Toxin | -0.47 | -1.06 | 0.38 | 2 | 11.01 | 676.88 | outside |
| 743 | 761 | CGDSTECSNLLLQYGSFC | 0.6024 | 18 | ABCPRED | 1 | NON-ALLERGEN | Non-Toxin | -0.05 | 0.14 | -0.35 | -2 | 3.67 | 1940.4 | outside |
| 427 | 441 | DDFTGCVIAWNSNN | -0.0284 | 14 | FBCPred | 1 | ALLERGEN | Non-Toxin | -0.08 | -0.32 | -0.3 | -2 | 3.57 | 1555.83 | outside |
| 366 | 382 | SVLYNSASFSTFKCYG | 0.1864 | 16 | ABCPRED | 1 | ALLERGEN | Non-Toxin | -0.01 | 0.23 | -0.65 | 1 | 8.51 | 1774.19 | outside |
| 1050 | 1068 | MSFPQSAPHGVVFLHVTY | 0.5609 | 18 | AAP | 1 | ALLERGEN | Non-Toxin | 0.1 | 0.48 | -0.89 | 1 | 7.25 | 2017.6 | outside |
| 879 | 899 | AGTITSGWTFGAGAALQIPF | 0.3757 | 20 | ABCPRED | 1 | ALLERGEN | Non-Toxin | 0.18 | 0.75 | -0.83 | 0 | 5.88 | 1966.52 | outside |
| 310 | 318 | KGIYQTSN | 0.2441 | 8 | literature | 1 | ALLERGEN | Non-Toxin | -0.24 | -1.2 | -0.1 | 1 | 8.94 | 910.11 | outside |
| 1191 | 1197 | KNLNES | 0.863 | 6 | IEDB | 1 | ALLERGEN | Non-Toxin | -0.46 | -1.9 | 0.82 | 0 | 6.35 | 703.83 | outside |
| 1231 | 1249 | TIMLCCMTSCCSCLKGCC | -0.2868 | 18 | ABCPRED | 1 | NON-ALLERGEN | Toxin | 0.04 | 1.45 | -0.68 | 1 | 7.99 | 1903.69 | transmembrane-region_inside |
| 331 | 347 | NITNLCPFGEVFNATR | 0.652 | 16 | ABCPRED | 1 | NON-ALLERGEN | Non-Toxin | -0.08 | 0.03 | -0.36 | 0 | 6.32 | 1796.26 | outside |
| 54 | 66 | LFLPFFSNVTWF | 0.2841 | 12 | ABCPRED | 1 | NON-ALLERGEN | Non-Toxin | 0.27 | 1.29 | -1.53 | 0 | 5.88 | 1517.96 | outside |
| 675 | 692 | QTQTNSPRRARSVASQS | 0.1698 | 17 | IEDB | 1 | NON-ALLERGEN | Non-Toxin | -0.5 | -1.52 | 0.45 | 3 | 12.31 | 1874.23 | outside |
| 475 | 491 | AGSTPCNGVEGFNCYF | 0.2658 | 16 | ABCPRED | 1 | NON-ALLERGEN | Non-Toxin | 0.01 | 0.03 | -0.5 | -1 | 4 | 1666.03 | outside |
| 688 | 708 | ASQSIIAYTMSLGAENSVAY | 0.4441 | 20 | ABCPRED | 1 | ALLERGEN | Non-Toxin | 0.04 | 0.44 | -0.53 | -1 | 4 | 2076.59 | outside |
| 535 | 547 | KNKCVNFNFNGL | 1.7395 | 12 | ABCPRED | 1 | NON-ALLERGEN | Non-Toxin | -0.19 | -0.51 | -0.21 | 2 | 9.36 | 1397.79 | outside |
| 675 | 693 | QTQTNSPRRARSVASQSI | 0.1602 | 18 | ABCPRED | 1 | NON-ALLERGEN | Non-Toxin | -0.43 | -1.19 | 0.33 | 3 | 12.31 | 1987.41 | outside |
| 676 | 694 | TQTNSPRRARSVASQSII | 0.0657 | 18 | ABCPRED | 1 | NON-ALLERGEN | Non-Toxin | -0.35 | -0.74 | 0.22 | 3 | 12.31 | 1972.44 | outside |
| 469 | 487 | STEIYQAGSTPCNGVEGF | -0.0535 | 18 | ABCPRED | 1 | NON-ALLERGEN | Non-Toxin | -0.05 | -0.29 | -0.19 | -2 | 3.8 | 1860.24 | outside |
| 1216 | 1236 | IWLGFIAGLIAIVMVTIMLC | 0.4687 | 20 | ABCPRED | 1 | NON-ALLERGEN | Non-Toxin | 0.42 | 2.63 | -1.42 | 0 | 5.85 | 2178.17 | transmembrane-region |
| 279 | 291 | YNENGTITDAVD | 0.4295 | 12 | ABCPRED,FBCPred | 2 | NON-ALLERGEN | Non-Toxin | -0.17 | -0.84 | 0.21 | -3 | 3.5 | 1311.49 | outside |
| 239 | 249 | QTLLALHRSY | 0.5596 | 10 | ABCPRED,IEDB,literature | 3 | ALLERGEN | Non-Toxin | -0.14 | -0.08 | -0.56 | 1.5 | 9.1 | 1201.54 | outside |

| | | | | | | | | | | | | | | | |
|---|---|---|---|---|---|---|---|---|---|---|---|---|---|---|---|
| 110 | 126 | LDSKTQSLLIVNNATN | 0.6519 | 16 | ABCPRED | 1 | ALLERGEN | Non-Toxin | -0.14 | -0.16 | -0.16 | 0 | 6.19 | 1731.18 | outside |
| 467 | 483 | DISTEIYQAGSTPCNG | -0.1091 | 16 | ABCPRED | 1 | NON-ALLERGEN | Non-Toxin | -0.1 | -0.46 | -0.07 | -2 | 3.67 | 1655.98 | outside |
| 768 | 782 | TGIAVEQDKNTQEV | 0.741 | 14 | ABCPRED,AAP | 2 | NON-ALLERGEN | Non-Toxin | -0.23 | -0.86 | 0.46 | -2 | 4.14 | 1531.85 | outside |
| 730 | 740 | SMTKTSVDCT | 1.4047 | 10 | ABCPRED | 1 | NON-ALLERGEN | Non-Toxin | -0.2 | -0.25 | 0.16 | 0 | 6.16 | 1072.34 | outside |
| 535 | 545 | KNKCVNFNFN | 2.2331 | 10 | ABCPRED | 1 | ALLERGEN | Non-Toxin | -0.3 | -0.95 | -0.07 | 2 | 9.36 | 1227.54 | outside |
| 511 | 527 | VVLSFELLHAPATVCG | 0.6184 | 16 | ABCPRED | 1 | NON-ALLERGEN | Non-Toxin | 0.19 | 1.42 | -0.75 | -0.5 | 5.25 | 1656.21 | outside |
| 15 | 24 | CVNLTTRTQ | 1.8792 | 9 | literature | 1 | NON-ALLERGEN | Non-Toxin | -0.28 | -0.34 | -0.23 | 1 | 8.6 | 1035.31 | outside |
| 217 | 227 | PQGFSALEPL | 0.8327 | 10 | ABCPRED | 1 | ALLERGEN | Non-Toxin | 0.04 | 0.08 | -0.31 | -1 | 4 | 1058.34 | outside |
| 523 | 535 | TVCGPKKSTNLV | 0.2615 | 12 | ABCPRED,AAP | 2 | NON-ALLERGEN | Non-Toxin | -0.14 | -0.07 | -0.01 | 2 | 9.36 | 1246.66 | outside |
| 160 | 176 | YSSANNCTFEYVSQPF | 0.1223 | 16 | ABCPRED | 1 | ALLERGEN | Non-Toxin | -0.1 | -0.45 | -0.53 | -1 | 4 | 1857.19 | outside |
| 356 | 361 | KRISN | short | 5 | literature | 1 | ALLERGEN | Non-Toxin | -0.61 | -1.64 | 0.94 | 2 | 11.01 | 616.78 | outside |
| 472 | 490 | IYQAGSTPCNGVEGFNCY | 0.1822 | 18 | ABCPRED,BCPREDS | 2 | NON-ALLERGEN | Non-Toxin | -0.02 | -0.14 | -0.52 | -1 | 4 | 1923.36 | outside |
| 1236 | 1252 | CMTSCCSCLKGCCSCG | -0.1747 | 16 | ABCPRED | 1 | ALLERGEN | Non-Toxin | -0.04 | 0.96 | -0.41 | 1 | 7.99 | 1589.16 | transmembrane-region_inside |
| 880 | 900 | GTITSGWTFGAGAALQIPFA | 0.4039 | 20 | ABCPRED | 1 | ALLERGEN | Non-Toxin | 0.18 | 0.76 | -0.83 | 0 | 5.88 | 1966.52 | outside |
| 678 | 690 | TNSPRRARSVAS | 0.2168 | 12 | ABCPRED | 1 | ALLERGEN | Non-Toxin | -0.49 | -1.16 | 0.6 | 3 | 12.31 | 1301.57 | outside |
| 610 | 622 | VLYQDVNCTEVP | 0.739 | 12 | ABCPRED | 1 | ALLERGEN | Non-Toxin | -0.06 | 0.11 | -0.3 | -2 | 3.67 | 1379.72 | outside |
| 579 | 591 | PQTLEILDITPC | 1.4158 | 12 | ABCPRED | 1 | NON-ALLERGEN | Non-Toxin | 0 | 0.33 | -0.23 | -2 | 3.67 | 1342.75 | outside |
| 732 | 742 | TKTSVDCTMY | 1.2485 | 10 | ABCPRED | 1 | ALLERGEN | Non-Toxin | -0.18 | -0.3 | -0.1 | 0 | 6.15 | 1148.44 | outside |
| 223 | 234 | LEPLVDLPIGI | 0.8222 | 11 | BCEPRED | 1 | NON-ALLERGEN | Non-Toxin | 0.21 | 1.27 | -0.41 | -2 | 3.67 | 1178.61 | outside |
| 938 | 956 | LSSTASALGKLQDVVNQN | 0.6299 | 18 | ABCPRED | 1 | NON-ALLERGEN | Non-Toxin | -0.12 | -0.08 | -0.12 | 0 | 6.19 | 1845.31 | outside |
| 868 | 884 | EMIAQYTSALLAGTIT | 0.4102 | 16 | ABCPRED | 1 | NON-ALLERGEN | Non-Toxin | 0.1 | 0.77 | -0.62 | -1 | 4 | 1683.19 | outside |
| 736 | 745 | VDCTMYICG | -0.7197 | 9 | BCEPRED | 1 | ALLERGEN | Non-Toxin | 0.1 | 1.08 | -0.7 | -1 | 3.8 | 1004.32 | outside |
| 770 | 780 | IAVEQDKNTQ | 0.5274 | 10 | ABCPRED | 1 | NON-ALLERGEN | Non-Toxin | -0.31 | -1.16 | 0.54 | -1 | 4.38 | 1145.38 | outside |
| 946 | 957 | GKLQDVVNQNA | 0.3966 | 11 | IEDB | 1 | NON-ALLERGEN | Non-Toxin | -0.22 | -0.71 | 0.14 | 0 | 6.19 | 1185.47 | outside |
| 739 | 749 | TMYICGDSTE | -0.5262 | 10 | ABCPRED | 1 | ALLERGEN | Toxin | -0.07 | -0.2 | -0.09 | -2 | 3.67 | 1119.36 | outside |
| 525 | 538 | CGPKKSTNLVKNK | 0.1411 | 13 | BCEPRED | 1 | NON-ALLERGEN | Non-Toxin | -0.38 | -1.2 | 0.62 | 4 | 10.05 | 1416.89 | outside |
| 677 | 695 | QTNSPRRARSVASQSIIA | 0.0681 | 18 | ABCPRED | 1 | NON-ALLERGEN | Non-Toxin | -0.32 | -0.61 | 0.21 | 3 | 12.31 | 1942.41 | outside |
| 583 | 593 | EILDITPCSF | 1.2698 | 10 | ABCPRED | 1 | ALLERGEN | Non-Toxin | 0.08 | 0.8 | -0.3 | -2 | 3.67 | 1137.45 | outside |
| 1105 | 1112 | TQRNFYE | 0.219 | 7 | IEDB | 1 | ALLERGEN | Non-Toxin | -0.47 | -2.03 | 0.17 | 0 | 6.35 | 957.1 | outside |
| 10 | 26 | LVSSQCVNLTTRTQLP | 0.999 | 16 | ABCPRED | 1 | NON-ALLERGEN | Non-Toxin | -0.14 | 0.13 | -0.4 | 1 | 8.6 | 1760.28 | outside |
| 198 | 210 | DGYFKIYSKHTP | 0.4818 | 12 | ABCPRED | 1 | ALLERGEN | Non-Toxin | -0.19 | -1.11 | -0.04 | 1.5 | 8.76 | 1455.8 | outside |
| 680 | 692 | SPRRARSVASQS | 0.4331 | 12 | ABCPRED | 1 | NON-ALLERGEN | Non-Toxin | -0.5 | -1.17 | 0.66 | 3 | 12.31 | 1301.57 | outside |
| 252 | 272 | GDSSSGWTAGAAAYYVGYLQ | 0.3899 | 20 | ABCPRED | 1 | NON-ALLERGEN | Non-Toxin | 0.04 | -0.07 | -0.6 | -1 | 3.8 | 2024.41 | outside |
| 81 | 93 | NPVLPFNDGVYF | 0.3899 | 12 | ABCPRED | 1 | NON-ALLERGEN | Non-Toxin | 0.07 | 0.2 | -0.72 | -1 | 3.8 | 1381.72 | outside |
| 442 | 454 | DSKVGGNYNYLY | 0.786 | 12 | ABCPRED | 1 | ALLERGEN | Non-Toxin | -0.16 | -0.99 | -0.29 | 0 | 6.18 | 1392.66 | outside |
| 166 | 178 | CTFEYVSQPFLM | 0.2955 | 12 | BCEPRED | 1 | NON-ALLERGEN | Non-Toxin | 0.07 | 0.55 | -0.82 | -1 | 4 | 1464.88 | outside |
| 943 | 955 | SALGKLQDVVNQ | 0.3782 | 12 | ABCPRED | 1 | NON-ALLERGEN | Non-Toxin | -0.13 | -0.11 | -0.02 | 0 | 6.19 | 1271.62 | outside |
| 364 | 382 | DYSVLYNSASFSTFKCYG | 0.3039 | 18 | ABCPRED | 1 | NON-ALLERGEN | Non-Toxin | -0.05 | -0.07 | -0.54 | 0 | 6.15 | 2052.48 | outside |
| 49 | 65 | HSTQDLFLPFFSNVTW | 0.5305 | 16 | ABCPRED | 1 | NON-ALLERGEN | Non-Toxin | 0.03 | 0.06 | -0.83 | -0.5 | 5.09 | 1939.39 | outside |
| 440 | 450 | NLDSKVGGNY | 0.7882 | 10 | ABCPRED | 1 | NON-ALLERGEN | Non-Toxin | -0.2 | -0.93 | 0.11 | 0 | 6.18 | 1066.28 | outside |
| 654 | 664 | EHVNNSYECD | 0.827 | 10 | ABCPRED,BCEPRED | 2 | NON-ALLERGEN | Non-Toxin | -0.33 | -1.61 | 0.44 | -2.5 | 4.14 | 1209.34 | outside |
| 615 | 631 | VNCTEVPVAIHADQLT | 0.4288 | 16 | ABCPRED | 1 | NON-ALLERGEN | Non-Toxin | 0 | 0.43 | -0.31 | -1.5 | 4.36 | 1710.17 | outside |
| 513 | 525 | LSFELLHAPATV | 0.9969 | 12 | ABCPRED | 1 | ALLERGEN | Non-Toxin | 0.14 | 1.02 | -0.67 | -0.5 | 5.25 | 1297.69 | outside |
| 332 | 342 | ITNLCPFGEV | 0.8724 | 10 | ABCPRED | 1 | NON-ALLERGEN | Non-Toxin | 0.11 | 0.81 | -0.58 | -1 | 4 | 1092.42 | outside |
| 524 | 536 | VCGPKKSTNLVK | -0.0923 | 12 | ABCPRED | 1 | ALLERGEN | Non-Toxin | -0.22 | -0.33 | 0.27 | 3 | 9.81 | 1273.73 | outside |
| 768 | 778 | TGIAVEQDKN | 1.0371 | 10 | ABCPRED | 1 | ALLERGEN | Non-Toxin | -0.23 | -0.85 | 0.52 | -1 | 4.38 | 1074.3 | outside |
| 200 | 210 | YFKIYSKHTP | 0.7972 | 10 | ABCPRED | 1 | ALLERGEN | Non-Toxin | -0.17 | -0.94 | -0.35 | 2.5 | 9.55 | 1283.63 | outside |
| 796 | 814 | DFGGFNFSQILPDPSKPS | 0.5036 | 18 | BCPREDS | 1 | NON-ALLERGEN | Non-Toxin | -0.08 | -0.51 | -0.04 | -1 | 4.21 | 1953.39 | outside |
| 679 | 689 | NSPRRARSVA | 0.2502 | 10 | BCEPRED | 1 | ALLERGEN | Non-Toxin | -0.55 | -1.24 | 0.73 | 3 | 12.31 | 1113.36 | outside |
| 483 | 493 | VEGFNCYFPL | 0.283 | 10 | ABCPRED | 1 | NON-ALLERGEN | Non-Toxin | 0.12 | 0.58 | -0.84 | -1 | 4 | 1188.5 | outside |
| 523 | 541 | TVCGPKKSTNLVKNKCVN | 0.2372 | 18 | AAP | 1 | NON-ALLERGEN | Non-Toxin | -0.26 | -0.49 | 0.21 | 4 | 9.72 | 1933.58 | outside |
| 1031 | 1040 | ECVLGQSKR | 0.9377 | 9 | BCEPRED | 1 | NON-ALLERGEN | Non-Toxin | -0.35 | -0.68 | 0.58 | 1 | 8.57 | 1019.31 | outside |
| 957 | 974 | QALNTLVKQLSSNFGAI | 0.0872 | 17 | COVIDep,literature | 2 | NON-ALLERGEN | Non-Toxin | -0.02 | 0.35 | -0.48 | 1 | 9.11 | 1804.34 | outside |
| 804 | 816 | QILPDPSKPSKR | 0.2594 | 12 | BCEPRED,IEDB | 2 | NON-ALLERGEN | Toxin | -0.4 | -1.45 | 0.77 | 2 | 10.01 | 1365.76 | outside |
| 437 | 450 | NSNNLDSKVGGNY | 0.6657 | 13 | IEDB | 1 | ALLERGEN | Non-Toxin | -0.27 | -1.32 | 0.14 | 0 | 6.18 | 1381.61 | outside |
| 515 | 525 | FELLHAPATV | 0.5982 | 10 | ABCPRED | 1 | ALLERGEN | Non-Toxin | 0.14 | 0.92 | -0.65 | -0.5 | 5.25 | 1097.42 | outside |
| 136 | 148 | CNDPFLGVYYHK | 0.6143 | 12 | ABCPRED | 1 | NON-ALLERGEN | Non-Toxin | -0.08 | -0.45 | -0.47 | 0.5 | 7.05 | 1455.82 | outside |
| 483 | 495 | VEGFNCYFPLQS | 0.5106 | 12 | ABCPRED | 1 | NON-ALLERGEN | Non-Toxin | 0.02 | 0.12 | -0.66 | -1 | 4 | 1403.74 | outside |
| 949 | 967 | QDVVNQNAQALNTLVKQL | -0.0557 | 18 | ABCPRED | 1 | ALLERGEN | Non-Toxin | -0.16 | -0.28 | -0.22 | 0 | 6.19 | 1996.54 | outside |
| 247 | 256 | SYLTPGDSS | 0.7324 | 9 | BCEPRED | 1 | ALLERGEN | Non-Toxin | -0.12 | -0.68 | -0.07 | -1 | 3.8 | 926.06 | outside |
| 603 | 621 | NTSNQVAVLYQDVNCTEV | 0.7978 | 18 | ABCPRED | 1 | NON-ALLERGEN | Non-Toxin | -0.13 | -0.17 | -0.28 | -2 | 3.67 | 1997.43 | outside |
| 697 | 715 | MSLGAENSVAYSNNSIAI | 0.9071 | 18 | ABCPRED | 1 | ALLERGEN | Non-Toxin | 0.01 | 0.3 | -0.4 | -1 | 4 | 1841.28 | outside |
| 439 | 449 | NNLDSKVGGN | 0.6721 | 10 | ABCPRED | 1 | NON-ALLERGEN | Non-Toxin | -0.26 | -1.15 | 0.36 | 0 | 6.19 | 1017.21 | outside |
| 939 | 955 | SSTASALGKLQDVVNQ | 0.4821 | 16 | ABCPRED | 1 | NON-ALLERGEN | Non-Toxin | -0.12 | -0.11 | -0.03 | 0 | 6.19 | 1618.01 | outside |
| 144 | 154 | YYHKNNKSWM | 0.3777 | 10 | BCEPRED,IEDB,literature | 3 | NON-ALLERGEN | Non-Toxin | -0.35 | -2.04 | -0.31 | 2.5 | 9.55 | 1370.69 | outside |
| 929 | 947 | SAIGKIQDSLSSTASALG | 0.6359 | 18 | ABCPRED | 1 | NON-ALLERGEN | Non-Toxin | -0.02 | 0.31 | -0.08 | 0 | 6.19 | 1706.14 | outside |
| 439 | 448 | NNLDSKVGG | 0.8904 | 9 | IEDB | 1 | NON-ALLERGEN | Non-Toxin | -0.22 | -0.89 | 0.38 | 0 | 6.19 | 903.09 | outside |
| 202 | 209 | KIYSKHT | 0.7773 | 7 | IEDB | 1 | NON-ALLERGEN | Non-Toxin | -0.33 | -1.33 | 0.19 | 2.5 | 9.72 | 876.12 | outside |
| 472 | 488 | IYQAGSTPCNGVEGFN | 0.0278 | 16 | ABCPRED | 1 | ALLERGEN | Non-Toxin | -0.03 | -0.24 | -0.38 | -1 | 4 | 1657.02 | outside |
| 599 | 617 | TPGTNTSNQVAVLYQDVN | 0.4335 | 18 | ABCPRED | 1 | NON-ALLERGEN | Non-Toxin | -0.13 | -0.5 | -0.33 | -1 | 3.8 | 1921.32 | outside |
| 1047 | 1065 | YHLMSFPQSAPHGVVFLH | 0.2078 | 18 | ABCPRED | 1 | ALLERGEN | Non-Toxin | 0.08 | 0.32 | -0.91 | 1.5 | 7.35 | 2067.67 | outside |
| 585 | 598 | LDITPCSFGGVSV | 1.882 | 13 | literature | 1 | NON-ALLERGEN | Non-Toxin | 0.14 | 1.06 | -0.53 | -1 | 3.8 | 1294.67 | outside |
| 248 | 259 | YLTPGDSSSGW | 0.624 | 11 | IEDB | 1 | ALLERGEN | Non-Toxin | -0.05 | -0.67 | -0.36 | -1 | 3.8 | 1169.36 | outside |
| 957 | 975 | QALNTLVKQLSSNFGAIS | 0.2655 | 18 | ABCPRED | 1 | NON-ALLERGEN | Non-Toxin | -0.03 | 0.28 | -0.44 | 1 | 9.11 | 1891.43 | outside |
| 695 | 713 | YTMSLGAENSVAYSNNSI | 0.6376 | 18 | ABCPRED,FBCPred | 2 | ALLERGEN | Non-Toxin | -0.06 | -0.16 | -0.42 | -1 | 4 | 1921.32 | outside |
| 743 | 763 | CGDSTECSNLLLQYGSFCTQ | 0.5547 | 20 | AAP | 1 | NON-ALLERGEN | Non-Toxin | -0.09 | -0.08 | -0.33 | -2 | 3.67 | 2169.67 | outside |
| 613 | 631 | QDVNCTEVPVAIHADQLT | 0.6219 | 18 | ABCPRED | 1 | NON-ALLERGEN | Non-Toxin | -0.08 | -0.01 | -0.1 | -2.5 | 4.03 | 1953.42 | outside |

| | | | | | | | | | | | | | | | |
|---|---|---|---|---|---|---|---|---|---|---|---|---|---|---|---|
| 798 | 814 | GGFNFSQILPDPSKPS | 0.3714 | 16 | ABCPRED | 1 | NON-ALLERGEN | Non-Toxin | -0.08 | -0.53 | -0.08 | 0 | 6.19 | 1691.1 | outside |
| 56 | 72 | LPFFSNVTWFHAIHVS | 0.7606 | 16 | ABCPRED | 1 | NON-ALLERGEN | Non-Toxin | 0.16 | 0.76 | -1.16 | 1 | 7.26 | 1902.42 | outside |
| 693 | 709 | IAYTMSLGAENSVAYS | 0.7484 | 16 | ABCPRED | 1 | ALLERGEN | Non-Toxin | 0.05 | 0.42 | -0.55 | -1 | 4 | 1677.08 | outside |
| 867 | 883 | DEMIAQYTSALLAGTI | 0.2175 | 16 | ABCPRED | 1 | NON-ALLERGEN | Non-Toxin | 0.07 | 0.59 | -0.41 | -2 | 3.67 | 1697.17 | outside |
| 437 | 449 | NSNNLDSKVGGN | 0.6962 | 12 | BCEPRED,IEDB | 2 | NON-ALLERGEN | Non-Toxin | -0.29 | -1.32 | 0.34 | 0 | 6.19 | 1218.42 | outside |
| 556 | 568 | NKKFLPFQQFGR | 0.6233 | 12 | ABCPRED | 1 | ALLERGEN | Non-Toxin | -0.29 | -1.05 | 0.03 | 3 | 11.17 | 1509.95 | outside |
| 947 | 959 | KLQDVVNQNAQA | 0.5943 | 12 | ABCPRED,AAP | 2 | NON-ALLERGEN | Non-Toxin | -0.26 | -0.76 | 0.1 | 0 | 6.19 | 1327.64 | outside |
| 555 | 563 | SNKKFLPF | 1.3952 | 8 | IEDB,literature | 2 | ALLERGEN | Non-Toxin | -0.18 | -0.54 | -0.04 | 2 | 10.02 | 980.28 | outside |
| 222 | 234 | ALEPLVDLPIGI | 0.7479 | 12 | ABCPRED | 1 | NON-ALLERGEN | Non-Toxin | 0.21 | 1.32 | -0.42 | -2 | 3.67 | 1249.7 | outside |
| 595 | 615 | VSVITPGTNTSNQVAVLYQD | 0.3742 | 20 | ABCPRED | 1 | NON-ALLERGEN | Non-Toxin | -0.03 | 0.12 | -0.46 | -1 | 3.8 | 2106.62 | outside |
| 55 | 67 | FLPFFSNVTWFH | 0.637 | 12 | ABCPRED | 1 | NON-ALLERGEN | Toxin | 0.19 | 0.71 | -1.43 | 0.5 | 7.1 | 1541.94 | outside |
| 941 | 959 | TASALGKLQDVVNQNAQA | 0.5539 | 18 | ABCPRED | 1 | ALLERGEN | Non-Toxin | -0.13 | -0.2 | -0.09 | 0 | 6.19 | 1828.28 | outside |
| 933 | 945 | KIQDSLSSTASA | 0.5795 | 12 | ABCPRED | 1 | ALLERGEN | Non-Toxin | -0.16 | -0.24 | 0.2 | 0 | 6.19 | 1207.46 | outside |
| 282 | 300 | NGTITDAVDCALDPLSET | 0.558 | 18 | ABCPRED | 1 | ALLERGEN | Non-Toxin | -0.07 | 0 | 0.13 | -4 | 3.38 | 1835.22 | outside |
| 134 | 146 | QFCNDPFLGVYY | 0.3132 | 12 | ABCPRED | 1 | NON-ALLERGEN | Non-Toxin | 0.03 | 0.08 | -0.88 | -1 | 3.8 | 1465.81 | outside |
| 616 | 633 | NCTEVPVAIHADQLTPT | 0.3987 | 17 | IEDB,literature | 2 | NON-ALLERGEN | Non-Toxin | -0.05 | 0.02 | -0.23 | -1.5 | 4.36 | 1809.27 | outside |
| 617 | 633 | CTEVPVAIHADQLTPT | 0.2794 | 16 | ABCPRED | 1 | NON-ALLERGEN | Non-Toxin | -0.01 | 0.24 | -0.26 | -1.5 | 4.36 | 1695.15 | outside |
| 1128 | 1144 | VVIGIVNNTVYDPLQP | 0.7712 | 16 | ABCPRED | 1 | NON-ALLERGEN | Non-Toxin | 0.08 | 0.62 | -0.66 | -1 | 3.8 | 1741.27 | outside |
| 849 | 861 | LICAQKFNGLTV | 0.3561 | 12 | ABCPRED | 1 | NON-ALLERGEN | Non-Toxin | 0.07 | 0.95 | -0.66 | 1 | 8.57 | 1306.77 | outside |
| 142 | 152 | GVYYHKNNKS | 0.7157 | 10 | ABCPRED | 1 | NON-ALLERGEN | Toxin | -0.34 | -1.76 | 0.01 | 2.5 | 9.55 | 1209.47 | outside |
| 956 | 974 | AQALNTLVKQLSSNFGAI | 0.1254 | 18 | ABCPRED | 1 | NON-ALLERGEN | Non-Toxin | 0 | 0.43 | -0.48 | 1 | 9.11 | 1875.43 | outside |
| 223 | 235 | LEPLVDLPIGIN | 0.8347 | 12 | ABCPRED | 1 | NON-ALLERGEN | Non-Toxin | 0.14 | 0.87 | -0.36 | -2 | 3.67 | 1292.73 | outside |
| 619 | 631 | EVPVAIHADQLT | 0.0344 | 12 | ABCPRED | 1 | ALLERGEN | Non-Toxin | 0.01 | 0.36 | -0.19 | -1.5 | 4.36 | 1292.63 | outside |
| 478 | 490 | TPCNGVEGFNCY | 0.2717 | 12 | ABCPRED | 1 | ALLERGEN | Non-Toxin | -0.05 | -0.24 | -0.44 | -1 | 4 | 1303.59 | outside |
| 771 | 782 | AVEQDKNTQEV | 0.3969 | 11 | BCEPRED,IEDB | 2 | ALLERGEN | Non-Toxin | -0.36 | -1.4 | 0.79 | -2 | 4.14 | 1260.48 | outside |
| 143 | 153 | VYYHKNNKSW | 0.4497 | 10 | ABCPRED | 1 | NON-ALLERGEN | Non-Toxin | -0.32 | -1.81 | -0.33 | 2.5 | 9.55 | 1338.63 | outside |
| 941 | 953 | TASALGKLQDVV | 0.7262 | 12 | ABCPRED | 1 | NON-ALLERGEN | Non-Toxin | -0.01 | 0.57 | -0.13 | 0 | 6.19 | 1201.56 | outside |
| 677 | 686 | QTNSPRRAR | -0.118 | 9 | BCEPRED | 1 | ALLERGEN | Non-Toxin | -0.76 | -2.42 | 0.98 | 3 | 12.31 | 1085.3 | outside |
| 44 | 52 | RSSVLHST | 0.5459 | 8 | IEDB,literature | 2 | ALLERGEN | Non-Toxin | -0.26 | -0.35 | -0.04 | 1.5 | 10.11 | 886.08 | outside |
| 932 | 944 | GKIQDSLSSTAS | 0.5659 | 12 | ABCPRED,IEDB | 2 | NON-ALLERGEN | Non-Toxin | -0.17 | -0.42 | 0.24 | 0 | 6.19 | 1193.44 | outside |
| 248 | 258 | YLTPGDSSSG | 0.661 | 10 | ABCPRED | 1 | ALLERGEN | Non-Toxin | -0.09 | -0.65 | -0.06 | -1 | 3.8 | 983.13 | outside |
| 585 | 601 | LDITPCSFGGVSVITP | 1.3502 | 16 | ABCPRED | 1 | ALLERGEN | Non-Toxin | 0.14 | 1 | -0.57 | -1 | 3.8 | 1606.1 | outside |
| 584 | 594 | ILDITPCSFG | 1.5391 | 10 | ABCPRED | 1 | NON-ALLERGEN | Non-Toxin | 0.16 | 1.11 | -0.6 | -1 | 3.8 | 1065.39 | outside |
| 596 | 616 | SVITPGTNTSNQVAVLYQDV | 0.3129 | 20 | ABCPRED,FBCPred | 2 | ALLERGEN | Non-Toxin | -0.03 | 0.12 | -0.46 | -1 | 3.8 | 2106.62 | outside |
| 325 | 335 | SIVRFPNITN | 0.2809 | 10 | ABCPRED | 1 | NON-ALLERGEN | Non-Toxin | -0.09 | 0.14 | -0.43 | 1 | 10.11 | 1160.48 | outside |
| 310 | 320 | KGIYQTSNFR | 0.3992 | 10 | ABCPRED | 1 | NON-ALLERGEN | Non-Toxin | -0.31 | -1.13 | -0.03 | 2 | 10.01 | 1213.5 | outside |
| 580 | 592 | QTLEILDITPCS | 1.2575 | 12 | ABCPRED | 1 | NON-ALLERGEN | Non-Toxin | -0.01 | 0.4 | -0.21 | -2 | 3.67 | 1332.71 | outside |
| 180 | 189 | EGKQGNFKN | 1.1232 | 9 | IEDB | 1 | NON-ALLERGEN | Non-Toxin | -0.43 | -2.2 | 0.79 | 1 | 8.94 | 1021.23 | outside |
| 701 | 720 | AENSVAYSNNSIAIPTNFT | 0.5643 | 19 | ellipro | 1 | NON-ALLERGEN | Non-Toxin | -0.05 | -0.15 | -0.39 | -1 | 4 | 2013.41 | outside |
| 783 | 793 | AQVKQIYKTP | 0.2356 | 10 | ABCPRED | 1 | ALLERGEN | Non-Toxin | -0.23 | -0.79 | -0.01 | 2 | 9.72 | 1175.54 | outside |
| 333 | 345 | TNLCPFGEVFNA | 0.2856 | 12 | ABCPRED | 1 | NON-ALLERGEN | Non-Toxin | 0.05 | 0.39 | -0.57 | -1 | 4 | 1311.64 | outside |
| 584 | 602 | ILDITPCSFGGVSVITPG | 1.1031 | 18 | BCEPRED | 1 | NON-ALLERGEN | Non-Toxin | 0.18 | 1.12 | -0.61 | -1 | 3.8 | 1776.35 | outside |
| 109 | 116 | TLDSKTQ | 1.2584 | 7 | IEDB | 1 | NON-ALLERGEN | Non-Toxin | -0.37 | -1.33 | 0.56 | 0 | 6.19 | 791.95 | outside |
| 688 | 706 | ASQSIIAYTMSLGAENSV | 0.4324 | 18 | ABCPRED | 1 | ALLERGEN | Toxin | 0.03 | 0.46 | -0.43 | -1 | 4 | 1842.31 | outside |
| 407 | 417 | VRQIAPGQTG | 1.3856 | 10 | ABCPRED | 1 | ALLERGEN | Non-Toxin | -0.15 | -0.41 | -0.08 | 1 | 10.11 | 1026.31 | outside |
| 247 | 257 | SYLTPGDSSS | 0.4882 | 10 | ABCPRED | 1 | ALLERGEN | Non-Toxin | -0.13 | -0.69 | -0.03 | -1 | 3.8 | 1013.15 | outside |
| 688 | 704 | ASQSIIAYTMSLGAEN | 0.583 | 16 | ABCPRED | 1 | ALLERGEN | Non-Toxin | 0.02 | 0.3 | -0.41 | -1 | 4 | 1656.07 | outside |
| 331 | 343 | NITNLCPFGEVF | 0.6782 | 12 | ABCPRED | 1 | NON-ALLERGEN | Non-Toxin | 0.09 | 0.62 | -0.67 | -1 | 4 | 1353.73 | outside |
| 108 | 117 | TTLDSKTQS | 1.0106 | 9 | IEDB | 1 | NON-ALLERGEN | Non-Toxin | -0.34 | -1.2 | 0.42 | 0 | 6.19 | 980.16 | outside |
| 514 | 526 | SFELLHAPATVC | 0.7389 | 12 | ABCPRED | 1 | NON-ALLERGEN | Non-Toxin | 0.1 | 0.91 | -0.6 | -0.5 | 5.25 | 1287.66 | outside |
| 678 | 696 | TNSPRRARSVASQSIIAY | 0.1229 | 18 | ABCPRED | 1 | ALLERGEN | Non-Toxin | -0.28 | -0.48 | 0.07 | 3 | 11.72 | 1977.45 | outside |
| 1229 | 1247 | MVTIMLCCMTSCCSCLKG | -0.039 | 18 | ABCPRED | 1 | NON-ALLERGEN | Non-Toxin | 0.08 | 1.51 | -0.72 | 1 | 8.1 | 1927.75 | transmembrane-region_inside |
| 469 | 485 | STEIYQAGSTPCNGVE | 0.0539 | 16 | ABCPRED | 1 | NON-ALLERGEN | Non-Toxin | -0.1 | -0.48 | -0.06 | -2 | 3.8 | 1655.98 | outside |
| 473 | 493 | YQAGSTPCNGVEGFNCYFPL | 0.4046 | 20 | FBCPred | 1 | NON-ALLERGEN | Non-Toxin | 0 | -0.11 | -0.59 | -1 | 4 | 2167.68 | outside |
| 141 | 153 | LGVYYHKNNKSW | 0.8156 | 12 | AAP | 1 | NON-ALLERGEN | Non-Toxin | -0.21 | -1.23 | -0.42 | 2.5 | 9.55 | 1508.88 | outside |
| 367 | 385 | VLYNSASFSTFKCYGVSP | 0.5346 | 18 | ABCPRED | 1 | NON-ALLERGEN | Non-Toxin | 0.02 | 0.34 | -0.66 | 1 | 8.51 | 1970.47 | outside |
| 135 | 147 | FCNDPFLGVYYH | 0.2914 | 12 | ABCPRED | 1 | ALLERGEN | Toxin | 0.06 | 0.11 | -0.93 | -0.5 | 5.09 | 1474.82 | outside |
| 1168 | 1186 | DISGINASVVNIQKEIDR | 0.3309 | 18 | ABCPRED | 1 | NON-ALLERGEN | Non-Toxin | -0.17 | -0.18 | 0.31 | -1 | 4.56 | 1971.47 | outside |
| 1213 | 1229 | PWYIWLGFIAGLIAIV | 0.6318 | 16 | ABCPRED | 1 | NON-ALLERGEN | Non-Toxin | 0.41 | 1.92 | -1.56 | 0 | 5.88 | 1832.52 | transmembrane-region |
| 84 | 96 | LPFNDGVYFAST | 0.5137 | 12 | ABCPRED | 1 | NON-ALLERGEN | Non-Toxin | 0.07 | 0.3 | -0.67 | -1 | 3.8 | 1330.62 | outside |
| 941 | 957 | TASALGKLQDVVNQNA | 0.5097 | 16 | ABCPRED | 1 | ALLERGEN | Non-Toxin | -0.12 | -0.12 | -0.09 | 0 | 6.19 | 1629.04 | outside |
| 85 | 95 | PFNDGVYFAS | 0.1769 | 10 | ABCPRED | 1 | ALLERGEN | Non-Toxin | 0.05 | 0.05 | -0.58 | -1 | 3.8 | 1116.32 | outside |
| 108 | 128 | TTLDSKTQSLLIVNNATNVV | 0.6842 | 20 | ABCPRED | 1 | ALLERGEN | Non-Toxin | -0.07 | 0.23 | -0.32 | 0 | 6.19 | 2131.72 | outside |
| 694 | 712 | AYTMSLGAENSVAYSNNS | 0.5996 | 18 | ABCPRED | 1 | NON-ALLERGEN | Non-Toxin | -0.08 | -0.31 | -0.35 | -1 | 4 | 1879.23 | outside |
| 472 | 486 | IYQAGSTPCNGVEG | -0.0612 | 14 | FBCPred,IEDB | 2 | NON-ALLERGEN | Non-Toxin | -0.03 | -0.22 | -0.27 | -1 | 4 | 1395.71 | outside |
| 552 | 560 | LTESNKKF | 0.7708 | 8 | BCEPRED | 1 | ALLERGEN | Non-Toxin | -0.35 | -1.21 | 0.6 | 1 | 8.94 | 966.21 | outside |
| 584 | 600 | ILDITPCSFGGVSVIT | 1.2689 | 16 | ABCPRED | 1 | NON-ALLERGEN | Non-Toxin | 0.19 | 1.38 | -0.68 | -1 | 3.8 | 1622.15 | outside |
| 371 | 383 | SASFSTFKCYGV | 0.5926 | 12 | ABCPRED | 1 | NON-ALLERGEN | Non-Toxin | 0.01 | 0.45 | -0.57 | 1 | 8.54 | 1296.61 | outside |
| 1013 | 1021 | IRAAEIRA | 0.2791 | 8 | BCEPRED | 1 | ALLERGEN | Non-Toxin | -0.24 | 0.24 | 0.49 | 1 | 9.95 | 899.16 | outside |
| 410 | 422 | IAPGQTGKIADY | 1.6626 | 12 | ABCPRED,literature | 2 | NON-ALLERGEN | Non-Toxin | -0.04 | -0.23 | -0.09 | 0 | 6.18 | 1233.56 | outside |
| 221 | 231 | SALEPLVDLP | -0.2366 | 10 | ABCPRED | 1 | NON-ALLERGEN | Non-Toxin | 0.06 | 0.64 | -0.11 | -2 | 3.67 | 1053.36 | outside |
| 695 | 705 | YTMSLGAENS | 0.8273 | 10 | ABCPRED | 1 | ALLERGEN | Non-Toxin | -0.07 | -0.35 | -0.25 | -1 | 4 | 1072.29 | outside |
| 1204 | 1210 | GKYEQY | 1.2821 | 6 | literature | 1 | ALLERGEN | Non-Toxin | -0.37 | -2.32 | 0.27 | 0 | 6.35 | 786.92 | outside |
| 160 | 178 | YSSANNCTFEYVSQPFLM | 0.1607 | 18 | ABCPRED | 1 | NON-ALLERGEN | Non-Toxin | -0.04 | -0.08 | -0.64 | -1 | 4 | 2101.58 | outside |
| 381 | 389 | GVSPTKLN | 1.9197 | 8 | IEDB | 1 | ALLERGEN | Non-Toxin | -0.13 | -0.36 | -0.03 | 1 | 9.11 | 815.05 | outside |
| 258 | 274 | WTAGAAAYYVGYLQPR | 0.9545 | 16 | ABCPRED | 1 | NON-ALLERGEN | Non-Toxin | 0.01 | -0.04 | -0.8 | 1 | 8.83 | 1787.23 | outside |
| 216 | 226 | LPQGFSALEP | 0.8025 | 10 | ABCPRED | 1 | ALLERGEN | Non-Toxin | 0.04 | 0.08 | -0.31 | -1 | 4 | 1058.34 | outside |

| | | | | | | | | | | | | | | | |
|---|---|---|---|---|---|---|---|---|---|---|---|---|---|---|---|
| 224 | 236 | EPLVDLPIGINI | 0.7968 | 12 | literature | 1 | NON-ALLERGEN | Non-Toxin | 0.15 | 0.93 | -0.36 | -2 | 3.67 | 1292.73 | outside |
| 738 | 748 | CTMYICGDST | -1.1142 | 10 | ABCPRED | 1 | ALLERGEN | Non-Toxin | -0.01 | 0.4 | -0.49 | -1 | 3.8 | 1093.38 | outside |
| 79 | 97 | FDNPVLPFNDGVYFASTE | 0.2213 | 18 | ABCPRED | 1 | ALLERGEN | Non-Toxin | 0 | -0.08 | -0.32 | -3 | 3.5 | 2032.44 | outside |
| 555 | 565 | SNKKFLPFQQ | 0.9488 | 10 | ABCPRED | 1 | ALLERGEN | Non-Toxin | -0.28 | -1.13 | 0.01 | 2 | 10.02 | 1236.58 | outside |
| 80 | 92 | DNPVLPFNDGVY | 0.1476 | 12 | AAP | 1 | ALLERGEN | Non-Toxin | -0.04 | -0.33 | -0.27 | -2 | 3.57 | 1349.63 | outside |
| 482 | 494 | GVEGFNCYFPLQ | 0.9537 | 12 | ABCPRED | 1 | NON-ALLERGEN | Non-Toxin | 0.05 | 0.16 | -0.68 | -1 | 4 | 1373.72 | outside |
| 678 | 688 | TNSPRRARSV | 0.221 | 10 | ABCPRED | 1 | ALLERGEN | Non-Toxin | -0.59 | -1.49 | 0.74 | 3 | 12.31 | 1143.39 | outside |
| 797 | 809 | FGGFNFSQILPD | 0.6291 | 12 | ABCPRED | 1 | NON-ALLERGEN | Non-Toxin | 0.09 | 0.25 | -0.62 | -1 | 3.8 | 1341.66 | outside |
| 771 | 783 | AVEQDKNTQEVF | 0.2192 | 12 | ABCPRED,BCEPRED,literature | 3 | NON-ALLERGEN | Non-Toxin | -0.28 | -1.05 | 0.52 | -2 | 4.14 | 1407.67 | outside |
| 532 | 544 | NLVKNKCVNFNF | 1.3432 | 12 | ABCPRED | 1 | NON-ALLERGEN | Non-Toxin | -0.16 | -0.12 | -0.33 | 2 | 9.36 | 1439.87 | outside |
| 135 | 143 | FCNDPFLG | 0.1982 | 8 | IEDB | 1 | NON-ALLERGEN | Non-Toxin | 0.07 | 0.36 | -0.57 | -1 | 3.8 | 912.13 | outside |
| 737 | 747 | DCTMYICGDS | -1.4237 | 10 | ABCPRED | 1 | NON-ALLERGEN | Non-Toxin | -0.06 | 0.12 | -0.15 | -2 | 3.57 | 1107.36 | outside |
| 369 | 381 | YNSASFSTFKCY | 0.3813 | 12 | ABCPRED | 1 | NON-ALLERGEN | Non-Toxin | -0.1 | -0.27 | -0.62 | 1 | 8.51 | 1417.7 | outside |
| 773 | 783 | EQDKNTQEVF | 0.093 | 10 | ABCPRED | 1 | ALLERGEN | Non-Toxin | -0.41 | -1.86 | 0.82 | -2 | 4.14 | 1237.43 | outside |
| 140 | 152 | FLGVYYHKNNKS | 0.7295 | 12 | ABCPRED | 1 | ALLERGEN | Non-Toxin | -0.19 | -0.92 | -0.35 | 2.5 | 9.55 | 1469.84 | outside |
| 798 | 810 | GGFNFSQILPDP | 0.6221 | 12 | ABCPRED | 1 | NON-ALLERGEN | Non-Toxin | 0.03 | -0.12 | -0.41 | -1 | 3.8 | 1291.6 | outside |
| 1237 | 1253 | MTSCCSCLKGCCSCGS | -0.1073 | 16 | ABCPRED | 1 | ALLERGEN | Non-Toxin | -0.06 | 0.76 | -0.33 | 1 | 8.04 | 1573.1 | inside |
| 201 | 209 | FKIYSKHT | 1.1661 | 8 | BCEPRED | 1 | ALLERGEN | Non-Toxin | -0.21 | -0.81 | -0.15 | 2.5 | 9.72 | 1023.31 | outside |
| 1235 | 1253 | CCMTSCCSCLKGCCSCGS | -0.198 | 18 | ABCPRED | 1 | ALLERGEN | Non-Toxin | -0.05 | 0.95 | -0.41 | 1 | 7.94 | 1779.4 | transmembrane-region_inside |
| 598 | 618 | ITPGTNTSNQVAVLYQDVNC | 0.7787 | 20 | ABCPRED | 1 | ALLERGEN | Non-Toxin | -0.08 | -0.1 | -0.44 | -1 | 3.8 | 2137.65 | outside |
| 581 | 597 | TLEILDITPCSFGGVS | 1.6878 | 16 | ABCPRED | 1 | ALLERGEN | Non-Toxin | 0.11 | 0.86 | -0.4 | -2 | 3.67 | 1652.13 | outside |
| 994 | 1012 | DRLITGRLQSLQTYVTQQ | -0.2656 | 18 | IEDB | 1 | ALLERGEN | Non-Toxin | -0.26 | -0.61 | -0.12 | 1 | 9.1 | 2120.68 | outside |
| 634 | 652 | RVYSTGSNVFQTRAGCLI | 0.6847 | 18 | ABCPRED | 1 | NON-ALLERGEN | Non-Toxin | -0.12 | 0.15 | -0.37 | 2 | 9.55 | 1972.51 | outside |
| 328 | 338 | RFPNITNLCP | 1.4779 | 10 | ABCPRED | 1 | NON-ALLERGEN | Non-Toxin | -0.14 | -0.18 | -0.41 | 1 | 8.6 | 1174.52 | outside |
| 1165 | 1185 | DLGDISGINASVVNIQKEID | 0.8227 | 20 | BCPREDS | 1 | NON-ALLERGEN | Non-Toxin | -0.06 | 0.06 | 0.18 | -3 | 3.84 | 2100.62 | outside |
| 16 | 28 | VNLTTRTQLPPA | 1.4268 | 12 | ABCPRED | 1 | NON-ALLERGEN | Non-Toxin | -0.16 | -0.27 | -0.28 | 1 | 10.11 | 1310.69 | outside |
| 423 | 433 | YKLPDDFTGC | -0.4848 | 10 | ABCPRED | 1 | NON-ALLERGEN | Non-Toxin | -0.14 | -0.58 | 0.1 | -1 | 4.21 | 1158.42 | outside |
| 910 | 926 | GVTQNVLYENQKLIAN | 0.1015 | 16 | ABCPRED | 1 | ALLERGEN | Non-Toxin | -0.12 | -0.31 | -0.29 | 0 | 6.35 | 1804.29 | outside |
| 1110 | 1128 | YEPQIITTDNTFVSGNCD | 0.3995 | 18 | ABCPRED,BCPREDS,FBCPred | 3 | NON-ALLERGEN | Non-Toxin | -0.11 | -0.48 | -0.12 | -3 | 3.5 | 2017.41 | outside |
| 326 | 338 | IVRFPNITNLCP | 0.9596 | 12 | ABCPRED | 1 | NON-ALLERGEN | Non-Toxin | -0.02 | 0.57 | -0.62 | 1 | 8.6 | 1386.85 | outside |
| 635 | 653 | VYSTGSNVFQTRAGCLIG | 0.6447 | 18 | ABCPRED | 1 | NON-ALLERGEN | Non-Toxin | -0.01 | 0.38 | -0.54 | 1 | 8.57 | 1873.38 | outside |
| 607 | 625 | QVAVLYQDVNCTEVPVAI | 0.7832 | 18 | ABCPRED | 1 | ALLERGEN | Non-Toxin | 0.05 | 0.79 | -0.51 | -2 | 3.67 | 1961.53 | outside |
| 914 | 930 | NVLYENQKLIANQFNS | 0.0011 | 16 | ABCPRED | 1 | NON-ALLERGEN | Non-Toxin | -0.17 | -0.6 | -0.29 | 0 | 6.35 | 1895.35 | outside |
| 1262 | 1271 | EPVLKGVKL | 1.2301 | 9 | IEDB | 1 | NON-ALLERGEN | Non-Toxin | -0.07 | 0.3 | 0.27 | 1 | 8.94 | 982.37 | inside |
| 582 | 600 | LEILDITPCSFGGVSVIT | 1.428 | 18 | ABCPRED | 1 | ALLERGEN | Non-Toxin | 0.17 | 1.24 | -0.54 | -2 | 3.67 | 1864.46 | outside |
| 830 | 848 | DAGFIKQYGDCLGDIAAR | 0.3839 | 18 | ABCPRED | 1 | NON-ALLERGEN | Non-Toxin | -0.1 | -0.08 | 0.14 | -1 | 4.43 | 1913.39 | outside |
| 402 | 410 | IRGDEVRQ | -0.6918 | 8 | BCEPRED | 1 | ALLERGEN | Non-Toxin | -0.51 | -1.4 | 1.11 | 0 | 6.42 | 972.18 | outside |
| 1111 | 1123 | EPQIITTDNTFV | 0.6005 | 12 | AAP | 1 | NON-ALLERGEN | Non-Toxin | -0.06 | -0.14 | -0.2 | -2 | 3.67 | 1377.69 | outside |
| 234 | 252 | NITRFQTLLALHRSYLTP | 0.3091 | 18 | ABCPRED | 1 | NON-ALLERGEN | Non-Toxin | -0.13 | -0.04 | -0.52 | 2.5 | 10.84 | 2144.78 | outside |
| 723 | 737 | TTEILPVSMTKTSV | 1.2571 | 14 | AAP | 1 | NON-ALLERGEN | Non-Toxin | -0.03 | 0.37 | -0.21 | 0 | 6.35 | 1506.98 | outside |
| 322 | 334 | PTESIVRFPNIT | -0.0307 | 12 | BCPREDS | 1 | NON-ALLERGEN | Non-Toxin | -0.1 | -0.07 | -0.16 | 0 | 6.36 | 1373.74 | outside |
| 200 | 212 | YFKIYSKHTPIN | 0.7893 | 12 | BCEPRED | 1 | ALLERGEN | Non-Toxin | -0.14 | -0.7 | -0.42 | 2.5 | 9.55 | 1510.93 | outside |
| 383 | 390 | SPTKLND | 0.839 | 7 | IEDB | 1 | ALLERGEN | Non-Toxin | -0.35 | -1.46 | 0.61 | 0 | 6.19 | 773.93 | outside |
| 11 | 27 | VSSQCVNLTTRTQLPP | 1.409 | 16 | ABCPRED | 1 | NON-ALLERGEN | Non-Toxin | -0.17 | -0.21 | -0.29 | 1 | 8.6 | 1744.23 | outside |
| 193 | 209 | VFKNIDGYFKIYSKHT | 0.0763 | 16 | ABCPRED | 1 | NON-ALLERGEN | Non-Toxin | -0.13 | -0.48 | -0.19 | 2.5 | 9.43 | 1960.5 | outside |
| 250 | 270 | TPGDSSSGWTAGAAAYYVGY | 0.3433 | 20 | ABCPRED,AAP | 2 | NON-ALLERGEN | Non-Toxin | 0.03 | -0.19 | -0.53 | -1 | 3.8 | 1981.33 | outside |
| 302 | 312 | TLKSFTVEKG | 0.2495 | 10 | ABCPRED | 1 | NON-ALLERGEN | Non-Toxin | -0.16 | -0.31 | 0.27 | 1 | 8.94 | 1109.43 | outside |
| 578 | 596 | DPQTLEILDITPCSFGGV | 1.4948 | 18 | ABCPRED | 1 | NON-ALLERGEN | Non-Toxin | 0.03 | 0.33 | -0.19 | -3 | 3.5 | 1905.42 | outside |
| 449 | 458 | YNYLYRLFR | -0.8692 | 9 | BCEPRED | 1 | ALLERGEN | Non-Toxin | -0.27 | -0.67 | -0.76 | 2 | 9.72 | 1307.64 | outside |
| 516 | 532 | ELLHAPATVCGPKKST | -0.0643 | 16 | ABCPRED | 1 | NON-ALLERGEN | Non-Toxin | -0.11 | -0.15 | 0.06 | 1.5 | 8.54 | 1652.17 | outside |
| 866 | 882 | TDEMIAQYTSALLAGT | 0.262 | 16 | ABCPRED | 1 | NON-ALLERGEN | Non-Toxin | 0.01 | 0.27 | -0.33 | -2 | 3.67 | 1685.11 | outside |
| 237 | 247 | RFQTLLALHR | 0.3353 | 10 | ABCPRED | 1 | ALLERGEN | Non-Toxin | -0.23 | -0.04 | -0.31 | 2.5 | 12.01 | 1254.65 | outside |
| 1202 | 1211 | ELGKYEQYI | 0.5415 | 9 | IEDB | 1 | NON-ALLERGEN | Non-Toxin | -0.17 | -1.01 | 0.11 | -1 | 4.54 | 1142.41 | outside |
| 372 | 388 | ASFSTFKCYGVSPTKL | 0.912 | 16 | ABCPRED | 1 | NON-ALLERGEN | Non-Toxin | -0.04 | 0.19 | -0.38 | 2 | 9.25 | 1736.23 | outside |
| 250 | 262 | TPGDSSSGWTAG | 0.0746 | 12 | ABCPRED,BCPREDS,FBCPred | 3 | ALLERGEN | Non-Toxin | -0.07 | -0.77 | -0.07 | -1 | 3.8 | 1122.27 | outside |
| 1110 | 1130 | YEPQIITTDNTFVSGNCDVV | 0.3685 | 20 | ABCPRED,FBCPred | 2 | NON-ALLERGEN | Toxin | -0.05 | -0.01 | -0.26 | -3 | 3.5 | 2215.71 | outside |
| 854 | 866 | KFNGLTVLPPLL | 0.3861 | 12 | BCPREDS | 1 | ALLERGEN | Non-Toxin | 0.11 | 0.88 | -0.7 | 1 | 9.11 | 1311.82 | outside |
| 374 | 386 | FSTFKCYGVSPT | 0.8097 | 12 | ABCPRED,BCEPRED | 2 | ALLERGEN | Non-Toxin | -0.01 | 0.18 | -0.58 | 1 | 8.54 | 1336.68 | outside |
| 1254 | 1262 | CKFDEDDS | -0.1581 | 8 | BCEPRED | 1 | ALLERGEN | Non-Toxin | -0.44 | -1.68 | 1.48 | -3 | 3.84 | 958.05 | inside |
| 386 | 395 | KLNDLCFTN | 2.9016 | 9 | literature | 1 | ALLERGEN | Non-Toxin | -0.17 | -0.24 | -0.12 | 0 | 6.16 | 1067.35 | outside |
| 1010 | 1028 | QQLIRAAEIRASANLAAT | 0.3283 | 18 | ABCPRED | 1 | NON-ALLERGEN | Non-Toxin | -0.14 | 0.16 | -0.04 | 1 | 9.95 | 1897.42 | outside |
| 1240 | 1258 | CCSCLKGCCSCGSCCKFD | 0.1122 | 18 | ABCPRED | 1 | ALLERGEN | Non-Toxin | -0.11 | 0.67 | -0.13 | 1 | 7.94 | 1850.46 | inside |
| 1010 | 1026 | QQLIRAAEIRASANLA | 0.2032 | 16 | ABCPRED | 1 | NON-ALLERGEN | Non-Toxin | -0.17 | 0.11 | 0.01 | 1 | 9.95 | 1725.21 | outside |
| 299 | 311 | TKCTLKSFTVEK | 0.2706 | 12 | ABCPRED | 1 | NON-ALLERGEN | Non-Toxin | -0.25 | -0.4 | 0.36 | 2 | 9.25 | 1384.82 | outside |
| 611 | 623 | LYQDVNCTEVPV | 0.5295 | 12 | ABCPRED,BCPREDS | 2 | NON-ALLERGEN | Non-Toxin | -0.06 | 0.11 | -0.3 | -2 | 3.67 | 1379.72 | outside |
| 413 | 425 | GQTGKIADYNYK | 1.7308 | 12 | ABCPRED | 1 | NON-ALLERGEN | Non-Toxin | -0.26 | -1.34 | 0.18 | 1 | 8.76 | 1357.66 | outside |
| 634 | 644 | RVYSTGSNVF | -0.1018 | 10 | ABCPRED | 1 | NON-ALLERGEN | Non-Toxin | -0.12 | -0.08 | -0.44 | 1 | 9.1 | 1129.37 | outside |
| 821 | 841 | LLFNKVTLADAGFIKQYGDC | 0.3957 | 20 | ABCPRED | 1 | NON-ALLERGEN | Non-Toxin | -0.01 | 0.36 | -0.3 | 0 | 6.27 | 2216.88 | outside |
| 249 | 262 | LTPGDSSSGWTAG | 0.495 | 13 | IEDB,literature | 2 | NON-ALLERGEN | Non-Toxin | -0.02 | -0.42 | -0.2 | -1 | 3.8 | 1235.45 | outside |
| 1134 | 1146 | NNTVYDPLQPEL | 0.444 | 12 | AAP,BCPREDS | 2 | ALLERGEN | Non-Toxin | -0.17 | -0.91 | -0.1 | -2 | 3.67 | 1402.7 | outside |
| 8 | 26 | LPLVSSQCVNLTTRTQLP | 1.0804 | 18 | ABCPRED | 1 | NON-ALLERGEN | Non-Toxin | -0.1 | 0.23 | -0.46 | 1 | 8.6 | 1970.59 | outside |
| 835 | 843 | KQYGDCLG | 0.4129 | 8 | IEDB | 1 | ALLERGEN | Non-Toxin | -0.2 | -0.84 | 0.14 | 0 | 6.15 | 883.1 | outside |
| 309 | 319 | EKGIYQTSNF | 0.4514 | 10 | ABCPRED | 1 | NON-ALLERGEN | Non-Toxin | -0.2 | -1.03 | -0.03 | 0 | 6.35 | 1186.43 | outside |
| 569 | 581 | IADTTDAVRDPQ | 0.4702 | 12 | ABCPRED | 1 | NON-ALLERGEN | Non-Toxin | -0.27 | -0.77 | 0.59 | -2 | 3.94 | 1301.53 | outside |
| 964 | 980 | KQLSSNFGAISSVLND | 0.4079 | 16 | ABCPRED | 1 | NON-ALLERGEN | Non-Toxin | -0.09 | -0.04 | -0.13 | 0 | 6.19 | 1680.08 | outside |
| 233 | 245 | INITRFQTLLAL | 0.4263 | 12 | ABCPRED | 1 | NON-ALLERGEN | Non-Toxin | 0.04 | 1.01 | -0.78 | 1 | 10.11 | 1402.89 | outside |
| 5 | 23 | LVLLPLVSSQCVNLTTRT | 1.1125 | 18 | ABCPRED | 1 | NON-ALLERGEN | Non-Toxin | 0.01 | 0.96 | -0.65 | 1 | 8.6 | 1957.64 | outside |

| | | | | | | | | | | | | | | | |
|---|---|---|---|---|---|---|---|---|---|---|---|---|---|---|---|
| 1020 | 1038 | ASANLAATKMSECVLGQS | 0.3745 | 18 | ABCPRED | 1 | ALLERGEN | Non-Toxin | -0.05 | 0.31 | -0.14 | 0 | 6.32 | 1781.28 | outside |
| 404 | 410 | GDEVRQ | 0.6701 | 6 | IEDB | 1 | ALLERGEN | Non-Toxin | -0.52 | -1.87 | 1.28 | -1 | 4.38 | 702.8 | outside |
| 775 | 789 | DKNTQEVFAQVKQI | 0.3493 | 14 | ABCPRED,FBCPred | 2 | NON-ALLERGEN | Non-Toxin | -0.27 | -0.86 | 0.33 | 0 | 6.42 | 1648.06 | outside |
| 89 | 99 | GVYFASTEKS | 0.8053 | 10 | ABCPRED | 1 | NON-ALLERGEN | Non-Toxin | -0.08 | -0.26 | -0.06 | 0 | 6.35 | 1088.31 | outside |
| 962 | 982 | LVKQLSSNFGAISSVLNDIL | 0.0834 | 20 | ABCPRED | 1 | NON-ALLERGEN | Non-Toxin | 0.04 | 0.79 | -0.45 | 0 | 6.19 | 2118.77 | outside |
| 567 | 583 | RDIADTTDAVRDPQTL | 0.3543 | 16 | ABCPRED | 1 | NON-ALLERGEN | Non-Toxin | -0.34 | -0.88 | 0.68 | -2 | 4.14 | 1787.13 | outside |
| 424 | 434 | KLPDDFTGCV | -0.5308 | 10 | ABCPRED | 1 | NON-ALLERGEN | Non-Toxin | -0.09 | -0.03 | 0.18 | -1 | 4.21 | 1094.38 | outside |
| 1245 | 1261 | KGCCSCGSCCKFDEDD | 0.2965 | 16 | ABCPRED | 1 | ALLERGEN | Non-Toxin | -0.27 | -0.56 | 0.69 | -2 | 4.23 | 1700.07 | inside |
| 824 | 836 | NKVTLADAGFIK | 0.7032 | 12 | ABCPRED | 1 | ALLERGEN | Non-Toxin | -0.06 | 0.25 | 0.02 | 1 | 8.94 | 1276.67 | outside |
| 545 | 561 | GLTGTGVLTESNKKFL | 0.659 | 16 | ABCPRED | 1 | NON-ALLERGEN | Non-Toxin | -0.07 | -0.03 | -0.07 | 1 | 8.94 | 1665.17 | outside |
| 1108 | 1126 | NFYEPQIITTDNTFVSGN | 0.3232 | 18 | ABCPRED | 1 | NON-ALLERGEN | Non-Toxin | -0.08 | -0.47 | -0.36 | -2 | 3.67 | 2060.47 | outside |
| 1108 | 1122 | NFYEPQIITTDNTF | 0.4308 | 14 | AAP | 1 | ALLERGEN | Non-Toxin | -0.09 | -0.56 | -0.39 | -2 | 3.67 | 1703.04 | outside |
| 740 | 756 | MYICGDSTECSNLLLQ | -0.1087 | 16 | ABCPRED | 1 | NON-ALLERGEN | Non-Toxin | -0.03 | 0.3 | -0.39 | -2 | 3.67 | 1790.29 | outside |
| 998 | 1014 | TGRLQSLQTYVTQQLI | -0.0444 | 16 | ABCPRED | 1 | ALLERGEN | Non-Toxin | -0.14 | -0.19 | -0.51 | 1 | 9.1 | 1849.38 | outside |
| 1009 | 1029 | TQQLIRAAEIRASANLAATK | 0.3191 | 20 | ABCPRED | 1 | NON-ALLERGEN | Non-Toxin | -0.19 | -0.09 | 0.09 | 2 | 10.84 | 2126.73 | outside |
| 248 | 260 | YLTPGDSSSGWT | 0.6069 | 12 | ABCPRED,literature | 2 | ALLERGEN | Non-Toxin | -0.06 | -0.67 | -0.37 | -1 | 3.8 | 1270.48 | outside |
| 908 | 928 | GIGVTQNVLYENQKLIANQF | 0.3693 | 20 | ABCPRED | 1 | NON-ALLERGEN | Non-Toxin | -0.05 | -0.08 | -0.43 | 0 | 6.35 | 2249.88 | outside |
| 865 | 883 | LTDEMIAQYTSALLAGTI | 0.2975 | 18 | ABCPRED | 1 | NON-ALLERGEN | Non-Toxin | 0.08 | 0.7 | -0.49 | -2 | 3.67 | 1911.47 | outside |
| 553 | 565 | TESNKKFLPFQQ | 0.7903 | 12 | ABCPRED,BCPREDS,ellipro | 3 | NON-ALLERGEN | Non-Toxin | -0.3 | -1.29 | 0.23 | 1 | 8.94 | 1466.83 | outside |
| 197 | 209 | IDGYFKIYSKHT | 0.4695 | 12 | ABCPRED | 1 | NON-ALLERGEN | Non-Toxin | -0.12 | -0.6 | -0.19 | 1.5 | 8.76 | 1471.85 | outside |
| 385 | 397 | TKLNDLCFTNVY | 2.0035 | 12 | ABCPRED | 1 | NON-ALLERGEN | Non-Toxin | -0.1 | 0 | -0.44 | 0 | 6.15 | 1430.81 | outside |
| 425 | 441 | LPDDFTGCVIAWNSNN | -0.0956 | 16 | ABCPRED | 1 | NON-ALLERGEN | Non-Toxin | -0.04 | -0.14 | -0.38 | -2 | 3.57 | 1766.14 | outside |
| 610 | 623 | VLYQDVNCTEVPV | 0.619 | 13 | BCEPRED | 1 | NON-ALLERGEN | Toxin | -0.01 | 0.42 | -0.39 | -2 | 3.67 | 1478.87 | outside |
| 234 | 246 | NITRFQTLLALH | 0.0939 | 12 | ABCPRED | 1 | NON-ALLERGEN | Non-Toxin | -0.06 | 0.37 | -0.67 | 1.5 | 10.11 | 1426.87 | outside |
| 573 | 583 | TDAVRDPQTL | 0.0334 | 10 | ABCPRED | 1 | ALLERGEN | Non-Toxin | -0.3 | -0.82 | 0.46 | -1 | 4.21 | 1115.34 | outside |
| 660 | 678 | YECDIPIGAGICASYQTQ | 0.7927 | 18 | ABCPRED | 1 | ALLERGEN | Non-Toxin | -0.01 | 0.09 | -0.37 | -2 | 3.67 | 1932.41 | outside |
| 1028 | 1039 | KMSECVLGQSK | 0.1391 | 11 | BCEPRED | 1 | NON-ALLERGEN | Non-Toxin | -0.23 | -0.4 | 0.38 | 1 | 8.54 | 1209.6 | outside |
| 636 | 652 | YSTGSNVFQTRAGCLI | 0.8063 | 16 | ABCPRED | 1 | NON-ALLERGEN | Non-Toxin | -0.06 | 0.19 | -0.51 | 1 | 8.57 | 1717.16 | outside |
| 296 | 314 | LSETKCTLKSFTVEKGIY | 0.428 | 18 | ABCPRED | 1 | NON-ALLERGEN | Non-Toxin | -0.14 | -0.14 | 0.09 | 1 | 8.48 | 2047.66 | outside |
| 125 | 135 | NVVIKVCEFQ | -0.1498 | 10 | IEDB,literature | 2 | NON-ALLERGEN | Non-Toxin | -0.01 | 0.8 | -0.34 | 0 | 6.32 | 1178.56 | outside |
| 106 | 124 | FGTTLDSKTQSLLIVNNA | 0.6645 | 18 | AAP | 1 | ALLERGEN | Non-Toxin | -0.05 | 0.15 | -0.32 | 0 | 6.19 | 1922.44 | outside |
| 366 | 386 | SVLYNSASFSTFKCYGVSPT | 0.5807 | 20 | FBCPred | 1 | ALLERGEN | Non-Toxin | -0.01 | 0.24 | -0.6 | 1 | 8.51 | 2158.68 | outside |
| 832 | 844 | GFIKQYGDCLGD | -0.3836 | 12 | BCPREDS | 1 | NON-ALLERGEN | Non-Toxin | -0.07 | -0.27 | -0.02 | -1 | 4.21 | 1315.64 | outside |
| 421 | 433 | YNYKLPDDFTGC | 0.0777 | 12 | ABCPRED,BCPREDS | 2 | NON-ALLERGEN | Non-Toxin | -0.17 | -0.88 | -0.09 | -1 | 4.21 | 1435.73 | outside |
| 1040 | 1049 | VDFCGKGYH | 0.375 | 9 | IEDB | 1 | ALLERGEN | Non-Toxin | -0.08 | -0.36 | -0.2 | 0.5 | 7.06 | 1025.27 | outside |
| 825 | 843 | KVTLADAGFIKQYGDCLG | 0.445 | 18 | ABCPRED | 1 | NON-ALLERGEN | Non-Toxin | -0.03 | 0.21 | -0.11 | 0 | 6.27 | 1899.46 | outside |
| 736 | 756 | VDCTMYICGDSTECSNLLLQ | -0.0759 | 20 | ABCPRED | 1 | ALLERGEN | Non-Toxin | -0.04 | 0.37 | -0.3 | -3 | 3.5 | 2208.81 | outside |
| 1111 | 1127 | EPQIITTDNTFVSGNC | 0.3844 | 16 | ABCPRED | 1 | NON-ALLERGEN | Non-Toxin | -0.09 | -0.24 | -0.18 | -2 | 3.67 | 1739.12 | outside |
| 44 | 54 | RSSVLHSTQD | 0.5404 | 10 | ABCPRED | 1 | NON-ALLERGEN | Non-Toxin | -0.35 | -0.98 | 0.29 | 0.5 | 7.1 | 1129.33 | outside |
| 312 | 322 | IYQTSNFRVQ | 0.6567 | 10 | ABCPRED | 1 | NON-ALLERGEN | Non-Toxin | -0.23 | -0.63 | -0.46 | 1 | 9.1 | 1255.54 | outside |
| 658 | 668 | NSYECDIPIG | 0.6533 | 10 | ABCPRED | 1 | NON-ALLERGEN | Non-Toxin | -0.06 | -0.31 | -0.04 | -2 | 3.67 | 1110.34 | outside |
| 603 | 623 | NTSNQVAVLYQDVNCTEVPV | 0.6145 | 20 | ABCPRED,AAP | 2 | NON-ALLERGEN | Non-Toxin | -0.09 | -0.02 | -0.33 | -2 | 3.67 | 2193.71 | outside |
| 1002 | 1018 | QSLQTYVTQQLIRAAE | -0.1462 | 16 | ABCPRED | 1 | NON-ALLERGEN | Non-Toxin | -0.18 | -0.35 | -0.24 | 0 | 6.35 | 1849.32 | outside |
| 407 | 420 | VRQIAPGQTGKIA | 1.2181 | 13 | literature | 1 | NON-ALLERGEN | Non-Toxin | -0.13 | -0.13 | -0.01 | 2 | 11.01 | 1338.77 | outside |
| 776 | 788 | KNTQEVFAQVKQ | 0.5382 | 12 | ABCPRED | 1 | NON-ALLERGEN | Non-Toxin | -0.31 | -1.08 | 0.28 | 1 | 8.94 | 1419.78 | outside |
| 612 | 628 | YQDVNCTEVPVAIHAD | 0.7832 | 16 | ABCPRED | 1 | NON-ALLERGEN | Non-Toxin | -0.07 | -0.07 | -0.13 | -2.5 | 4.03 | 1774.16 | outside |
| 383 | 393 | SPTKLNDLCF | 2.2412 | 10 | ABCPRED | 1 | ALLERGEN | Non-Toxin | -0.13 | -0.11 | -0.1 | 0 | 6.16 | 1137.45 | outside |
| 904 | 916 | YRFNGIGVTQNV | 1.1043 | 12 | ABCPRED,FBCPred | 2 | NON-ALLERGEN | Non-Toxin | -0.1 | -0.17 | -0.53 | 1 | 9.1 | 1367.71 | outside |
| 313 | 323 | YQTSNFRVQP | 1.1866 | 10 | ABCPRED | 1 | ALLERGEN | Non-Toxin | -0.31 | -1.24 | -0.28 | 1 | 9.1 | 1239.49 | outside |
| 838 | 850 | GDCLGDIAARDL | 0.5734 | 12 | ABCPRED | 1 | ALLERGEN | Non-Toxin | -0.11 | 0.2 | 0.38 | -2 | 3.94 | 1218.51 | outside |
| 354 | 366 | NRKRISNCVADY | 0.0942 | 12 | ABCPRED | 1 | ALLERGEN | Non-Toxin | -0.44 | -1.04 | 0.47 | 2 | 9.36 | 1438.78 | outside |
| 139 | 151 | PFLGVYYHKNNK | 0.8563 | 12 | ABCPRED | 1 | NON-ALLERGEN | Non-Toxin | -0.17 | -0.98 | -0.37 | 2.5 | 9.55 | 1479.88 | outside |
| 1001 | 1017 | LQSLQTYVTQQLIRAA | -0.1502 | 16 | ABCPRED | 1 | NON-ALLERGEN | Non-Toxin | -0.11 | 0.11 | -0.54 | 1 | 9.1 | 1833.37 | outside |
| 915 | 935 | VLYENQKLIANQFNSAIGKI | 0.0153 | 20 | ABCPRED | 1 | NON-ALLERGEN | Non-Toxin | -0.07 | 0.02 | -0.3 | 1 | 8.83 | 2263.94 | outside |
| 250 | 268 | TPGDSSSGWTAGAAAYYV | 0.2539 | 18 | ABCPRED | 1 | ALLERGEN | Non-Toxin | 0.03 | -0.12 | -0.47 | -1 | 3.8 | 1761.07 | outside |
| 654 | 672 | EHVNNSYECDIPIGAGIC | 0.8002 | 18 | ABCPRED | 1 | NON-ALLERGEN | Non-Toxin | -0.03 | -0.04 | -0.14 | -2.5 | 4.14 | 1934.39 | outside |
| 829 | 847 | ADAGFIKQYGDCLGDIAA | 0.1842 | 18 | BCPREDS,FBCPred | 2 | NON-ALLERGEN | Non-Toxin | 0.01 | 0.27 | -0.06 | -2 | 3.94 | 1828.28 | outside |
| 757 | 767 | GSFCTQLNRA | 0.8748 | 10 | ABCPRED | 1 | ALLERGEN | Non-Toxin | -0.19 | -0.25 | -0.25 | 1 | 8.6 | 1096.36 | outside |
| 80 | 98 | DNPVLPFNDGVYFASTEK | 0.3451 | 18 | ABCPRED | 1 | ALLERGEN | Non-Toxin | -0.1 | -0.46 | -0.02 | -2 | 4.03 | 2013.44 | outside |
| 158 | 170 | RVYSSANNCTFE | -0.2207 | 12 | ABCPRED | 1 | ALLERGEN | Non-Toxin | -0.24 | -0.61 | -0.1 | 0 | 6.32 | 1390.64 | outside |
| 886 | 904 | WTFGAGAALQIPFAMQMA | 0.748 | 18 | ABCPRED | 1 | ALLERGEN | Non-Toxin | 0.18 | 0.87 | -0.95 | 0 | 5.88 | 1911.53 | outside |
| 806 | 817 | LPDPSKPSKRS | 0.5972 | 11 | IEDB | 1 | NON-ALLERGEN | Non-Toxin | -0.47 | -1.75 | 1.01 | 2 | 10.01 | 1211.52 | outside |
| 634 | 645 | RVYSTGSNVFQ | -0.1 | 11 | IEDB,literature | 2 | NON-ALLERGEN | Non-Toxin | -0.17 | -0.39 | -0.38 | 1 | 9.1 | 1257.52 | outside |
| 1187 | 1205 | NEVAKNLNESLIDLQELG | 0.5337 | 18 | ABCPRED | 1 | ALLERGEN | Non-Toxin | -0.15 | -0.41 | 0.28 | -3 | 4.01 | 1999.49 | outside |
| 536 | 556 | NKCVNFNFNGLTGTGVLTES | 1.0131 | 20 | ABCPRED | 1 | ALLERGEN | Non-Toxin | -0.06 | -0.07 | -0.34 | 0 | 6.32 | 2115.65 | outside |
| 652 | 670 | GAEHVNNSYECDIPIGAG | 0.9847 | 18 | ABCPRED,AAP,BCPREDS | 3 | NON-ALLERGEN | Non-Toxin | -0.05 | -0.35 | -0.01 | -2.5 | 4.14 | 1846.22 | outside |
| 381 | 391 | GVSPTKLNDL | 1.6525 | 10 | ABCPRED | 1 | ALLERGEN | Non-Toxin | -0.12 | -0.26 | 0.1 | 0 | 6.19 | 1043.33 | outside |
| 638 | 650 | TGSNVFQTRAGC | 0.8604 | 12 | ABCPRED | 1 | ALLERGEN | Non-Toxin | -0.16 | -0.27 | -0.22 | 1 | 8.6 | 1240.52 | outside |
| 1190 | 1204 | AKNLNESLIDLQEL | 0.8403 | 14 | FBCPred | 1 | NON-ALLERGEN | Non-Toxin | -0.16 | -0.3 | 0.24 | -2 | 4.14 | 1600.02 | outside |
| 779 | 791 | QEVFAQVKQIYK | -0.0944 | 12 | FBCPred | 1 | NON-ALLERGEN | Non-Toxin | -0.18 | -0.47 | -0.04 | 1 | 8.83 | 1480.91 | outside |
| 1108 | 1128 | NFYEPQIITTDNTFVSGNCD | 0.2402 | 20 | BCPREDS | 1 | NON-ALLERGEN | Non-Toxin | -0.1 | -0.47 | -0.23 | -3 | 3.5 | 2278.72 | outside |
| 563 | 569 | QQFGRD | -0.55 | 6 | COVIDep,literature | 2 | ALLERGEN | Non-Toxin | -0.52 | -2.1 | 0.65 | 0 | 6.19 | 749.86 | outside |
| 826 | 844 | VTLADAGFIKQYGDCLGD | 0.3176 | 18 | ABCPRED | 1 | NON-ALLERGEN | Non-Toxin | -0.01 | 0.23 | -0.11 | -2 | 3.94 | 1886.37 | outside |
| 738 | 752 | CTMYICGDSTECSN | -0.5258 | 14 | ABCPRED,AAP | 2 | NON-ALLERGEN | Non-Toxin | -0.11 | -0.09 | -0.17 | -2 | 3.67 | 1526.87 | outside |
| 736 | 752 | VDCTMYICGDSTECSN | -0.1722 | 16 | ABCPRED | 1 | NON-ALLERGEN | Non-Toxin | -0.11 | -0.04 | -0.06 | -3 | 3.5 | 1741.12 | outside |
| 639 | 655 | GSNVFQTRAGCLIGAE | 0.9074 | 16 | ABCPRED | 1 | NON-ALLERGEN | Non-Toxin | -0.04 | 0.23 | -0.21 | 0 | 6.32 | 1623.05 | outside |
| 674 | 684 | YQTQTNSPRR | -0.2647 | 10 | IEDB | 1 | NON-ALLERGEN | Non-Toxin | -0.62 | -2.46 | 0.38 | 2 | 10.84 | 1250.47 | outside |

| | | | | | | | | | | | | | | | |
|---|---|---|---|---|---|---|---|---|---|---|---|---|---|---|---|
| 922 | 938 | LIANQFNSAIGKIQDS | 0.1678 | 16 | ABCPRED | 1 | NON-ALLERGEN | Non-Toxin | -0.06 | 0.02 | -0.21 | 0 | 6.19 | 1719.17 | outside |
| 735 | 751 | SVDCTMYICGDSTECS | -0.0934 | 16 | ABCPRED | 1 | NON-ALLERGEN | Non-Toxin | -0.09 | 0.13 | -0.05 | -3 | 3.5 | 1714.09 | outside |
| 403 | 413 | RGDEVRQIAP | 0.1209 | 10 | ABCPRED | 1 | NON-ALLERGEN | Non-Toxin | -0.39 | -1.1 | 0.84 | 0 | 6.42 | 1140.4 | outside |
| 158 | 168 | RVYSSANNCT | -0.4227 | 10 | ABCPRED | 1 | NON-ALLERGEN | Non-Toxin | -0.29 | -0.66 | -0.17 | 1 | 8.57 | 1114.32 | outside |
| 386 | 396 | KLNDLCFTNV | 2.6927 | 10 | ABCPRED,literature | 2 | NON-ALLERGEN | Non-Toxin | -0.1 | 0.2 | -0.26 | 0 | 6.16 | 1166.5 | outside |
| 685 | 697 | RSVASQSIIAYT | 0.1222 | 12 | ABCPRED | 1 | NON-ALLERGEN | Non-Toxin | -0.07 | 0.37 | -0.39 | 1 | 9.1 | 1295.62 | outside |
| 835 | 847 | KQYGDCLGDIAA | 0.2179 | 12 | FBCPred | 1 | ALLERGEN | Non-Toxin | -0.09 | -0.18 | 0.11 | -1 | 4.21 | 1253.56 | outside |
| 119 | 137 | IVNNATNVVIKVCEFQFC | -0.1975 | 18 | FBCPred | 1 | ALLERGEN | Non-Toxin | 0.04 | 0.89 | -0.59 | 0 | 6.29 | 2041.68 | outside |
| 195 | 205 | KNIDGYFKIY | -0.3249 | 10 | ABCPRED | 1 | ALLERGEN | Non-Toxin | -0.13 | -0.6 | -0.15 | 1 | 8.76 | 1260.6 | outside |
| 826 | 842 | VTLADAGFIKQYGDCL | 0.4712 | 16 | ABCPRED | 1 | NON-ALLERGEN | Non-Toxin | 0.03 | 0.5 | -0.31 | -1 | 4.21 | 1714.2 | outside |
| 279 | 297 | YNENGTITDAVDCALDPL | 0.5757 | 18 | ABCPRED | 1 | NON-ALLERGEN | Non-Toxin | -0.08 | -0.18 | 0.02 | -4 | 3.38 | 1924.32 | outside |
| 63 | 70 | KNLNSSRV | 0.3752 | 8 | literature | 1 | ALLERGEN | Non-Toxin | -0.45 | -1.12 | 0.46 | 2 | 11.01 | 917.14 | outside |
| 322 | 332 | PTESIVRFPN | -0.6977 | 10 | ABCPRED | 1 | ALLERGEN | Non-Toxin | -0.17 | -0.47 | 0.03 | 0 | 6.36 | 1159.44 | outside |
| 1250 | 1264 | CGSCCKFDEDDSEP | 0.4214 | 14 | FBCPred | 1 | NON-ALLERGEN | Non-Toxin | -0.3 | -1.05 | 0.94 | -4 | 3.77 | 1534.77 | inside |
| 656 | 674 | VNNSYECDIPIGAGICAS | 0.7019 | 18 | ABCPRED | 1 | NON-ALLERGEN | Non-Toxin | 0.02 | 0.39 | -0.29 | -2 | 3.67 | 1826.28 | outside |
| 370 | 388 | NSASFSTFKCYGVSPTKL | 0.8108 | 18 | ABCPRED,AAP | 2 | NON-ALLERGEN | Non-Toxin | -0.09 | -0.07 | -0.31 | 2 | 9.25 | 1937.44 | outside |
| 609 | 623 | AVLYQDVNCTEVPV | 0.6208 | 14 | AAP | 1 | NON-ALLERGEN | Non-Toxin | 0.01 | 0.52 | -0.4 | -2 | 3.67 | 1549.96 | outside |
| 327 | 337 | VRFPNITNLC | 1.1924 | 10 | ABCPRED | 1 | NON-ALLERGEN | Non-Toxin | -0.08 | 0.4 | -0.56 | 1 | 8.6 | 1176.54 | outside |
| 786 | 795 | KQIYKTPPI | 0.2705 | 9 | IEDB | 1 | ALLERGEN | Non-Toxin | -0.19 | -0.83 | -0.01 | 2 | 9.72 | 1087.46 | outside |
| 368 | 386 | LYNSASFSTFKCYGVSPT | 0.6517 | 18 | FBCPred | 1 | ALLERGEN | Non-Toxin | -0.02 | 0.07 | -0.6 | 1 | 8.51 | 1972.44 | outside |
| 926 | 945 | QFNSAIGKIQDSLSSTASA | 0.3983 | 19 | IEDB | 1 | NON-ALLERGEN | Non-Toxin | -0.1 | -0.11 | -0.09 | 0 | 6.19 | 1925.35 | outside |
| 192 | 210 | FVFKNIDGYFKIYSKHTP | 0.4273 | 18 | ABCPRED | 1 | NON-ALLERGEN | Toxin | -0.09 | -0.36 | -0.31 | 2.5 | 9.43 | 2204.82 | outside |
| 251 | 269 | PGDSSSGWTAGAAAYYVG | 0.2668 | 18 | ABCPRED | 1 | NON-ALLERGEN | Non-Toxin | 0.05 | -0.11 | -0.44 | -1 | 3.8 | 1717.02 | outside |
| 832 | 848 | GFIKQYGDCLGDIAAR | 0.1391 | 16 | ABCPRED | 1 | ALLERGEN | Non-Toxin | -0.08 | 0.02 | 0 | 0 | 6.28 | 1727.2 | outside |
| 424 | 431 | KLPDDFT | -0.051 | 7 | IEDB | 1 | ALLERGEN | Non-Toxin | -0.24 | -0.94 | 0.61 | -1 | 4.21 | 835.01 | outside |
| 737 | 755 | DCTMYICGDSTECSNLLL | -0.3363 | 18 | ABCPRED | 1 | ALLERGEN | Non-Toxin | -0.04 | 0.37 | -0.27 | -3 | 3.5 | 1981.51 | outside |
| 295 | 303 | PLSETKCT | 1.2573 | 8 | IEDB | 1 | NON-ALLERGEN | Non-Toxin | -0.23 | -0.61 | 0.34 | 0 | 6.32 | 878.11 | outside |
| 779 | 789 | QEVFAQVKQI | 0.1829 | 10 | ABCPRED | 1 | NON-ALLERGEN | Non-Toxin | -0.11 | -0.04 | -0.12 | 0 | 6.35 | 1189.53 | outside |
| 157 | 169 | FRVYSSANNCTF | -0.1749 | 12 | ABCPRED | 1 | ALLERGEN | Non-Toxin | -0.14 | -0.08 | -0.56 | 1 | 8.57 | 1408.7 | outside |
| 274 | 284 | TFLLKYNENG | 0.426 | 10 | ABCPRED | 1 | ALLERGEN | Non-Toxin | -0.13 | -0.64 | -0.24 | 0 | 6.35 | 1198.49 | outside |
| 1109 | 1127 | FYEPQIITTDNTFVSGNC | 0.2738 | 18 | ABCPRED,AAP | 2 | ALLERGEN | Non-Toxin | -0.04 | -0.13 | -0.43 | -2 | 3.67 | 2049.5 | outside |
| 239 | 251 | QTLLALHRSYLT | 0.3939 | 12 | ABCPRED | 1 | NON-ALLERGEN | Non-Toxin | -0.09 | 0.19 | -0.65 | 1.5 | 9.1 | 1415.84 | outside |
| 822 | 840 | LFNKVTLADAGFIKQYGD | 0.2555 | 18 | ABCPRED | 1 | NON-ALLERGEN | Toxin | -0.04 | 0.05 | -0.18 | 0 | 6.3 | 2000.55 | outside |
| 717 | 733 | NFTISVTTEILPVSMT | 1.3573 | 16 | ABCPRED | 1 | NON-ALLERGEN | Non-Toxin | 0.09 | 0.81 | -0.63 | -1 | 4 | 1753.28 | outside |
| 719 | 737 | TISVTTEILPVSMTKTSV | 1.1598 | 18 | ABCPRED | 1 | NON-ALLERGEN | Non-Toxin | 0.02 | 0.69 | -0.35 | 0 | 6.35 | 1907.52 | outside |
| 906 | 926 | FNGIGVTQNVLYENQKLIAN | 0.3916 | 20 | ABCPRED | 1 | NON-ALLERGEN | Non-Toxin | -0.05 | -0.08 | -0.43 | 0 | 6.35 | 2235.85 | outside |
| 1188 | 1205 | EVAKNLNESLIDLQELG | 0.6224 | 17 | COVIDep,literature | 2 | NON-ALLERGEN | Non-Toxin | -0.12 | -0.23 | 0.29 | -3 | 4.01 | 1885.37 | outside |
| 864 | 882 | LLTDEMIAQYTSALLAGT | 0.1617 | 18 | ABCPRED | 1 | NON-ALLERGEN | Non-Toxin | 0.07 | 0.66 | -0.49 | -2 | 3.67 | 1911.47 | outside |
| 386 | 398 | KLNDLCFTNVYA | 1.9268 | 12 | literature | 1 | NON-ALLERGEN | Non-Toxin | -0.06 | 0.21 | -0.45 | 0 | 6.15 | 1400.78 | outside |
| 861 | 873 | LPPLLTDEMIAQ | 0.3701 | 12 | ABCPRED | 1 | NON-ALLERGEN | Non-Toxin | 0.04 | 0.43 | -0.27 | -2 | 3.67 | 1340.78 | outside |
| 368 | 388 | LYNSASFSTFKCYGVSPTKL | 0.6975 | 20 | ABCPRED | 1 | NON-ALLERGEN | Non-Toxin | -0.05 | 0.06 | -0.48 | 2 | 9.17 | 2213.81 | outside |
| 415 | 427 | TGKIADYNYKLP | 1.1956 | 12 | ABCPRED | 1 | ALLERGEN | Non-Toxin | -0.18 | -0.83 | 0.01 | 1 | 8.76 | 1382.75 | outside |
| 319 | 331 | RVQPTESIVRFP | -0.0823 | 12 | ABCPRED | 1 | NON-ALLERGEN | Non-Toxin | -0.25 | -0.42 | 0.15 | 1 | 9.95 | 1428.82 | outside |
| 248 | 268 | YLTPGDSSSGWTAGAAAYYV | 0.4852 | 20 | ABCPRED | 1 | ALLERGEN | Non-Toxin | 0.05 | 0.01 | -0.62 | -1 | 3.8 | 2037.44 | outside |
| 42 | 52 | VFRSSVLHST | -0.0642 | 10 | ABCPRED | 1 | NON-ALLERGEN | Non-Toxin | -0.09 | 0.42 | -0.43 | 1.5 | 10.11 | 1132.42 | outside |
| 862 | 874 | PPLLTDEMIAQY | 0.2239 | 12 | ABCPRED | 1 | NON-ALLERGEN | Non-Toxin | 0 | 0.01 | -0.31 | -2 | 3.67 | 1390.79 | outside |
| 341 | 353 | VFNATRFASVYA | 0.1817 | 12 | ABCPRED | 1 | NON-ALLERGEN | Non-Toxin | 0.02 | 0.72 | -0.72 | 1 | 9.1 | 1345.67 | outside |
| 90 | 100 | VYFASTEKSN | 1.0231 | 10 | literature | 1 | ALLERGEN | Non-Toxin | -0.16 | -0.57 | -0.04 | 0 | 6.35 | 1145.36 | outside |
| 1128 | 1148 | VVIGIVNNTVYDPLQPELDS | 0.655 | 20 | ABCPRED | 1 | NON-ALLERGEN | Non-Toxin | 0.01 | 0.3 | -0.3 | -3 | 3.5 | 2185.77 | outside |
| 864 | 876 | LLTDEMIAQYTS | -0.0323 | 12 | ABCPRED | 1 | NON-ALLERGEN | Non-Toxin | -0.03 | 0.15 | -0.32 | -2 | 3.67 | 1384.74 | outside |
| 552 | 561 | LTESNKKFL | 0.3167 | 9 | BCEPRED | 1 | NON-ALLERGEN | Non-Toxin | -0.25 | -0.66 | 0.33 | 1 | 8.94 | 1079.39 | outside |
| 611 | 629 | LYQDVNCTEVPVAIHADQ | 0.4111 | 18 | ABCPRED | 1 | NON-ALLERGEN | Non-Toxin | -0.07 | -0.04 | -0.21 | -2.5 | 4.03 | 2015.49 | outside |
| 994 | 1010 | DRLITGRLQSLQTYVT | -0.3647 | 16 | ABCPRED | 1 | NON-ALLERGEN | Non-Toxin | -0.21 | -0.25 | -0.16 | 1 | 9.1 | 1864.38 | outside |
| 786 | 796 | KQIYKTPPIK | -0.0905 | 10 | ABCPRED | 1 | ALLERGEN | Non-Toxin | -0.28 | -1.14 | 0.29 | 3 | 10.01 | 1215.65 | outside |
| 1157 | 1168 | KNHTSPDVDLG | 1.4039 | 11 | IEDB | 1 | NON-ALLERGEN | Non-Toxin | -0.26 | -1.19 | 0.48 | -0.5 | 5.22 | 1182.41 | outside |
| 661 | 677 | ECDIPIGAGICASYQT | 0.7167 | 16 | ABCPRED | 1 | NON-ALLERGEN | Toxin | 0.04 | 0.4 | -0.29 | -2 | 3.67 | 1641.07 | outside |
| 14 | 28 | QCVNLTTRTQLPPA | 1.4983 | 14 | BCPREDS | 1 | ALLERGEN | Non-Toxin | -0.18 | -0.3 | -0.3 | 1 | 8.6 | 1541.99 | outside |
| 554 | 566 | ESNKKFLPFQQF | 0.4277 | 12 | ABCPRED | 1 | NON-ALLERGEN | Non-Toxin | -0.23 | -1 | 0.05 | 1 | 8.94 | 1512.9 | outside |
| 831 | 845 | AGFIKQYGDCLGDI | -0.0812 | 14 | ABCPRED,BCPREDS,FBCPred | 3 | ALLERGEN | Non-Toxin | 0.01 | 0.21 | -0.18 | -1 | 4.21 | 1499.91 | outside |
| 740 | 752 | MYICGDSTECSN | -0.2368 | 12 | ABCPRED | 1 | NON-ALLERGEN | Non-Toxin | -0.12 | -0.26 | -0.08 | -2 | 3.67 | 1322.6 | outside |
| 215 | 229 | DLPQGFSALEPLVD | 0.3503 | 14 | FBCPred | 1 | ALLERGEN | Non-Toxin | 0 | 0.13 | -0.03 | -3 | 3.5 | 1500.87 | outside |
| 371 | 387 | SASFSTFKCYGVSPTK | 0.908 | 16 | ABCPRED | 1 | NON-ALLERGEN | Non-Toxin | -0.09 | -0.1 | -0.24 | 2 | 9.25 | 1710.14 | outside |
| 653 | 667 | AEHVNNSYECDIPI | 0.6687 | 14 | BCPREDS,FBCPred | 2 | NON-ALLERGEN | Non-Toxin | -0.12 | -0.49 | 0.02 | -2.5 | 4.14 | 1603.92 | outside |
| 655 | 667 | HVNNSYECDIPI | 0.6752 | 12 | FBCPred,IEDB | 2 | NON-ALLERGEN | Non-Toxin | -0.11 | -0.43 | -0.18 | -1.5 | 4.36 | 1403.7 | outside |
| 822 | 834 | LFNKVTLADAGF | 0.845 | 12 | ABCPRED | 1 | NON-ALLERGEN | Non-Toxin | 0.07 | 0.75 | -0.44 | 0 | 6.19 | 1295.67 | outside |
| 727 | 741 | LPVSMTKTSVDCTM | 1.4435 | 14 | BCPREDS | 1 | NON-ALLERGEN | Non-Toxin | -0.06 | 0.41 | -0.21 | 0 | 6.16 | 1513.01 | outside |
| 201 | 213 | FKIYSKHTPINL | 1.1557 | 12 | ABCPRED | 1 | ALLERGEN | Non-Toxin | -0.09 | -0.28 | -0.38 | 2.5 | 9.72 | 1460.92 | outside |
| 297 | 313 | SETKCTLKSFTVEKGI | 0.4345 | 16 | ABCPRED | 1 | NON-ALLERGEN | Non-Toxin | -0.19 | -0.31 | 0.36 | 1 | 8.51 | 1771.29 | outside |
| 437 | 453 | NSNNLDSKVGGNYNYL | 0.7804 | 16 | ABCPRED | 1 | NON-ALLERGEN | Non-Toxin | -0.22 | -1.13 | -0.13 | 0 | 6.18 | 1772.1 | outside |
| 344 | 354 | ATRFASVYAW | -0.1278 | 10 | ABCPRED | 1 | NON-ALLERGEN | Non-Toxin | 0.01 | 0.42 | -0.83 | 1 | 9.1 | 1171.44 | outside |
| 860 | 872 | VLPPLLTDEMIA | 0.3229 | 12 | ABCPRED | 1 | ALLERGEN | Non-Toxin | 0.14 | 1.08 | -0.41 | -2 | 3.67 | 1311.78 | outside |
| 1157 | 1175 | KNHTSPDVDLGDISGINA | 1.0631 | 18 | ABCPRED | 1 | ALLERGEN | Non-Toxin | -0.14 | -0.58 | 0.26 | -1.5 | 4.42 | 1853.24 | outside |
| 94 | 104 | STEKSNIIRG | -0.3168 | 10 | ABCPRED | 1 | NON-ALLERGEN | Non-Toxin | -0.32 | -0.91 | 0.58 | 1 | 9.1 | 1104.37 | outside |
| 145 | 155 | YHKNNKSWME | 0.1657 | 10 | ABCPRED | 1 | NON-ALLERGEN | Non-Toxin | -0.41 | -2.26 | 0.22 | 1.5 | 8.84 | 1336.63 | outside |
| 386 | 397 | KLNDLCFTNVY | 2.2239 | 11 | literature | 1 | NON-ALLERGEN | Non-Toxin | -0.09 | 0.06 | -0.45 | 0 | 6.15 | 1329.69 | outside |
| 217 | 235 | PQGFSALEPLVDLPIGIN | 0.749 | 18 | ABCPRED | 1 | NON-ALLERGEN | Non-Toxin | 0.09 | 0.49 | -0.38 | -2 | 3.67 | 1880.45 | outside |
| 544 | 562 | NGLTGTGVLTESNKKFLP | 0.5984 | 18 | ABCPRED | 1 | NON-ALLERGEN | Non-Toxin | -0.1 | -0.31 | -0.05 | 1 | 8.94 | 1876.42 | outside |

| | | | | | | | | | | | | | | | |
|---|---|---|---|---|---|---|---|---|---|---|---|---|---|---|---|
| 218 | 230 | QGFSALEPLVDL | 0.4856 | 12 | ABCPRED | 1 | NON-ALLERGEN | Non-Toxin | 0.07 | 0.57 | -0.28 | -2 | 3.67 | 1288.64 | outside |
| 1013 | 1027 | IRAAEIRASANLAA | 0.5209 | 14 | AAP | 1 | ALLERGEN | Toxin | -0.11 | 0.49 | 0.08 | 1 | 9.95 | 1426.82 | outside |
| 778 | 790 | TQEVFAQVKQIY | 0.2454 | 12 | ABCPRED | 1 | NON-ALLERGEN | Non-Toxin | -0.11 | -0.2 | -0.33 | 0 | 6.35 | 1453.84 | outside |
| 830 | 842 | DAGFIKQYGDCL | 0.2155 | 12 | ABCPRED | 1 | ALLERGEN | Non-Toxin | -0.06 | -0.09 | -0.06 | -1 | 4.21 | 1329.66 | outside |
| 921 | 939 | KLIANQFNSAIGKIQDSL | -0.1166 | 18 | ABCPRED | 1 | NON-ALLERGEN | Non-Toxin | -0.09 | 0.01 | -0.12 | 1 | 8.94 | 1960.54 | outside |
| 908 | 924 | GIGVTQNVLYENQKLI | 0.6869 | 16 | ABCPRED | 1 | NON-ALLERGEN | Non-Toxin | -0.04 | 0.05 | -0.38 | 0 | 6.35 | 1789.33 | outside |
| 783 | 795 | AQVKQIYKTPPI | 0.5488 | 12 | ABCPRED | 1 | ALLERGEN | Non-Toxin | -0.14 | -0.42 | -0.16 | 2 | 9.72 | 1385.85 | outside |
| 125 | 137 | NVVIKVCEFQFC | -0.4813 | 12 | ABCPRED | 1 | NON-ALLERGEN | Non-Toxin | 0.05 | 1.11 | -0.57 | 0 | 6.29 | 1428.9 | outside |
| 847 | 854 | RDLICAQ | 1.1443 | 7 | IEDB,literature | 2 | ALLERGEN | Non-Toxin | -0.23 | 0.16 | 0.16 | 0 | 6.16 | 818.05 | outside |
| 323 | 335 | TESIVRFPNITN | 0.0758 | 12 | FBCPred | 1 | NON-ALLERGEN | Non-Toxin | -0.14 | -0.23 | -0.14 | 0 | 6.36 | 1390.73 | outside |
| 550 | 562 | GVLTESNKKFLP | 0.6053 | 12 | ABCPRED | 1 | NON-ALLERGEN | Non-Toxin | -0.13 | -0.31 | 0.13 | 1 | 8.94 | 1332.74 | outside |
| 1012 | 1028 | LIRAAEIRASANLAAT | 0.4496 | 16 | ABCPRED | 1 | ALLERGEN | Non-Toxin | -0.07 | 0.62 | -0.07 | 1 | 9.95 | 1641.12 | outside |
| 379 | 389 | CYGVSPTKLN | 1.5759 | 10 | ABCPRED | 1 | NON-ALLERGEN | Non-Toxin | -0.1 | -0.17 | -0.35 | 1 | 8.54 | 1081.39 | outside |
| 219 | 235 | GFSALEPLVDLPIGIN | 0.8838 | 16 | ABCPRED | 1 | NON-ALLERGEN | Non-Toxin | 0.15 | 0.87 | -0.44 | -2 | 3.67 | 1655.17 | outside |
| 403 | 409 | RGDEVR | -2.2527 | 6 | IEDB | 1 | ALLERGEN | Non-Toxin | -0.69 | -2.03 | 1.75 | 0 | 6.42 | 730.85 | outside |
| 1008 | 1026 | VTQQLIRAAEIRASANLA | 0.2378 | 18 | ABCPRED | 1 | NON-ALLERGEN | Non-Toxin | -0.13 | 0.29 | -0.09 | 1 | 9.95 | 1925.48 | outside |
| 294 | 306 | DPLSETKCTLKS | 0.9204 | 12 | FBCPred,IEDB | 2 | NON-ALLERGEN | Non-Toxin | -0.28 | -0.77 | 0.6 | 0 | 6.38 | 1321.67 | outside |
| 735 | 755 | SVDCTMYICGDSTECSNLLL | -0.019 | 20 | ABCPRED | 1 | ALLERGEN | Non-Toxin | -0.02 | 0.5 | -0.3 | -3 | 3.5 | 2167.75 | outside |
| 1137 | 1149 | VYDPLQPELDSF | 0.0903 | 12 | IEDB | 1 | ALLERGEN | Non-Toxin | -0.08 | -0.39 | -0.03 | -3 | 3.5 | 1422.72 | outside |
| 249 | 269 | LTPGDSSSGWTAGAAAYYVG | 0.5229 | 20 | ABCPRED | 1 | NON-ALLERGEN | Non-Toxin | 0.06 | 0.06 | -0.51 | -1 | 3.8 | 1931.32 | outside |
| 376 | 388 | TFKCYGVSPTKL | 1.4293 | 12 | ABCPRED | 1 | NON-ALLERGEN | Non-Toxin | -0.08 | 0 | -0.3 | 2 | 9.25 | 1343.77 | outside |
| 883 | 903 | TSGWTFGAGAALQIPFAMQM | 0.462 | 20 | ABCPRED | 1 | NON-ALLERGEN | Non-Toxin | 0.14 | 0.6 | -0.84 | 0 | 5.88 | 2085.72 | outside |
| 581 | 599 | TLEILDITPCSFGGVSVI | 1.4847 | 18 | ABCPRED | 1 | NON-ALLERGEN | Non-Toxin | 0.17 | 1.24 | -0.54 | -2 | 3.67 | 1864.46 | outside |
| 1011 | 1029 | QLIRAAEIRASANLAATK | 0.3527 | 18 | ABCPRED | 1 | NON-ALLERGEN | Non-Toxin | -0.17 | 0.14 | 0.12 | 2 | 10.84 | 1897.46 | outside |
| 1246 | 1264 | GCCSCGSCCKFDEDDSEP | 0.1272 | 18 | ABCPRED,FBCPred | 2 | NON-ALLERGEN | Non-Toxin | -0.23 | -0.61 | 0.63 | -4 | 3.77 | 1885.23 | inside |
| 18 | 28 | LTTRTQLPPA | 1.3398 | 10 | ABCPRED | 1 | NON-ALLERGEN | Non-Toxin | -0.18 | -0.39 | -0.21 | 1 | 10.11 | 1097.42 | outside |
| 719 | 735 | TISVTTEILPVSMTKT | 1.2262 | 16 | ABCPRED | 1 | NON-ALLERGEN | Non-Toxin | 0.01 | 0.56 | -0.32 | 0 | 6.35 | 1721.28 | outside |
| 1009 | 1025 | TQQLIRAAEIRASANL | 0.1489 | 16 | ABCPRED | 1 | NON-ALLERGEN | Non-Toxin | -0.19 | -0.04 | 0.02 | 1 | 9.95 | 1755.24 | outside |
| 311 | 323 | GIYQTSNFRVQP | 0.9986 | 12 | ABCPRED | 1 | NON-ALLERGEN | Non-Toxin | -0.19 | -0.69 | -0.38 | 1 | 9.1 | 1409.74 | outside |
| 634 | 650 | RVYSTGSNVFQTRAGC | 0.5772 | 16 | ABCPRED | 1 | NON-ALLERGEN | Non-Toxin | -0.21 | -0.35 | -0.19 | 2 | 9.55 | 1746.15 | outside |
| 362 | 380 | VADYSVLYNSASFSTFKC | 0.1972 | 18 | ABCPRED | 1 | ALLERGEN | Non-Toxin | -0.02 | 0.36 | -0.52 | 0 | 6.15 | 2002.46 | outside |
| 838 | 854 | GDCLGDIAARDLICAQ | 0.6192 | 16 | ABCPRED | 1 | NON-ALLERGEN | Non-Toxin | -0.06 | 0.48 | 0.09 | -2 | 3.94 | 1634.08 | outside |
| 231 | 249 | IGINITRFQTLLALHRSY | 0.7573 | 18 | ABCPRED | 1 | NON-ALLERGEN | Non-Toxin | -0.06 | 0.36 | -0.59 | 2.5 | 10.84 | 2116.78 | outside |
| 927 | 943 | FNSAIGKIQDSLSSTA | 0.3375 | 16 | ABCPRED | 1 | NON-ALLERGEN | Non-Toxin | -0.07 | 0.03 | -0.11 | 0 | 6.19 | 1639.02 | outside |
| 582 | 598 | LEILDITPCSFGGVSV | 1.7357 | 16 | ABCPRED | 1 | ALLERGEN | Non-Toxin | 0.15 | 1.16 | -0.47 | -2 | 3.67 | 1650.16 | outside |
| 909 | 929 | IGVTQNVLYENQKLIANQFN | 0.3425 | 20 | ABCPRED | 1 | ALLERGEN | Non-Toxin | -0.09 | -0.23 | -0.42 | 0 | 6.35 | 2306.93 | outside |
| 1134 | 1150 | NNTVYDPLQPELDSFK | 0.0462 | 16 | ABCPRED | 1 | NON-ALLERGEN | Non-Toxin | -0.22 | -1.02 | 0.16 | -2 | 4.03 | 1880.27 | outside |
| 1156 | 1164 | FKNHTSPD | 0.5003 | 8 | BCEPRED | 1 | NON-ALLERGEN | Non-Toxin | -0.34 | -1.8 | 0.39 | 0.5 | 7.09 | 945.1 | outside |
| 766 | 778 | ALTGIAVEQDKN | 0.9459 | 12 | ABCPRED | 1 | NON-ALLERGEN | Non-Toxin | -0.12 | -0.24 | 0.24 | -1 | 4.38 | 1258.57 | outside |
| 828 | 843 | LADAGFIKQYGDCLG | 0.2071 | 15 | IEDB,literature | 2 | NON-ALLERGEN | Non-Toxin | 0.01 | 0.27 | -0.2 | -1 | 4.21 | 1571 | outside |
| 924 | 938 | ANQFNSAIGKIQDS | 0.1577 | 14 | ABCPRED,AAP | 2 | NON-ALLERGEN | Non-Toxin | -0.16 | -0.57 | 0.02 | 0 | 6.19 | 1492.81 | outside |
| 540 | 560 | NFNFNGLTGTGVLTESNKKF | 0.9432 | 20 | ABCPRED | 1 | NON-ALLERGEN | Non-Toxin | -0.11 | -0.46 | -0.19 | 1 | 8.94 | 2188.73 | outside |
| 192 | 204 | FVFKNIDGYFKI | 0.2353 | 12 | ABCPRED | 1 | NON-ALLERGEN | Non-Toxin | 0.04 | 0.42 | -0.47 | 1 | 8.83 | 1490.94 | outside |
| 441 | 451 | LDSKVGGNYN | 0.91 | 10 | ABCPRED | 1 | ALLERGEN | Non-Toxin | -0.2 | -0.93 | 0.11 | 0 | 6.18 | 1066.28 | outside |
| 9 | 23 | PLVSSQCVNLTTRT | 1.2427 | 14 | ABCPRED,FBCPred | 2 | NON-ALLERGEN | Non-Toxin | -0.15 | 0.12 | -0.34 | 1 | 8.6 | 1518.95 | outside |
| 118 | 138 | LIVNNATNVVIKVCEFQFCN | -0.1541 | 20 | ABCPRED | 1 | NON-ALLERGEN | Non-Toxin | 0.03 | 0.82 | -0.61 | 0 | 6.29 | 2268.98 | outside |
| 416 | 426 | GKIADYNYKL | 1.6079 | 10 | ABCPRED | 1 | ALLERGEN | Non-Toxin | -0.18 | -0.77 | 0.05 | 1 | 8.76 | 1184.5 | outside |
| 777 | 789 | NTQEVFAQVKQI | 0.4262 | 12 | ABCPRED | 1 | NON-ALLERGEN | Non-Toxin | -0.16 | -0.38 | -0.12 | 0 | 6.35 | 1404.77 | outside |
| 251 | 271 | PGDSSSGWTAGAAAYYVGYL | 0.3382 | 20 | ABCPRED | 1 | ALLERGEN | Non-Toxin | 0.07 | 0.03 | -0.61 | -1 | 3.8 | 1993.39 | outside |
| 305 | 317 | SFTVEKGIYQTS | 0.2911 | 12 | ABCPRED | 1 | NON-ALLERGEN | Non-Toxin | -0.1 | -0.34 | -0.17 | 0 | 6.35 | 1359.67 | outside |
| 829 | 845 | ADAGFIKQYGDCLGDI | 0.1759 | 16 | ABCPRED | 1 | NON-ALLERGEN | Non-Toxin | -0.02 | 0.08 | 0 | -2 | 3.94 | 1686.1 | outside |
| 757 | 769 | GSFCTQLNRALT | 0.5729 | 12 | ABCPRED | 1 | NON-ALLERGEN | Non-Toxin | -0.13 | 0.05 | -0.39 | 1 | 8.6 | 1310.66 | outside |
| 568 | 582 | DIADTTDAVRDPQT | 0.5256 | 14 | ABCPRED,FBCPred | 2 | NON-ALLERGEN | Non-Toxin | -0.3 | -0.96 | 0.69 | -3 | 3.77 | 1517.75 | outside |
| 766 | 782 | ALTGIAVEQDKNTQEV | 0.7299 | 16 | ABCPRED | 1 | NON-ALLERGEN | Toxin | -0.15 | -0.4 | 0.26 | -2 | 4.14 | 1716.12 | outside |
| 913 | 924 | QNVLYENQKLI | 0.2774 | 11 | BCEPRED | 1 | NON-ALLERGEN | Non-Toxin | -0.18 | -0.58 | -0.22 | 0 | 6.35 | 1361.74 | outside |
| 327 | 339 | VRFPNITNLCPF | 1.2901 | 12 | ABCPRED | 1 | NON-ALLERGEN | Non-Toxin | -0.02 | 0.43 | -0.67 | 1 | 8.6 | 1420.86 | outside |
| 658 | 674 | NSYECDIPIGAGICAS | 0.5433 | 16 | ABCPRED | 1 | ALLERGEN | Non-Toxin | 0.03 | 0.39 | -0.24 | -2 | 3.67 | 1613.01 | outside |
| 238 | 250 | FQTLLALHRSYL | 0.2592 | 12 | ABCPRED | 1 | NON-ALLERGEN | Non-Toxin | -0.02 | 0.48 | -0.83 | 1.5 | 9.1 | 1461.91 | outside |
| 546 | 562 | LTGTGVLTESNKKFLP | 0.7066 | 16 | ABCPRED | 1 | NON-ALLERGEN | Non-Toxin | -0.08 | -0.11 | -0.07 | 1 | 8.94 | 1705.23 | outside |
| 885 | 903 | GWTFGAGAALQIPFAMQM | 0.626 | 18 | ABCPRED | 1 | ALLERGEN | Non-Toxin | 0.18 | 0.75 | -0.92 | 0 | 5.88 | 1897.51 | outside |
| 236 | 248 | TRFQTLLALHRS | 0.3559 | 12 | ABCPRED | 1 | NON-ALLERGEN | Non-Toxin | -0.23 | -0.16 | -0.27 | 2.5 | 12.01 | 1442.86 | outside |
| 300 | 312 | KCTLKSFTVEKG | 0.2904 | 12 | ABCPRED | 1 | NON-ALLERGEN | Non-Toxin | -0.22 | -0.38 | 0.39 | 2 | 9.25 | 1340.77 | outside |
| 402 | 412 | IRGDEVRQIA | -0.3641 | 10 | ABCPRED | 1 | ALLERGEN | Non-Toxin | -0.31 | -0.49 | 0.66 | 0 | 6.42 | 1156.45 | outside |
| 249 | 265 | LTPGDSSSGWTAGAAA | 0.5315 | 16 | ABCPRED | 1 | NON-ALLERGEN | Non-Toxin | 0.03 | 0 | -0.26 | -1 | 3.8 | 1448.72 | outside |
| 276 | 286 | LLKYNENGTI | 0.4428 | 10 | ABCPRED | 1 | NON-ALLERGEN | Non-Toxin | -0.12 | -0.47 | -0.17 | 0 | 6.35 | 1164.48 | outside |
| 754 | 772 | LQYGSFCTQLNRALTGIA | 0.9721 | 18 | ABCPRED | 1 | ALLERGEN | Non-Toxin | -0.03 | 0.31 | -0.61 | 1 | 8.57 | 1956.52 | outside |
| 383 | 391 | SPTKLNDL | 1.0358 | 8 | BCEPRED | 1 | ALLERGEN | Non-Toxin | -0.24 | -0.8 | 0.31 | 0 | 6.19 | 887.11 | outside |
| 420 | 430 | DYNYKLPDDF | 0.7529 | 10 | ABCPRED | 1 | ALLERGEN | Non-Toxin | -0.28 | -1.55 | 0.33 | -2 | 3.94 | 1289.49 | outside |
| 775 | 785 | DKNTQEVFAQ | 0.0859 | 10 | ABCPRED | 1 | NON-ALLERGEN | Non-Toxin | -0.32 | -1.33 | 0.47 | -1 | 4.38 | 1179.39 | outside |
| 445 | 457 | VGGNYNYLYRLF | -0.0113 | 12 | FBCPred | 1 | NON-ALLERGEN | Non-Toxin | -0.04 | -0.13 | -0.92 | 1 | 8.83 | 1478.85 | outside |
| 180 | 191 | EGKQGNFKNLR | 1.0042 | 11 | IEDB | 1 | ALLERGEN | Non-Toxin | -0.46 | -1.86 | 0.75 | 2 | 10.01 | 1290.61 | outside |
| 95 | 105 | TEKSNIIRGW | -0.7651 | 10 | ABCPRED | 1 | ALLERGEN | Non-Toxin | -0.26 | -0.92 | 0.21 | 1 | 9.1 | 1203.51 | outside |
| 1128 | 1146 | VVIGIVNNTVYDPLQPEL | 0.7941 | 18 | ABCPRED | 1 | NON-ALLERGEN | Non-Toxin | 0.07 | 0.57 | -0.52 | -2 | 3.67 | 1983.58 | outside |
| 218 | 236 | QGFSALEPLVDLPIGINI | 0.7918 | 18 | ABCPRED | 1 | NON-ALLERGEN | Non-Toxin | 0.13 | 0.83 | -0.48 | -2 | 3.67 | 1896.5 | outside |
| 28 | 40 | YTNSFTRGVYYP | -0.1085 | 12 | ABCPRED | 1 | NON-ALLERGEN | Non-Toxin | -0.14 | -0.76 | -0.68 | 1 | 8.83 | 1467.76 | outside |
| 456 | 461 | FRKSN | short | 5 | literature | 1 | ALLERGEN | Non-Toxin | -0.63 | -1.98 | 0.8 | 2 | 11.01 | 650.79 | outside |
| 580 | 598 | QTLEILDITPCSFGGVSV | 1.5916 | 18 | ABCPRED | 1 | NON-ALLERGEN | Non-Toxin | 0.09 | 0.8 | -0.43 | -2 | 3.67 | 1879.43 | outside |

| | | | | | | | | | | | | | | | |
|---|---|---|---|---|---|---|---|---|---|---|---|---|---|---|---|
| 1136 | 1147 | TVYDPLQPELD | 0.3454 | 11 | IEDB | 1 | ALLERGEN | Non-Toxin | -0.13 | -0.67 | 0.13 | -3 | 3.5 | 1289.56 | outside |
| 659 | 675 | SYECDIPIGAGICASY | 0.7437 | 16 | ABCPRED | 1 | ALLERGEN | Non-Toxin | 0.07 | 0.53 | -0.4 | -2 | 3.67 | 1662.08 | outside |
| 235 | 247 | ITRFQTLLALHR | 0.0169 | 12 | ABCPRED | 1 | NON-ALLERGEN | Non-Toxin | -0.15 | 0.28 | -0.44 | 2.5 | 12.01 | 1468.95 | outside |
| 297 | 315 | SETKCTLKSFTVEKGIYQ | 0.2727 | 18 | ABCPRED | 1 | ALLERGEN | Non-Toxin | -0.2 | -0.54 | 0.21 | 1 | 8.48 | 2062.63 | outside |
| 639 | 651 | GSNVFQTRAGCL | 1.0436 | 12 | ABCPRED | 1 | NON-ALLERGEN | Non-Toxin | -0.1 | 0.11 | -0.33 | 1 | 8.6 | 1252.58 | outside |
| 639 | 649 | GSNVFQTRAG | 0.6448 | 10 | ABCPRED | 1 | ALLERGEN | Non-Toxin | -0.18 | -0.5 | -0.12 | 1 | 10.11 | 1036.25 | outside |
| 408 | 420 | RQIAPGQTGKIA | 1.4558 | 12 | BCPREDS,FBCPred | 2 | NON-ALLERGEN | Non-Toxin | -0.18 | -0.49 | 0.12 | 2 | 11.01 | 1239.62 | outside |
| 583 | 603 | EILDITPCSFGGVSVITPGT | 1.0825 | 20 | ABCPRED | 1 | NON-ALLERGEN | Non-Toxin | 0.12 | 0.79 | -0.42 | -2 | 3.67 | 2006.6 | outside |
| 608 | 626 | VAVLYQDVNCTEVPVAIH | 0.7935 | 18 | ABCPRED | 1 | NON-ALLERGEN | Non-Toxin | 0.07 | 0.81 | -0.55 | -1.5 | 4.36 | 1970.54 | outside |
| 242 | 252 | LALHRSYLTP | 0.6697 | 10 | ABCPRED | 1 | NON-ALLERGEN | Non-Toxin | -0.08 | 0.11 | -0.58 | 1.5 | 9.1 | 1170.52 | outside |
| 736 | 754 | VDCTMYICGDSTECSNLL | -0.1588 | 18 | ABCPRED | 1 | NON-ALLERGEN | Non-Toxin | -0.04 | 0.39 | -0.25 | -3 | 3.5 | 1967.48 | outside |
| 907 | 923 | NGIGVTQNVLYENQKL | 0.5842 | 16 | ABCPRED | 1 | NON-ALLERGEN | Non-Toxin | -0.12 | -0.45 | -0.26 | 0 | 6.35 | 1790.27 | outside |
| 298 | 308 | ETKCTLKSFT | 0.6923 | 10 | ABCPRED | 1 | NON-ALLERGEN | Non-Toxin | -0.24 | -0.51 | 0.28 | 1 | 8.54 | 1157.48 | outside |
| 572 | 582 | TTDAVRDPQT | -0.0079 | 10 | ABCPRED | 1 | ALLERGEN | Non-Toxin | -0.37 | -1.27 | 0.6 | -1 | 4.21 | 1103.28 | outside |
| 582 | 594 | LEILDITPCSFG | 1.759 | 12 | ABCPRED | 1 | NON-ALLERGEN | Non-Toxin | 0.12 | 0.95 | -0.4 | -2 | 3.67 | 1307.7 | outside |
| 374 | 388 | FSTFKCYGVSPTKL | 0.8708 | 14 | ABCPRED,BCPREDS | 2 | NON-ALLERGEN | Non-Toxin | -0.05 | 0.14 | -0.41 | 2 | 9.25 | 1578.05 | outside |
| 911 | 929 | VTQNVLYENQKLIANQFN | 0.0578 | 18 | ABCPRED | 1 | ALLERGEN | Non-Toxin | -0.15 | -0.49 | -0.37 | 0 | 6.35 | 2136.68 | outside |
| 829 | 841 | ADAGFIKQYGDC | 0.2355 | 12 | ABCPRED | 1 | NON-ALLERGEN | Non-Toxin | -0.08 | -0.26 | 0.05 | -1 | 4.21 | 1287.57 | outside |
| 824 | 840 | NKVTLADAGFIKQYGD | 0.3248 | 16 | ABCPRED | 1 | NON-ALLERGEN | Non-Toxin | -0.12 | -0.36 | 0.07 | 0 | 6.3 | 1740.18 | outside |
| 195 | 207 | KNIDGYFKIYSK | 0.1412 | 12 | ABCPRED | 1 | ALLERGEN | Non-Toxin | -0.22 | -0.89 | 0.15 | 2 | 9.43 | 1475.88 | outside |
| 321 | 333 | QPTESIVRFPNI | -0.1214 | 12 | ABCPRED | 1 | NON-ALLERGEN | Non-Toxin | -0.14 | -0.31 | -0.11 | 0 | 6.36 | 1400.77 | outside |
| 382 | 392 | VSPTKLNDLC | 1.9573 | 10 | ABCPRED | 1 | ALLERGEN | Non-Toxin | -0.13 | 0.03 | 0 | 0 | 6.16 | 1089.41 | outside |
| 425 | 445 | LPDDFTGCVIAWNSNNLDSK | 0.2532 | 20 | FBCPred | 1 | NON-ALLERGEN | Non-Toxin | -0.11 | -0.33 | -0.07 | -2 | 3.94 | 2209.7 | outside |
| 752 | 770 | LLLQYGSFCTQLNRALTG | 0.6849 | 18 | ABCPRED | 1 | ALLERGEN | Non-Toxin | -0.03 | 0.38 | -0.68 | 1 | 8.57 | 1998.61 | outside |
| 401 | 411 | VIRGDEVRQI | -0.3175 | 10 | ABCPRED | 1 | ALLERGEN | Non-Toxin | -0.28 | -0.25 | 0.56 | 0 | 6.42 | 1184.51 | outside |
| 1202 | 1208 | ELGKYE | 0.4595 | 6 | IEDB | 1 | ALLERGEN | Non-Toxin | -0.27 | -1.47 | 0.82 | -1 | 4.54 | 737.89 | outside |
| 567 | 581 | RDIADTTDAVRDPQ | 0.44 | 14 | IEDB,literature | 2 | NON-ALLERGEN | Non-Toxin | -0.41 | -1.23 | 0.94 | -2 | 4.14 | 1572.83 | outside |
| 739 | 757 | TMYICGDSTECSNLLLQY | -0.0875 | 18 | ABCPRED | 1 | ALLERGEN | Non-Toxin | -0.04 | 0.16 | -0.49 | -2 | 3.67 | 2054.6 | outside |
| 385 | 403 | TKLNDLCFTNVYADSFVI | 1.0271 | 18 | ABCPRED | 1 | NON-ALLERGEN | Non-Toxin | 0 | 0.5 | -0.46 | -1 | 4.21 | 2063.61 | outside |
| 994 | 1014 | DRLITGRLQSLQTYVTQQLI | -0.3472 | 20 | ABCPRED | 1 | NON-ALLERGEN | Non-Toxin | -0.17 | -0.14 | -0.28 | 1 | 9.1 | 2347.04 | outside |
| 231 | 241 | IGINITRFQT | 0.8941 | 10 | ABCPRED | 1 | NON-ALLERGEN | Non-Toxin | -0.05 | 0.3 | -0.53 | 1 | 10.11 | 1162.51 | outside |
| 753 | 767 | LLQYGSFCTQLNRA | 0.9077 | 14 | ABCPRED,AAP | 2 | NON-ALLERGEN | Non-Toxin | -0.11 | 0.02 | -0.59 | 1 | 8.57 | 1614.06 | outside |
| 734 | 752 | TSVDCTMYICGDSTECSN | -0.0104 | 18 | ABCPRED | 1 | NON-ALLERGEN | Non-Toxin | -0.12 | -0.12 | -0.06 | -3 | 3.5 | 1929.33 | outside |
| 657 | 673 | NNSYECDIPIGAGICA | 0.6817 | 16 | ABCPRED | 1 | NON-ALLERGEN | Non-Toxin | 0.01 | 0.22 | -0.25 | -2 | 3.67 | 1640.04 | outside |
| 827 | 839 | TLADAGFIKQYG | 0.2851 | 12 | ABCPRED | 1 | NON-ALLERGEN | Non-Toxin | 0 | 0.08 | -0.3 | 0 | 6.18 | 1283.62 | outside |
| 807 | 817 | PDPSKPSKRS | 0.8374 | 10 | BCEPRED | 1 | ALLERGEN | Non-Toxin | -0.57 | -2.3 | 1.29 | 2 | 10.01 | 1098.34 | outside |
| 927 | 945 | FNSAIGKIQDSLSSTASA | 0.3582 | 18 | AAP | 1 | NON-ALLERGEN | Non-Toxin | -0.06 | 0.08 | -0.11 | 0 | 6.19 | 1797.2 | outside |
| 301 | 317 | CTLKSFTVEKGIYQTS | 0.0691 | 16 | ABCPRED | 1 | ALLERGEN | Non-Toxin | -0.12 | -0.15 | -0.14 | 1 | 8.51 | 1805.31 | outside |
| 852 | 864 | AQKFNGLTVLPP | 0.2224 | 12 | ABCPRED | 1 | NON-ALLERGEN | Non-Toxin | -0.01 | 0.1 | -0.42 | 1 | 9.11 | 1284.7 | outside |
| 426 | 444 | PDDFTGCVIAWNSNNLDS | 0.1508 | 18 | ABCPRED,BCPREDS | 2 | NON-ALLERGEN | Non-Toxin | -0.09 | -0.37 | -0.15 | -3 | 3.43 | 1968.33 | outside |
| 839 | 855 | DCLGDIAARDLICAQK | 0.7894 | 16 | ABCPRED | 1 | NON-ALLERGEN | Non-Toxin | -0.14 | 0.26 | 0.28 | -1 | 4.43 | 1705.2 | outside |
| 566 | 582 | GRDIADTTDAVRDPQT | 0.3229 | 16 | ABCPRED | 1 | NON-ALLERGEN | Toxin | -0.36 | -1.14 | 0.79 | -2 | 4.14 | 1731.02 | outside |
| 194 | 204 | FKNIDGYFKI | 0.0261 | 10 | ABCPRED | 1 | ALLERGEN | Toxin | -0.07 | -0.19 | -0.17 | 1 | 8.83 | 1244.6 | outside |
| 437 | 451 | NSNNLDSKVGGNYN | 0.7538 | 14 | AAP | 1 | ALLERGEN | Non-Toxin | -0.29 | -1.47 | 0.14 | 0 | 6.18 | 1495.73 | outside |
| 194 | 210 | FKNIDGYFKIYSKHTP | 0.3569 | 16 | ABCPRED | 1 | NON-ALLERGEN | Toxin | -0.17 | -0.84 | -0.1 | 2.5 | 9.43 | 1958.48 | outside |
| 194 | 206 | FKNIDGYFKIYS | -0.07 | 12 | ABCPRED,BCPREDS | 2 | ALLERGEN | Non-Toxin | -0.08 | -0.33 | -0.31 | 1 | 8.76 | 1494.88 | outside |
| 650 | 668 | LIGAEHVNNSYECDIPIG | 0.9717 | 18 | ABCPRED | 1 | NON-ALLERGEN | Non-Toxin | 0 | 0.03 | -0.18 | -2.5 | 4.14 | 1944.42 | outside |
| 914 | 934 | NVLYENQKLIANQFNSAIGK | 0.0231 | 20 | ABCPRED | 1 | ALLERGEN | Non-Toxin | -0.13 | -0.38 | -0.2 | 1 | 8.83 | 2264.88 | outside |
| 726 | 740 | ILPVSMTKTSVDCT | 1.606 | 14 | ABCPRED | 1 | NON-ALLERGEN | Non-Toxin | -0.02 | 0.6 | -0.25 | 0 | 6.16 | 1494.98 | outside |
| 247 | 261 | SYLTPGDSSSGWTA | 0.5658 | 14 | ABCPRED,AAP | 2 | ALLERGEN | Non-Toxin | -0.05 | -0.51 | -0.33 | -1 | 3.8 | 1428.66 | outside |
| 63 | 79 | TWFHAIHVSGTNGTKR | 0.5247 | 16 | ABCPRED | 1 | ALLERGEN | Non-Toxin | -0.14 | -0.6 | -0.34 | 3 | 11.01 | 1812.26 | outside |
| 651 | 669 | IGAEHVNNSYECDIPIGA | 1.0231 | 18 | ABCPRED | 1 | NON-ALLERGEN | Non-Toxin | -0.02 | -0.08 | -0.11 | -2.5 | 4.14 | 1902.33 | outside |
| 27 | 38 | AYTNSFTRGVY | -0.3374 | 11 | ellipro,literature | 2 | ALLERGEN | Non-Toxin | -0.13 | -0.4 | -0.58 | 1 | 8.93 | 1278.53 | outside |
| 1157 | 1174 | KNHTSPDVDLGDISGIN | 1.1116 | 17 | COVIDep,literature | 2 | ALLERGEN | Non-Toxin | -0.17 | -0.72 | 0.31 | -1.5 | 4.42 | 1782.15 | outside |
| 659 | 677 | SYECDIPIGAGICASYQT | 0.7709 | 18 | ABCPRED | 1 | ALLERGEN | Toxin | 0.02 | 0.24 | -0.37 | -2 | 3.67 | 1891.35 | outside |
| 1011 | 1028 | QLIRAAEIRASANLAAT | 0.3714 | 17 | COVIDep,literature | 2 | ALLERGEN | Non-Toxin | -0.11 | 0.38 | -0.05 | 1 | 9.95 | 1769.27 | outside |
| 132 | 144 | EFQFCNDPFLGV | 0.5987 | 12 | ABCPRED | 1 | NON-ALLERGEN | Non-Toxin | 0.03 | 0.24 | -0.45 | -2 | 3.67 | 1415.75 | outside |
| 11 | 29 | VSSQCVNLTTRTQLPPAY | 1.3205 | 18 | ABCPRED | 1 | ALLERGEN | Non-Toxin | -0.14 | -0.16 | -0.41 | 1 | 8.57 | 1978.51 | outside |
| 999 | 1013 | GRLQSLQTYVTQQL | -0.0073 | 14 | ABCPRED,BCPREDS | 2 | NON-ALLERGEN | Non-Toxin | -0.2 | -0.49 | -0.42 | 1 | 9.1 | 1635.08 | outside |
| 866 | 884 | TDEMIAQYTSALLAGTIT | 0.352 | 18 | ABCPRED | 1 | NON-ALLERGEN | Non-Toxin | 0.04 | 0.45 | -0.41 | -2 | 3.67 | 1899.41 | outside |
| 371 | 389 | SASFSTFKCYGVSPTKLN | 0.9891 | 18 | ABCPRED | 1 | NON-ALLERGEN | Non-Toxin | -0.09 | -0.07 | -0.31 | 2 | 9.25 | 1937.44 | outside |
| 414 | 424 | QTGKIADYNY | 1.5116 | 10 | ABCPRED | 1 | ALLERGEN | Non-Toxin | -0.21 | -1.18 | -0.09 | 0 | 6.18 | 1172.4 | outside |
| 638 | 648 | TGSNVFQTRA | 0.5406 | 10 | ABCPRED | 1 | ALLERGEN | Non-Toxin | -0.21 | -0.53 | -0.16 | 1 | 10.11 | 1080.3 | outside |
| 210 | 217 | INLVRDL | -0.3198 | 7 | IEDB,literature | 2 | NON-ALLERGEN | Non-Toxin | -0.11 | 0.69 | -0.1 | 0 | 6.19 | 842.11 | outside |
| 775 | 793 | DKNTQEVFAQVKQIYKTP | 0.1935 | 18 | ABCPRED | 1 | NON-ALLERGEN | Non-Toxin | -0.28 | -1.08 | 0.27 | 1 | 8.76 | 2137.69 | outside |
| 827 | 845 | TLADAGFIKQYGDCLGDI | 0.2181 | 18 | ABCPRED | 1 | NON-ALLERGEN | Non-Toxin | 0 | 0.24 | -0.12 | -2 | 3.94 | 1900.4 | outside |
| 344 | 350 | ATRFAS | -0.1511 | 6 | literature | 1 | ALLERGEN | Non-Toxin | -0.18 | 0.07 | -0.1 | 1 | 10.11 | 651.78 | outside |
| 542 | 560 | NFNGLTGTGVLTESNKKF | 0.6243 | 18 | ABCPRED | 1 | NON-ALLERGEN | Non-Toxin | -0.12 | -0.47 | -0.08 | 1 | 8.94 | 1927.42 | outside |
| 775 | 787 | DKNTQEVFAQVK | 0.4392 | 12 | ABCPRED | 1 | NON-ALLERGEN | Non-Toxin | -0.32 | -1.08 | 0.52 | 0 | 6.42 | 1406.73 | outside |
| 784 | 796 | QVKQIYKTPPIK | 0.299 | 12 | ABCPRED | 1 | ALLERGEN | Non-Toxin | -0.25 | -0.89 | 0.13 | 3 | 10.01 | 1442.95 | outside |
| 724 | 736 | TEILPVSMTKTS | 1.4845 | 12 | ABCPRED | 1 | NON-ALLERGEN | Non-Toxin | -0.07 | 0.14 | -0.08 | 0 | 6.35 | 1306.71 | outside |
| 376 | 386 | TFKCYGVSPT | 1.5059 | 10 | ABCPRED,IEDB,literature | 3 | ALLERGEN | Non-Toxin | -0.04 | 0.01 | -0.48 | 1 | 8.54 | 1102.4 | outside |
| 636 | 654 | YSTGSNVFQTRAGCLIGA | 0.7538 | 18 | ABCPRED | 1 | NON-ALLERGEN | Non-Toxin | -0.03 | 0.24 | -0.48 | 1 | 8.57 | 1845.32 | outside |
| 375 | 389 | STFKCYGVSPTKLN | 1.2695 | 14 | AAP | 1 | NON-ALLERGEN | Non-Toxin | -0.14 | -0.31 | -0.22 | 2 | 9.25 | 1544.98 | outside |
| 297 | 309 | SETKCTLKSFTV | 0.6191 | 12 | BCPREDS | 1 | NON-ALLERGEN | Non-Toxin | -0.18 | -0.14 | 0.13 | 1 | 8.54 | 1343.72 | outside |
| 420 | 432 | DYNYKLPDDFTG | 0.6394 | 12 | ABCPRED | 1 | NON-ALLERGEN | Non-Toxin | -0.23 | -1.38 | 0.24 | -2 | 3.94 | 1447.68 | outside |
| 606 | 624 | NQVAVLYQDVNCTEVPVA | 0.6999 | 18 | ABCPRED | 1 | ALLERGEN | Non-Toxin | -0.03 | 0.35 | -0.4 | -2 | 3.67 | 1962.47 | outside |

| | | | | | | | | | | | | | | | |
|---|---|---|---|---|---|---|---|---|---|---|---|---|---|---|---|
| 923 | 941 | IANQFNSAIGKIQDSLSS | 0.1989 | 18 | ABCPRED | 1 | NON-ALLERGEN | Non-Toxin | -0.09 | -0.07 | -0.15 | 0 | 6.19 | 1893.35 | outside |
| 825 | 841 | KVTLADAGFIKQYGDC | 0.4827 | 16 | ABCPRED | 1 | NON-ALLERGEN | Non-Toxin | -0.08 | 0.02 | -0.01 | 0 | 6.27 | 1729.21 | outside |
| 216 | 232 | LPQGFSALEPLVDLPI | 0.267 | 16 | ABCPRED | 1 | NON-ALLERGEN | Non-Toxin | 0.12 | 0.75 | -0.44 | -2 | 3.67 | 1709.26 | outside |
| 300 | 318 | KCTLKSFTVEKGIYQTSN | 0.1792 | 18 | ABCPRED | 1 | NON-ALLERGEN | Non-Toxin | -0.2 | -0.54 | 0.05 | 2 | 9.17 | 2047.62 | outside |
| 785 | 795 | VKQIYKTPPI | 0.4633 | 10 | ABCPRED | 1 | NON-ALLERGEN | Non-Toxin | -0.12 | -0.33 | -0.16 | 2 | 9.72 | 1186.61 | outside |
| 417 | 427 | KIADYNYKLP | 1.2565 | 10 | ABCPRED | 1 | NON-ALLERGEN | Non-Toxin | -0.21 | -0.89 | 0.05 | 1 | 8.76 | 1224.56 | outside |
| 356 | 365 | KRISNCVAD | 0.0005 | 9 | IEDB | 1 | ALLERGEN | Non-Toxin | -0.32 | -0.36 | 0.52 | 1 | 8.57 | 1005.27 | outside |
| 640 | 650 | SNVFQTRAGC | 0.7569 | 10 | ABCPRED | 1 | NON-ALLERGEN | Non-Toxin | -0.19 | -0.21 | -0.22 | 1 | 8.6 | 1082.33 | outside |
| 828 | 844 | LADAGFIKQYGDCLGD | 0.0965 | 16 | ABCPRED | 1 | NON-ALLERGEN | Toxin | -0.03 | 0.04 | 0 | -2 | 3.94 | 1686.1 | outside |
| 715 | 733 | PTNFTISVTTEILPVSMT | 1.3036 | 18 | ABCPRED | 1 | ALLERGEN | Non-Toxin | 0.06 | 0.59 | -0.58 | -1 | 4 | 1951.53 | outside |
| 91 | 103 | YFASTEKSNIIR | -0.0942 | 12 | BCEPRED | 1 | NON-ALLERGEN | Non-Toxin | -0.21 | -0.45 | 0.04 | 1 | 8.93 | 1428.77 | outside |
| 405 | 417 | DEVRQIAPGQTG | 1.0397 | 12 | ABCPRED | 1 | ALLERGEN | Non-Toxin | -0.24 | -0.93 | 0.43 | -1 | 4.38 | 1270.54 | outside |
| 82 | 98 | PVLPFNDGVYFASTEK | 0.4819 | 16 | ABCPRED | 1 | NON-ALLERGEN | Non-Toxin | -0.02 | -0.07 | -0.22 | -1 | 4.38 | 1784.22 | outside |
| 910 | 928 | GVTQNVLYENQKLIANQF | -0.0266 | 18 | ABCPRED | 1 | ALLERGEN | Non-Toxin | -0.11 | -0.32 | -0.38 | 0 | 6.35 | 2079.63 | outside |
| 320 | 336 | VQPTESIVRFPNITNL | 0.4454 | 16 | ABCPRED | 1 | NON-ALLERGEN | Non-Toxin | -0.09 | 0.01 | -0.3 | 0 | 6.36 | 1828.34 | outside |
| 1245 | 1263 | KGCCSCGSCCKFDEDDSE | 0.2445 | 18 | ABCPRED | 1 | NON-ALLERGEN | Toxin | -0.29 | -0.73 | 0.8 | -3 | 4.11 | 1916.29 | inside |
| 295 | 305 | PLSETKCTLK | 0.8717 | 10 | ABCPRED | 1 | NON-ALLERGEN | Non-Toxin | -0.24 | -0.5 | 0.39 | 1 | 8.54 | 1119.48 | outside |
| 249 | 261 | LTPGDSSSGWTA | 0.689 | 12 | ABCPRED,AAP | 2 | NON-ALLERGEN | Non-Toxin | -0.04 | -0.42 | -0.22 | -1 | 3.8 | 1178.38 | outside |
| 1023 | 1039 | NLAATKMSECVLGQSK | 0.4846 | 16 | ABCPRED | 1 | NON-ALLERGEN | Non-Toxin | -0.14 | -0.08 | 0.08 | 1 | 8.54 | 1680.2 | outside |
| 298 | 311 | ETKCTLKSFTVEK | 0.5711 | 13 | BCEPRED | 1 | NON-ALLERGEN | Non-Toxin | -0.28 | -0.64 | 0.56 | 1 | 8.51 | 1513.95 | outside |
| 857 | 873 | GLTVLPPLLTDEMIAQ | 0.4069 | 16 | ABCPRED | 1 | NON-ALLERGEN | Non-Toxin | 0.1 | 0.76 | -0.43 | -2 | 3.67 | 1711.3 | outside |
| 1250 | 1262 | CGSCCKFDEDDS | 0.3274 | 12 | BCPREDS | 1 | NON-ALLERGEN | Non-Toxin | -0.29 | -0.8 | 0.84 | -3 | 3.84 | 1308.51 | inside |
| 859 | 871 | TVLPPLLTDEMI | 0.2131 | 12 | ABCPRED | 1 | ALLERGEN | Non-Toxin | 0.11 | 0.87 | -0.4 | -2 | 3.67 | 1341.81 | outside |
| 1256 | 1266 | FDEDDSEPVL | 0.3154 | 10 | IEDB | 1 | ALLERGEN | Non-Toxin | -0.2 | -0.91 | 0.95 | -5 | 3.34 | 1165.3 | inside |
| 414 | 426 | QTGKIADYNYKL | 1.4851 | 12 | ABCPRED | 1 | ALLERGEN | Non-Toxin | -0.23 | -0.99 | 0.03 | 1 | 8.76 | 1413.77 | outside |
| 1000 | 1012 | RLQSLQTYVTQQ | 0.1103 | 12 | BCEPRED | 1 | NON-ALLERGEN | Non-Toxin | -0.29 | -0.85 | -0.34 | 1 | 9.1 | 1464.83 | outside |
| 307 | 317 | TVEKGIYQTS | 0.0601 | 10 | ABCPRED | 1 | NON-ALLERGEN | Non-Toxin | -0.16 | -0.61 | 0.01 | 0 | 6.35 | 1125.39 | outside |
| 319 | 335 | RVQPTESIVRFPNITN | 0.4049 | 16 | ABCPRED | 1 | NON-ALLERGEN | Non-Toxin | -0.23 | -0.51 | 0 | 1 | 9.95 | 1871.36 | outside |
| 828 | 840 | LADAGFIKQYGD | 0.0636 | 12 | ABCPRED | 1 | NON-ALLERGEN | Non-Toxin | -0.04 | -0.15 | -0.02 | -1 | 4.21 | 1297.6 | outside |
| 1020 | 1036 | ASANLAATKMSECVLG | 0.4586 | 16 | ABCPRED | 1 | ALLERGEN | Non-Toxin | 0 | 0.61 | -0.19 | 0 | 6.32 | 1566.04 | outside |
| 369 | 387 | YNSASFSTFKCYGVSPTK | 0.8466 | 18 | BCPREDS | 1 | ALLERGEN | Non-Toxin | -0.11 | -0.36 | -0.33 | 2 | 9.17 | 1987.45 | outside |
| 118 | 136 | LIVNNATNVVIKVCEFQF | 0.1148 | 18 | ABCPRED | 1 | ALLERGEN | Non-Toxin | 0.06 | 0.97 | -0.64 | 0 | 6.32 | 2051.71 | outside |
| 17 | 27 | NLTTRTQLPP | 1.2832 | 10 | ABCPRED | 1 | ALLERGEN | Non-Toxin | -0.27 | -0.92 | -0.14 | 1 | 10.11 | 1140.45 | outside |
| 248 | 261 | YLTPGDSSSGWTA | 0.627 | 13 | IEDB | 1 | NON-ALLERGEN | Non-Toxin | -0.03 | -0.48 | -0.38 | -1 | 3.8 | 1341.57 | outside |
| 29 | 39 | TNSFTRGVYY | -0.251 | 10 | ABCPRED | 1 | ALLERGEN | Non-Toxin | -0.17 | -0.62 | -0.59 | 1 | 8.93 | 1207.44 | outside |
| 656 | 667 | VNNSYECDIPI | 0.6124 | 11 | IEDB,literature | 2 | NON-ALLERGEN | Non-Toxin | -0.08 | -0.18 | -0.15 | -2 | 3.67 | 1266.54 | outside |
| 423 | 429 | YKLPDD | -0.9691 | 6 | IEDB | 1 | NON-ALLERGEN | Non-Toxin | -0.34 | -1.67 | 0.82 | -1 | 4.21 | 749.89 | outside |
| 204 | 214 | YSKHTPINLV | 1.0391 | 10 | ABCPRED | 1 | NON-ALLERGEN | Non-Toxin | -0.08 | -0.25 | -0.48 | 1.5 | 8.94 | 1171.51 | outside |
| 15 | 27 | CVNLTTRTQLPP | 1.649 | 12 | BCPREDS,FBCPred | 2 | NON-ALLERGEN | Non-Toxin | -0.18 | -0.21 | -0.33 | 1 | 8.6 | 1342.75 | outside |
| 1109 | 1125 | FYEPQIITTDNTFVSG | 0.299 | 16 | ABCPRED | 1 | ALLERGEN | Non-Toxin | -0.01 | -0.09 | -0.43 | -2 | 3.67 | 1832.23 | outside |
| 673 | 685 | SYQTQTNSPRRA | 0.0145 | 12 | ABCPRED,AAP,BCEPRED | 3 | NON-ALLERGEN | Non-Toxin | -0.52 | -1.97 | 0.3 | 2 | 10.84 | 1408.65 | outside |
| 247 | 267 | SYLTPGDSSSGWTAGAAAYY | 0.4845 | 20 | ABCPRED | 1 | ALLERGEN | Non-Toxin | 0.01 | -0.23 | -0.53 | -1 | 3.8 | 2025.38 | outside |
| 196 | 206 | NIDGYFKIYS | -0.1497 | 10 | ABCPRED | 1 | ALLERGEN | Non-Toxin | -0.05 | -0.29 | -0.42 | 0 | 6.18 | 1219.5 | outside |
| 93 | 105 | ASTEKSNIIRGW | -0.5092 | 12 | BCPREDS | 1 | NON-ALLERGEN | Non-Toxin | -0.21 | -0.68 | 0.16 | 1 | 9.1 | 1361.69 | outside |
| 1252 | 1264 | SCCKFDEDDSEP | 0.271 | 12 | FBCPred | 1 | NON-ALLERGEN | Non-Toxin | -0.37 | -1.4 | 1.18 | -4 | 3.77 | 1374.55 | inside |
| 757 | 773 | GSFCTQLNRALTGIAV | 0.6508 | 16 | ABCPRED | 1 | ALLERGEN | Non-Toxin | 0.01 | 0.67 | -0.53 | 1 | 8.6 | 1651.15 | outside |
| 216 | 234 | LPQGFSALEPLVDLPIGI | 0.6706 | 18 | AAP | 1 | NON-ALLERGEN | Non-Toxin | 0.16 | 0.89 | -0.49 | -2 | 3.67 | 1879.51 | outside |
| 526 | 544 | GPKKSTNLVKNKCVNFNF | 1.1626 | 18 | ABCPRED | 1 | NON-ALLERGEN | Non-Toxin | -0.25 | -0.71 | 0.11 | 4 | 10.05 | 2038.66 | outside |
| 127 | 137 | VIKVCEFQFC | -0.8173 | 10 | ABCPRED | 1 | NON-ALLERGEN | Non-Toxin | 0.07 | 1.26 | -0.56 | 0 | 6.29 | 1215.63 | outside |
| 128 | 138 | IKVCEFQFCN | -0.6653 | 10 | ABCPRED | 1 | ALLERGEN | Non-Toxin | -0.05 | 0.49 | -0.39 | 0 | 6.29 | 1230.6 | outside |
| 749 | 767 | CSNLLLQYGSFCTQLNRA | 0.6966 | 18 | ABCPRED | 1 | NON-ALLERGEN | Non-Toxin | -0.1 | 0.13 | -0.58 | 1 | 8.38 | 2031.6 | outside |
| 516 | 537 | ELLHAPATVCGPKKSTNLVKN | 0.0029 | 21 | IEDB,literature | 2 | NON-ALLERGEN | Non-Toxin | -0.14 | -0.25 | 0.05 | 2.5 | 9.25 | 2220.93 | outside |
| 1203 | 1210 | LGKYEQY | 0.9561 | 7 | literature | 1 | ALLERGEN | Non-Toxin | -0.24 | -1.44 | -0.03 | 0 | 6.35 | 900.1 | outside |
| 302 | 320 | TLKSFTVEKGIYQTSNFR | 0.2601 | 18 | ABCPRED | 1 | ALLERGEN | Non-Toxin | -0.21 | -0.56 | -0.03 | 2 | 9.72 | 2119.67 | outside |
| 1243 | 1259 | CLKGCCSCGSCCKFDE | 0.2145 | 16 | ABCPRED | 1 | NON-ALLERGEN | Non-Toxin | -0.15 | 0.27 | 0.14 | 0 | 6.27 | 1686.2 | inside |
| 910 | 930 | GVTQNVLYENQKLIANQFNS | 0.0293 | 20 | ABCPRED | 1 | NON-ALLERGEN | Non-Toxin | -0.14 | -0.5 | -0.32 | 0 | 6.35 | 2280.84 | outside |
| 909 | 925 | IGVTQNVLYENQKLIA | 0.5664 | 16 | ABCPRED | 1 | NON-ALLERGEN | Non-Toxin | -0.03 | 0.19 | -0.41 | 0 | 6.35 | 1803.35 | outside |
| 907 | 925 | NGIGVTQNVLYENQKLIA | 0.4669 | 18 | ABCPRED | 1 | NON-ALLERGEN | Non-Toxin | -0.06 | -0.05 | -0.36 | 0 | 6.35 | 1974.54 | outside |
| 401 | 413 | VIRGDEVRQIAP | 0.0486 | 12 | ABCPRED | 1 | NON-ALLERGEN | Non-Toxin | -0.22 | -0.19 | 0.43 | 0 | 6.42 | 1352.73 | outside |
| 657 | 675 | NNSYECDIPIGAGICASY | 0.7153 | 18 | ABCPRED | 1 | NON-ALLERGEN | Non-Toxin | 0 | 0.08 | -0.33 | -2 | 3.67 | 1890.32 | outside |
| 249 | 263 | LTPGDSSSGWTAGA | 0.5017 | 14 | BCPREDS | 1 | NON-ALLERGEN | Non-Toxin | 0 | -0.26 | -0.22 | -1 | 3.8 | 1306.54 | outside |
| 580 | 596 | QTLEILDITPCSFGGV | 1.5382 | 16 | ABCPRED | 1 | NON-ALLERGEN | Non-Toxin | 0.08 | 0.69 | -0.41 | -2 | 3.67 | 1693.19 | outside |
| 1201 | 1212 | QELGKYEQYIK | 0.2086 | 11 | BCEPRED | 1 | NON-ALLERGEN | Non-Toxin | -0.31 | -1.5 | 0.38 | 0 | 6.48 | 1398.75 | outside |
| 321 | 335 | QPTESIVRFPNITN | 0.055 | 14 | ABCPRED,BCPREDS | 2 | NON-ALLERGEN | Non-Toxin | -0.18 | -0.56 | -0.11 | 0 | 6.36 | 1616.01 | outside |
| 859 | 873 | TVLPPLLTDEMIAQ | 0.2417 | 14 | ABCPRED,FBCPred | 2 | NON-ALLERGEN | Non-Toxin | 0.06 | 0.62 | -0.36 | -2 | 3.67 | 1541.05 | outside |
| 637 | 655 | STGSNVFQTRAGCLIGAE | 0.8825 | 18 | ABCPRED | 1 | ALLERGEN | Non-Toxin | -0.06 | 0.12 | -0.19 | 0 | 6.32 | 1811.26 | outside |
| 856 | 874 | NGLTVLPPLLTDEMIAQY | 0.1888 | 18 | ABCPRED | 1 | NON-ALLERGEN | Non-Toxin | 0.05 | 0.41 | -0.5 | -2 | 3.67 | 1988.61 | outside |
| 203 | 213 | IYSKHTPINL | 1.2064 | 10 | ABCPRED | 1 | ALLERGEN | Non-Toxin | -0.06 | -0.22 | -0.51 | 1.5 | 8.94 | 1185.54 | outside |
| 824 | 842 | NKVTLADAGFIKQYGDCL | 0.4094 | 18 | ABCPRED | 1 | ALLERGEN | Non-Toxin | -0.07 | 0.03 | -0.09 | 0 | 6.27 | 1956.51 | outside |
| 884 | 902 | SGWTFGAGAALQIPFAMQ | 0.5141 | 18 | ABCPRED | 1 | NON-ALLERGEN | Non-Toxin | 0.15 | 0.6 | -0.83 | 0 | 5.88 | 1853.39 | outside |
| 516 | 530 | ELLHAPATVCGPKK | -0.1481 | 14 | BCPREDS | 1 | NON-ALLERGEN | Toxin | -0.09 | -0.06 | 0.07 | 1.5 | 8.54 | 1463.96 | outside |
| 386 | 404 | KLNDLCFTNVYADSFVIR | 1.0694 | 18 | ABCPRED | 1 | ALLERGEN | Non-Toxin | -0.09 | 0.29 | -0.27 | 0 | 6.28 | 2118.69 | outside |
| 220 | 230 | FSALEPLVDL | 0.3757 | 10 | ABCPRED | 1 | ALLERGEN | Non-Toxin | 0.13 | 1.08 | -0.36 | -2 | 3.67 | 1103.42 | outside |
| 636 | 648 | YSTGSNVFQTRA | 0.4054 | 12 | BCPREDS | 1 | NON-ALLERGEN | Non-Toxin | -0.2 | -0.62 | -0.3 | 1 | 9.1 | 1330.58 | outside |
| 820 | 832 | DLLFNKVTLADA | 0.7587 | 12 | ABCPRED | 1 | NON-ALLERGEN | Non-Toxin | -0.01 | 0.57 | -0.13 | -1 | 4.21 | 1319.69 | outside |
| 541 | 559 | FNFNGLTGTGVLTESNKK | 0.9066 | 18 | ABCPRED | 1 | NON-ALLERGEN | Non-Toxin | -0.12 | -0.47 | -0.08 | 1 | 8.94 | 1927.42 | outside |
| 583 | 599 | EILDITPCSFGGVSVI | 1.3971 | 16 | ABCPRED | 1 | ALLERGEN | Non-Toxin | 0.17 | 1.21 | -0.47 | -2 | 3.67 | 1650.16 | outside |

| | | | | | | | | | | | | | | | |
|---|---|---|---|---|---|---|---|---|---|---|---|---|---|---|---|
| 637 | 648 | STGSNVFQTRA | 0.502 | 11 | IEDB | 1 | NON-ALLERGEN | Non-Toxin | -0.22 | -0.55 | -0.12 | 1 | 10.11 | 1167.39 | outside |
| 823 | 841 | FNKVTLADAGFIKQYGDC | 0.3771 | 18 | AAP | 1 | ALLERGEN | Non-Toxin | -0.07 | -0.02 | -0.13 | 0 | 6.27 | 1990.52 | outside |
| 740 | 751 | MYICGDSTECS | -0.3346 | 11 | IEDB | 1 | NON-ALLERGEN | Non-Toxin | -0.07 | 0.04 | -0.11 | -2 | 3.67 | 1208.48 | outside |
| 609 | 625 | AVLYQDVNCTEVPVAI | 0.7384 | 16 | ABCPRED | 1 | NON-ALLERGEN | Non-Toxin | 0.07 | 0.85 | -0.49 | -2 | 3.67 | 1734.23 | outside |
| 569 | 582 | IADTTDAVRDPQT | 0.3018 | 13 | IEDB | 1 | NON-ALLERGEN | Non-Toxin | -0.27 | -0.76 | 0.52 | -2 | 3.94 | 1402.65 | outside |
| 105 | 115 | IFGTTLDSKT | 0.4027 | 10 | ABCPRED | 1 | NON-ALLERGEN | Non-Toxin | -0.06 | 0.04 | -0.1 | 0 | 6.19 | 1082.36 | outside |
| 755 | 767 | QYGSFCTQLNRA | 1.1892 | 12 | ABCPRED | 1 | ALLERGEN | Non-Toxin | -0.22 | -0.61 | -0.38 | 1 | 8.57 | 1387.7 | outside |
| 737 | 753 | DCTMYICGDSTECSNL | -0.2636 | 16 | ABCPRED | 1 | NON-ALLERGEN | Non-Toxin | -0.11 | -0.06 | -0.07 | -3 | 3.5 | 1755.15 | outside |
| 554 | 564 | ESNKKFLPFQ | 0.883 | 10 | ellipro | 1 | ALLERGEN | Non-Toxin | -0.27 | -1.13 | 0.29 | 1 | 8.94 | 1237.56 | outside |
| 826 | 846 | VTLADAGFIKQYGDCLGDIA | 0.4018 | 20 | ABCPRED | 1 | NON-ALLERGEN | Non-Toxin | 0.04 | 0.52 | -0.21 | -2 | 3.94 | 2070.64 | outside |
| 866 | 878 | TDEMIAQYTSAL | 0.0429 | 12 | ABCPRED | 1 | ALLERGEN | Non-Toxin | -0.05 | -0.02 | -0.21 | -2 | 3.67 | 1342.65 | outside |
| 303 | 315 | LKSFTVEKGIYQ | -0.236 | 12 | ABCPRED | 1 | NON-ALLERGEN | Non-Toxin | -0.11 | -0.22 | -0.07 | 1 | 8.83 | 1412.83 | outside |
| 106 | 118 | FGTTLDSKTQSL | 0.7789 | 12 | ABCPRED | 1 | ALLERGEN | Non-Toxin | -0.14 | -0.38 | -0.04 | 0 | 6.19 | 1297.6 | outside |
| 298 | 310 | ETKCTLKSFTVE | 0.7594 | 12 | ABCPRED | 1 | NON-ALLERGEN | Non-Toxin | -0.21 | -0.37 | 0.36 | 0 | 6.46 | 1385.76 | outside |
| 439 | 451 | NNLDSKVGGNYN | 0.7538 | 12 | AAP | 1 | NON-ALLERGEN | Non-Toxin | -0.27 | -1.36 | 0.13 | 0 | 6.18 | 1294.52 | outside |
| 355 | 365 | RKRISNCVAD | -0.0428 | 10 | ABCPRED | 1 | NON-ALLERGEN | Non-Toxin | -0.47 | -0.77 | 0.77 | 2 | 9.55 | 1161.47 | outside |
| 543 | 561 | FNGLTGTGVLTESNKKFL | 0.3938 | 18 | ABCPRED | 1 | NON-ALLERGEN | Non-Toxin | -0.06 | -0.07 | -0.19 | 1 | 8.94 | 1926.48 | outside |
| 248 | 264 | YLTPGDSSSGWTAGAA | 0.465 | 16 | ABCPRED,IEDB | 2 | NON-ALLERGEN | Non-Toxin | 0.01 | -0.19 | -0.37 | -1 | 3.8 | 1540.82 | outside |
| 756 | 774 | YGSFCTQLNRALTGIAVE | 0.7509 | 18 | ABCPRED | 1 | NON-ALLERGEN | Non-Toxin | -0.03 | 0.33 | -0.43 | 0 | 6.32 | 1943.47 | outside |
| 612 | 624 | YQDVNCTEVPVA | 1.0646 | 12 | AAP,FBCPred | 2 | NON-ALLERGEN | Non-Toxin | -0.08 | -0.06 | -0.19 | -2 | 3.67 | 1337.63 | outside |
| 1009 | 1027 | TQQLIRAAEIRASANLAA | 0.2 | 18 | AAP | 1 | NON-ALLERGEN | Non-Toxin | -0.14 | 0.16 | -0.04 | 1 | 9.95 | 1897.42 | outside |
| 7 | 25 | LLPLVSSQCVNLTTRTQL | 1.1517 | 18 | ABCPRED | 1 | NON-ALLERGEN | Non-Toxin | -0.06 | 0.53 | -0.56 | 1 | 8.6 | 1986.64 | outside |
| 1200 | 1212 | LQELGKYEQYIK | 0.2102 | 12 | AAP | 1 | ALLERGEN | Non-Toxin | -0.24 | -1.06 | 0.2 | 0 | 6.48 | 1511.93 | outside |
| 131 | 147 | CEFQFCNDPFLGVYYH | 0.0771 | 16 | ABCPRED | 1 | NON-ALLERGEN | Non-Toxin | 0 | -0.02 | -0.72 | -1.5 | 4.36 | 1982.44 | outside |
| 751 | 769 | NLLLQYGSFCTQLNRALT | 0.6949 | 18 | ABCPRED | 1 | ALLERGEN | Non-Toxin | -0.07 | 0.21 | -0.67 | 1 | 8.57 | 2055.66 | outside |
| 10 | 28 | LVSSQCVNLTTRTQLPPA | 1.2068 | 18 | ABCPRED | 1 | ALLERGEN | Non-Toxin | -0.11 | 0.12 | -0.38 | 1 | 8.6 | 1928.5 | outside |
| 178 | 188 | DLEGKQGNFK | 1.3065 | 10 | ABCPRED | 1 | NON-ALLERGEN | Non-Toxin | -0.34 | -1.6 | 0.81 | 0 | 6.42 | 1135.39 | outside |
| 995 | 1012 | RLITGRLQSLQTYVTQQ | -0.2774 | 17 | COVIDep,literature | 2 | NON-ALLERGEN | Non-Toxin | -0.24 | -0.44 | -0.3 | 2 | 10.84 | 2005.58 | outside |
| 1187 | 1203 | NEVAKNLNESLIDLQE | 0.3993 | 16 | ABCPRED | 1 | NON-ALLERGEN | Non-Toxin | -0.21 | -0.67 | 0.43 | -3 | 4.01 | 1829.24 | outside |
| 406 | 416 | EVRQIAPGQT | 1.0655 | 10 | ABCPRED | 1 | NON-ALLERGEN | Non-Toxin | -0.23 | -0.72 | 0.22 | 0 | 6.36 | 1098.37 | outside |
| 997 | 1017 | ITGRLQSLQTYVTQQLIRAA | 0.0539 | 20 | ABCPRED | 1 | NON-ALLERGEN | Non-Toxin | -0.14 | 0.03 | -0.4 | 2 | 10.84 | 2260.94 | outside |
| 908 | 926 | GIGVTQNVLYENQKLIAN | 0.5302 | 18 | ABCPRED | 1 | NON-ALLERGEN | Non-Toxin | -0.06 | -0.05 | -0.36 | 0 | 6.35 | 1974.54 | outside |
| 777 | 793 | NTQEVFAQVKQIYKTP | 0.2234 | 16 | ABCPRED | 1 | NON-ALLERGEN | Non-Toxin | -0.2 | -0.76 | -0.07 | 1 | 8.83 | 1894.4 | outside |
| 646 | 658 | RAGCLIGAEHVN | 1.3707 | 12 | ABCPRED | 1 | NON-ALLERGEN | Non-Toxin | -0.06 | 0.26 | -0.12 | 0.5 | 7.07 | 1239.59 | outside |
| 841 | 853 | LGDIAARDLICA | 1.1914 | 12 | ABCPRED | 1 | ALLERGEN | Non-Toxin | 0.02 | 1.05 | -0.06 | -1 | 4.21 | 1230.61 | outside |
| 231 | 240 | IGINITRFQ | 1.3386 | 9 | literature | 1 | NON-ALLERGEN | Non-Toxin | -0.03 | 0.41 | -0.54 | 1 | 10.11 | 1061.39 | outside |
| 1070 | 1079 | AQEKNFTTA | 0.7576 | 9 | IEDB | 1 | NON-ALLERGEN | Non-Toxin | -0.26 | -1.04 | 0.23 | 0 | 6.35 | 1009.2 | outside |
| 552 | 562 | LTESNKKFLP | 0.6681 | 10 | BCEPRED | 1 | ALLERGEN | Non-Toxin | -0.23 | -0.75 | 0.3 | 1 | 8.94 | 1176.52 | outside |
| 300 | 316 | KCTLKSFTVEKGIYQT | 0.11 | 16 | ABCPRED | 1 | NON-ALLERGEN | Non-Toxin | -0.17 | -0.34 | 0.02 | 2 | 9.17 | 1846.41 | outside |
| 903 | 915 | AYRFNGIGVTQN | 1.2149 | 12 | ABCPRED | 1 | ALLERGEN | Non-Toxin | -0.12 | -0.38 | -0.45 | 1 | 9.1 | 1339.65 | outside |
| 553 | 564 | TESNKKFLPFQ | 0.9243 | 11 | ellipro | 1 | NON-ALLERGEN | Non-Toxin | -0.26 | -1.09 | 0.23 | 1 | 8.94 | 1338.68 | outside |
| 1110 | 1126 | YEPQIITTDNTFVSGN | 0.5227 | 16 | ABCPRED | 1 | NON-ALLERGEN | Non-Toxin | -0.09 | -0.48 | -0.26 | -2 | 3.67 | 1799.16 | outside |
| 651 | 671 | IGAEHVNNSYECDIPIGAGI | 1.1141 | 20 | ABCPRED,BCPREDS,literature | 3 | NON-ALLERGEN | Non-Toxin | 0.03 | 0.14 | -0.19 | -2.5 | 4.14 | 2072.58 | outside |
| 217 | 229 | PQGFSALEPLVD | 0.3583 | 12 | BCPREDS,FBCPred | 2 | NON-ALLERGEN | Non-Toxin | 0.02 | 0.12 | -0.13 | -2 | 3.67 | 1272.59 | outside |
| 649 | 667 | CLIGAEHVNNSYECDIPI | 0.9191 | 18 | ABCPRED,FBCPred | 2 | NON-ALLERGEN | Non-Toxin | -0.01 | 0.19 | -0.24 | -2.5 | 4.14 | 1990.5 | outside |
| 738 | 754 | CTMYICGDSTECSNLL | -0.4585 | 16 | ABCPRED | 1 | NON-ALLERGEN | Toxin | -0.03 | 0.39 | -0.38 | -2 | 3.67 | 1753.23 | outside |
| 1008 | 1028 | VTQQLIRAAEIRASANLAAT | 0.3436 | 20 | ABCPRED | 1 | ALLERGEN | Non-Toxin | -0.11 | 0.32 | -0.13 | 1 | 9.95 | 2097.69 | outside |
| 842 | 854 | GDIAARDLICAQ | 1.1569 | 12 | ABCPRED | 1 | ALLERGEN | Non-Toxin | -0.08 | 0.44 | 0.11 | -1 | 4.21 | 1245.58 | outside |
| 1132 | 1150 | IVNNTVYDPLQPELDSFK | 0.0258 | 18 | ABCPRED | 1 | NON-ALLERGEN | Non-Toxin | -0.12 | -0.42 | -0.04 | -2 | 4.03 | 2092.6 | outside |
| 295 | 311 | PLSETKCTLKSFTVEK | 0.5718 | 16 | ABCPRED | 1 | NON-ALLERGEN | Non-Toxin | -0.21 | -0.43 | 0.36 | 1 | 8.51 | 1811.35 | outside |
| 547 | 563 | TGTGVLTESNKKFLPF | 0.9925 | 16 | ABCPRED | 1 | NON-ALLERGEN | Non-Toxin | -0.07 | -0.17 | -0.11 | 1 | 8.94 | 1739.24 | outside |
| 771 | 787 | AVEQDKNTQEVFAQVK | 0.4031 | 16 | ABCPRED | 1 | NON-ALLERGEN | Non-Toxin | -0.27 | -0.88 | 0.46 | -1 | 4.68 | 1834.25 | outside |
| 71 | 82 | SGTNGTKRFDN | 0.5906 | 11 | IEDB,literature | 2 | NON-ALLERGEN | Non-Toxin | -0.41 | -1.74 | 0.58 | 1 | 9.1 | 1196.39 | outside |
| 783 | 799 | AQVKQIYKTPPIKDFG | 0.1976 | 16 | ABCPRED | 1 | NON-ALLERGEN | Non-Toxin | -0.17 | -0.63 | 0.1 | 2 | 9.55 | 1833.4 | outside |
| 672 | 691 | ASYQTQTNSPRRARSVASQ | 0.2556 | 19 | IEDB,literature | 2 | NON-ALLERGEN | Non-Toxin | -0.43 | -1.34 | 0.26 | 3 | 11.72 | 2108.51 | outside |
| 21 | 32 | RTQLPPAYTNS | 0.871 | 11 | IEDB,literature | 2 | NON-ALLERGEN | Non-Toxin | -0.28 | -1.15 | -0.15 | 1 | 9.1 | 1247.52 | outside |
| 632 | 642 | TWRVYSTGSN | 0.4797 | 10 | ABCPRED | 1 | ALLERGEN | Non-Toxin | -0.22 | -0.94 | -0.42 | 1 | 9.1 | 1170.38 | outside |
| 1208 | 1226 | QYIKWPWYIWLGFIAGLI | 0.7751 | 18 | ABCPRED | 1 | NON-ALLERGEN | Non-Toxin | 0.25 | 0.84 | -1.41 | 1 | 8.83 | 2268.04 | outside_transmembrane-region |
| 446 | 464 | GGNYNYLYRLFRKSNLKP | 0.3059 | 18 | ABCPRED | 1 | NON-ALLERGEN | Non-Toxin | -0.3 | -1.12 | -0.11 | 4 | 10.17 | 2203.8 | outside |
| 788 | 802 | IYKTPPIKDFGGFN | -0.2461 | 14 | ABCPRED,literature | 2 | NON-ALLERGEN | Non-Toxin | -0.06 | -0.44 | -0.15 | 1 | 8.83 | 1597.05 | outside |
| 851 | 869 | CAQKFNGLTVLPPLLTDE | 0.0951 | 18 | ABCPRED | 1 | NON-ALLERGEN | Non-Toxin | -0.03 | 0.2 | -0.23 | -1 | 4.38 | 1959.56 | outside |
| 1064 | 1082 | HVTYVPAQEKNFTTAPAI | 0.8421 | 18 | ABCPRED | 1 | NON-ALLERGEN | Non-Toxin | -0.05 | -0.17 | -0.36 | 0.5 | 7.1 | 1987.5 | outside |
| 674 | 692 | YQTQTNSPRRARSVASQS | 0.1763 | 18 | ABCPRED,literature | 2 | NON-ALLERGEN | Non-Toxin | -0.47 | -1.51 | 0.3 | 3 | 11.72 | 2037.42 | outside |
| 671 | 689 | CASYQTQTNSPRRARSVA | 0.3684 | 18 | BCPREDS,literature | 2 | ALLERGEN | Non-Toxin | -0.4 | -1.03 | 0.19 | 3 | 10.77 | 1996.42 | outside |
| 817 | 835 | FIEDLLFNKVTLADAGFI | 0.5025 | 18 | ABCPRED | 1 | ALLERGEN | Non-Toxin | 0.12 | 0.98 | -0.4 | -2 | 4.03 | 2026.63 | outside |
| 445 | 461 | VGGNYNYLYRLFRKSN | 0.1574 | 16 | ABCPRED | 1 | NON-ALLERGEN | Non-Toxin | -0.26 | -0.89 | -0.29 | 3 | 10 | 1964.45 | outside |
| 17 | 35 | NLTTRTQLPPAYTNSFTR | 0.45 | 18 | ABCPRED | 1 | NON-ALLERGEN | Non-Toxin | -0.27 | -0.89 | -0.22 | 2 | 10.84 | 2081.57 | outside |
| 290 | 306 | DCALDPLSETKCTLKS | 0.8025 | 16 | ABCPRED | 1 | NON-ALLERGEN | Non-Toxin | -0.21 | -0.29 | 0.43 | -1 | 4.56 | 1724.19 | outside |
| 724 | 740 | TEILPVSMTKTSVDCT | 1.5042 | 16 | ABCPRED | 1 | NON-ALLERGEN | Non-Toxin | -0.07 | 0.26 | -0.06 | -1 | 4.38 | 1725.23 | outside |
| 19 | 37 | TTRTQLPPAYTNSFTRGV | 0.3886 | 18 | ABCPRED | 1 | NON-ALLERGEN | Non-Toxin | -0.22 | -0.7 | -0.22 | 2 | 10.84 | 2010.49 | outside |
| 1144 | 1161 | ELDSFKEELDKYFKNHT | -0.8623 | 17 | BCEPRED | 1 | NON-ALLERGEN | Non-Toxin | -0.34 | -1.5 | 0.75 | -1.5 | 4.9 | 2143.58 | outside |
| 1262 | 1274 | EPVLKGVKLHYT | 1.4118 | 12 | BCEPRED | 1 | NON-ALLERGEN | Non-Toxin | -0.1 | -0.21 | -0.07 | 1.5 | 8.84 | 1383.84 | inside |
| 786 | 801 | KQIYKTPPIKDFGGF | -0.3896 | 15 | IEDB,literature | 2 | NON-ALLERGEN | Non-Toxin | -0.13 | -0.67 | 0.06 | 2 | 9.55 | 1739.27 | outside |
| 1093 | 1111 | GVFVSNGTHWFVTQRNFY | 0.3991 | 18 | ABCPRED | 1 | NON-ALLERGEN | Non-Toxin | -0.03 | -0.13 | -0.84 | 1.5 | 9.1 | 2159.66 | outside |
| 13 | 33 | SQCVNLTTRTQLPPAYTNSF | 0.9351 | 20 | ABCPRED | 1 | ALLERGEN | Non-Toxin | -0.16 | -0.42 | -0.43 | 1 | 8.57 | 2241.79 | outside |
| 992 | 1010 | QIDRLITGRLQSLQTYVT | -0.3906 | 18 | ABCPRED | 1 | ALLERGEN | Non-Toxin | -0.19 | -0.17 | -0.23 | 1 | 9.1 | 2105.71 | outside |
| 857 | 875 | GLTVLPPLLTDEMIAQYT | 0.4125 | 18 | ABCPRED | 1 | NON-ALLERGEN | Non-Toxin | 0.08 | 0.56 | -0.53 | -2 | 3.67 | 1975.61 | outside |

| | | | | | | | | | | | | | | | |
|---|---|---|---|---|---|---|---|---|---|---|---|---|---|---|---|
| 99 | 109 | NIIRGWIFGT | -0.5963 | 10 | ABCPRED | 1 | ALLERGEN | Non-Toxin | 0.09 | 0.59 | -0.85 | 1 | 10.11 | 1176.54 | outside |
| 899 | 917 | AMQMAYRFNGIGVTQNVL | 0.8921 | 18 | AAP | 1 | NON-ALLERGEN | Non-Toxin | -0.02 | 0.31 | -0.64 | 1 | 9.1 | 2013.64 | outside |
| 286 | 302 | TDAVDCALDPLSETKC | 0.8013 | 16 | ABCPRED | 1 | ALLERGEN | Non-Toxin | -0.15 | -0.08 | 0.4 | -3 | 3.84 | 1681.07 | outside |
| 771 | 785 | AVEQDKNTQEVFAQ | 0.1637 | 14 | ABCPRED,literature | 2 | ALLERGEN | Non-Toxin | -0.27 | -1.02 | 0.42 | -2 | 4.14 | 1606.91 | outside |
| 380 | 392 | YGVSPTKLNDLC | 1.8347 | 12 | ABCPRED,AAP | 2 | ALLERGEN | Non-Toxin | -0.1 | -0.12 | -0.19 | 0 | 6.15 | 1309.67 | outside |
| 982 | 989 | SRLDKVE | 0.0966 | 7 | IEDB | 1 | ALLERGEN | Non-Toxin | -0.48 | -1.17 | 1.29 | 0 | 6.42 | 846.04 | outside |
| 277 | 293 | LKYNENGTITDAVDCA | 0.7352 | 16 | ABCPRED | 1 | NON-ALLERGEN | Non-Toxin | -0.14 | -0.37 | 0.14 | -2 | 4.03 | 1727.1 | outside |
| 160 | 180 | YSSANNCTFEYVSQPFLMDL | 0.2507 | 20 | ABCPRED | 1 | ALLERGEN | Non-Toxin | -0.05 | -0.06 | -0.52 | -2 | 3.67 | 2329.86 | outside |
| 731 | 747 | MTKTSVDCTMYICGDS | -0.0634 | 16 | ABCPRED | 1 | ALLERGEN | Non-Toxin | -0.1 | 0.07 | -0.11 | -1 | 4.21 | 1755.24 | outside |
| 293 | 303 | LDPLSETKCT | 1.2586 | 10 | ABCPRED | 1 | ALLERGEN | Non-Toxin | -0.2 | -0.46 | 0.39 | -1 | 4.38 | 1106.39 | outside |
| 229 | 241 | LPIGINITRFQT | 0.9919 | 12 | ABCPRED | 1 | NON-ALLERGEN | Non-Toxin | 0 | 0.43 | -0.59 | 1 | 10.11 | 1372.82 | outside |
| 314 | 324 | QTSNFRVQPT | 1.405 | 10 | ABCPRED,IEDB | 2 | ALLERGEN | Non-Toxin | -0.33 | -1.18 | -0.09 | 1 | 10.11 | 1177.42 | outside |
| 993 | 1013 | IDRLITGRLQSLQTYVTQQL | -0.371 | 20 | ABCPRED | 1 | NON-ALLERGEN | Non-Toxin | -0.17 | -0.14 | -0.28 | 1 | 9.1 | 2347.04 | outside |
| 761 | 777 | TQLNRALTGIAVEQDK | 0.4291 | 16 | ABCPRED | 1 | NON-ALLERGEN | Non-Toxin | -0.22 | -0.49 | 0.24 | 0 | 6.42 | 1757.22 | outside |
| 147 | 159 | KNNKSWMESEFR | 0.6205 | 12 | ABCPRED | 1 | NON-ALLERGEN | Non-Toxin | -0.48 | -2.01 | 0.73 | 1 | 8.93 | 1555.89 | outside |
| 138 | 154 | DPFLGVYYHKNNKSWM | 0.6956 | 16 | ABCPRED | 1 | ALLERGEN | Non-Toxin | -0.15 | -0.94 | -0.37 | 1.5 | 8.76 | 1999.51 | outside |
| 137 | 149 | NDPFLGVYYHKN | 0.9849 | 12 | ABCPRED | 1 | ALLERGEN | Non-Toxin | -0.14 | -0.95 | -0.37 | 0.5 | 7.09 | 1466.79 | outside |
| 668 | 686 | AGICASYQTQTNSPRRAR | 0.4638 | 18 | ABCPRED | 1 | NON-ALLERGEN | Non-Toxin | -0.36 | -0.99 | 0.16 | 3 | 10.77 | 1980.43 | outside |
| 345 | 357 | TRFASVYAWNRK | 0.4594 | 12 | ABCPRED,FBCPred | 2 | NON-ALLERGEN | Non-Toxin | -0.3 | -0.79 | -0.13 | 3 | 11.01 | 1498.86 | outside |
| 748 | 768 | ECSNLLLQYGSFCTQLNRAL | 0.626 | 20 | ABCPRED | 1 | NON-ALLERGEN | Non-Toxin | -0.1 | 0.13 | -0.47 | 0 | 6.29 | 2273.91 | outside |
| 450 | 462 | NYLYRLFRKSNL | 0.0104 | 12 | ABCPRED | 1 | NON-ALLERGEN | Non-Toxin | -0.33 | -0.76 | -0.23 | 3 | 10.29 | 1587.03 | outside |
| 902 | 919 | MAYRFNGIGVTQNVLYE | 0.759 | 17 | COVIDep,literature | 2 | NON-ALLERGEN | Non-Toxin | -0.04 | 0.04 | -0.55 | 0 | 6.35 | 1975.51 | outside |
| 268 | 278 | GYLQPRTFLL | 0.7535 | 10 | ABCPRED | 1 | ALLERGEN | Non-Toxin | -0.03 | 0.22 | -0.74 | 1 | 9.1 | 1207.59 | outside |
| 1153 | 1171 | DKYFKNHTSPDVDLGDIS | 0.847 | 18 | ABCPRED | 1 | ALLERGEN | Non-Toxin | -0.24 | -1.04 | 0.44 | -1.5 | 4.56 | 2051.45 | outside |
| 496 | 512 | GFQPTNGVGYQPYRVV | 0.7343 | 16 | ABCPRED | 1 | NON-ALLERGEN | Non-Toxin | -0.08 | -0.46 | -0.52 | 1 | 8.93 | 1782.23 | outside |
| 207 | 217 | HTPINLVRDL | 0.1368 | 10 | ABCPRED | 1 | NON-ALLERGEN | Non-Toxin | -0.14 | -0.07 | -0.16 | 0.5 | 7.1 | 1177.52 | outside |
| 278 | 290 | KYNENGTITDAV | 0.5998 | 12 | ABCPRED | 1 | NON-ALLERGEN | Non-Toxin | -0.2 | -0.87 | 0.21 | -1 | 4.38 | 1324.58 | outside |
| 13 | 38 | SQCVNLTTRTQLPPAYTNSFTRGVY | 0.686 | 25 | IEDB,literature | 2 | NON-ALLERGEN | Non-Toxin | -0.18 | -0.45 | -0.39 | 2 | 9.36 | 2818.52 | outside |
| 773 | 791 | EQDKNTQEVFAQVKQIYK | 0.105 | 18 | FBCPred | 1 | NON-ALLERGEN | Toxin | -0.34 | -1.34 | 0.47 | 0 | 6.52 | 2196.72 | outside |
| 972 | 989 | AISSVLNDILSRLDKVE | -0.0619 | 17 | COVIDep,literature | 2 | ALLERGEN | Non-Toxin | -0.12 | 0.34 | 0.21 | -1 | 4.56 | 1872.4 | outside |
| 733 | 749 | KTSVDCTMYICGDSTE | 0.029 | 16 | ABCPRED | 1 | ALLERGEN | Non-Toxin | -0.15 | -0.26 | 0.16 | -2 | 4.03 | 1753.16 | outside |
| 381 | 401 | GVSPTKLNDLCFTNVYADSF | 1.2617 | 20 | ABCPRED | 1 | NON-ALLERGEN | Non-Toxin | -0.05 | 0.08 | -0.31 | -1 | 4.21 | 2191.72 | outside |
| 1042 | 1058 | FCGKGYHLMSFPQSAP | 0.2518 | 16 | ABCPRED | 1 | ALLERGEN | Non-Toxin | -0.01 | -0.12 | -0.54 | 1.5 | 8.54 | 1770.28 | outside |
| 24 | 40 | LPPAYTNSFTRGVYYP | 0.2263 | 16 | ABCPRED | 1 | NON-ALLERGEN | Toxin | -0.07 | -0.42 | -0.66 | 1 | 8.83 | 1846.29 | outside |
| 72 | 82 | GTNGTKRFDN | 0.6665 | 10 | ABCPRED | 1 | NON-ALLERGEN | Non-Toxin | -0.43 | -1.83 | 0.61 | 1 | 9.1 | 1109.3 | outside |
| 103 | 123 | GWIFGTTLDSKTQSLLIVNN | 0.5113 | 20 | ABCPRED | 1 | NON-ALLERGEN | Non-Toxin | 0 | 0.2 | -0.52 | 0 | 6.19 | 2207.83 | outside |
| 1063 | 1079 | LHVTYVPAQEKNFTTA | 1.0276 | 16 | ABCPRED | 1 | NON-ALLERGEN | Non-Toxin | -0.08 | -0.25 | -0.37 | 0.5 | 7.1 | 1819.28 | outside |
| 790 | 800 | KTPPIKDFGG | -0.6798 | 10 | ABCPRED | 1 | ALLERGEN | Non-Toxin | -0.16 | -0.87 | 0.43 | 1 | 8.94 | 1059.37 | outside |
| 418 | 434 | IADYNYKLPDDFTGCV | 0.3963 | 16 | ABCPRED | 1 | ALLERGEN | Non-Toxin | -0.08 | -0.22 | -0.12 | -2 | 3.94 | 1834.25 | outside |
| 1069 | 1078 | PAQEKNFTT | 0.382 | 9 | IEDB | 1 | ALLERGEN | Non-Toxin | -0.29 | -1.42 | 0.29 | 0 | 6.35 | 1035.24 | outside |
| 725 | 741 | EILPVSMTKTSVDCTM | 1.5372 | 16 | ABCPRED | 1 | NON-ALLERGEN | Non-Toxin | -0.04 | 0.43 | -0.11 | -1 | 4.38 | 1755.32 | outside |
| 377 | 393 | FKCYGVSPTKLNDLCF | 2.0332 | 16 | ABCPRED | 1 | ALLERGEN | Non-Toxin | -0.06 | 0.17 | -0.33 | 1 | 8.34 | 1835.39 | outside |
| 1094 | 1110 | VFVSNGTHWFVTQRNF | 0.3321 | 16 | ABCPRED | 1 | ALLERGEN | Non-Toxin | -0.05 | -0.04 | -0.8 | 1.5 | 10.11 | 1939.4 | outside |
| 784 | 800 | QVKQIYKTPPIKDFGG | -0.1495 | 16 | ABCPRED | 1 | ALLERGEN | Non-Toxin | -0.17 | -0.76 | 0.13 | 2 | 9.55 | 1819.38 | outside |
| 805 | 817 | ILPDPSKPSKRS | 0.5322 | 12 | IEDB,literature | 2 | NON-ALLERGEN | Non-Toxin | -0.37 | -1.22 | 0.78 | 2 | 10.01 | 1324.7 | outside |
| 615 | 635 | VNCTEVPVAIHADQLTPTWR | 0.6253 | 20 | ABCPRED | 1 | NON-ALLERGEN | Non-Toxin | -0.09 | -0.04 | -0.29 | -0.5 | 5.33 | 2250.85 | outside |
| 276 | 288 | LLKYNENGTITD | 0.6261 | 12 | ABCPRED | 1 | NON-ALLERGEN | Non-Toxin | -0.18 | -0.74 | 0.08 | -1 | 4.38 | 1380.7 | outside |
| 263 | 279 | AAYYVGYLQPRTFLLK | 0.383 | 16 | ABCPRED | 1 | ALLERGEN | Non-Toxin | -0.02 | 0.22 | -0.72 | 2 | 9.55 | 1903.49 | outside |
| 294 | 310 | DPLSETKCTLKSFTVE | 0.6579 | 16 | ABCPRED | 1 | ALLERGEN | Non-Toxin | -0.19 | -0.41 | 0.36 | -1 | 4.68 | 1798.26 | outside |
| 495 | 513 | YGFQPTNGVGYQPYRVVV | 0.6768 | 18 | ABCPRED,BCPREDS | 2 | NON-ALLERGEN | Non-Toxin | -0.04 | -0.24 | -0.68 | 1 | 8.83 | 2044.57 | outside |
| 513 | 533 | LSFELLHAPATVCGPKKSTN | 0.4763 | 20 | ABCPRED,AAP | 2 | ALLERGEN | Non-Toxin | -0.07 | -0.01 | -0.14 | 1.5 | 8.54 | 2113.75 | outside |
| 642 | 660 | VFQTRAGCLIGAEHVNNS | 1.1368 | 18 | ABCPRED | 1 | NON-ALLERGEN | Non-Toxin | -0.08 | 0.09 | -0.28 | 0.5 | 7.07 | 1916.41 | outside |
| 1201 | 1214 | QELGKYEQYIKWP | 0.3709 | 13 | BCEPRED | 1 | NON-ALLERGEN | Non-Toxin | -0.24 | -1.46 | 0.06 | 0 | 6.48 | 1682.11 | outside_transmembrane-region |
| 21 | 33 | RTQLPPAYTNSF | 0.5931 | 12 | ABCPRED | 1 | NON-ALLERGEN | Non-Toxin | -0.2 | -0.82 | -0.35 | 1 | 9.1 | 1394.71 | outside |
| 436 | 446 | WNSNNLDSKV | 0.7059 | 10 | ABCPRED | 1 | NON-ALLERGEN | Non-Toxin | -0.28 | -1.24 | 0.05 | 0 | 6.19 | 1176.39 | outside |
| 967 | 985 | SSNFGAISSVLNDILSRL | 0.1356 | 18 | ABCPRED | 1 | NON-ALLERGEN | Non-Toxin | -0.02 | 0.54 | -0.31 | 0 | 6.19 | 1893.39 | outside |
| 413 | 429 | GQTGKIADYNYKLPDD | 1.0071 | 16 | ABCPRED | 1 | ALLERGEN | Non-Toxin | -0.25 | -1.31 | 0.39 | -1 | 4.43 | 1798.17 | outside |
| 443 | 461 | SKVGGNYNYLYRLFRKSN | 0.156 | 18 | ABCPRED,AAP | 2 | NON-ALLERGEN | Non-Toxin | -0.31 | -1.06 | -0.07 | 4 | 10.17 | 2179.73 | outside |
| 374 | 390 | FSTFKCYGVSPTKLND | 0.965 | 16 | ABCPRED | 1 | NON-ALLERGEN | Non-Toxin | -0.12 | -0.31 | -0.16 | 1 | 8.51 | 1807.27 | outside |
| 1257 | 1273 | DEDDSEPVLKGVKLHY | 1.0101 | 16 | ABCPRED | 1 | NON-ALLERGEN | Toxin | -0.25 | -1.04 | 0.74 | -2.5 | 4.43 | 1844.24 | inside |
| 969 | 985 | NFGAISSVLNDILSRL | 0.1779 | 16 | ABCPRED | 1 | NON-ALLERGEN | Non-Toxin | 0 | 0.71 | -0.39 | 0 | 6.19 | 1719.21 | outside |
| 98 | 110 | SNIIRGWIFGTT | -0.3537 | 12 | ABCPRED | 1 | NON-ALLERGEN | Non-Toxin | 0.04 | 0.37 | -0.72 | 1 | 10.11 | 1364.75 | outside |
| 448 | 464 | NYNYLYRLFRKSNLKP | 0.4204 | 16 | ABCPRED | 1 | NON-ALLERGEN | Non-Toxin | -0.36 | -1.21 | -0.12 | 4 | 10.17 | 2089.66 | outside |
| 1003 | 1020 | SLQTYVTQQLIRAAEIR | -0.012 | 17 | COVIDep,literature | 2 | ALLERGEN | Non-Toxin | -0.19 | -0.12 | -0.17 | 1 | 9.1 | 1990.55 | outside |
| 404 | 416 | GDEVRQIAPGQT | 0.7073 | 12 | ABCPRED | 1 | ALLERGEN | Non-Toxin | -0.24 | -0.92 | 0.43 | -1 | 4.38 | 1270.54 | outside |
| 674 | 686 | YQTQTNSPRRAR | 0.076 | 12 | BCPREDS,IEDB,literature | 3 | NON-ALLERGEN | Non-Toxin | -0.64 | -2.27 | 0.53 | 3 | 11.72 | 1477.76 | outside |
| 503 | 512 | VGYQPYRVV | 1.4383 | 9 | BCEPRED | 1 | ALLERGEN | Non-Toxin | -0.08 | 0 | -0.66 | 1 | 8.93 | 1080.38 | outside |
| 383 | 395 | SPTKLNDLCFTN | 2.0157 | 12 | literature | 1 | ALLERGEN | Non-Toxin | -0.17 | -0.44 | -0.1 | 0 | 6.16 | 1352.69 | outside |
| 989 | 1005 | AEVQIDRLITGRLQSL | -0.2062 | 16 | ABCPRED | 1 | NON-ALLERGEN | Non-Toxin | -0.17 | 0.09 | 0.08 | 0 | 6.42 | 1812.35 | outside |
| 722 | 740 | VTTEILPVSMTKTSVDCT | 1.3791 | 18 | ABCPRED,AAP,FBCPred | 3 | NON-ALLERGEN | Non-Toxin | -0.04 | 0.43 | -0.16 | -1 | 4.38 | 1925.5 | outside |
| 404 | 418 | GDEVRQIAPGQTGK | 1.1397 | 14 | ABCPRED,AAP,FBCPred | 3 | NON-ALLERGEN | Non-Toxin | -0.27 | -1.1 | 0.59 | 0 | 6.42 | 1455.8 | outside |
| 442 | 458 | DSKVGGNYNYLYRLFR | 0.018 | 16 | ABCPRED | 1 | NON-ALLERGEN | Non-Toxin | -0.27 | -0.89 | -0.11 | 2 | 9.55 | 1965.43 | outside |
| 638 | 656 | TGSNVFQTRAGCLIGAEH | 0.9162 | 18 | ABCPRED | 1 | NON-ALLERGEN | Non-Toxin | -0.07 | -0.01 | -0.23 | 0.5 | 7.07 | 1861.33 | outside |
| 768 | 784 | TGIAVEQDKNTQEVFA | 0.5614 | 16 | ABCPRED | 1 | ALLERGEN | Non-Toxin | -0.15 | -0.46 | 0.22 | -2 | 4.14 | 1750.13 | outside |
| 723 | 741 | TTEILPVSMTKTSVDCTM | 1.3906 | 18 | ABCPRED | 1 | ALLERGEN | Non-Toxin | -0.06 | 0.3 | -0.14 | -1 | 4.38 | 1957.56 | outside |
| 42 | 54 | VFRSSVLHSTQD | 0.0591 | 12 | ABCPRED | 1 | NON-ALLERGEN | Non-Toxin | -0.19 | -0.23 | -0.09 | 0.5 | 7.1 | 1375.67 | outside |
| 293 | 305 | LDPLSETKCTLK | 0.9712 | 12 | ABCPRED | 1 | NON-ALLERGEN | Non-Toxin | -0.22 | -0.39 | 0.42 | 0 | 6.38 | 1347.76 | outside |

| | | | | | | | | | | | | | | | |
|---|---|---|---|---|---|---|---|---|---|---|---|---|---|---|---|
| 18 | 33 | LTTRTQLPPAYTNSF | 0.79 | 15 | IEDB | 1 | NON-ALLERGEN | Non-Toxin | -0.15 | -0.49 | -0.45 | 1 | 9.1 | 1710.13 | outside |
| 380 | 396 | YGVSPTKLNDLCFTNV | 2.0255 | 16 | ABCPRED | 1 | NON-ALLERGEN | Non-Toxin | -0.05 | 0.09 | -0.41 | 0 | 6.15 | 1771.25 | outside |
| 419 | 431 | ADYNYKLPDDFT | 0.8392 | 12 | ABCPRED,AAP | 2 | ALLERGEN | Non-Toxin | -0.23 | -1.2 | 0.2 | -2 | 3.94 | 1461.7 | outside |
| 228 | 246 | DLPIGINITRFQTLLALH | 0.9638 | 18 | ABCPRED | 1 | NON-ALLERGEN | Toxin | 0.04 | 0.65 | -0.58 | 0.5 | 7.1 | 2035.71 | outside |
| 378 | 396 | KCYGVSPTKLNDLCFTNV | 1.8687 | 18 | ABCPRED | 1 | NON-ALLERGEN | Non-Toxin | -0.1 | 0 | -0.25 | 1 | 8.34 | 2002.59 | outside |
| 1012 | 1030 | LIRAAEIRASANLAATKM | 0.4049 | 18 | ABCPRED | 1 | ALLERGEN | Non-Toxin | -0.11 | 0.44 | 0.03 | 2 | 10.84 | 1900.52 | outside |
| 1068 | 1077 | VPAQEKNFT | 1.0107 | 9 | IEDB | 1 | ALLERGEN | Non-Toxin | -0.21 | -0.88 | 0.17 | 0 | 6.35 | 1033.27 | outside |
| 1155 | 1171 | YFKNHTSPDVDLGDIS | 1.1715 | 16 | ABCPRED | 1 | ALLERGEN | Non-Toxin | -0.15 | -0.71 | 0.12 | -1.5 | 4.42 | 1808.16 | outside |
| 789 | 801 | YKTPPIKDFGGF | -0.3433 | 12 | ABCPRED,BCPREDS | 2 | ALLERGEN | Non-Toxin | -0.08 | -0.6 | -0.04 | 1 | 8.83 | 1369.75 | outside |
| 410 | 424 | IAPGQTGKIADYNY | 1.724 | 14 | ABCPRED,BCPREDS | 2 | NON-ALLERGEN | Non-Toxin | -0.08 | -0.54 | -0.23 | 0 | 6.18 | 1510.87 | outside |
| 154 | 174 | ESEFRVYSSANNCTFEYVSQ | 0.1805 | 20 | ABCPRED,FBCPred | 2 | NON-ALLERGEN | Non-Toxin | -0.21 | -0.68 | -0.03 | -2 | 4.26 | 2360.76 | outside |
| 1146 | 1162 | DSFKEELDKYFKNHTS | -0.9728 | 16 | ABCPRED | 1 | NON-ALLERGEN | Non-Toxin | -0.37 | -1.66 | 0.74 | -0.5 | 5.5 | 1988.36 | outside |
| 1190 | 1208 | AKNLNESLIDLQELGKYE | 0.7136 | 18 | ABCPRED | 1 | NON-ALLERGEN | Non-Toxin | -0.21 | -0.74 | 0.39 | -2 | 4.41 | 2077.6 | outside |
| 1153 | 1173 | DKYFKNHTSPDVDLGDISGI | 0.5734 | 20 | ABCPRED,literature | 2 | NON-ALLERGEN | Non-Toxin | -0.17 | -0.73 | 0.31 | -1.5 | 4.56 | 2221.7 | outside |
| 1154 | 1172 | KYFKNHTSPDVDLGDISG | 0.5442 | 18 | ABCPRED | 1 | ALLERGEN | Non-Toxin | -0.19 | -0.87 | 0.28 | -0.5 | 5.31 | 1993.42 | outside |
| 454 | 464 | RLFRKSNLKP | 0.3804 | 10 | ABCPRED | 1 | NON-ALLERGEN | Non-Toxin | -0.5 | -1.23 | 0.64 | 4 | 12.02 | 1258.67 | outside |
| 816 | 828 | SFIEDLLFNKVT | -0.033 | 12 | ABCPRED | 1 | NON-ALLERGEN | Non-Toxin | 0 | 0.5 | -0.23 | -1 | 4.38 | 1425.82 | outside |
| 17 | 33 | NLTTRTQLPPAYTNSF | 0.7508 | 16 | ABCPRED | 1 | NON-ALLERGEN | Non-Toxin | -0.18 | -0.68 | -0.41 | 1 | 9.1 | 1824.25 | outside |
| 172 | 182 | SQPFLMDLEG | 0.1895 | 10 | ABCPRED | 1 | ALLERGEN | Non-Toxin | -0.03 | -0.1 | -0.09 | -2 | 3.67 | 1136.43 | outside |
| 384 | 395 | PTKLNDLCFTN | 2.1999 | 11 | literature | 1 | ALLERGEN | Non-Toxin | -0.17 | -0.41 | -0.14 | 0 | 6.16 | 1265.6 | outside |
| 414 | 430 | QTGKIADYNYKLPDDF | 0.986 | 16 | ABCPRED | 1 | NON-ALLERGEN | Non-Toxin | -0.23 | -1.11 | 0.24 | -1 | 4.43 | 1888.29 | outside |
| 493 | 513 | QSYGFQPTNGVGYQPYRVVV | 0.6459 | 20 | ABCPRED,BCPREDS | 2 | ALLERGEN | Non-Toxin | -0.09 | -0.44 | -0.58 | 1 | 8.83 | 2259.81 | outside |
| 672 | 685 | ASYQTQTNSPRRA | 0.107 | 13 | BCEPRED | 1 | NON-ALLERGEN | Non-Toxin | -0.46 | -1.68 | 0.24 | 2 | 10.84 | 1479.74 | outside |
| 384 | 394 | PTKLNDLCFT | 2.5246 | 10 | ABCPRED | 1 | ALLERGEN | Non-Toxin | -0.12 | -0.1 | -0.17 | 0 | 6.16 | 1151.48 | outside |
| 673 | 691 | SYQTQTNSPRRARSVASQ | 0.2129 | 18 | ABCPRED | 1 | NON-ALLERGEN | Non-Toxin | -0.47 | -1.51 | 0.3 | 3 | 11.72 | 2037.42 | outside |
| 339 | 351 | GEVFNATRFASV | 0.1647 | 12 | ABCPRED | 1 | ALLERGEN | Non-Toxin | -0.04 | 0.35 | -0.24 | 0 | 6.36 | 1297.59 | outside |
| 35 | 45 | GVYYPDKVFR | -0.1378 | 10 | ABCPRED | 1 | NON-ALLERGEN | Non-Toxin | -0.18 | -0.53 | -0.11 | 1 | 8.83 | 1243.56 | outside |
| 264 | 276 | AYYVGYLQPRTF | 0.8536 | 12 | ABCPRED | 1 | NON-ALLERGEN | Non-Toxin | -0.05 | -0.17 | -0.87 | 1 | 8.83 | 1477.85 | outside |
| 134 | 152 | QFCNDPFLGVYYHKNNKS | 0.5089 | 18 | ABCPRED | 1 | NON-ALLERGEN | Non-Toxin | -0.21 | -0.99 | -0.24 | 1.5 | 8.48 | 2174.68 | outside |
| 129 | 139 | KVCEFQFCND | -1.0165 | 10 | ABCPRED | 1 | ALLERGEN | Non-Toxin | -0.19 | -0.31 | 0.09 | -1 | 4.38 | 1232.52 | outside |
| 1251 | 1265 | GSCCKFDEDDSEPV | 0.344 | 14 | AAP | 1 | NON-ALLERGEN | Non-Toxin | -0.26 | -0.93 | 0.9 | -4 | 3.77 | 1530.77 | inside |
| 228 | 244 | DLPIGINITRFQTLLA | 0.8861 | 16 | ABCPRED | 1 | NON-ALLERGEN | Non-Toxin | 0.03 | 0.69 | -0.51 | 0 | 6.19 | 1785.37 | outside |
| 338 | 348 | FGEVFNATRF | -0.0941 | 10 | ABCPRED | 1 | NON-ALLERGEN | Non-Toxin | -0.04 | 0.18 | -0.37 | 0 | 6.36 | 1187.45 | outside |
| 1200 | 1214 | LQELGKYEQYIKWP | 0.3532 | 14 | AAP | 1 | NON-ALLERGEN | Non-Toxin | -0.18 | -1.09 | -0.07 | 0 | 6.48 | 1795.29 | outside |
| 323 | 339 | TESIVRFPNITNLCPF | 0.5756 | 16 | ABCPRED | 1 | NON-ALLERGEN | Non-Toxin | -0.04 | 0.29 | -0.44 | 0 | 6.32 | 1851.38 | outside |
| 546 | 566 | LTGTGVLTESNKKFLPFQQF | 0.6894 | 20 | ABCPRED | 1 | NON-ALLERGEN | Non-Toxin | -0.07 | -0.16 | -0.28 | 1 | 8.94 | 2255.91 | outside |
| 781 | 799 | VFAQVKQIYKTPPIKDFG | 0.2068 | 18 | ABCPRED | 1 | NON-ALLERGEN | Non-Toxin | -0.09 | -0.17 | -0.13 | 2 | 9.55 | 2079.74 | outside |
| 313 | 325 | YQTSNFRVQPTE | 1.1569 | 12 | literature | 1 | ALLERGEN | Non-Toxin | -0.33 | -1.38 | -0.02 | 0 | 6.35 | 1469.74 | outside |
| 1156 | 1169 | FKNHTSPDVDLGD | 1.0616 | 13 | IEDB | 1 | NON-ALLERGEN | Non-Toxin | -0.23 | -1.06 | 0.45 | -1.5 | 4.42 | 1444.7 | outside |
| 412 | 424 | PGQTGKIADYNY | 1.642 | 12 | ABCPRED | 1 | NON-ALLERGEN | Non-Toxin | -0.17 | -1.15 | -0.07 | 0 | 6.18 | 1326.6 | outside |
| 1073 | 1087 | KNFTTAPAICHDGK | 0.0999 | 14 | AAP | 1 | ALLERGEN | Non-Toxin | -0.17 | -0.57 | 0.11 | 1.5 | 8.54 | 1502.9 | outside |
| 122 | 140 | NATNVVIKVCEFQFCNDP | -0.1889 | 18 | ABCPRED | 1 | NON-ALLERGEN | Non-Toxin | -0.08 | 0.13 | -0.24 | -1 | 4.38 | 2041.58 | outside |
| 863 | 881 | PLLTDEMIAQYTSALLAG | 0.1924 | 18 | ABCPRED | 1 | ALLERGEN | Non-Toxin | 0.07 | 0.61 | -0.47 | -2 | 3.67 | 1907.48 | outside |
| 412 | 428 | PGQTGKIADYNYKLPD | 1.4039 | 16 | ABCPRED | 1 | NON-ALLERGEN | Non-Toxin | -0.21 | -1.19 | 0.21 | 0 | 6.3 | 1780.2 | outside |
| 21 | 31 | RTQLPPAYTN | 1.1291 | 10 | ABCPRED | 1 | NON-ALLERGEN | Toxin | -0.28 | -1.18 | -0.2 | 1 | 9.1 | 1160.43 | outside |
| 452 | 462 | LYRLFRKSNL | -0.1612 | 10 | ABCPRED | 1 | ALLERGEN | Non-Toxin | -0.33 | -0.43 | -0.07 | 3 | 11.01 | 1309.72 | outside |
| 1194 | 1210 | NESLIDLQELGKYEQY | 0.7595 | 16 | ABCPRED | 1 | NON-ALLERGEN | Non-Toxin | -0.21 | -1.02 | 0.26 | -3 | 4.01 | 1942.36 | outside |
| 904 | 922 | YRFNGIGVTQNVLYENQK | 0.8334 | 18 | BCPREDS | 1 | ALLERGEN | Non-Toxin | -0.2 | -0.78 | -0.23 | 1 | 8.83 | 2143.67 | outside |
| 379 | 391 | CYGVSPTKLNDL | 1.4808 | 12 | ABCPRED | 1 | ALLERGEN | Non-Toxin | -0.1 | -0.12 | -0.19 | 0 | 6.15 | 1309.67 | outside |
| 448 | 460 | NYNYLYRLFRKS | -0.1108 | 12 | ABCPRED | 1 | NON-ALLERGEN | Non-Toxin | -0.37 | -1.18 | -0.27 | 3 | 10 | 1637.04 | outside |
| 340 | 352 | EVFNATRFASVY | 0.0753 | 12 | ABCPRED | 1 | ALLERGEN | Non-Toxin | -0.05 | 0.27 | -0.43 | 0 | 6.35 | 1403.71 | outside |
| 171 | 183 | VSQPFLMDLEGK | 0.518 | 12 | ABCPRED | 1 | ALLERGEN | Non-Toxin | -0.07 | -0.06 | 0.05 | -1 | 4.38 | 1363.77 | outside |
| 447 | 463 | GNYNYLYRLFRKSNLK | 0.0625 | 16 | ABCPRED | 1 | NON-ALLERGEN | Non-Toxin | -0.34 | -1.14 | -0.12 | 4 | 10.17 | 2049.6 | outside |
| 25 | 39 | PPAYTNSFTRGVYY | -0.006 | 14 | ABCPRED,literature | 2 | NON-ALLERGEN | Non-Toxin | -0.11 | -0.64 | -0.62 | 1 | 8.83 | 1635.98 | outside |
| 760 | 776 | CTQLNRALTGIAVEQD | 0.6579 | 16 | ABCPRED | 1 | NON-ALLERGEN | Non-Toxin | -0.15 | -0.09 | -0.01 | -1 | 4.38 | 1732.18 | outside |
| 1145 | 1163 | LDSFKEELDKYFKNHTSP | -0.6917 | 18 | ABCPRED | 1 | NON-ALLERGEN | Non-Toxin | -0.3 | -1.36 | 0.56 | -0.5 | 5.5 | 2198.67 | outside |
| 383 | 399 | SPTKLNDLCFTNVYAD | 1.3666 | 16 | ABCPRED | 1 | NON-ALLERGEN | Non-Toxin | -0.12 | -0.26 | -0.16 | -1 | 4.21 | 1801.22 | outside |
| 772 | 784 | VEQDKNTQEVFA | 0.1188 | 12 | AAP,BCPREDS | 2 | NON-ALLERGEN | Non-Toxin | -0.28 | -1.05 | 0.52 | -2 | 4.14 | 1407.67 | outside |
| 381 | 397 | GVSPTKLNDLCFTNVY | 1.9127 | 16 | ABCPRED | 1 | NON-ALLERGEN | Non-Toxin | -0.05 | 0.09 | -0.41 | 0 | 6.15 | 1771.25 | outside |
| 789 | 799 | YKTPPIKDFG | 0.02 | 10 | ABCPRED,IEDB | 2 | NON-ALLERGEN | Non-Toxin | -0.17 | -0.96 | 0.2 | 1 | 8.83 | 1165.49 | outside |
| 1044 | 1062 | GKGYHLMSFPQSAPHGVV | 0.2218 | 18 | ABCPRED | 1 | NON-ALLERGEN | Non-Toxin | 0 | -0.13 | -0.48 | 2 | 8.94 | 1912.47 | outside |
| 991 | 1009 | VQIDRLITGRLQSLQTYV | -0.388 | 18 | ABCPRED | 1 | ALLERGEN | Non-Toxin | -0.15 | 0.11 | -0.29 | 1 | 9.1 | 2103.74 | outside |
| 1096 | 1110 | VSNGTHWFVTQRNF | 0.4144 | 14 | ABCPRED,BCPREDS | 2 | NON-ALLERGEN | Non-Toxin | -0.14 | -0.55 | -0.63 | 1.5 | 10.11 | 1693.06 | outside |
| 550 | 566 | GVLTESNKKFLPFQQF | 0.5985 | 16 | ABCPRED | 1 | NON-ALLERGEN | Non-Toxin | -0.11 | -0.32 | -0.19 | 1 | 8.94 | 1883.42 | outside |
| 791 | 803 | TPPIKDFGGFNF | 0.2723 | 12 | ABCPRED | 1 | ALLERGEN | Non-Toxin | 0.01 | -0.23 | -0.29 | 0 | 6.19 | 1339.68 | outside |
| 500 | 514 | TNGVGYQPYRVVVL | 0.5763 | 14 | ABCPRED,FBCPred | 2 | NON-ALLERGEN | Non-Toxin | -0.02 | 0.24 | -0.67 | 1 | 8.93 | 1565.02 | outside |
| 646 | 662 | RAGCLIGAEHVNNSYE | 1.1613 | 16 | ABCPRED | 1 | NON-ALLERGEN | Non-Toxin | -0.14 | -0.38 | -0.01 | -0.5 | 5.41 | 1733.12 | outside |
| 642 | 658 | VFQTRAGCLIGAEHVN | 1.2576 | 16 | ABCPRED | 1 | NON-ALLERGEN | Non-Toxin | -0.03 | 0.37 | -0.35 | 0.5 | 7.07 | 1715.2 | outside |
| 26 | 38 | PAYTNSFTRGVY | -0.2209 | 12 | ABCPRED | 1 | NON-ALLERGEN | Non-Toxin | -0.12 | -0.5 | -0.53 | 1 | 8.93 | 1375.66 | outside |
| 988 | 1005 | EAEVQIDRLITGRLQSL | -0.1784 | 17 | COVIDep,literature | 2 | NON-ALLERGEN | Non-Toxin | -0.19 | -0.12 | 0.25 | -1 | 4.68 | 1941.48 | outside |
| 416 | 430 | GKIADYNYKLPDDF | 0.9776 | 14 | ABCPRED | 1 | NON-ALLERGEN | Non-Toxin | -0.2 | -0.96 | 0.29 | -1 | 4.43 | 1659.02 | outside |
| 423 | 435 | YKLPDDFTGCVI | -0.3083 | 12 | ABCPRED | 1 | NON-ALLERGEN | Non-Toxin | -0.01 | 0.24 | -0.19 | -1 | 4.21 | 1370.75 | outside |
| 508 | 520 | YRVVVLSFELLH | 1.1085 | 12 | ABCPRED | 1 | NON-ALLERGEN | Non-Toxin | 0.07 | 1.13 | -0.74 | 0.5 | 7.1 | 1474.95 | outside |
| 1064 | 1084 | HVTYVPAQEKNFTTAPAICH | 0.7635 | 20 | ABCPRED,FBCPred | 2 | NON-ALLERGEN | Non-Toxin | -0.07 | -0.19 | -0.39 | 1 | 7.22 | 2227.81 | outside |
| 816 | 834 | SFIEDLLFNKVTLADAGF | 0.5167 | 18 | ABCPRED | 1 | ALLERGEN | Non-Toxin | 0.06 | 0.68 | -0.28 | -2 | 4.03 | 2000.54 | outside |
| 562 | 580 | FQQFGRDIADTTDAVRDP | 0.0641 | 18 | ABCPRED | 1 | NON-ALLERGEN | Non-Toxin | -0.28 | -0.86 | 0.46 | -2 | 4.14 | 2052.43 | outside |
| 230 | 242 | PIGINITRFQTL | 1.0169 | 12 | ABCPRED | 1 | NON-ALLERGEN | Non-Toxin | 0 | 0.43 | -0.59 | 1 | 10.11 | 1372.82 | outside |

| | | | | | | | | | | | | | | | |
|---|---|---|---|---|---|---|---|---|---|---|---|---|---|---|---|
| 1007 | 1023 | YVTQQLIRAAEIRASA | 0.1572 | 16 | ABCPRED | 1 | NON-ALLERGEN | Non-Toxin | -0.15 | 0.12 | -0.12 | 1 | 9.1 | 1790.28 | outside |
| 1147 | 1159 | SFKEELDKYFKN | -0.8791 | 12 | BCPREDS | 1 | NON-ALLERGEN | Non-Toxin | -0.37 | -1.53 | 0.78 | 0 | 6.52 | 1547.89 | outside |
| 771 | 789 | AVEQDKNTQEVFAQVKQI | 0.3411 | 18 | ABCPRED | 1 | NON-ALLERGEN | Non-Toxin | -0.24 | -0.72 | 0.32 | -1 | 4.68 | 2075.58 | outside |
| 494 | 514 | SYGFQPTNGVGYQPYRVVVL | 0.8007 | 20 | ABCPRED,FBCPred | 2 | NON-ALLERGEN | Non-Toxin | -0.03 | -0.07 | -0.69 | 1 | 8.83 | 2244.84 | outside |
| 547 | 565 | TGTGVLTESNKKFLPFQQ | 0.8567 | 18 | ABCPRED | 1 | ALLERGEN | Non-Toxin | -0.14 | -0.54 | -0.08 | 1 | 8.94 | 1995.54 | outside |
| 267 | 279 | VGYLQPRTFLLK | 0.523 | 12 | ABCPRED | 1 | ALLERGEN | Non-Toxin | -0.07 | 0.21 | -0.49 | 2 | 10.01 | 1434.93 | outside |
| 565 | 575 | FGRDIADTTD | 0.1447 | 10 | ABCPRED | 1 | ALLERGEN | Non-Toxin | -0.25 | -0.77 | 0.64 | -2 | 3.94 | 1110.27 | outside |
| 277 | 295 | LKYNENGTITDAVDCALD | 0.6554 | 18 | ABCPRED | 1 | NON-ALLERGEN | Non-Toxin | -0.14 | -0.31 | 0.19 | -3 | 3.84 | 1955.38 | outside |
| 777 | 795 | NTQEVFAQVKQIYKTPPI | 0.4221 | 18 | ABCPRED | 1 | ALLERGEN | Non-Toxin | -0.14 | -0.51 | -0.16 | 1 | 8.83 | 2104.71 | outside |
| 143 | 155 | VYYHKNNKSWME | 0.2319 | 12 | ABCPRED,BCPREDS | 2 | NON-ALLERGEN | Non-Toxin | -0.3 | -1.64 | -0.13 | 1.5 | 8.77 | 1598.97 | outside |
| 43 | 57 | FRSSVLHSTQDLFL | 0.2631 | 14 | AAP | 1 | NON-ALLERGEN | Non-Toxin | -0.08 | 0.24 | -0.41 | 0.5 | 7.1 | 1650.07 | outside |
| 177 | 187 | MDLEGKQGNF | 1.7852 | 10 | ABCPRED | 1 | ALLERGEN | Non-Toxin | -0.2 | -1.02 | 0.38 | -1 | 4.38 | 1138.41 | outside |
| 429 | 447 | FTGCVIAWNSNNLDSKVG | 0.5604 | 18 | ABCPRED | 1 | NON-ALLERGEN | Non-Toxin | -0.03 | 0.11 | -0.4 | 0 | 6.16 | 1925.41 | outside |
| 436 | 454 | WNSNNLDSKVGGNYNYLY | 0.7515 | 18 | ABCPRED,literature | 2 | NON-ALLERGEN | Non-Toxin | -0.18 | -1.13 | -0.43 | 0 | 6.18 | 2121.52 | outside |
| 817 | 829 | FIEDLLFNKVTL | 0.0704 | 12 | ABCPRED | 1 | ALLERGEN | Non-Toxin | 0.07 | 0.88 | -0.41 | -1 | 4.38 | 1451.91 | outside |
| 902 | 914 | MAYRFNGIGVTQ | 1.3822 | 12 | ABCPRED | 1 | NON-ALLERGEN | Non-Toxin | -0.05 | 0.08 | -0.57 | 1 | 9.1 | 1356.74 | outside |
| 515 | 533 | FELLHAPATVCGPKKSTN | 0.1994 | 18 | ABCPRED | 1 | ALLERGEN | Non-Toxin | -0.1 | -0.17 | -0.08 | 1.5 | 8.54 | 1913.48 | outside |
| 411 | 427 | APGQTGKIADYNYKLP | 1.2303 | 16 | ABCPRED | 1 | NON-ALLERGEN | Non-Toxin | -0.15 | -0.86 | -0.01 | 1 | 8.76 | 1736.19 | outside |
| 776 | 794 | KNTQEVFAQVKQIYKTPP | 0.3432 | 18 | ABCPRED,BCPREDS | 2 | NON-ALLERGEN | Non-Toxin | -0.25 | -0.98 | 0.11 | 2 | 9.55 | 2119.72 | outside |
| 262 | 278 | AAAYYVGYLQPRTFLL | 0.569 | 16 | ABCPRED | 1 | NON-ALLERGEN | Non-Toxin | 0.06 | 0.58 | -0.94 | 1 | 8.83 | 1846.39 | outside |
| 104 | 116 | WIFGTTLDSKTQ | 0.6072 | 12 | ABCPRED | 1 | NON-ALLERGEN | Non-Toxin | -0.08 | -0.33 | -0.35 | 0 | 6.19 | 1396.74 | outside |
| 162 | 180 | SANNCTFEYVSQPFLMDL | 0.2814 | 18 | ABCPRED | 1 | NON-ALLERGEN | Toxin | -0.04 | 0.05 | -0.47 | -2 | 3.67 | 2079.58 | outside |
| 858 | 876 | LTVLPPLLTDEMIAQYTS | 0.3595 | 18 | ABCPRED | 1 | NON-ALLERGEN | Non-Toxin | 0.05 | 0.54 | -0.52 | -2 | 3.67 | 2005.63 | outside |
| 207 | 219 | HTPINLVRDLPQ | -0.1335 | 12 | ABCPRED,BCEPRED | 2 | ALLERGEN | Non-Toxin | -0.18 | -0.48 | -0.12 | 0.5 | 7.1 | 1402.8 | outside |
| 394 | 406 | NVYADSFVIRGD | -0.2771 | 12 | ABCPRED | 1 | NON-ALLERGEN | Non-Toxin | -0.1 | 0 | -0.05 | -1 | 4.21 | 1355.63 | outside |
| 573 | 584 | TDAVRDPQTLE | 0.2349 | 11 | BCEPRED | 1 | ALLERGEN | Non-Toxin | -0.33 | -1.06 | 0.69 | -2 | 4.03 | 1244.47 | outside |
| 454 | 462 | RLFRKSNL | 0.2128 | 8 | BCEPRED | 1 | ALLERGEN | Non-Toxin | -0.48 | -0.85 | 0.43 | 3 | 12.01 | 1033.35 | outside |
| 375 | 393 | STFKCYGVSPTKLNDLCF | 1.7606 | 18 | ABCPRED | 1 | NON-ALLERGEN | Non-Toxin | -0.08 | 0.07 | -0.3 | 1 | 8.34 | 2023.6 | outside |
| 546 | 564 | LTGTGVLTESNKKFLPFQ | 0.941 | 18 | ABCPRED | 1 | NON-ALLERGEN | Non-Toxin | -0.08 | -0.13 | -0.19 | 1 | 8.94 | 1980.57 | outside |
| 379 | 397 | CYGVSPTKLNDLCFTNVY | 1.7815 | 18 | ABCPRED | 1 | ALLERGEN | Non-Toxin | -0.04 | 0.14 | -0.54 | 0 | 6.12 | 2037.59 | outside |
| 973 | 989 | ISSVLNDILSRLDKVE | -0.1004 | 16 | ABCPRED | 1 | NON-ALLERGEN | Non-Toxin | -0.14 | 0.25 | 0.26 | -1 | 4.56 | 1801.31 | outside |
| 988 | 1008 | EAEVQIDRLITGRLQSLQTY | -0.2507 | 20 | ABCPRED | 1 | NON-ALLERGEN | Non-Toxin | -0.21 | -0.38 | 0.09 | -1 | 4.68 | 2333.94 | outside |
| 720 | 740 | ISVTTEILPVSMTKTSVDCT | 1.4126 | 20 | ABCPRED,FBCPred | 2 | NON-ALLERGEN | Non-Toxin | -0.01 | 0.57 | -0.22 | -1 | 4.38 | 2125.77 | outside |
| 387 | 405 | LNDLCFTNVYADSFVIRG | 0.7651 | 18 | ABCPRED | 1 | NON-ALLERGEN | Non-Toxin | -0.02 | 0.48 | -0.44 | -1 | 4.21 | 2047.57 | outside |
| 1063 | 1083 | LHVTYVPAQEKNFTTAPAIC | 0.9251 | 20 | BCPREDS | 1 | NON-ALLERGEN | Non-Toxin | -0.02 | 0.16 | -0.46 | 0.5 | 7.06 | 2203.83 | outside |
| 1244 | 1262 | LKGCCSCGSCCKFDEDDS | 0.0887 | 18 | ABCPRED,BCPREDS | 2 | ALLERGEN | Non-Toxin | -0.23 | -0.33 | 0.53 | -2 | 4.23 | 1900.34 | inside |
| 633 | 651 | WRVYSTGSNVFQTRAGCL | 0.774 | 18 | ABCPRED | 1 | ALLERGEN | Non-Toxin | -0.14 | -0.15 | -0.46 | 2 | 9.55 | 2045.56 | outside |
| 773 | 789 | EQDKNTQEVFAQVKQI | 0.2993 | 16 | ABCPRED | 1 | NON-ALLERGEN | Non-Toxin | -0.32 | -1.19 | 0.49 | -1 | 4.68 | 1905.34 | outside |
| 1155 | 1174 | YFKNHTSPDVDLGDISGIN | 0.88 | 19 | IEDB | 1 | NON-ALLERGEN | Non-Toxin | -0.12 | -0.57 | 0.02 | -1.5 | 4.42 | 2092.53 | outside |
| 1097 | 1109 | SNGTHWFVTQRN | 0.2775 | 12 | BCPREDS,FBCPred | 2 | NON-ALLERGEN | Non-Toxin | -0.26 | -1.23 | -0.4 | 1.5 | 10.11 | 1446.72 | outside |
| 13 | 31 | SQCVNLTTRTQLPPAYTN | 1.2205 | 18 | ABCPRED | 1 | NON-ALLERGEN | Toxin | -0.2 | -0.58 | -0.36 | 1 | 8.57 | 2007.51 | outside |
| 731 | 745 | MTKTSVDCTMYICG | 0.196 | 14 | ABCPRED,FBCPred | 2 | NON-ALLERGEN | Non-Toxin | -0.04 | 0.39 | -0.36 | 0 | 6.12 | 1553.05 | outside |
| 402 | 414 | IRGDEVRQIAPG | 0.281 | 12 | ABCPRED | 1 | NON-ALLERGEN | Non-Toxin | -0.25 | -0.58 | 0.55 | 0 | 6.42 | 1310.65 | outside |
| 413 | 431 | GQTGKIADYNYKLPDDFT | 1.0564 | 18 | ABCPRED | 1 | ALLERGEN | Non-Toxin | -0.2 | -1.04 | 0.19 | -1 | 4.43 | 2046.48 | outside |
| 381 | 395 | GVSPTKLNDLCFTN | 2.2123 | 14 | literature | 1 | NON-ALLERGEN | Non-Toxin | -0.1 | -0.11 | -0.19 | 0 | 6.16 | 1508.91 | outside |
| 901 | 913 | QMAYRFNGIGVT | 1.4219 | 12 | ABCPRED | 1 | ALLERGEN | Non-Toxin | -0.05 | 0.08 | -0.58 | 1 | 9.1 | 1356.74 | outside |
| 782 | 802 | FAQVKQIYKTPPIKDFGGFN | 0.0841 | 20 | ABCPRED,BCPREDS | 2 | ALLERGEN | Non-Toxin | -0.1 | -0.42 | -0.16 | 2 | 9.55 | 2298.97 | outside |
| 41 | 51 | KVFRSSVLHS | -0.6506 | 10 | ABCPRED | 1 | ALLERGEN | Non-Toxin | -0.18 | 0.1 | -0.09 | 2.5 | 11.01 | 1159.49 | outside |
| 195 | 213 | KNIDGYFKIYSKHTPINL | 0.5622 | 18 | ABCPRED | 1 | NON-ALLERGEN | Non-Toxin | -0.15 | -0.63 | -0.14 | 2.5 | 9.43 | 2151.77 | outside |
| 1007 | 1025 | YVTQQLIRAAEIRASANL | 0.196 | 18 | ABCPRED | 1 | ALLERGEN | Non-Toxin | -0.14 | 0.12 | -0.19 | 1 | 9.1 | 2017.58 | outside |
| 321 | 339 | QPTESIVRFPNITNLCPF | 0.4966 | 18 | ABCPRED | 1 | NON-ALLERGEN | Non-Toxin | -0.08 | -0.02 | -0.38 | 0 | 6.32 | 2076.66 | outside |
| 729 | 747 | VSMTKTSVDCTMYICGDS | 0.2082 | 18 | ABCPRED | 1 | NON-ALLERGEN | Non-Toxin | -0.07 | 0.26 | -0.17 | -1 | 4.21 | 1941.48 | outside |
| 667 | 687 | GAGICASYQTQTNSPRRARS | 0.3404 | 20 | FBCPred | 1 | ALLERGEN | Non-Toxin | -0.33 | -0.95 | 0.15 | 3 | 10.77 | 2124.59 | outside |
| 501 | 513 | NGVGYQPYRVVV | 0.5253 | 12 | ABCPRED | 1 | NON-ALLERGEN | Non-Toxin | -0.05 | 0.02 | -0.6 | 1 | 8.93 | 1350.72 | outside |
| 1185 | 1197 | RLNEVAKNLNES | 0.3486 | 12 | AAP | 1 | NON-ALLERGEN | Non-Toxin | -0.37 | -1.09 | 0.61 | 0 | 6.49 | 1386.7 | outside |
| 278 | 296 | KYNENGTITDAVDCALDP | 0.7089 | 18 | ABCPRED | 1 | NON-ALLERGEN | Non-Toxin | -0.17 | -0.61 | 0.29 | -3 | 3.84 | 1939.33 | outside |
| 1156 | 1174 | FKNHTSPDVDLGDISGIN | 0.9468 | 18 | ABCPRED | 1 | ALLERGEN | Non-Toxin | -0.12 | -0.53 | 0.15 | -1.5 | 4.42 | 1929.34 | outside |
| 418 | 436 | IADYNYKLPDDFTGCVIA | 0.4349 | 18 | ABCPRED | 1 | NON-ALLERGEN | Non-Toxin | -0.02 | 0.15 | -0.23 | -2 | 3.94 | 2018.52 | outside |
| 632 | 650 | TWRVYSTGSNVFQTRAGC | 0.6157 | 18 | ABCPRED | 1 | NON-ALLERGEN | Non-Toxin | -0.18 | -0.4 | -0.38 | 2 | 9.55 | 2033.5 | outside |
| 1062 | 1078 | FLHVTYVPAQEKNFTT | 0.9556 | 16 | ABCPRED | 1 | ALLERGEN | Non-Toxin | -0.06 | -0.19 | -0.49 | 0.5 | 7.1 | 1895.38 | outside |
| 434 | 452 | IAWNSNNLDSKVGGNYNY | 1.0683 | 18 | ABCPRED | 1 | NON-ALLERGEN | Non-Toxin | -0.15 | -0.92 | -0.33 | 0 | 6.18 | 2029.42 | outside |
| 417 | 429 | KIADYNYKLPDD | 0.8714 | 12 | ABCPRED | 1 | ALLERGEN | Non-Toxin | -0.29 | -1.32 | 0.54 | -1 | 4.43 | 1454.76 | outside |
| 259 | 277 | TAGAAAYYVGYLQPRTFL | 0.5955 | 18 | AAP | 1 | NON-ALLERGEN | Non-Toxin | 0.04 | 0.34 | -0.78 | 1 | 8.83 | 1962.49 | outside |
| 898 | 914 | FAMQMAYRFNGIGVTQ | 1.3096 | 16 | ABCPRED,literature | 2 | ALLERGEN | Non-Toxin | -0.01 | 0.24 | -0.69 | 1 | 9.1 | 1834.38 | outside |
| 730 | 746 | SMTKTSVDCTMYICGD | 0.048 | 16 | ABCPRED | 1 | ALLERGEN | Non-Toxin | -0.1 | 0.07 | -0.11 | -1 | 4.21 | 1755.24 | outside |
| 312 | 324 | IYQTSNFRVQPT | 1.1283 | 12 | ABCPRED,AAP | 2 | NON-ALLERGEN | Non-Toxin | -0.21 | -0.72 | -0.42 | 1 | 9.1 | 1453.79 | outside |
| 378 | 394 | KCYGVSPTKLNDLCFT | 2.0395 | 16 | ABCPRED | 1 | NON-ALLERGEN | Non-Toxin | -0.11 | -0.04 | -0.2 | 1 | 8.34 | 1789.32 | outside |
| 378 | 390 | KCYGVSPTKLND | 1.3195 | 12 | ABCPRED,BCPREDS | 2 | ALLERGEN | Non-Toxin | -0.23 | -0.76 | 0.21 | 1 | 8.51 | 1324.68 | outside |
| 294 | 312 | DPLSETKCTLKSFTVEKG | 0.5951 | 18 | AAP | 1 | NON-ALLERGEN | Non-Toxin | -0.22 | -0.6 | 0.49 | 0 | 6.49 | 1983.52 | outside |
| 844 | 856 | IAARDLICAQKF | 0.5718 | 12 | ABCPRED | 1 | ALLERGEN | Non-Toxin | -0.07 | 0.67 | -0.1 | 1 | 8.57 | 1348.79 | outside |
| 1059 | 1077 | GVVFLHVTYVPAQEKNFT | 1.1197 | 18 | ABCPRED | 1 | NON-ALLERGEN | Non-Toxin | 0.03 | 0.32 | -0.58 | 0.5 | 7.1 | 2049.63 | outside |
| 772 | 790 | VEQDKNTQEVFAQVKQIY | 0.2311 | 18 | ABCPRED | 1 | NON-ALLERGEN | Non-Toxin | -0.25 | -0.89 | 0.22 | -1 | 4.68 | 2167.68 | outside |
| 836 | 854 | QYGDCLGDIAARDLICAQ | 0.6649 | 18 | ABCPRED | 1 | ALLERGEN | Non-Toxin | -0.09 | 0.16 | -0.03 | -2 | 3.94 | 1925.42 | outside |
| 263 | 275 | AAYYVGYLQPRT | 0.9255 | 12 | BCPREDS | 1 | NON-ALLERGEN | Non-Toxin | -0.08 | -0.25 | -0.7 | 1 | 8.83 | 1401.75 | outside |
| 1064 | 1080 | HVTYVPAQEKNFTTAP | 0.8933 | 16 | ABCPRED,literature | 2 | NON-ALLERGEN | Non-Toxin | -0.12 | -0.59 | -0.26 | 0.5 | 7.1 | 1803.23 | outside |
| 276 | 294 | LLKYNENGTITDAVDCAL | 0.493 | 18 | ABCPRED | 1 | NON-ALLERGEN | Non-Toxin | -0.07 | 0.09 | -0.08 | -2 | 4.03 | 1953.46 | outside |
| 157 | 175 | FRVYSSANNCTFEYVSQP | -0.0112 | 18 | ABCPRED | 1 | NON-ALLERGEN | Non-Toxin | -0.15 | -0.42 | -0.39 | 0 | 6.32 | 2112.54 | outside |

| | | | | | | | | | | | | | | | |
|---|---|---|---|---|---|---|---|---|---|---|---|---|---|---|---|
| 1209 | 1225 | YIKWPWYIWLGFIAGL | 0.8934 | 16 | ABCPRED | 1 | NON-ALLERGEN | Non-Toxin | 0.28 | 0.88 | -1.49 | 1 | 8.83 | 2026.71 | outside_transmembrane-region |
| 900 | 912 | MQMAYRFNGIGV | 1.3524 | 12 | ABCPRED | 1 | NON-ALLERGEN | Non-Toxin | -0.01 | 0.29 | -0.65 | 1 | 9.1 | 1386.83 | outside |
| 15 | 31 | CVNLTTRTQLPPAYTN | 1.382 | 16 | literature | 1 | NON-ALLERGEN | Non-Toxin | -0.17 | -0.39 | -0.43 | 1 | 8.57 | 1792.27 | outside |
| 276 | 296 | LLKYNENGTITDAVDCALDP | 0.5659 | 20 | ABCPRED,literature | 2 | NON-ALLERGEN | Non-Toxin | -0.1 | -0.17 | 0.08 | -3 | 3.84 | 2165.69 | outside |
| 395 | 407 | VYADSFVIRGDE | -0.1343 | 12 | ABCPRED | 1 | NON-ALLERGEN | Non-Toxin | -0.1 | 0 | 0.18 | -2 | 4.03 | 1370.64 | outside |
| 163 | 181 | ANNCTFEYVSQPFLMDLE | 0.4285 | 18 | ABCPRED | 1 | NON-ALLERGEN | Non-Toxin | -0.06 | -0.1 | -0.32 | -3 | 3.58 | 2121.62 | outside |
| 785 | 803 | VKQIYKTPPIKDFGGFNF | 0.1499 | 18 | ABCPRED | 1 | ALLERGEN | Non-Toxin | -0.08 | -0.37 | -0.16 | 2 | 9.55 | 2099.73 | outside |
| 1063 | 1081 | LHVTYVPAQEKNFTTAPA | 0.9567 | 18 | BCPREDS,FBCPred | 2 | NON-ALLERGEN | Non-Toxin | -0.06 | -0.21 | -0.36 | 0.5 | 7.1 | 1987.5 | outside |
| 1244 | 1264 | LKGCCSCGSCCKFDEDDSEP | 0.1896 | 20 | FBCPred | 1 | NON-ALLERGEN | Non-Toxin | -0.24 | -0.55 | 0.63 | -3 | 4.11 | 2126.6 | inside |
| 71 | 81 | SGTNGTKRFD | 0.7387 | 10 | ABCPRED | 1 | ALLERGEN | Non-Toxin | -0.39 | -1.56 | 0.62 | 1 | 9.1 | 1082.27 | outside |
| 145 | 159 | YHKNNKSWMESEFR | 0.5416 | 14 | AAP | 1 | ALLERGEN | Non-Toxin | -0.44 | -2.04 | 0.43 | 1.5 | 8.84 | 1856.24 | outside |
| 128 | 140 | IKVCEFQFCNDP | -0.6957 | 12 | ABCPRED | 1 | NON-ALLERGEN | Non-Toxin | -0.11 | -0.02 | -0.07 | -1 | 4.38 | 1442.83 | outside |
| 35 | 44 | GVYYPDKVF | 0.0652 | 9 | IEDB | 1 | NON-ALLERGEN | Non-Toxin | 0 | -0.09 | -0.46 | 0 | 6.18 | 1087.36 | outside |
| 990 | 1008 | EVQIDRLITGRLQSLQTY | -0.3309 | 18 | ABCPRED | 1 | NON-ALLERGEN | Non-Toxin | -0.21 | -0.32 | -0.04 | 0 | 6.42 | 2133.72 | outside |
| 1193 | 1211 | LNESLIDLQELGKYEQYI | 0.5807 | 18 | ABCPRED | 1 | NON-ALLERGEN | Non-Toxin | -0.12 | -0.44 | 0.03 | -3 | 4.01 | 2168.72 | outside |
| 783 | 797 | AQVKQIYKTPPIKD | 0.3637 | 14 | ABCPRED,AAP | 2 | NON-ALLERGEN | Toxin | -0.25 | -0.89 | 0.29 | 2 | 9.55 | 1629.14 | outside |
| 415 | 433 | TGKIADYNYKLPDDFTGC | 0.6148 | 18 | ABCPRED | 1 | ALLERGEN | Non-Toxin | -0.16 | -0.71 | 0.12 | -1 | 4.43 | 2021.48 | outside |
| 1253 | 1265 | CCKFDEDDSEPV | 0.4603 | 12 | BCEPRED | 1 | NON-ALLERGEN | Non-Toxin | -0.3 | -0.98 | 1.03 | -4 | 3.77 | 1386.61 | inside |
| 901 | 919 | QMAYRFNGIGVTQNVLYE | 0.7768 | 18 | ABCPRED | 1 | ALLERGEN | Non-Toxin | -0.08 | -0.16 | -0.51 | 0 | 6.35 | 2103.66 | outside |
| 382 | 395 | VSPTKLNDLCFTN | 2.1787 | 13 | literature | 1 | ALLERGEN | Non-Toxin | -0.12 | -0.08 | -0.21 | 0 | 6.16 | 1451.84 | outside |
| 1148 | 1161 | FKEELDKYFKNHT | -0.967 | 13 | BCEPRED | 1 | NON-ALLERGEN | Non-Toxin | -0.36 | -1.65 | 0.63 | 0.5 | 7.1 | 1699.08 | outside |
| 495 | 511 | YGFQPTNGVGYQPYRV | 0.7985 | 16 | ABCPRED | 1 | NON-ALLERGEN | Non-Toxin | -0.12 | -0.8 | -0.57 | 1 | 8.83 | 1846.27 | outside |
| 1057 | 1075 | PHGVVFLHVTYVPAQEKN | 0.9072 | 18 | ABCPRED | 1 | ALLERGEN | Non-Toxin | -0.02 | -0.07 | -0.45 | 1 | 7.25 | 2035.61 | outside |
| 318 | 325 | FRVQPTE | 1.6729 | 7 | IEDB | 1 | ALLERGEN | Non-Toxin | -0.31 | -0.97 | 0.26 | 0 | 6.36 | 876.07 | outside |
| 794 | 812 | IKDFGGFNFSQILPDPSK | 0.1651 | 18 | ABCPRED | 1 | NON-ALLERGEN | Non-Toxin | -0.08 | -0.34 | 0.01 | 0 | 6.31 | 2010.54 | outside |
| 401 | 417 | VIRGDEVRQIAPGQTG | 0.6398 | 16 | ABCPRED | 1 | NON-ALLERGEN | Non-Toxin | -0.2 | -0.46 | 0.31 | 0 | 6.42 | 1696.14 | outside |
| 40 | 52 | DKVFRSSVLHST | -0.2832 | 12 | ABCPRED | 1 | NON-ALLERGEN | Non-Toxin | -0.23 | -0.27 | 0.14 | 1.5 | 9.1 | 1375.71 | outside |
| 347 | 357 | FASVYAWNRK | 0.5868 | 10 | ABCPRED | 1 | NON-ALLERGEN | Non-Toxin | -0.17 | -0.43 | -0.42 | 2 | 10.01 | 1241.54 | outside |
| 43 | 55 | FRSSVLHSTQDL | 0.4159 | 12 | ABCPRED | 1 | NON-ALLERGEN | Non-Toxin | -0.19 | -0.27 | -0.12 | 0.5 | 7.1 | 1389.7 | outside |
| 502 | 512 | GVGYQPYRVV | 1.1915 | 10 | ABCPRED | 1 | ALLERGEN | Non-Toxin | -0.05 | -0.04 | -0.59 | 1 | 8.93 | 1137.45 | outside |
| 774 | 786 | QDKNTQEVFAQV | 0.2877 | 12 | ABCPRED | 1 | NON-ALLERGEN | Non-Toxin | -0.28 | -1.05 | 0.28 | -1 | 4.38 | 1406.69 | outside |
| 784 | 797 | QVKQIYKTPPIKD | 0.3607 | 13 | BCEPRED | 1 | ALLERGEN | Non-Toxin | -0.28 | -1.09 | 0.35 | 2 | 9.55 | 1558.05 | outside |
| 372 | 398 | ASFSTFKCYGVSPTKLNDLCFTNVYA | 1.288 | 26 | literature | 1 | NON-ALLERGEN | Toxin | -0.03 | 0.22 | -0.48 | 1 | 8.33 | 2877.64 | outside |
| 515 | 531 | FELLHAPATVCGPKKS | 0.1215 | 16 | ABCPRED | 1 | NON-ALLERGEN | Non-Toxin | -0.06 | 0.07 | -0.07 | 1.5 | 8.54 | 1698.24 | outside |
| 145 | 157 | YHKNNKSWMESE | 0.3182 | 12 | FBCPred | 1 | NON-ALLERGEN | Non-Toxin | -0.42 | -2.24 | 0.46 | 0.5 | 7.1 | 1552.85 | outside |
| 135 | 153 | FCNDPFLGVYYHKNNKSW | 0.5084 | 18 | ABCPRED | 1 | NON-ALLERGEN | Toxin | -0.15 | -0.84 | -0.44 | 1.5 | 8.48 | 2232.76 | outside |
| 682 | 700 | RRARSVASQSIIAYTMSL | 0.4521 | 18 | ABCPRED | 1 | NON-ALLERGEN | Non-Toxin | -0.2 | 0.12 | -0.11 | 3 | 11.72 | 2010.59 | outside |
| 778 | 796 | TQEVFAQVKQIYKTPPIK | 0.2322 | 18 | ABCPRED | 1 | NON-ALLERGEN | Non-Toxin | -0.17 | -0.53 | -0.01 | 2 | 9.55 | 2118.78 | outside |
| 260 | 276 | AGAAAYYVGYLQPRTF | 0.7711 | 16 | ABCPRED | 1 | NON-ALLERGEN | Non-Toxin | 0.02 | 0.19 | -0.74 | 1 | 8.83 | 1748.19 | outside |
| 685 | 703 | RSVASQSIIAYTMSLGAE | 0.4149 | 18 | ABCPRED | 1 | NON-ALLERGEN | Non-Toxin | -0.03 | 0.4 | -0.28 | 0 | 6.35 | 1884.39 | outside |
| 133 | 149 | FQFCNDPFLGVYYHKN | 0.5081 | 16 | ABCPRED | 1 | NON-ALLERGEN | Toxin | -0.07 | -0.42 | -0.64 | 0.5 | 7.05 | 1992.47 | outside |
| 861 | 879 | LPPLLTDEMIAQYTSALL | 0.3058 | 18 | ABCPRED | 1 | NON-ALLERGEN | Non-Toxin | 0.08 | 0.66 | -0.54 | -2 | 3.67 | 1989.63 | outside |
| 1002 | 1022 | QSLQTYVTQQLIRAAEIRAS | 0.0298 | 20 | AAP | 1 | NON-ALLERGEN | Non-Toxin | -0.2 | -0.23 | -0.15 | 1 | 9.1 | 2276.88 | outside |
| 1209 | 1227 | YIKWPWYIWLGFIAGLIA | 0.8171 | 18 | ABCPRED | 1 | ALLERGEN | Non-Toxin | 0.3 | 1.13 | -1.45 | 1 | 8.83 | 2210.98 | outside_transmembrane-region |
| 493 | 511 | QSYGFQPTNGVGYQPYRV | 0.7469 | 18 | ABCPRED | 1 | ALLERGEN | Non-Toxin | -0.16 | -0.95 | -0.48 | 1 | 8.83 | 2061.51 | outside |
| 904 | 920 | YRFNGIGVTQNVLYEN | 0.7167 | 16 | ABCPRED | 1 | NON-ALLERGEN | Non-Toxin | -0.12 | -0.41 | -0.46 | 0 | 6.35 | 1887.33 | outside |
| 197 | 215 | IDGYFKIYSKHTPINLVR | 0.5724 | 18 | ABCPRED | 1 | NON-ALLERGEN | Non-Toxin | -0.12 | -0.24 | -0.23 | 2.5 | 9.55 | 2164.81 | outside |
| 770 | 786 | IAVEQDKNTQEVFAQV | 0.2917 | 16 | ABCPRED | 1 | NON-ALLERGEN | Non-Toxin | -0.16 | -0.35 | 0.16 | -2 | 4.14 | 1819.24 | outside |
| 818 | 830 | IEDLLFNKVTLA | 0.5555 | 12 | ABCPRED | 1 | ALLERGEN | Non-Toxin | 0.04 | 0.8 | -0.24 | -1 | 4.38 | 1375.81 | outside |
| 798 | 816 | GGFNFSQILPDPSKPSKR | 0.5239 | 18 | ABCPRED | 1 | ALLERGEN | Non-Toxin | -0.23 | -0.94 | 0.26 | 2 | 10.01 | 1975.49 | outside |
| 167 | 179 | TFEYVSQPFLMD | 0.5408 | 12 | ABCPRED | 1 | ALLERGEN | Non-Toxin | 0 | 0.05 | -0.48 | -2 | 3.67 | 1476.83 | outside |
| 1041 | 1057 | DFCGKGYHLMSFPQSA | 0.2205 | 16 | ABCPRED | 1 | ALLERGEN | Non-Toxin | -0.05 | -0.24 | -0.35 | 0.5 | 7.06 | 1788.25 | outside |
| 403 | 415 | RGDEVRQIAPGQ | 0.4088 | 12 | ABCPRED | 1 | NON-ALLERGEN | Non-Toxin | -0.37 | -1.24 | 0.72 | 0 | 6.42 | 1325.62 | outside |
| 442 | 460 | DSKVGGNYNYLYRLFRKS | 0.1448 | 18 | ABCPRED | 1 | NON-ALLERGEN | Non-Toxin | -0.31 | -1.06 | 0.08 | 3 | 9.83 | 2180.71 | outside |
| 319 | 337 | RVQPTESIVRFPNITNLC | 0.571 | 18 | ABCPRED | 1 | NON-ALLERGEN | Non-Toxin | -0.18 | -0.11 | -0.16 | 1 | 8.6 | 2087.69 | outside |
| 20 | 32 | TRTQLPPAYTNS | 0.9332 | 12 | ABCPRED,literature | 2 | NON-ALLERGEN | Non-Toxin | -0.27 | -1.11 | -0.17 | 1 | 9.1 | 1348.64 | outside |
| 670 | 686 | ICASYQTQTNSPRRAR | 0.3821 | 16 | ABCPRED | 1 | NON-ALLERGEN | Non-Toxin | -0.43 | -1.21 | 0.21 | 3 | 10.77 | 1852.27 | outside |
| 549 | 567 | TGVLTESNKKFLPFQQFG | 0.6749 | 18 | ABCPRED | 1 | NON-ALLERGEN | Non-Toxin | -0.1 | -0.34 | -0.19 | 1 | 8.94 | 2041.61 | outside |
| 430 | 446 | TGCVIAWNSNNLDSKV | 0.5086 | 16 | ABCPRED | 1 | NON-ALLERGEN | Non-Toxin | -0.08 | -0.03 | -0.29 | 0 | 6.16 | 1721.15 | outside |
| 73 | 81 | TNGTKRFD | 0.2041 | 8 | IEDB | 1 | ALLERGEN | Non-Toxin | -0.48 | -1.8 | 0.74 | 1 | 9.1 | 938.11 | outside |
| 499 | 517 | PTNGVGYQPYRVVVLSFE | 0.722 | 18 | ABCPRED | 1 | ALLERGEN | Non-Toxin | -0.03 | 0.02 | -0.48 | 0 | 6.35 | 2025.56 | outside |
| 124 | 140 | TNVVIKVCEFQFCNDP | -0.3969 | 16 | ABCPRED | 1 | ALLERGEN | Non-Toxin | -0.06 | 0.25 | -0.26 | -1 | 4.38 | 1856.37 | outside |
| 788 | 800 | IYKTPPIKDFGG | -0.5719 | 12 | ABCPRED | 1 | ALLERGEN | Non-Toxin | -0.07 | -0.46 | 0.02 | 1 | 8.83 | 1335.74 | outside |
| 501 | 511 | NGVGYQPYRV | 0.7324 | 10 | ABCPRED | 1 | ALLERGEN | Non-Toxin | -0.17 | -0.81 | -0.42 | 1 | 8.93 | 1152.42 | outside |
| 323 | 337 | TESIVRFPNITNLC | 0.3641 | 14 | FBCPred | 1 | ALLERGEN | Toxin | -0.08 | 0.25 | -0.32 | 0 | 6.32 | 1607.06 | outside |
| 789 | 805 | YKTPPIKDFGGFNFSQ | 0.2144 | 16 | ABCPRED | 1 | ALLERGEN | Non-Toxin | -0.12 | -0.76 | -0.14 | 1 | 8.83 | 1846.3 | outside |
| 21 | 39 | RTQLPPAYTNSFTRGVYY | 0.3371 | 18 | ABCPRED | 1 | NON-ALLERGEN | Non-Toxin | -0.2 | -0.77 | -0.43 | 2 | 9.72 | 2134.63 | outside |
| 560 | 570 | LPFQQFGRDI | 0.0043 | 10 | ABCPRED | 1 | NON-ALLERGEN | Non-Toxin | -0.13 | -0.31 | -0.22 | 0 | 6.19 | 1220.54 | outside |
| 505 | 515 | YQPYRVVVLS | 0.7612 | 10 | ABCPRED | 1 | NON-ALLERGEN | Non-Toxin | -0.06 | 0.34 | -0.74 | 1 | 8.93 | 1223.58 | outside |
| 733 | 743 | KTSVDCTMYI | 0.9345 | 10 | ABCPRED | 1 | ALLERGEN | Non-Toxin | -0.08 | 0.22 | -0.24 | 0 | 6.15 | 1160.5 | outside |
| 898 | 910 | FAMQMAYRFNGI | 0.8395 | 12 | ABCPRED | 1 | NON-ALLERGEN | Non-Toxin | 0.01 | 0.36 | -0.77 | 1 | 9.1 | 1448.89 | outside |
| 134 | 150 | QFCNDPFLGVYYHKNN | 0.5475 | 16 | ABCPRED | 1 | NON-ALLERGEN | Non-Toxin | -0.15 | -0.82 | -0.47 | 0.5 | 7.05 | 1959.4 | outside |
| 191 | 209 | EFVFKNIDGYFKIYSKHT | 0.5302 | 18 | ABCPRED | 1 | ALLERGEN | Non-Toxin | -0.12 | -0.46 | -0.14 | 1.5 | 8.71 | 2236.82 | outside |
| 496 | 514 | GFQPTNGVGYQPYRVVVL | 0.7542 | 18 | FBCPred | 1 | NON-ALLERGEN | Non-Toxin | -0.02 | 0.04 | -0.65 | 1 | 8.93 | 1994.56 | outside |
| 617 | 635 | CTEVPVAIHADQLTPTWR | 0.3701 | 18 | ABCPRED | 1 | NON-ALLERGEN | Non-Toxin | -0.09 | -0.09 | -0.25 | -0.5 | 5.33 | 2037.58 | outside |
| 787 | 803 | QIYKTPPIKDFGGFNF | 0.0547 | 16 | ABCPRED | 1 | ALLERGEN | Non-Toxin | -0.06 | -0.43 | -0.27 | 1 | 8.83 | 1872.39 | outside |
| 374 | 392 | FSTFKCYGVSPTKLNDLC | 1.2396 | 18 | ABCPRED | 1 | NON-ALLERGEN | Non-Toxin | -0.08 | 0.07 | -0.3 | 1 | 8.34 | 2023.6 | outside |

| | | | | | | | | | | | | | | | |
|---|---|---|---|---|---|---|---|---|---|---|---|---|---|---|---|
| 505 | 517 | YQPYRVVVLSFE | 0.9911 | 12 | ABCPRED | 1 | NON-ALLERGEN | Non-Toxin | -0.05 | 0.23 | -0.57 | 0 | 6.35 | 1499.9 | outside |
| 643 | 659 | FQTRAGCLIGAEHVNN | 1.2294 | 16 | ABCPRED | 1 | ALLERGEN | Non-Toxin | -0.1 | -0.11 | -0.24 | 0.5 | 7.07 | 1730.17 | outside |
| 410 | 426 | IAPGQTGKIADYNYKL | 1.6707 | 16 | ABCPRED | 1 | ALLERGEN | Non-Toxin | -0.1 | -0.48 | -0.12 | 1 | 8.76 | 1752.24 | outside |
| 672 | 692 | ASYQTQTNSPRRARSVASQS | 0.2704 | 20 | ABCPRED,AAP,BCPREDS | 3 | ALLERGEN | Non-Toxin | -0.42 | -1.31 | 0.26 | 3 | 11.72 | 2195.6 | outside |
| 514 | 532 | SFELLHAPATVCGPKKST | 0.2697 | 18 | ABCPRED | 1 | ALLERGEN | Non-Toxin | -0.07 | -0.02 | -0.07 | 1.5 | 8.54 | 1886.45 | outside |
| 562 | 572 | FQQFGRDIAD | -0.2747 | 10 | ABCPRED | 1 | NON-ALLERGEN | Non-Toxin | -0.22 | -0.7 | 0.21 | -1 | 4.21 | 1196.42 | outside |
| 278 | 288 | KYNENGTITD | 0.9589 | 10 | BCEPRED | 1 | NON-ALLERGEN | Non-Toxin | -0.32 | -1.65 | 0.45 | -1 | 4.38 | 1154.34 | outside |
| 208 | 218 | TPINLVRDLP | 0.2894 | 10 | ABCPRED | 1 | NON-ALLERGEN | Non-Toxin | -0.11 | 0.09 | -0.11 | 0 | 6.19 | 1137.49 | outside |
| 40 | 50 | DKVFRSSVLH | -0.5398 | 10 | ABCPRED | 1 | ALLERGEN | Non-Toxin | -0.23 | -0.17 | 0.18 | 1.5 | 9.1 | 1187.5 | outside |
| 119 | 139 | IVNNATNVVIKVCEFQFCND | -0.3479 | 20 | FBCPred | 1 | ALLERGEN | Non-Toxin | -0.03 | 0.46 | -0.38 | -1 | 4.38 | 2270.9 | outside |
| 169 | 181 | EYVSQPFLMDLE | 0.6161 | 12 | ABCPRED | 1 | ALLERGEN | Non-Toxin | -0.04 | -0.1 | -0.14 | -3 | 3.58 | 1470.83 | outside |
| 1062 | 1082 | FLHVTYVPAQEKNFTTAPAI | 0.9351 | 20 | AAP | 1 | ALLERGEN | Non-Toxin | 0.01 | 0.18 | -0.54 | 0.5 | 7.1 | 2247.87 | outside |
| 289 | 305 | VDCALDPLSETKCTLK | 0.7752 | 16 | ABCPRED | 1 | NON-ALLERGEN | Non-Toxin | -0.16 | 0.02 | 0.32 | -1 | 4.56 | 1736.25 | outside |
| 288 | 304 | AVDCALDPLSETKCTL | 0.7905 | 16 | ABCPRED | 1 | ALLERGEN | Non-Toxin | -0.07 | 0.38 | 0.1 | -2 | 4.03 | 1679.15 | outside |
| 1007 | 1019 | YVTQQLIRAAEI | 0.2847 | 12 | AAP | 1 | NON-ALLERGEN | Non-Toxin | -0.07 | 0.3 | -0.35 | 0 | 6.35 | 1404.81 | outside |
| 633 | 645 | WRVYSTGSNVFQ | 0.0616 | 12 | ABCPRED | 1 | NON-ALLERGEN | Non-Toxin | -0.13 | -0.43 | -0.63 | 1 | 9.1 | 1443.75 | outside |
| 721 | 739 | SVTTEILPVSMTKTSVDC | 1.2841 | 18 | ABCPRED | 1 | NON-ALLERGEN | Non-Toxin | -0.05 | 0.42 | -0.12 | -1 | 4.38 | 1911.47 | outside |
| 988 | 1004 | EAEVQIDRLITGRLQS | -0.2814 | 16 | ABCPRED | 1 | NON-ALLERGEN | Non-Toxin | -0.24 | -0.36 | 0.38 | -1 | 4.68 | 1828.3 | outside |
| 549 | 565 | TGVLTESNKKFLPFQQ | 0.7878 | 16 | ABCPRED | 1 | NON-ALLERGEN | Non-Toxin | -0.16 | -0.54 | -0.06 | 1 | 8.94 | 1837.35 | outside |
| 859 | 879 | TVLPPLLTDEMIAQYTSALL | 0.2357 | 20 | ABCPRED,FBCPred | 2 | NON-ALLERGEN | Non-Toxin | 0.09 | 0.77 | -0.58 | -2 | 3.67 | 2189.9 | outside |
| 1254 | 1265 | CKFDEDDSEPV | 0.2159 | 11 | BCEPRED | 1 | NON-ALLERGEN | Non-Toxin | -0.33 | -1.3 | 1.21 | -4 | 3.77 | 1283.46 | inside |
| 633 | 643 | WRVYSTGSNV | 0.4326 | 10 | ABCPRED | 1 | NON-ALLERGEN | Non-Toxin | -0.15 | -0.45 | -0.53 | 1 | 9.1 | 1168.41 | outside |
| 377 | 395 | FKCYGVSPTKLNDLCFTN | 1.9296 | 18 | ABCPRED | 1 | ALLERGEN | Non-Toxin | -0.1 | -0.08 | -0.31 | 1 | 8.34 | 2050.63 | outside |
| 787 | 797 | QIYKTPPIKD | 0.0772 | 10 | ABCPRED | 1 | ALLERGEN | Non-Toxin | -0.25 | -1.1 | 0.29 | 1 | 8.83 | 1202.56 | outside |
| 645 | 663 | TRAGCLIGAEHVNNSYEC | 1.1982 | 18 | ABCPRED | 1 | NON-ALLERGEN | Non-Toxin | -0.13 | -0.23 | -0.09 | -0.5 | 5.41 | 1937.39 | outside |
| 446 | 460 | GGNYNYLYRLFRKS | -0.1832 | 14 | FBCPred | 1 | NON-ALLERGEN | Non-Toxin | -0.29 | -1.07 | -0.24 | 3 | 10 | 1751.18 | outside |
| 669 | 687 | GICASYQTQTNSPRRARS | 0.3898 | 18 | ABCPRED,FBCPred | 2 | NON-ALLERGEN | Non-Toxin | -0.39 | -1.14 | 0.2 | 3 | 10.77 | 1996.43 | outside |
| 672 | 690 | ASYQTQTNSPRRARSVAS | 0.2963 | 18 | IEDB | 1 | NON-ALLERGEN | Non-Toxin | -0.41 | -1.22 | 0.26 | 3 | 11.72 | 1980.36 | outside |
| 667 | 685 | GAGICASYQTQTNSPRRA | 0.31 | 18 | ABCPRED | 1 | NON-ALLERGEN | Non-Toxin | -0.26 | -0.77 | -0.01 | 2 | 9.55 | 1881.3 | outside |
| 968 | 986 | SNFGAISSVLNDILSRLD | 0.2827 | 18 | ABCPRED | 1 | NON-ALLERGEN | Non-Toxin | -0.05 | 0.39 | -0.16 | -1 | 4.21 | 1921.4 | outside |
| 191 | 207 | EFVFKNIDGYFKIYSK | 0.4475 | 16 | ABCPRED | 1 | NON-ALLERGEN | Non-Toxin | -0.09 | -0.28 | -0.11 | 1 | 8.71 | 1998.54 | outside |
| 673 | 689 | SYQTQTNSPRRARSVA | 0.3218 | 16 | ABCPRED | 1 | NON-ALLERGEN | Non-Toxin | -0.47 | -1.43 | 0.31 | 3 | 11.72 | 1822.18 | outside |
| 269 | 279 | YLQPRTFLLK | -0.0183 | 10 | ABCPRED | 1 | ALLERGEN | Non-Toxin | -0.16 | -0.13 | -0.44 | 2 | 10.01 | 1278.71 | outside |
| 863 | 875 | PLLTDEMIAQYT | 0.153 | 12 | ABCPRED | 1 | ALLERGEN | Non-Toxin | -0.01 | 0.08 | -0.34 | -2 | 3.67 | 1394.78 | outside |
| 838 | 856 | GDCLGDIAARDLICAQKF | 0.3629 | 18 | ABCPRED | 1 | NON-ALLERGEN | Non-Toxin | -0.08 | 0.37 | 0.11 | -1 | 4.43 | 1909.46 | outside |
| 789 | 806 | YKTPPIKDFGGFNFSQI | 0.3455 | 17 | ellipro,literature | 2 | ALLERGEN | Non-Toxin | -0.07 | -0.45 | -0.24 | 1 | 8.83 | 1959.48 | outside |
| 380 | 390 | YGVSPTKLND | 1.4531 | 10 | IEDB,literature | 2 | NON-ALLERGEN | Non-Toxin | -0.17 | -0.77 | 0.05 | 0 | 6.18 | 1093.34 | outside |
| 357 | 367 | RISNCVADYS | -0.3956 | 10 | ABCPRED | 1 | NON-ALLERGEN | Non-Toxin | -0.21 | -0.14 | -0.03 | 0 | 6.15 | 1127.36 | outside |
| 992 | 1012 | QIDRLITGRLQSLQTYVTQQ | -0.3021 | 20 | ABCPRED | 1 | ALLERGEN | Non-Toxin | -0.24 | -0.5 | -0.18 | 1 | 9.1 | 2362.01 | outside |
| 792 | 808 | PPIKDFGGFNFSQILP | 0.3404 | 16 | ABCPRED | 1 | NON-ALLERGEN | Non-Toxin | 0.03 | 0.02 | -0.39 | 0 | 6.19 | 1777.29 | outside |
| 1194 | 1212 | NESLIDLQELGKYEQYIK | 0.5691 | 18 | ABCPRED | 1 | NON-ALLERGEN | Non-Toxin | -0.21 | -0.87 | 0.29 | -2 | 4.41 | 2183.73 | outside |
| 783 | 801 | AQVKQIYKTPPIKDFGGF | -0.0606 | 18 | AAP | 1 | NON-ALLERGEN | Non-Toxin | -0.11 | -0.42 | -0.05 | 2 | 9.55 | 2037.66 | outside |
| 788 | 798 | IYKTPPIKDF | -0.1626 | 10 | IEDB | 1 | NON-ALLERGEN | Non-Toxin | -0.12 | -0.47 | 0.02 | 1 | 8.83 | 1221.6 | outside |
| 504 | 516 | GYQPYRVVVLSF | 1.1235 | 12 | ABCPRED | 1 | NON-ALLERGEN | Non-Toxin | 0.01 | 0.48 | -0.82 | 1 | 8.93 | 1427.84 | outside |
| 382 | 394 | VSPTKLNDLCFT | 2.4268 | 12 | ABCPRED | 1 | NON-ALLERGEN | Non-Toxin | -0.08 | 0.2 | -0.24 | 0 | 6.16 | 1337.72 | outside |
| 14 | 30 | QCVNLTTRTQLPPAYT | 1.4165 | 16 | ABCPRED | 1 | NON-ALLERGEN | Non-Toxin | -0.17 | -0.39 | -0.43 | 1 | 8.57 | 1806.3 | outside |
| 36 | 44 | VYYPDKVF | 0.0301 | 8 | IEDB | 1 | NON-ALLERGEN | Non-Toxin | -0.02 | -0.05 | -0.51 | 0 | 6.18 | 1030.29 | outside |
| 563 | 573 | QQFGRDIADT | -0.2223 | 10 | ABCPRED | 1 | ALLERGEN | Toxin | -0.3 | -1.05 | 0.42 | -1 | 4.21 | 1150.35 | outside |
| 1243 | 1261 | CLKGCCSCGSCCKFDEDD | 0.141 | 18 | ABCPRED | 1 | ALLERGEN | Non-Toxin | -0.21 | -0.14 | 0.46 | -2 | 4.23 | 1916.4 | inside |
| 1060 | 1076 | VVFLHVTYVPAQEKNF | 1.1471 | 16 | ABCPRED | 1 | NON-ALLERGEN | Non-Toxin | 0.03 | 0.42 | -0.63 | 0.5 | 7.1 | 1891.44 | outside |
| 682 | 702 | RRARSVASQSIIAYTMSLGA | 0.5611 | 20 | ABCPRED | 1 | ALLERGEN | Non-Toxin | -0.16 | 0.17 | -0.12 | 3 | 11.72 | 2138.75 | outside |
| 508 | 518 | YRVVVLSFEL | 1.1494 | 10 | ABCPRED | 1 | NON-ALLERGEN | Non-Toxin | 0.07 | 1.29 | -0.66 | 0 | 6.35 | 1224.61 | outside |
| 260 | 278 | AGAAAYYVGYLQPRTFLL | 0.6528 | 18 | ABCPRED | 1 | NON-ALLERGEN | Non-Toxin | 0.08 | 0.59 | -0.86 | 1 | 8.83 | 1974.55 | outside |
| 1210 | 1228 | IKWPWYIWLGFIAGLIAI | 0.9041 | 18 | ABCPRED | 1 | NON-ALLERGEN | Non-Toxin | 0.34 | 1.46 | -1.42 | 1 | 8.94 | 2160.97 | outside_transmembrane-region |
| 786 | 798 | KQIYKTPPIKDF | -0.1444 | 12 | AAP | 1 | NON-ALLERGEN | Non-Toxin | -0.24 | -1.01 | 0.28 | 2 | 9.55 | 1477.94 | outside |
| 500 | 516 | TNGVGYQPYRVVVLSF | 0.7668 | 16 | ABCPRED | 1 | NON-ALLERGEN | Non-Toxin | 0 | 0.34 | -0.72 | 1 | 8.93 | 1799.3 | outside |
| 378 | 398 | KCYGVSPTKLNDLCFTNVYA | 1.5664 | 20 | ABCPRED,AAP | 2 | ALLERGEN | Non-Toxin | -0.08 | 0.02 | -0.36 | 1 | 8.33 | 2236.87 | outside |
| 431 | 451 | GCVIAWNSNNLDSKVGGNYN | 0.604 | 20 | ABCPRED | 1 | NON-ALLERGEN | Non-Toxin | -0.1 | -0.44 | -0.31 | 0 | 6.15 | 2125.6 | outside |
| 860 | 876 | VLPPLLTDEMIAQYTS | 0.274 | 16 | ABCPRED | 1 | NON-ALLERGEN | Non-Toxin | 0.04 | 0.41 | -0.44 | -2 | 3.67 | 1791.33 | outside |
| 792 | 810 | PPIKDFGGFNFSQILPDP | 0.2576 | 18 | ABCPRED | 1 | NON-ALLERGEN | Non-Toxin | -0.01 | -0.26 | -0.18 | -1 | 4.21 | 1989.52 | outside |
| 272 | 279 | PRTFLLK | -1.3917 | 7 | IEDB,literature | 2 | ALLERGEN | Non-Toxin | -0.21 | -0.04 | -0.07 | 2 | 11.01 | 874.19 | outside |
| 632 | 644 | TWRVYSTGSNVF | 0.1158 | 12 | ABCPRED | 1 | NON-ALLERGEN | Non-Toxin | -0.09 | -0.2 | -0.68 | 1 | 9.1 | 1416.72 | outside |
| 1067 | 1081 | YVPAQEKNFTTAPA | 0.6045 | 14 | ABCPRED,BCPREDS,FBCPred | 3 | NON-ALLERGEN | Non-Toxin | -0.12 | -0.56 | -0.16 | 0 | 6.35 | 1536.89 | outside |
| 732 | 744 | TKTSVDCTMYIC | 0.5424 | 12 | ABCPRED | 1 | NON-ALLERGEN | Non-Toxin | -0.08 | 0.33 | -0.32 | 0 | 6.12 | 1364.77 | outside |
| 416 | 428 | GKIADYNYKLPD | 1.3553 | 12 | ABCPRED | 1 | NON-ALLERGEN | Non-Toxin | -0.22 | -1.07 | 0.29 | 0 | 6.3 | 1396.73 | outside |
| 837 | 853 | YGDCLGDIAARDLICA | 0.5066 | 16 | ABCPRED | 1 | NON-ALLERGEN | Non-Toxin | -0.01 | 0.62 | -0.06 | -2 | 3.94 | 1669.12 | outside |
| 730 | 748 | SMTKTSVDCTMYICGDST | 0.1426 | 18 | ABCPRED | 1 | NON-ALLERGEN | Non-Toxin | -0.11 | -0.02 | -0.11 | -1 | 4.21 | 1943.45 | outside |
| 559 | 569 | FLPFQQFGRD | 0.287 | 10 | ABCPRED | 1 | NON-ALLERGEN | Non-Toxin | -0.14 | -0.48 | -0.29 | 0 | 6.19 | 1254.55 | outside |
| 20 | 30 | TRTQLPPAYT | 1.2416 | 10 | ABCPRED | 1 | NON-ALLERGEN | Non-Toxin | -0.23 | -0.9 | -0.26 | 1 | 9.1 | 1147.43 | outside |
| 1067 | 1083 | YVPAQEKNFTTAPAIC | 0.6145 | 16 | ABCPRED | 1 | ALLERGEN | Non-Toxin | -0.05 | -0.06 | -0.31 | 0 | 6.32 | 1753.22 | outside |
| 572 | 584 | TTDAVRDPQTLE | 0.2917 | 12 | AAP | 1 | ALLERGEN | Non-Toxin | -0.32 | -1.03 | 0.6 | -2 | 4.03 | 1345.59 | outside |
| 1147 | 1163 | SFKEELDKYFKNHTSP | -0.4485 | 16 | ABCPRED | 1 | NON-ALLERGEN | Non-Toxin | -0.33 | -1.54 | 0.55 | 0.5 | 7.1 | 1970.39 | outside |
| 1179 | 1187 | IQKEIDRL | -1.0253 | 8 | BCEPRED,IEDB | 2 | ALLERGEN | Non-Toxin | -0.36 | -0.76 | 0.85 | 0 | 6.42 | 1014.31 | outside |
| 788 | 806 | IYKTPPIKDFGGFNFSQI | 0.2158 | 18 | ABCPRED | 1 | ALLERGEN | Non-Toxin | -0.03 | -0.18 | -0.33 | 1 | 8.83 | 2072.66 | outside |
| 1192 | 1210 | NLNESLIDLQELGKYEQY | 0.7268 | 18 | ABCPRED | 1 | NON-ALLERGEN | Non-Toxin | -0.2 | -0.89 | 0.14 | -3 | 4.01 | 2169.66 | outside |
| 724 | 742 | TEILPVSMTKTSVDCTMY | 1.2922 | 18 | ABCPRED | 1 | NON-ALLERGEN | Non-Toxin | -0.05 | 0.27 | -0.25 | -1 | 4.38 | 2019.63 | outside |
| 773 | 785 | EQDKNTQEVFAQ | 0.0568 | 12 | ABCPRED | 1 | NON-ALLERGEN | Non-Toxin | -0.38 | -1.69 | 0.66 | -2 | 4.14 | 1436.67 | outside |

| | | | | | | | | | | | | | | | |
|---|---|---|---|---|---|---|---|---|---|---|---|---|---|---|---|
| 1097 | 1111 | SNGTHWFVTQRNFY | 0.26 | 14 | FBCPred | 1 | NON-ALLERGEN | Non-Toxin | -0.17 | -0.94 | -0.69 | 1.5 | 9.1 | 1757.1 | outside |
| 436 | 453 | WNSNNLDSKVGGNYNYL | 0.8074 | 17 | IEDB | 1 | NON-ALLERGEN | Non-Toxin | -0.19 | -1.12 | -0.32 | 0 | 6.18 | 1958.33 | outside |
| 1043 | 1063 | CGKGYHLMSFPQSAPHGVVF | 0.0669 | 20 | ABCPRED | 1 | NON-ALLERGEN | Non-Toxin | 0.03 | 0.14 | -0.6 | 2 | 8.54 | 2162.81 | outside |
| 71 | 83 | SGTNGTKRFDNP | 0.4821 | 12 | ABCPRED,FBCPred | 2 | NON-ALLERGEN | Toxin | -0.39 | -1.73 | 0.53 | 1 | 9.1 | 1293.52 | outside |
| 819 | 837 | EDLLFNKVTLADAGFIKQ | 0.6262 | 18 | ABCPRED | 1 | NON-ALLERGEN | Non-Toxin | -0.06 | 0.16 | 0.02 | -1 | 4.56 | 2022.6 | outside |
| 165 | 183 | NCTFEYVSQPFLMDLEGK | 0.6452 | 18 | ABCPRED | 1 | NON-ALLERGEN | Non-Toxin | -0.09 | -0.24 | -0.13 | -2 | 4.14 | 2121.67 | outside |
| 22 | 40 | TQLPPAYTNSFTRGVYYP | 0.2481 | 18 | ABCPRED | 1 | NON-ALLERGEN | Non-Toxin | -0.11 | -0.61 | -0.59 | 1 | 8.83 | 2075.56 | outside |
| 498 | 516 | QPTNGVGYQPYRVVVLSF | 0.6604 | 18 | ABCPRED,AAP | 2 | NON-ALLERGEN | Non-Toxin | -0.04 | 0.02 | -0.63 | 1 | 8.93 | 2024.58 | outside |
| 719 | 739 | TISVTTEILPVSMTKTSVDC | 1.3041 | 20 | ABCPRED,AAP | 2 | ALLERGEN | Non-Toxin | -0.01 | 0.57 | -0.22 | -1 | 4.38 | 2125.77 | outside |
| 121 | 139 | NNATNVVIKVCEFQFCND | -0.3653 | 18 | ABCPRED | 1 | NON-ALLERGEN | Non-Toxin | -0.11 | 0.02 | -0.23 | -1 | 4.38 | 2058.57 | outside |
| 228 | 240 | DLPIGINITRFQ | 1.3916 | 12 | ABCPRED | 1 | NON-ALLERGEN | Non-Toxin | -0.05 | 0.2 | -0.31 | 0 | 6.19 | 1386.8 | outside |
| 899 | 907 | AMQMAYRF | 0.9776 | 8 | COVIDep,literature | 2 | NON-ALLERGEN | Non-Toxin | -0.1 | 0.11 | -0.65 | 1 | 9.1 | 1017.33 | outside |
| 456 | 464 | FRKSNLKP | 1.1111 | 8 | BCEPRED | 1 | ALLERGEN | Non-Toxin | -0.47 | -1.45 | 0.65 | 3 | 11.17 | 989.29 | outside |
| 790 | 804 | KTPPIKDFGGFNFS | 0.2323 | 14 | ABCPRED,BCPREDS | 2 | NON-ALLERGEN | Non-Toxin | -0.09 | -0.53 | -0.01 | 1 | 8.94 | 1554.96 | outside |
| 1067 | 1080 | YVPAQEKNFTTAP | 0.6578 | 13 | BCEPRED | 1 | NON-ALLERGEN | Non-Toxin | -0.14 | -0.75 | -0.13 | 0 | 6.35 | 1465.8 | outside |
| 562 | 574 | FQQFGRDIADTT | 0.0707 | 12 | ABCPRED | 1 | ALLERGEN | Non-Toxin | -0.21 | -0.7 | 0.11 | -1 | 4.21 | 1398.66 | outside |
| 970 | 986 | FGAISSVLNDILSRLD | 0.1184 | 16 | ABCPRED | 1 | NON-ALLERGEN | Non-Toxin | 0 | 0.71 | -0.21 | -1 | 4.21 | 1720.19 | outside |
| 22 | 38 | TQLPPAYTNSFTRGVY | 0.1724 | 16 | ABCPRED | 1 | ALLERGEN | Non-Toxin | -0.12 | -0.5 | -0.53 | 1 | 8.93 | 1815.24 | outside |
| 729 | 745 | VSMTKTSVDCTMYICG | 0.4705 | 16 | ABCPRED | 1 | ALLERGEN | Non-Toxin | -0.02 | 0.56 | -0.39 | 0 | 6.12 | 1739.29 | outside |
| 435 | 449 | AWNSNNLDSKVGGN | 0.7597 | 14 | BCPREDS | 1 | NON-ALLERGEN | Non-Toxin | -0.21 | -1.06 | 0.01 | 0 | 6.19 | 1475.74 | outside |
| 819 | 831 | EDLLFNKVTLAD | 0.7333 | 12 | ABCPRED | 1 | NON-ALLERGEN | Non-Toxin | -0.08 | 0.13 | 0.16 | -2 | 4.03 | 1377.73 | outside |
| 632 | 648 | TWRVYSTGSNVFQTRA | 0.4092 | 16 | ABCPRED | 1 | NON-ALLERGEN | Non-Toxin | -0.21 | -0.58 | -0.37 | 2 | 10.84 | 1873.28 | outside |
| 502 | 514 | GVGYQPYRVVVL | 1.1207 | 12 | BCPREDS,FBCPred | 2 | ALLERGEN | Non-Toxin | 0.04 | 0.63 | -0.77 | 1 | 8.93 | 1349.78 | outside |
| 315 | 325 | TSNFRVQPTE | 1.3571 | 10 | ABCPRED,literature | 2 | ALLERGEN | Non-Toxin | -0.33 | -1.18 | 0.19 | 0 | 6.36 | 1178.4 | outside |
| 18 | 36 | LTTRTQLPPAYTNSFTRG | 0.5312 | 18 | ABCPRED,AAP | 2 | NON-ALLERGEN | Non-Toxin | -0.22 | -0.72 | -0.23 | 2 | 10.84 | 2024.52 | outside |
| 640 | 658 | SNVFQTRAGCLIGAEHVN | 0.9399 | 18 | ABCPRED | 1 | NON-ALLERGEN | Non-Toxin | -0.08 | 0.09 | -0.28 | 0.5 | 7.07 | 1916.41 | outside |
| 859 | 877 | TVLPPLLTDEMIAQYTSA | 0.2552 | 18 | ABCPRED,FBCPred | 2 | NON-ALLERGEN | Non-Toxin | 0.04 | 0.43 | -0.44 | -2 | 3.67 | 1963.54 | outside |
| 420 | 436 | DYNYKLPDDFTGCVIA | 0.3579 | 16 | ABCPRED | 1 | NON-ALLERGEN | Non-Toxin | -0.08 | -0.23 | -0.12 | -2 | 3.94 | 1834.25 | outside |
| 1075 | 1091 | FTTAPAICHDGKAHFP | 0.1129 | 16 | ABCPRED | 1 | ALLERGEN | Non-Toxin | -0.01 | -0.05 | -0.32 | 1 | 7.22 | 1713.16 | outside |
| 671 | 687 | CASYQTQTNSPRRARS | 0.2042 | 16 | ABCPRED | 1 | NON-ALLERGEN | Non-Toxin | -0.5 | -1.54 | 0.34 | 3 | 10.77 | 1826.18 | outside |
| 384 | 400 | PTKLNDLCFTNVYADS | 1.2303 | 16 | ABCPRED | 1 | NON-ALLERGEN | Non-Toxin | -0.12 | -0.26 | -0.16 | -1 | 4.21 | 1801.22 | outside |
| 1060 | 1078 | VVFLHVTYVPAQEKNFTT | 0.9224 | 18 | ABCPRED | 1 | NON-ALLERGEN | Non-Toxin | 0.01 | 0.3 | -0.61 | 0.5 | 7.1 | 2093.68 | outside |
| 417 | 433 | KIADYNYKLPDDFTGC | 0.5559 | 16 | ABCPRED | 1 | ALLERGEN | Non-Toxin | -0.18 | -0.73 | 0.16 | -1 | 4.43 | 1863.29 | outside |
| 1064 | 1076 | HVTYVPAQEKNF | 1.1353 | 12 | AAP,BCPREDS,FBCPred | 3 | NON-ALLERGEN | Non-Toxin | -0.15 | -0.68 | -0.23 | 0.5 | 7.1 | 1432.77 | outside |
| 766 | 784 | ALTGIAVEQDKNTQEVFA | 0.5729 | 18 | ABCPRED | 1 | NON-ALLERGEN | Non-Toxin | -0.09 | -0.1 | 0.07 | -2 | 4.14 | 1934.4 | outside |
| 161 | 179 | SSANNCTFEYVSQPFLMD | 0.1629 | 18 | ABCPRED | 1 | NON-ALLERGEN | Non-Toxin | -0.08 | -0.21 | -0.35 | -2 | 3.67 | 2053.49 | outside |
| 375 | 391 | STFKCYGVSPTKLNDL | 1.2492 | 16 | ABCPRED | 1 | NON-ALLERGEN | Non-Toxin | -0.13 | -0.25 | -0.12 | 1 | 8.51 | 1773.26 | outside |
| 990 | 1006 | EVQIDRLITGRLQSLQ | -0.2543 | 16 | ABCPRED | 1 | ALLERGEN | Non-Toxin | -0.23 | -0.24 | 0.13 | 0 | 6.42 | 1869.41 | outside |
| 720 | 738 | ISVTTEILPVSMTKTSVD | 1.1887 | 18 | ABCPRED | 1 | NON-ALLERGEN | Non-Toxin | -0.01 | 0.53 | -0.16 | -1 | 4.38 | 1921.5 | outside |
| 23 | 39 | QLPPAYTNSFTRGVYY | 0.1744 | 16 | ABCPRED | 1 | NON-ALLERGEN | Non-Toxin | -0.11 | -0.54 | -0.64 | 1 | 8.83 | 1877.31 | outside |
| 19 | 39 | TTRTQLPPAYTNSFTRGVYY | 0.434 | 20 | BCPREDS | 1 | NON-ALLERGEN | Non-Toxin | -0.2 | -0.76 | -0.42 | 2 | 9.72 | 2336.87 | outside |
| 450 | 460 | NYLYRLFRKS | -0.4937 | 10 | ABCPRED | 1 | ALLERGEN | Non-Toxin | -0.38 | -0.94 | -0.12 | 3 | 10.29 | 1359.73 | outside |
| 903 | 921 | AYRFNGIGVTQNVLYENQ | 0.6989 | 18 | ABCPRED,FBCPred | 2 | ALLERGEN | Non-Toxin | -0.13 | -0.46 | -0.42 | 0 | 6.35 | 2086.57 | outside |
| 778 | 794 | TQEVFAQVKQIYKTPP | 0.2211 | 16 | ABCPRED | 1 | NON-ALLERGEN | Non-Toxin | -0.17 | -0.64 | -0.08 | 1 | 8.83 | 1877.41 | outside |
| 262 | 276 | AAAYYVGYLQPRTF | 0.6905 | 14 | ABCPRED,AAP | 2 | NON-ALLERGEN | Non-Toxin | 0 | 0.11 | -0.81 | 1 | 8.83 | 1620.03 | outside |
| 790 | 802 | KTPPIKDFGGFN | -0.2264 | 12 | ABCPRED | 1 | NON-ALLERGEN | Non-Toxin | -0.13 | -0.78 | 0.17 | 1 | 8.94 | 1320.68 | outside |
| 515 | 535 | FELLHAPATVCGPKKSTNLV | 0.3229 | 20 | ABCPRED | 1 | NON-ALLERGEN | Non-Toxin | -0.03 | 0.24 | -0.24 | 1.5 | 8.54 | 2125.81 | outside |
| 19 | 33 | TTRTQLPPAYTNSF | 0.6884 | 14 | AAP | 1 | NON-ALLERGEN | Non-Toxin | -0.2 | -0.8 | -0.36 | 1 | 9.1 | 1596.95 | outside |
| 103 | 115 | GWIFGTTLDSKT | 0.3472 | 12 | ABCPRED | 1 | ALLERGEN | Non-Toxin | 0 | -0.08 | -0.37 | 0 | 6.19 | 1325.66 | outside |
| 343 | 355 | NATRFASVYAWN | 0.0524 | 12 | ABCPRED | 1 | NON-ALLERGEN | Non-Toxin | -0.1 | -0.23 | -0.66 | 1 | 9.1 | 1399.68 | outside |
| 500 | 512 | TNGVGYQPYRVV | 0.5085 | 12 | ABCPRED | 1 | NON-ALLERGEN | Non-Toxin | -0.11 | -0.38 | -0.51 | 1 | 8.93 | 1352.69 | outside |
| 434 | 450 | IAWNSNNLDSKVGGNY | 0.906 | 16 | ABCPRED | 1 | NON-ALLERGEN | Non-Toxin | -0.13 | -0.73 | -0.24 | 0 | 6.18 | 1752.11 | outside |
| 1043 | 1059 | CGKGYHLMSFPQSAPH | 0.0624 | 16 | ABCPRED | 1 | NON-ALLERGEN | Non-Toxin | -0.08 | -0.49 | -0.41 | 2 | 8.54 | 1760.25 | outside |
| 787 | 800 | QIYKTPPIKDFGG | -0.5194 | 13 | literature | 1 | NON-ALLERGEN | Non-Toxin | -0.12 | -0.69 | 0.03 | 1 | 8.83 | 1463.89 | outside |
| 1073 | 1091 | KNFTTAPAICHDGKAHFP | 0.139 | 18 | ABCPRED | 1 | ALLERGEN | Non-Toxin | -0.11 | -0.46 | -0.11 | 2 | 8.54 | 1955.47 | outside |
| 1247 | 1265 | CCSCGSCCKFDEDDSEPV | 0.2781 | 18 | ABCPRED | 1 | NON-ALLERGEN | Non-Toxin | -0.21 | -0.35 | 0.55 | -4 | 3.77 | 1927.31 | inside |
| 289 | 307 | VDCALDPLSETKCTLKSF | 0.7161 | 18 | ABCPRED,FBCPred | 2 | NON-ALLERGEN | Non-Toxin | -0.12 | 0.13 | 0.16 | -1 | 4.56 | 1970.53 | outside |
| 294 | 308 | DPLSETKCTLKSFT | 0.59 | 14 | FBCPred | 1 | NON-ALLERGEN | Non-Toxin | -0.21 | -0.51 | 0.31 | 0 | 6.38 | 1569.98 | outside |
| 574 | 584 | DAVRDPQTLE | 0.0812 | 10 | ABCPRED | 1 | NON-ALLERGEN | Non-Toxin | -0.34 | -1.1 | 0.8 | -2 | 4.03 | 1143.35 | outside |
| 144 | 156 | YYHKNNKSWMES | 0.178 | 12 | ABCPRED | 1 | NON-ALLERGEN | Non-Toxin | -0.36 | -2.06 | 0.02 | 1.5 | 8.77 | 1586.91 | outside |
| 672 | 689 | ASYQTQTNSPRRARSVA | 0.3628 | 17 | BCEPRED | 1 | NON-ALLERGEN | Non-Toxin | -0.42 | -1.24 | 0.26 | 3 | 11.72 | 1893.27 | outside |
| 287 | 307 | DAVDCALDPLSETKCTLKSF | 0.682 | 20 | ABCPRED,FBCPred | 2 | ALLERGEN | Non-Toxin | -0.13 | 0.03 | 0.27 | -2 | 4.23 | 2156.72 | outside |
| 1074 | 1090 | NFTTAPAICHDGKAHF | 0.3817 | 16 | ABCPRED | 1 | ALLERGEN | Non-Toxin | -0.05 | -0.17 | -0.31 | 1 | 7.22 | 1730.15 | outside |
| 566 | 584 | GRDIADTTDAVRDPQTLE | 0.461 | 18 | FBCPred | 1 | NON-ALLERGEN | Non-Toxin | -0.33 | -1 | 0.77 | -3 | 4.04 | 1973.33 | outside |
| 154 | 172 | ESEFRVYSSANNCTFEYV | 0.1152 | 18 | FBCPred | 1 | NON-ALLERGEN | Non-Toxin | -0.18 | -0.52 | -0.07 | -2 | 4.26 | 2145.52 | outside |
| 817 | 834 | FIEDLLFNKVTLADAGF | 0.5301 | 17 | COVIDep | 1 | NON-ALLERGEN | Non-Toxin | 0.08 | 0.77 | -0.32 | -2 | 4.03 | 1913.45 | outside |
| 419 | 429 | ADYNYKLPDD | 0.6956 | 10 | IEDB | 1 | NON-ALLERGEN | Non-Toxin | -0.31 | -1.65 | 0.53 | -2 | 3.94 | 1213.39 | outside |
| 432 | 448 | CVIAWNSNNLDSKVGG | 0.7723 | 16 | ABCPRED | 1 | NON-ALLERGEN | Non-Toxin | -0.06 | -0.01 | -0.27 | 0 | 6.16 | 1677.1 | outside |
| 684 | 702 | ARSVASQSIIAYTMSLGA | 0.3963 | 18 | ABCPRED | 1 | NON-ALLERGEN | Non-Toxin | 0.02 | 0.69 | -0.47 | 1 | 9.1 | 1826.35 | outside |
| 679 | 697 | NSPRRARSVASQSIIAYT | 0.1489 | 18 | AAP | 1 | NON-ALLERGEN | Non-Toxin | -0.28 | -0.48 | 0.07 | 3 | 11.72 | 1977.45 | outside |
| 672 | 688 | ASYQTQTNSPRRARSV | 0.3062 | 16 | ABCPRED | 1 | ALLERGEN | Non-Toxin | -0.47 | -1.43 | 0.31 | 3 | 11.72 | 1822.18 | outside |
| 178 | 198 | DLEGKQGNFKNLREFVFKNI | 0.5535 | 20 | ABCPRED,literature | 2 | NON-ALLERGEN | Non-Toxin | -0.26 | -0.84 | 0.37 | 1 | 8.83 | 2397.04 | outside |
| 577 | 585 | RDPQTLEI | 0.9248 | 8 | ellipro | 1 | ALLERGEN | Non-Toxin | -0.35 | -1.12 | 0.65 | -1 | 4.38 | 971.19 | outside |
| 506 | 524 | QPYRVVVLSFELLHAPAT | 0.69 | 18 | ABCPRED | 1 | NON-ALLERGEN | Non-Toxin | 0.02 | 0.54 | -0.56 | 0.5 | 7.1 | 2040.66 | outside |
| 1016 | 1034 | AEIRASANLAATKMSECV | 0.5374 | 18 | ABCPRED | 1 | NON-ALLERGEN | Non-Toxin | -0.12 | 0.26 | 0.14 | 0 | 6.46 | 1865.39 | outside |
| 564 | 582 | QFGRDIADTTDAVRDPQT | 0.0807 | 18 | ABCPRED | 1 | NON-ALLERGEN | Non-Toxin | -0.32 | -1.06 | 0.58 | -2 | 4.14 | 2006.36 | outside |
| 401 | 419 | VIRGDEVRQIAPGQTGKI | 0.6123 | 18 | AAP | 1 | ALLERGEN | Non-Toxin | -0.2 | -0.37 | 0.34 | 1 | 9.1 | 1937.51 | outside |

| | | | | | | | | | | | | | | | |
|---|---|---|---|---|---|---|---|---|---|---|---|---|---|---|---|
| 897 | 914 | PFAMQMAYRFNGIGVTQ | 1.3306 | 17 | COVIDep,literature | 2 | NON-ALLERGEN | Non-Toxin | -0.01 | 0.14 | -0.65 | 1 | 9.1 | 1931.51 | outside |
| 893 | 909 | ALQIPFAMQMAYRFNG | 0.6875 | 16 | ABCPRED | 1 | NON-ALLERGEN | Non-Toxin | 0 | 0.3 | -0.71 | 1 | 9.1 | 1858.44 | outside |
| 835 | 851 | KQYGDCLGDIAARDLI | 0.4125 | 16 | ABCPRED | 1 | NON-ALLERGEN | Non-Toxin | -0.14 | -0.11 | 0.23 | -1 | 4.43 | 1751.22 | outside |
| 1134 | 1173 | TVYDPLQPELDSFKEELDKYFKNHTSPDVDLGDI | 0.1406 | 39 | IEDB | 1 | NON-ALLERGEN | Non-Toxin | -0.19 | -0.88 | 0.33 | -5.5 | 4.13 | 4455.41 | outside |
| 44 | 62 | RSSVLHSTQDLFLPFFSN | 0.2233 | 18 | ABCPRED | 1 | NON-ALLERGEN | Non-Toxin | -0.09 | 0.02 | -0.43 | 0.5 | 7.1 | 2095.6 | outside |
| 205 | 217 | SKHTPINLVRDL | 0.4662 | 12 | ABCPRED | 1 | NON-ALLERGEN | Non-Toxin | -0.23 | -0.45 | 0.14 | 1.5 | 9.1 | 1392.8 | outside |
| 292 | 310 | ALDPLSETKCTLKSFTVE | 0.7706 | 18 | ABCPRED | 1 | ALLERGEN | Non-Toxin | -0.13 | -0.05 | 0.19 | -1 | 4.68 | 1982.53 | outside |
| 349 | 361 | SVYAWNRKRISN | 0.4858 | 12 | ABCPRED | 1 | NON-ALLERGEN | Non-Toxin | -0.38 | -1.1 | 0.04 | 3 | 11.01 | 1493.85 | outside |
| 975 | 995 | SVLNDILSRLDKVEAEVQID | 0.0524 | 20 | ABCPRED | 1 | ALLERGEN | Non-Toxin | -0.16 | 0.02 | 0.4 | -3 | 4.11 | 2256.84 | outside |
| 22 | 42 | TQLPPAYTNSFTRGVYYPDK | 0.2871 | 20 | ABCPRED,AAP | 2 | NON-ALLERGEN | Toxin | -0.19 | -0.92 | -0.23 | 1 | 8.76 | 2318.85 | outside |
| 628 | 644 | QLTPTWRVYSTGSNVF | 0.6678 | 16 | ABCPRED | 1 | ALLERGEN | Non-Toxin | -0.09 | -0.27 | -0.64 | 1 | 9.1 | 1856.3 | outside |
| 1134 | 1152 | NNTVYDPLQPELDSFKEE | 0.1001 | 18 | ABCPRED | 1 | NON-ALLERGEN | Non-Toxin | -0.26 | -1.29 | 0.48 | -4 | 3.84 | 2138.53 | outside |
| 226 | 244 | LVDLPIGINITRFQTLLA | 0.786 | 18 | ABCPRED | 1 | NON-ALLERGEN | Non-Toxin | 0.09 | 1.06 | -0.64 | 0 | 6.19 | 1997.7 | outside |
| 508 | 526 | YRVVVLSFELLHAPATVC | 0.7718 | 18 | ABCPRED | 1 | NON-ALLERGEN | Non-Toxin | 0.09 | 1.19 | -0.71 | 0.5 | 7.06 | 2017.68 | outside |
| 574 | 590 | DAVRDPQTLEILDITP | 1.1133 | 16 | ABCPRED | 1 | NON-ALLERGEN | Non-Toxin | -0.15 | -0.25 | 0.33 | -3 | 3.84 | 1796.24 | outside |
| 1018 | 1036 | IRASANLAATKMSECVLG | 0.4521 | 18 | ABCPRED | 1 | NON-ALLERGEN | Non-Toxin | -0.06 | 0.54 | -0.1 | 1 | 8.57 | 1835.42 | outside |
| 1069 | 1087 | PAQEKNFTTAPAICHDGK | 0.3179 | 18 | ABCPRED | 1 | NON-ALLERGEN | Non-Toxin | -0.19 | -0.82 | 0.24 | 0.5 | 7.06 | 1928.4 | outside |
| 84 | 102 | LPFNDGVYFASTEKSNII | 0.6088 | 18 | ABCPRED | 1 | NON-ALLERGEN | Non-Toxin | -0.02 | 0.05 | -0.28 | -1 | 4.38 | 2015.51 | outside |
| 347 | 363 | FASVYAWNRKRISNCV | 0.3371 | 16 | ABCPRED | 1 | NON-ALLERGEN | Non-Toxin | -0.19 | -0.12 | -0.31 | 3 | 10.07 | 1914.43 | outside |
| 902 | 920 | MAYRFNGIGVTQNVLYEN | 0.7259 | 18 | ABCPRED | 1 | NON-ALLERGEN | Non-Toxin | -0.08 | -0.16 | -0.51 | 0 | 6.35 | 2089.63 | outside |
| 1019 | 1037 | RASANLAATKMSECVLGQ | 0.4001 | 18 | ABCPRED | 1 | NON-ALLERGEN | Non-Toxin | -0.14 | 0.1 | 0.01 | 1 | 8.57 | 1850.39 | outside |
| 354 | 372 | NRKRISNCVADYSVLYNS | 0.0152 | 18 | ABCPRED | 1 | NON-ALLERGEN | Non-Toxin | -0.3 | -0.61 | 0.04 | 2 | 9.25 | 2102.6 | outside |
| 978 | 996 | NDILSRLDKVEAEVQIDR | -0.0673 | 18 | FBCPred | 1 | NON-ALLERGEN | Non-Toxin | -0.32 | -0.63 | 0.78 | -2 | 4.44 | 2113.62 | outside |
| 974 | 994 | SSVLNDILSRLDKVEAEVQI | 0.0516 | 20 | ABCPRED,FBCPred | 2 | NON-ALLERGEN | Non-Toxin | -0.14 | 0.15 | 0.26 | -2 | 4.32 | 2228.83 | outside |
| 355 | 373 | RKRISNCVADYSVLYNSA | -0.0939 | 18 | ABCPRED | 1 | NON-ALLERGEN | Non-Toxin | -0.25 | -0.31 | 0.01 | 2 | 9.25 | 2059.57 | outside |
| 842 | 860 | GDIAARDLICAQKFNGLT | 0.5523 | 18 | ABCPRED | 1 | NON-ALLERGEN | Non-Toxin | -0.09 | 0.19 | -0.01 | 0 | 6.28 | 1906.45 | outside |
| 1179 | 1195 | IQKEIDRLNEVAKNLN | 0.0407 | 16 | ABCPRED | 1 | NON-ALLERGEN | Non-Toxin | -0.33 | -0.89 | 0.6 | 0 | 6.53 | 1897.41 | outside |
| 392 | 410 | FTNVYADSFVIRGDEVRQ | -0.1517 | 18 | ABCPRED | 1 | NON-ALLERGEN | Non-Toxin | -0.19 | -0.29 | 0.07 | -1 | 4.56 | 2116.57 | outside |
| 304 | 322 | KSFTVEKGIYQTSNFRVQ | 0.379 | 18 | ABCPRED | 1 | NON-ALLERGEN | Non-Toxin | -0.24 | -0.69 | 0.02 | 2 | 9.72 | 2132.67 | outside |
| 405 | 421 | DEVRQIAPGQTGKIAD | 1.0388 | 16 | ABCPRED | 1 | NON-ALLERGEN | Non-Toxin | -0.23 | -0.76 | 0.56 | -1 | 4.56 | 1698.1 | outside |
| 628 | 640 | QLTPTWRVYSTG | 1.1857 | 12 | ABCPRED | 1 | NON-ALLERGEN | Non-Toxin | -0.14 | -0.59 | -0.56 | 1 | 9.1 | 1408.75 | outside |
| 1206 | 1220 | YEQYIKWPWYIWLG | 0.8422 | 14 | BCPREDS | 1 | NON-ALLERGEN | Non-Toxin | 0.06 | -0.48 | -1.16 | 0 | 6.34 | 1945.47 | outside_transmembrane-region |
| 840 | 858 | CLGDIAARDLICAQKFNG | 0.5394 | 18 | ABCPRED | 1 | NON-ALLERGEN | Non-Toxin | -0.07 | 0.37 | -0.04 | 0 | 6.25 | 1908.48 | outside |
| 322 | 342 | PTESIVRFPNITNLCPFGEV | 0.4532 | 20 | FBCPred | 1 | ALLERGEN | Non-Toxin | -0.03 | 0.17 | -0.28 | -1 | 4.54 | 2233.86 | outside |
| 504 | 522 | GYQPYRVVVLSFELLHAP | 0.8989 | 18 | ABCPRED | 1 | ALLERGEN | Non-Toxin | 0.02 | 0.38 | -0.64 | 0.5 | 7.1 | 2088.71 | outside |
| 577 | 587 | RDPQTLEILD | 1.0339 | 10 | ABCPRED | 1 | ALLERGEN | Non-Toxin | -0.3 | -0.87 | 0.64 | -2 | 4.03 | 1199.47 | outside |
| 559 | 577 | FLPFQQFGRDIADTTDAV | 0.3733 | 18 | ABCPRED | 1 | NON-ALLERGEN | Non-Toxin | -0.08 | -0.05 | -0.11 | -2 | 3.94 | 2041.5 | outside |
| 629 | 645 | LTPTWRVYSTGSNVFQ | 0.5257 | 16 | ABCPRED | 1 | NON-ALLERGEN | Non-Toxin | -0.09 | -0.28 | -0.64 | 1 | 9.1 | 1856.3 | outside |
| 977 | 995 | LNDILSRLDKVEAEVQID | 0.1894 | 18 | ABCPRED | 1 | NON-ALLERGEN | Non-Toxin | -0.19 | -0.17 | 0.51 | -3 | 4.11 | 2070.6 | outside |
| 567 | 587 | RDIADTTDAVRDPQTLEILD | 0.643 | 20 | ABCPRED | 1 | NON-ALLERGEN | Non-Toxin | -0.27 | -0.64 | 0.66 | -4 | 3.91 | 2257.72 | outside |
| 304 | 323 | KSFTVEKGIYQTSNFRVQP | 0.5729 | 19 | IEDB,literature | 2 | NON-ALLERGEN | Non-Toxin | -0.23 | -0.74 | 0.02 | 2 | 9.72 | 2229.8 | outside |
| 395 | 413 | VYADSFVIRGDEVRQIAP | 0.0171 | 18 | ABCPRED | 1 | NON-ALLERGEN | Non-Toxin | -0.12 | 0.05 | 0.09 | -1 | 4.56 | 2035.54 | outside |
| 70 | 81 | VSGTNGTKRFD | 0.8493 | 11 | IEDB | 1 | NON-ALLERGEN | Non-Toxin | -0.31 | -1.04 | 0.43 | 1 | 9.1 | 1181.42 | outside |
| 350 | 360 | VYAWNRKRIS | 0.6137 | 10 | ABCPRED | 1 | NON-ALLERGEN | Non-Toxin | -0.36 | -0.89 | 0 | 3 | 11.01 | 1292.64 | outside |
| 506 | 518 | QPYRVVVLSFEL | 0.9785 | 12 | ABCPRED | 1 | NON-ALLERGEN | Non-Toxin | -0.01 | 0.65 | -0.53 | 0 | 6.35 | 1449.89 | outside |
| 1238 | 1265 | TSCCSCLKGCCSCGSCCKFDEDDSEPV | 0.1337 | 27 | IEDB | 1 | NON-ALLERGEN | Toxin | -0.18 | -0.06 | 0.31 | -3 | 4.11 | 2810.5 | inside |
| 575 | 585 | AVRDPQTLEI | 0.7083 | 10 | ABCPRED | 1 | ALLERGEN | Non-Toxin | -0.2 | -0.3 | 0.32 | -1 | 4.38 | 1141.43 | outside |
| 184 | 194 | GNFKNLREFV | -0.2373 | 10 | ABCPRED | 1 | NON-ALLERGEN | Non-Toxin | -0.23 | -0.57 | 0.11 | 1 | 9.1 | 1223.54 | outside |
| 679 | 699 | NSPRRARSVASQSIIAYTMS | 0.1656 | 20 | ABCPRED | 1 | NON-ALLERGEN | Non-Toxin | -0.26 | -0.38 | 0.02 | 3 | 11.72 | 2195.75 | outside |
| 627 | 639 | DQLTPTWRVYST | 0.7559 | 12 | AAP | 1 | NON-ALLERGEN | Non-Toxin | -0.21 | -0.85 | -0.31 | 0 | 6.19 | 1466.78 | outside |
| 441 | 459 | LDSKVGGNYNYLYRLFRK | 0.1027 | 18 | ABCPRED | 1 | NON-ALLERGEN | Non-Toxin | -0.27 | -0.8 | -0.03 | 3 | 9.83 | 2206.8 | outside |
| 1199 | 1217 | DLQELGKYEQYIKWPWYI | 0.8917 | 18 | ABCPRED | 1 | NON-ALLERGEN | Non-Toxin | -0.12 | -0.91 | -0.31 | -1 | 4.68 | 2372.99 | outside_transmembrane-region |
| 974 | 992 | SSVLNDILSRLDKVEAEV | -0.1243 | 18 | ABCPRED | 1 | NON-ALLERGEN | Non-Toxin | -0.15 | 0.11 | 0.38 | -2 | 4.32 | 1987.5 | outside |
| 141 | 157 | LGVYYHKNNKSWMESE | 0.5923 | 16 | ABCPRED | 1 | ALLERGEN | Non-Toxin | -0.23 | -1.29 | -0.01 | 0.5 | 7.09 | 1985.44 | outside |
| 79 | 99 | FDNPVLPFNDGVYFASTEKS | 0.3218 | 20 | ABCPRED,AAP | 2 | NON-ALLERGEN | Non-Toxin | -0.07 | -0.31 | -0.12 | -2 | 4.03 | 2247.72 | outside |
| 182 | 198 | KQGNFKNLREFVFKNI | 0.28 | 16 | ABCPRED | 1 | ALLERGEN | Non-Toxin | -0.28 | -0.83 | 0.2 | 3 | 10.3 | 1982.56 | outside |
| 763 | 779 | LNRALTGIAVEQDKNT | 0.3795 | 16 | ABCPRED | 1 | NON-ALLERGEN | Non-Toxin | -0.22 | -0.49 | 0.24 | 0 | 6.42 | 1743.19 | outside |
| 356 | 376 | KRISNCVADYSVLYNSASFS | 0.1401 | 20 | ABCPRED | 1 | NON-ALLERGEN | Non-Toxin | -0.13 | 0.01 | -0.24 | 1 | 8.51 | 2224.74 | outside |
| 1179 | 1192 | IQKEIDRLNEVAK | -0.1773 | 13 | BCEPRED | 1 | NON-ALLERGEN | Non-Toxin | -0.34 | -0.85 | 0.85 | 0 | 6.53 | 1555.99 | outside |
| 835 | 855 | KQYGDCLGDIAARDLICAQK | 0.6711 | 20 | ABCPRED | 1 | NON-ALLERGEN | Non-Toxin | -0.19 | -0.25 | 0.27 | 0 | 6.32 | 2181.8 | outside |
| 71 | 84 | SGTNGTKRFDNPV | 0.3648 | 13 | BCEPRED,IEDB | 2 | NON-ALLERGEN | Non-Toxin | -0.31 | -1.27 | 0.38 | 1 | 9.1 | 1392.67 | outside |
| 424 | 442 | KLPDDFTGCVIAWNSNNL | -0.0227 | 18 | ABCPRED | 1 | ALLERGEN | Non-Toxin | -0.07 | -0.13 | -0.27 | -1 | 4.21 | 2007.51 | outside |
| 627 | 637 | DQLTPTWRVY | 0.6489 | 10 | ABCPRED,IEDB | 2 | ALLERGEN | Non-Toxin | -0.21 | -0.87 | -0.36 | 0 | 6.19 | 1278.57 | outside |
| 985 | 1005 | DKVEAEVQIDRLITGRLQSL | -0.0848 | 20 | ABCPRED | 1 | NON-ALLERGEN | Non-Toxin | -0.23 | -0.26 | 0.44 | -1 | 4.79 | 2283.92 | outside |
| 762 | 780 | QLNRALTGIAVEQDKNTQ | 0.396 | 18 | ABCPRED | 1 | NON-ALLERGEN | Non-Toxin | -0.27 | -0.82 | 0.24 | 0 | 6.42 | 1999.49 | outside |
| 891 | 909 | GAALQIPFAMQMAYRFNG | 0.586 | 18 | ABCPRED | 1 | NON-ALLERGEN | Non-Toxin | 0.03 | 0.34 | -0.66 | 1 | 9.1 | 1986.6 | outside |
| 344 | 362 | ATRFASVYAWNRKRISNC | 0.2851 | 18 | ABCPRED | 1 | NON-ALLERGEN | Non-Toxin | -0.29 | -0.53 | -0.08 | 4 | 10.92 | 2143.69 | outside |
| 1206 | 1224 | YEQYIKWPWYIWLGFIAG | 0.7812 | 18 | ABCPRED | 1 | ALLERGEN | Non-Toxin | 0.14 | 0.11 | -1.17 | 0 | 6.34 | 2334 | outside_transmembrane-region |
| 31 | 41 | SFTRGVYYPD | 0.5224 | 10 | ABCPRED | 1 | NON-ALLERGEN | Non-Toxin | -0.16 | -0.71 | -0.27 | 0 | 6.18 | 1204.43 | outside |
| 764 | 782 | NRALTGIAVEQDKNTQEV | 0.5339 | 18 | ABCPRED | 1 | ALLERGEN | Toxin | -0.27 | -0.8 | 0.41 | -1 | 4.68 | 1986.44 | outside |
| 1176 | 1194 | VVNIQKEIDRLNEVAKNL | 0.1409 | 18 | ABCPRED | 1 | ALLERGEN | Non-Toxin | -0.23 | -0.32 | 0.37 | 0 | 6.53 | 2095.71 | outside |
| 1032 | 1050 | CVLGQSKRVDFCGKGYHL | 1.1216 | 18 | ABCPRED | 1 | NON-ALLERGEN | Non-Toxin | -0.15 | -0.11 | -0.08 | 2.5 | 8.94 | 2010.63 | outside |
| 975 | 991 | SVLNDILSRLDKVEAE | -0.0959 | 16 | ABCPRED | 1 | NON-ALLERGEN | Non-Toxin | -0.19 | -0.09 | 0.51 | -2 | 4.32 | 1801.26 | outside |
| 835 | 849 | KQYGDCLGDIAARD | 0.6965 | 14 | AAP | 1 | NON-ALLERGEN | Non-Toxin | -0.26 | -0.72 | 0.52 | -1 | 4.43 | 1524.86 | outside |
| 179 | 197 | LEGKQGNFKNLREFVFKN | 0.4969 | 18 | ABCPRED | 1 | ALLERGEN | Toxin | -0.29 | -0.99 | 0.34 | 2 | 9.72 | 2168.76 | outside |
| 843 | 861 | DIAARDLICAQKFNGLTV | 0.5841 | 18 | ABCPRED | 1 | NON-ALLERGEN | Non-Toxin | -0.07 | 0.44 | -0.09 | 0 | 6.28 | 1948.53 | outside |
| 1071 | 1089 | QEKNFTTAPAICHDGKAH | 0.3398 | 18 | ABCPRED | 1 | NON-ALLERGEN | Non-Toxin | -0.21 | -0.91 | 0.21 | 1 | 7.22 | 1968.43 | outside |

| | | | | | | | | | | | | | | | |
|---|---|---|---|---|---|---|---|---|---|---|---|---|---|---|---|
| 629 | 639 | LTPTWRVYST | 1.1346 | 10 | ABCPRED | 1 | ALLERGEN | Non-Toxin | -0.12 | -0.32 | -0.69 | 1 | 9.1 | 1223.53 | outside |
| 1139 | 1163 | DPLQPELDSFKEELDKYFKNHTSP | -0.3378 | 24 | IEDB | 1 | NON-ALLERGEN | Non-Toxin | -0.3 | -1.43 | 0.6 | -2.5 | 4.59 | 2878.49 | outside |
| 631 | 641 | PTWRVYSTGS | 0.6815 | 10 | ABCPRED | 1 | ALLERGEN | Non-Toxin | -0.16 | -0.75 | -0.44 | 1 | 9.1 | 1153.39 | outside |
| 330 | 348 | PNITNLCPFGEVFNATRF | 0.6006 | 18 | ABCPRED | 1 | NON-ALLERGEN | Non-Toxin | -0.04 | 0.09 | -0.46 | 0 | 6.32 | 2040.58 | outside |
| 317 | 327 | NFRVQPTESI | 1.2412 | 10 | ABCPRED | 1 | ALLERGEN | Non-Toxin | -0.23 | -0.66 | 0.05 | 0 | 6.36 | 1190.46 | outside |
| 264 | 282 | AYYVGYLQPRTFLLKYNE | 0.4672 | 18 | ABCPRED | 1 | ALLERGEN | Non-Toxin | -0.1 | -0.37 | -0.56 | 1 | 8.71 | 2238.84 | outside |
| 101 | 113 | IRGWIFGTTLDS | -0.3858 | 12 | ABCPRED,AAP | 2 | ALLERGEN | Non-Toxin | 0.02 | 0.31 | -0.48 | 0 | 6.19 | 1365.73 | outside |
| 1015 | 1033 | AAEIRASANLAATKMSEC | 0.5496 | 18 | ABCPRED | 1 | NON-ALLERGEN | Non-Toxin | -0.13 | 0.13 | 0.19 | 0 | 6.46 | 1837.33 | outside |
| 1203 | 1219 | LGKYEQYIKWPWYIWL | 0.735 | 16 | ABCPRED | 1 | ALLERGEN | Non-Toxin | 0.02 | -0.43 | -0.94 | 1 | 8.71 | 2186.84 | outside_transmembrane-region |
| 1202 | 1220 | ELGKYEQYIKWPWYIWLG | 0.8715 | 18 | ABCPRED | 1 | NON-ALLERGEN | Non-Toxin | -0.01 | -0.59 | -0.67 | 0 | 6.48 | 2373.04 | outside_transmembrane-region |
| 352 | 362 | AWNRKRISNC | 0.6063 | 10 | ABCPRED | 1 | ALLERGEN | Non-Toxin | -0.48 | -1.28 | 0.3 | 3 | 10.87 | 1247.57 | outside |
| 98 | 114 | SNIIRGWIFGTTLDSK | -0.0513 | 16 | ABCPRED | 1 | NON-ALLERGEN | Non-Toxin | -0.07 | 0 | -0.26 | 1 | 9.1 | 1808.31 | outside |
| 1251 | 1271 | GSCCKFDEDDSEPVLKGVKL | 0.6126 | 20 | AAP | 1 | NON-ALLERGEN | Non-Toxin | -0.21 | -0.47 | 0.68 | -2 | 4.44 | 2169.73 | inside |
| 505 | 525 | YQPYRVVVLSFELLHAPATV | 0.7299 | 20 | ABCPRED | 1 | ALLERGEN | Non-Toxin | 0.04 | 0.63 | -0.7 | 0.5 | 7.1 | 2303 | outside |
| 982 | 993 | SRLDKVEAEVQ | 0.3878 | 11 | BCEPRED | 1 | NON-ALLERGEN | Non-Toxin | -0.36 | -0.84 | 0.93 | -1 | 4.68 | 1273.56 | outside |
| 1135 | 1152 | NTVYDPLQPELDSFKEE | 0.0864 | 17 | literature | 1 | ALLERGEN | Non-Toxin | -0.24 | -1.16 | 0.49 | -4 | 3.84 | 2024.41 | outside |
| 311 | 331 | GIYQTSNFRVQPTESIVRFP | 0.2931 | 20 | BCPREDS | 1 | ALLERGEN | Non-Toxin | -0.16 | -0.4 | -0.23 | 1 | 9.1 | 2339.93 | outside |
| 393 | 411 | TNVYADSFVIRGDEVRQI | -0.2055 | 18 | ABCPRED | 1 | NON-ALLERGEN | Non-Toxin | -0.18 | -0.19 | 0.11 | -1 | 4.56 | 2082.56 | outside |
| 184 | 196 | GNFKNLREFVFK | -0.1011 | 12 | ABCPRED | 1 | ALLERGEN | Non-Toxin | -0.23 | -0.57 | 0.13 | 2 | 10.01 | 1498.92 | outside |
| 980 | 997 | ILSRLDKVEAEVQIDRL | 0.052 | 17 | COVIDep,literature | 2 | NON-ALLERGEN | Non-Toxin | -0.23 | -0.04 | 0.53 | -1 | 4.79 | 1997.58 | outside |
| 290 | 308 | DCALDPLSETKCTLKSFT | 0.5804 | 18 | ABCPRED | 1 | NON-ALLERGEN | Non-Toxin | -0.16 | -0.14 | 0.22 | -1 | 4.56 | 1972.5 | outside |
| 844 | 860 | IAARDLICAQKFNGLT | 0.4407 | 16 | ABCPRED | 1 | ALLERGEN | Non-Toxin | -0.06 | 0.46 | -0.2 | 1 | 8.57 | 1734.28 | outside |
| 627 | 647 | DQLTPTWRVYSTGSNVFQTR | 0.5975 | 20 | BCPREDS | 1 | ALLERGEN | Non-Toxin | -0.24 | -0.83 | -0.22 | 1 | 9.1 | 2356.87 | outside |
| 681 | 699 | PRRARSVASQSIIAYTMS | 0.3249 | 18 | ABCPRED | 1 | NON-ALLERGEN | Non-Toxin | -0.23 | -0.18 | -0.01 | 3 | 11.72 | 1994.54 | outside |
| 1256 | 1270 | FDEDDSEPVLKGVK | 0.6734 | 14 | IEDB | 1 | NON-ALLERGEN | Non-Toxin | -0.25 | -0.94 | 1 | -3 | 4.11 | 1577.9 | inside |
| 986 | 1006 | KVEAEVQIDRLITGRLQSLQ | -0.0616 | 20 | ABCPRED | 1 | NON-ALLERGEN | Non-Toxin | -0.23 | -0.26 | 0.3 | 0 | 6.53 | 2296.97 | outside |
| 1104 | 1114 | VTQRNFYEPQ | 0.6509 | 10 | BCEPRED | 1 | NON-ALLERGEN | Non-Toxin | -0.35 | -1.51 | -0.01 | 0 | 6.35 | 1281.53 | outside |
| 318 | 330 | FRVQPTESIVRF | 0.3221 | 12 | ABCPRED | 1 | NON-ALLERGEN | Non-Toxin | -0.19 | -0.05 | -0.06 | 1 | 9.95 | 1478.88 | outside |
| 314 | 332 | QTSNFRVQPTESIVRFPN | 0.3397 | 18 | ABCPRED,AAP,BCPREDS | 3 | ALLERGEN | Non-Toxin | -0.27 | -0.79 | -0.01 | 1 | 9.95 | 2120.61 | outside |
| 348 | 364 | ASVYAWNRKRISNCVA | 0.3032 | 16 | ABCPRED | 1 | NON-ALLERGEN | Non-Toxin | -0.21 | -0.18 | -0.19 | 3 | 10.07 | 1838.33 | outside |
| 984 | 1000 | LDKVEAEVQIDRLITG | -0.2015 | 16 | ABCPRED | 1 | NON-ALLERGEN | Non-Toxin | -0.15 | -0.01 | 0.44 | -2 | 4.32 | 1799.3 | outside |
| 314 | 328 | QTSNFRVQPTESIV | 0.7811 | 14 | AAP | 1 | NON-ALLERGEN | Non-Toxin | -0.21 | -0.53 | -0.06 | 0 | 6.36 | 1605.97 | outside |
| 1082 | 1094 | CHDGKAHFPREG | -0.4455 | 12 | BCPREDS | 1 | NON-ALLERGEN | Non-Toxin | -0.32 | -1.43 | 0.58 | 1 | 7.22 | 1353.64 | outside |
| 897 | 915 | PFAMQMAYRFNGIGVTQN | 1.2715 | 18 | ABCPRED | 1 | NON-ALLERGEN | Non-Toxin | -0.05 | -0.07 | -0.6 | 1 | 9.1 | 2045.63 | outside |
| 44 | 64 | RSSVLHSTQDLFLPFFSNVT | 0.3684 | 20 | ABCPRED | 1 | NON-ALLERGEN | Non-Toxin | -0.06 | 0.19 | -0.48 | 0.5 | 7.1 | 2295.87 | outside |
| 180 | 194 | EGKQGNFKNLREFV | 0.3419 | 14 | ABCPRED,AAP | 2 | NON-ALLERGEN | Non-Toxin | -0.33 | -1.21 | 0.52 | 1 | 8.93 | 1666.08 | outside |
| 572 | 590 | TTDAVRDPQTLEILDITP | 1.1067 | 18 | AAP | 1 | NON-ALLERGEN | Non-Toxin | -0.15 | -0.3 | 0.24 | -3 | 3.84 | 1998.48 | outside |
| 1176 | 1192 | VVNIQKEIDRLNEVAK | 0.1859 | 16 | ABCPRED | 1 | NON-ALLERGEN | Non-Toxin | -0.25 | -0.38 | 0.51 | 0 | 6.53 | 1868.41 | outside |
| 1249 | 1267 | SCGSCCKFDEDDSEPVLK | 0.3129 | 18 | ABCPRED | 1 | NON-ALLERGEN | Non-Toxin | -0.25 | -0.63 | 0.73 | -3 | 4.11 | 1962.38 | inside |
| 313 | 329 | YQTSNFRVQPTESIVR | 0.4863 | 16 | ABCPRED | 1 | NON-ALLERGEN | Non-Toxin | -0.29 | -0.83 | -0.01 | 1 | 9.1 | 1925.36 | outside |
| 406 | 422 | EVRQIAPGQTGKIADY | 1.3837 | 16 | ABCPRED,literature | 2 | NON-ALLERGEN | Non-Toxin | -0.19 | -0.62 | 0.23 | 0 | 6.42 | 1746.19 | outside |
| 1147 | 1165 | SFKEELDKYFKNHTSPDV | -0.3025 | 18 | ABCPRED | 1 | NON-ALLERGEN | Non-Toxin | -0.3 | -1.33 | 0.57 | -0.5 | 5.5 | 2184.64 | outside |
| 465 | 477 | ERDISTEIYQAG | -0.1112 | 12 | ABCPRED | 1 | NON-ALLERGEN | Non-Toxin | -0.25 | -0.91 | 0.47 | -2 | 4.14 | 1381.63 | outside |
| 894 | 912 | LQIPFAMQMAYRFNGIGV | 1.1951 | 18 | ABCPRED | 1 | ALLERGEN | Non-Toxin | 0.07 | 0.63 | -0.79 | 1 | 9.1 | 2056.75 | outside |
| 185 | 197 | NFKNLREFVFKN | 0.1722 | 12 | BCEPRED | 1 | NON-ALLERGEN | Non-Toxin | -0.3 | -0.83 | 0.15 | 2 | 10.01 | 1555.97 | outside |
| 34 | 42 | RGVYYPDK | 1.0191 | 8 | IEDB,literature | 2 | NON-ALLERGEN | Non-Toxin | -0.36 | -1.54 | 0.36 | 1 | 8.83 | 997.22 | outside |
| 354 | 370 | NRKRISNCVADYSVLY | 0.0823 | 16 | ABCPRED | 1 | NON-ALLERGEN | Non-Toxin | -0.28 | -0.41 | 0.02 | 2 | 9.25 | 1901.39 | outside |
| 178 | 194 | DLEGKQGNFKNLREFV | 0.527 | 16 | ABCPRED | 1 | NON-ALLERGEN | Toxin | -0.3 | -1.04 | 0.53 | 0 | 6.53 | 1894.36 | outside |
| 1205 | 1219 | KYEQYIKWPWYIWL | 0.7469 | 14 | FBCPred | 1 | NON-ALLERGEN | Non-Toxin | -0.03 | -0.73 | -0.95 | 1 | 8.71 | 2016.59 | outside_transmembrane-region |
| 125 | 143 | NVVIKVCEFQFCNDPFLG | -0.1997 | 18 | ABCPRED | 1 | NON-ALLERGEN | Toxin | 0.03 | 0.61 | -0.44 | -1 | 4.38 | 2072.69 | outside |
| 1198 | 1214 | IDLQELGKYEQYIKWP | 0.7389 | 16 | ABCPRED | 1 | NON-ALLERGEN | Non-Toxin | -0.16 | -0.89 | 0.01 | -1 | 4.68 | 2023.57 | outside |
| 1136 | 1154 | TVYDPLQPELDSFKEELD | -0.0374 | 18 | ABCPRED | 1 | NON-ALLERGEN | Non-Toxin | -0.2 | -0.89 | 0.52 | -5 | 3.72 | 2138.57 | outside |
| 1040 | 1058 | VDFCGKGYHLMSFPQSAP | 0.2391 | 18 | ABCPRED | 1 | NON-ALLERGEN | Non-Toxin | -0.02 | -0.07 | -0.39 | 0.5 | 7.06 | 1984.53 | outside |
| 1143 | 1159 | PELDSFKEELDKYFKN | -0.8708 | 16 | ABCPRED | 1 | NON-ALLERGEN | Non-Toxin | -0.33 | -1.45 | 0.85 | -2 | 4.52 | 2002.43 | outside |
| 235 | 255 | ITRFQTLLALHRSYLTPGDS | 0.2654 | 20 | ABCPRED | 1 | ALLERGEN | Non-Toxin | -0.13 | -0.1 | -0.31 | 1.5 | 9.1 | 2289.92 | outside |
| 1200 | 1220 | LQELGKYEQYIKWPWYIWLG | 0.7323 | 20 | ABCPRED,BCPREDS | 2 | NON-ALLERGEN | Non-Toxin | -0.02 | -0.52 | -0.69 | 0 | 6.48 | 2614.37 | outside_transmembrane-region |
| 399 | 409 | SFVIRGDEVR | 0.1752 | 10 | ABCPRED | 1 | NON-ALLERGEN | Non-Toxin | -0.25 | -0.15 | 0.5 | 0 | 6.42 | 1177.46 | outside |
| 242 | 254 | LALHRSYLTPGD | 0.7249 | 12 | ABCPRED | 1 | NON-ALLERGEN | Non-Toxin | -0.11 | -0.23 | -0.23 | 0.5 | 7.09 | 1342.69 | outside |
| 336 | 348 | CPFGEVFNATRF | 0.0965 | 12 | ABCPRED | 1 | ALLERGEN | Non-Toxin | -0.04 | 0.22 | -0.39 | 0 | 6.32 | 1387.73 | outside |
| 464 | 471 | FERDIST | -1.2261 | 7 | IEDB | 1 | ALLERGEN | Non-Toxin | -0.31 | -0.81 | 0.66 | -1 | 4.38 | 867.01 | outside |
| 43 | 61 | FRSSVLHSTQDLFLPFFS | 0.2312 | 18 | ABCPRED | 1 | NON-ALLERGEN | Non-Toxin | -0.02 | 0.37 | -0.58 | 0.5 | 7.1 | 2128.67 | outside |
| 717 | 735 | NFTISVTTEILPVSMTKT | 1.3127 | 18 | ABCPRED | 1 | ALLERGEN | Non-Toxin | 0.01 | 0.46 | -0.41 | 0 | 6.35 | 1982.59 | outside |
| 317 | 333 | NFRVQPTESIVRFPNI | 0.6366 | 16 | ABCPRED | 1 | ALLERGEN | Non-Toxin | -0.18 | -0.29 | -0.13 | 1 | 9.95 | 1917.43 | outside |
| 143 | 157 | VYYHKNNKSWMESE | 0.3303 | 14 | ABCPRED,BCPREDS,FBCPred | 3 | NON-ALLERGEN | Non-Toxin | -0.32 | -1.71 | 0.12 | 0.5 | 7.09 | 1815.19 | outside |
| 1178 | 1198 | NIQKEIDRLNEVAKNLNESL | -0.0099 | 20 | BCEPRED | 1 | ALLERGEN | Non-Toxin | -0.31 | -0.91 | 0.56 | -1 | 4.88 | 2340.93 | outside |
| 313 | 331 | YQTSNFRVQPTESIVRFP | 0.2985 | 18 | ABCPRED | 1 | NON-ALLERGEN | Non-Toxin | -0.23 | -0.67 | -0.15 | 1 | 9.1 | 2169.68 | outside |
| 40 | 58 | DKVFRSSVLHSTQDLFLP | -0.13 | 18 | ABCPRED | 1 | NON-ALLERGEN | Non-Toxin | -0.14 | -0.08 | -0.07 | 0.5 | 7.1 | 2089.64 | outside |
| 829 | 849 | ADAGFIKQYGDCLGDIAARD | 0.4894 | 20 | ABCPRED,FBCPred | 2 | ALLERGEN | Non-Toxin | -0.11 | -0.15 | 0.25 | -2 | 4.14 | 2099.58 | outside |
| 1199 | 1219 | DLQELGKYEQYIKWPWYIWL | 0.6838 | 20 | FBCPred | 1 | ALLERGEN | Non-Toxin | -0.06 | -0.67 | -0.53 | -1 | 4.68 | 2672.4 | outside_transmembrane-region |
| 466 | 478 | RDISTEIYQAGS | -0.0903 | 12 | ABCPRED | 1 | NON-ALLERGEN | Non-Toxin | -0.22 | -0.68 | 0.25 | -1 | 4.38 | 1339.59 | outside |
| 139 | 155 | PFLGVYYHKNNKSWME | 0.5447 | 16 | ABCPRED | 1 | NON-ALLERGEN | Non-Toxin | -0.14 | -0.94 | -0.37 | 1.5 | 8.77 | 2013.54 | outside |
| 844 | 862 | IAARDLICAQKFNGLTVL | 0.5307 | 18 | ABCPRED | 1 | ALLERGEN | Non-Toxin | 0 | 0.85 | -0.36 | 1 | 8.57 | 1946.61 | outside |
| 351 | 364 | YAWNRKRISNCVA | 0.3936 | 13 | literature | 1 | ALLERGEN | Non-Toxin | -0.3 | -0.62 | -0.1 | 3 | 10.07 | 1581 | outside |
| 82 | 102 | PVLPFNDGVYFASTEKSNII | 0.5207 | 20 | ABCPRED | 1 | NON-ALLERGEN | Non-Toxin | 0.01 | 0.17 | -0.33 | -1 | 4.38 | 2211.79 | outside |
| 1206 | 1222 | YEQYIKWPWYIWLGFI | 0.951 | 16 | ABCPRED,literature | 2 | NON-ALLERGEN | Non-Toxin | 0.14 | 0.04 | -1.29 | 0 | 6.34 | 2205.84 | outside_transmembrane-region |
| 257 | 275 | GWTAGAAAYYVGYLQPRT | 0.7921 | 18 | ABCPRED | 1 | NON-ALLERGEN | Non-Toxin | 0.01 | -0.1 | -0.73 | 1 | 8.83 | 1945.42 | outside |
| 355 | 371 | RKRISNCVADYSVLYN | -0.0708 | 16 | ABCPRED | 1 | NON-ALLERGEN | Non-Toxin | -0.28 | -0.41 | 0.02 | 2 | 9.25 | 1901.39 | outside |

| | | | | | | | | | | | | | | | |
|---|---|---|---|---|---|---|---|---|---|---|---|---|---|---|---|
| 842 | 862 | GDIAARDLICAQKFNGLTVL | 0.6162 | 20 | ABCPRED | 1 | NON-ALLERGEN | Non-Toxin | -0.03 | 0.57 | -0.17 | 0 | 6.28 | 2118.78 | outside |
| 271 | 281 | QPRTFLLKYN | -0.28 | 10 | ABCPRED | 1 | ALLERGEN | Non-Toxin | -0.28 | -0.86 | -0.24 | 2 | 10.01 | 1279.65 | outside |
| 573 | 589 | TDAVRDPQTLEILDIT | 0.9832 | 16 | ABCPRED | 1 | NON-ALLERGEN | Non-Toxin | -0.16 | -0.19 | 0.3 | -3 | 3.84 | 1800.23 | outside |
| 972 | 990 | AISSVLNDILSRLDKVEA | -0.0411 | 18 | ABCPRED | 1 | ALLERGEN | Non-Toxin | -0.1 | 0.42 | 0.17 | -1 | 4.56 | 1943.49 | outside |
| 394 | 412 | NVYADSFVIRGDEVRQIA | -0.1912 | 18 | ABCPRED | 1 | NON-ALLERGEN | Non-Toxin | -0.16 | -0.06 | 0.1 | -1 | 4.56 | 2052.53 | outside |
| 397 | 409 | ADSFVIRGDEVR | -0.1593 | 12 | ABCPRED | 1 | ALLERGEN | Non-Toxin | -0.25 | -0.27 | 0.62 | -1 | 4.56 | 1363.65 | outside |
| 1148 | 1164 | FKEELDKYFKNHTSPD | -0.5072 | 16 | ABCPRED | 1 | NON-ALLERGEN | Non-Toxin | -0.36 | -1.71 | 0.72 | -0.5 | 5.5 | 1998.4 | outside |
| 1207 | 1223 | EQYIKWPWYIWLGFIA | 1.0127 | 16 | ABCPRED | 1 | NON-ALLERGEN | Non-Toxin | 0.15 | 0.23 | -1.18 | 0 | 6.35 | 2113.74 | outside_transmembrane-region |
| 438 | 456 | SNNLDSKVGGNYNYLYRL | 0.4318 | 18 | ABCPRED | 1 | ALLERGEN | Non-Toxin | -0.23 | -0.92 | -0.19 | 1 | 8.76 | 2090.55 | outside |
| 316 | 326 | SNFRVQPTES | 1.2078 | 10 | IEDB | 1 | ALLERGEN | Non-Toxin | -0.33 | -1.19 | 0.26 | 0 | 6.36 | 1164.37 | outside |
| 81 | 101 | NPVLPFNDGVYFASTEKSNI | 0.5637 | 20 | ABCPRED | 1 | NON-ALLERGEN | Non-Toxin | -0.06 | -0.23 | -0.23 | -1 | 4.38 | 2212.73 | outside |
| 356 | 372 | KRISNCVADYSVLYNS | -0.1093 | 16 | ABCPRED | 1 | NON-ALLERGEN | Non-Toxin | -0.19 | -0.18 | -0.15 | 1 | 8.51 | 1832.28 | outside |
| 577 | 586 | RDPQTLEIL | 0.7101 | 9 | ellipro,literature | 2 | NON-ALLERGEN | Non-Toxin | -0.25 | -0.58 | 0.38 | -1 | 4.38 | 1084.37 | outside |
| 1204 | 1220 | GKYEQYIKWPWYIWLG | 0.9643 | 16 | ABCPRED | 1 | NON-ALLERGEN | Non-Toxin | -0.01 | -0.69 | -0.83 | 1 | 8.71 | 2130.73 | outside_transmembrane-region |
| 390 | 408 | LCFTNVYADSFVIRGDEV | 0.4869 | 18 | ABCPRED | 1 | ALLERGEN | Non-Toxin | -0.02 | 0.51 | -0.27 | -2 | 4.03 | 2048.55 | outside |
| 841 | 859 | LGDIAARDLICAQKFNGL | 0.7076 | 18 | ABCPRED | 1 | NON-ALLERGEN | Non-Toxin | -0.05 | 0.44 | -0.09 | 0 | 6.28 | 1918.51 | outside |
| 1179 | 1188 | IQKEIDRLN | -0.4173 | 9 | IEDB | 1 | ALLERGEN | Non-Toxin | -0.39 | -1.07 | 0.78 | 0 | 6.42 | 1128.43 | outside |
| 29 | 41 | TNSFTRGVYYPD | 0.0072 | 12 | FBCPred | 1 | NON-ALLERGEN | Non-Toxin | -0.2 | -0.94 | -0.24 | 0 | 6.18 | 1419.67 | outside |
| 450 | 466 | NYLYRLFRKSNLKPFE | 0.1234 | 16 | ABCPRED | 1 | NON-ALLERGEN | Non-Toxin | -0.32 | -0.96 | 0.04 | 3 | 10 | 2088.67 | outside |
| 270 | 280 | LQPRTFLLKY | 0.0494 | 10 | ABCPRED | 1 | ALLERGEN | Non-Toxin | -0.16 | -0.13 | -0.44 | 2 | 10.01 | 1278.71 | outside |
| 333 | 349 | TNLCPFGEVFNATRFA | 0.388 | 16 | ABCPRED | 1 | NON-ALLERGEN | Non-Toxin | -0.03 | 0.26 | -0.45 | 0 | 6.32 | 1787.24 | outside |
| 352 | 358 | AWNRKR | 1.632 | 6 | IEDB,literature | 2 | ALLERGEN | Non-Toxin | -0.77 | -2.58 | 0.88 | 3 | 12.01 | 830.03 | outside |
| 124 | 148 | TNVVIKVCEFQFCNDPFLGVYYHK | 0.072 | 24 | BCEPRED | 1 | NON-ALLERGEN | Non-Toxin | -0.03 | 0.2 | -0.5 | 0.5 | 7.03 | 2864.69 | outside |
| 1138 | 1152 | YDPLQPELDSFKEE | 0.1899 | 14 | AAP | 1 | NON-ALLERGEN | Non-Toxin | -0.27 | -1.41 | 0.72 | -4 | 3.84 | 1710.02 | outside |
| 565 | 581 | FGRDIADTTDAVRDPQ | 0.1853 | 16 | ABCPRED | 1 | NON-ALLERGEN | Non-Toxin | -0.31 | -0.92 | 0.66 | -2 | 4.14 | 1777.09 | outside |
| 183 | 195 | QGNFKNLREFVF | 0.3002 | 12 | ABCPRED | 1 | NON-ALLERGEN | Non-Toxin | -0.2 | -0.53 | -0.1 | 1 | 9.1 | 1498.88 | outside |
| 973 | 993 | ISSVLNDILSRLDKVEAEVQ | 0.083 | 20 | ABCPRED | 1 | NON-ALLERGEN | Non-Toxin | -0.14 | 0.15 | 0.27 | -2 | 4.32 | 2228.83 | outside |
| 85 | 103 | PFNDGVYFASTEKSNIIR | 0.0658 | 18 | ABCPRED | 1 | NON-ALLERGEN | Non-Toxin | -0.14 | -0.41 | -0.02 | 0 | 6.42 | 2058.53 | outside |
| 1206 | 1218 | YEQYIKWPWYIW | 0.6453 | 12 | BCPREDS,FBCPred | 2 | ALLERGEN | Non-Toxin | 0.01 | -0.84 | -1.21 | 0 | 6.34 | 1775.22 | outside_transmembrane-region |
| 560 | 572 | LPFQQFGRDIAD | 0.0048 | 12 | ABCPRED | 1 | NON-ALLERGEN | Non-Toxin | -0.15 | -0.4 | 0.03 | -1 | 4.21 | 1406.73 | outside |
| 74 | 84 | NGTKRFDNPV | -0.1686 | 10 | ABCPRED | 1 | NON-ALLERGEN | Non-Toxin | -0.38 | -1.46 | 0.5 | 1 | 9.1 | 1147.39 | outside |
| 765 | 783 | RALTGIAVEQDKNTQEVF | 0.3708 | 18 | ABCPRED | 1 | NON-ALLERGEN | Non-Toxin | -0.2 | -0.45 | 0.26 | -1 | 4.68 | 2019.51 | outside |
| 311 | 329 | GIYQTSNFRVQPTESIVR | 0.4574 | 18 | ABCPRED | 1 | NON-ALLERGEN | Non-Toxin | -0.21 | -0.51 | -0.11 | 1 | 9.1 | 2095.61 | outside |
| 575 | 587 | AVRDPQTLEILD | 0.8287 | 12 | ABCPRED | 1 | NON-ALLERGEN | Non-Toxin | -0.18 | -0.23 | 0.37 | -2 | 4.03 | 1369.71 | outside |
| 345 | 359 | TRFASVYAWNRKRI | 0.3964 | 14 | ABCPRED,BCPREDS,FBCPred | 3 | NON-ALLERGEN | Non-Toxin | -0.33 | -0.68 | -0.03 | 4 | 11.73 | 1768.24 | outside |
| 901 | 921 | QMAYRFNGIGVTQNVLYENQ | 0.7935 | 20 | ABCPRED,FBCPred | 2 | ALLERGEN | Non-Toxin | -0.14 | -0.49 | -0.44 | 0 | 6.35 | 2345.93 | outside |
| 317 | 335 | NFRVQPTESIVRFPNITN | 0.6761 | 18 | ABCPRED | 1 | NON-ALLERGEN | Non-Toxin | -0.21 | -0.49 | -0.13 | 1 | 9.95 | 2132.67 | outside |
| 1196 | 1214 | SLIDLQELGKYEQYIKWP | 0.7503 | 18 | ABCPRED | 1 | NON-ALLERGEN | Non-Toxin | -0.12 | -0.62 | -0.07 | -1 | 4.68 | 2223.84 | outside |
| 400 | 412 | FVIRGDEVRQIA | 0.1243 | 12 | ABCPRED | 1 | NON-ALLERGEN | Non-Toxin | -0.17 | 0.17 | 0.22 | 0 | 6.42 | 1402.79 | outside |
| 1017 | 1033 | EIRASANLAATKMSEC | 0.5562 | 16 | ABCPRED | 1 | NON-ALLERGEN | Non-Toxin | -0.18 | -0.08 | 0.28 | 0 | 6.46 | 1695.15 | outside |
| 347 | 359 | FASVYAWNRKRI | 0.4842 | 12 | ABCPRED | 1 | NON-ALLERGEN | Non-Toxin | -0.23 | -0.36 | -0.25 | 3 | 11.01 | 1510.92 | outside |
| 244 | 256 | LHRSYLTPGDSS | 0.5772 | 12 | ABCPRED | 1 | NON-ALLERGEN | Non-Toxin | -0.22 | -0.83 | 0.01 | 0.5 | 7.09 | 1332.6 | outside |
| 346 | 362 | RFASVYAWNRKRISNC | 0.3389 | 16 | ABCPRED | 1 | NON-ALLERGEN | Non-Toxin | -0.34 | -0.66 | -0.03 | 4 | 10.92 | 1971.48 | outside |
| 65 | 81 | FHAIHVSGTNGTKRFD | 0.798 | 16 | ABCPRED | 1 | ALLERGEN | Toxin | -0.16 | -0.54 | -0.07 | 2 | 9.11 | 1787.2 | outside |
| 1025 | 1041 | AATKMSECVLGQSKRV | 0.4298 | 16 | ABCPRED | 1 | NON-ALLERGEN | Non-Toxin | -0.21 | -0.11 | 0.27 | 2 | 9.36 | 1708.25 | outside |
| 1076 | 1094 | TTAPAICHDGKAHFPREG | -0.156 | 18 | ABCPRED | 1 | NON-ALLERGEN | Non-Toxin | -0.17 | -0.67 | 0.19 | 1 | 7.22 | 1908.37 | outside |
| 214 | 224 | RDLPQGFSAL | 0.8917 | 10 | ABCPRED | 1 | NON-ALLERGEN | Non-Toxin | -0.14 | -0.21 | -0.01 | 0 | 6.19 | 1103.38 | outside |
| 245 | 257 | HRSYLTPGDSSS | 0.5714 | 12 | ABCPRED | 1 | NON-ALLERGEN | Non-Toxin | -0.29 | -1.22 | 0.18 | 0.5 | 7.09 | 1306.51 | outside |
| 83 | 101 | VLPFNDGVYFASTEKSNI | 0.6202 | 18 | ABCPRED | 1 | NON-ALLERGEN | Non-Toxin | -0.03 | 0.03 | -0.27 | -1 | 4.38 | 2001.48 | outside |
| 565 | 582 | FGRDIADTTDAVRDPQT | 0.0859 | 17 | IEDB | 1 | NON-ALLERGEN | Non-Toxin | -0.3 | -0.91 | 0.6 | -2 | 4.14 | 1878.21 | outside |
| 1140 | 1158 | PLQPELDSFKEELDKYFK | -0.5178 | 18 | ABCPRED | 1 | NON-ALLERGEN | Non-Toxin | -0.27 | -1.17 | 0.66 | -2 | 4.52 | 2226.77 | outside |
| 832 | 850 | GFIKQYGDCLGDIAARDL | 0.2759 | 18 | ABCPRED | 1 | NON-ALLERGEN | Non-Toxin | -0.09 | 0.03 | 0.07 | -1 | 4.43 | 1955.48 | outside |
| 630 | 650 | TPTWRVYSTGSNVFQTRAGC | 0.5926 | 20 | ABCPRED,AAP | 2 | NON-ALLERGEN | Non-Toxin | -0.17 | -0.47 | -0.37 | 2 | 9.55 | 2231.75 | outside |
| 262 | 280 | AAAYYVGYLQPRTFLLKY | 0.3441 | 18 | ABCPRED | 1 | NON-ALLERGEN | Non-Toxin | 0 | 0.22 | -0.79 | 2 | 9.43 | 2137.77 | outside |
| 1091 | 1109 | REGVFVSNGTHWFVTQRN | 0.6133 | 18 | ABCPRED | 1 | NON-ALLERGEN | Non-Toxin | -0.2 | -0.66 | -0.24 | 1.5 | 9.95 | 2134.61 | outside |
| 89 | 105 | GVYFASTEKSNIIRGW | -0.0792 | 16 | ABCPRED | 1 | NON-ALLERGEN | Non-Toxin | -0.08 | -0.18 | -0.28 | 1 | 8.93 | 1828.29 | outside |
| 315 | 327 | TSNFRVQPTESI | 0.9265 | 12 | ABCPRED | 1 | NON-ALLERGEN | Non-Toxin | -0.23 | -0.67 | 0.03 | 0 | 6.36 | 1378.67 | outside |
| 153 | 163 | MESEFRVYSS | 0.2776 | 10 | ABCPRED | 1 | NON-ALLERGEN | Non-Toxin | -0.23 | -0.63 | 0.23 | -1 | 4.54 | 1234.47 | outside |
| 322 | 340 | PTESIVRFPNITNLCPFG | 0.4873 | 18 | FBCPred | 1 | ALLERGEN | Non-Toxin | -0.03 | 0.15 | -0.39 | 0 | 6.32 | 2005.58 | outside |
| 1038 | 1050 | KRVDFCGKGYHL | 1.3047 | 12 | BCPREDS | 1 | NON-ALLERGEN | Non-Toxin | -0.25 | -0.65 | 0.2 | 2.5 | 9.25 | 1422.84 | outside |
| 453 | 465 | YRLFRKSNLKPF | 0.0409 | 12 | ABCPRED | 1 | NON-ALLERGEN | Non-Toxin | -0.37 | -0.9 | 0.13 | 4 | 11.1 | 1569.05 | outside |
| 1023 | 1041 | NLAATKMSECVLGQSKRV | 0.561 | 18 | ABCPRED | 1 | NON-ALLERGEN | Non-Toxin | -0.19 | -0.08 | 0.15 | 2 | 9.36 | 1935.55 | outside |
| 339 | 355 | GEVFNATRFASVYAWN | 0.0053 | 16 | ABCPRED | 1 | ALLERGEN | Toxin | -0.03 | 0.02 | -0.56 | 0 | 6.35 | 1832.22 | outside |
| 315 | 333 | TSNFRVQPTESIVRFPNI | 0.5132 | 18 | ABCPRED | 1 | ALLERGEN | Non-Toxin | -0.19 | -0.34 | -0.12 | 1 | 9.95 | 2105.64 | outside |
| 1082 | 1095 | CHDGKAHFPREGV | -0.4141 | 13 | BCEPRED | 1 | NON-ALLERGEN | Non-Toxin | -0.25 | -0.99 | 0.42 | 1 | 7.22 | 1452.79 | outside |
| 1140 | 1152 | PLQPELDSFKEE | 0.1495 | 12 | FBCPred | 1 | ALLERGEN | Non-Toxin | -0.26 | -1.25 | 0.78 | -3 | 4.01 | 1431.73 | outside |
| 895 | 907 | QIPFAMQMAYRF | 1.1307 | 12 | ABCPRED | 1 | ALLERGEN | Non-Toxin | -0.02 | 0.26 | -0.77 | 1 | 9.1 | 1502.98 | outside |
| 1205 | 1221 | KYEQYIKWPWYIWLGF | 1.1402 | 16 | ABCPRED | 1 | NON-ALLERGEN | Non-Toxin | 0.02 | -0.49 | -0.99 | 1 | 8.71 | 2220.85 | outside_transmembrane-region |
| 348 | 360 | ASVYAWNRKRIS | 0.5771 | 12 | ABCPRED,BCPREDS | 2 | NON-ALLERGEN | Non-Toxin | -0.3 | -0.66 | -0.02 | 3 | 11.01 | 1450.82 | outside |
| 1207 | 1225 | EQYIKWPWYIWLGFIAGL | 0.7646 | 18 | ABCPRED | 1 | NON-ALLERGEN | Non-Toxin | 0.17 | 0.39 | -1.14 | 0 | 6.35 | 2283.99 | outside_transmembrane-region |
| 70 | 82 | VSGTNGTKRFDN | 0.7085 | 12 | ABCPRED | 1 | NON-ALLERGEN | Non-Toxin | -0.33 | -1.24 | 0.41 | 1 | 9.1 | 1295.54 | outside |
| 774 | 794 | QDKNTQEVFAQVKQIYKTPP | 0.3359 | 20 | ABCPRED,FBCPred | 2 | NON-ALLERGEN | Toxin | -0.29 | -1.23 | 0.26 | 1 | 8.76 | 2362.97 | outside |
| 780 | 798 | EVFAQVKQIYKTPPIKDF | 0.1004 | 18 | ABCPRED | 1 | NON-ALLERGEN | Non-Toxin | -0.13 | -0.34 | 0.03 | 1 | 8.76 | 2151.8 | outside |
| 236 | 254 | TRFQTLLALHRSYLTPGD | 0.5595 | 18 | ABCPRED | 1 | ALLERGEN | Non-Toxin | -0.17 | -0.31 | -0.26 | 1.5 | 9.1 | 2089.65 | outside |
| 333 | 351 | TNLCPFGEVFNATRFASV | 0.4365 | 18 | ABCPRED | 1 | NON-ALLERGEN | Non-Toxin | -0.01 | 0.42 | -0.47 | 0 | 6.32 | 1973.48 | outside |
| 1252 | 1268 | SCCKFDEDDSEPVLKG | 0.4347 | 16 | ABCPRED,IEDB,literature | 3 | NON-ALLERGEN | Non-Toxin | -0.27 | -0.82 | 0.86 | -3 | 4.11 | 1772.14 | inside |
| 290 | 310 | DCALDPLSETKCTLKSFTVE | 0.6362 | 20 | ABCPRED | 1 | NON-ALLERGEN | Non-Toxin | -0.15 | -0.1 | 0.28 | -2 | 4.32 | 2200.78 | outside |

| | | | | | | | | | | | | | | | |
|---|---|---|---|---|---|---|---|---|---|---|---|---|---|---|---|
| 1135 | 1155 | NTVYDPLQPELDSFKEELDK | -0.3936 | 20 | ABCPRED,BCPREDS,FBCPred | 3 | NON-ALLERGEN | Non-Toxin | -0.27 | -1.17 | 0.63 | -4 | 4.02 | 2380.88 | outside |
| 81 | 99 | NPVLPFNDGVYFASTEKS | 0.5539 | 18 | ABCPRED | 1 | NON-ALLERGEN | Non-Toxin | -0.07 | -0.31 | -0.17 | -1 | 4.38 | 1985.43 | outside |
| 346 | 364 | RFASVYAWNRKRISNCVA | 0.3101 | 18 | ABCPRED,BCPREDS | 2 | NON-ALLERGEN | Non-Toxin | -0.25 | -0.26 | -0.14 | 4 | 10.92 | 2141.72 | outside |
| 337 | 349 | PFGEVFNATRFA | 0.0745 | 12 | ABCPRED | 1 | ALLERGEN | Non-Toxin | -0.02 | 0.17 | -0.35 | 0 | 6.36 | 1355.67 | outside |
| 351 | 362 | YAWNRKRISNC | 0.4749 | 11 | BCEPRED | 1 | NON-ALLERGEN | Non-Toxin | -0.43 | -1.28 | 0.06 | 3 | 10.07 | 1410.76 | outside |
| 227 | 245 | VDLPIGINITRFQTLLAL | 0.8733 | 18 | ABCPRED | 1 | NON-ALLERGEN | Non-Toxin | 0.09 | 1.06 | -0.64 | 0 | 6.19 | 1997.7 | outside |
| 1033 | 1051 | VLGQSKRVDFCGKGYHLM | 1.0561 | 18 | ABCPRED | 1 | NON-ALLERGEN | Non-Toxin | -0.14 | -0.14 | -0.09 | 2.5 | 9.25 | 2038.69 | outside |
| 1134 | 1164 | NNTVYDPLQPELDSFKEELDKYFKNHTSPD | -0.2097 | 30 | BCEPRED | 1 | NON-ALLERGEN | Non-Toxin | -0.29 | -1.42 | 0.45 | -3.5 | 4.38 | 3585.29 | outside |
| 100 | 114 | IIRGWIFGTTLDSK | 0.0158 | 14 | ABCPRED,AAP | 2 | NON-ALLERGEN | Non-Toxin | -0.01 | 0.31 | -0.33 | 1 | 9.1 | 1607.1 | outside |
| 631 | 649 | PTWRVYSTGSNVFQTRAG | 0.562 | 18 | ABCPRED | 1 | ALLERGEN | Non-Toxin | -0.18 | -0.63 | -0.33 | 2 | 10.84 | 2027.48 | outside |
| 1137 | 1155 | VYDPLQPELDSFKEELDK | -0.5091 | 18 | BCPREDS,FBCPred | 2 | NON-ALLERGEN | Non-Toxin | -0.25 | -1.07 | 0.71 | -4 | 4.02 | 2165.64 | outside |
| 305 | 321 | SFTVEKGIYQTSNFRV | 0.3631 | 16 | ABCPRED | 1 | ALLERGEN | Non-Toxin | -0.16 | -0.32 | -0.18 | 1 | 8.93 | 1876.33 | outside |
| 97 | 107 | KSNIIRGWIF | -1.2675 | 10 | ABCPRED | 1 | ALLERGEN | Non-Toxin | -0.04 | 0.23 | -0.48 | 2 | 11.01 | 1233.63 | outside |
| 177 | 190 | MDLEGKQGNFKNL | 1.2592 | 13 | IEDB,literature | 2 | NON-ALLERGEN | Non-Toxin | -0.25 | -1.06 | 0.4 | 0 | 6.42 | 1493.9 | outside |
| 328 | 344 | RFPNITNLCPFGEVFN | 0.6318 | 16 | ABCPRED | 1 | ALLERGEN | Non-Toxin | -0.05 | 0.04 | -0.46 | 0 | 6.32 | 1868.37 | outside |
| 440 | 458 | NLDSKVGGNYNYLYRLFR | 0.1518 | 18 | ABCPRED | 1 | NON-ALLERGEN | Non-Toxin | -0.24 | -0.78 | -0.19 | 2 | 9.55 | 2192.73 | outside |
| 982 | 990 | SRLDKVEA | 0.1615 | 8 | BCEPRED | 1 | NON-ALLERGEN | Non-Toxin | -0.39 | -0.8 | 1.06 | 0 | 6.42 | 917.13 | outside |
| 326 | 344 | IVRFPNITNLCPFGEVFN | 0.4325 | 18 | ABCPRED | 1 | ALLERGEN | Non-Toxin | 0.03 | 0.52 | -0.59 | 0 | 6.32 | 2080.7 | outside |
| 891 | 908 | GAALQIPFAMQMAYRFN | 0.8583 | 17 | COVIDep,literature | 2 | ALLERGEN | Non-Toxin | 0.02 | 0.39 | -0.7 | 1 | 9.1 | 1929.53 | outside |
| 136 | 154 | CNDPFLGVYYHKNNKSWM | 0.531 | 18 | ABCPRED | 1 | NON-ALLERGEN | Non-Toxin | -0.17 | -0.89 | -0.37 | 1.5 | 8.48 | 2216.78 | outside |
| 1205 | 1223 | KYEQYIKWPWYIWLGFIA | 1.0289 | 18 | ABCPRED | 1 | NON-ALLERGEN | Non-Toxin | 0.07 | -0.08 | -1.01 | 1 | 8.71 | 2405.12 | outside_transmembrane-region |
| 181 | 193 | GKQGNFKNLREF | 0.5402 | 12 | BCPREDS | 1 | NON-ALLERGEN | Non-Toxin | -0.37 | -1.47 | 0.48 | 2 | 10.01 | 1437.8 | outside |
| 66 | 82 | HAIHVSGTNGTKRFDN | 0.5909 | 16 | ABCPRED | 1 | NON-ALLERGEN | Non-Toxin | -0.24 | -0.94 | 0.1 | 2 | 9.11 | 1754.13 | outside |
| 447 | 465 | GNYNYLYRLFRKSNLKPF | 0.197 | 18 | ABCPRED | 1 | NON-ALLERGEN | Non-Toxin | -0.27 | -0.94 | -0.24 | 4 | 10.17 | 2293.92 | outside |
| 131 | 149 | CEFQFCNDPFLGVYYHKN | 0.3057 | 18 | ABCPRED | 1 | NON-ALLERGEN | Non-Toxin | -0.09 | -0.43 | -0.46 | -0.5 | 5.33 | 2224.75 | outside |
| 351 | 361 | YAWNRKRISN | 0.5855 | 10 | BCEPRED | 1 | ALLERGEN | Non-Toxin | -0.48 | -1.66 | 0.17 | 3 | 11.01 | 1307.61 | outside |
| 398 | 410 | DSFVIRGDEVRQ | -0.0659 | 12 | ABCPRED,AAP | 2 | ALLERGEN | Non-Toxin | -0.33 | -0.71 | 0.68 | -1 | 4.56 | 1420.71 | outside |
| 1255 | 1266 | KFDEDDSEPVL | 0.29 | 11 | literature | 1 | NON-ALLERGEN | Non-Toxin | -0.29 | -1.18 | 1.14 | -4 | 3.77 | 1293.49 | inside |
| 1254 | 1270 | CKFDEDDSEPVLKGVK | 0.5198 | 16 | ABCPRED | 1 | NON-ALLERGEN | Non-Toxin | -0.29 | -0.91 | 1 | -2 | 4.44 | 1809.24 | inside |
| 1015 | 1031 | AAEIRASANLAATKMS | 0.8103 | 16 | ABCPRED | 1 | ALLERGEN | Non-Toxin | -0.11 | 0.21 | 0.09 | 1 | 9.1 | 1605.05 | outside |
| 1080 | 1096 | AICHDGKAHFPREGVF | -0.2375 | 16 | ABCPRED | 1 | NON-ALLERGEN | Non-Toxin | -0.11 | -0.24 | 0.04 | 1 | 7.22 | 1784.25 | outside |
| 1141 | 1159 | LQPELDSFKEELDKYFKN | -0.6174 | 18 | ABCPRED | 1 | NON-ALLERGEN | Non-Toxin | -0.3 | -1.27 | 0.67 | -2 | 4.52 | 2243.76 | outside |
| 1069 | 1085 | PAQEKNFTTAPAICHD | 0.3653 | 16 | ABCPRED | 1 | NON-ALLERGEN | Non-Toxin | -0.16 | -0.66 | 0.08 | -0.5 | 5.33 | 1743.14 | outside |
| 206 | 216 | KHTPINLVRD | 0.6867 | 10 | ABCPRED | 1 | ALLERGEN | Non-Toxin | -0.31 | -0.84 | 0.32 | 1.5 | 9.1 | 1192.53 | outside |
| 568 | 586 | DIADTTDAVRDPQTLEIL | 0.6732 | 18 | ABCPRED | 1 | ALLERGEN | Non-Toxin | -0.17 | -0.27 | 0.41 | -4 | 3.72 | 1986.42 | outside |
| 398 | 408 | DSFVIRGDEV | 0.3245 | 10 | ABCPRED | 1 | NON-ALLERGEN | Non-Toxin | -0.15 | -0.05 | 0.5 | -2 | 4.03 | 1136.36 | outside |
| 1254 | 1274 | CKFDEDDSEPVLKGVKLHYT | 0.8247 | 20 | COVIDep | 1 | NON-ALLERGEN | Non-Toxin | -0.23 | -0.8 | 0.55 | -1.5 | 4.83 | 2323.89 | inside |
| 1133 | 1173 | NTVYDPLQPELDSFKEELDKYFKNHTSPDVDLGD | 0.1613 | 40 | IEDB,literature | 2 | NON-ALLERGEN | Non-Toxin | -0.18 | -0.75 | 0.29 | -5.5 | 4.13 | 4554.56 | outside |
| 1014 | 1032 | RAAEIRASANLAATKMSE | 0.5976 | 18 | ABCPRED | 1 | NON-ALLERGEN | Non-Toxin | -0.23 | -0.26 | 0.42 | 1 | 9.1 | 1890.38 | outside |
| 338 | 356 | FGEVFNATRFASVYAWNR | -0.0164 | 18 | ABCPRED | 1 | ALLERGEN | Non-Toxin | -0.09 | -0.08 | -0.47 | 1 | 9.1 | 2135.61 | outside |
| 423 | 439 | YKLPDDFTGCVIAWNS | -0.2253 | 16 | ABCPRED | 1 | ALLERGEN | Non-Toxin | -0.03 | -0.03 | -0.36 | -1 | 4.21 | 1829.28 | outside |
| 564 | 583 | QFGRDIADTTDAVRDPQTL | 0.1344 | 19 | IEDB | 1 | NON-ALLERGEN | Non-Toxin | -0.28 | -0.8 | 0.45 | -2 | 4.14 | 2119.54 | outside |
| 1142 | 1160 | QPELDSFKEELDKYFKNH | -0.7418 | 18 | ABCPRED | 1 | NON-ALLERGEN | Non-Toxin | -0.35 | -1.66 | 0.74 | -1.5 | 4.9 | 2267.74 | outside |
| 331 | 349 | NITNLCPFGEVFNATRFA | 0.611 | 18 | ABCPRED | 1 | NON-ALLERGEN | Toxin | -0.02 | 0.28 | -0.49 | 0 | 6.32 | 2014.54 | outside |
| 240 | 256 | TLLALHRSYLTPGDSS | 0.7029 | 16 | ABCPRED | 1 | NON-ALLERGEN | Non-Toxin | -0.1 | -0.08 | -0.28 | 0.5 | 7.09 | 1731.17 | outside |
| 309 | 321 | EKGIYQTSNFRV | 0.3081 | 12 | FBCPred | 1 | NON-ALLERGEN | Non-Toxin | -0.27 | -0.88 | 0.1 | 1 | 8.93 | 1441.78 | outside |
| 1181 | 1199 | KEIDRLNEVAKNLNESLI | 0.2064 | 18 | ABCPRED | 1 | ALLERGEN | Non-Toxin | -0.27 | -0.62 | 0.61 | -1 | 4.88 | 2098.66 | outside |
| 129 | 147 | KVCEFQFCNDPFLGVYYH | -0.0382 | 18 | ABCPRED | 1 | NON-ALLERGEN | Non-Toxin | -0.03 | -0.01 | -0.56 | -0.5 | 5.33 | 2209.78 | outside |
| 779 | 799 | QEVFAQVKQIYKTPPIKDFG | 0.1047 | 20 | ABCPRED | 1 | NON-ALLERGEN | Non-Toxin | -0.14 | -0.5 | 0.04 | 1 | 8.76 | 2337.02 | outside |
| 631 | 643 | PTWRVYSTGSNV | 0.5085 | 12 | ABCPRED | 1 | NON-ALLERGEN | Non-Toxin | -0.14 | -0.57 | -0.48 | 1 | 9.1 | 1366.66 | outside |
| 454 | 466 | RLFRKSNLKPFE | 0.1904 | 12 | ABCPRED | 1 | ALLERGEN | Non-Toxin | -0.42 | -1.08 | 0.58 | 3 | 11.01 | 1534.99 | outside |
| 631 | 647 | PTWRVYSTGSNVFQTR | 0.4819 | 16 | ABCPRED | 1 | ALLERGEN | Non-Toxin | -0.23 | -0.79 | -0.34 | 2 | 10.84 | 1899.32 | outside |
| 807 | 819 | PDPSKPSKRSFI | 0.4605 | 12 | BCPREDS | 1 | NON-ALLERGEN | Non-Toxin | -0.36 | -1.31 | 0.72 | 2 | 10.01 | 1358.71 | outside |
| 272 | 282 | PRTFLLKYNE | -0.4103 | 10 | ABCPRED | 1 | ALLERGEN | Non-Toxin | -0.27 | -0.86 | 0.04 | 1 | 8.93 | 1280.63 | outside |
| 833 | 849 | FIKQYGDCLGDIAARD | 0.2183 | 16 | ABCPRED | 1 | NON-ALLERGEN | Non-Toxin | -0.14 | -0.18 | 0.19 | -1 | 4.43 | 1785.23 | outside |
| 1253 | 1269 | CCKFDEDDSEPVLKGV | 0.5963 | 16 | ABCPRED | 1 | NON-ALLERGEN | Toxin | -0.22 | -0.51 | 0.75 | -3 | 4.11 | 1784.2 | inside |
| 1078 | 1094 | APAICHDGKAHFPREG | -0.3045 | 16 | ABCPRED | 1 | ALLERGEN | Non-Toxin | -0.17 | -0.66 | 0.26 | 1 | 7.22 | 1706.13 | outside |
| 630 | 640 | TPTWRVYSTG | 0.5628 | 10 | ABCPRED | 1 | ALLERGEN | Non-Toxin | -0.15 | -0.74 | -0.51 | 1 | 9.1 | 1167.42 | outside |
| 1251 | 1269 | GSCCKFDEDDSEPVLKGV | 0.4944 | 18 | ABCPRED | 1 | NON-ALLERGEN | Toxin | -0.2 | -0.52 | 0.68 | -3 | 4.11 | 1928.36 | inside |
| 242 | 258 | LALHRSYLTPGDSSSG | 0.5369 | 16 | ABCPRED | 1 | NON-ALLERGEN | Non-Toxin | -0.12 | -0.35 | -0.12 | 0.5 | 7.09 | 1661.03 | outside |
| 349 | 359 | SVYAWNRKRI | 0.5509 | 10 | ABCPRED | 1 | ALLERGEN | Non-Toxin | -0.36 | -0.89 | 0 | 3 | 11.01 | 1292.64 | outside |
| 727 | 745 | LPVSMTKTSVDCTMYICG | 0.5623 | 18 | ABCPRED | 1 | NON-ALLERGEN | Non-Toxin | 0.01 | 0.62 | -0.45 | 0 | 6.12 | 1949.6 | outside |
| 976 | 994 | VLNDILSRLDKVEAEVQI | 0.0567 | 18 | ABCPRED | 1 | NON-ALLERGEN | Non-Toxin | -0.12 | 0.26 | 0.26 | -2 | 4.32 | 2054.65 | outside |
| 1019 | 1036 | RASANLAATKMSECVLG | 0.4414 | 17 | COVIDep,literature | 2 | NON-ALLERGEN | Non-Toxin | -0.1 | 0.31 | 0 | 1 | 8.57 | 1722.24 | outside |
| 777 | 797 | NTQEVFAQVKQIYKTPPIKD | 0.3305 | 20 | ABCPRED,AAP | 2 | ALLERGEN | Non-Toxin | -0.22 | -0.83 | 0.16 | 1 | 8.76 | 2348 | outside |
| 981 | 995 | LSRLDKVEAEVQID | 0.4233 | 14 | ABCPRED,AAP,FBCPred | 3 | NON-ALLERGEN | Non-Toxin | -0.24 | -0.31 | 0.69 | -2 | 4.32 | 1615.02 | outside |
| 1200 | 1218 | LQELGKYEQYIKWPWYIW | 0.5918 | 18 | ABCPRED,BCPREDS,FBCPred | 3 | NON-ALLERGEN | Non-Toxin | -0.06 | -0.77 | -0.66 | 0 | 6.48 | 2444.12 | outside_transmembrane-region |
| 571 | 585 | DTTDAVRDPQTLEI | 0.5633 | 14 | ABCPRED,AAP | 2 | NON-ALLERGEN | Non-Toxin | -0.27 | -0.81 | 0.6 | -3 | 3.84 | 1573.87 | outside |
| 1248 | 1266 | CSCGSCCKFDEDDSEPVL | 0.172 | 18 | ABCPRED | 1 | NON-ALLERGEN | Non-Toxin | -0.19 | -0.28 | 0.51 | -4 | 3.77 | 1937.34 | inside |
| 327 | 344 | VRFPNITNLCPFGEVFN | 0.6787 | 17 | COVIDep | 1 | NON-ALLERGEN | Non-Toxin | -0.01 | 0.28 | -0.52 | 0 | 6.32 | 1967.52 | outside |
| 969 | 989 | NFGAISSVLNDILSRLDKVE | 0.2922 | 20 | ABCPRED | 1 | NON-ALLERGEN | Non-Toxin | -0.09 | 0.24 | 0.07 | -1 | 4.56 | 2190.78 | outside |
| 403 | 419 | RGDEVRQIAPGQTGKI | 0.7748 | 16 | ABCPRED | 1 | ALLERGEN | Non-Toxin | -0.3 | -0.96 | 0.59 | 1 | 9.1 | 1725.18 | outside |
| 341 | 359 | VFNATRFASVYAWNRKRI | 0.3322 | 18 | ABCPRED | 1 | ALLERGEN | Non-Toxin | -0.22 | -0.23 | -0.26 | 4 | 11.73 | 2199.79 | outside |
| 778 | 798 | TQEVFAQVKQIYKTPPIKDF | 0.1649 | 20 | ABCPRED | 1 | ALLERGEN | Toxin | -0.16 | -0.52 | 0.02 | 1 | 8.76 | 2381.07 | outside |
| 627 | 645 | DQLTPTWRVYSTGSNVFQ | 0.386 | 18 | ABCPRED | 1 | ALLERGEN | Non-Toxin | -0.16 | -0.63 | -0.39 | 0 | 6.19 | 2099.55 | outside |
| 126 | 146 | VVIKVCEFQFCNDPFLGVYY | -0.063 | 20 | ABCPRED | 1 | NON-ALLERGEN | Non-Toxin | 0.08 | 0.8 | -0.72 | -1 | 4.38 | 2384.1 | outside |
| 977 | 993 | LNDILSRLDKVEAEVQ | 0.1686 | 16 | ABCPRED | 1 | NON-ALLERGEN | Non-Toxin | -0.22 | -0.26 | 0.5 | -2 | 4.32 | 1842.32 | outside |

| | | | | | | | | | | | | | | | |
|---|---|---|---|---|---|---|---|---|---|---|---|---|---|---|---|
| 1034 | 1050 | LGQSKRVDFCGKGYHL | 1.2157 | 16 | ABCPRED | 1 | ALLERGEN | Non-Toxin | -0.2 | -0.54 | 0.07 | 2.5 | 9.25 | 1808.33 | outside |
| 1072 | 1090 | EKNFTTAPAICHDGKAHF | 0.323 | 18 | ABCPRED | 1 | ALLERGEN | Non-Toxin | -0.14 | -0.56 | 0.06 | 1 | 7.22 | 1987.47 | outside |
| 1142 | 1156 | QPELDSFKEELDKY | -0.9397 | 14 | BCPREDS | 1 | NON-ALLERGEN | Non-Toxin | -0.35 | -1.58 | 0.94 | -3 | 4.18 | 1741.08 | outside |
| 421 | 441 | YNYKLPDDFTGCVIAWNSNN | 0.1744 | 20 | ABCPRED,AAP | 2 | ALLERGEN | Non-Toxin | -0.12 | -0.61 | -0.37 | -1 | 4.21 | 2334.83 | outside |
| 206 | 218 | KHTPINLVRDLP | 0.4683 | 12 | ABCPRED | 1 | ALLERGEN | Non-Toxin | -0.22 | -0.52 | 0.12 | 1.5 | 9.1 | 1402.84 | outside |
| 356 | 368 | KRISNCVADYSV | -0.0398 | 12 | ABCPRED | 1 | ALLERGEN | Non-Toxin | -0.22 | -0.09 | 0.1 | 1 | 8.53 | 1354.7 | outside |
| 389 | 409 | DLCFTNVYADSFVIRGDEVR | 0.4322 | 20 | ABCPRED,FBCPred | 2 | NON-ALLERGEN | Non-Toxin | -0.14 | 0.06 | 0.06 | -2 | 4.23 | 2319.85 | outside |
| 42 | 62 | VFRSSVLHSTQDLFLPFFSN | -0.0083 | 20 | ABCPRED | 1 | NON-ALLERGEN | Non-Toxin | -0.02 | 0.36 | -0.58 | 0.5 | 7.1 | 2341.94 | outside |
| 210 | 222 | INLVRDLPQGFS | 0.3874 | 12 | ABCPRED | 1 | NON-ALLERGEN | Non-Toxin | -0.09 | 0.11 | -0.23 | 0 | 6.19 | 1358.74 | outside |
| 246 | 258 | RSYLTPGDSSSG | 0.5374 | 12 | ABCPRED | 1 | NON-ALLERGEN | Non-Toxin | -0.24 | -0.98 | 0.22 | 0 | 6.19 | 1226.42 | outside |
| 394 | 410 | NVYADSFVIRGDEVRQ | -0.2502 | 16 | ABCPRED | 1 | ALLERGEN | Non-Toxin | -0.24 | -0.46 | 0.26 | -1 | 4.56 | 1868.26 | outside |
| 91 | 105 | YFASTEKSNIIRGW | -0.2178 | 14 | ABCPRED,BCPREDS | 2 | NON-ALLERGEN | Non-Toxin | -0.14 | -0.48 | -0.21 | 1 | 8.93 | 1672.07 | outside |
| 1149 | 1167 | KEELDKYFKNHTSPDVDL | -0.1276 | 18 | ABCPRED | 1 | NON-ALLERGEN | Non-Toxin | -0.33 | -1.43 | 0.76 | -1.5 | 4.83 | 2178.64 | outside |
| 351 | 359 | YAWNRKRI | 0.7206 | 8 | BCEPRED | 1 | ALLERGEN | Non-Toxin | -0.49 | -1.54 | 0.15 | 3 | 11.01 | 1106.4 | outside |
| 1019 | 1035 | RASANLAATKMSECVL | 0.3735 | 16 | ABCPRED | 1 | NON-ALLERGEN | Non-Toxin | -0.12 | 0.36 | 0 | 1 | 8.57 | 1665.17 | outside |
| 841 | 857 | LGDIAARDLICAQKFN | 0.9412 | 16 | ABCPRED | 1 | ALLERGEN | Non-Toxin | -0.1 | 0.28 | 0.01 | 0 | 6.28 | 1748.26 | outside |
| 67 | 83 | AIHVSGTNGTKRFDNP | 0.5163 | 16 | ABCPRED | 1 | NON-ALLERGEN | Non-Toxin | -0.22 | -0.84 | 0.13 | 1.5 | 9.1 | 1714.1 | outside |
| 1032 | 1048 | CVLGQSKRVDFCGKGY | 1.0648 | 16 | ABCPRED | 1 | ALLERGEN | Non-Toxin | -0.18 | -0.16 | 0.06 | 2 | 8.94 | 1760.29 | outside |
| 316 | 332 | SNFRVQPTESIVRFPN | 0.3243 | 16 | ABCPRED | 1 | NON-ALLERGEN | Non-Toxin | -0.25 | -0.63 | 0 | 1 | 9.95 | 1891.34 | outside |
| 316 | 328 | SNFRVQPTESIV | 0.8573 | 12 | ABCPRED | 1 | NON-ALLERGEN | Non-Toxin | -0.17 | -0.27 | -0.06 | 0 | 6.36 | 1376.7 | outside |
| 30 | 42 | NSFTRGVYYPDK | 0.1289 | 12 | ABCPRED | 1 | NON-ALLERGEN | Non-Toxin | -0.28 | -1.21 | 0.04 | 1 | 8.83 | 1446.74 | outside |
| 128 | 144 | IKVCEFQFCNDPFLGV | -0.1003 | 16 | ABCPRED | 1 | NON-ALLERGEN | Non-Toxin | 0.04 | 0.64 | -0.42 | -1 | 4.38 | 1859.42 | outside |
| 26 | 42 | PAYTNSFTRGVYYPDK | 0.0619 | 16 | ABCPRED | 1 | ALLERGEN | Non-Toxin | -0.21 | -1.02 | -0.17 | 1 | 8.76 | 1879.27 | outside |
| 728 | 746 | PVSMTKTSVDCTMYICGD | 0.4071 | 18 | ABCPRED | 1 | ALLERGEN | Non-Toxin | -0.06 | 0.21 | -0.18 | -1 | 4.21 | 1951.52 | outside |
| 423 | 441 | YKLPDDFTGCVIAWNSNN | -0.0911 | 18 | ABCPRED | 1 | NON-ALLERGEN | Non-Toxin | -0.1 | -0.42 | -0.29 | -1 | 4.21 | 2057.52 | outside |
| 342 | 360 | FNATRFASVYAWNRKRIS | 0.4806 | 18 | FBCPred | 1 | NON-ALLERGEN | Non-Toxin | -0.26 | -0.51 | -0.16 | 4 | 11.73 | 2187.73 | outside |
| 388 | 408 | NDLCFTNVYADSFVIRGDEV | 0.6104 | 20 | ABCPRED | 1 | ALLERGEN | Non-Toxin | -0.08 | 0.11 | -0.08 | -3 | 3.84 | 2277.77 | outside |
| 846 | 862 | ARDLICAQKFNGLTVL | 0.341 | 16 | ABCPRED | 1 | NON-ALLERGEN | Non-Toxin | -0.06 | 0.56 | -0.26 | 1 | 8.57 | 1762.34 | outside |
| 181 | 197 | GKQGNFKNLREFVFKN | 0.368 | 16 | ABCPRED | 1 | ALLERGEN | Toxin | -0.32 | -1.13 | 0.31 | 3 | 10.3 | 1926.45 | outside |
| 1141 | 1155 | LQPELDSFKEELDK | -0.7047 | 14 | FBCPred,IEDB | 2 | NON-ALLERGEN | Toxin | -0.31 | -1.21 | 0.97 | -3 | 4.18 | 1691.07 | outside |
| 138 | 155 | DPFLGVYYHKNNKSWME | 0.5821 | 17 | IEDB,literature | 2 | NON-ALLERGEN | Non-Toxin | -0.18 | -1.09 | -0.17 | 0.5 | 7.09 | 2128.64 | outside |
| 335 | 353 | LCPFGEVFNATRFASVYA | 0.179 | 18 | ABCPRED | 1 | NON-ALLERGEN | Non-Toxin | 0.05 | 0.68 | -0.61 | 0 | 6.32 | 1992.52 | outside |
| 628 | 646 | QLTPTWRVYSTGSNVFQT | 0.6238 | 18 | ABCPRED | 1 | ALLERGEN | Non-Toxin | -0.13 | -0.48 | -0.58 | 1 | 9.1 | 2085.57 | outside |
| 43 | 59 | FRSSVLHSTQDLFLPF | 0.4298 | 16 | ABCPRED | 1 | ALLERGEN | Non-Toxin | -0.04 | 0.29 | -0.51 | 0.5 | 7.1 | 1894.39 | outside |
| 563 | 583 | QQFGRDIADTTDAVRDPQTL | 0.0914 | 20 | ABCPRED,literature | 2 | NON-ALLERGEN | Non-Toxin | -0.3 | -0.93 | 0.44 | -2 | 4.14 | 2247.69 | outside |
| 393 | 413 | TNVYADSFVIRGDEVRQIAP | -0.0186 | 20 | ABCPRED | 1 | NON-ALLERGEN | Non-Toxin | -0.15 | -0.16 | 0.07 | -1 | 4.56 | 2250.78 | outside |
| 728 | 744 | PVSMTKTSVDCTMYIC | 0.886 | 16 | ABCPRED | 1 | ALLERGEN | Non-Toxin | -0.03 | 0.48 | -0.39 | 0 | 6.12 | 1779.35 | outside |
| 561 | 573 | PFQQFGRDIADT | -0.175 | 12 | ABCPRED | 1 | NON-ALLERGEN | Non-Toxin | -0.21 | -0.78 | 0.14 | -1 | 4.21 | 1394.67 | outside |
| 1033 | 1047 | VLGQSKRVDFCGKG | 1.3582 | 14 | ABCPRED,IEDB | 2 | NON-ALLERGEN | Non-Toxin | -0.21 | -0.27 | 0.3 | 2 | 9.36 | 1493.95 | outside |
| 500 | 518 | TNGVGYQPYRVVVLSFEL | 0.8694 | 18 | ABCPRED | 1 | NON-ALLERGEN | Non-Toxin | 0 | 0.32 | -0.58 | 0 | 6.35 | 2041.61 | outside |
| 27 | 41 | AYTNSFTRGVYYPD | -0.0257 | 14 | FBCPred | 1 | NON-ALLERGEN | Non-Toxin | -0.16 | -0.77 | -0.41 | 0 | 6.18 | 1653.95 | outside |
| 760 | 780 | CTQLNRALTGIAVEQDKNTQ | 0.6396 | 20 | ABCPRED | 1 | NON-ALLERGEN | Non-Toxin | -0.25 | -0.65 | 0.15 | 0 | 6.39 | 2203.76 | outside |
| 212 | 222 | LVRDLPQGFS | 0.29 | 10 | ABCPRED | 1 | NON-ALLERGEN | Non-Toxin | -0.11 | 0.03 | -0.11 | 0 | 6.19 | 1131.44 | outside |
| 758 | 776 | SFCTQLNRALTGIAVEQD | 0.548 | 18 | ABCPRED | 1 | ALLERGEN | Non-Toxin | -0.12 | 0.03 | -0.13 | -1 | 4.38 | 1966.46 | outside |
| 391 | 409 | CFTNVYADSFVIRGDEVR | 0.0008 | 18 | ABCPRED,FBCPred | 2 | ALLERGEN | Non-Toxin | -0.15 | 0.04 | 0 | -1 | 4.56 | 2091.57 | outside |
| 1087 | 1103 | AHFPREGVFVSNGTHW | 0.4732 | 16 | ABCPRED | 1 | NON-ALLERGEN | Non-Toxin | -0.07 | -0.43 | -0.42 | 1 | 7.26 | 1841.25 | outside |
| 1250 | 1266 | CGSCCKFDEDDSEPVL | 0.3941 | 16 | ABCPRED | 1 | NON-ALLERGEN | Toxin | -0.2 | -0.42 | 0.61 | -4 | 3.77 | 1747.1 | inside |
| 345 | 361 | TRFASVYAWNRKRISN | 0.3968 | 16 | ABCPRED | 1 | NON-ALLERGEN | Non-Toxin | -0.35 | -0.86 | 0.01 | 4 | 11.73 | 1969.45 | outside |
| 422 | 438 | NYKLPDDFTGCVIAWN | 0.0557 | 16 | ABCPRED | 1 | NON-ALLERGEN | Non-Toxin | -0.05 | -0.2 | -0.36 | -1 | 4.21 | 1856.31 | outside |
| 732 | 750 | TKTSVDCTMYICGDSTEC | 0.1577 | 18 | ABCPRED | 1 | NON-ALLERGEN | Non-Toxin | -0.14 | -0.13 | 0.06 | -2 | 4.03 | 1957.43 | outside |
| 832 | 852 | GFIKQYGDCLGDIAARDLIC | 0.2316 | 20 | ABCPRED | 1 | NON-ALLERGEN | Non-Toxin | -0.04 | 0.38 | -0.08 | -1 | 4.43 | 2171.81 | outside |
| 564 | 584 | QFGRDIADTTDAVRDPQTLE | 0.2315 | 20 | ABCPRED,FBCPred | 2 | NON-ALLERGEN | Non-Toxin | -0.3 | -0.93 | 0.58 | -3 | 4.04 | 2248.67 | outside |
| 343 | 363 | NATRFASVYAWNRKRISNCV | 0.235 | 20 | FBCPred | 1 | NON-ALLERGEN | Non-Toxin | -0.27 | -0.44 | -0.13 | 4 | 10.92 | 2356.96 | outside |
| 455 | 465 | LFRKSNLKPF | 0.379 | 10 | ABCPRED | 1 | NON-ALLERGEN | Non-Toxin | -0.27 | -0.5 | 0.09 | 3 | 11.17 | 1249.66 | outside |
| 1251 | 1267 | GSCCKFDEDDSEPVLK | 0.2562 | 16 | ABCPRED | 1 | NON-ALLERGEN | Non-Toxin | -0.27 | -0.82 | 0.86 | -3 | 4.11 | 1772.14 | inside |
| 981 | 993 | LSRLDKVEAEVQ | 0.4604 | 12 | FBCPred | 1 | NON-ALLERGEN | Non-Toxin | -0.28 | -0.45 | 0.7 | -1 | 4.68 | 1386.74 | outside |
| 566 | 586 | GRDIADTTDAVRDPQTLEIL | 0.5014 | 20 | ABCPRED | 1 | NON-ALLERGEN | Non-Toxin | -0.23 | -0.48 | 0.51 | -3 | 4.04 | 2199.69 | outside |
| 765 | 781 | RALTGIAVEQDKNTQE | 0.4494 | 16 | ABCPRED | 1 | ALLERGEN | Non-Toxin | -0.3 | -0.94 | 0.54 | -1 | 4.68 | 1773.17 | outside |
| 1255 | 1271 | KFDEDDSEPVLKGVKL | 0.7015 | 16 | ABCPRED | 1 | ALLERGEN | Non-Toxin | -0.26 | -0.82 | 0.95 | -2 | 4.44 | 1819.27 | inside |
| 1037 | 1046 | SKRVDFCGK | 1.7321 | 9 | literature | 1 | NON-ALLERGEN | Non-Toxin | -0.4 | -0.83 | 0.81 | 2 | 9.36 | 1039.33 | outside |
| 1150 | 1168 | EELDKYFKNHTSPDVDLG | 0.4312 | 18 | ABCPRED | 1 | NON-ALLERGEN | Non-Toxin | -0.26 | -1.23 | 0.59 | -2.5 | 4.43 | 2107.52 | outside |
| 808 | 820 | DPSKPSKRSFIE | 0.3737 | 12 | ABCPRED | 1 | NON-ALLERGEN | Non-Toxin | -0.41 | -1.47 | 0.97 | 1 | 8.93 | 1390.71 | outside |
| 837 | 857 | YGDCLGDIAARDLICAQKFN | 0.4753 | 20 | ABCPRED | 1 | ALLERGEN | Non-Toxin | -0.1 | 0.09 | 0 | -1 | 4.43 | 2186.77 | outside |
| 760 | 778 | CTQLNRALTGIAVEQDKN | 0.71 | 18 | ABCPRED | 1 | ALLERGEN | Non-Toxin | -0.23 | -0.49 | 0.17 | 0 | 6.39 | 1974.49 | outside |
| 1173 | 1189 | NASVVNIQKEIDRLNE | 0.2791 | 16 | ABCPRED | 1 | NON-ALLERGEN | Non-Toxin | -0.27 | -0.67 | 0.45 | -1 | 4.68 | 1842.28 | outside |
| 1136 | 1153 | TVYDPLQPELDSFKEEL | -0.0634 | 17 | COVIDep | 1 | NON-ALLERGEN | Non-Toxin | -0.17 | -0.74 | 0.38 | -4 | 3.84 | 2023.47 | outside |
| 499 | 519 | PTNGVGYQPYRVVVLSFELL | 0.6919 | 20 | ABCPRED | 1 | NON-ALLERGEN | Non-Toxin | 0.02 | 0.39 | -0.61 | 0 | 6.35 | 2251.92 | outside |
| 727 | 743 | LPVSMTKTSVDCTMYI | 1.1469 | 16 | ABCPRED | 1 | ALLERGEN | Non-Toxin | 0 | 0.56 | -0.44 | 0 | 6.15 | 1789.38 | outside |
| 1252 | 1270 | SCCKFDEDDSEPVLKGVK | 0.5409 | 18 | ABCPRED | 1 | NON-ALLERGEN | Non-Toxin | -0.27 | -0.71 | 0.85 | -2 | 4.44 | 1999.48 | inside |
| 1170 | 1190 | SGINASVVNIQKEIDRLNEV | 0.3526 | 20 | ABCPRED | 1 | NON-ALLERGEN | Non-Toxin | -0.16 | -0.16 | 0.21 | -1 | 4.68 | 2198.77 | outside |
| 350 | 362 | VYAWNRKRISNC | 0.3253 | 12 | BCEPRED | 1 | NON-ALLERGEN | Non-Toxin | -0.35 | -0.83 | -0.07 | 3 | 10.07 | 1509.91 | outside |
| 1256 | 1274 | FDEDDSEPVLKGVKLHYT | 0.9755 | 18 | AAP | 1 | NON-ALLERGEN | Non-Toxin | -0.2 | -0.81 | 0.5 | -2.5 | 4.43 | 2092.55 | inside |
| 239 | 257 | QTLLALHRSYLTPGDSSS | 0.4919 | 18 | ABCPRED | 1 | NON-ALLERGEN | Non-Toxin | -0.14 | -0.31 | -0.22 | 0.5 | 7.09 | 1946.41 | outside |
| 188 | 198 | NLREFVFKNI | -0.0209 | 10 | ABCPRED | 1 | NON-ALLERGEN | Non-Toxin | -0.17 | -0.08 | -0.07 | 1 | 9.1 | 1279.65 | outside |
| 221 | 239 | SALEPLVDLPIGINITRF | 0.671 | 18 | ABCPRED | 1 | NON-ALLERGEN | Non-Toxin | 0.06 | 0.76 | -0.34 | -1 | 4.38 | 1968.6 | outside |
| 327 | 345 | VRFPNITNLCPFGEVFNA | 0.639 | 18 | ABCPRED | 1 | NON-ALLERGEN | Non-Toxin | 0 | 0.37 | -0.52 | 0 | 6.32 | 2038.61 | outside |
| 1079 | 1095 | PAICHDGKAHFPREGV | -0.35 | 16 | ABCPRED | 1 | NON-ALLERGEN | Non-Toxin | -0.15 | -0.51 | 0.2 | 1 | 7.22 | 1734.19 | outside |

| | | | | | | | | | | | | | | | |
|---|---|---|---|---|---|---|---|---|---|---|---|---|---|---|---|
| 291 | 309 | CALDPLSETKCTLKSFTV | 0.6686 | 18 | ABCPRED | 1 | ALLERGEN | Non-Toxin | -0.09 | 0.28 | -0.03 | 0 | 6.36 | 1956.55 | outside |
| 1133 | 1151 | VNNTVYDPLQPELDSFKE | 0.0831 | 18 | AAP | 1 | NON-ALLERGEN | Toxin | -0.2 | -0.87 | 0.23 | -3 | 3.92 | 2108.55 | outside |
| 630 | 648 | TPTWRVYSTGSNVFQTRA | 0.4094 | 18 | BCPREDS | 1 | NON-ALLERGEN | Non-Toxin | -0.2 | -0.64 | -0.35 | 2 | 10.84 | 2071.53 | outside |
| 132 | 150 | EFQFCNDPFLGVYYHKNN | 0.6373 | 18 | ABCPRED | 1 | ALLERGEN | Toxin | -0.13 | -0.77 | -0.39 | -0.5 | 5.33 | 2235.72 | outside |
| 138 | 156 | DPFLGVYYHKNNKSWMES | 0.5191 | 18 | ABCPRED | 1 | NON-ALLERGEN | Toxin | -0.18 | -1.08 | -0.14 | 0.5 | 7.09 | 2215.73 | outside |
| 400 | 416 | FVIRGDEVRQIAPGQT | 0.5147 | 16 | ABCPRED | 1 | ALLERGEN | Non-Toxin | -0.17 | -0.26 | 0.15 | 0 | 6.42 | 1786.26 | outside |
| 1104 | 1122 | VTQRNFYEPQIITTDNTF | 0.6261 | 18 | ABCPRED | 1 | ALLERGEN | Non-Toxin | -0.18 | -0.69 | -0.23 | -1 | 4.38 | 2187.66 | outside |
| 352 | 372 | AWNRKRISNCVADYSVLYNS | 0.1387 | 20 | ABCPRED | 1 | NON-ALLERGEN | Non-Toxin | -0.24 | -0.5 | -0.15 | 2 | 9.25 | 2359.92 | outside |
| 224 | 240 | EPLVDLPIGINITRFQ | 0.8017 | 16 | ABCPRED | 1 | NON-ALLERGEN | Non-Toxin | -0.01 | 0.33 | -0.25 | -1 | 4.38 | 1825.39 | outside |
| 93 | 111 | ASTEKSNIIRGWIFGTTL | -0.2071 | 18 | BCPREDS | 1 | NON-ALLERGEN | Non-Toxin | -0.05 | 0.06 | -0.28 | 1 | 9.1 | 1994.55 | outside |
| 304 | 324 | KSFTVEKGIYQTSNFRVQPT | 0.658 | 20 | ABCPRED | 1 | NON-ALLERGEN | Non-Toxin | -0.23 | -0.74 | 0 | 2 | 9.72 | 2330.92 | outside |
| 294 | 326 | DPLSETKCTLKSFTVEKGIYQTSNFRVQPTES | 0.6225 | 32 | BCEPRED | 1 | NON-ALLERGEN | Non-Toxin | -0.23 | -0.74 | 0.22 | 0 | 6.56 | 3635.53 | outside |
| 95 | 107 | TEKSNIIRGWIF | -0.7069 | 12 | ABCPRED | 1 | ALLERGEN | Non-Toxin | -0.1 | -0.16 | -0.18 | 1 | 9.1 | 1463.88 | outside |
| 847 | 865 | RDLICAQKFNGLTVLPPL | 0.4425 | 18 | ABCPRED | 1 | ALLERGEN | Non-Toxin | -0.04 | 0.43 | -0.31 | 1 | 8.57 | 1998.69 | outside |
| 25 | 43 | PPAYTNSFTRGVYYPDKV | 0.1859 | 18 | ABCPRED,BCPREDS,FBCPred | 3 | NON-ALLERGEN | Non-Toxin | -0.16 | -0.76 | -0.23 | 1 | 8.76 | 2075.55 | outside |
| 86 | 104 | FNDGVYFASTEKSNIIRG | 0.0569 | 18 | ABCPRED | 1 | NON-ALLERGEN | Non-Toxin | -0.13 | -0.34 | -0.02 | 0 | 6.42 | 2018.47 | outside |
| 17 | 47 | NLTTRTQLPPAYTNSFTRGVYYPDKVFRSS | 0.3475 | 30 | BCEPRED | 1 | NON-ALLERGEN | Non-Toxin | -0.24 | -0.77 | -0.15 | 3 | 10 | 3481.31 | outside |
| 1026 | 1042 | ATKMSECVLGQSKRVD | 0.6341 | 16 | ABCPRED | 1 | NON-ALLERGEN | Non-Toxin | -0.27 | -0.44 | 0.49 | 1 | 8.53 | 1752.26 | outside |
| 36 | 63 | VYYPDKVFRSSVLHSTQDLFLPFFSNV | 0.0327 | 27 | BCEPRED | 1 | NON-ALLERGEN | Toxin | -0.04 | 0.15 | -0.49 | 0.5 | 7.09 | 3207.04 | outside |
| 1088 | 1106 | HFPREGVFVSNGTHWFVT | 0.2217 | 18 | ABCPRED | 1 | NON-ALLERGEN | Non-Toxin | -0.02 | -0.13 | -0.59 | 1 | 7.26 | 2117.62 | outside |
| 246 | 264 | RSYLTPGDSSSGWTAGAA | 0.4145 | 18 | ABCPRED | 1 | NON-ALLERGEN | Non-Toxin | -0.1 | -0.47 | -0.14 | 0 | 6.19 | 1784.11 | outside |
| 23 | 43 | QLPPAYTNSFTRGVYYPDKV | 0.2964 | 20 | FBCPred | 1 | NON-ALLERGEN | Non-Toxin | -0.15 | -0.67 | -0.29 | 1 | 8.76 | 2316.88 | outside |
| 625 | 643 | HADQLTPTWRVYSTGSNV | 0.5334 | 18 | FBCPred | 1 | NON-ALLERGEN | Non-Toxin | -0.16 | -0.67 | -0.32 | 0.5 | 7.09 | 2032.46 | outside |
| 246 | 260 | RSYLTPGDSSSGWT | 0.512 | 14 | ABCPRED,FBCPred | 2 | NON-ALLERGEN | Non-Toxin | -0.19 | -0.96 | -0.08 | 0 | 6.19 | 1513.77 | outside |
| 395 | 415 | VYADSFVIRGDEVRQIAPGQ | 0.1515 | 20 | ABCPRED | 1 | ALLERGEN | Non-Toxin | -0.14 | -0.15 | 0.09 | -1 | 4.56 | 2220.76 | outside |
| 59 | 82 | FSNVTWFHAIHVSGTNGTKRFDN | 0.6767 | 23 | IEDB,literature | 2 | ALLERGEN | Non-Toxin | -0.12 | -0.48 | -0.36 | 2 | 9.11 | 2636.22 | outside |
| 626 | 640 | ADQLTPTWRVYSTG | 0.6906 | 14 | ABCPRED,FBCPred | 2 | NON-ALLERGEN | Non-Toxin | -0.15 | -0.63 | -0.3 | 0 | 6.19 | 1594.94 | outside |
| 211 | 229 | NLVRDLPQGFSALEPLVD | 0.1726 | 18 | FBCPred | 1 | ALLERGEN | Non-Toxin | -0.07 | 0.1 | -0.03 | -2 | 4.03 | 1983.52 | outside |
| 89 | 109 | GVYFASTEKSNIIRGWIFGT | -0.0328 | 20 | ABCPRED | 1 | NON-ALLERGEN | Non-Toxin | 0 | 0.16 | -0.45 | 1 | 8.93 | 2246.85 | outside |
| 449 | 467 | YNYLYRLFRKSNLKPFER | 0.0951 | 18 | ABCPRED | 1 | ALLERGEN | Non-Toxin | -0.38 | -1.17 | 0.08 | 4 | 10.17 | 2408.06 | outside |
| 1031 | 1049 | ECVLGQSKRVDFCGKGYH | 1.009 | 18 | ABCPRED | 1 | NON-ALLERGEN | Non-Toxin | -0.21 | -0.52 | 0.19 | 1.5 | 8.35 | 2026.58 | outside |
| 815 | 831 | RSFIEDLLFNKVTLAD | 0.1028 | 16 | ABCPRED | 1 | ALLERGEN | Non-Toxin | -0.1 | 0.22 | 0.06 | -1 | 4.56 | 1881.39 | outside |
| 310 | 328 | KGIYQTSNFRVQPTESIV | 0.6139 | 18 | ABCPRED | 1 | ALLERGEN | Non-Toxin | -0.17 | -0.47 | -0.11 | 1 | 8.93 | 2067.6 | outside |
| 200 | 218 | YFKIYSKHTPINLVRDLP | 0.5291 | 18 | ABCPRED | 1 | NON-ALLERGEN | Toxin | -0.14 | -0.34 | -0.23 | 2.5 | 9.55 | 2204.87 | outside |
| 67 | 85 | AIHVSGTNGTKRFDNPVL | 0.4566 | 18 | ABCPRED | 1 | ALLERGEN | Non-Toxin | -0.13 | -0.3 | -0.07 | 1.5 | 9.1 | 1926.43 | outside |
| 407 | 423 | VRQIAPGQTGKIADYN | 1.3372 | 16 | ABCPRED | 1 | NON-ALLERGEN | Non-Toxin | -0.19 | -0.62 | 0.05 | 1 | 8.93 | 1731.18 | outside |
| 267 | 285 | VGYLQPRTFLLKYNENGT | 0.5538 | 18 | ABCPRED | 1 | NON-ALLERGEN | Toxin | -0.15 | -0.58 | -0.29 | 1 | 8.83 | 2113.68 | outside |
| 1030 | 1046 | SECVLGQSKRVDFCGK | 1.0135 | 16 | ABCPRED | 1 | NON-ALLERGEN | Non-Toxin | -0.24 | -0.33 | 0.41 | 1 | 8.36 | 1756.25 | outside |
| 244 | 262 | LHRSYLTPGDSSSGWTAG | 0.3768 | 18 | FBCPred | 1 | NON-ALLERGEN | Non-Toxin | -0.12 | -0.63 | -0.22 | 0.5 | 7.09 | 1892.27 | outside |
| 179 | 199 | LEGKQGNFKNLREFVFKNID | 0.7083 | 20 | ABCPRED | 1 | NON-ALLERGEN | Toxin | -0.26 | -0.84 | 0.37 | 1 | 8.83 | 2397.04 | outside |
| 309 | 329 | EKGIYQTSNFRVQPTESIVR | 0.4085 | 20 | ABCPRED | 1 | NON-ALLERGEN | Non-Toxin | -0.27 | -0.83 | 0.2 | 1 | 8.93 | 2352.93 | outside |
| 202 | 220 | KIYSKHTPINLVRDLPQG | 0.5086 | 18 | ABCPRED | 1 | NON-ALLERGEN | Non-Toxin | -0.21 | -0.64 | 0.04 | 2.5 | 9.72 | 2079.71 | outside |
| 813 | 825 | SKRSFIEDLLFN | -0.0939 | 12 | ABCPRED | 1 | NON-ALLERGEN | Non-Toxin | -0.2 | -0.23 | 0.2 | 0 | 6.42 | 1468.84 | outside |
| 800 | 818 | FNFSQILPDPSKPSKRSF | 0.6041 | 18 | ABCPRED | 1 | ALLERGEN | Non-Toxin | -0.23 | -0.78 | 0.14 | 2 | 10.01 | 2095.63 | outside |
| 71 | 85 | SGTNGTKRFDNPVL | 0.3982 | 14 | FBCPred | 1 | NON-ALLERGEN | Non-Toxin | -0.25 | -0.91 | 0.22 | 1 | 9.1 | 1505.85 | outside |
| 459 | 471 | SNLKPFERDIST | 0.2978 | 12 | ABCPRED | 1 | NON-ALLERGEN | Non-Toxin | -0.31 | -0.97 | 0.52 | 0 | 6.42 | 1406.72 | outside |
| 187 | 199 | KNLREFVFKNID | 0.2991 | 12 | ABCPRED | 1 | NON-ALLERGEN | Non-Toxin | -0.3 | -0.68 | 0.44 | 1 | 8.93 | 1522.94 | outside |
| 802 | 818 | FSQILPDPSKPSKRSF | 0.2789 | 16 | ABCPRED | 1 | ALLERGEN | Non-Toxin | -0.26 | -0.84 | 0.3 | 2 | 10.01 | 1834.32 | outside |
| 68 | 88 | IHVSGTNGTKRFDNPVLPFN | 0.5334 | 20 | ABCPRED,literature | 2 | NON-ALLERGEN | Non-Toxin | -0.14 | -0.47 | -0.15 | 1.5 | 9.1 | 2213.78 | outside |
| 463 | 475 | PFERDISTEIYQ | -0.4749 | 12 | ABCPRED | 1 | NON-ALLERGEN | Non-Toxin | -0.24 | -0.93 | 0.31 | -2 | 4.14 | 1497.79 | outside |
| 100 | 118 | IIRGWIFGTTLDSKTQSL | 0.1411 | 18 | ABCPRED | 1 | NON-ALLERGEN | Non-Toxin | -0.04 | 0.17 | -0.35 | 1 | 9.1 | 2036.64 | outside |
| 151 | 163 | SWMESEFRVYSS | 0.3082 | 12 | ABCPRED | 1 | NON-ALLERGEN | Non-Toxin | -0.19 | -0.67 | -0.07 | -1 | 4.54 | 1507.79 | outside |
| 172 | 188 | SQPFLMDLEGKQGNFK | 0.9284 | 16 | ABCPRED | 1 | NON-ALLERGEN | Toxin | -0.19 | -0.84 | 0.19 | 0 | 6.42 | 1839.34 | outside |
| 208 | 226 | TPINLVRDLPQGFSALEP | 0.7363 | 18 | ABCPRED | 1 | NON-ALLERGEN | Non-Toxin | -0.07 | -0.03 | -0.13 | -1 | 4.38 | 1967.52 | outside |
| 814 | 830 | KRSFIEDLLFNKVTLA | 0.1475 | 16 | ABCPRED | 1 | NON-ALLERGEN | Non-Toxin | -0.13 | 0.2 | 0.06 | 1 | 8.93 | 1894.48 | outside |
| 349 | 367 | SVYAWNRKRISNCVADYS | 0.2293 | 18 | ABCPRED | 1 | NON-ALLERGEN | Non-Toxin | -0.26 | -0.57 | -0.08 | 2 | 9.25 | 2132.62 | outside |
| 95 | 111 | TEKSNIIRGWIFGTTL | -0.2839 | 16 | ABCPRED | 1 | NON-ALLERGEN | Non-Toxin | -0.06 | 0.01 | -0.3 | 1 | 9.1 | 1836.37 | outside |
| 209 | 229 | PINLVRDLPQGFSALEPLVD | 0.4744 | 20 | FBCPred | 1 | NON-ALLERGEN | Non-Toxin | -0.03 | 0.23 | -0.12 | -2 | 4.03 | 2193.83 | outside |
| 1106 | 1124 | QRNFYEPQIITTDNTFVS | 0.4782 | 18 | ABCPRED | 1 | NON-ALLERGEN | Non-Toxin | -0.19 | -0.69 | -0.19 | -1 | 4.38 | 2173.63 | outside |
| 206 | 222 | KHTPINLVRDLPQGFS | 0.6403 | 16 | IEDB,literature | 2 | NON-ALLERGEN | Non-Toxin | -0.17 | -0.51 | -0.04 | 1.5 | 9.1 | 1822.34 | outside |
| 36 | 48 | VYYPDKVFRSSV | -0.3137 | 12 | ABCPRED | 1 | ALLERGEN | Non-Toxin | -0.16 | -0.19 | -0.17 | 1 | 8.83 | 1459.82 | outside |
| 801 | 817 | NFSQILPDPSKPSKRS | 0.505 | 16 | ABCPRED | 1 | ALLERGEN | Toxin | -0.34 | -1.23 | 0.47 | 2 | 10.01 | 1801.25 | outside |
| 577 | 597 | RDPQTLEILDITPCSFGGVS | 1.4479 | 20 | ABCPRED | 1 | NON-ALLERGEN | Non-Toxin | -0.08 | 0.03 | -0.01 | -2 | 4.03 | 2148.71 | outside |
| 436 | 456 | WNSNNLDSKVGGNYNYLYRL | 0.5304 | 20 | ABCPRED | 1 | NON-ALLERGEN | Non-Toxin | -0.22 | -1.05 | -0.33 | 1 | 8.76 | 2390.9 | outside |
| 351 | 367 | YAWNRKRISNCVADYS | 0.2628 | 16 | ABCPRED | 1 | NON-ALLERGEN | Non-Toxin | -0.31 | -0.86 | -0.02 | 2 | 9.25 | 1946.38 | outside |
| 35 | 51 | GVYYPDKVFRSSVLHS | -0.1113 | 16 | ABCPRED | 1 | ALLERGEN | Non-Toxin | -0.12 | -0.18 | -0.25 | 1.5 | 8.84 | 1854.32 | outside |
| 35 | 53 | GVYYPDKVFRSSVLHSTQ | 0.0785 | 18 | ABCPRED | 1 | NON-ALLERGEN | Toxin | -0.15 | -0.39 | -0.23 | 1.5 | 8.84 | 2083.59 | outside |
| 190 | 200 | REFVFKNIDG | -0.325 | 10 | ABCPRED | 1 | ALLERGEN | Non-Toxin | -0.22 | -0.5 | 0.39 | 0 | 6.42 | 1224.52 | outside |
| 27 | 45 | AYTNSFTRGVYYPDKVFR | -0.172 | 18 | ABCPRED | 1 | NON-ALLERGEN | Non-Toxin | -0.22 | -0.68 | -0.21 | 2 | 9.55 | 2184.68 | outside |
| 223 | 241 | LEPLVDLPIGINITRFQT | 0.6852 | 18 | ABCPRED | 1 | NON-ALLERGEN | Non-Toxin | 0.01 | 0.47 | -0.34 | -1 | 4.38 | 2039.69 | outside |
| 1025 | 1042 | AATKMSECVLGQSKRVD | 0.6159 | 17 | COVIDep,literature | 2 | NON-ALLERGEN | Non-Toxin | -0.24 | -0.31 | 0.43 | 1 | 8.53 | 1823.35 | outside |
| 1084 | 1102 | DGKAHFPREGVFVSNGTH | 0.2385 | 18 | AAP | 1 | NON-ALLERGEN | Non-Toxin | -0.17 | -0.77 | 0.14 | 1 | 7.26 | 1955.38 | outside |
| 621 | 641 | PVAIHADQLTPTWRVYSTGS | 0.5993 | 20 | ABCPRED | 1 | ALLERGEN | Non-Toxin | -0.07 | -0.2 | -0.41 | 0.5 | 7.09 | 2199.74 | outside |
| 76 | 88 | TKRFDNPVLPFN | 0.3512 | 12 | ABCPRED | 1 | NON-ALLERGEN | Toxin | -0.24 | -0.77 | 0.06 | 1 | 9.1 | 1447.82 | outside |
| 174 | 186 | PFLMDLEGKQGN | 1.2067 | 12 | ABCPRED | 1 | ALLERGEN | Non-Toxin | -0.13 | -0.67 | 0.17 | -1 | 4.38 | 1348.72 | outside |
| 348 | 366 | ASVYAWNRKRISNCVADY | 0.1958 | 18 | ABCPRED,AAP | 2 | NON-ALLERGEN | Non-Toxin | -0.23 | -0.43 | -0.13 | 2 | 9.25 | 2116.62 | outside |
| 347 | 365 | FASVYAWNRKRISNCVAD | 0.2502 | 18 | ABCPRED | 1 | NON-ALLERGEN | Non-Toxin | -0.2 | -0.2 | -0.14 | 2 | 9.36 | 2100.62 | outside |

| | | | | | | | | | | | | | | | |
|---|---|---|---|---|---|---|---|---|---|---|---|---|---|---|---|
| 91 | 109 | YFASTEKSNIIRGWIFGT | -0.1278 | 18 | ABCPRED | 1 | NON-ALLERGEN | Non-Toxin | -0.04 | -0.03 | -0.42 | 1 | 8.93 | 2090.63 | outside |
| 39 | 48 | PDKVFRSSV | -0.7314 | 9 | IEDB | 1 | NON-ALLERGEN | Non-Toxin | -0.28 | -0.43 | 0.46 | 1 | 9.1 | 1034.29 | outside |
| 625 | 641 | HADQLTPTWRVYSTGS | 0.6093 | 16 | ABCPRED | 1 | NON-ALLERGEN | Non-Toxin | -0.18 | -0.8 | -0.28 | 0.5 | 7.09 | 1819.19 | outside |
| 28 | 46 | YTNSFTRGVYYPDKVFRS | -0.1806 | 18 | ABCPRED | 1 | ALLERGEN | Non-Toxin | -0.24 | -0.82 | -0.16 | 2 | 9.55 | 2200.68 | outside |
| 148 | 160 | NNKSWMESEFRV | 0.6015 | 12 | ABCPRED | 1 | NON-ALLERGEN | Non-Toxin | -0.34 | -1.33 | 0.36 | 0 | 6.49 | 1526.85 | outside |
| 204 | 216 | YSKHTPINLVRD | 0.5522 | 12 | ABCPRED | 1 | NON-ALLERGEN | Non-Toxin | -0.28 | -0.87 | 0.1 | 1.5 | 8.94 | 1442.81 | outside |
| 174 | 190 | PFLMDLEGKQGNFKNL | 1.0418 | 16 | ABCPRED | 1 | NON-ALLERGEN | Toxin | -0.14 | -0.55 | 0.06 | 0 | 6.42 | 1851.4 | outside |
| 723 | 743 | TTEILPVSMTKTSVDCTMYI | 1.1726 | 20 | ABCPRED,BCPREDS | 2 | ALLERGEN | Non-Toxin | -0.01 | 0.43 | -0.34 | -1 | 4.38 | 2233.93 | outside |
| 98 | 116 | SNIIRGWIFGTTLDSKTQ | 0.003 | 18 | ABCPRED | 1 | NON-ALLERGEN | Non-Toxin | -0.11 | -0.23 | -0.24 | 1 | 9.1 | 2037.58 | outside |
| 69 | 89 | HVSGTNGTKRFDNPVLPFND | 0.4802 | 20 | FBCPred | 1 | ALLERGEN | Non-Toxin | -0.21 | -0.88 | 0.09 | 0.5 | 7.1 | 2215.7 | outside |
| 31 | 43 | SFTRGVYYPDKV | 0.548 | 12 | ABCPRED,AAP | 2 | ALLERGEN | Non-Toxin | -0.18 | -0.57 | -0.1 | 1 | 8.83 | 1431.77 | outside |
| 172 | 190 | SQPFLMDLEGKQGNFKNL | 0.8829 | 18 | ABCPRED | 1 | NON-ALLERGEN | Non-Toxin | -0.17 | -0.73 | 0.08 | 0 | 6.42 | 2066.64 | outside |
| 1097 | 1115 | SNGTHWFVTQRNFYEPQI | 0.2295 | 18 | ABCPRED | 1 | ALLERGEN | Non-Toxin | -0.17 | -0.96 | -0.46 | 0.5 | 7.1 | 2224.69 | outside |
| 1090 | 1108 | PREGVFVSNGTHWFVTQR | 0.4436 | 18 | ABCPRED | 1 | NON-ALLERGEN | Non-Toxin | -0.17 | -0.56 | -0.25 | 1.5 | 9.95 | 2117.62 | outside |
| 457 | 467 | RKSNLKPFER | 0.8187 | 10 | ABCPRED | 1 | NON-ALLERGEN | Non-Toxin | -0.62 | -1.96 | 1.12 | 3 | 11.01 | 1274.62 | outside |
| 408 | 424 | RQIAPGQTGKIADYNY | 1.5799 | 16 | ABCPRED | 1 | NON-ALLERGEN | Non-Toxin | -0.22 | -0.97 | 0 | 1 | 8.83 | 1795.22 | outside |
| 408 | 428 | RQIAPGQTGKIADYNYKLPD | 1.4102 | 20 | ABCPRED,BCPREDS,literature | 3 | NON-ALLERGEN | Non-Toxin | -0.24 | -1.03 | 0.21 | 1 | 8.76 | 2248.82 | outside |
| 1107 | 1125 | RNFYEPQIITTDNTFVSG | 0.2808 | 18 | ABCPRED | 1 | ALLERGEN | Non-Toxin | -0.14 | -0.52 | -0.21 | -1 | 4.38 | 2102.55 | outside |
| 337 | 355 | PFGEVFNATRFASVYAWN | 0.0554 | 18 | ABCPRED | 1 | NON-ALLERGEN | Non-Toxin | 0 | 0.08 | -0.63 | 0 | 6.35 | 2076.54 | outside |
| 76 | 94 | TKRFDNPVLPFNDGVYFA | 0.2892 | 18 | ABCPRED | 1 | NON-ALLERGEN | Non-Toxin | -0.11 | -0.31 | -0.17 | 0 | 6.31 | 2100.61 | outside |
| 41 | 59 | KVFRSSVLHSTQDLFLPF | 0.0218 | 18 | ABCPRED | 1 | ALLERGEN | Non-Toxin | -0.07 | 0.27 | -0.37 | 1.5 | 9.1 | 2121.73 | outside |
| 77 | 95 | KRFDNPVLPFNDGVYFAS | 0.2197 | 18 | ABCPRED | 1 | NON-ALLERGEN | Non-Toxin | -0.12 | -0.32 | -0.13 | 0 | 6.31 | 2086.58 | outside |
| 1237 | 1265 | MTSCCSCLKGCCSCGSCCKFDEDDSEPV | 0.1043 | 28 | IEDB | 1 | ALLERGEN | Non-Toxin | -0.16 | 0.01 | 0.25 | -3 | 4.11 | 2941.71 | inside |
| 1035 | 1053 | GQSKRVDFCGKGYHLMSF | 0.9773 | 18 | ABCPRED | 1 | NON-ALLERGEN | Non-Toxin | -0.18 | -0.48 | -0.03 | 2.5 | 9.25 | 2060.64 | outside |
| 243 | 261 | ALHRSYLTPGDSSSGWTA | 0.4894 | 18 | ABCPRED | 1 | NON-ALLERGEN | Non-Toxin | -0.12 | -0.51 | -0.24 | 0.5 | 7.09 | 1906.29 | outside |
| 1098 | 1114 | NGTHWFVTQRNFYEPQ | 0.222 | 16 | ABCPRED | 1 | NON-ALLERGEN | Non-Toxin | -0.22 | -1.31 | -0.42 | 0.5 | 7.1 | 2024.42 | outside |
| 799 | 817 | GFNFSQILPDPSKPSKRS | 0.6974 | 18 | ABCPRED | 1 | ALLERGEN | Non-Toxin | -0.26 | -0.96 | 0.28 | 2 | 10.01 | 2005.51 | outside |
| 71 | 87 | SGTNGTKRFDNPVLPF | 0.4673 | 16 | ABCPRED | 1 | NON-ALLERGEN | Non-Toxin | -0.19 | -0.72 | 0.04 | 1 | 9.1 | 1750.17 | outside |
| 202 | 218 | KIYSKHTPINLVRDLP | 0.4252 | 16 | ABCPRED | 1 | NON-ALLERGEN | Non-Toxin | -0.2 | -0.48 | 0.04 | 2.5 | 9.72 | 1894.49 | outside |
| 74 | 86 | NGTKRFDNPVLP | 0.0178 | 12 | ABCPRED,BCPREDS | 2 | NON-ALLERGEN | Non-Toxin | -0.28 | -1.03 | 0.27 | 1 | 9.1 | 1357.7 | outside |
| 400 | 418 | FVIRGDEVRQIAPGQTGK | 0.8644 | 18 | ABCPRED | 1 | NON-ALLERGEN | Non-Toxin | -0.21 | -0.47 | 0.3 | 1 | 9.1 | 1971.52 | outside |
| 814 | 832 | KRSFIEDLLFNKVTLADA | 0.2279 | 18 | ABCPRED | 1 | ALLERGEN | Non-Toxin | -0.14 | 0.08 | 0.19 | 0 | 6.47 | 2080.67 | outside |
| 196 | 216 | NIDGYFKIYSKHTPINLVRD | 0.4217 | 20 | ABCPRED | 1 | NON-ALLERGEN | Non-Toxin | -0.18 | -0.56 | -0.05 | 1.5 | 8.76 | 2394.03 | outside |
| 166 | 186 | CTFEYVSQPFLMDLEGKQGN | 0.748 | 20 | ABCPRED | 1 | ALLERGEN | Non-Toxin | -0.11 | -0.42 | -0.11 | -2 | 4.14 | 2306.89 | outside |
| 624 | 642 | IHADQLTPTWRVYSTGSN | 0.4812 | 18 | ABCPRED | 1 | NON-ALLERGEN | Non-Toxin | -0.15 | -0.66 | -0.33 | 0.5 | 7.09 | 2046.49 | outside |
| 190 | 202 | REFVFKNIDGYF | -0.2327 | 12 | ABCPRED | 1 | NON-ALLERGEN | Non-Toxin | -0.13 | -0.29 | -0.07 | 0 | 6.42 | 1534.9 | outside |
| 34 | 44 | RGVYYPDKVF | 0.5225 | 10 | ABCPRED | 1 | ALLERGEN | Non-Toxin | -0.18 | -0.53 | -0.11 | 1 | 8.83 | 1243.56 | outside |
| 353 | 365 | WNRKRISNCVAD | 0.2338 | 12 | ABCPRED | 1 | NON-ALLERGEN | Non-Toxin | -0.41 | -1.01 | 0.38 | 2 | 9.55 | 1461.82 | outside |
| 246 | 266 | RSYLTPGDSSSGWTAGAAAY | 0.4627 | 20 | ABCPRED | 1 | ALLERGEN | Non-Toxin | -0.08 | -0.4 | -0.27 | 0 | 6.18 | 2018.39 | outside |
| 32 | 48 | FTRGVYYPDKVFRSSV | 0.0228 | 16 | ABCPRED | 1 | NON-ALLERGEN | Non-Toxin | -0.19 | -0.32 | -0.12 | 2 | 9.72 | 1921.4 | outside |
| 171 | 191 | VSQPFLMDLEGKQGNFKNLR | 0.7645 | 20 | ABCPRED | 1 | ALLERGEN | Non-Toxin | -0.22 | -0.67 | 0.15 | 1 | 8.93 | 2321.99 | outside |
| 152 | 162 | WMESEFRVYS | 0.3518 | 10 | ABCPRED | 1 | NON-ALLERGEN | Non-Toxin | -0.17 | -0.64 | -0.14 | -1 | 4.54 | 1333.61 | outside |
| 309 | 327 | EKGIYQTSNFRVQPTESI | 0.6643 | 18 | ABCPRED | 1 | ALLERGEN | Non-Toxin | -0.24 | -0.9 | 0.14 | 0 | 6.49 | 2097.58 | outside |
| 168 | 186 | FEYVSQPFLMDLEGKQGN | 0.9891 | 18 | ABCPRED | 1 | ALLERGEN | Non-Toxin | -0.11 | -0.56 | -0.04 | -2 | 4.14 | 2102.62 | outside |
| 34 | 54 | RGVYYPDKVFRSSVLHSTQD | 0.2586 | 20 | ABCPRED | 1 | NON-ALLERGEN | Non-Toxin | -0.26 | -0.76 | 0.09 | 1.5 | 8.84 | 2354.89 | outside |
| 308 | 324 | VEKGIYQTSNFRVQPT | 0.8907 | 16 | ABCPRED | 1 | NON-ALLERGEN | Non-Toxin | -0.22 | -0.76 | -0.03 | 1 | 8.93 | 1867.33 | outside |
| 75 | 85 | GTKRFDNPVL | 0.2188 | 10 | ABCPRED | 1 | ALLERGEN | Non-Toxin | -0.26 | -0.73 | 0.3 | 1 | 9.1 | 1146.45 | outside |
| 803 | 819 | SQILPDPSKPSKRSFI | 0.2672 | 16 | ABCPRED | 1 | ALLERGEN | Non-Toxin | -0.25 | -0.73 | 0.34 | 2 | 10.01 | 1800.31 | outside |
| 397 | 415 | ADSFVIRGDEVRQIAPGQ | 0.2599 | 18 | ABCPRED | 1 | NON-ALLERGEN | Non-Toxin | -0.18 | -0.33 | 0.31 | -1 | 4.56 | 1958.42 | outside |
| 352 | 370 | AWNRKRISNCVADYSVLY | 0.2161 | 18 | ABCPRED | 1 | ALLERGEN | Toxin | -0.21 | -0.32 | -0.2 | 2 | 9.25 | 2158.71 | outside |
| 308 | 328 | VEKGIYQTSNFRVQPTESIV | 0.6006 | 20 | ABCPRED,AAP | 2 | ALLERGEN | Non-Toxin | -0.16 | -0.39 | -0.03 | 0 | 6.49 | 2295.88 | outside |
| 171 | 187 | VSQPFLMDLEGKQGNF | 0.9279 | 16 | ABCPRED | 1 | ALLERGEN | Non-Toxin | -0.09 | -0.33 | -0.09 | -1 | 4.38 | 1810.3 | outside |
| 1196 | 1216 | SLIDLQELGKYEQYIKWPWY | 1.0282 | 20 | ABCPRED,AAP | 2 | NON-ALLERGEN | Non-Toxin | -0.09 | -0.67 | -0.35 | -1 | 4.68 | 2573.26 | outside_transmembrane-region |
| 26 | 44 | PAYTNSFTRGVYYPDKVF | -0.0258 | 18 | ABCPRED | 1 | NON-ALLERGEN | Non-Toxin | -0.12 | -0.52 | -0.37 | 1 | 8.76 | 2125.61 | outside |
| 853 | 871 | QKFNGLTVLPPLLTDEMI | 0.1787 | 18 | ABCPRED | 1 | ALLERGEN | Non-Toxin | 0.01 | 0.32 | -0.32 | -1 | 4.38 | 2029.71 | outside |
| 78 | 98 | RFDNPVLPFNDGVYFASTEK | 0.3267 | 20 | ABCPRED | 1 | NON-ALLERGEN | Non-Toxin | -0.15 | -0.49 | 0.01 | -1 | 4.56 | 2316.83 | outside |
| 167 | 187 | TFEYVSQPFLMDLEGKQGNF | 1.009 | 20 | ABCPRED | 1 | NON-ALLERGEN | Non-Toxin | -0.08 | -0.4 | -0.18 | -2 | 4.14 | 2350.93 | outside |
| 206 | 224 | KHTPINLVRDLPQGFSAL | 0.6498 | 18 | ABCPRED | 1 | NON-ALLERGEN | Non-Toxin | -0.11 | -0.14 | -0.16 | 1.5 | 9.1 | 2006.61 | outside |
| 350 | 368 | VYAWNRKRISNCVADYSV | 0.1551 | 18 | ABCPRED | 1 | NON-ALLERGEN | Non-Toxin | -0.21 | -0.29 | -0.18 | 2 | 9.25 | 2144.68 | outside |
| 1030 | 1048 | SECVLGQSKRVDFCGKGY | 0.8392 | 18 | ABCPRED | 1 | NON-ALLERGEN | Non-Toxin | -0.21 | -0.38 | 0.23 | 1 | 8.34 | 1976.51 | outside |
| 201 | 217 | FKIYSKHTPINLVRDL | 0.6568 | 16 | ABCPRED | 1 | NON-ALLERGEN | Non-Toxin | -0.16 | -0.21 | -0.12 | 2.5 | 9.72 | 1944.55 | outside |
| 459 | 479 | SNLKPFERDISTEIYQAGST | 0.1665 | 20 | ABCPRED,literature | 2 | ALLERGEN | Non-Toxin | -0.22 | -0.78 | 0.24 | -1 | 4.68 | 2256.74 | outside |
| 32 | 44 | FTRGVYYPDKVF | 0.3305 | 12 | ABCPRED | 1 | ALLERGEN | Non-Toxin | -0.11 | -0.27 | -0.33 | 1 | 8.83 | 1491.87 | outside |
| 846 | 864 | ARDLICAQKFNGLTVLPP | 0.3804 | 18 | ABCPRED | 1 | NON-ALLERGEN | Non-Toxin | -0.06 | 0.32 | -0.23 | 1 | 8.57 | 1956.6 | outside |
| 139 | 157 | PFLGVYYHKNNKSWMESE | 0.5664 | 18 | ABCPRED | 1 | NON-ALLERGEN | Toxin | -0.18 | -1.08 | -0.14 | 0.5 | 7.09 | 2229.76 | outside |
| 462 | 478 | KPFERDISTEIYQAGS | -0.2463 | 16 | ABCPRED | 1 | NON-ALLERGEN | Non-Toxin | -0.24 | -0.9 | 0.41 | -1 | 4.68 | 1841.23 | outside |
| 66 | 84 | HAIHVSGTNGTKRFDNPV | 0.4408 | 18 | ABCPRED | 1 | ALLERGEN | Non-Toxin | -0.19 | -0.69 | 0.01 | 2 | 9.11 | 1950.41 | outside |
| 801 | 819 | NFSQILPDPSKPSKRSFI | 0.3238 | 18 | ABCPRED | 1 | NON-ALLERGEN | Non-Toxin | -0.23 | -0.69 | 0.18 | 2 | 10.01 | 2061.62 | outside |
| 210 | 226 | INLVRDLPQGFSALEP | 0.6332 | 16 | ABCPRED | 1 | NON-ALLERGEN | Toxin | -0.06 | 0.11 | -0.13 | -1 | 4.38 | 1769.27 | outside |
| 198 | 216 | DGYFKIYSKHTPINLVRD | 0.4887 | 18 | ABCPRED | 1 | NON-ALLERGEN | Non-Toxin | -0.2 | -0.68 | 0.03 | 1.5 | 8.76 | 2166.73 | outside |
| 96 | 108 | EKSNIIRGWIFG | -0.7155 | 12 | ABCPRED | 1 | NON-ALLERGEN | Non-Toxin | -0.07 | -0.13 | -0.15 | 1 | 9.1 | 1419.83 | outside |
| 94 | 106 | STEKSNIIRGWI | -0.8185 | 12 | ABCPRED | 1 | NON-ALLERGEN | Non-Toxin | -0.18 | -0.46 | 0.05 | 1 | 9.1 | 1403.78 | outside |
| 623 | 643 | AIHADQLTPTWRVYSTGSNV | 0.5062 | 20 | FBCPred | 1 | NON-ALLERGEN | Non-Toxin | -0.1 | -0.29 | -0.4 | 0.5 | 7.09 | 2216.73 | outside |
| 1024 | 1042 | LAATKMSECVLGQSKRVD | 0.73 | 18 | ABCPRED | 1 | NON-ALLERGEN | Non-Toxin | -0.2 | -0.08 | 0.31 | 1 | 8.53 | 1936.53 | outside |
| 102 | 118 | RGWIFGTTLDSKTQSL | 0.5054 | 16 | ABCPRED | 1 | NON-ALLERGEN | Non-Toxin | -0.14 | -0.37 | -0.17 | 1 | 9.1 | 1810.28 | outside |
| 182 | 202 | KQGNFKNLREFVFKNIDGYF | 0.1077 | 20 | ABCPRED | 1 | ALLERGEN | Non-Toxin | -0.22 | -0.78 | 0.07 | 2 | 9.55 | 2465.11 | outside |
| 168 | 184 | FEYVSQPFLMDLEGKQ | 0.8052 | 16 | ABCPRED | 1 | NON-ALLERGEN | Non-Toxin | -0.09 | -0.39 | -0.06 | -2 | 4.14 | 1931.43 | outside |

| | | | | | | | | | | | | | | | |
|---|---|---|---|---|---|---|---|---|---|---|---|---|---|---|---|
| 624 | 640 | IHADQLTPTWRVYSTG | 0.5299 | 16 | ABCPRED | 1 | NON-ALLERGEN | Non-Toxin | -0.12 | -0.47 | -0.41 | 0.5 | 7.09 | 1845.28 | outside |
| 309 | 325 | EKGIYQTSNFRVQPTE | 0.8559 | 16 | ABCPRED | 1 | NON-ALLERGEN | Non-Toxin | -0.3 | -1.24 | 0.25 | 0 | 6.49 | 1897.31 | outside |
| 33 | 47 | TRGVYYPDKVFRSS | 0.3326 | 14 | BCPREDS | 1 | NON-ALLERGEN | Non-Toxin | -0.3 | -0.86 | 0.15 | 2 | 9.72 | 1675.06 | outside |
| 1037 | 1053 | SKRVDFCGKGYHLMSF | 1.0195 | 16 | ABCPRED | 1 | ALLERGEN | Non-Toxin | -0.17 | -0.29 | -0.05 | 2.5 | 9.25 | 1875.42 | outside |
| 1099 | 1115 | GTHWFVTQRNFYEPQI | 0.259 | 16 | ABCPRED | 1 | ALLERGEN | Non-Toxin | -0.14 | -0.81 | -0.54 | 0.5 | 7.1 | 2023.48 | outside |
| 34 | 50 | RGVYYPDKVFRSSVLH | 0.1595 | 16 | ABCPRED | 1 | NON-ALLERGEN | Non-Toxin | -0.21 | -0.41 | -0.08 | 2.5 | 9.72 | 1923.43 | outside |
| 269 | 287 | YLQPRTFLLKYNENGTIT | 0.4452 | 18 | ABCPRED,FBCPred | 2 | NON-ALLERGEN | Non-Toxin | -0.16 | -0.58 | -0.33 | 1 | 8.83 | 2171.76 | outside |
| 68 | 84 | IHVSGTNGTKRFDNPV | 0.4407 | 16 | ABCPRED | 1 | NON-ALLERGEN | Non-Toxin | -0.2 | -0.69 | 0.07 | 1.5 | 9.1 | 1742.16 | outside |
| 73 | 91 | TNGTKRFDNPVLPFNDGV | 0.1605 | 18 | ABCPRED | 1 | NON-ALLERGEN | Non-Toxin | -0.2 | -0.75 | 0.11 | 0 | 6.31 | 1991.45 | outside |
| 626 | 638 | ADQLTPTWRVYS | 0.5764 | 12 | FBCPred | 1 | ALLERGEN | Non-Toxin | -0.18 | -0.64 | -0.32 | 0 | 6.19 | 1436.75 | outside |
| 1198 | 1216 | IDLQELGKYEQYIKWPWY | 1.0558 | 18 | ABCPRED | 1 | NON-ALLERGEN | Non-Toxin | -0.12 | -0.91 | -0.31 | -1 | 4.68 | 2372.99 | outside_transmembrane-region |
| 71 | 89 | SGTNGTKRFDNPVLPFND | 0.4296 | 18 | ABCPRED,BCPREDS,FBCPred | 3 | NON-ALLERGEN | Non-Toxin | -0.24 | -1.03 | 0.21 | 0 | 6.31 | 1979.39 | outside |
| 802 | 820 | FSQILPDPSKPSKRSFIE | 0.0988 | 18 | literature | 1 | NON-ALLERGEN | Non-Toxin | -0.22 | -0.69 | 0.33 | 1 | 8.93 | 2076.63 | outside |
| 150 | 160 | KSWMESEFRV | 0.2861 | 10 | ABCPRED | 1 | NON-ALLERGEN | Non-Toxin | -0.28 | -0.9 | 0.39 | 0 | 6.49 | 1298.61 | outside |
| 70 | 84 | VSGTNGTKRFDNPV | 0.4841 | 14 | ABCPRED,AAP | 2 | NON-ALLERGEN | Non-Toxin | -0.25 | -0.88 | 0.24 | 1 | 9.1 | 1491.82 | outside |
| 1105 | 1121 | TQRNFYEPQIITTDNT | 0.5783 | 16 | ABCPRED | 1 | ALLERGEN | Non-Toxin | -0.28 | -1.21 | -0.01 | -1 | 4.38 | 1941.32 | outside |
| 174 | 184 | PFLMDLEGKQ | 0.915 | 10 | ABCPRED | 1 | ALLERGEN | Non-Toxin | -0.11 | -0.41 | 0.18 | -1 | 4.38 | 1177.53 | outside |
| 459 | 469 | SNLKPFERDI | 0.5791 | 10 | ABCPRED | 1 | ALLERGEN | Non-Toxin | -0.33 | -1.02 | 0.64 | 0 | 6.42 | 1218.51 | outside |
| 268 | 285 | GYLQPRTFLLKYNENGT | 0.4239 | 17 | BCEPRED | 1 | NON-ALLERGEN | Non-Toxin | -0.2 | -0.86 | -0.22 | 1 | 8.83 | 2014.53 | outside |
| 398 | 414 | DSFVIRGDEVRQIAPG | 0.3572 | 16 | ABCPRED | 1 | NON-ALLERGEN | Non-Toxin | -0.18 | -0.26 | 0.37 | -1 | 4.56 | 1759.18 | outside |
| 69 | 87 | HVSGTNGTKRFDNPVLPF | 0.5183 | 18 | AAP | 1 | ALLERGEN | Non-Toxin | -0.16 | -0.58 | -0.08 | 1.5 | 9.1 | 1986.48 | outside |
| 1107 | 1119 | RNFYEPQIITTD | 0.3529 | 12 | IEDB,literature | 2 | NON-ALLERGEN | Non-Toxin | -0.23 | -0.92 | 0.02 | -1 | 4.38 | 1496.81 | outside |
| 396 | 414 | YADSFVIRGDEVRQIAPG | 0.2888 | 18 | ABCPRED | 1 | NON-ALLERGEN | Non-Toxin | -0.14 | -0.21 | 0.17 | -1 | 4.56 | 1993.46 | outside |
| 463 | 481 | PFERDISTEIYQAGSTPC | -0.1057 | 18 | ABCPRED | 1 | ALLERGEN | Non-Toxin | -0.16 | -0.57 | 0.12 | -2 | 4.14 | 2014.44 | outside |
| 463 | 473 | PFERDISTEI | -0.3446 | 10 | ABCPRED | 1 | NON-ALLERGEN | Non-Toxin | -0.22 | -0.63 | 0.58 | -2 | 4.14 | 1206.45 | outside |
| 460 | 478 | NLKPFERDISTEIYQAGS | 0.2279 | 18 | ABCPRED | 1 | ALLERGEN | Non-Toxin | -0.22 | -0.78 | 0.27 | -1 | 4.68 | 2068.53 | outside |
| 28 | 44 | YTNSFTRGVYYPDKVF | -0.0699 | 16 | ABCPRED | 1 | ALLERGEN | Non-Toxin | -0.15 | -0.59 | -0.39 | 1 | 8.76 | 1957.39 | outside |
| 270 | 286 | LQPRTFLLKYNENGTI | 0.3917 | 16 | ABCPRED | 1 | NON-ALLERGEN | Non-Toxin | -0.17 | -0.53 | -0.2 | 1 | 8.93 | 1907.45 | outside |
| 1101 | 1117 | HWFVTQRNFYEPQIIT | 0.4571 | 16 | ABCPRED | 1 | NON-ALLERGEN | Non-Toxin | -0.1 | -0.51 | -0.66 | 0.5 | 7.1 | 2079.59 | outside |
| 622 | 638 | VAIHADQLTPTWRVYS | 0.4782 | 16 | ABCPRED | 1 | ALLERGEN | Non-Toxin | -0.06 | -0.03 | -0.51 | 0.5 | 7.09 | 1857.33 | outside |
| 558 | 576 | KFLPFQQFGRDIADTTDA | 0.375 | 18 | ABCPRED | 1 | NON-ALLERGEN | Non-Toxin | -0.17 | -0.5 | 0.14 | -1 | 4.43 | 2070.54 | outside |
| 244 | 260 | LHRSYLTPGDSSSGWT | 0.4647 | 16 | ABCPRED | 1 | ALLERGEN | Non-Toxin | -0.16 | -0.8 | -0.21 | 0.5 | 7.09 | 1764.11 | outside |
| 245 | 263 | HRSYLTPGDSSSGWTAGA | 0.4794 | 18 | ABCPRED,BCPREDS | 2 | NON-ALLERGEN | Non-Toxin | -0.14 | -0.74 | -0.14 | 0.5 | 7.09 | 1850.18 | outside |
| 188 | 206 | NLREFVFKNIDGYFKIYS | -0.0865 | 18 | ABCPRED | 1 | ALLERGEN | Non-Toxin | -0.13 | -0.26 | -0.18 | 1 | 8.76 | 2253.85 | outside |
| 184 | 202 | GNFKNLREFVFKNIDGYF | -0.0622 | 18 | ABCPRED | 1 | NON-ALLERGEN | Non-Toxin | -0.15 | -0.46 | -0.1 | 1 | 8.83 | 2208.77 | outside |
| 101 | 119 | IRGWIFGTTLDSKTQSLL | 0.152 | 18 | ABCPRED | 1 | NON-ALLERGEN | Non-Toxin | -0.05 | 0.13 | -0.35 | 1 | 9.1 | 2036.64 | outside |
| 63 | 81 | TWFHAIHVSGTNGTKRFD | 0.5104 | 18 | ABCPRED | 1 | NON-ALLERGEN | Non-Toxin | -0.13 | -0.57 | -0.27 | 2 | 9.11 | 2074.55 | outside |
| 77 | 89 | KRFDNPVLPFND | 0.3102 | 12 | ABCPRED | 1 | ALLERGEN | Non-Toxin | -0.29 | -1 | 0.34 | 0 | 6.31 | 1461.8 | outside |
| 188 | 200 | NLREFVFKNIDG | -0.2284 | 12 | ABCPRED | 1 | NON-ALLERGEN | Non-Toxin | -0.19 | -0.39 | 0.19 | 0 | 6.42 | 1451.82 | outside |
| 75 | 93 | GTKRFDNPVLPFNDGVYF | 0.2732 | 18 | ABCPRED | 1 | ALLERGEN | Non-Toxin | -0.12 | -0.43 | -0.14 | 0 | 6.31 | 2086.59 | outside |
| 623 | 641 | AIHADQLTPTWRVYSTGS | 0.5664 | 18 | ABCPRED,literature | 2 | NON-ALLERGEN | Non-Toxin | -0.1 | -0.36 | -0.37 | 0.5 | 7.09 | 2003.46 | outside |
| 1027 | 1043 | TKMSECVLGQSKRVDF | 1.0151 | 16 | ABCPRED | 1 | NON-ALLERGEN | Non-Toxin | -0.25 | -0.38 | 0.36 | 1 | 8.53 | 1828.36 | outside |
| 1088 | 1104 | HFPREGVFVSNGTHWF | 0.265 | 16 | ABCPRED | 1 | NON-ALLERGEN | Non-Toxin | -0.05 | -0.37 | -0.55 | 1 | 7.26 | 1917.35 | outside |
| 73 | 85 | TNGTKRFDNPVL | 0.0607 | 12 | ABCPRED | 1 | ALLERGEN | Non-Toxin | -0.29 | -0.96 | 0.23 | 1 | 9.1 | 1361.69 | outside |
| 813 | 831 | SKRSFIEDLLFNKVTLAD | 0.3091 | 18 | ABCPRED | 1 | ALLERGEN | Non-Toxin | -0.17 | -0.06 | 0.23 | 0 | 6.47 | 2096.67 | outside |
| 725 | 743 | EILPVSMTKTSVDCTMYI | 1.2689 | 18 | ABCPRED | 1 | NON-ALLERGEN | Non-Toxin | 0 | 0.56 | -0.33 | -1 | 4.38 | 2031.69 | outside |
| 243 | 263 | ALHRSYLTPGDSSSGWTAGA | 0.3957 | 20 | BCPREDS,FBCPred | 2 | ALLERGEN | Non-Toxin | -0.08 | -0.39 | -0.24 | 0.5 | 7.09 | 2034.45 | outside |
| 349 | 365 | SVYAWNRKRISNCVAD | 0.2357 | 16 | ABCPRED | 1 | NON-ALLERGEN | Non-Toxin | -0.27 | -0.51 | 0.03 | 2 | 9.36 | 1882.34 | outside |
| 74 | 90 | NGTKRFDNPVLPFNDG | 0.1375 | 16 | ABCPRED | 1 | NON-ALLERGEN | Non-Toxin | -0.25 | -1.06 | 0.24 | 0 | 6.31 | 1791.18 | outside |
| 93 | 109 | ASTEKSNIIRGWIFGT | -0.3043 | 16 | ABCPRED | 1 | NON-ALLERGEN | Non-Toxin | -0.08 | -0.12 | -0.17 | 1 | 9.1 | 1780.25 | outside |
| 399 | 415 | SFVIRGDEVRQIAPGQ | 0.5366 | 16 | ABCPRED | 1 | NON-ALLERGEN | Non-Toxin | -0.18 | -0.26 | 0.19 | 0 | 6.42 | 1772.23 | outside |
| 209 | 227 | PINLVRDLPQGFSALEPL | 0.7044 | 18 | ABCPRED | 1 | NON-ALLERGEN | Non-Toxin | -0.03 | 0.22 | -0.21 | -1 | 4.38 | 1979.58 | outside |
| 455 | 467 | LFRKSNLKPFER | 0.4123 | 12 | ABCPRED | 1 | ALLERGEN | Non-Toxin | -0.42 | -1.08 | 0.58 | 3 | 11.01 | 1534.99 | outside |
| 70 | 88 | VSGTNGTKRFDNPVLPFN | 0.571 | 18 | ABCPRED | 1 | ALLERGEN | Non-Toxin | -0.17 | -0.6 | -0.04 | 1 | 9.1 | 1963.44 | outside |
| 31 | 47 | SFTRGVYYPDKVFRSS | 0.1599 | 16 | ABCPRED | 1 | ALLERGEN | Non-Toxin | -0.24 | -0.63 | -0.01 | 2 | 9.72 | 1909.34 | outside |
| 465 | 481 | ERDISTEIYQAGSTPC | 0.0412 | 16 | ABCPRED | 1 | NON-ALLERGEN | Non-Toxin | -0.21 | -0.72 | 0.29 | -2 | 4.14 | 1770.12 | outside |
| 337 | 357 | PFGEVFNATRFASVYAWNRK | 0.2378 | 20 | ABCPRED | 1 | NON-ALLERGEN | Non-Toxin | -0.14 | -0.35 | -0.27 | 2 | 10 | 2360.93 | outside |
| 340 | 358 | EVFNATRFASVYAWNRKR | 0.3543 | 18 | ABCPRED | 1 | NON-ALLERGEN | Non-Toxin | -0.29 | -0.68 | 0.01 | 3 | 10.91 | 2215.74 | outside |
| 71 | 92 | SGTNGTKRFDNPVLPFNDGVY | 0.3544 | 21 | IEDB | 1 | NON-ALLERGEN | Non-Toxin | -0.17 | -0.76 | 0 | 0 | 6.31 | 2298.8 | outside |
| 1100 | 1118 | THWFVTQRNFYEPQIITT | 0.4386 | 18 | ABCPRED | 1 | ALLERGEN | Non-Toxin | -0.11 | -0.53 | -0.63 | 0.5 | 7.1 | 2281.83 | outside |
| 1105 | 1123 | TQRNFYEPQIITTDNTFV | 0.5081 | 18 | ABCPRED | 1 | ALLERGEN | Non-Toxin | -0.18 | -0.69 | -0.23 | -1 | 4.38 | 2187.66 | outside |
| 557 | 575 | KKFLPFQQFGRDIADTTD | 0.2409 | 18 | ABCPRED | 1 | ALLERGEN | Non-Toxin | -0.25 | -0.82 | 0.33 | 0 | 6.39 | 2127.64 | outside |
| 33 | 45 | TRGVYYPDKVFR | 0.3643 | 12 | BCPREDS | 1 | ALLERGEN | Non-Toxin | -0.31 | -0.88 | 0.13 | 2 | 9.72 | 1500.88 | outside |
| 143 | 161 | VYYHKNNKSWMESEFRVY | 0.3309 | 18 | ABCPRED,BCPREDS | 2 | ALLERGEN | Non-Toxin | -0.28 | -1.27 | -0.09 | 1.5 | 8.71 | 2380.92 | outside |
| 147 | 167 | KNNKSWMESEFRVYSSANNC | 0.2134 | 20 | ABCPRED | 1 | NON-ALLERGEN | Non-Toxin | -0.34 | -1.28 | 0.23 | 1 | 8.51 | 2394.89 | outside |
| 805 | 823 | ILPDPSKPSKRSFIEDLL | -0.0841 | 18 | ABCPRED,literature | 2 | NON-ALLERGEN | Non-Toxin | -0.19 | -0.38 | 0.41 | 0 | 6.47 | 2055.66 | outside |
| 552 | 571 | LTESNKKFLPFQQFGRDIA | 0.3962 | 19 | literature | 1 | NON-ALLERGEN | Non-Toxin | -0.2 | -0.58 | 0.11 | 1 | 8.93 | 2239.84 | outside |
| 272 | 288 | PRTFLLKYNENGTITD | 0.2056 | 16 | ABCPRED | 1 | ALLERGEN | Non-Toxin | -0.22 | -0.81 | 0.06 | 0 | 6.42 | 1882.34 | outside |
| 272 | 292 | PRTFLLKYNENGTITDAVDC | 0.2329 | 20 | ABCPRED | 1 | ALLERGEN | Non-Toxin | -0.17 | -0.39 | 0.05 | -1 | 4.56 | 2270.83 | outside |
| 440 | 502 | [illegible]RLFRKSNLKPFERDISTEIYQAGSTPCNGVEGFN | 0.3951 | 62 | IEDB,literature | 2 | NON-ALLERGEN | too long | | | | | | | outside |
| 144 | 161 | YYHKNNKSWMESEFRVY | 0.3439 | 17 | BCEPRED | 1 | NON-ALLERGEN | Non-Toxin | -0.33 | -1.59 | -0.01 | 1.5 | 8.71 | 2281.77 | outside |
| 406 | 427 | EVRQIAPGQTGKIADYNYKLP | 1.3005 | 21 | literature | 1 | ALLERGEN | Non-Toxin | -0.2 | -0.79 | 0.13 | 1 | 8.76 | 2362 | outside |
| 456 | 474 | FRKSNLKPFERDISTEIY | 0.2315 | 18 | ABCPRED | 1 | ALLERGEN | Non-Toxin | -0.33 | -0.98 | 0.48 | 1 | 8.83 | 2243.8 | outside |
| 453 | 469 | YRLFRKSNLKPFERDI | 0.1133 | 16 | ABCPRED | 1 | ALLERGEN | Non-Toxin | -0.42 | -1.11 | 0.55 | 3 | 10.28 | 2082.66 | outside |
| 142 | 160 | GVYYHKNNKSWMESEFRV | 0.5457 | 18 | ABCPRED | 1 | ALLERGEN | Toxin | -0.27 | -1.22 | 0.04 | 1.5 | 8.76 | 2274.8 | outside |
| 453 | 471 | YRLFRKSNLKPFERDIST | 0.0127 | 18 | ABCPRED | 1 | NON-ALLERGEN | Non-Toxin | -0.4 | -1.07 | 0.48 | 3 | 10.28 | 2270.87 | outside |
| 812 | 832 | PSKRSFIEDLLFNKVTLADA | 0.2946 | 20 | ABCPRED | 1 | NON-ALLERGEN | Non-Toxin | -0.14 | -0.05 | 0.19 | 0 | 6.47 | 2264.89 | outside |

| | | | | | | | | | | | | | | | |
|---|---|---|---|---|---|---|---|---|---|---|---|---|---|---|---|
| 455 | 473 | LFRKSNLKPFERDISTEI | 0.2087 | 18 | ABCPRED | 1 | NON-ALLERGEN | Non-Toxin | -0.3 | -0.69 | 0.51 | 1 | 8.93 | 2193.79 | outside |
| 93 | 113 | ASTEKSNIIRGWIFGTTLDS | -0.2274 | 20 | ABCPRED,BCPREDS | 2 | NON-ALLERGEN | Non-Toxin | -0.09 | -0.16 | -0.08 | 0 | 6.42 | 2196.74 | outside |
| 206 | 226 | KHTPINLVRDLPQGFSALEP | 0.7726 | 20 | ABCPRED | 1 | NON-ALLERGEN | Non-Toxin | -0.13 | -0.38 | 0.01 | 0.5 | 7.1 | 2232.87 | outside |
| 84 | 104 | LPFNDGVYFASTEKSNIIRG | 0.2665 | 20 | ABCPRED | 1 | NON-ALLERGEN | Non-Toxin | -0.1 | -0.2 | -0.1 | 0 | 6.42 | 2228.78 | outside |
| 457 | 473 | RKSNLKPFERDISTEI | 0.4298 | 16 | ABCPRED | 1 | NON-ALLERGEN | Non-Toxin | -0.41 | -1.19 | 0.84 | 1 | 8.93 | 1933.42 | outside |
| 457 | 472 | RKSNLKPFERDISTE | 0.4847 | 15 | IEDB | 1 | NON-ALLERGEN | Non-Toxin | -0.48 | -1.57 | 1.02 | 1 | 8.93 | 1820.24 | outside |
| 146 | 162 | HKNNKSWMESEFRVYS | 0.4069 | 16 | ABCPRED | 1 | ALLERGEN | Toxin | -0.37 | -1.58 | 0.3 | 1.5 | 8.84 | 2042.48 | outside |
| 92 | 112 | FASTEKSNIIRGWIFGTTLD | -0.0019 | 20 | ABCPRED | 1 | NON-ALLERGEN | Non-Toxin | -0.05 | 0.02 | -0.23 | 0 | 6.42 | 2256.84 | outside |
| 1079 | 1097 | PAICHDGKAHFPREGVFV | -0.1254 | 18 | ABCPRED | 1 | NON-ALLERGEN | Non-Toxin | -0.07 | -0.07 | -0.04 | 1 | 7.22 | 1980.53 | outside |
| 150 | 168 | KSWMESEFRVYSSANNCT | -0.0348 | 18 | ABCPRED | 1 | NON-ALLERGEN | Non-Toxin | -0.25 | -0.85 | 0.04 | 0 | 6.46 | 2139.58 | outside |
| 271 | 291 | QPRTFLLKYNENGTITDAVD | 0.2221 | 20 | ABCPRED,AAP | 2 | ALLERGEN | Non-Toxin | -0.21 | -0.69 | 0.11 | -1 | 4.56 | 2295.83 | outside |
| 447 | 472 | GNYNYLYRLFRKSNLKPFERDISTE | 0.1595 | 25 | BCEPRED | 1 | ALLERGEN | Non-Toxin | -0.33 | -1.16 | 0.23 | 2 | 9.43 | 3124.87 | outside |
| 149 | 161 | NKSWMESEFRVY | 0.332 | 12 | ABCPRED | 1 | ALLERGEN | Non-Toxin | -0.29 | -1.15 | 0.15 | 0 | 6.49 | 1575.92 | outside |
| 406 | 424 | EVRQIAPGQTGKIADYNY | 1.4691 | 18 | BCPREDS | 1 | NON-ALLERGEN | Non-Toxin | -0.2 | -0.82 | 0.08 | 0 | 6.41 | 2023.5 | outside |
| 399 | 419 | SFVIRGDEVRQIAPGQTGKI | 0.7803 | 20 | ABCPRED,AAP | 2 | NON-ALLERGEN | Non-Toxin | -0.16 | -0.24 | 0.2 | 1 | 9.1 | 2171.79 | outside |
| 810 | 830 | SKPSKRSFIEDLLFNKVTLA | 0.3115 | 20 | ABCPRED | 1 | ALLERGEN | Non-Toxin | -0.19 | -0.19 | 0.22 | 2 | 9.72 | 2293.98 | outside |
| 808 | 824 | DPSKPSKRSFIEDLLF | 0.2203 | 16 | ABCPRED | 1 | NON-ALLERGEN | Non-Toxin | -0.25 | -0.67 | 0.53 | 0 | 6.47 | 1879.36 | outside |
| 142 | 161 | GVYYHKNNKSWMESEFRVY | 0.4302 | 19 | BCEPRED | 1 | NON-ALLERGEN | Non-Toxin | -0.26 | -1.22 | -0.08 | 1.5 | 8.71 | 2437.99 | outside |
| 177 | 193 | MDLEGKQGNFKNLREF | 0.8263 | 16 | BCEPRED | 1 | ALLERGEN | Non-Toxin | -0.31 | -1.19 | 0.54 | 0 | 6.53 | 1926.42 | outside |
| 1029 | 1045 | MSECVLGQSKRVDFCG | 0.6471 | 16 | ABCPRED | 1 | NON-ALLERGEN | Non-Toxin | -0.16 | 0.04 | 0.14 | 0 | 6.36 | 1759.27 | outside |
| 175 | 193 | FLMDLEGKQGNFKNLREF | 0.5902 | 18 | ABCPRED | 1 | NON-ALLERGEN | Non-Toxin | -0.22 | -0.69 | 0.24 | 0 | 6.53 | 2186.79 | outside |
| 146 | 164 | HKNNKSWMESEFRVYSSA | 0.3352 | 18 | ABCPRED | 1 | NON-ALLERGEN | Non-Toxin | -0.33 | -1.34 | 0.26 | 1.5 | 8.84 | 2200.66 | outside |
| 143 | 163 | VYYHKNNKSWMESEFRVYSS | 0.3042 | 20 | ABCPRED,literature | 2 | NON-ALLERGEN | Non-Toxin | -0.28 | -1.22 | -0.05 | 1.5 | 8.71 | 2555.1 | outside |
| 457 | 475 | RKSNLKPFERDISTEIYQ | 0.2263 | 18 | ABCPRED | 1 | ALLERGEN | Non-Toxin | -0.4 | -1.33 | 0.63 | 1 | 8.83 | 2224.76 | outside |
| 176 | 194 | LMDLEGKQGNFKNLREFV | 0.5821 | 18 | ABCPRED | 1 | ALLERGEN | Non-Toxin | -0.22 | -0.61 | 0.3 | 0 | 6.53 | 2138.75 | outside |
| 809 | 821 | PSKPSKRSFIED | 0.3363 | 12 | ABCPRED,BCEPRED | 2 | NON-ALLERGEN | Non-Toxin | -0.41 | -1.47 | 0.97 | 1 | 8.93 | 1390.71 | outside |
| 148 | 164 | NNKSWMESEFRVYSSA | 0.3638 | 16 | ABCPRED | 1 | NON-ALLERGEN | Non-Toxin | -0.27 | -1.07 | 0.13 | 0 | 6.49 | 1935.31 | outside |
| 150 | 166 | KSWMESEFRVYSSANN | 0.1223 | 16 | ABCPRED | 1 | NON-ALLERGEN | Non-Toxin | -0.27 | -1.07 | 0.13 | 0 | 6.49 | 1935.31 | outside |
| 148 | 166 | NNKSWMESEFRVYSSANN | 0.3439 | 18 | ABCPRED,AAP | 2 | NON-ALLERGEN | Non-Toxin | -0.31 | -1.34 | 0.14 | 0 | 6.49 | 2163.55 | outside |
| 811 | 829 | KPSKRSFIEDLLFNKVTL | 0.2177 | 18 | ABCPRED | 1 | NON-ALLERGEN | Non-Toxin | -0.21 | -0.27 | 0.26 | 2 | 9.72 | 2135.8 | outside |
| 149 | 165 | NKSWMESEFRVYSSAN | 0.3261 | 16 | ABCPRED | 1 | NON-ALLERGEN | Non-Toxin | -0.27 | -1.07 | 0.13 | 0 | 6.49 | 1935.31 | outside |
| 1027 | 1047 | TKMSECVLGQSKRVDFCGKG | 0.8644 | 20 | AAP | 1 | NON-ALLERGEN | Non-Toxin | -0.24 | -0.41 | 0.39 | 2 | 8.94 | 2173.84 | outside |
| 1029 | 1047 | MSECVLGQSKRVDFCGKG | 0.8658 | 18 | AAP | 1 | NON-ALLERGEN | Non-Toxin | -0.19 | -0.21 | 0.29 | 1 | 8.36 | 1944.53 | outside |
| 1083 | 1103 | HDGKAHFPREGVFVSNGTHW | 0.2022 | 20 | AAP | 1 | NON-ALLERGEN | Non-Toxin | -0.16 | -0.89 | -0.06 | 1.5 | 7.35 | 2278.77 | outside |
| 552 | 570 | LTESNKKFLPFQQFGRDI | 0.4096 | 18 | ABCPRED | 1 | NON-ALLERGEN | Non-Toxin | -0.23 | -0.71 | 0.14 | 1 | 8.93 | 2168.75 | outside |
| 619 | 639 | EVPVAIHADQLTPTWRVYST | 0.4261 | 20 | ABCPRED | 1 | NON-ALLERGEN | Non-Toxin | -0.07 | -0.1 | -0.35 | -0.5 | 5.33 | 2283.86 | outside |
| 803 | 821 | SQILPDPSKPSKRSFIED | 0.0722 | 18 | AAP | 1 | NON-ALLERGEN | Non-Toxin | -0.3 | -1.04 | 0.64 | 0 | 6.47 | 2044.54 | outside |
| 148 | 168 | NNKSWMESEFRVYSSANNCT | 0.1806 | 20 | ABCPRED,AAP,BCPREDS | 3 | NON-ALLERGEN | Non-Toxin | -0.29 | -1.11 | 0.06 | 0 | 6.46 | 2367.82 | outside |
| 455 | 469 | LFRKSNLKPFERDI | 0.361 | 14 | IEDB | 1 | NON-ALLERGEN | Non-Toxin | -0.36 | -0.86 | 0.58 | 2 | 10 | 1763.27 | outside |
| 1101 | 1119 | HWFVTQRNFYEPQIITTD | 0.5142 | 18 | ABCPRED | 1 | NON-ALLERGEN | Non-Toxin | -0.14 | -0.68 | -0.44 | -0.5 | 5.33 | 2295.81 | outside |
| 1082 | 1098 | CHDGKAHFPREGVFVS | -0.034 | 16 | ABCPRED | 1 | ALLERGEN | Non-Toxin | -0.15 | -0.42 | 0.11 | 1 | 7.22 | 1786.22 | outside |
| 809 | 827 | PSKPSKRSFIEDLLFNKV | 0.2066 | 18 | literature | 1 | NON-ALLERGEN | Non-Toxin | -0.24 | -0.58 | 0.4 | 2 | 9.72 | 2105.72 | outside |
| 150 | 162 | KSWMESEFRVYS | 0.1763 | 12 | ABCPRED | 1 | ALLERGEN | Non-Toxin | -0.26 | -0.93 | 0.16 | 0 | 6.49 | 1548.89 | outside |
| 619 | 637 | EVPVAIHADQLTPTWRVY | 0.3281 | 18 | ABCPRED | 1 | ALLERGEN | Non-Toxin | -0.05 | -0.03 | -0.38 | -0.5 | 5.33 | 2095.65 | outside |
| 617 | 637 | CTEVPVAIHADQLTPTWRVY | 0.297 | 20 | ABCPRED | 1 | NON-ALLERGEN | Non-Toxin | -0.05 | 0.07 | -0.42 | -0.5 | 5.33 | 2299.92 | outside |
| 553 | 571 | TESNKKFLPFQQFGRDIA | 0.3035 | 18 | ABCPRED,literature | 2 | ALLERGEN | Non-Toxin | -0.24 | -0.82 | 0.22 | 1 | 8.93 | 2126.66 | outside |
| 85 | 105 | PFNDGVYFASTEKSNIIRGW | -0.0412 | 20 | ABCPRED | 1 | ALLERGEN | Non-Toxin | -0.1 | -0.44 | -0.18 | 0 | 6.42 | 2301.83 | outside |
| 812 | 824 | PSKRSFIEDLLF | 0.0348 | 12 | ABCPRED | 1 | NON-ALLERGEN | Non-Toxin | -0.15 | -0.08 | 0.18 | 0 | 6.42 | 1451.85 | outside |
| 94 | 112 | STEKSNIIRGWIFGTTLD | -0.2026 | 18 | ABCPRED | 1 | NON-ALLERGEN | Non-Toxin | -0.1 | -0.23 | -0.08 | 0 | 6.42 | 2038.56 | outside |
| 1083 | 1099 | HDGKAHFPREGVFVSN | 0.0493 | 16 | ABCPRED | 1 | NON-ALLERGEN | Non-Toxin | -0.19 | -0.79 | 0.19 | 1 | 7.26 | 1797.19 | outside |
| 144 | 162 | YYHKNNKSWMESEFRVYS | 0.3799 | 18 | ABCPRED | 1 | NON-ALLERGEN | Non-Toxin | -0.32 | -1.54 | 0.01 | 1.5 | 8.71 | 2368.86 | outside |
| 806 | 826 | LPDPSKPSKRSFIEDLLFNK | 0.033 | 20 | ABCPRED | 1 | NON-ALLERGEN | Non-Toxin | -0.26 | -0.79 | 0.49 | 1 | 8.83 | 2331.98 | outside |
| 329 | 364 | PNITNLCPFGEVFNATRFASVYAWNRKRISNCV | 0.4466 | 35 | IEDB,literature | 2 | NON-ALLERGEN | Non-Toxin | -0.1 | 0.05 | -0.39 | 3 | 9.53 | 4008.1 | outside |
| 177 | 195 | MDLEGKQGNFKNLREFVF | 0.8006 | 18 | ABCPRED | 1 | NON-ALLERGEN | Non-Toxin | -0.22 | -0.67 | 0.26 | 0 | 6.53 | 2172.76 | outside |
| 452 | 472 | LYRLFRKSNLKPFERDISTE | -0.0464 | 20 | BCEPRED | 1 | NON-ALLERGEN | Non-Toxin | -0.36 | -0.95 | 0.49 | 2 | 9.71 | 2513.18 | outside |
| 808 | 828 | DPSKPSKRSFIEDLLFNKVT | 0.2134 | 20 | ABCPRED | 1 | NON-ALLERGEN | Toxin | -0.27 | -0.73 | 0.49 | 1 | 8.83 | 2321.94 | outside |
| 458 | 476 | KSNLKPFERDISTEIYQA | 0.2293 | 18 | ABCPRED | 1 | ALLERGEN | Non-Toxin | -0.29 | -0.98 | 0.44 | 0 | 6.52 | 2139.65 | outside |
| 1081 | 1097 | ICHDGKAHFPREGVFV | -0.1447 | 16 | ABCPRED | 1 | NON-ALLERGEN | Non-Toxin | -0.09 | -0.09 | -0.02 | 1 | 7.22 | 1812.31 | outside |
| 95 | 113 | TEKSNIIRGWIFGTTLDS | -0.2974 | 18 | ABCPRED | 1 | NON-ALLERGEN | Non-Toxin | -0.1 | -0.23 | -0.08 | 0 | 6.42 | 2038.56 | outside |
| 806 | 822 | LPDPSKPSKRSFIEDL | 0.0302 | 16 | ABCPRED | 1 | NON-ALLERGEN | Non-Toxin | -0.29 | -0.94 | 0.69 | 0 | 6.47 | 1829.3 | outside |
| 637 | 643 | STGSNV | 0.3608 | 6 | IEDB | 1 | ALLERGEN | Non-Toxin | -0.11 | -0.33 | -0.18 | 0 | 5.88 | 563.64 | outside |
| 41 | 42 | K | short | 1 | literature | 1 | NON-ALLERGEN | Non-Toxin | -1.1 | -3.9 | 3 | 1 | 9.11 | 146.19 | outside |
| 602 | 607 | TNTSN | short | 5 | literature | 1 | NON-ALLERGEN | Non-Toxin | -0.38 | -1.84 | -0.02 | 0 | 5.88 | 535.57 | outside |
| 96 | 97 | E | short | 1 | literature | 1 | NON-ALLERGEN | Non-Toxin | -0.62 | -3.5 | 3 | -1 | 4 | 147.13 | outside |
| 601 | 609 | GTNTSNQV | 0.7179 | 8 | BCEPRED | 1 | ALLERGEN | Non-Toxin | -0.24 | -1.11 | -0.18 | 0 | 5.88 | 819.94 | outside |
